# Gene- and genome-centric dynamics shape the diversity of oral bacterial populations

**DOI:** 10.1101/2021.05.14.444208

**Authors:** Daniel R. Utter, Colleen M. Cavanaugh, Gary G. Borisy

## Abstract

Two major viewpoints have been put forward for how microbes adapt to a niche, differing in whether adaptation is driven principally by gene-centric or genome-centric processes. Longitudinal sampling at microbially-relevant timescales, i.e., days to weeks, is critical for distinguishing these mechanisms. Because of its significance for both microbial ecology and human health and its accessibility and high level of curation, we used the oral microbiota to evaluate evolutionary mechanisms. Metagenomes were generated by shotgun sequencing of total community DNA from the healthy tongues of 17 volunteers at four to seven timepoints obtained over intervals of days to weeks. We obtained 390 high-quality metagenome-assembled genomes (MAGs) defining population genomes from 55 genera, the majority of which were temporally stable at the MAG level. Decomposing MAG-defined populations by single nucleotide variant frequencies revealed MAGs were composed of up to 5 haplotypes, putatively distinct strain- or subpopulation-level genotypes. Most haplotypes were stable over time, yet we found examples of individual haplotypes sweeping from low abundance to dominance in a population over a period of days, a pattern suggestive of genome-centric adaptation. At the gene level, the vast majority of genes in each MAG were tightly linked over the two-week sampling window based on their frequency in the metagenomes of different mouths. The few genes that changed in abundance independently from nearby genes did not change in a directional manner, nor did nonsynonymous codon variants within such genes. Altogether, these observations characterize the intrapopulation genomic dynamics of the oral microbiota at microbially-relevant timescales. Our results demonstrate that both gene- and genome-wide sweeps occur on daily timescales but likely with different ecological ramifications. We infer that genome-wide selection of ecotypes is the dominant mode of adaptation in the oral populations, with short-term changes in gene frequency also occurring.

## Introduction

The mechanisms by which bacterial populations adapt to a niche is a central yet unresolved issue in microbial ecology. Two major viewpoints have been put forward, differing in whether adaptation is driven principally by gene-centric or by genome-centric processes. The gene-centric view hypothesizes that a niche defines selection for traits encoded by a few key genes. Over time, specific genes change in frequency as they sweep through the population more or less independently (Burke et al., 2011; Doolittle & Booth, 2016; Garud et al., 2019; Polz et al., 2013; Shapiro et al., 2012). The genome-centric view, also referred to as the ecotype hypothesis, holds that population members change as discrete eco-evolutionary units termed ‘ecotypes’, thus changing the frequency of an entire genome or set of related genomes as selection sweeps favored population members to higher abundance (Bendall et al., 2016; Cohan & Perry, 2007; Garud & Pollard, 2020). While both viewpoints agree that the genomic composition of bacterial populations change, the different processes suggested by each viewpoint imply different ecological dynamics.

Distinguishing gene-specific vs genome-wide sweeps in a complex natural microbiome necessitates analyzing genomes representative of the members living in that microbiome. Metagenomes, resulting from the shotgun sequencing of total DNA from a sample, capture a snapshot of the genomic composition of a community, yet are typically fragmented short reads lacking genomic context (Quince et al., 2017b). Metagenome-assembled genomes (MAGs) avoid cultivation bias and so are currently the best genomic representation of a population (Nayfach et al., 2019). Recent studies have generated hundreds of thousands of MAGs from published human metagenomes, underscoring the extent of genomic diversity within the human microbiome (Nayfach et al., 2019; Pasolli et al., 2019). However, most of these MAGs originate from populations sampled only at a single timepoint; thus, they provide no information about how each population varies over time.

Longitudinal sampling at microbially-relevant timescales is critical to distinguishing gene-centric vs genome-centric adaptation (Garud & Pollard, 2020). While both viewpoints represent genetic change in a population, they differ in the predicted dynamics of the adaptive process. Gene-centric adaptation predicts incremental change from gene-specific sweeps (Shapiro, 2018), while genome-centric adaptation predicts more drastic changes through pre-existing genomes sweeping to abundance (Cohan, 2016). Thus, the evolutionary processes can be distinguished by observing whether populations change through gene-by-gene fixation or by groups of genes changing together. Implicit in this approach is that the process must be monitored at timescales relevant to microbial growth and selection.

Longitudinal sampling is also methodologically advantageous as it can improve confidence in the quality of MAGs that are generated (Quince et al., 2017b). Since the same population is typically re-sampled over a longitudinal series, the process of binning assembled contiguous fragments (contigs) into MAGs is improved by the expectation that all parts of the same chromosome should have high correlation in relative abundance despite changes in the total population abundance (Alneberg et al., 2014; Chen et al., 2019). This expectation also allows a means of quality control by identifying misplaced contigs based on aberrant coverage relative to other MAGs (Chen et al., 2019).

Adaptation in complex microbiomes has been studied by a variety of sequence methods from marker gene sequencing to metagenomes to MAGs, generally focusing on the gut microbiome and at long timescales. Examples of intra-species phase shifts and habitat restriction were discovered via 16S ASVs in the human microbiome and presumed to reflect different ecotypes (Eren et al., 2014; Hall et al., 2017; Mark Welch et al., 2014; Mukherjee et al., 2018; Utter et al., 2016) although ASVs potentially could reflect ecologically neutral variants. Sharon et al. sequenced infant gut metagenomes and identified rapid changes among populations on the order of days (Sharon et al., 2013), and further analyses of the same data revealed changing single-nucleotide polymorphism (SNP) frequencies within those fluctuating populations (Eren et al., 2015). Zhao et al. (Zhao et al., 2019) cultivated and sequenced numerous *B. fragilis* isolate genomes in conjunction with metagenomic sequencing to reveal each volunteer’s gut *B. fragilis* population dominated by a few strains of *B. fragilis* differing by single-digit numbers of SNPs. Further, the strain composition changed as new strains arose via SNPs in a few genes under strong selection or mobile elements. Notably, they found strain coalescence within approximately 1 year.

Garud et al. (2019) found a few cases of strain replacement in the gut, but more commonly gene sweeps via recombination of existing genes. In the high-selection environment of hospitals, Evans et al. (Evans et al., 2020) documented frequent mobile element-mediated horizontal gene transfer between populations isolated from infections on the order of days to weeks. In a long-term longitudinal survey of the gut microbiome of a single human, treatment with an antibiotic seemingly induced genome-wide sweeps within some populations (Roodgar et al. 2020). Outside the human microbiome, long-term serial passaging of laboratory communities derived from pitcher plant phytotelmata revealed different ecological dynamics among related strains that sometimes resulted in genome-wide sweeps (Goyal et al. 2021). These reports emphasize different features of adaptation important to populations in complex microbiomes, particularly the human gut microbiome at long timescales.

The human oral microbiome represents a distinct microbial environment from the gut and a readily accessible site for investigating basic questions of microbial ecology. Approximately 700 recognized bacterial species inhabit the oral cavity (Escapa et al. 2019), the majority of which display sharply differential abundances among the different oral sites (Eren et al., 2014; Kraal et al., 2014; Lloyd-Price et al., 2017) leading to the hypothesis that the majority of oral bacterial populations are specialists adapted to a specific oral habitat (Mark Welch et al., 2019). In contrast, the gut microbiota is thought to harbor thousands of species (Almeida et al., 2019; Claesson et al., 2009). Further, oral microbes typically exist in highly ordered, discrete biofilms (Kim et al., 2020; Mark Welch et al., 2016; Wilbert et al., 2020), in contrast to the better-mixed gut bacteria (Earle et al., 2015; Mark Welch et al., 2017). Combined with the mouth’s direct exposure to exogenous factors, these ecological distinctions of the oral microbiome could result in different intra-population dynamics than elsewhere in the human microbiome.

These considerations of adaptation emphasize the importance of matching the timescale of analysis to the biology of the microbiome. Thus, we collected a metagenomic time series from the healthy human tongue dorsa of 17 volunteers, sampled over a period of days to two weeks, to investigate changes in microbial populations. Here we use ‘population’ to refer to a set of phylogenetically related lineages defined metagenomically. We constructed MAGs and tracked the abundance of genes and nucleotide variants within the MAGs to identify gene and genome-wide sweeps. Both gene- and genome-wide sweeps occurred on daily timescales but to different extents, suggesting that both modes occur in adaptation but differ in their relative impact. We suggest adaptation in the oral microbiome involves both gene- and genome-centric viewpoints to differing extents, where genome-wide selection of ecotypes is the most ecologically impactful mode of adaptation for oral bacteria, with short-term changes in gene frequency also occurring within populations to possibly generate new ecotypes over longer timescales.

## Methods

### Sample collection and library preparation

We sampled the tongue dorsum from 19 self-described healthy human volunteers at days 1, 2, 5, 8, and 15 under IRB oversight (IRB16-0367). Volunteers were between the ages of 20 and 29 years; 11 (58%) female, 8 (42%) male. Participants were sampled in the morning and were asked to abstain from eating or drinking anything besides water before sampling. Some donors were sampled at off days due to their availability (e.g., day 3 instead of day 2), or missed one of the 5 days; one donor was sampled 7 times (days 1, 2, 3, 5, 8, 13, 15). In total, 95 samples were self-collected by the volunteers with UV-sterilized disposable tongue scrapers (BreathRx, Philips; Stamford, CT). At each collection day a sampling control was collected, consisting of a sterile scraper being opened and transferred to an empty collection tube. Samples were immediately flash-frozen in liquid nitrogen and stored at −80°C until DNA extraction.

DNA was extracted with Qiagen DNeasy PowerLyzer kits (Germantown, MD) for all samples and sampling controls and stored at −20°C until used for library preparation. A blank extraction (nothing added to extraction kit) was prepared along with each extraction batch as an extraction control. Only one control, a single extraction control, had detectable DNA with a TapeStation HS D1000 tape (Agilent), but was too low for sequencing at 0.01 ng/μL and therefore was dropped. The 81 samples from 17 volunteers with DNA concentrations >1 ng/uL were used to prepare metagenomic libraries with Kapa DNA HyperPrep kit (Wilmington, MA) according to the manufacturer’s recommendations. Input total DNA was sheared with a Covaris sonicator to an average fragment size of 600 bp. Libraries were quantified with a TapeStation (Agilent; Lexington, MA), pooled at equimolar concentrations, and size selected with a Pippin Prep for 600-1200 bp fragments.

### Sequence analysis and MAG (metagenome-assembled genome) construction

The complete bioinformatic workflow, including all code with a full explanation, is provided in Supplemental Methods [URL].

The same library pool of 81 metagenomes was sequenced with 10 runs of Illumina NextSeq 550 with the High Output paired-end 150bp kit, producing over 6 billion read pairs. Metagenomic short reads were quality controlled with illumina-utils (Eren et al., 2013) following quality recommendations by Minoche et al (2011). Human reads were discarded by mapping with bowtie2 (Langmead & Salzberg, 2012) to the human genome (hg38), resulting in a total of 6,210,900,714 quality-controlled reads.

For each donor, the assembly program IDBA-UD used metagenomes from all of each donor’s timepoints to produce a single assembly, i.e., co-assembly by donor. Co-assembly by donor was chosen to avoid overloading the assembler with too much diversity from a co-assembly of all samples (Sczyrba et al., 2017) while still recovering low-abundance populations and retaining the advantages offered by longitudinal sampling of individuals. Each coassembly’s contigs were binned with CONCOCT (Alneberg et al., 2014) on the basis of tetranucleotide frequency and differential coverage of contigs across that co-assembly’s samples, and manually refined in anvi’o (Eren et al., 2015) using best practices for MAG refinement (Delmont et al., 2018; Shaiber et al., 2019). Manual refinement consisted of inspecting each bin to remove wrongly-included contigs identified by outlier coverage and/or nucleotide variability in the metagenomes from which the contigs were assembled (Chen et al., 2019; Shaiber et al., 2019). A set of 71 universal bacterial single-copy genes were used to assess genome completeness and redundancy (Bowers et al., 2017; Campbell et al., 2013). Genes were called with Prodigal (Hyatt et al., 2010) with default parameters. Gene functions were predicted with Interproscan for Pfam and TIGRFAM annotations, and emapper v2 for eggNOG annotations, including NCBI COGs (El-Gebali et al., 2019; Galperin et al., 2015; Haft et al., 2001; Huerta-Cepas et al., 2017; Huerta-Cepas et al., 2019; Jones et al., 2014).

From these bins, MAGs were defined as bins that are less than 10% redundant and either have a size of at least 2 Mbp or a completion estimate of >80%. After refinement, the 17 co-assemblies’ MAGs (n=409) were pooled and duplicate MAGs (e.g., substantially identical MAGs defined independently in different mouths) were identified as having ≥98% ANI over at least 75% of the smaller MAG’s length and a Pearson correlation coefficient of ≥0.85 for their coverage across all samples. 19 MAGs (5%) were identified as duplicate and discarded to retain a dereplicated set of 390 unique MAGs (Supplemental Table 1). These criteria for MAG inclusion span both the high- and medium-quality draft genome categories set out by the Genomic Standards Consortium (Bowers et al., 2017). 376 and 188 out of 390 MAGs had redundancy estimates of ≤5% and ≤1%, respectively; therefore, combined with the manual curation based on above parameters, the final 390 MAGs have a very low chance including contaminant contigs from distant populations that could affect their interpretability.

MAGs were assigned taxonomy by GTDB-Tk v1.40 (Chaumeil et al., 2019) using the Genomes Taxonomy Database release 95 (Parks et al., 2020). While the final set of MAGs included representatives from the majority of common oral taxa (Supplemental Table 1, Supplemental Figure 1), certain genera, namely *Streptococcus, Veillonella, Haemophilus*, and *Neisseria*, did not assemble or bin well and were underrepresented. Their exclusion was likely due to the challenges posed to assembly algorithms by highly diverse populations (Quince et al., 2017b; Sczyrba et al., 2017). Thus, our analyses focused on populations for which high-quality MAGs could be generated.

### Coverage-based definitions and metrics

The final coverage was determined using Bowtie2 (Langmead & Salzberg, 2012) to map the raw short reads to a single database containing all 390 non-redundant MAGs. Coverage for a collection of nucleotides, e.g., a gene or MAG, is reported as the average nucleotide’s coverage. All single nucleotide variants (SNVs) mapped by Bowtie2 were retained regardless of coverage depth to allow application-specific filtering. Detection for a collection of nucleotides, e.g., a gene or genome, is defined as the fraction of that collection’s nucleotides receiving any coverage at all. Relative abundance for a MAG in a sample is the percentage of reads recruited by a MAG relative to the total reads recruited by all MAGs.

### Haplotyping

We used the SNV counts and DESMAN (Quince et al., 2017a) following the developers’ recommendations to deconvolve each MAG into haplotypes. DESMAN attempts to split a consensus population genome (in this case, a MAG) into constitutive ‘haplotypes’ by tracking SNVs in core genes that co-vary, using a set of 71 core genes found in a single copy in all known bacteria and archaea (Alneberg et al., 2014). DESMAN requires prior information on the expected number of haplotypes in the population. To estimate this, we decomposed each MAG into 1, 2, 3, 4, 5, 6, 8, 10, 12 haplotypes for 500 iterations and plotted the mean posterior deviance for each MAG and haplotype combination. The optimal number of haplotypes for each MAG was determined by SNV uncertainty below 10% (Quince et al., 2017a) and a posterior deviance step of >10%.

### Detecting outlier genes

To identify genes that were outliers relative to the other genes in a MAG, we detrended each gene’s coverage by subtracting the gene’s coverage from the MAG coverage value. This detrending was performed for the 199 MAGs with at least 50% of the length of the MAG receiving coverage in at least 15 samples (approximately 3 donors). Outliers, termed decoupled genes, were then defined as genes with detrended coverages at least 5 standard deviations above the mean gene’s detrended coverage.

To estimate the linkage of genes within a population, we also correlated coverage of gene pairs for all possible pairs in a MAG. This method is sensitive to MAG-scale detection; if a MAG is truly present in only one donor but a few of its genes receive coverage in another donor’s samples, inclusion of the other donor’s metagenomes will bias the results as all gene pairs will be correlated yet they are not all equally represented. So, for each MAG, we correlated gene pairs using only the samples from mouths covering at least 75% of that MAG’s nucleotides for 40, 30, 20, 10, and 5 samples. Correlations were determined mouth-by-mouth, iterating mouth-by-mouth using all samples from one mouth at a time, and across all samples from all donors meeting the detection criteria. Mouth-by-mouth correlations were only performed for MAGs detected in 40, 30, 20, and 10 samples due to the lack of change and the computationally infeasible number of pairwise correlations for all MAGs meeting the detection criteria in 5 samples.

### Synonymity metric

Coupled and uncoupled genes were identified by first correlating each gene’s coverage against that of its parent MAG over all samples in which that MAG was detected, for all genes from the 25 MAGs with 75% of their nucleotides receiving coverage in at least 40 samples (see above). Coupled genes were defined as genes producing correlations in the ninetieth or ninety-ninth percentile while uncoupled were defined as those producing correlations in the first or tenth percentiles. For each percentile bracket, single codon variants (SCVs) were determined by identifying and translating SNVs to SCVs from only the metagenomic short reads that completely covered the entire codon, after Delmont et al. (2019). From these counts, the synonymity was calculated as the fraction of possible synonymous pairs from all SCVs detected per codon per metagenome (Delmont et al., 2019). Note that this approach uses MAGs only for defining codon windows; the resultant synonymity is not relative to the reference MAG sequence but relative to all possible observed variants in the metagenome.

Trends for each SCV were determined in R from the above SCV data by computing the least squares linear regression of each SCV’s proportion over time within each mouth for codon positions with at least >10x coverage and detectable SCVs in at least 4 samples.

### MAG and haplotype naming conventions

MAGs are named in the format MAG_[DONOR]_[NUMBER], where [DONOR] is an alphabetical identifier for the donor from which that MAG was assembled and [NUMBER] is a unique number to distinguish multiple MAGs assembled from the same donor. DESMAN haplotypes are designated by appending “_h[NUMBER]” to the MAG designation, e.g., MAG_AI_08_h4 is the fourth haplotype based off of the eighth MAG refined from donor AI.

## Results

### Assembly of390 MAGs allows investigation of the population dynamics of oral bacteria

Longitudinal studies using metagenome-assembled genomes (MAGs) allow the investigation of the genomic dynamics of microbial populations at unprecedented resolution. But, to have utility the MAGs must meet certain criteria: First, they must be consistently detected over time; second, the MAGs must be faithful representations of natural microbial populations. Therefore, attention must be paid to the quality of the MAGs generated. Ideally, these MAGs will sample many diverse oral taxa; however, all taxa need not be represented as our focus is on generating high quality MAGs to serve as indicators for the genome dynamics of microbial populations within individuals.

We generated 390 unique metagenome-assembled genomes (MAGs) from 81 tongue samples obtained from 17 volunteers sampled at 4-7 timepoints over a two-week period. Contiguous stretches of nucleotides (contigs) were assembled in a single step for each donor using all of that donor’s timepoints and binned with CONCOCT (Alneberg et al., 2014) to generate rough genome bins. Bins were manually refined in anvi’o (Eren et al., 2015) using differential coverage, tetranucleotide frequency, and SNV frequencies to remove misplaced contigs (Chen et al., 2019; Quince et al., 2017b; Shaiber et al., 2019) (see Methods). Taxonomic annotation according to the Genomes Taxonomy Database (Parks et al., 2020) assigned the MAGs to 55 genera, including the common oral genera *Prevotella, Actinomyces*, and *Porphyromonas* (Supplemental Figure 1, Supplemental Table 1). *Prevotella* was the most represented genus among the dataset with 77 MAGs (Supplemental Figure 1). These 390 MAGs form the basis of our analyses into the population and sub-population dynamics of the oral microbiome.

Operationally, we define a population as all the metagenomic reads that recruit to one MAG vs. other MAGs. The recruiting MAG serves as the reference genome for that population. By tracking single nucleotide variants (SNVs) throughout a MAG across samples, we can identify which SNVs are tightly correlated and therefore not shuffled by recombination. These distinct SNV combinations mark ‘haplotypes’ – putative major genotypes within each population (Quince et al., 2017a). These haplotypes represent subpopulations of a MAG-defined population.

### Many populations are consistently detected across multiple timepoints

To track the process of genetic change in a population, its representative MAG should be consistently detected across multiple timepoints while subpopulations or gene frequencies may vary. The fraction of nucleotides in a MAG receiving any coverage provides a metric to assess a MAG’s presence in a sample (Figure 1A). We consider a MAG to be detected in a sample if at least 50% of its genome is covered. With this criterion, the thirty MAGs receiving the highest mean percent of coverage across all samples were detected consistently within individual subjects and across samples from multiple mouths (Figure 1B, Supplemental Figure 2). Indeed, 241 MAGs were detected in 10 or more samples and 83 MAGs were detected in at least 40 samples (Supplemental Figure 3); thus, the majority of MAGs were detected in at least two mouths across multiple samples, and approximately one-fifth of MAGs were detected in half the samples. Therefore, when analyzing the genome dynamics of populations (MAGs), the same population can be analyzed in multiple donors, avoiding any possible biases associated with inferences based solely on samples from which a MAG is defined.

**Figure 1.**
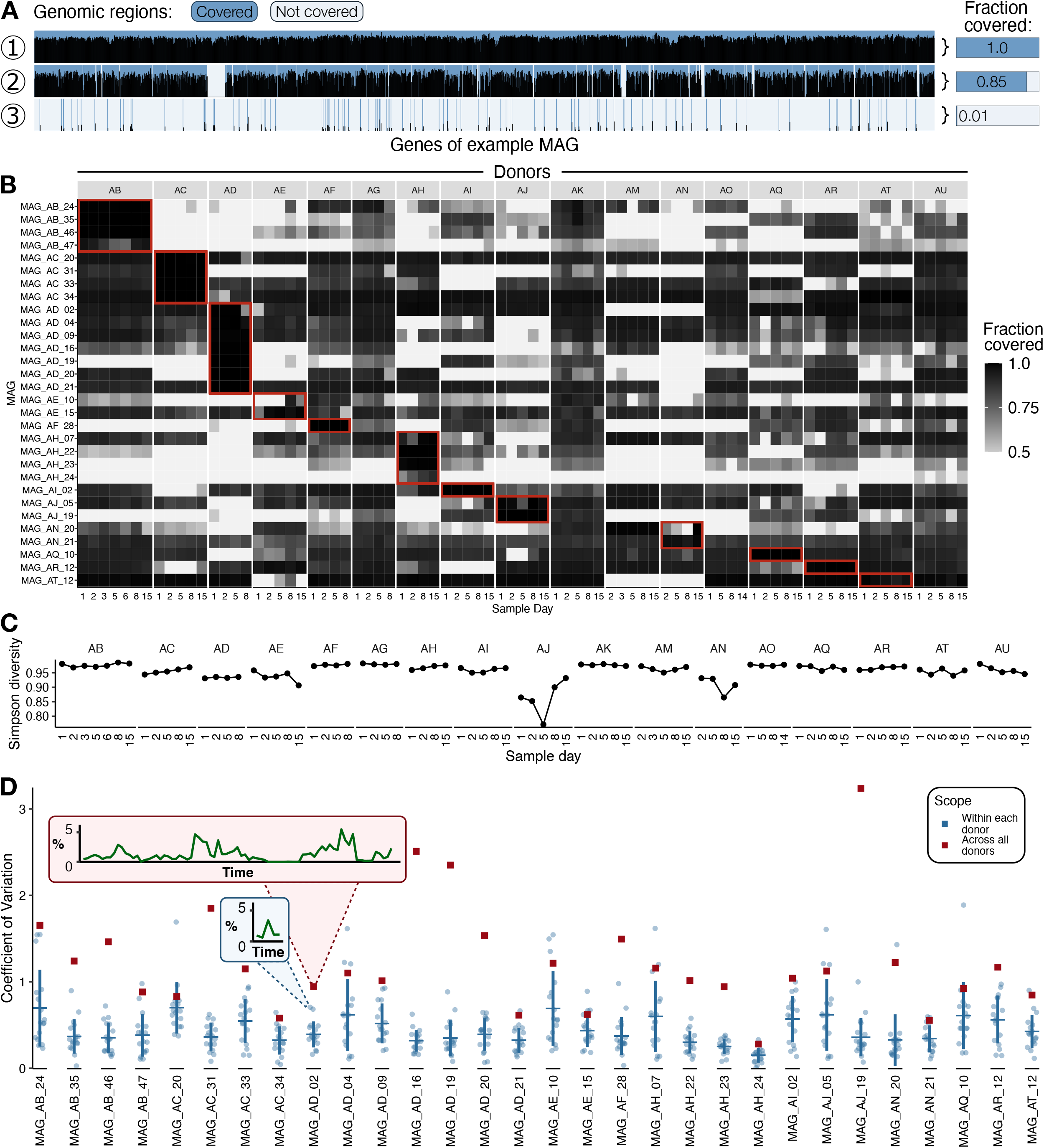
The 30 most abundant MAGs are detectable and stable over time within several individual mouths. **A)** The fraction of a MAG covered in a sample provides a useful detection metric. Here, for each of three example metagenomes (tracks 1, 2, and 3) a cartoon shows the coverage received (height of thin black bars) for each gene (x-axis) in a single MAG. The dark blue shading highlights regions receiving coverage from that metagenome while the pale blue shading highlights regions not receiving coverage. The fraction covered (right side) is calculated as the length of covered regions (dark blue) divided by the total length (dark blue + pale blue). **B)** Many MAGs are detected across timepoints and in multiple donors. A heatmap shows the fraction of MAG covered at all (shading darkness) for the 30 MAGs (rows) with the highest average relative abundance. Each column shows the fraction detected in a sample, with samples arranged in by donor (AB, AC, etc. listed on top) in chronological order. Red boxes outline the samples from which each MAG was co-assembled, revealing that many MAGs are detected outside their defining metagenomes. Only fractions >=0.5 are shown since we consider MAGs with less than half of their nucleotides covered to not be confidently detected. **C)** Simpson diversity (y-axis) of MAG relative abundances at each timepoint (x-axis) for each donor (groups of connected dots). **D)** Coefficient of variation (y-axis) of MAG relative abundance for the 30 MAGs with the highest average relative abundance. Along the x-axis unit are MAGs; for each is plotted the coefficients of variation for that MAG within each of the 17 donors (blue dots) or across all samples from all donors (red dot). The red and blue callout boxes show the underlying relative abundance data behind two coefficient of variation calculations, one across samples from all donors (red callout) and within a single donor (blue callout). The horizontal and vertical bars show the mean coefficient of variation ± 1 standard deviation, respectively.

Having confirmed that many MAGs are detected in multiple samples, we set out first to characterize the population-level diversity and dynamics before assessing the dynamics within populations. The diversity of all MAG-defined populations was broadly stable over a two-week period (Figure 1C), suggesting few or no major microbiome-level changes occurred within the time window analyzed. Similarly, among the 30 MAGs with the highest mean relative abundance, MAG relative abundance was generally stable within a mouth in that most of these MAGs had lower coefficients of variation within each donor than across donors (Figure 1D). For example, the blue callout box in Fig. 1D shows the relative abundances behind an arbitrary MAG/donor combination fluctuating up and down over time by approximately 2% of the total reads. This fluctuation was minor, however, compared to the approximately 5% change in relative abundance of the same MAG across the entire set of 81 samples from all donors (Figure 1D red callout). Thus, populations detectable and abundant at one timepoint are likely to be detectable with similar abundances at future timepoints of the same mouth, thereby allowing the investigation of sub-population dynamics, whether at the gene or genome level.

### Genome-wide sweeps of subpopulations occur at daily timescales

We investigated whether we could detect genome-wide sweeps within a MAG-defined population. DESMAN deconvolves a consensus or population genome, in this case each MAG, into ‘haplotypes’ that represent major strain types (Quince, et al., 2017a). DESMAN uses single nucleotide variant (SNV) frequencies that belong to single-copy core genes (Alneberg et al., 2014) in a MAG to identify combinations of co-occurring SNVs over multiple metagenome samples. Operationally, each DESMAN haplotype corresponds to a different subpopulation genotype.

All 390 MAGs were composed of one to five haplotypes (median = 3; Supplemental Figure 4), the majority of which were stable over time within a donor (Figure 2). One example of a MAG with a stable haplotype composition is illustrated in Figure 2B; all are presented in Supplemental Figure 5. For the example MAG in Fig. 2B, a 2.39 Mbp *Prevotella aurantiaca* MAG estimated to be 95% complete and 1.4% redundant, each haplotype was detectable in all 17 mouths at similar proportions on day 15 as at day 1. To place this example within the context of the entire dataset, the enlarged points in Figure 2A corresponding to MAG_AD_09_h1 in AM and across all donors are colored peach in the within- and across-donor comparisons, respectively.

**Figure 2.**
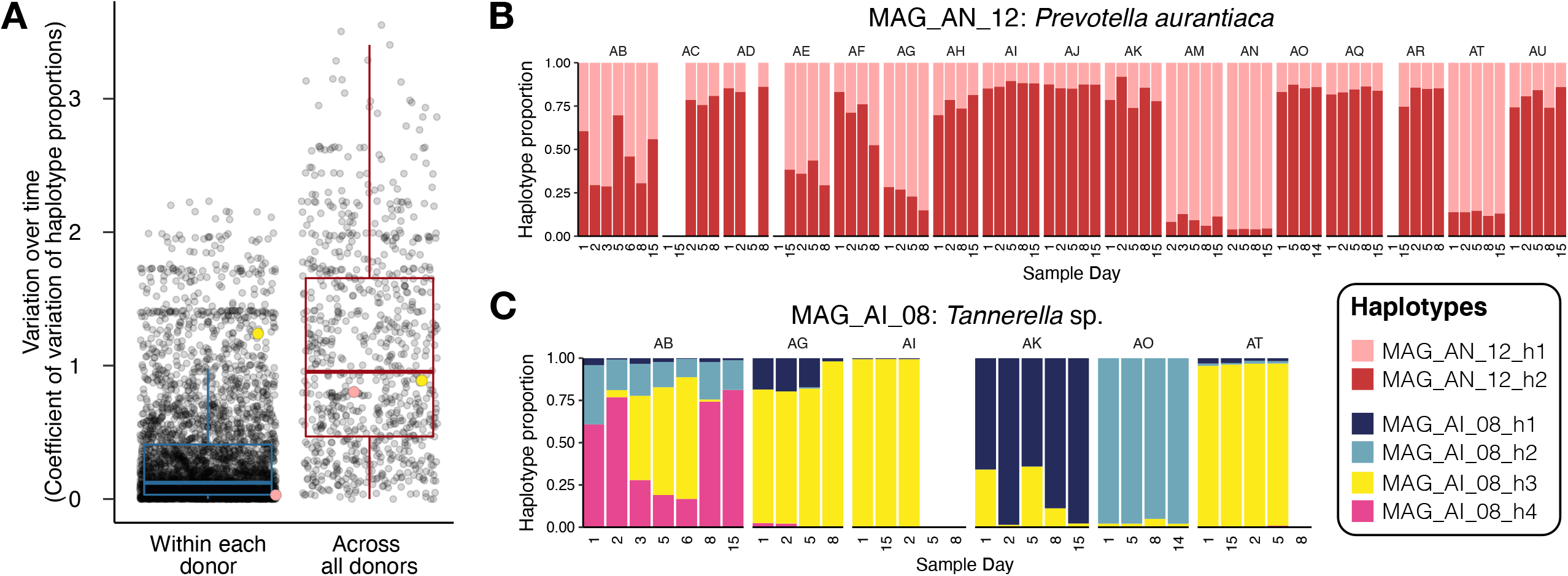
Haplotypes are generally stable but some sweep to dominance over a period of days. **A)** Coefficient of variation (y-axis) of haplotype proportions (proportion of parent MAG abundance) over time for each haplotype across all timepoints from all donors (red boxplot) or within a single donor for each donor with that haplotype (blue boxplot). Translucent points show the individual haplotype coefficients of variation summarized by the boxplots. **B)** Example of stable haplotype dynamics. The black bars on top show the coverage of the MAG_AN_12 while the colored bars show the proportions of the two haplotypes relative to total MAG_AN_12 abundance. Haplotype proportions are not shown for samples in which the MAG or its core single nucleotide variants could not be detected. The points in panel A corresponding to MAG_AN_12_h1 are colored peach; the peach point in the within-donor category is for MAG_AN_12_h1 in donor AM. **C)** Example of sweeping haplotype dynamics within MAG_AI_08. Same layout and detection criteria as in B. Yellow points in panel A correspond to MAG_AI_08_h3, with the yellow point in the within-donor category representing MAG_AI_08_h3 in donor AB.

Although haplotypes were generally more stable within a donor than across donors (Figure 2A), we also found examples of haplotypes that exchanged dominance over a period of days, e.g., those from MAG_AI_08, a 2.59 Mbp *Tannerella* sp. MAG with 95% completion and 0.7% redundancy (Figure 2C). For example, on Day 2, h4 accounted for approximately 75% of MAG_AI_08 coverage from volunteer AB, while haplotype 3 represented around 5% of the population. One day later the population was 50% h3 with just over 25% h4. By day 6, h3 was almost 75% of the population, and then by day 8, the relationship reverted and h4 was at 75% abundance with h3 barely detected. These haplotype dynamics are consistent with the ecotype hypothesis, which predicts that distinct genotypes exist within a population that change in abundance over time.

While some haplotypes dropped below the detection limit, some supplanted haplotypes remained at low abundance, indicative of a ‘soft sweep’ where a particular genotype rose to dominance without driving other genotypes to extinction, e.g., h3 and h4 of MAG_AI_08 in volunteer AB on day 6 (Figure 2C). This suggests that oral populations maintain a standing diversity of genotypes that may rise to dominance at a later time. The standing diversity of genotypes comprising a given population, i.e., the alpha diversity of subpopulations, differs both among populations and among mouths (Figure 2, Supplemental Figure 5) although sampling and sequencing depth may affect the resolution.

Correlating gene coverage to haplotype relative abundance within MAG_AI_08 allows inference into what genes are associated with a sweeping haplotype (Supplemental Figure 6A) to investigate whether the haplotype matches expectations for an ecotype. If a haplotype represents an ecotype, i.e., a distinct eco-evolutionary unit, then it should both have a unique set of genes that enable its success in a distinct niche. We define haplotype-associated genes to be those highly correlated (Pearson r > 0.8) with a haplotype but not so to other haplotypes (r < 0.2). Applying this threshold to the exemplar haplotype 4 of MAG_AI_08 revealed 155 genes associated specifically with this haplotype (Supplemental Table 2). These genes encompass a broad variety of functions from numerous COG categories, including metabolic genes such as a susD-like gene and an arginine decarboxylase, further supporting the hypothesis that this haplotype may represent a subpopulation-level ecotype with a unique complement of metabolic genes that allow for success in a specific niche (Supplemental Table 2, Supplemental Figure 6B). We also investigated whether the haplotype-associated genes were distributed throughout the genome, indicative of multiple independent events, or distributed as a single contiguous block, indicative of a single gene transfer event such as a phage or mobile element. Haplotype-associated genes in the case of MAG_AI_08_h4 were scattered across most of the 67 contigs with few haplotype-associated genes per contig (Supplemental Figure 6C). Altogether, the genes associated with this sweeping haplotype matched expectations for an ecotype.

### Gene-level sweeps are observed but rarely become fixed

Having detected genome-wide sweeps within the oral microbiota, we investigated the extent to which specific genes swept through populations. Operationally, we define gene sweeps as occasions where individual genes change in frequency within populations independent of their surrounding genomes. To detect such cases, we compared the coverage of each gene to the other genes in each MAG. The degree of correlation between genes informs about their linkage within the population, since genes on the same chromosome should correlate in coverage perfectly. Deviations in coverage between gene pairs, hereafter “decoupled” genes, implies the existence of features like copy number variation within the population or recent horizontal gene transfer into other population(s). Analogous to SNV patterns revealing linkage through recombination, the correlation of gene coverages over time reveals the linkage in frequency of genes within a population.

Gene abundances within populations tended to be stable and tightly correlated over days to weeks, with some notable exceptions (Figure 4). While the majority were stable, the frequencies of a small fraction of genes were strongly decoupled from those of other genes in the same MAG, as exemplified by MAG_AD_09 (Figure 4A), a *Gemella sanguinis* MAG detected across all donors (Supplemental Table 1). Forty-eight genes had coverages higher than the MAG by more than 5 standard deviations (Figure 4A red lines; Supplemental Figure 7). However, these changes were fleeting, with relatively few of the decoupled genes maintaining disproportionately high or low frequencies for the remainder of the sampling period, and genes were more tightly coupled within donors than across donors (Supplemental Figure 8). Thus, we infer that while most genes are tightly coupled within a population over time, some genes are decoupled from their populations and warrant further investigation.

**Figure 3.**
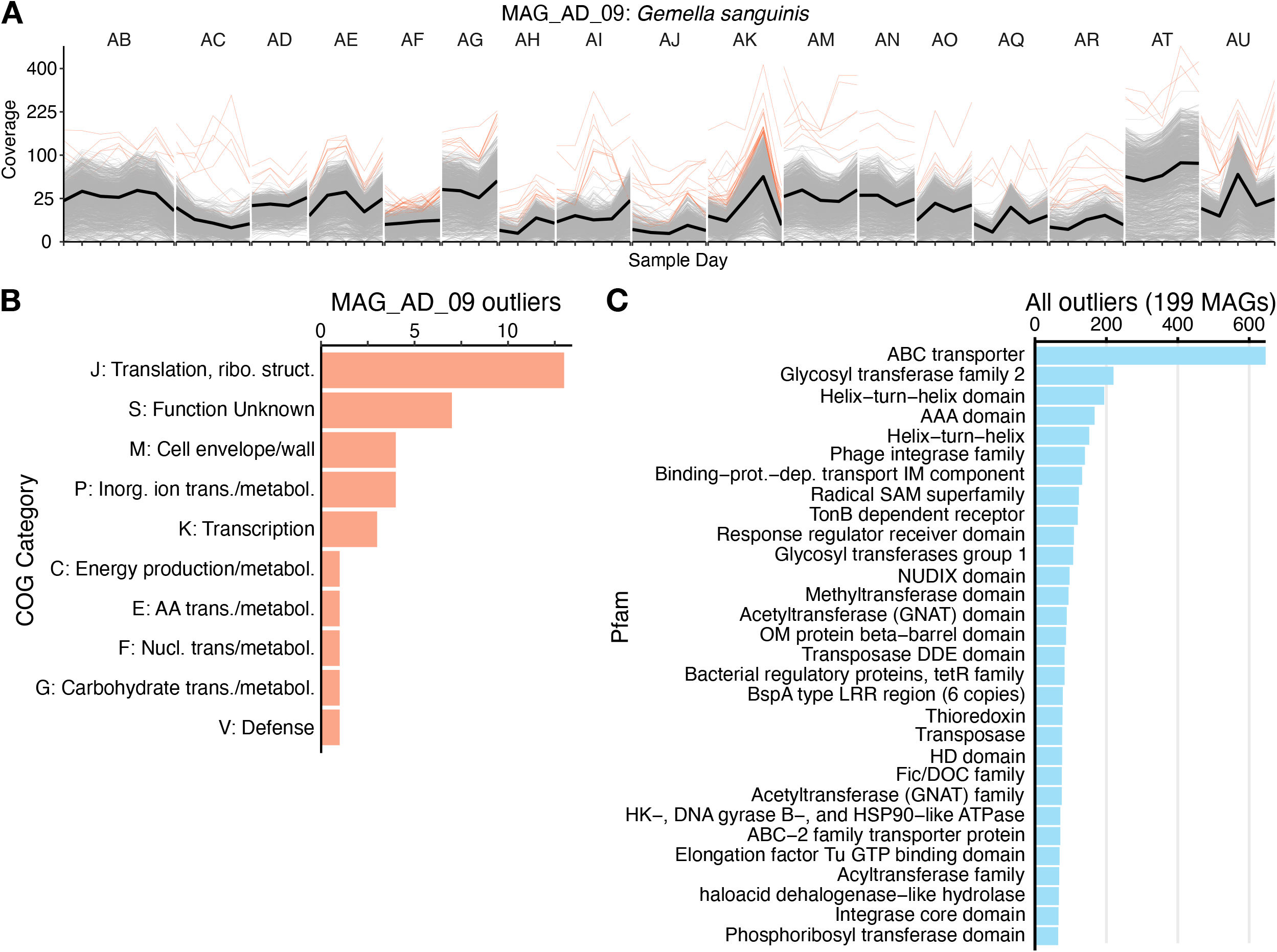
Most genes in a population are tightly correlated within a single mouth over time, but less so between mouths over time. **A)** Coverage of each gene (thin grey or orange lines) in an example MAG, MAG_AD_09, across timepoints (x-axis). The thick black line shows the mean MAG coverage. In orange are decoupled genes, being 5 standard deviations or more above the average gene’s coverage difference from the MAG. **B)** Counts (x-axis) of COG categories (y-axis) occurring in the 48 decoupled MAG_AD_09 genes (‘outliers’). Unannotated genes are omitted. **C)** Counts (x-axis) of 30 most frequent Pfam functions (y-axis) in decoupled genes (‘outliers’) obtained from all 199 MAGs with at least 50% of their nucleotides covered in at least 15 samples. Unannotated genes are omitted.

**Figure 4.**
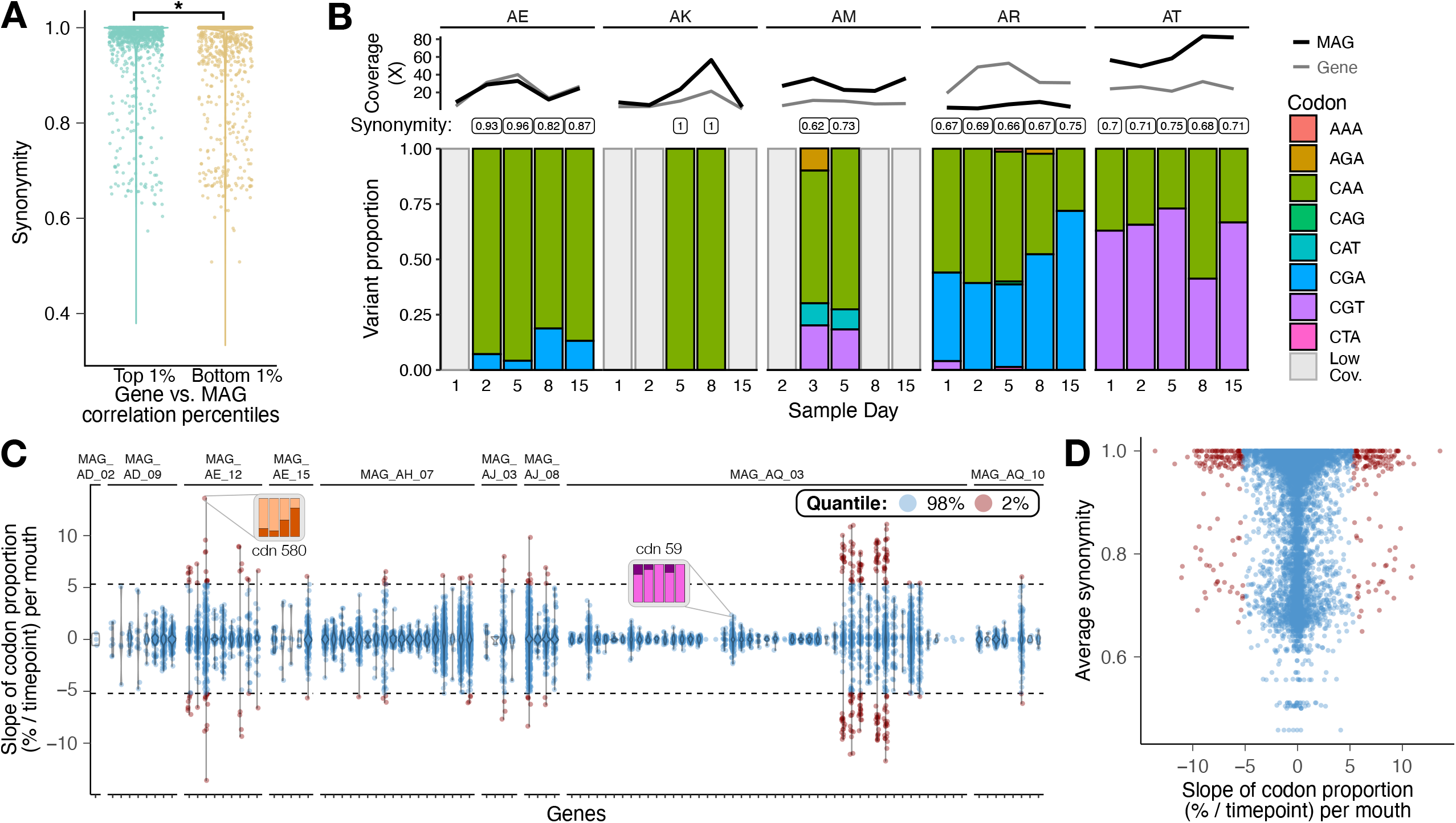
Changes in codon synonymity do not explain decoupled genes. **A)** Tightly coupled genes have higher synonymity than do uncoupled genes. Synonymity (y-axis), the fraction of synonymous pairs out of all codon variants in the metagenome, is shown for all single codon variants detected in each metagenome for the top (green) and bottom (brown) 1% of genes based on their correlation with their parent MAGs across mouths. Each dot corresponds to a single codon position in a single metagenome. **B)** The proportion of each SCV within a single codon variant site is stable over time. Each colored bar shows the proportion of a different codon triplet at each time point with >10x coverage for a single exemplar codon position in MAG_AD_09. Grey bars (“Low cov.”) mark samples with insufficient coverage for calling SCVs. The synonymity is shown in the text bubble above each bar. Above the colored bars, the black and grey lines show the coverage of the MAG and the parent gene, respectively. SCV proportions are only shown for donors with two or more samples meeting the SCV coverage threshold. **C)** SCV proportions are stable for most genes despite changes in gene frequency. The change in codon proportion over time was determined from the slope of the line of best fit for codon positions with at least 10x coverage in at least 4 timepoints. Each dot shows the slope (y-axis) of a single codon for one variant codon position in one donor. The slope is in units of proportion per timepoint. Slopes are separated by gene (x-axis), grouped by MAG. Dot color shows whether the slopes belong to the central 98^th^ percentile (blue) or the upper and lower 1^st^ percentiles (red). The changes in codon proportion are shown in call-outs for two arbitrary codons, displayed as in 4B, to place the slope values in context. **D)** Directionally changing SCVs generally have high synonymity. For each SCV slope in (C), the average synonymity (y-axis) for that codon position in that donor was determined. Quantiles and their colors reflect the distribution of slope proportions from (C).

Inspecting the predicted functions of the 48 decoupled genes in MAG_AD_09 revealed a combination of highly conserved genes like translation machinery along with a surprising diversity of other functions (Figure 4B). The highly conserved functions, e.g., ribosomal proteins, suggest the outlier coverage could be explained by reads originating from multiple closely related populations. Since the metagenomic read recruitment was competitive (i.e., each metagenomic read was mapped against all 390 MAGs), the different populations must be related such that these slowly evolving housekeeping genes are practically identical in nucleotide sequence while the surrounding genome has diverged enough to not be recruited. Thus, these decoupled housekeeping genes likely do not represent horizontal gene transfer or genes under positive selection. Beyond these housekeeping genes, cell-wall-related functions were the third-most frequent category, followed by other metabolic and/or membrane categories including carbohydrate metabolism and energy production (Figure 4B). Inspection of the Pfam annotations for the genes belonging to these categories confirms that they include functions like an FMN-dependent dehydrogenase and an acyltransferase (Supplemental Table 3). Potentially, these non-housekeeping genes could represent horizontal gene transfer (HGT) events into other populations recent enough that the transferred genes have not yet diverged. If so, these functions would represent the types of functions likely to undergo HGT in the tongue microbiome. Further, some of these categories may include phage- and mobile-element related functions, illuminating the diversity of mechanisms for HGT among oral populations.

To scale our inference of tight correlation between genes in a population from an example MAG to all MAGs in our dataset, we compared gene coverages to MAG coverages to identify similar outliers across the dataset (Figure 4C). Gene coverages were compared only for the 199 MAGs at least 50% of their length receiving coverage in at least 15 samples. Inspection of the 30 most frequent Pfam functions in the set of decoupled genes revealed an enrichment for phage and mobile elements, confirming their importance to transferring genes between populations. However, other functions like thioredoxins and ABC transporters appear across among decoupled genes. As these gene families are not generally as conserved as key housekeeping genes like ribosomal proteins or elongation factors, their membership in the decoupled set is likely not due to sequence-level conservation but instead recent horizontal gene transfer.

### Codon-level variation does not explain gene frequency changes

Having detected a subset of genes decoupled from the majority of genes within a population, i.e., within a MAG, we wanted to investigate whether the decoupled genes showed features associated with selection. To accomplish this we compared synonymity, the ratio of synonymous to non-synonymous single codon variants (SCVs) within each metagenome (see Methods) for the genes producing the top and bottom 1% of gene correlations vs. their parent MAGs across all mouths (Figure 5A), for all genes from the 25 MAGs with 75% of their nucleotides detected in ?40 samples. Synonymity was generally high, with the majority of codon variation being completely synonymous; however, the genes from the bottom 1% had slightly higher proportions of nonsynonymous SCV pairings within the metagenomes (lower synonymity) than the top 1% (Wilcoxon rank sum test, p = 1E-16). This result was robust to whether the top and bottom 1% or 10% of genes were used (Supplemental Figure 9). Thus, although changes in individual gene frequencies were not sustained, decoupled genes were slightly enriched in non-synonymous codon variants.

Although we did not observe sustained turnover at the level of gene coverage, certain nonsynonymous SCVs could confer selective advantage. If so, then such mutations might be predicted to sweep the population over time. To address this scenario, we compared the proportional changes of each codon within variant sites covered at least 10x in the highly decoupled genes producing the bottom 1% of pairwise correlations (Figure 5B-D). Focusing on an exemplar codon position from one such highly decoupled gene of unknown function from MAG_AD_09 (Figure 5B), synonymity was indeed generally high in mouths where the gene’s frequency matched the surrounding genome (donors AE, AK) but lower in mouths where gene and genome frequencies diverged (donors AM, AR, AT). However, despite lower synonymity, the proportion of each codon remained relatively constant. If such codon-level stability can be generalized beyond this example, then perhaps under these circumstances the nonsynonymous variants do not enjoy a selective advantage over the other variants.

To address the generality of this pattern of stable SCV proportions, we investigated the slopes of the proportion of each codon variant in each donor (Figure 5C), i.e., the rate of change for each codon variant. The slopes for each SCV were generally close to zero, with 98% of the slopes being between approximately −0.05 and 0.05. Thus, we infer that the majority of the observed codons in these decoupled genes are not in the process of sweeping but instead represent standing genetic variability within the population. To support this inference, we investigated the trend of synonymity with respect to each SCV’s slope (Figure 5D), reasoning that SCVs with the highest slopes, i.e., those changing in proportion the fastest, should also have low synonymity if the directional change of certain SCVs was due to a selective advantage over competing SCVs. However, we did not observe this pattern – instead, the majority of SCVs undergoing the strongest directional change had high synonymity, and the lowest synonymity occurred stable SCVs (Figure 5D). Therefore, we infer that the majority of observed changes in gene and nucleotide frequency are not due to direct selection.

## Discussion

Previous studies of the oral microbiome have introduced and supported the concept of ‘dynamic stability’ where populations defined by 16S rRNA amplicon sequence variants (ASVs) fluctuate on daily timescales around a mean that is stable over time (Hall et al., 2017; Utter et al., 2016; Wake et al., 2016). However, these studies were constrained by the resolution limits of 16S ASVs and the lack of genomic context to support or reject the extrapolation of an ASV to mark a distinct population. Metagenomic surveys have dramatically increased the analytical resolution to reveal increasing levels of diversity (Donati et al., 2016; Lloyd-Price et al., 2017); however, they have not yet been applied to investigate the pattern of dynamic stability within individuals at microbially-relevant, daily timescales. Our observations of fluctuating yet generally stable haplotypes within populations are consistent with the concept of ‘dynamic stability’ derived from amplicon studies, but now broaden the basis for that conclusion to entire genomes. For an individual tongue, most microbial genomes, although varying in relative abundance, were stably present over the two-week sampling period. Within populations, in cases where directional change of population variants within a MAG interrupted the pattern of dynamic stability, specific genes were associated with the divergent haplotypes, revealing the genetic distinction between these subpopulations.

The observed patterns of gene and genome-wide dynamics distinguish the oral microbiome from the healthy gut, where gene sweeps account for most of the change (Garud et al., 2019) and cultivation and SNV analyses reveal successive strain replacement or stable cooccurrence (Zhao et al., 2019). The timescales of Zhao et al. are longer than the two weeks surveyed here, so the possibility remains that the fluctuating haplotypes observed in our study may eventually completely sweep the population. In the antibiotic-challenged gut, however, entire subpopulations exchanged abundances within some populations over a long-term time course experiment (Roodgar et al. 2021), revealing the existence genome-centric population changes akin to those reported in the mouths of antibiotic-free individuals. Others have highlighted the distinct differences between the oral and gut microbiomes in terms of composition, dynamics, responsiveness to diet (David et al., 2014); the results presented here add sub-population dynamics to the list. The population dynamics of the human oral microbiota appear more similar to environmental systems in which genome-wide selection events are known to occur (Bendall et al., 2016; Goyal et al. 2021). Yet, adaptation in other non-human environmental microbes like *Rhizobium leguminosarum* are reported to be dominated by gene- or gene-cluster-specific recombination events (Klinger, Lau, & Heath, 2016), proving that no one-size-fits-all pattern exists for bacterial population adaptation.

Although metagenomics can identify and characterize haplotypes, existing bioinformatic approaches alone cannot definitively prove that the observed haplotypes represent ecotypes. Haplotypes with distinctive abundance patterns allow the identification of genes associated with a particular haplotype, from which genetic dissimilarity can be inferred, as was the case for MAG_AI_08_h3. However, haplotypes without distinctive abundance trajectories cannot be similarly addressed with confidence. As such, we did not similarly characterize the number and nature of genes associated with stable haplotypes, e.g., the genetic distinction between the MAG_AN_12 haplotypes. However, a similar study by Goyal et al. (2021) found that in pitcher-plant-derived communities, as few as 100 SNPs throughout the genome distinguished ecologically distinct subpopulations. Assuming these estimates hold true for oral-dwelling bacteria, many of these haplotypes likely represent ecologically distinct subpopulations, i.e., ecotypes. If haplotypes are assumed to represent ecotypes, then the stable co-occurrence of multiple haplotypes within a mouth over time implies co-existence or partitioning among ecotypes. Such stable co-occurrence among subpopulations has been documented previously in bacterial communities at varying phylogenetic scales (Good et al. 2017, Zhao et al. 2019, Roodgar et al. 2020, Goyal et al. 2021). Alternatively, the two-week time course may not have afforded the proper conditions to reveal divergent ecology or changes in the competitive advantage.

Recent metagenomic studies have highlighted the importance of mobile genetic elements like phage and transposons for population-level evolution (Hackl et al., 2020; Kent et al., 2020; Zlitni et al., 2020). While mobile elements did not feature strongly in our data, this may be due to methodological choices designed to increase our confidence in MAG quality at the expense of biasing against retaining mobile elements due to difficulties in confidently assigning them to a single population based solely on differential coverage and tetranucleotide frequency (Quince et al. 2017). Recent sequencing innovations such as Hi-C (chromatin conformation capture) offer promise for solving this problem (Roodgar et al., 2019).

Different definitions of ‘population’ for bacteria have been given in the literature (Delmont et al., 2019; Shapiro, 2018). Populations have been defined by the existence of a unifying niche (Cohan & Perry, 2007), sequence similarity thresholds obtained from marker gene or whole genome alignments (Jain et al., 2018; Konstantinidis & Tiedje, 2005; Stackebrandt & Goebel, 1994), measured or modeled recombination frequency (Marttinen et al., 2015; Shapiro et al., 2012), or operationally based on geographic or seasonal distribution (Hunt et al., 2008). Here, we use ‘population’ to refer to a set of phylogenetically-related lineages defined metagenomically. This definition has utility for our analyses as it allowed us a reference to relate populations across mouths, e.g., we found that the *Prevotella aurantiaca* population defined by MAG_AN_12 in donor AM was dominated by a different subpopulation than in donor AN. While each definition has strengths and weaknesses, the operational definition of a population as a MAG and the short reads that map to it is conceptually and practically useful – this definition is operationally clear, it is approximate to the species concept, and it allows for the existence of subpopulations with potentially adaptive and distinct genetic features.

The existence of subpopulations within our MAGs reveals not only the current technical limitations of contig-binning approaches to reconstructing MAGs but also the limitations in our conceptual understanding of bacterial genomes. While poorly constructed MAGs can have extreme, artificial the genetic heterogeneity arising from methodological errors during assembly, e.g., Shaiber et al. (2019), in many cases the heterogeneity occurs from the reality of multiple closely-related lineages coexisting within an ecosystem (Quince et al. 2017b). Given the amount of heterogeneity known from populations of cultured and environmental genomes, (Tettelin et al. 2005, Utter et al. 2020, Shaiber et al. 2020), the limitations of current metagenomic sequencing approaches force even the best MAGs to be composite to a small degree. Indeed, given the ability of microbial populations to rapidly diversify into distinct subpopulations (Good et al., 2017), perhaps the existence of a singular genome is an artifact of isolation and a consensus genome is the more useful genomic descriptor for populations in their native habitats. The question facing microbial ecologists is not so much which genome is an accurate single proxy for a population but rather how the extant variations of that genome impact microbial ecology.

The coexistence of soft sweeps both at the gene and genome level suggests that both processes work to shape oral bacterial populations. While gene-specific and genome-wide sweeps fall under different concepts for species and population evolution (Garud & Pollard, 2020), bacterial evolution is not constrained to the kinetics predicted by a single species concept (Shapiro, 2018). Our data suggest that both mechanisms of genomic change are relevant to the oral microbiome. We propose that the most impactful adaptive mode is genome centric – ecotypes sweep to dominance in response to environmental changes. In addition, changes in gene frequency also occur via horizontal gene transfer or other mechanisms, presenting a means of generating or modifying genetic diversity among ecotypes over longer timescales. While we suggest this model in the context of the oral microbiome, we anticipate the broader application of high temporal and genomic resolution to other microbial systems will bring about a more complete understanding of microbial adaptation.

---

**Supplemental Table 1**. Taxonomic assignment for each MAG along with summary metrics (length in bp, estimated completion and redundancy). The second tab (“Detection”) reports the fraction of each MAG recruiting any coverage for all samples, and the third tab (“Coverage”) reports the coverage of each MAG in each sample.

**Supplemental Table 2**. Predicted functions for the 155 genes associated with MAG_AI_08_h4.

**Supplemental Figure 1**. Number of MAGs binned by genus (y-axis).

**Supplemental Figure 2. A)** Fraction of MAG covered at all (shading intensity) for the all 390 MAGs (rows), with the rows (MAGs) arranged to cluster MAGs with similar fraction covered in each metagenome (columns). Each column shows an individual sample, grouped by donor. Only fractions >=0.5 are shown since we consider MAGs with less than half of their nucleotides covered to not be confidently detected. Red boxes mark the samples for each MAG that were obtained. **B)** Same data and organization as in A but not considering the fraction of MAG covered for samples used to bin each MAG prior to ordering.

**Supplemental Figure 3**. Distribution of MAGs recruiting reads over at least 50% of their length. Each of the 390 vertical bars is a different MAG, ranked in descending order, and the height of the bar shows number of samples providing 50% detection (up to 81 samples).

**Supplemental Figure 4**. Histogram showing the distribution of the number of DESMAN-deconvolved haplotypes across all MAGs.

**Supplemental Figure 5**. DESMAN haplotype decomposition for each MAG. The stacked, colored bars show the proportion (left y-axis) of each haplotype (colors) in each sample, grouped by participant. The black line shows the coverage of the parent MAG (right y-axis).

**Supplemental Figure 6**. **A)** Correlation of gene coverage vs. haplotype relative abundance for all genes (x-axis) and haplotypes (colors) in MAG_AI_08. Genes are arranged to group similar correlation patterns, not any chromosomal order. **B)** Histogram of COG categories for the 163 genes associated with MAG_AI_08_h4. C) Number of genes per MAG_AI_08 contig associated with haplotype 4 (pink) or not (grey) showing haplotype-associated genes are well distributed.

**Supplemental Figure 7**. Distribution of detrended gene coverages. The coverage deviation (x-axis) between each gene in MAG_AD_09 and the MAG’s average coverage is plotted as a density distribution using all samples from each donor (rows). Dashed red vertical lines mark 5 standard deviations above the mean, the threshold used in Figure 4.

**Supplemental Figure 8**. Full distribution of the pairwise gene correlations summarized in Figure 4B. For each MAG with 75% of their genome covered in 5, 10, 20, 30, and 40 samples (subpanel rows), the Pearson correlation (x-axis) over time for all gene pairs were calculated mouth-by-mouth (blue distributions; only for ?20 sample detections) or across mouths (orange). A random subset of 1 million correlations were subsampled from each set and used to calculate the probability density distribution (y-axis). The vertical line in each distribution marks the median correlation coefficient, and the translucent color spans the second to the third quartile.

**Supplemental Figure 9**. Differences in synonymity between coupled and uncoupled genes are robust to whether coupled and uncoupled are defined by the extreme 1% or 10% of pairwise gene correlations. Synonymity (y-axis), the fraction of synonymous pairs out of all codon variants in the metagenome, is shown for all single codon variants detected in each metagenome for the top and bottom 1% of genes based on their correlation with their parent MAGs across mouths (identical to Fig. 5A). Adjacent to these data are shown the synonymities from the top and bottom 10% of genes to show that the observation is robust to using 1% or 10% thresholds. Each dot corresponds to a single codon position in a single metagenome.

## Declarations

### Ethics approval and consent to participate

All samples were collected following IRB approval and oversight (IRB16-0367)

### Consent for publication

Not applicable

### Availability of data and materials

The raw metagenomic data used in this study will be publicly available at NIH NCBI SRA upon manuscript publication. Analyzed data in the form of anvi’o databases can be found at https://doi.org/10.6084/m9.figshare.14597832. A reproducible methods document at https://dutter.github.io/projects/diversity_dynamics provides the code used for all analyses.

### Author contributions

D.R.U., C.M.C., and G.G.B designed research; D.R.U, C.M.C., and G.G.B performed research and analyzed data; D.R.U, C.M.C., and G.G.B wrote the paper.

## Acknowledgements

We thank I-Ting Huang for helpful discussions. We thank the many volunteers who donated samples. The content is solely those of the authors and does not necessarily reflect the views of Harvard Catalyst, Harvard University and its affiliated academic health care centers, the National Science Foundation, or the National Institutes of Health.

