## Supplemental Figure 5 for "Gene- and genome-centric dynamics shape the diversity of oral bacterial populations"

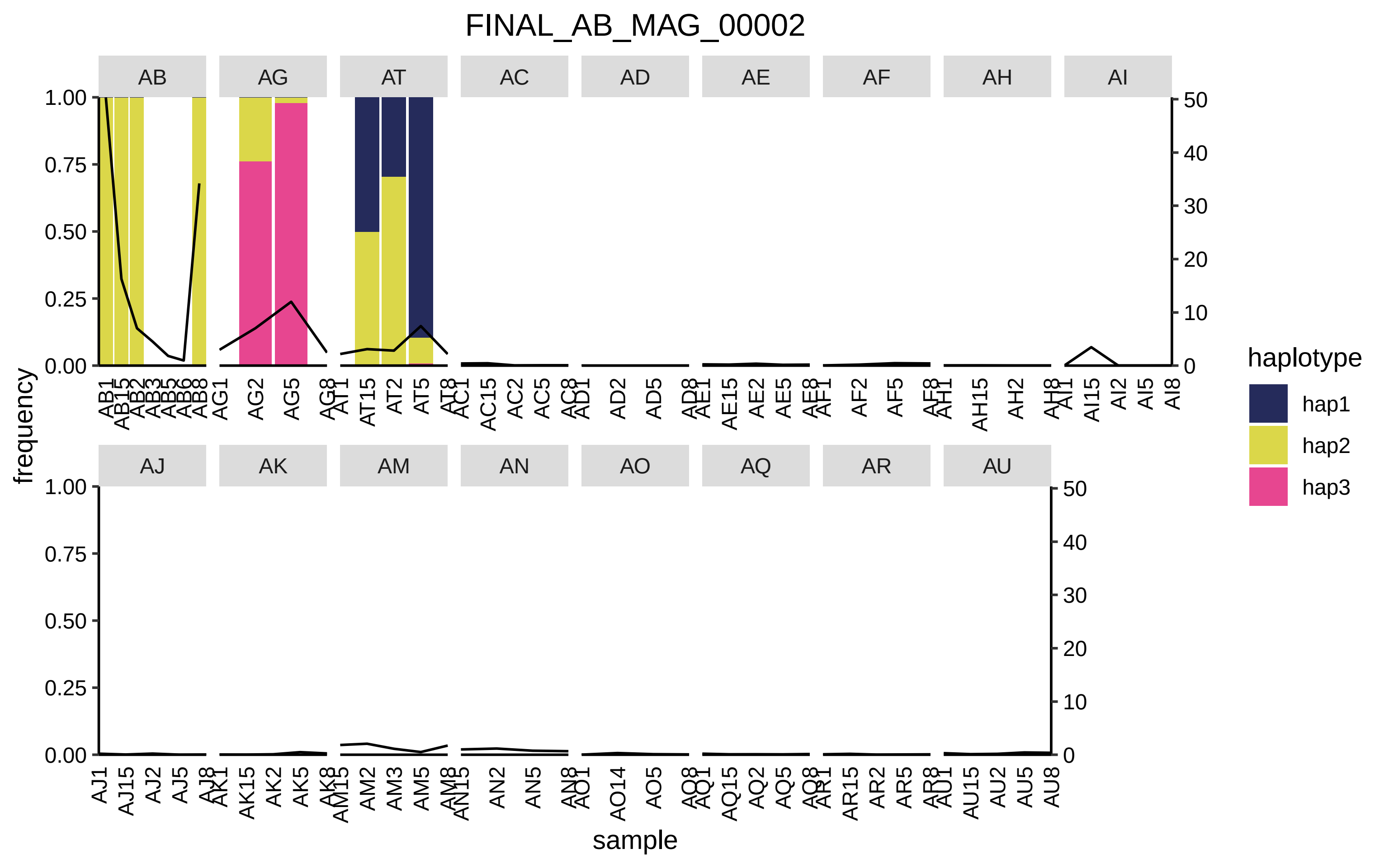

FINAL_AB_MAG_00002
AB
AG
AT
AC
AD
AE
AF
AH
AI
AJ
AK
AM
AN
AO
AQ
AR
AU
1.00
0.75
0.50
0.25
0.00
50
40
30
20
10
0
frequency
sample
haplotype
hap1
hap2
hap3

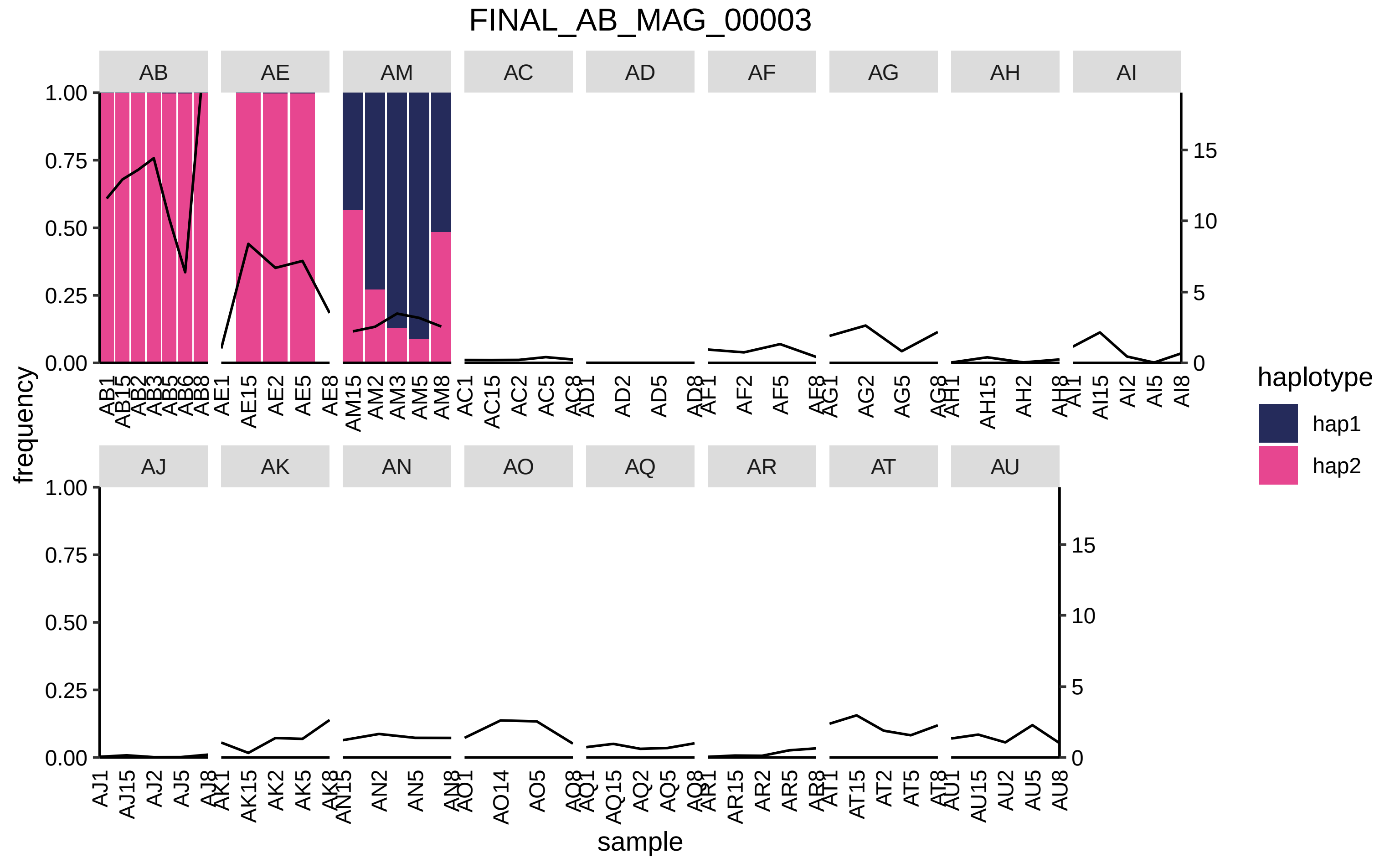
FINAL_AB_MAG_00003
AB
AE
AM
AC
AD
AF
AG
AH
AI
AJ
AK
AN
AO
AQ
AR
AT
AU
frequency
sample
haplotype
hap1
hap2

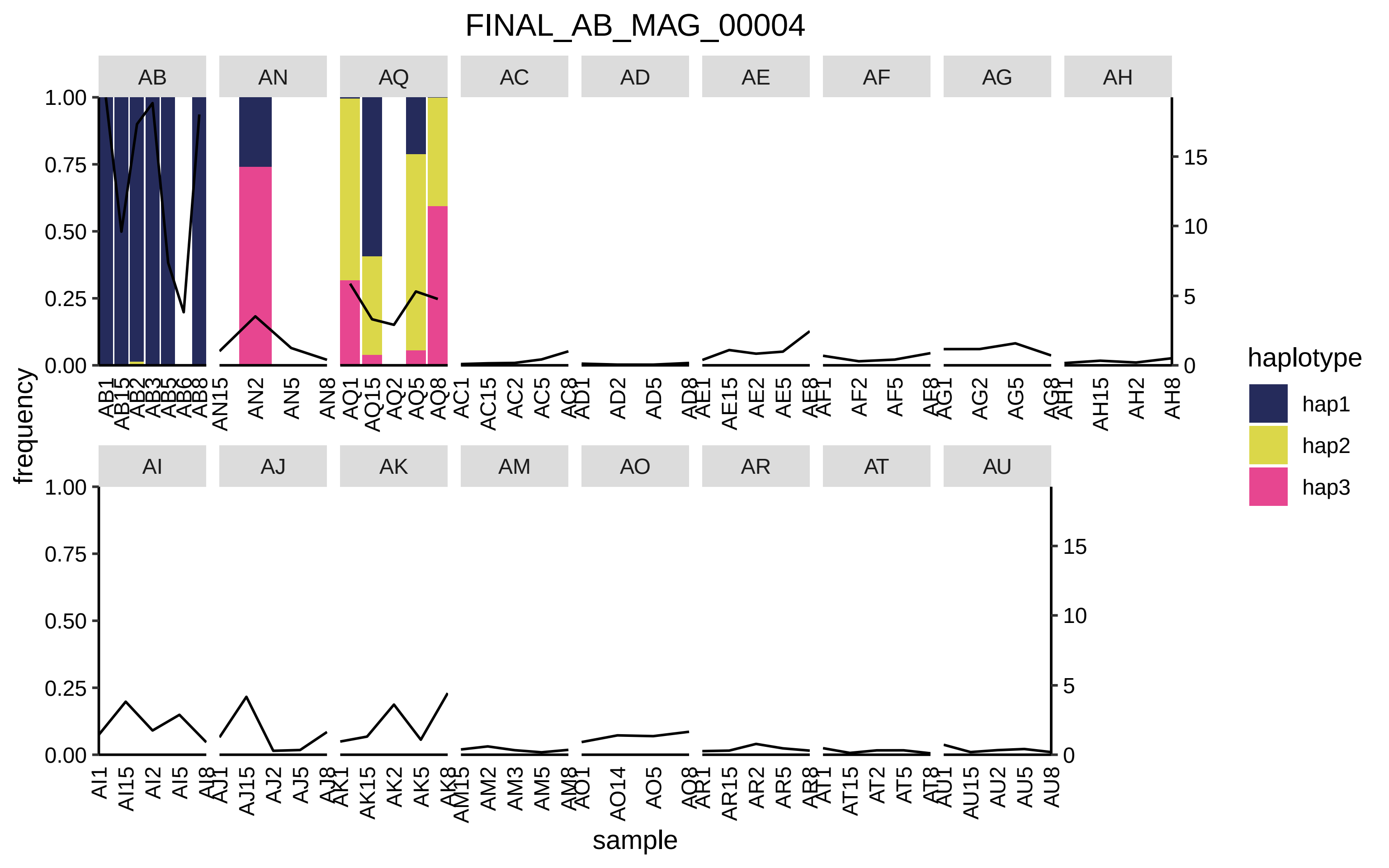
FINAL_AB_MAG_00004
AB
AN
AQ
AC
AD
AE
AF
AG
AH
AI
AJ
AK
AM
AO
AR
AT
AU
1.00
0.75
0.50
0.25
0.00
15
10
5
0
frequency
sample
haplotype
hap1
hap2
hap3
AB1 AB15 AB2 AB3 AB5 AB6 AB8
AN15 AN2 AN5 AN8
AQ1 AQ15 AQ2 AQ5 AQ8
AC1 AC15 AC2 AC5 AC8
AD1 AD2 AD5 AD8
AE1 AE15 AE2 AE5 AE8
AF1 AF2 AF5 AF8
AG1 AG2 AG5 AG8
AH1 AH15 AH2 AH8
AI1 AI15 AI2 AI5 AI8
AJ1 AJ15 AJ2 AJ5 AJ8
AK1 AK15 AK2 AK5 AK8
AM15 AM2 AM3 AM5 AM8
AO1 AO14 AO5 AO8
AR1 AR15 AR2 AR5 AR8
AT1 AT15 AT2 AT5 AT8
AU1 AU15 AU2 AU5 AU8

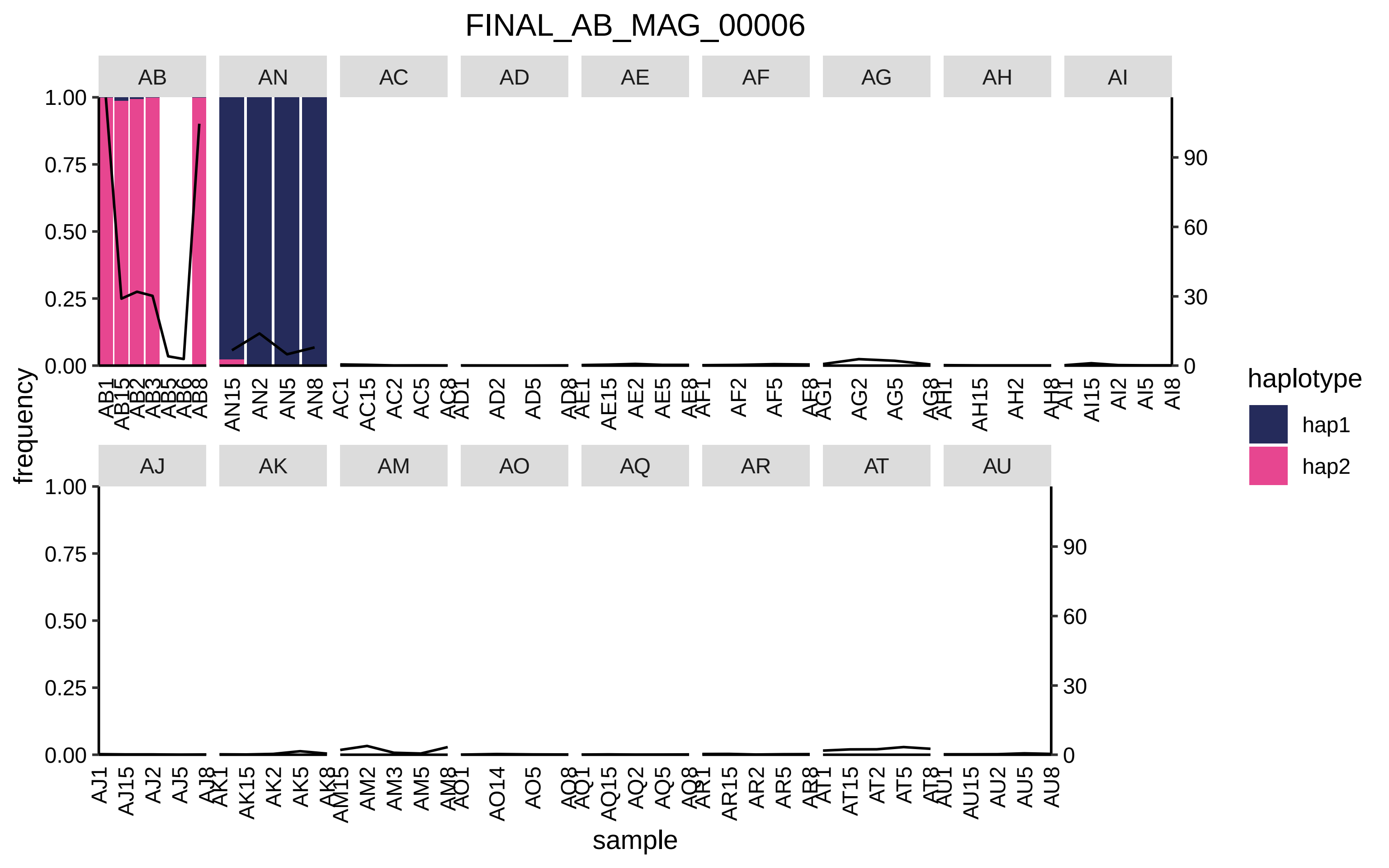

FINAL_AB_MAG_00006
AB
AN
AC
AD
AE
AF
AG
AH
AI
AJ
AK
AM
AO
AQ
AR
AT
AU
frequency
sample
haplotype
hap1
hap2

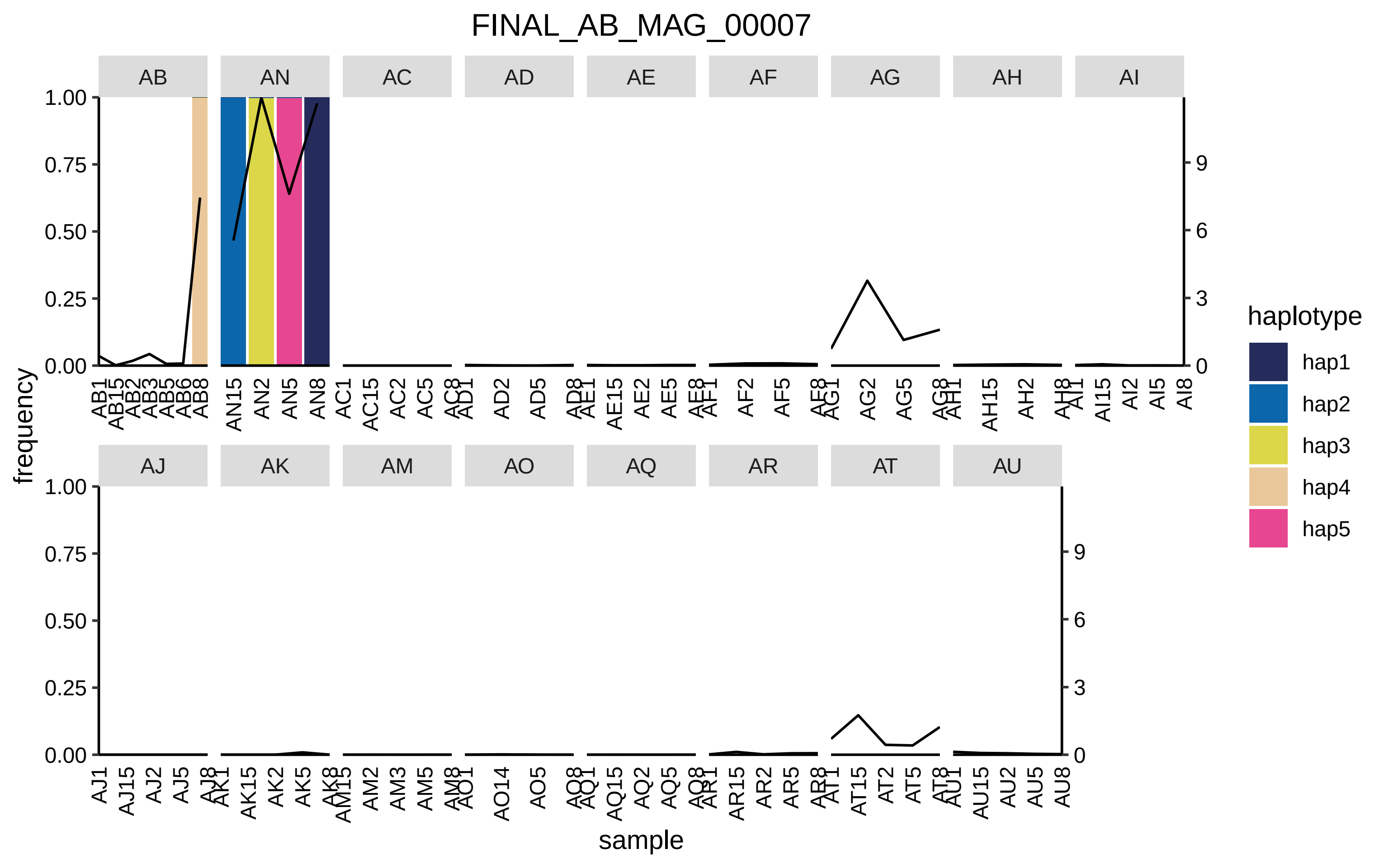

FINAL_AB_MAG_00007
AB
AN
AC
AD
AE
AF
AG
AH
AI
AJ
AK
AM
AO
AQ
AR
AT
AU
frequency
sample
haplotype
hap1
hap2
hap3
hap4
hap5
1.00
0.75
0.50
0.25
0.00
9
6
3
0

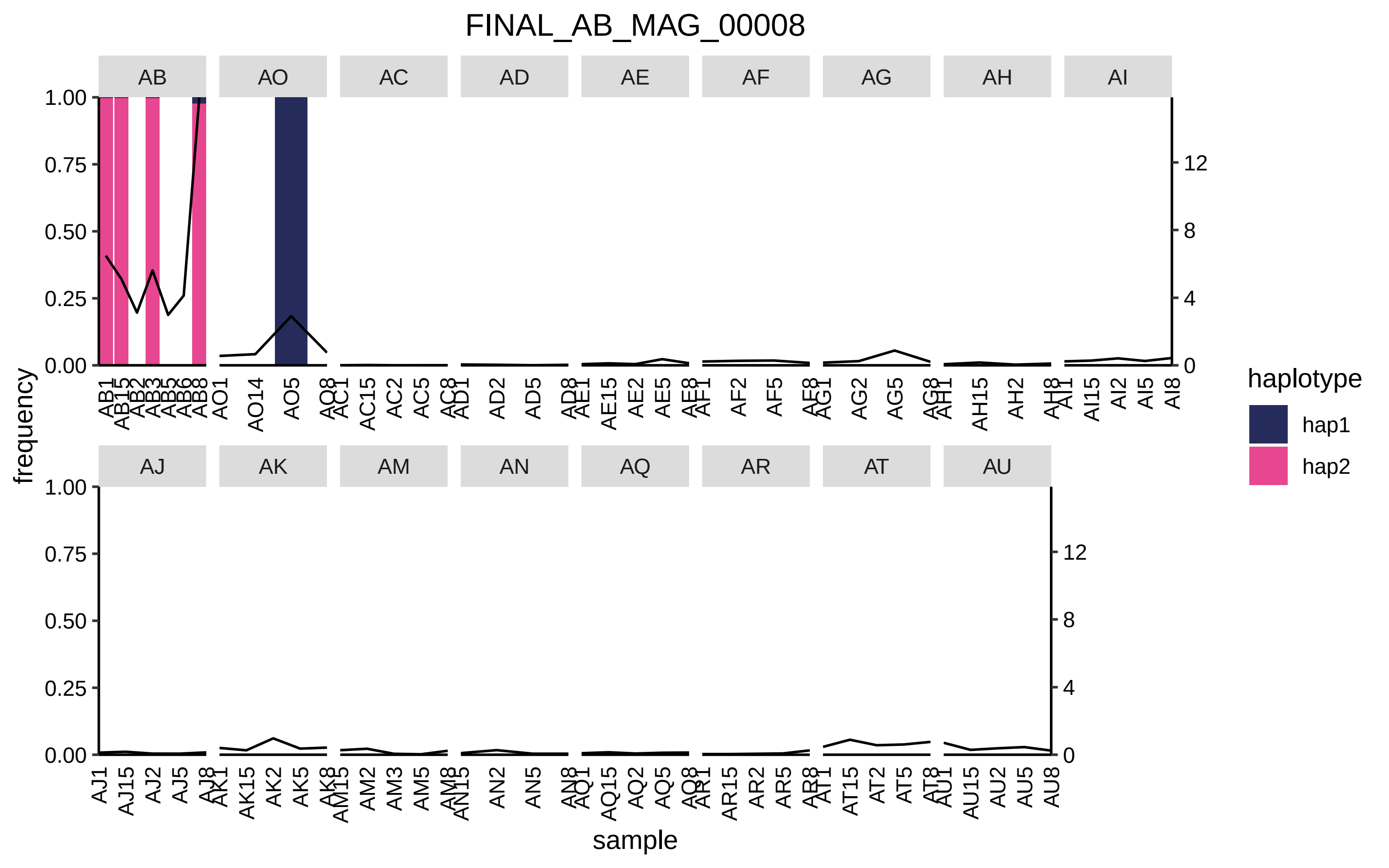
FINAL_AB_MAG_00008
AB
AO
AC
AD
AE
AF
AG
AH
AI
AJ
AK
AM
AN
AQ
AR
AT
AU
frequency
sample
haplotype
hap1
hap2

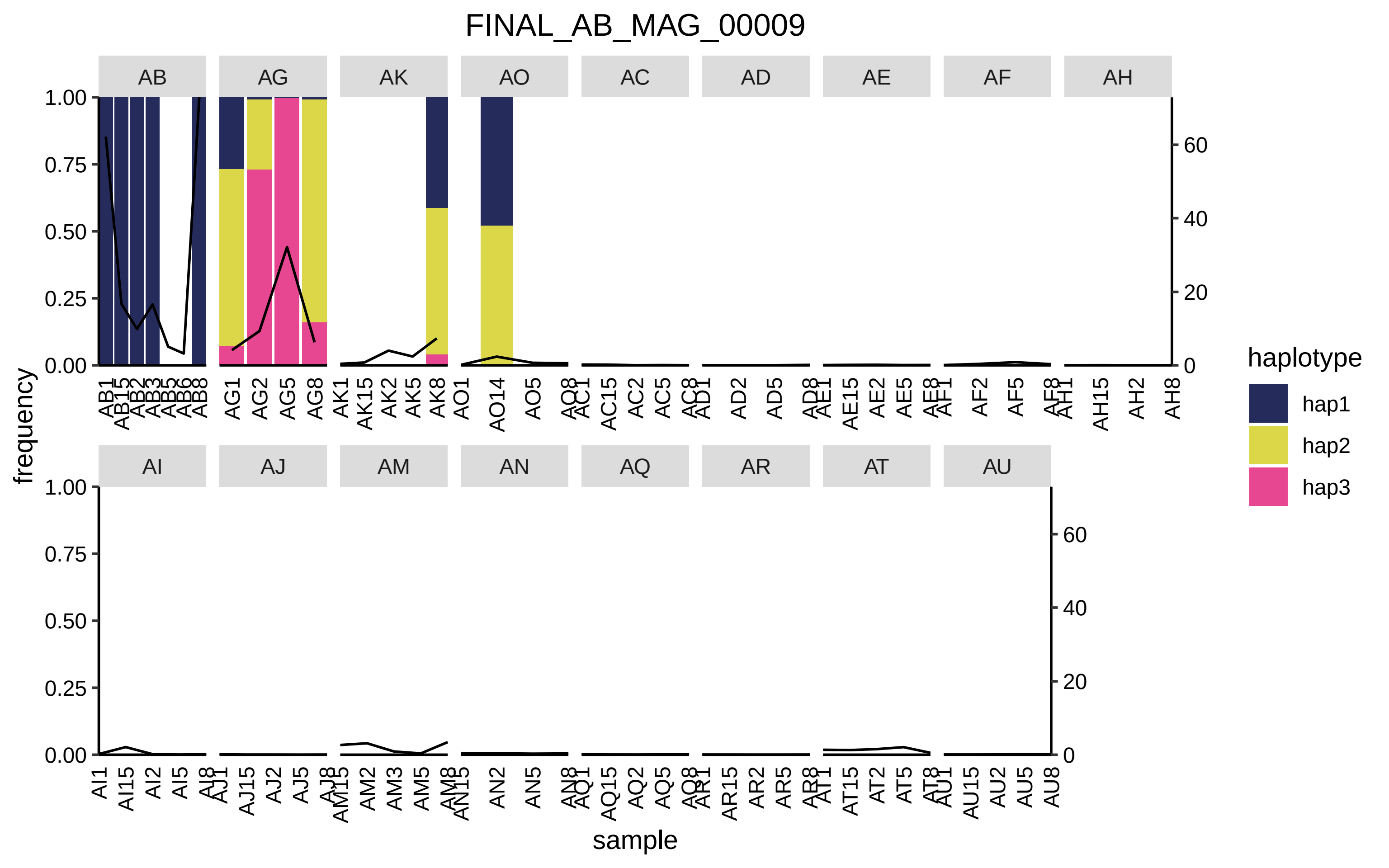
FINAL_AB_MAG_00009
AB
AG
AK
AO
AC
AD
AE
AF
AH
AI
AJ
AM
AN
AQ
AR
AT
AU
frequency
sample
haplotype
hap1
hap2
hap3

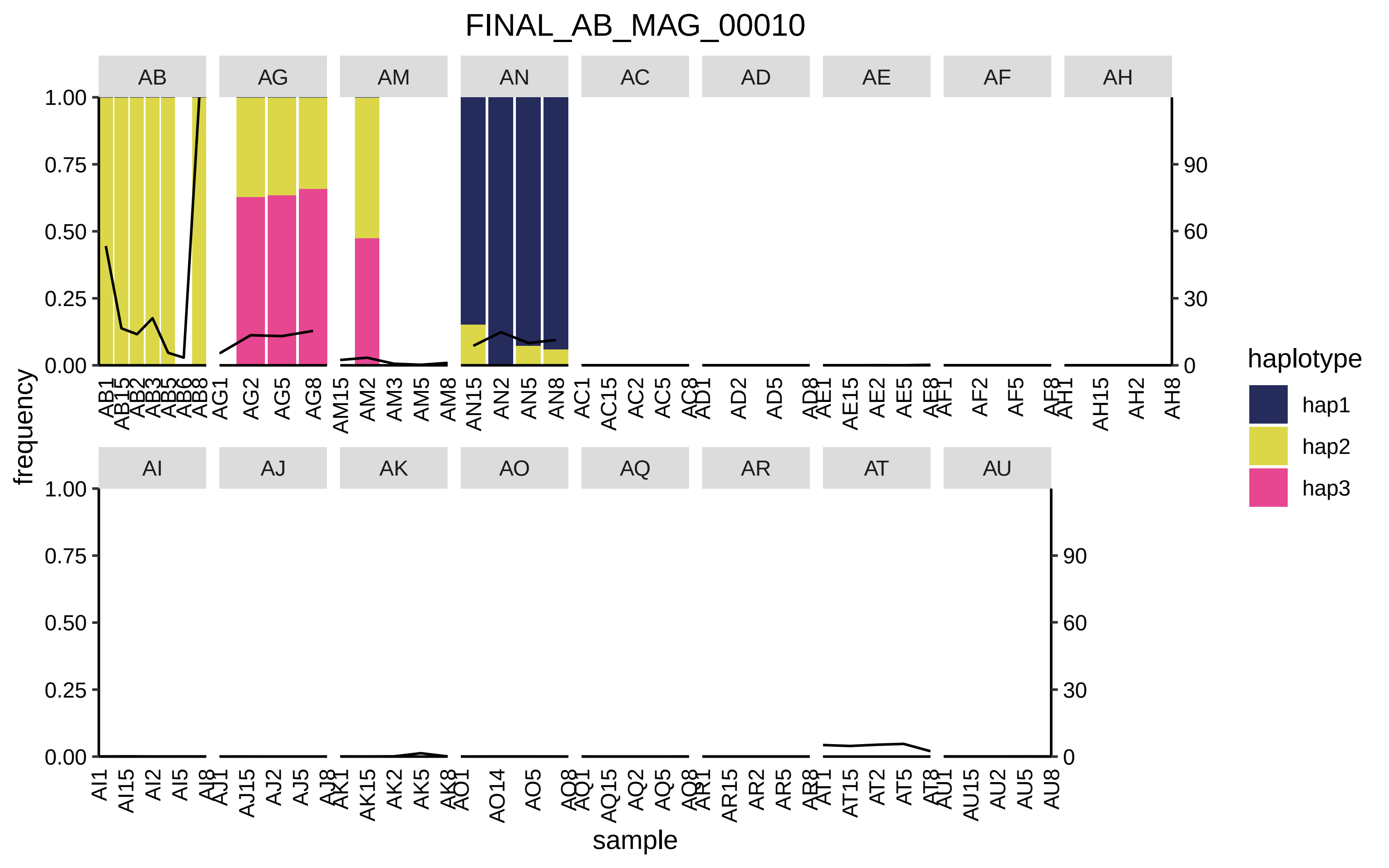

FINAL_AB_MAG_00010
AB
AG
AM
AN
AC
AD
AE
AF
AH
AI
AJ
AK
AO
AQ
AR
AT
AU
frequency
sample
haplotype
hap1
hap2
hap3

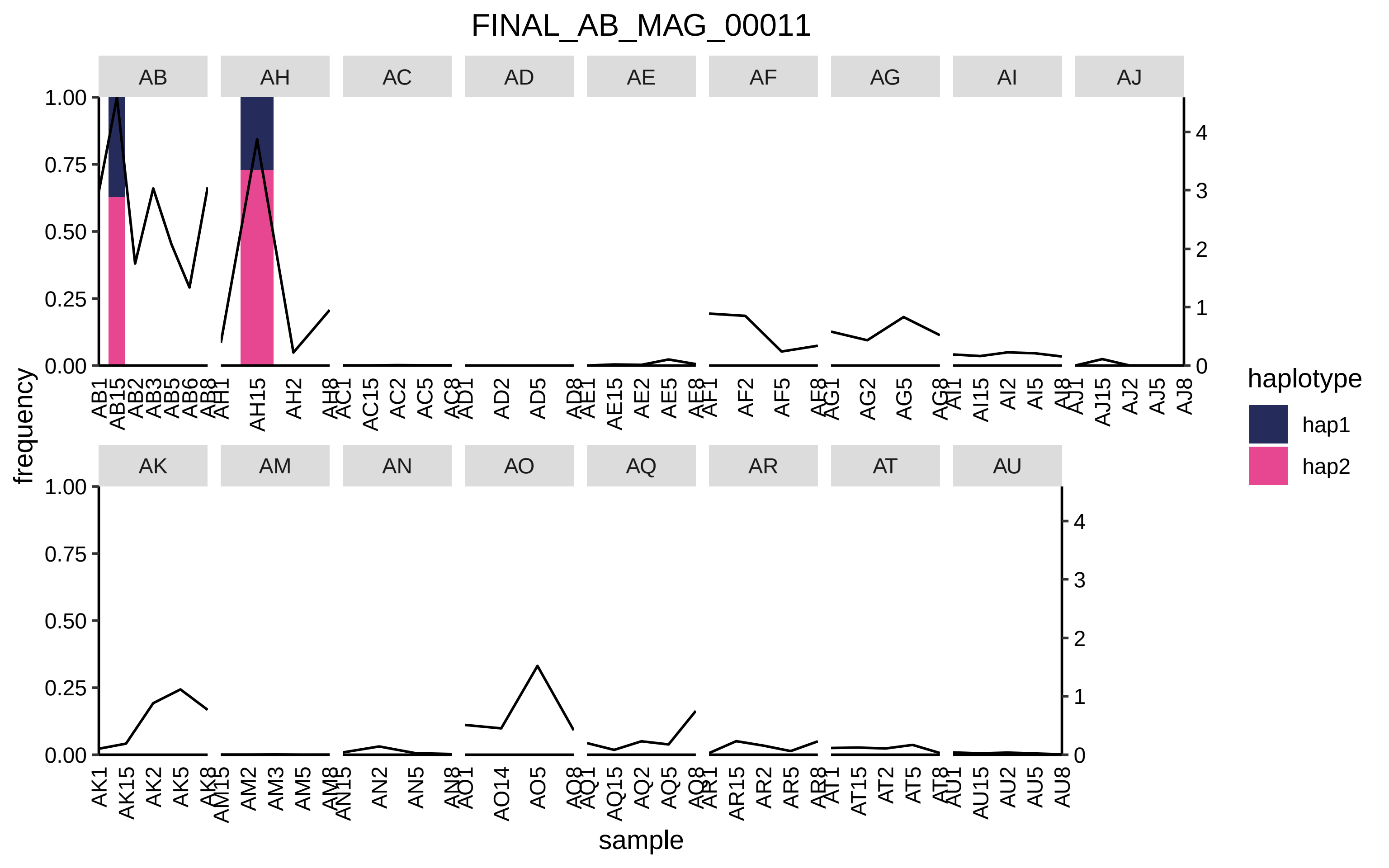

FINAL_AB_MAG_00011
AB
AH
AC
AD
AE
AF
AG
AI
AJ
AK
AM
AN
AO
AQ
AR
AT
AU
frequency
sample
haplotype
hap1
hap2
1.00
0.75
0.50
0.25
0.00
0
1
2
3
4

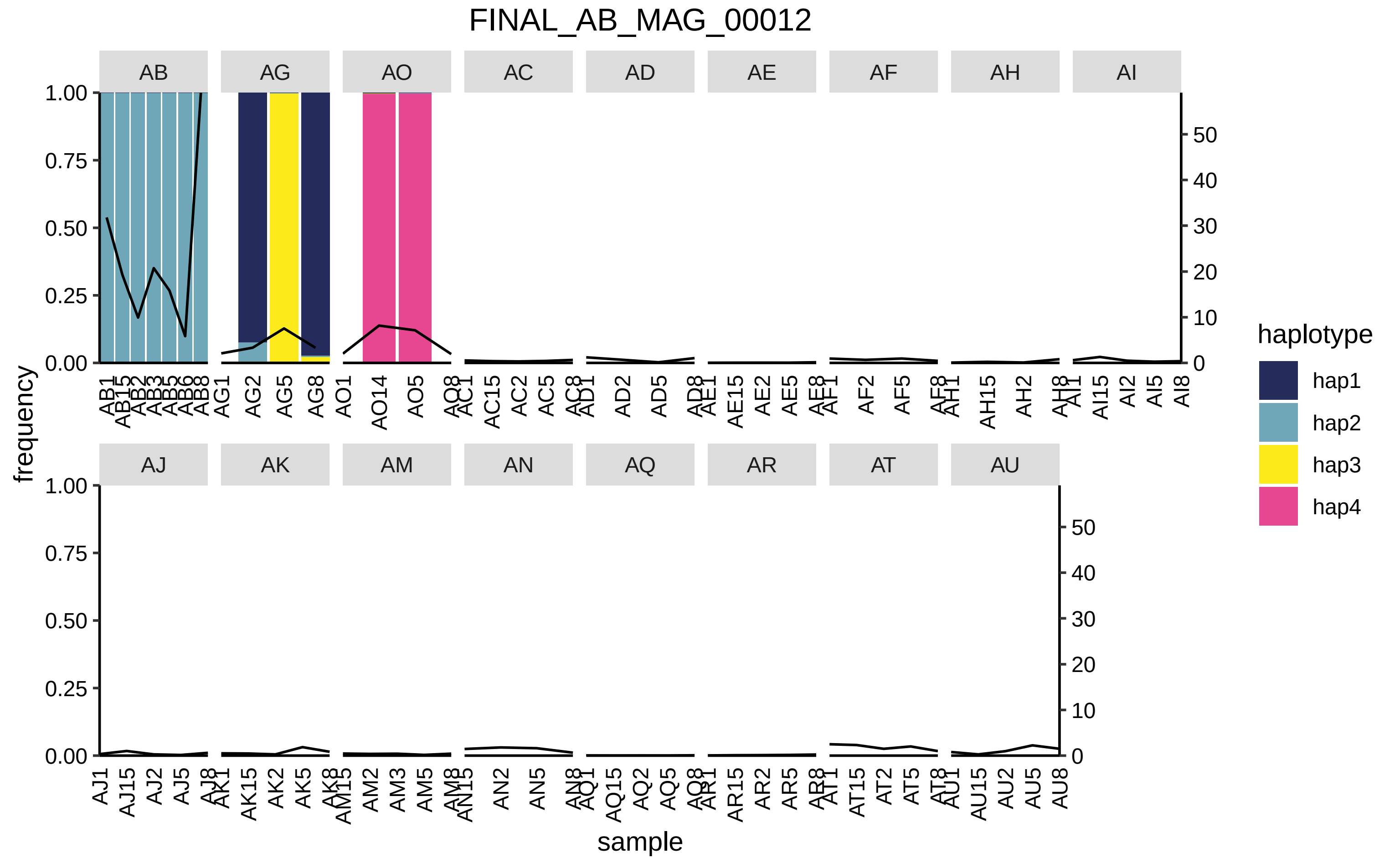
FINAL_AB_MAG_00012
AB
AG
AO
AC
AD
AE
AF
AH
AI
AJ
AK
AM
AN
AQ
AR
AT
AU
frequency
sample
haplotype
hap1
hap2
hap3
hap4

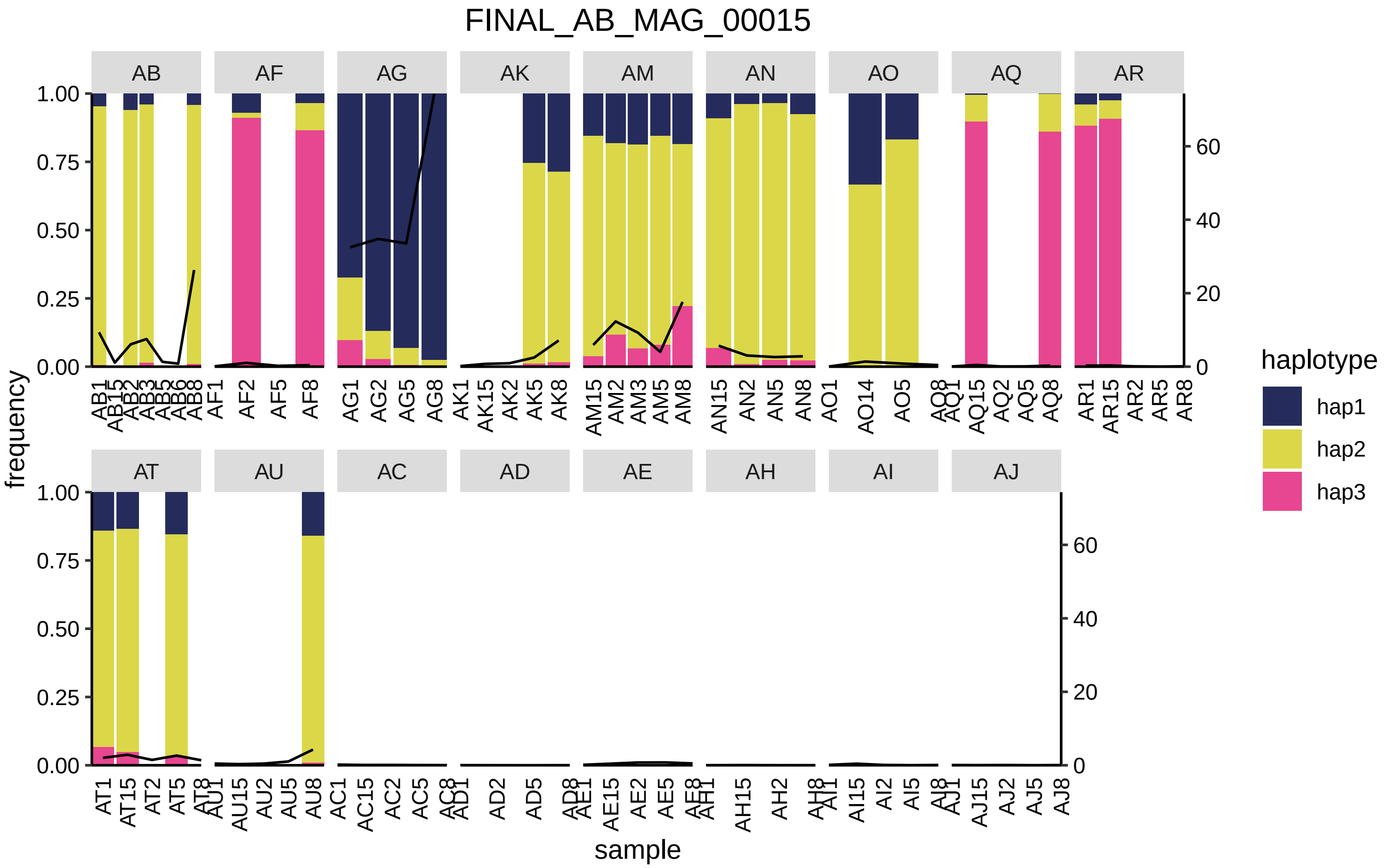

FINAL_AB_MAG_00015
AB
AF
AG
AK
AM
AN
AO
AQ
AR
AT
AU
AC
AD
AE
AH
AI
AJ
frequency
sample
haplotype
hap1
hap2
hap3

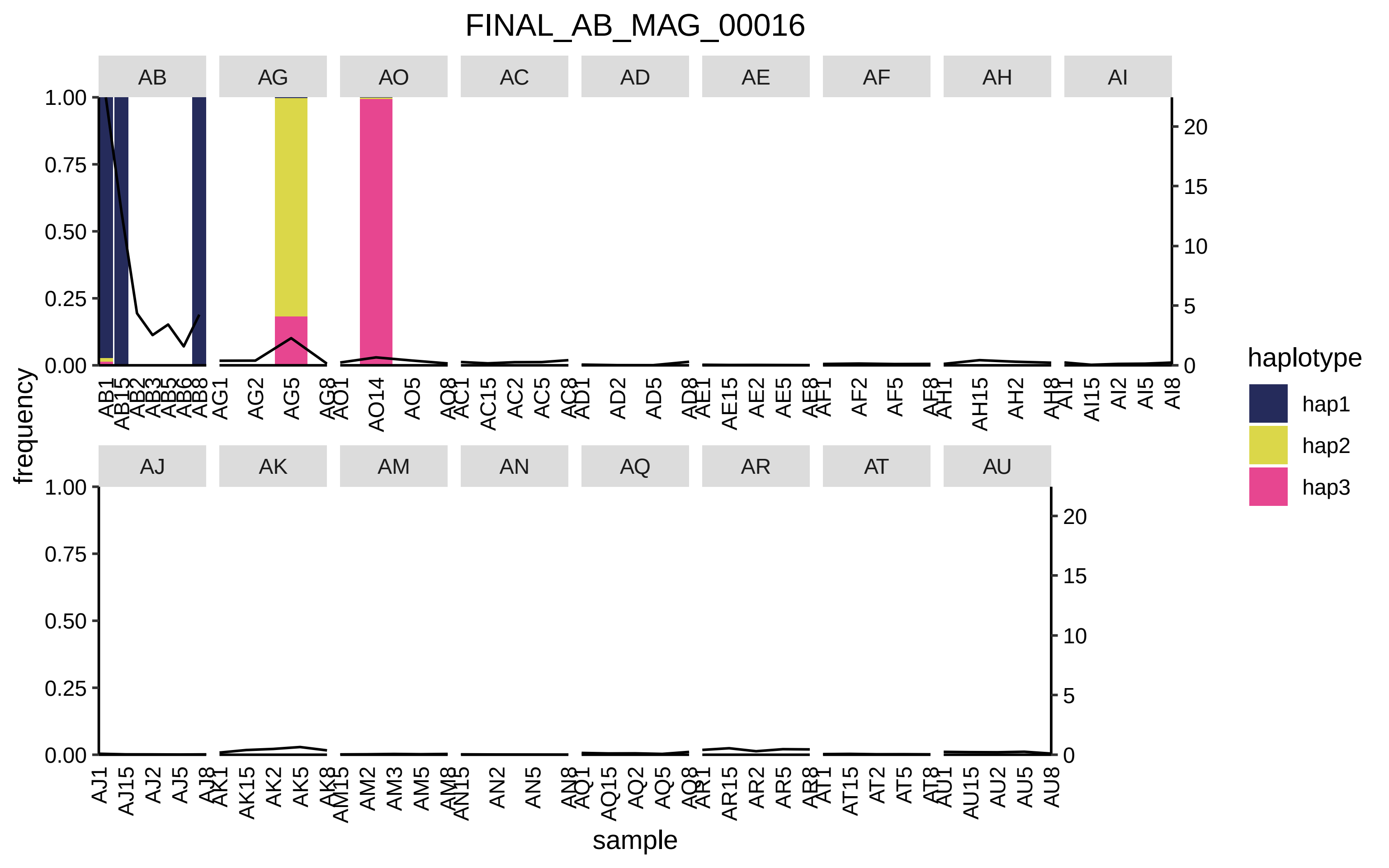
FINAL_AB_MAG_00016
AB
AG
AO
AC
AD
AE
AF
AH
AI
AJ
AK
AM
AN
AQ
AR
AT
AU
frequency
sample
haplotype
hap1
hap2
hap3

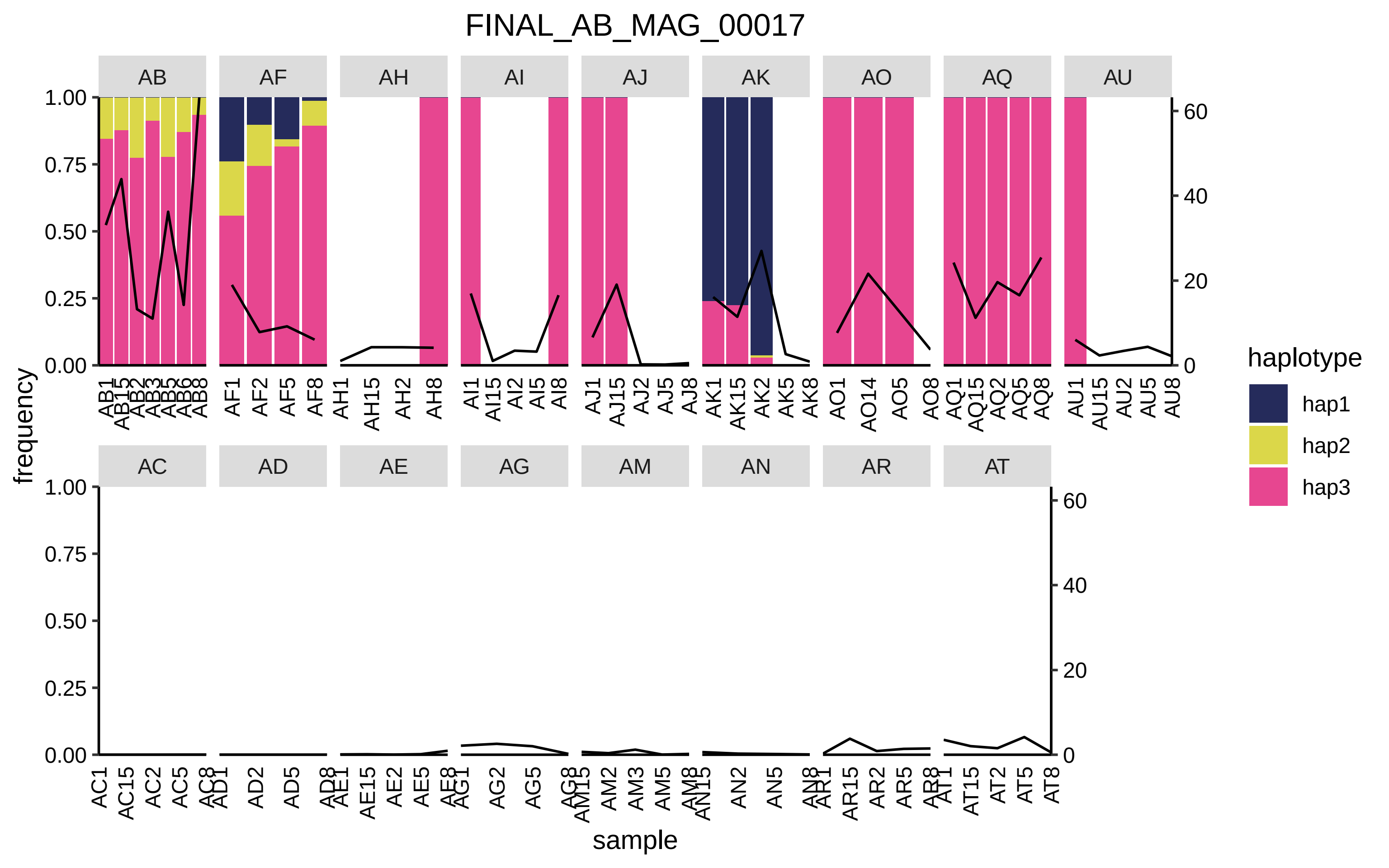
FINAL_AB_MAG_00017
AB
AF
AH
AI
AJ
AK
AO
AQ
AU
AC
AD
AE
AG
AM
AN
AR
AT
frequency
sample
haplotype
hap1
hap2
hap3
1.00
0.75
0.50
0.25
0.00
60
40
20
0

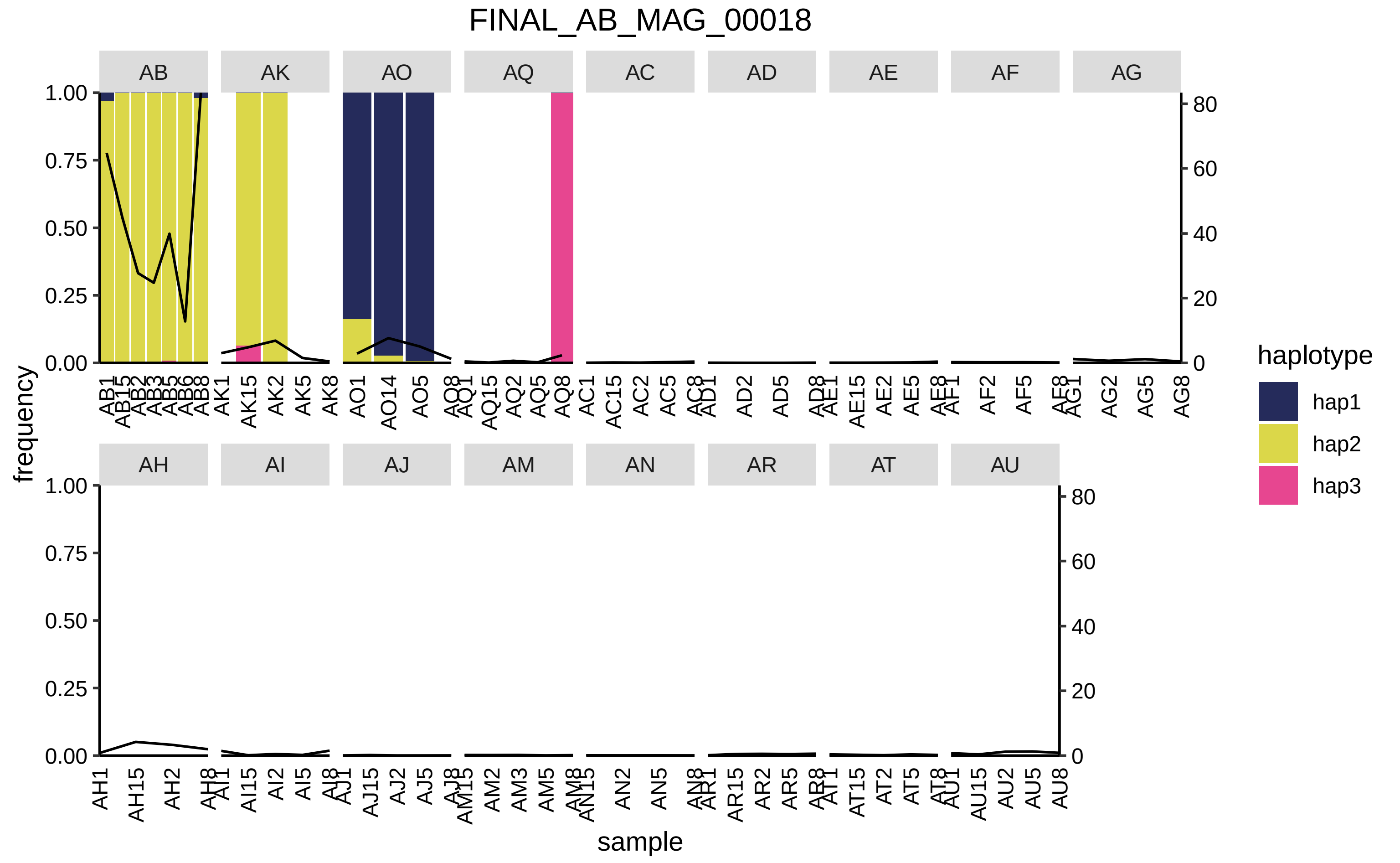
FINAL_AB_MAG_00018
AB
AK
AO
AQ
AC
AD
AE
AF
AG
AH
AI
AJ
AM
AN
AR
AT
AU
frequency
sample
haplotype
hap1
hap2
hap3

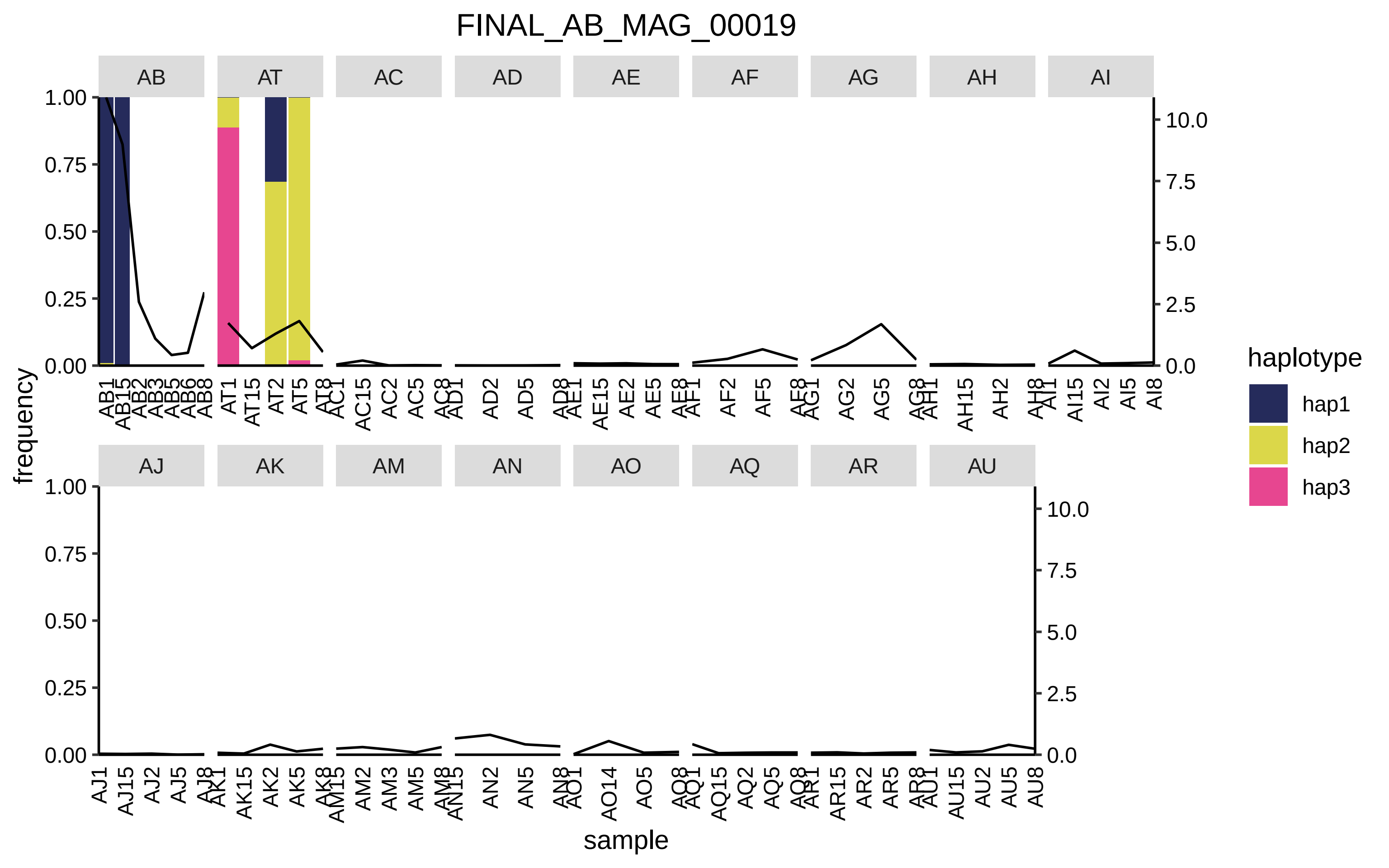

FINAL_AB_MAG_00019
AB
AT
AC
AD
AE
AF
AG
AH
AI
AJ
AK
AM
AN
AO
AQ
AR
AU
frequency
sample
haplotype
hap1
hap2
hap3

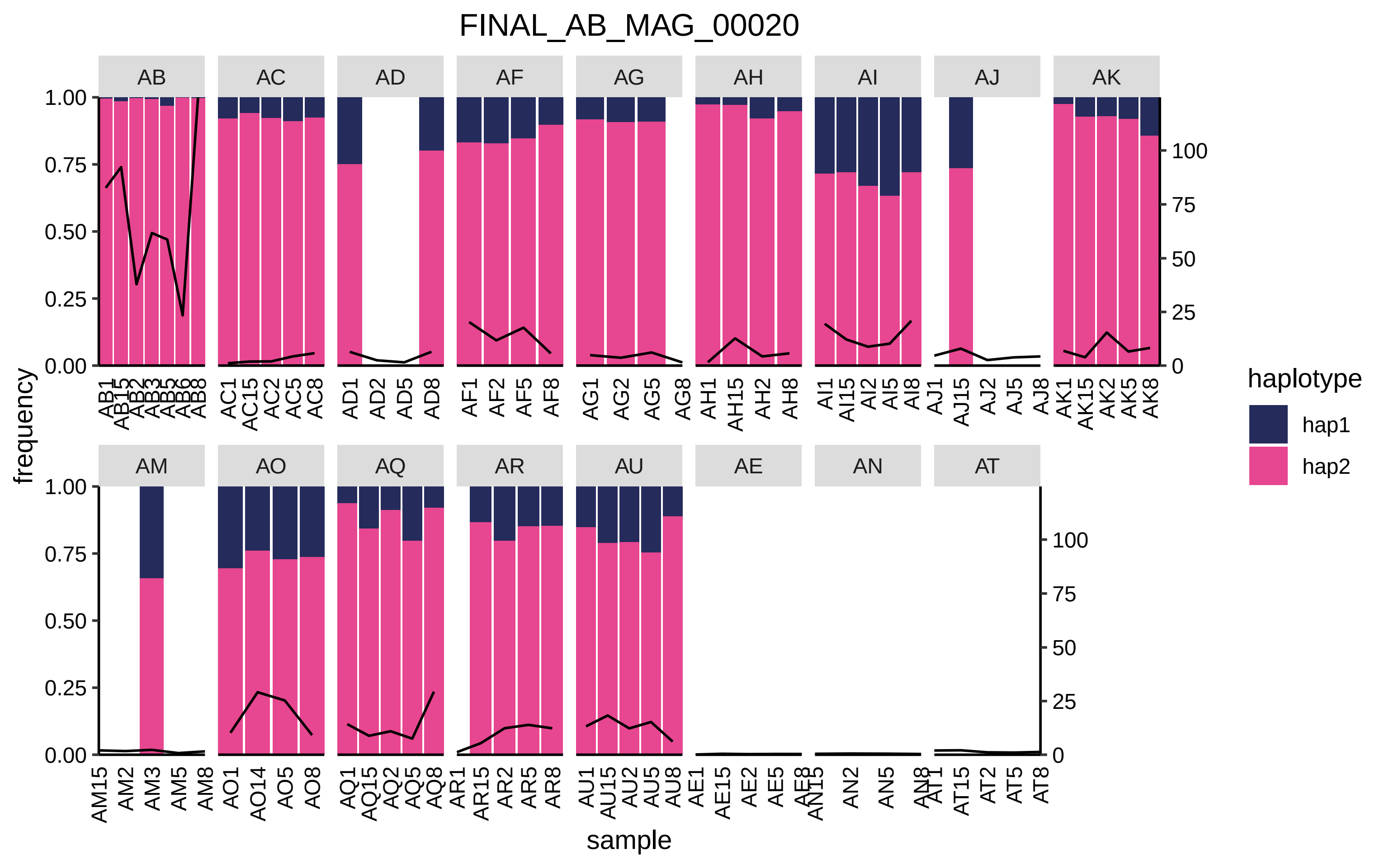

FINAL_AB_MAG_00020
AB
AC
AD
AF
AG
AH
AI
AJ
AK
AM
AO
AQ
AR
AU
AE
AN
AT
frequency
sample
haplotype
hap1
hap2

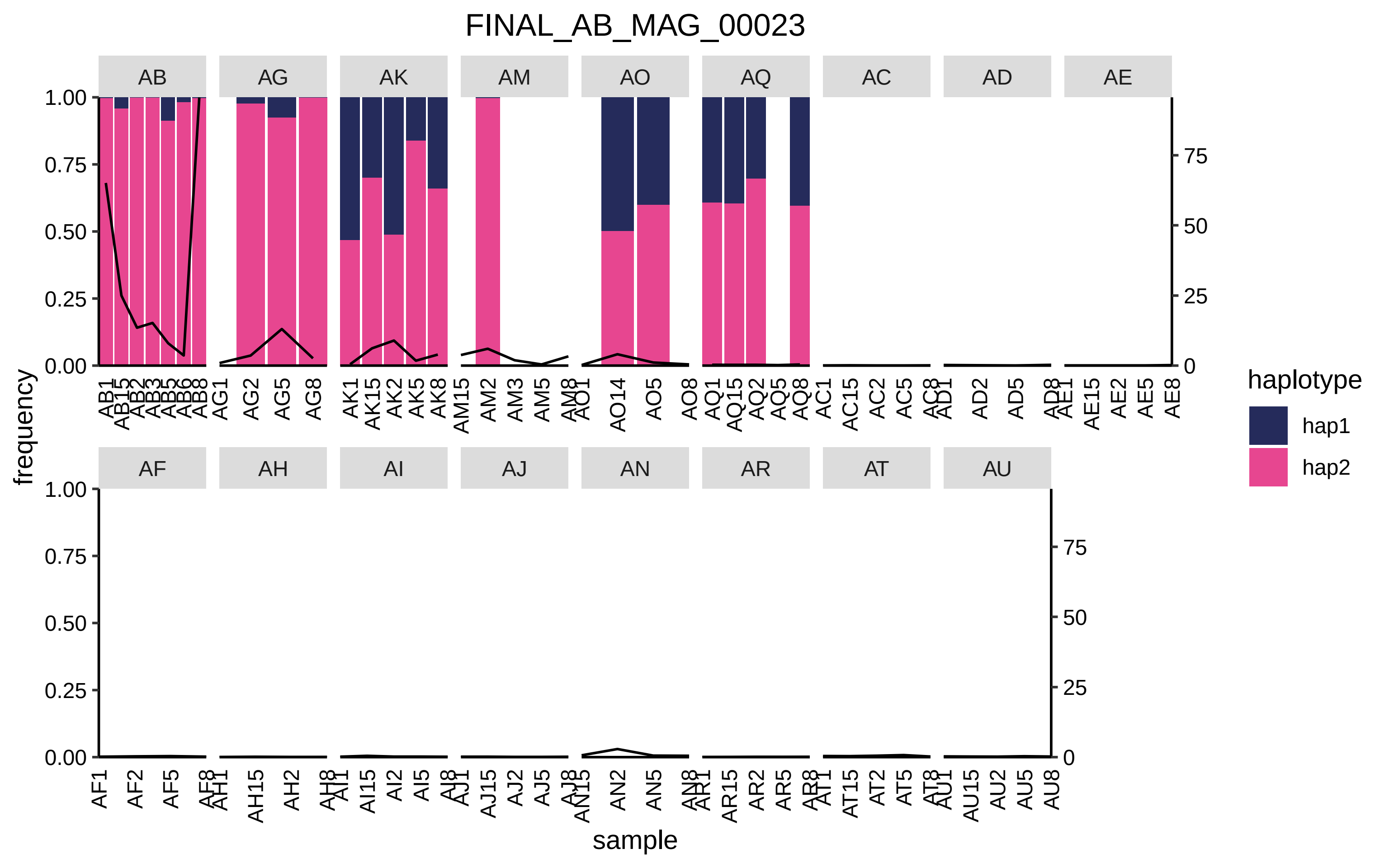
FINAL_AB_MAG_00023
AB
AG
AK
AM
AO
AQ
AC
AD
AE
AF
AH
AI
AJ
AN
AR
AT
AU
frequency
sample
haplotype
hap1
hap2
1.00
0.75
0.50
0.25
0.00
75
50
25
0

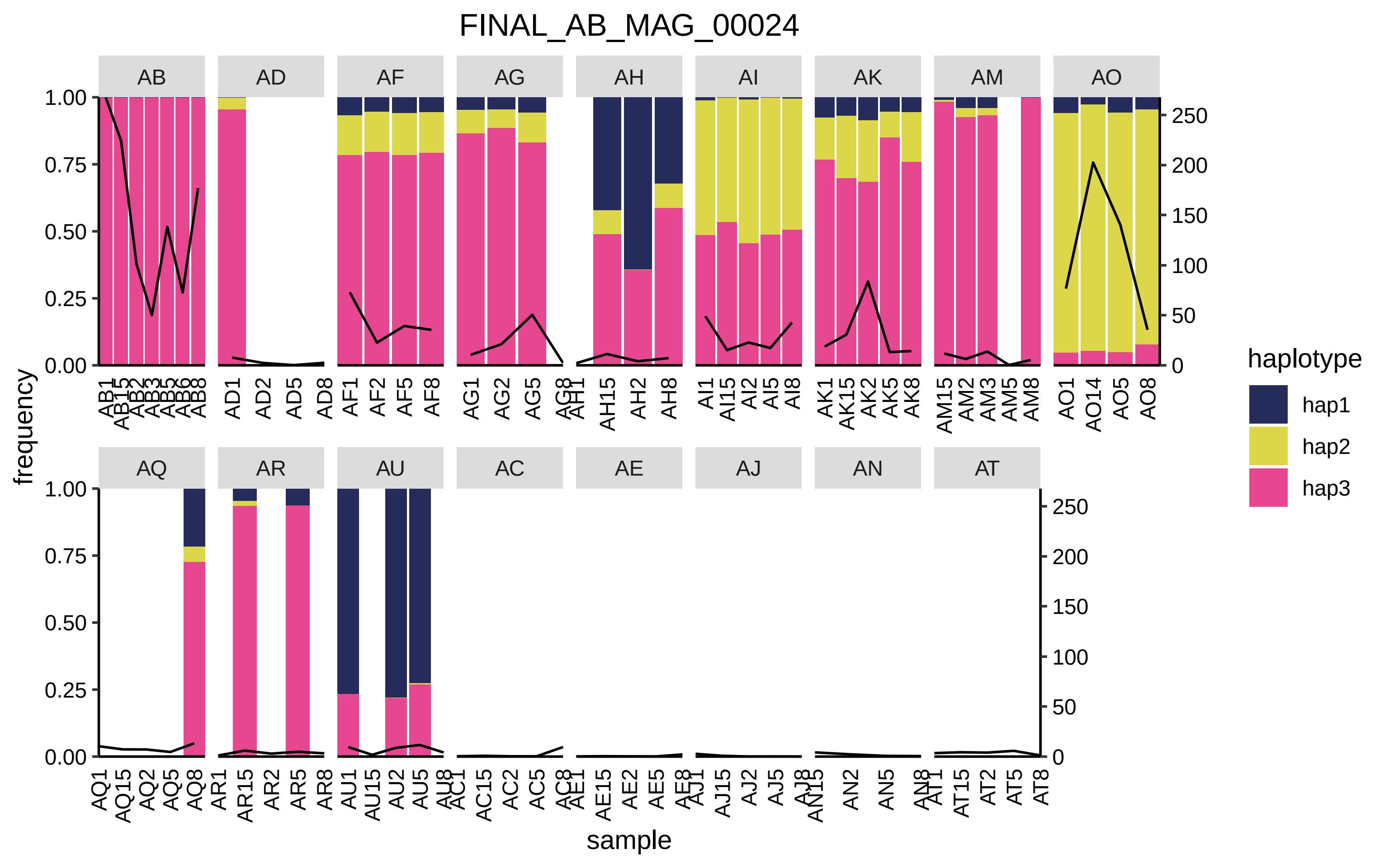

FINAL_AB_MAG_00024
AB
AD
AF
AG
AH
AI
AK
AM
AO
AQ
AR
AU
AC
AE
AJ
AN
AT
frequency
sample
haplotype
hap1
hap2
hap3
1.00
0.75
0.50
0.25
0.00
250
200
150
100
50
0

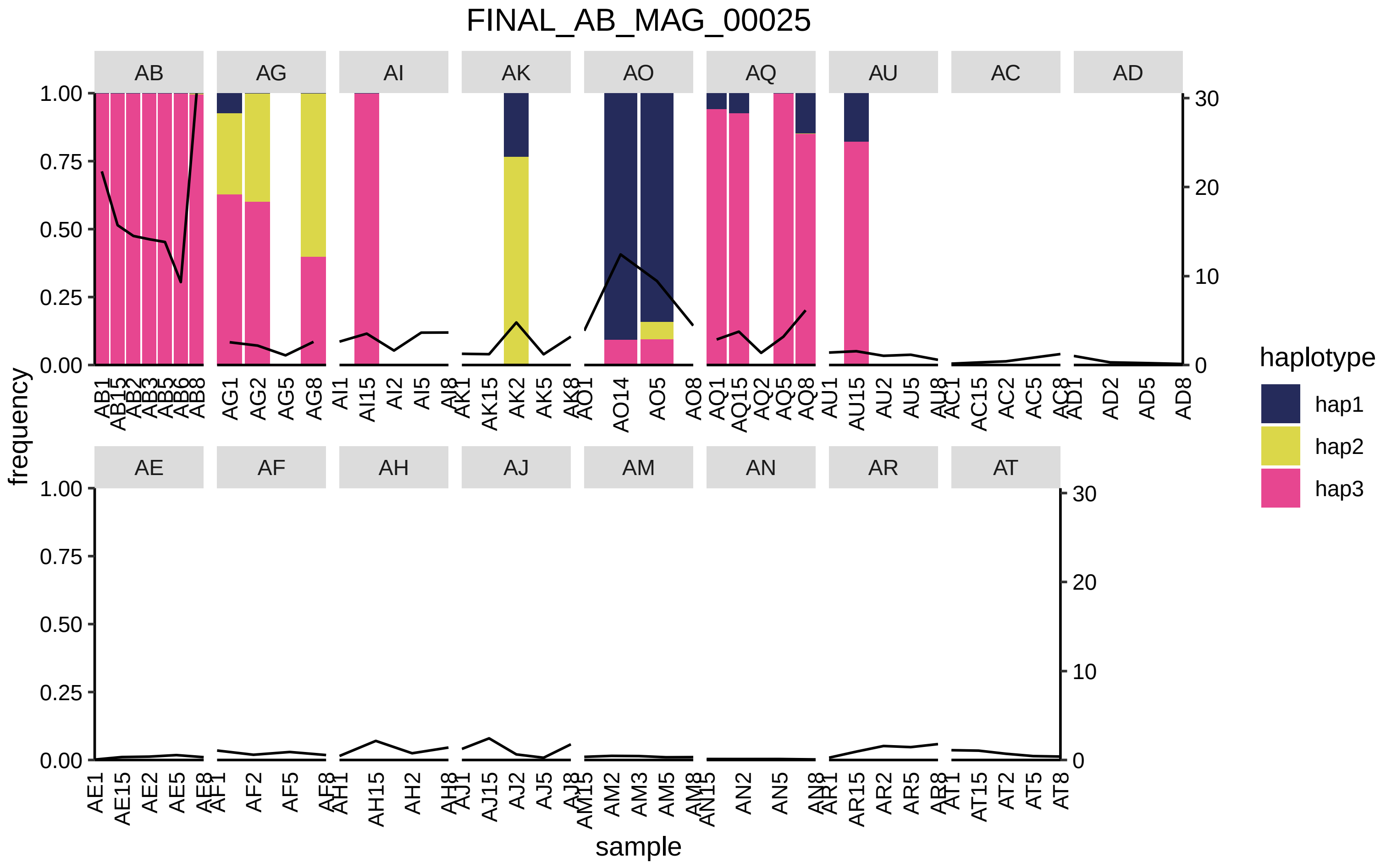

FINAL_AB_MAG_00025
AB
AG
AI
AK
AO
AQ
AU
AC
AD
AE
AF
AH
AJ
AM
AN
AR
AT
frequency
sample
haplotype
hap1
hap2
hap3
1.00
0.75
0.50
0.25
0.00
30
20
10
0

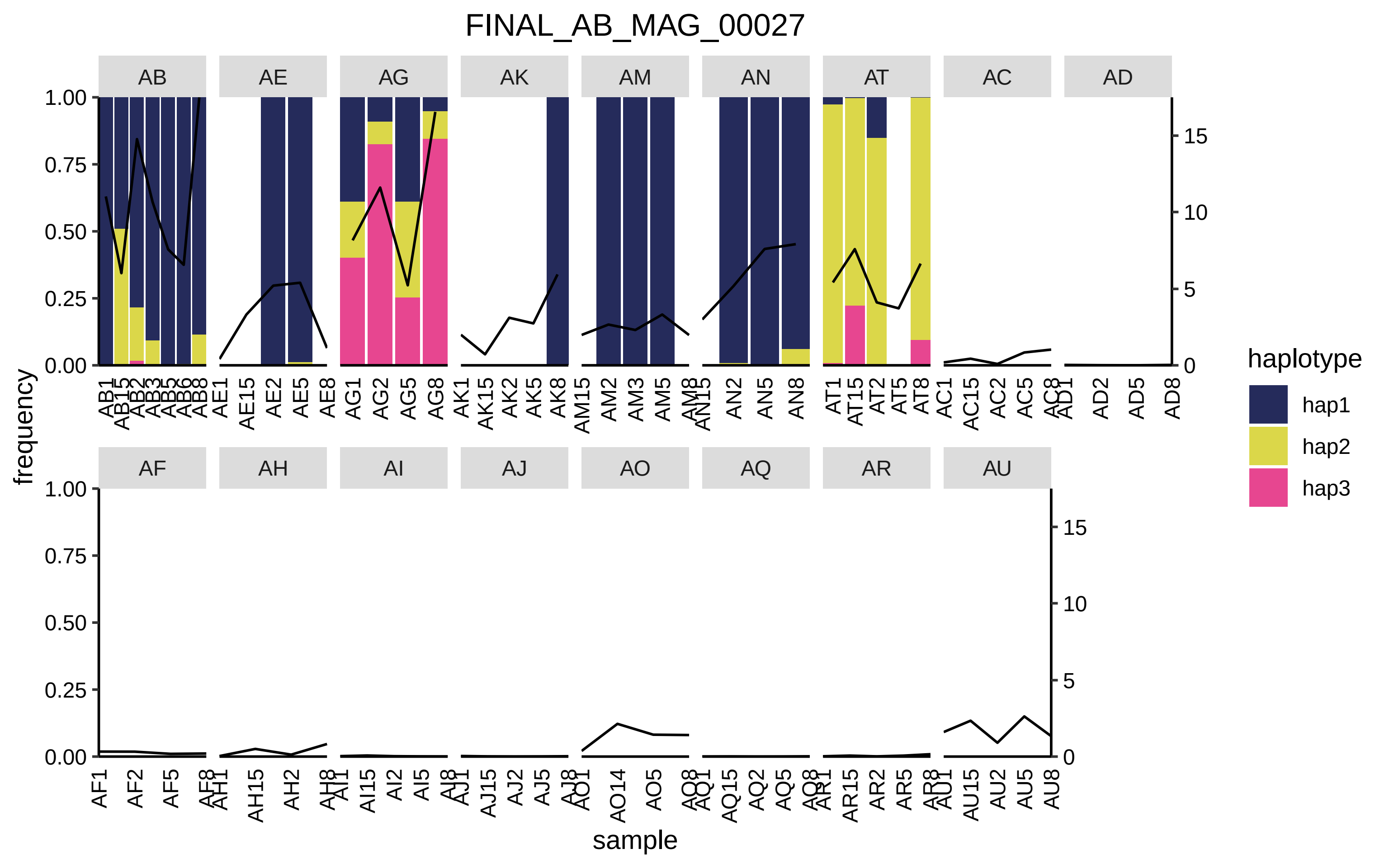

FINAL_AB_MAG_00027
AB
AE
AG
AK
AM
AN
AT
AC
AD
AF
AH
AI
AJ
AO
AQ
AR
AU
frequency
sample
haplotype
hap1
hap2
hap3
1.00
0.75
0.50
0.25
0.00
15
10
5
0
AB1 AB15 AB2 AB3 AB5 AB6 AB8
AE1 AE15 AE2 AE5 AE8
AG1 AG2 AG5 AG8
AK1 AK15 AK2 AK5 AK8
AM15 AM2 AM3 AM5 AM8
AN15 AN2 AN5 AN8
AT1 AT15 AT2 AT5 AT8
AC1 AC15 AC2 AC5 AC8
AD1 AD2 AD5 AD8
AF1 AF2 AF5 AF8
AH1 AH15 AH2 AH8
AI1 AI15 AI2 AI5 AI8
AJ1 AJ15 AJ2 AJ5 AJ8
AO1 AO14 AO5 AO8
AQ1 AQ15 AQ2 AQ5 AQ8
AR1 AR15 AR2 AR5 AR8
AU1 AU15 AU2 AU5 AU8

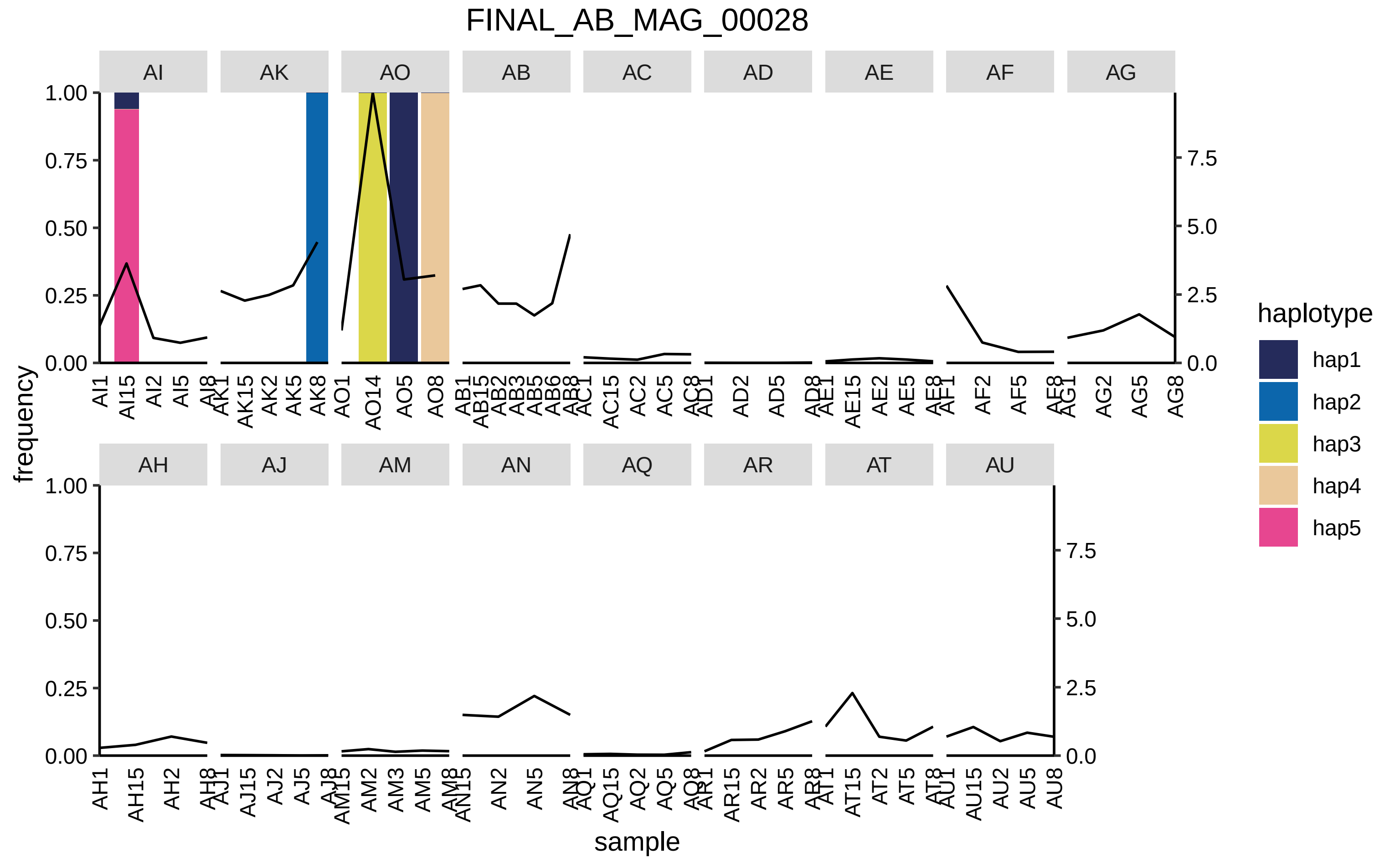
FINAL_AB_MAG_00028
AI
AK
AO
AB
AC
AD
AE
AF
AG
AH
AJ
AM
AN
AQ
AR
AT
AU
frequency
sample
haplotype
hap1
hap2
hap3
hap4
hap5

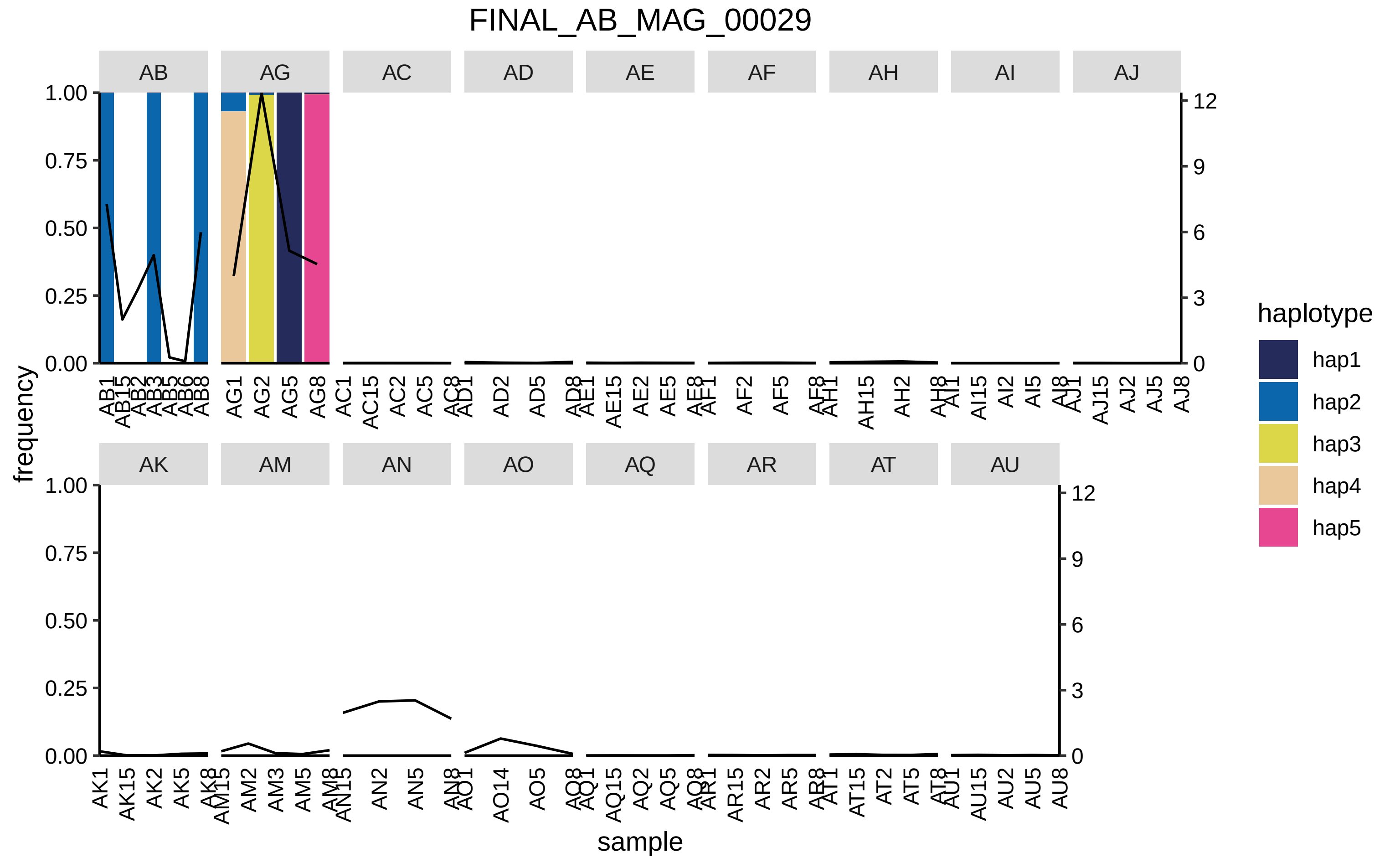
FINAL_AB_MAG_00029
AB
AG
AC
AD
AE
AF
AH
AI
AJ
AK
AM
AN
AO
AQ
AR
AT
AU
frequency
sample
haplotype
hap1
hap2
hap3
hap4
hap5

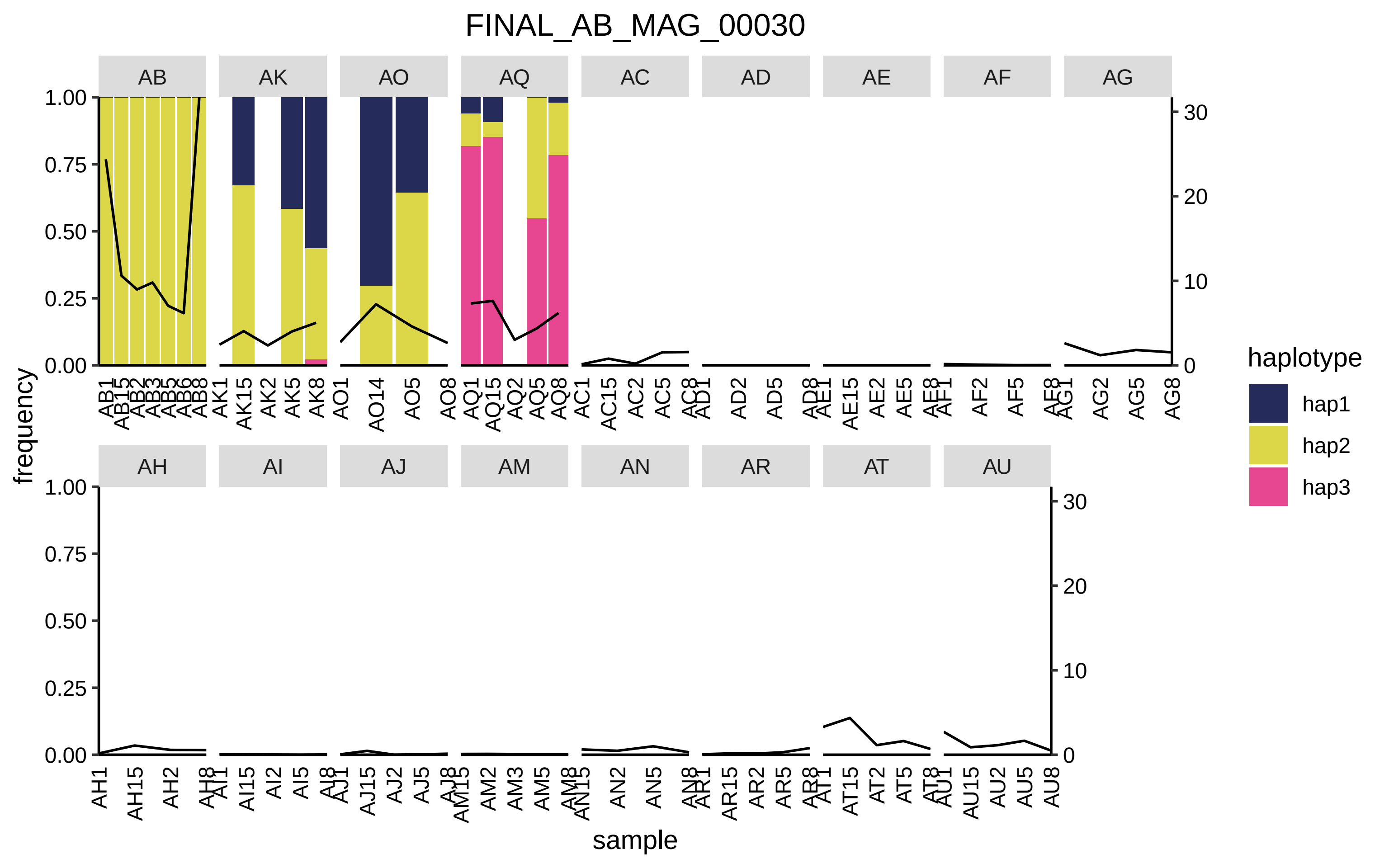

FINAL_AB_MAG_00030
AB
AK
AO
AQ
AC
AD
AE
AF
AG
AH
AI
AJ
AM
AN
AR
AT
AU
frequency
sample
haplotype
hap1
hap2
hap3

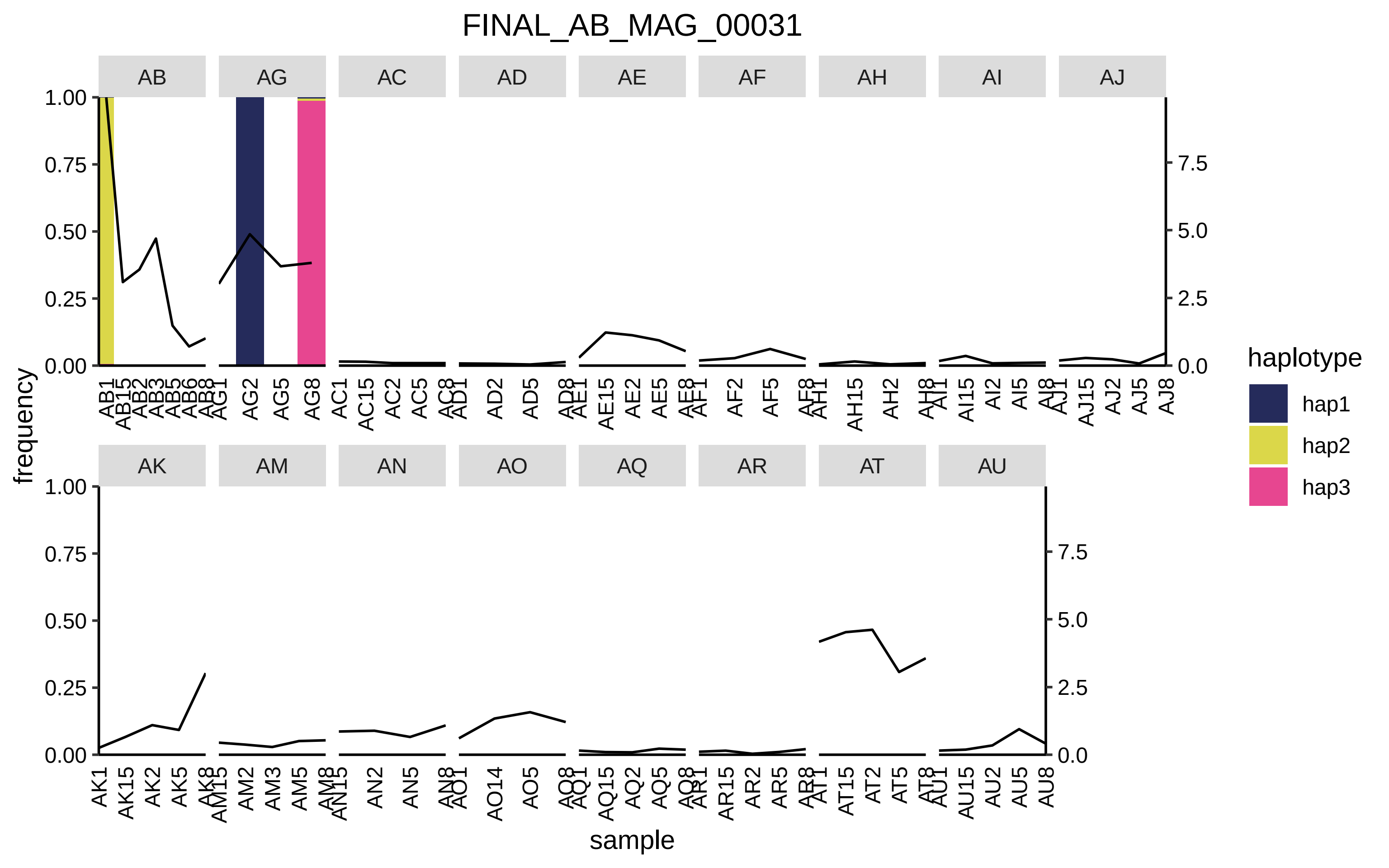
FINAL_AB_MAG_00031
AB
AG
AC
AD
AE
AF
AH
AI
AJ
AK
AM
AN
AO
AQ
AR
AT
AU
frequency
sample
haplotype
hap1
hap2
hap3
1.00
0.75
0.50
0.25
0.00
7.5
5.0
2.5
0.0

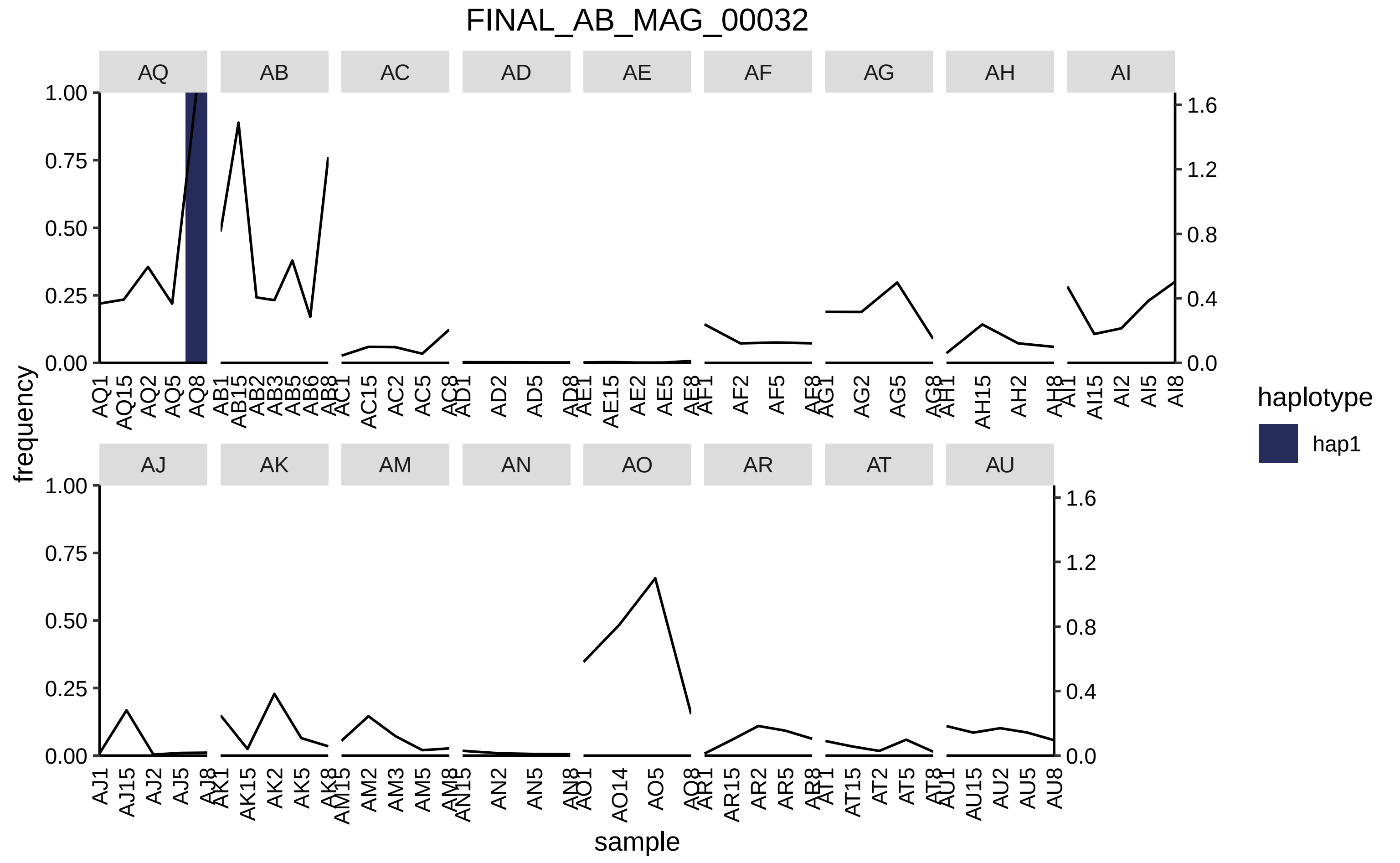
FINAL_AB_MAG_00032
AQ
AB
AC
AD
AE
AF
AG
AH
AI
AJ
AK
AM
AN
AO
AR
AT
AU
frequency
sample
haplotype
hap1
1.00
0.75
0.50
0.25
0.00
1.6
1.2
0.8
0.4
0.0

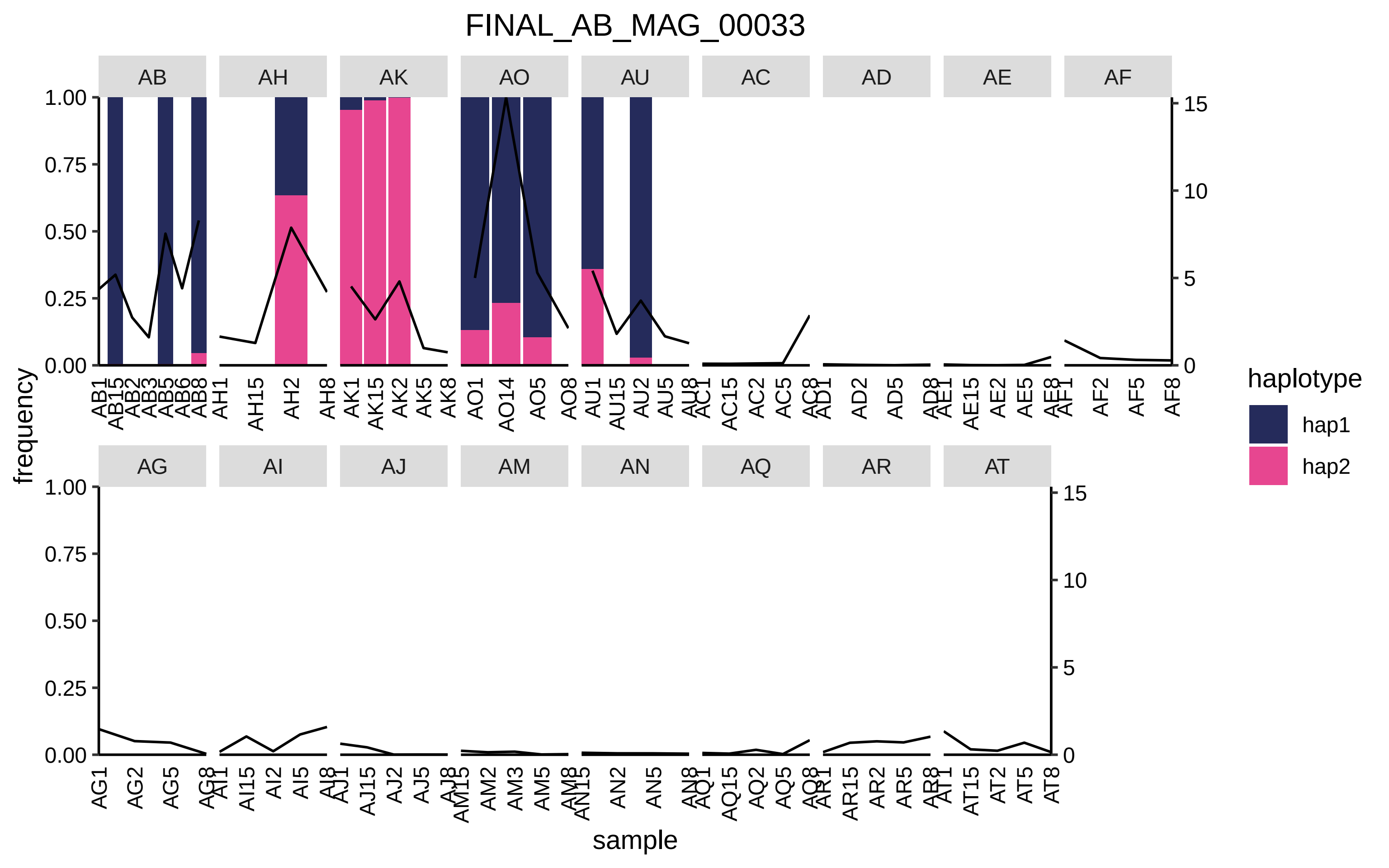
FINAL_AB_MAG_00033
AB
AH
AK
AO
AU
AC
AD
AE
AF
AG
AI
AJ
AM
AN
AQ
AR
AT
frequency
sample
haplotype
hap1
hap2

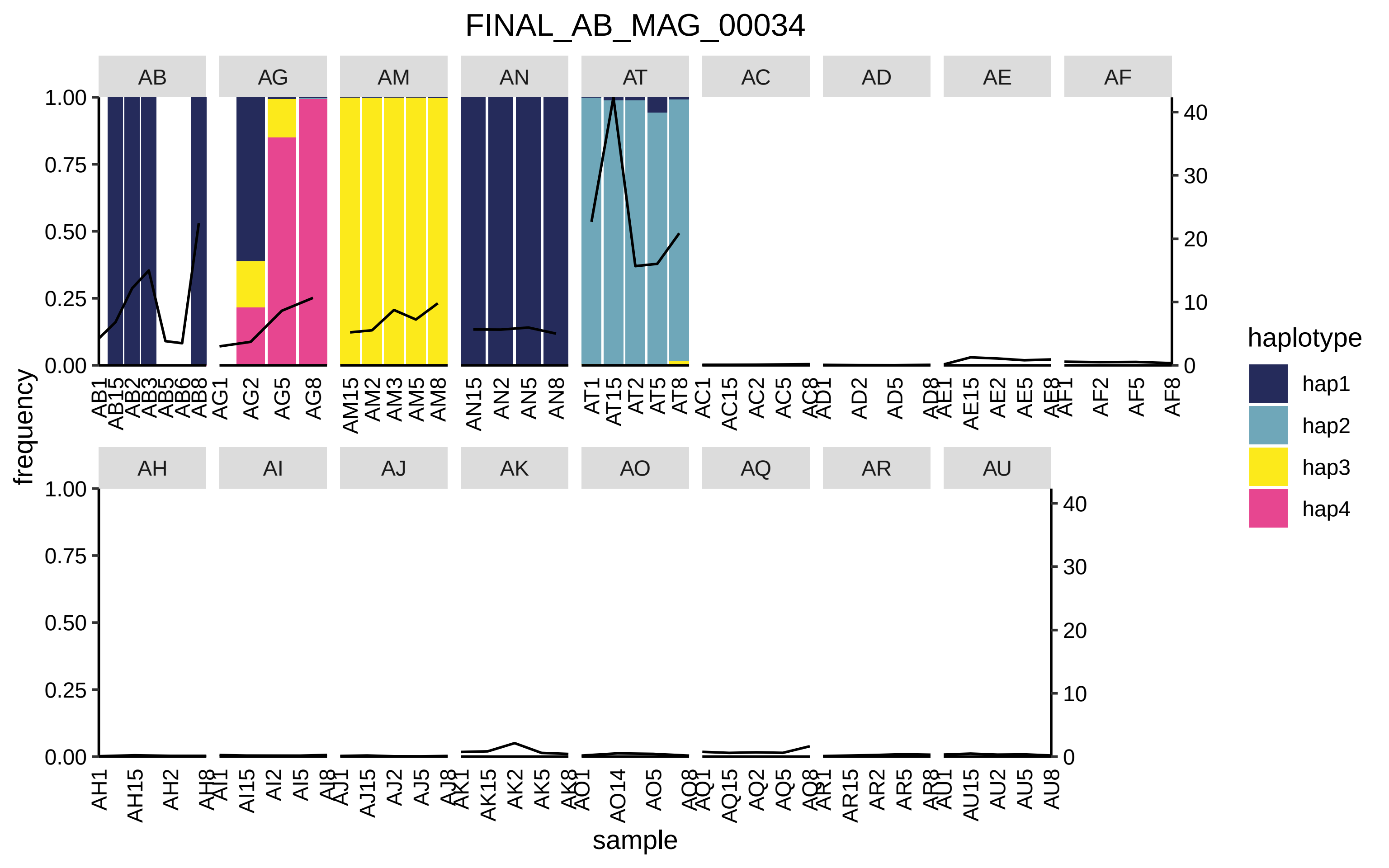
FINAL_AB_MAG_00034
AB
AG
AM
AN
AT
AC
AD
AE
AF
AH
AI
AJ
AK
AO
AQ
AR
AU
frequency
sample
haplotype
hap1
hap2
hap3
hap4

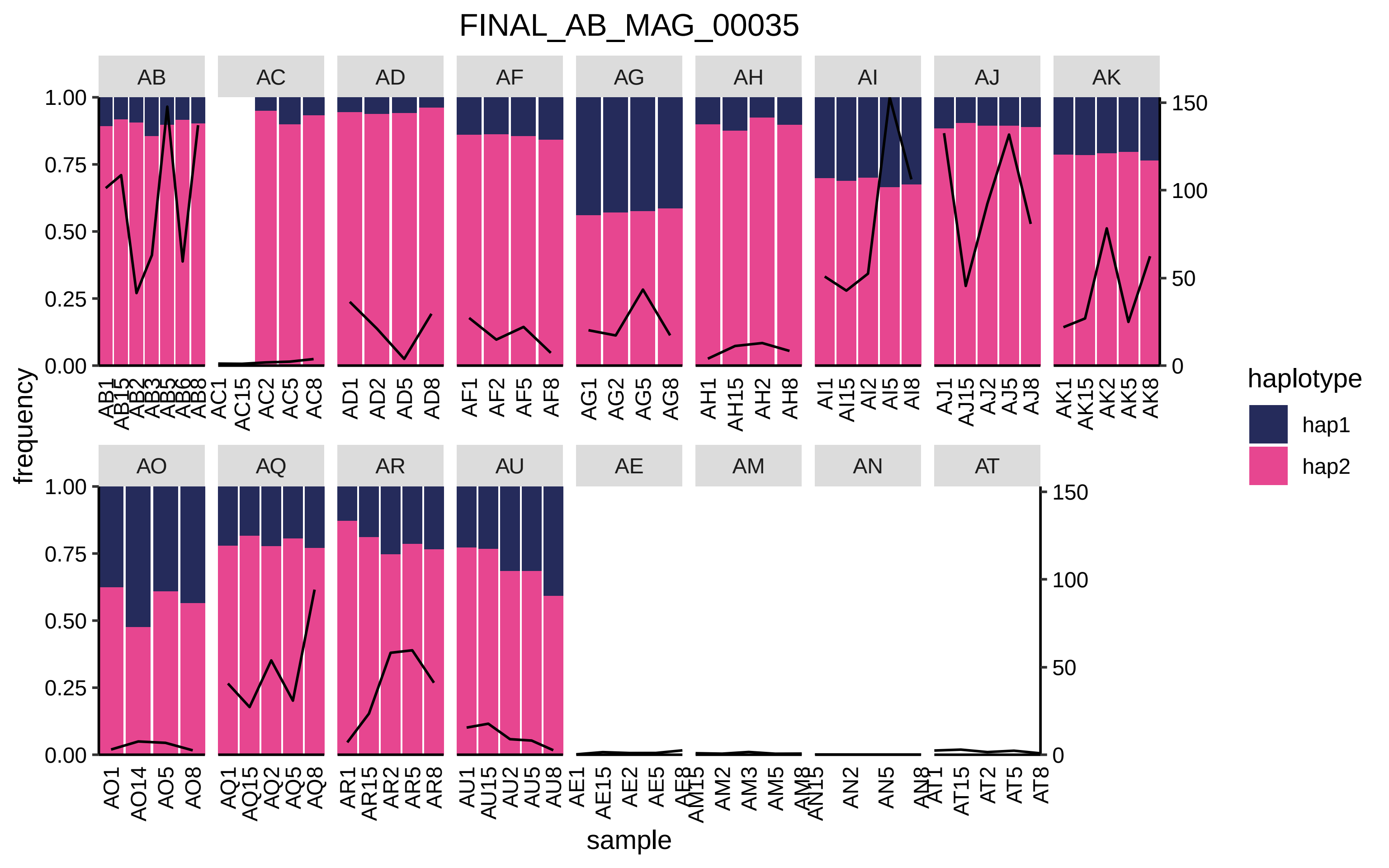
FINAL_AB_MAG_00035
AB
AC
AD
AF
AG
AH
AI
AJ
AK
AO
AQ
AR
AU
AE
AM
AN
AT
frequency
sample
haplotype
hap1
hap2
1.00
0.75
0.50
0.25
0.00
150
100
50
0

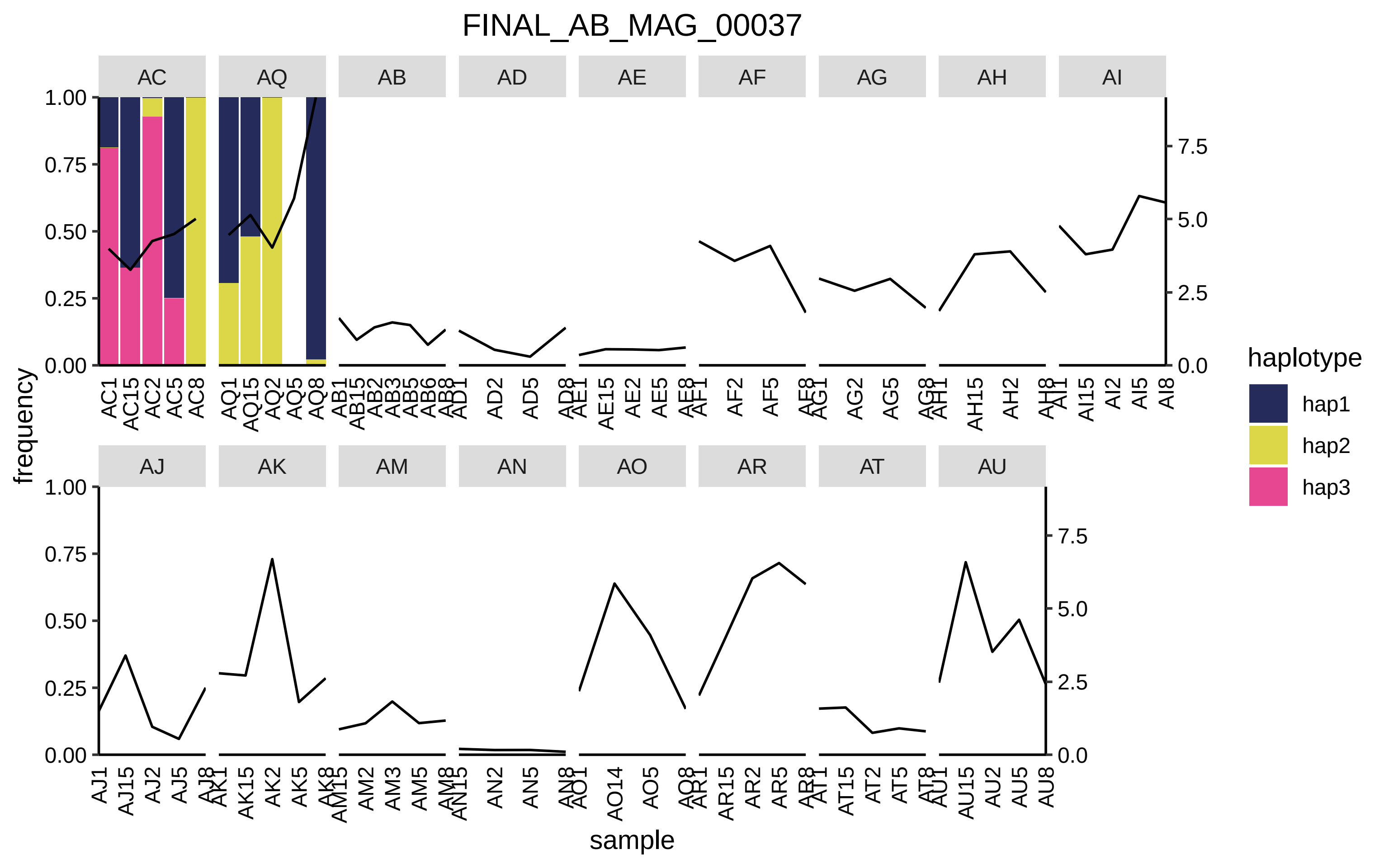

FINAL_AB_MAG_00037
AC
AQ
AB
AD
AE
AF
AG
AH
AI
AJ
AK
AM
AN
AO
AR
AT
AU
frequency
sample
1.00
0.75
0.50
0.25
0.00
7.5
5.0
2.5
0.0
AC1 AC15 AC2 AC5 AC8
AQ1 AQ15 AQ2 AQ5 AQ8
AB1 AB15 AB2 AB3 AB5 AB6 AB8
AD1 AD2 AD5 AD8
AE1 AE15 AE2 AE5 AE8
AF1 AF2 AF5 AF8
AG1 AG2 AG5 AG8
AH1 AH15 AH2 AH8
AI1 AI15 AI2 AI5 AI8
AJ1 AJ15 AJ2 AJ5 AJ8
AK1 AK15 AK2 AK5 AK8
AM15 AM2 AM3 AM5 AM8
AN15 AN2 AN5 AN8
AO1 AO14 AO5 AO8
AR1 AR15 AR2 AR5 AR8
AT1 AT15 AT2 AT5 AT8
AU1 AU15 AU2 AU5 AU8
haplotype
hap1
hap2
hap3

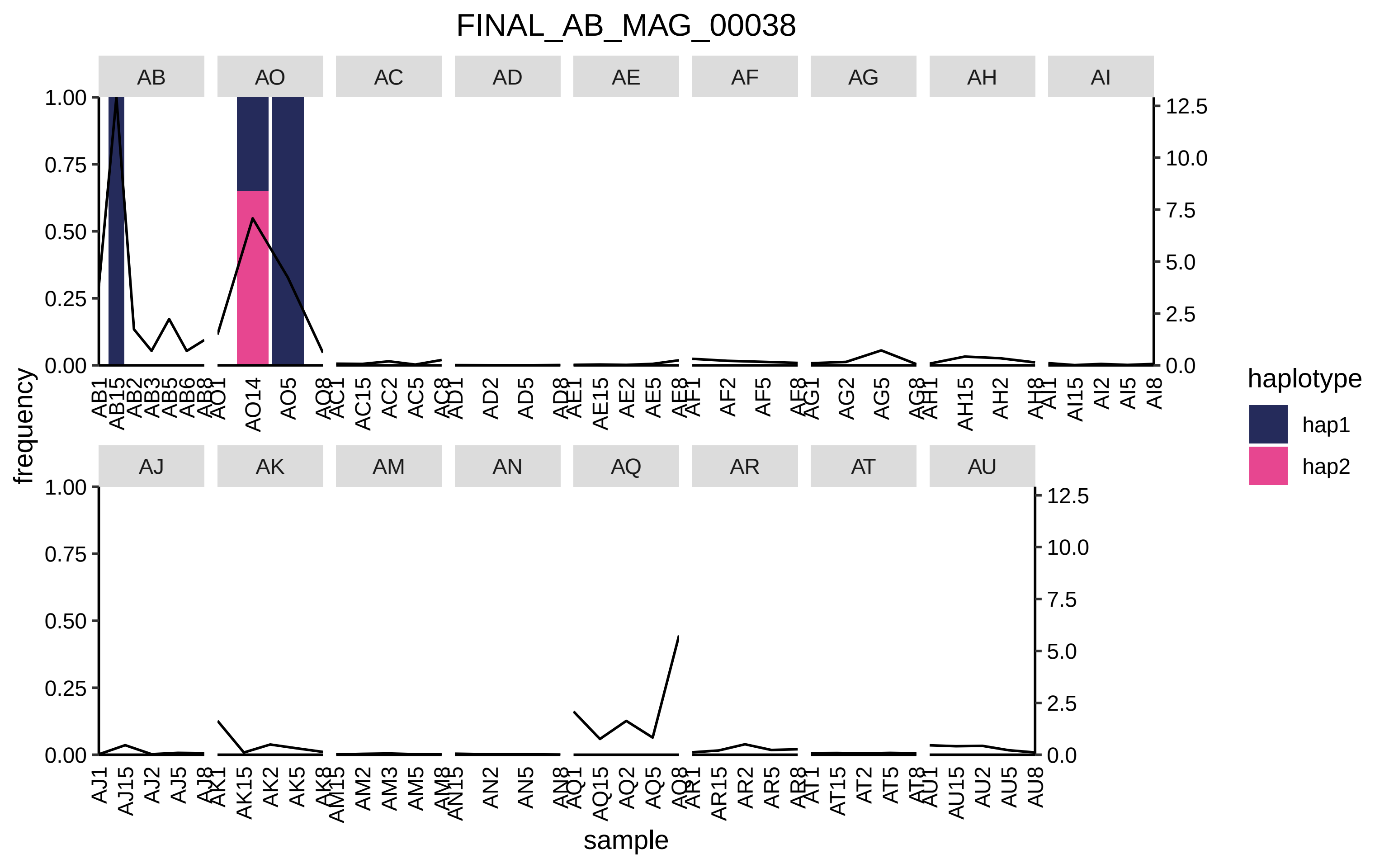
FINAL_AB_MAG_00038
AB
AO
AC
AD
AE
AF
AG
AH
AI
AJ
AK
AM
AN
AQ
AR
AT
AU
frequency
sample
haplotype
hap1
hap2
1.00
0.75
0.50
0.25
0.00
12.5
10.0
7.5
5.0
2.5
0.0

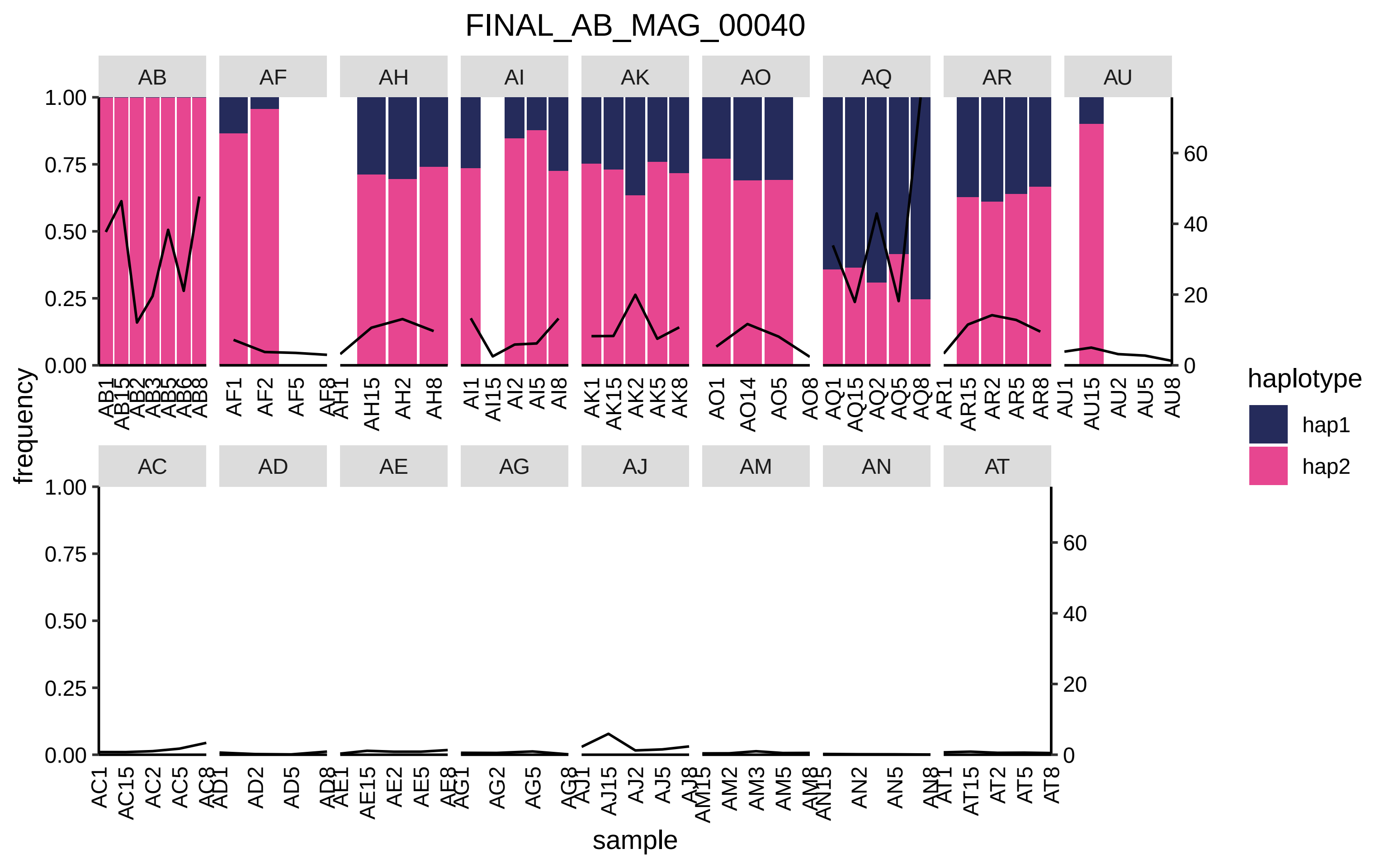

FINAL_AB_MAG_00040
AB
AF
AH
AI
AK
AO
AQ
AR
AU
AC
AD
AE
AG
AJ
AM
AN
AT
frequency
sample
haplotype
hap1
hap2

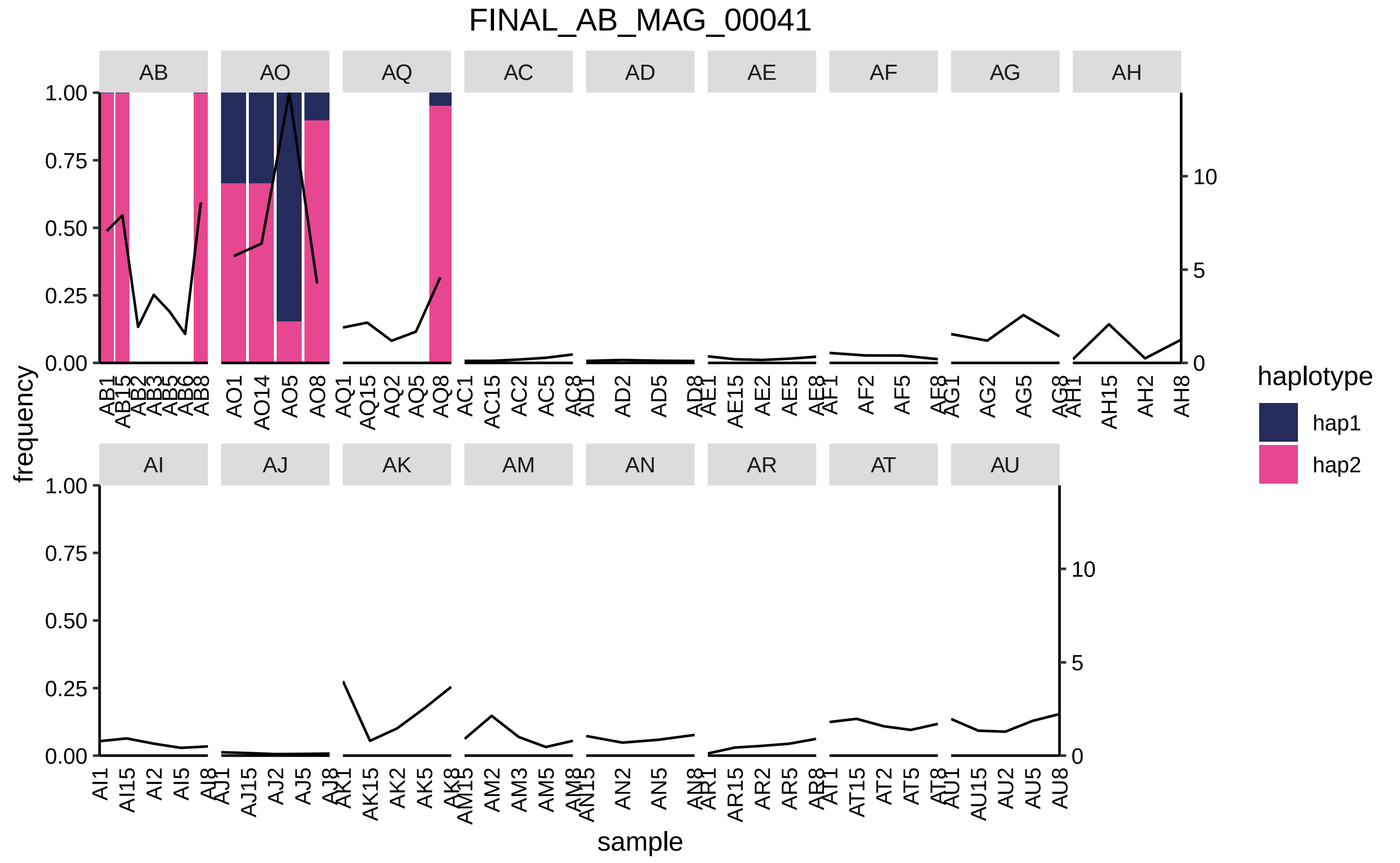

FINAL_AB_MAG_00041
AB
AO
AQ
AC
AD
AE
AF
AG
AH
AI
AJ
AK
AM
AN
AR
AT
AU
frequency
sample
haplotype
hap1
hap2
1.00
0.75
0.50
0.25
0.00
10
5
0
AO1
AO14
AO5
AO8
AQ1
AQ15
AQ2
AQ5
AQ8
AC1
AC15
AC2
AC5
AD2
AD5
AE15
AE2
AE5
AF2
AF5
AG2
AG5
AH15
AH2
AH8
AI15
AI2
AI5
AJ15
AJ2
AJ5
AK15
AK2
AK5
AM2
AM3
AM5
AN2
AN5
AR15
AR2
AR5
AT15
AT2
AT5
AU15
AU2
AU5
AU8

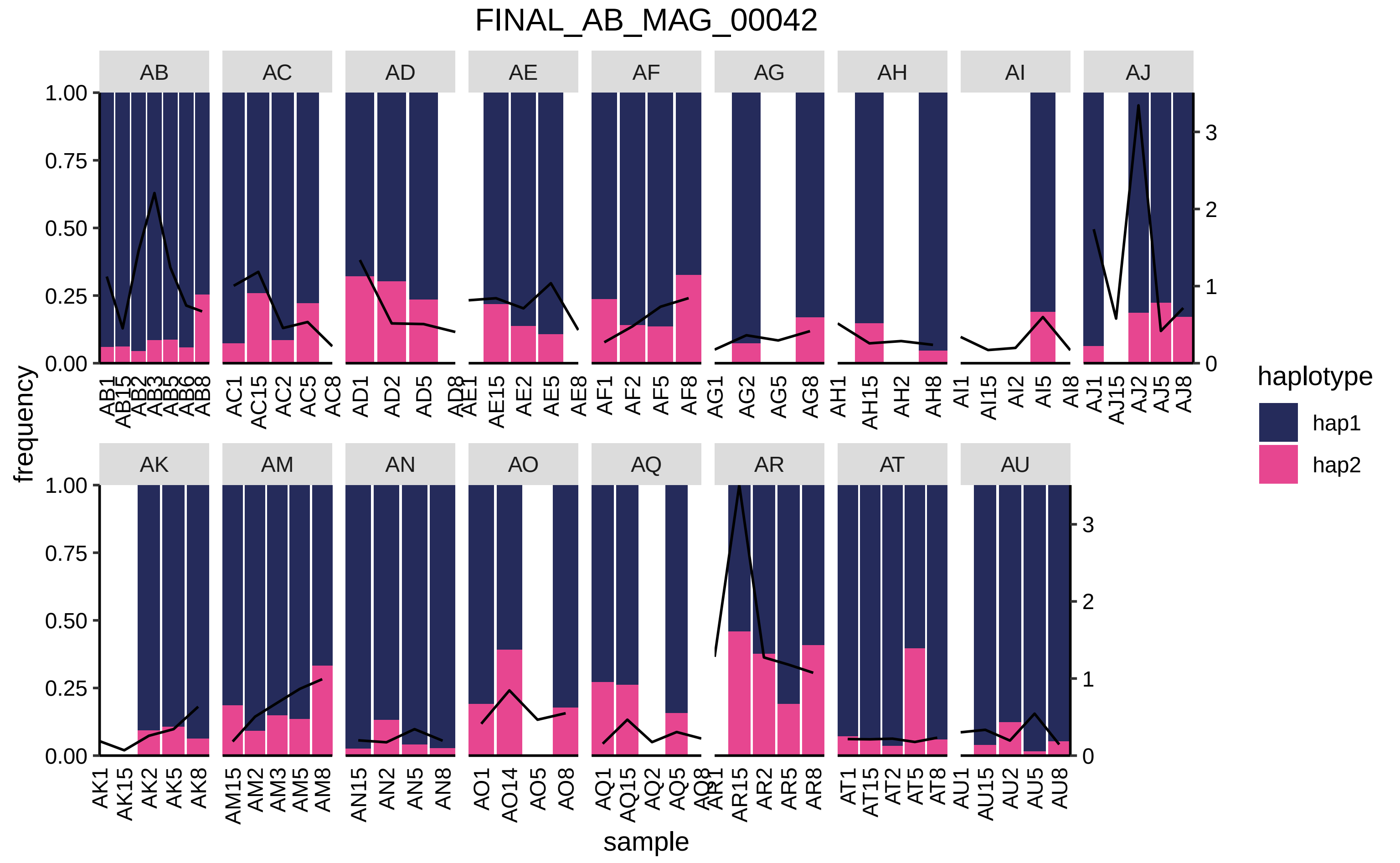

FINAL_AB_MAG_00042
AB
AC
AD
AE
AF
AG
AH
AI
AJ
AK
AM
AN
AO
AQ
AR
AT
AU
frequency
sample
haplotype
hap1
hap2
1.00
0.75
0.50
0.25
0.00
0
1
2
3
AB1 AB15 AB2 AB3 AB5 AB6 AB8
AC1 AC15 AC2 AC5 AC8
AD1 AD2 AD5 AD8
AE1 AE15 AE2 AE5 AE8
AF1 AF2 AF5 AF8
AG1 AG2 AG5 AG8
AH1 AH15 AH2 AH8
AI1 AI15 AI2 AI5 AI8
AJ1 AJ15 AJ2 AJ5 AJ8
AK1 AK15 AK2 AK5 AK8
AM15 AM2 AM3 AM5 AM8
AN15 AN2 AN5 AN8
AO1 AO14 AO5 AO8
AQ1 AQ15 AQ2 AQ5 AQ8
AR1 AR15 AR2 AR5 AR8
AT1 AT15 AT2 AT5 AT8
AU1 AU15 AU2 AU5 AU8

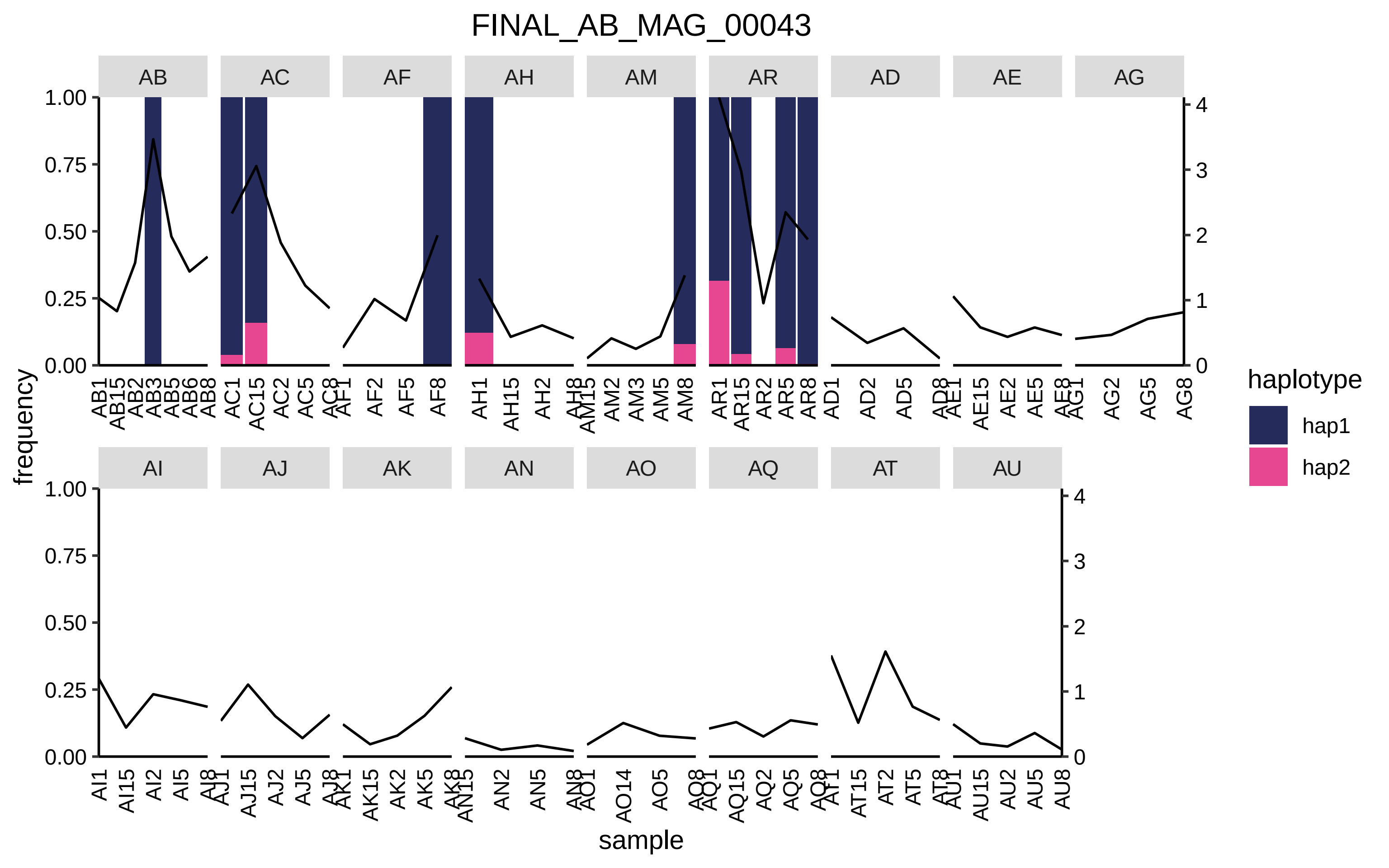
FINAL_AB_MAG_00043
AB
AC
AF
AH
AM
AR
AD
AE
AG
AI
AJ
AK
AN
AO
AQ
AT
AU
frequency
sample
haplotype
hap1
hap2

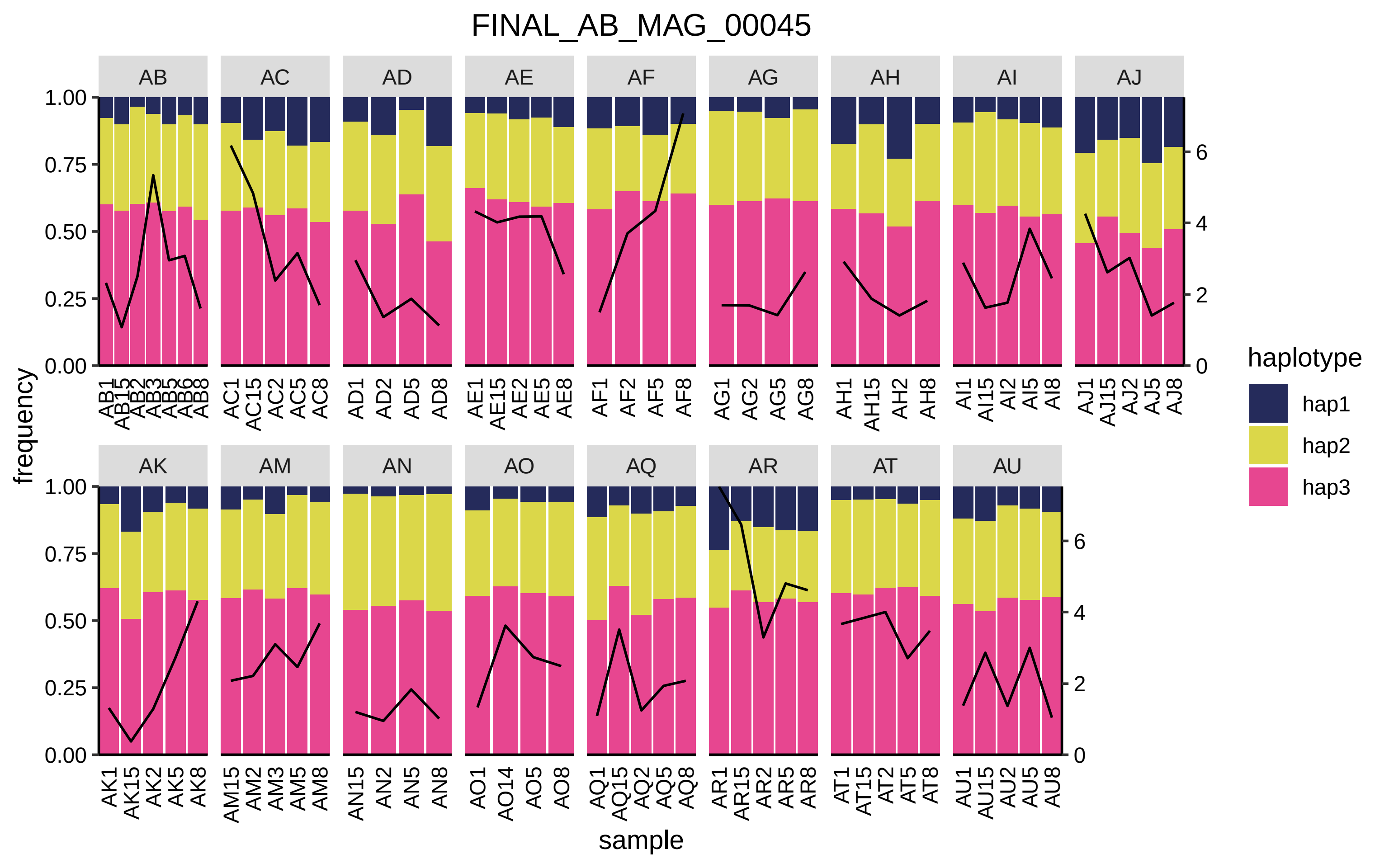

FINAL_AB_MAG_00045
AB
AC
AD
AE
AF
AG
AH
AI
AJ
AK
AM
AN
AO
AQ
AR
AT
AU
frequency
sample
haplotype
hap1
hap2
hap3

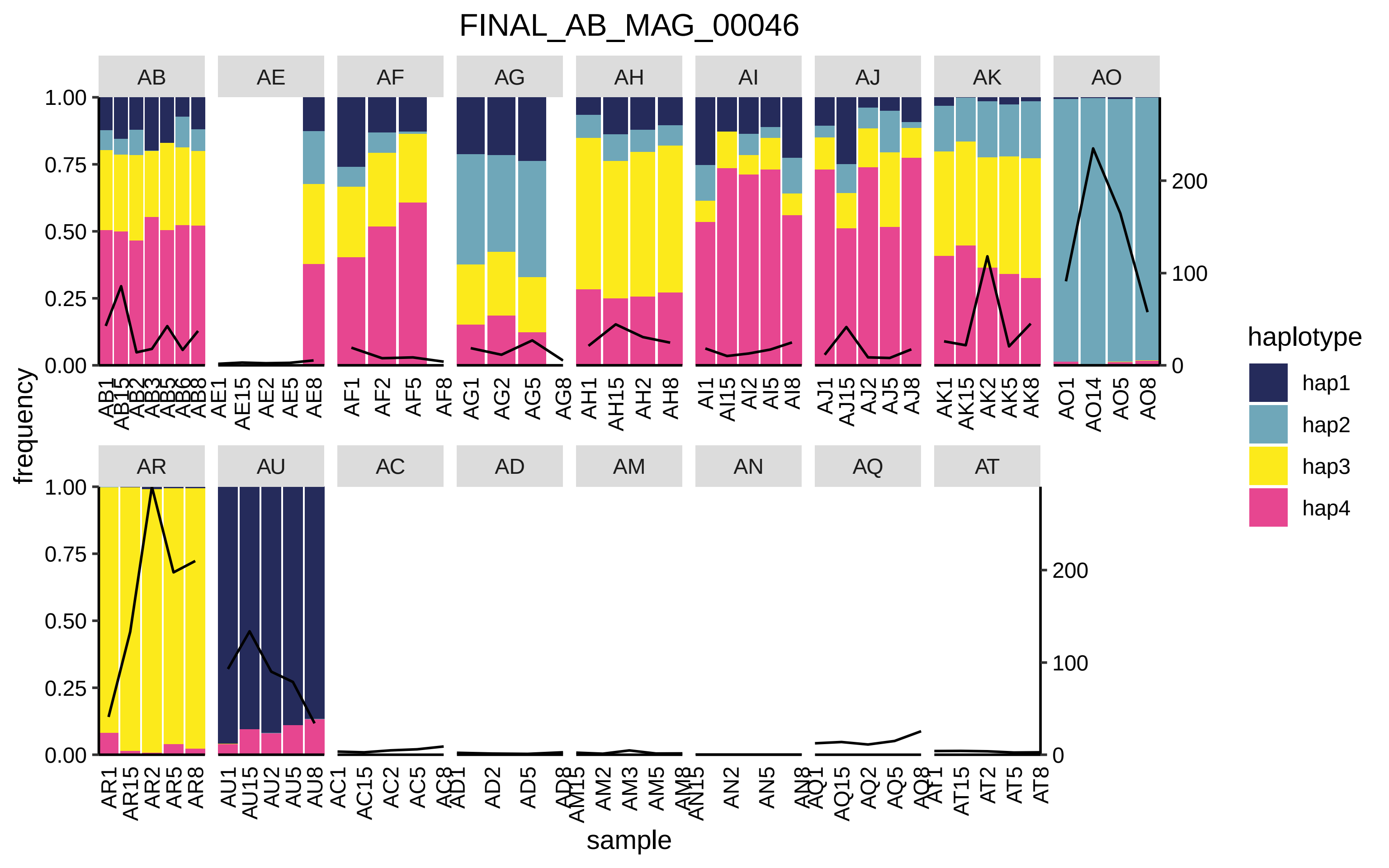
FINAL_AB_MAG_00046
AB
AE
AF
AG
AH
AI
AJ
AK
AO
AR
AU
AC
AD
AM
AN
AQ
AT
frequency
sample
haplotype
hap1
hap2
hap3
hap4

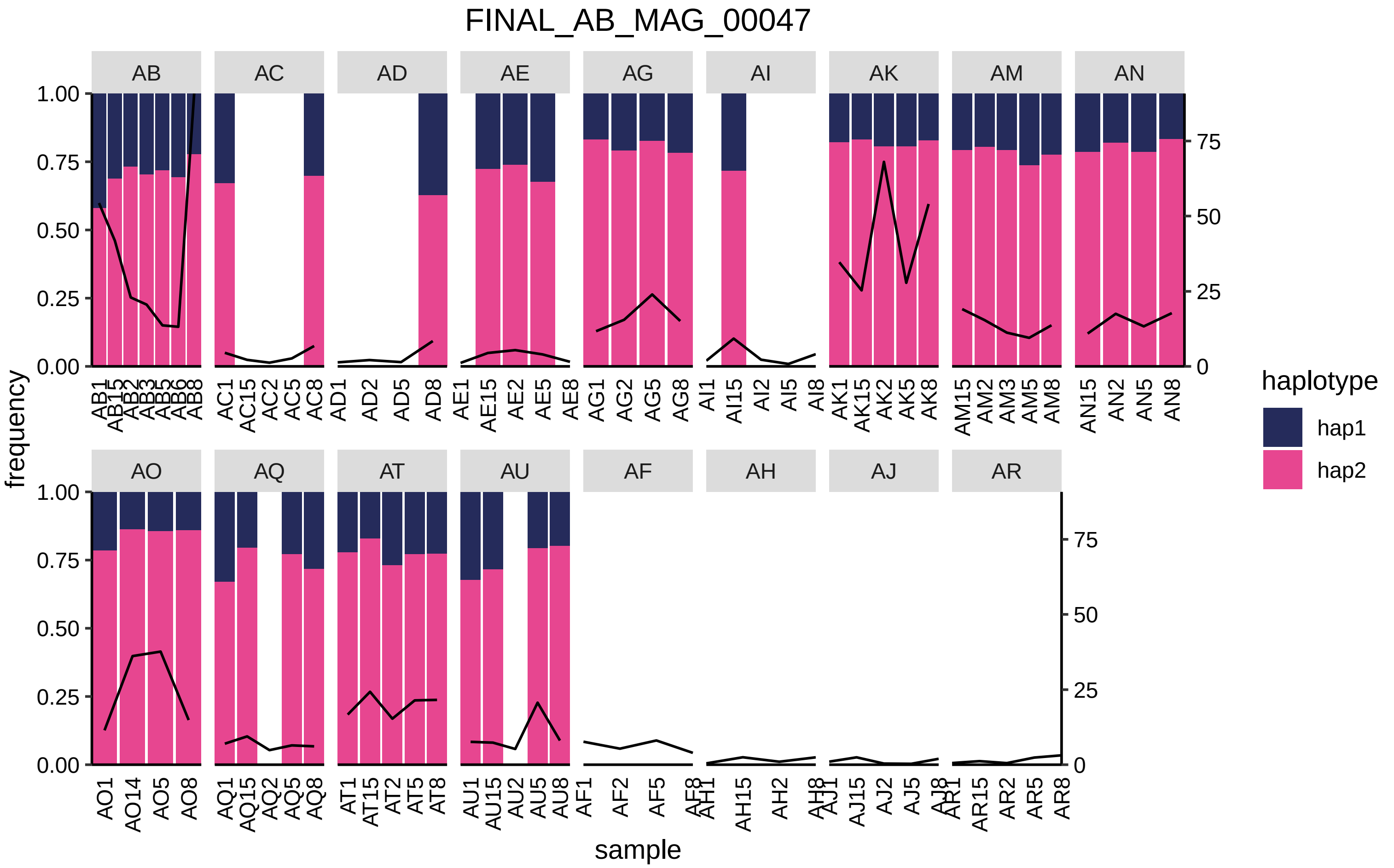

FINAL_AB_MAG_00047
AB
AC
AD
AE
AG
AI
AK
AM
AN
AO
AQ
AT
AU
AF
AH
AJ
AR
frequency
sample
1.00
0.75
0.50
0.25
0.00
75
50
25
0
AB1 AB15 AB2 AB3 AB5 AB6 AB8
AC1 AC15 AC2 AC5 AC8
AD1 AD2 AD5 AD8
AE1 AE15 AE2 AE5 AE8
AG1 AG2 AG5 AG8
AI1 AI15 AI2 AI5 AI8
AK1 AK15 AK2 AK5 AK8
AM15 AM2 AM3 AM5 AM8
AN15 AN2 AN5 AN8
AO1 AO14 AO5 AO8
AQ1 AQ15 AQ2 AQ5 AQ8
AT1 AT15 AT2 AT5 AT8
AU1 AU15 AU2 AU5 AU8
AF1 AF2 AF5 AF8
AH1 AH15 AH2 AH8
AJ1 AJ15 AJ2 AJ5 AJ8
AR1 AR15 AR2 AR5 AR8
haplotype
hap1
hap2

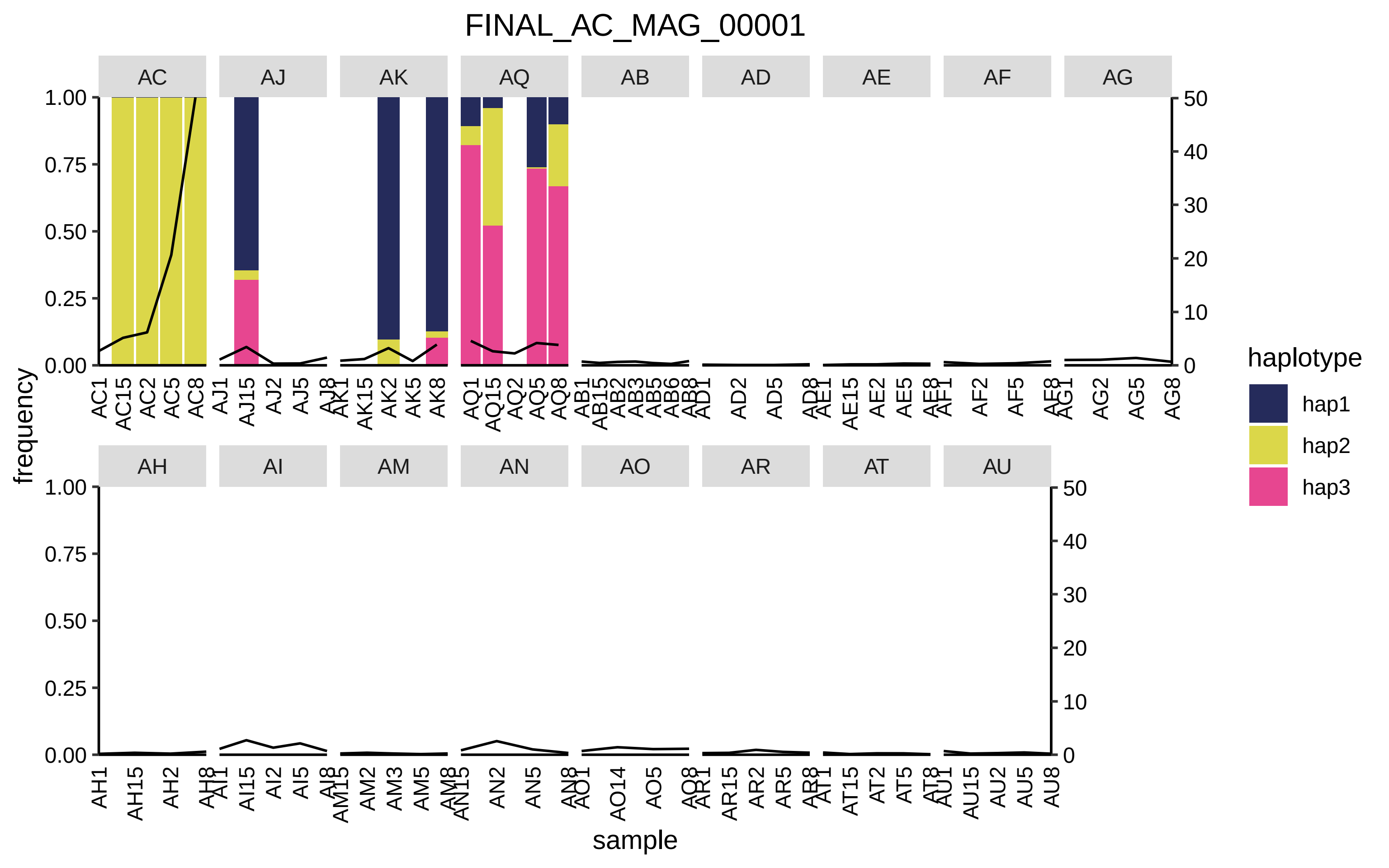

FINAL_AC_MAG_00001
AC
AJ
AK
AQ
AB
AD
AE
AF
AG
AH
AI
AM
AN
AO
AR
AT
AU
frequency
sample
haplotype
hap1
hap2
hap3

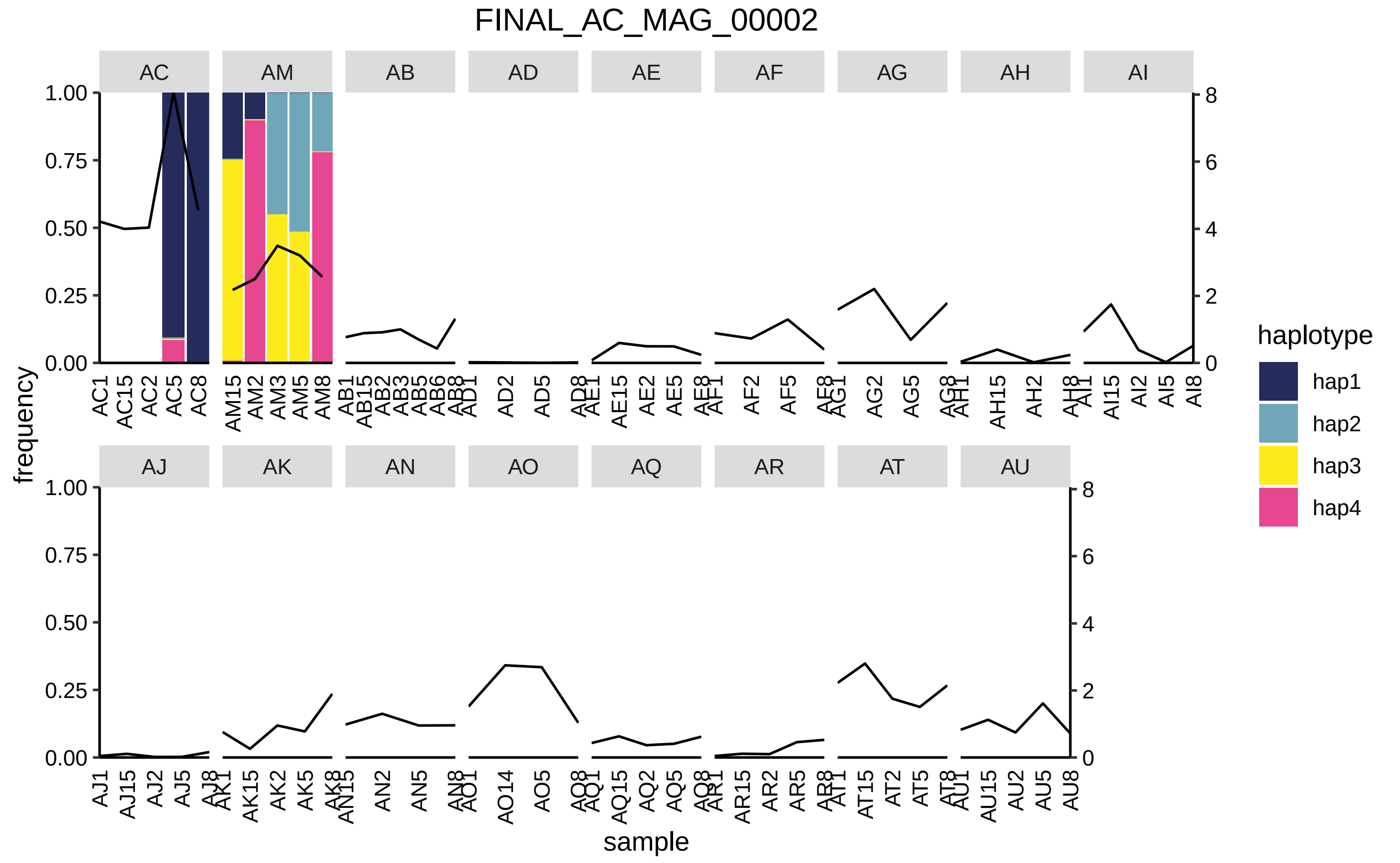
FINAL_AC_MAG_00002
AC
AM
AB
AD
AE
AF
AG
AH
AI
AJ
AK
AN
AO
AQ
AR
AT
AU
frequency
sample
haplotype
hap1
hap2
hap3
hap4
1.00
0.75
0.50
0.25
0.00
8
6
4
2
0

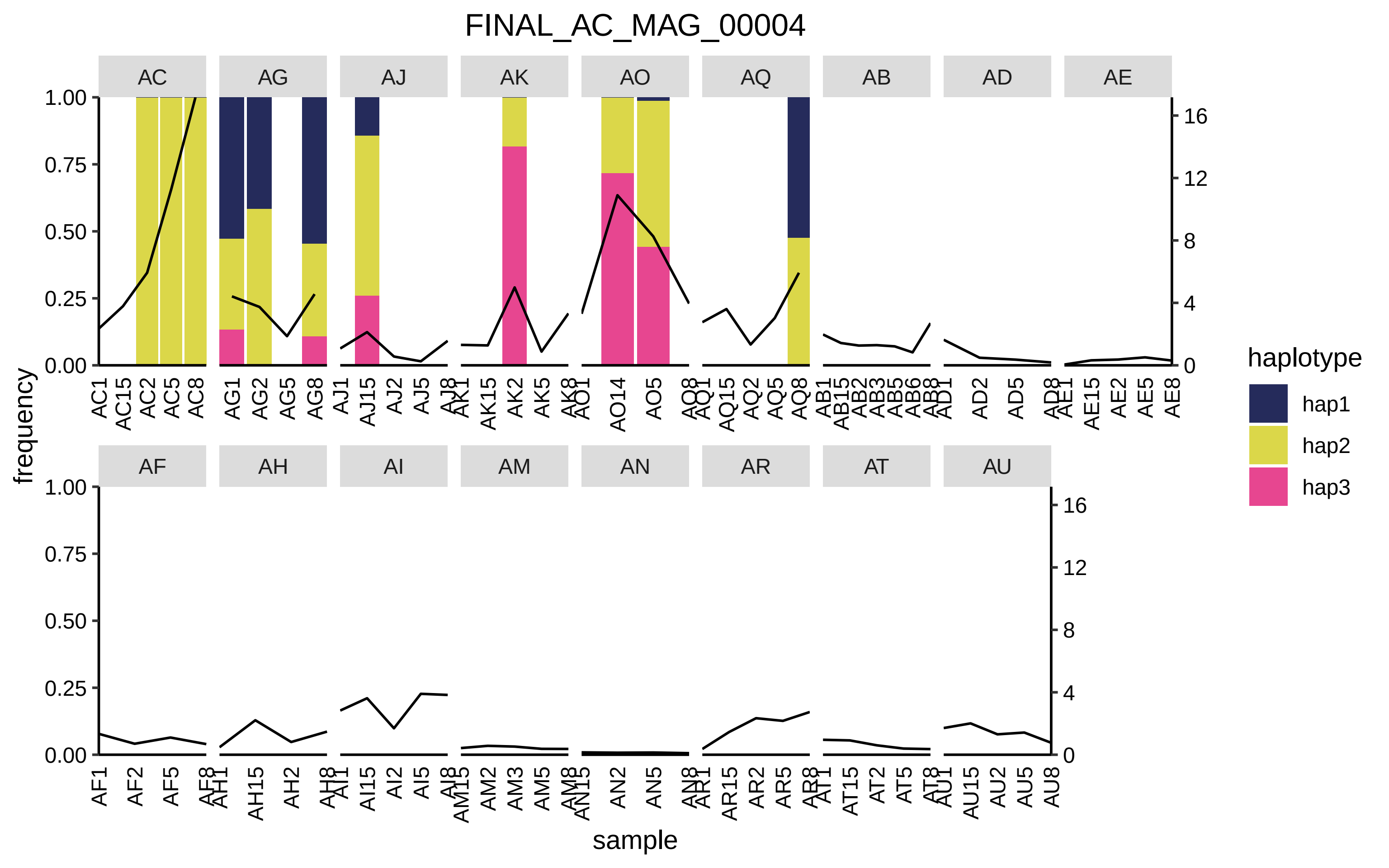

FINAL_AC_MAG_00004
AC
AG
AJ
AK
AO
AQ
AB
AD
AE
AF
AH
AI
AM
AN
AR
AT
AU
frequency
sample
haplotype
hap1
hap2
hap3
1.00
0.75
0.50
0.25
0.00
16
12
8
4
0

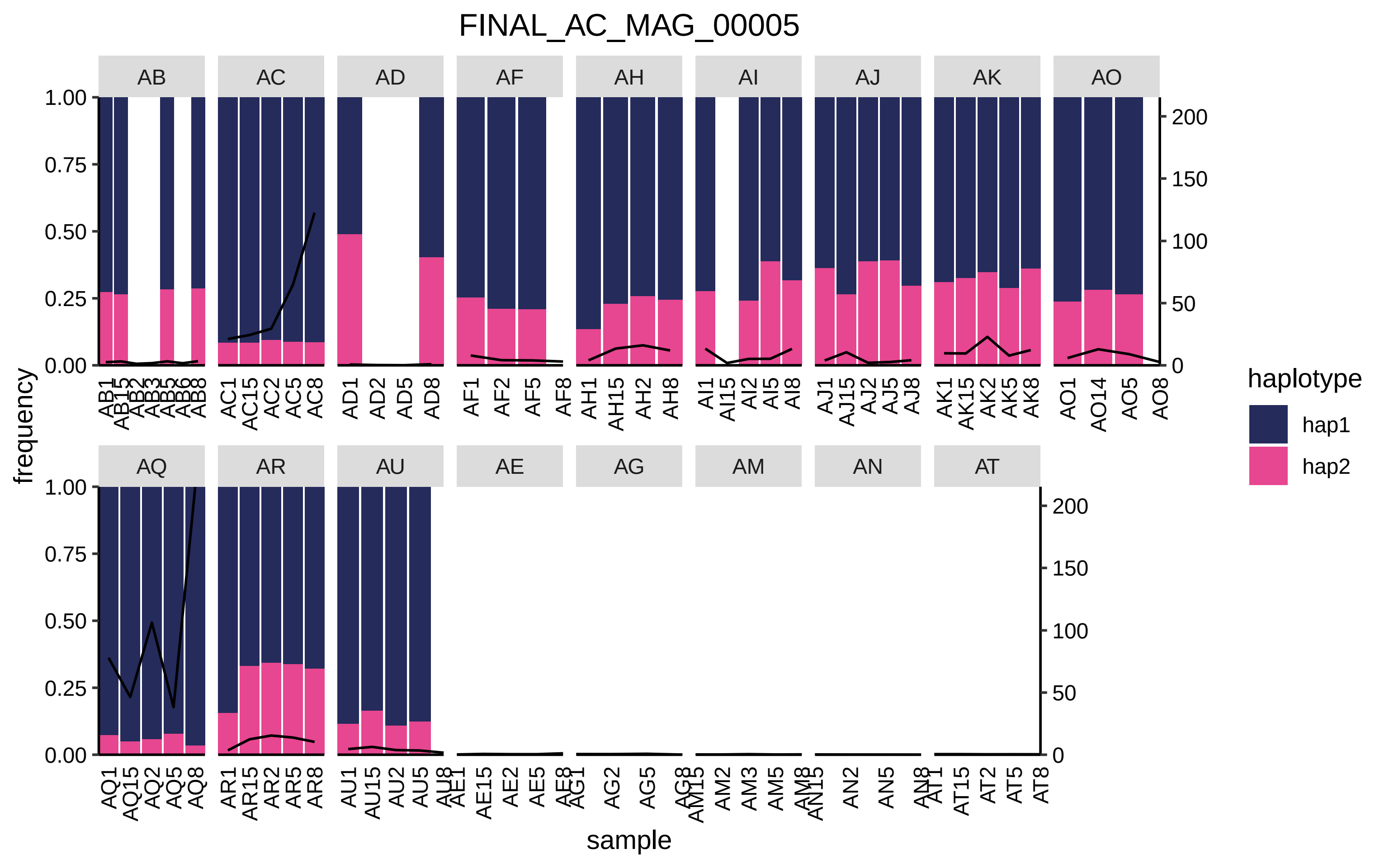

FINAL_AC_MAG_00005
AB
AC
AD
AF
AH
AI
AJ
AK
AO
AQ
AR
AU
AE
AG
AM
AN
AT
frequency
sample
haplotype
hap1
hap2
1.00
0.75
0.50
0.25
0.00
200
150
100
50
0
AC1
AC15
AC2
AC5
AC8
AD1
AD2
AD5
AD8
AF1
AF2
AF5
AF8
AH1
AH15
AH2
AH8
AI1
AI15
AI2
AI5
AI8
AJ1
AJ15
AJ2
AJ5
AJ8
AK1
AK15
AK2
AK5
AK8
AO1
AO14
AO5
AO8
AQ1
AQ15
AQ2
AQ5
AQ8
AR1
AR15
AR2
AR5
AR8
AU1
AU15
AU2
AU5
AE15
AE2
AE5
AG2
AG5
AM2
AM3
AM5
AN2
AN5
AT15
AT2
AT5
AT8

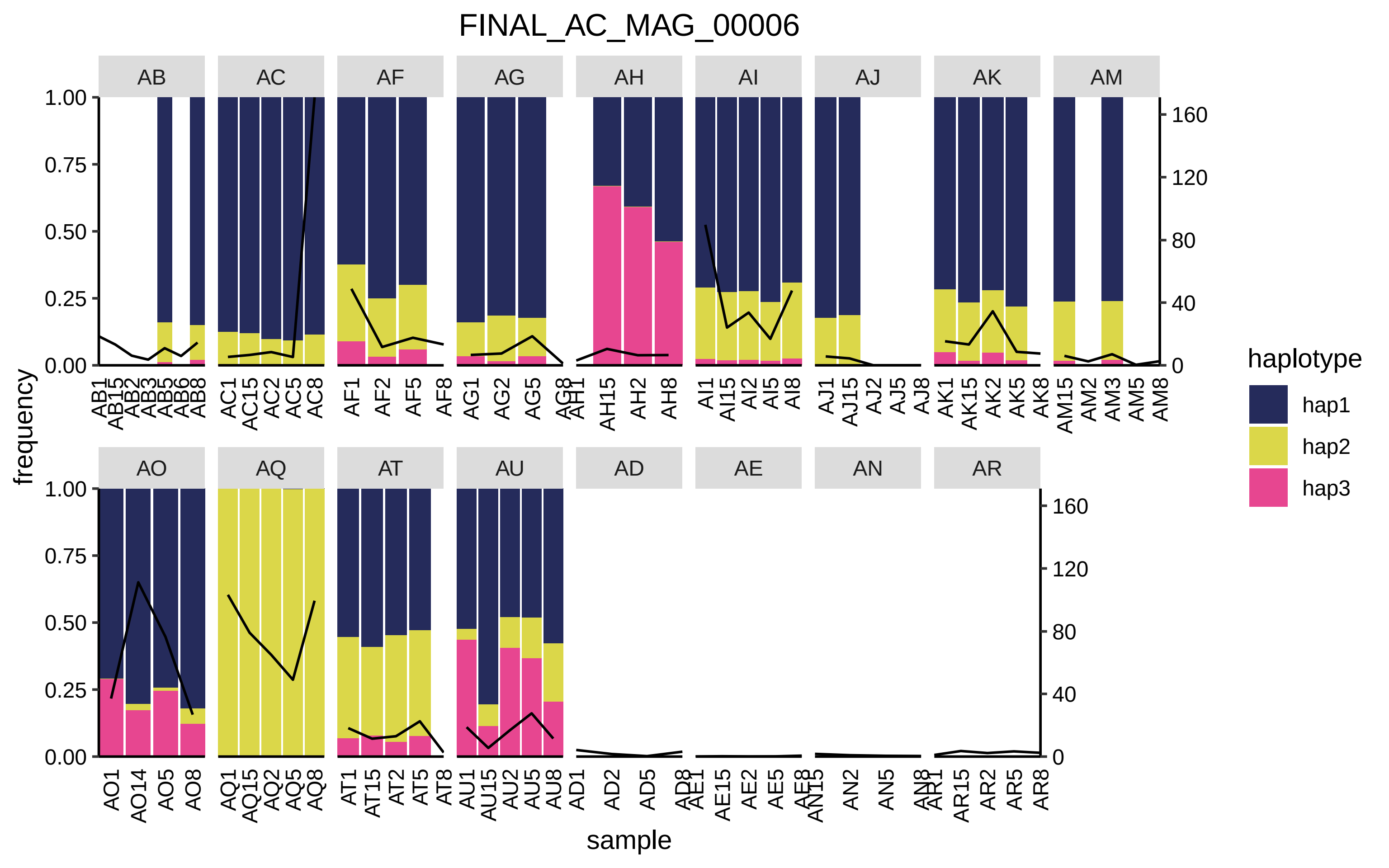

FINAL_AC_MAG_00006
AB
AC
AF
AG
AH
AI
AJ
AK
AM
AO
AQ
AT
AU
AD
AE
AN
AR
1.00
0.75
0.50
0.25
0.00
160
120
80
40
0
frequency
sample
haplotype
hap1
hap2
hap3

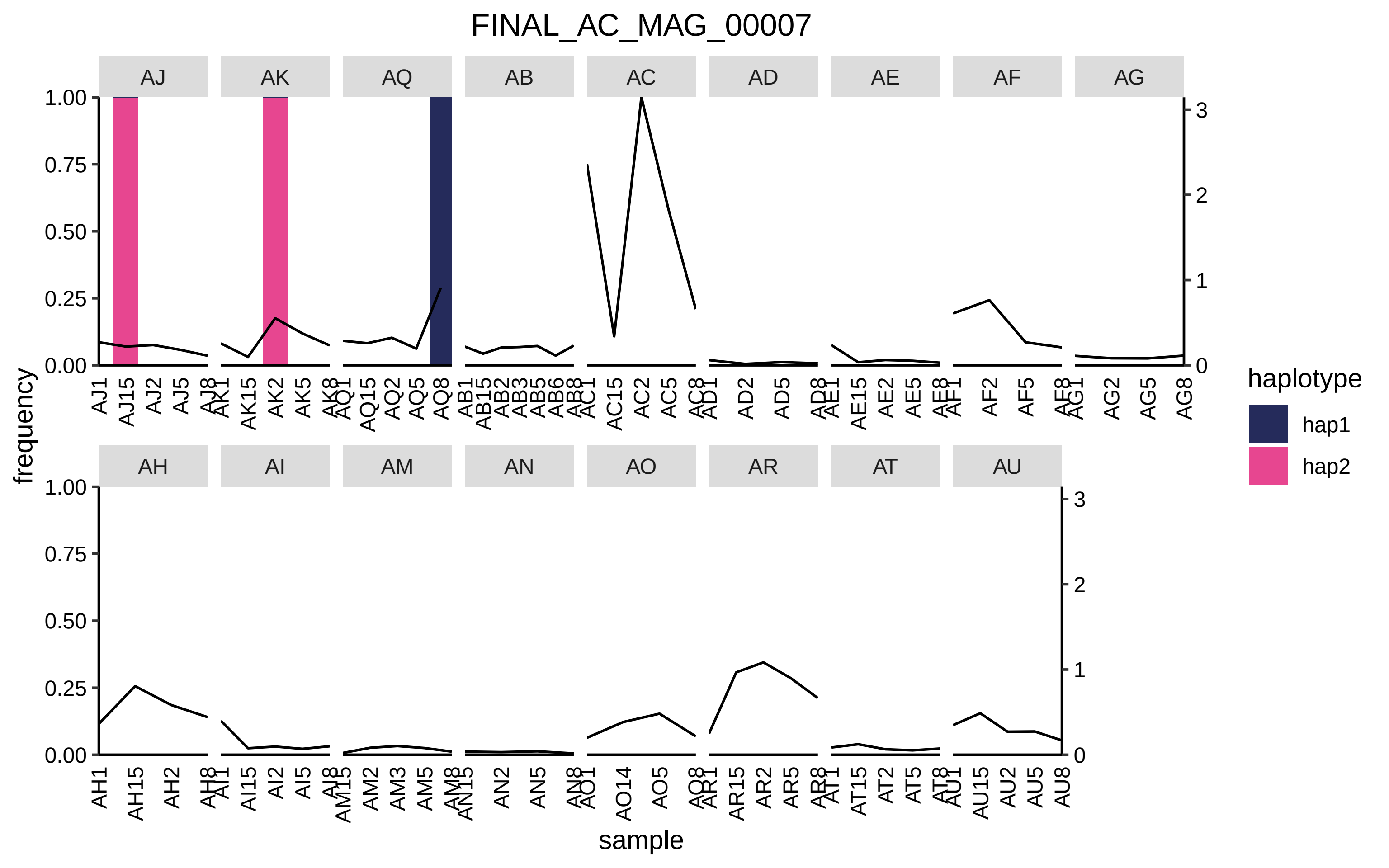

FINAL_AC_MAG_00007
AJ
AK
AQ
AB
AC
AD
AE
AF
AG
AH
AI
AM
AN
AO
AR
AT
AU
frequency
sample
haplotype
hap1
hap2
1.00
0.75
0.50
0.25
0.00
0
1
2
3

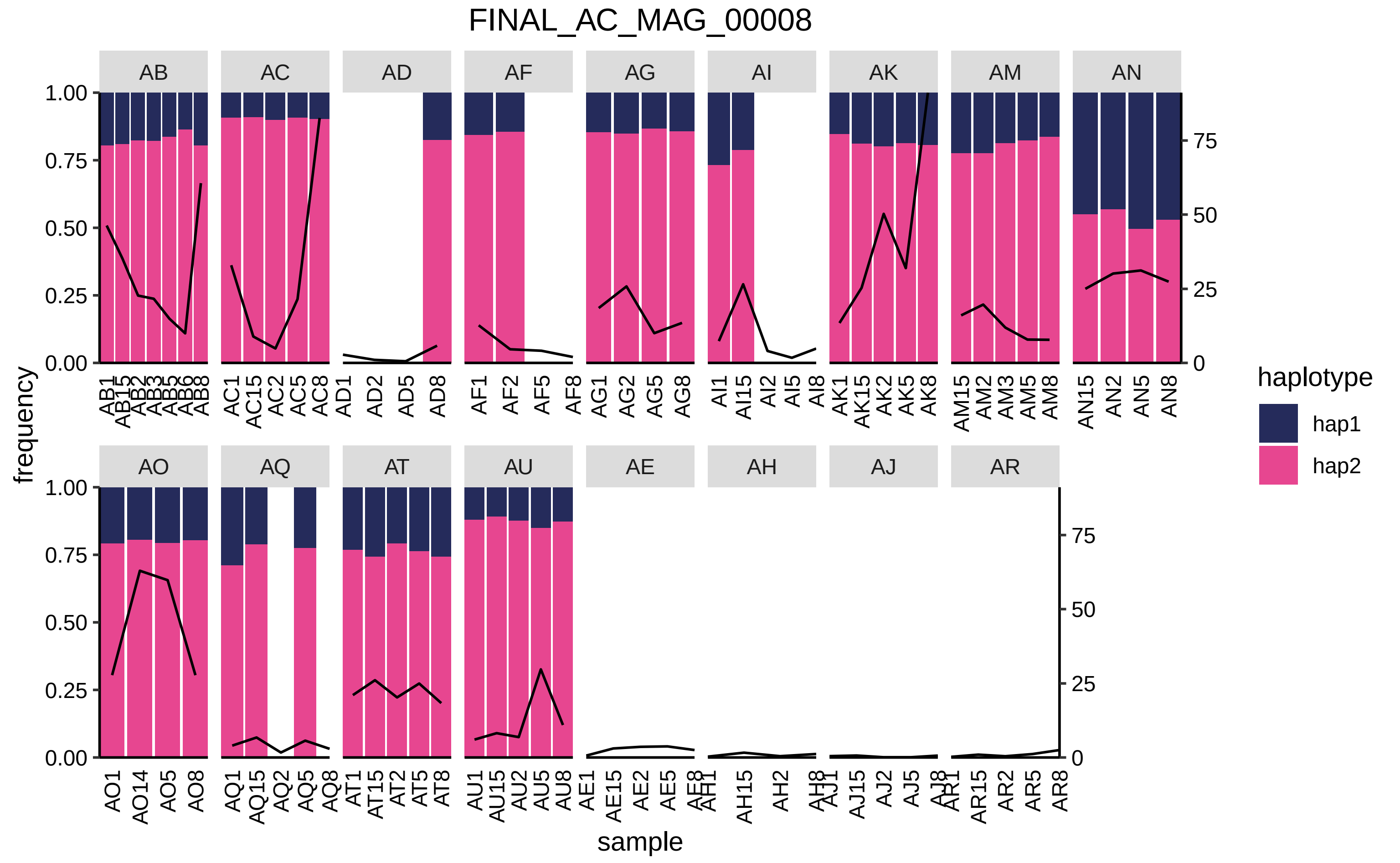
FINAL_AC_MAG_00008
AB
AC
AD
AF
AG
AI
AK
AM
AN
AO
AQ
AT
AU
AE
AH
AJ
AR
frequency
sample
haplotype
hap1
hap2
1.00
0.75
0.50
0.25
0.00
75
50
25
0
AB1 AB15 AB2 AB3 AB5 AB6 AB8
AC1 AC15 AC2 AC5 AC8
AD1 AD2 AD5 AD8
AF1 AF2 AF5 AF8
AG1 AG2 AG5 AG8
AI1 AI15 AI2 AI5 AI8
AK1 AK15 AK2 AK5 AK8
AM15 AM2 AM3 AM5 AM8
AN15 AN2 AN5 AN8
AO1 AO14 AO5 AO8
AQ1 AQ15 AQ2 AQ5 AQ8
AT1 AT15 AT2 AT5 AT8
AU1 AU15 AU2 AU5 AU8
AE1 AE15 AE2 AE5 AE8
AH1 AH15 AH2 AH8
AJ1 AJ15 AJ2 AJ5 AJ8
AR1 AR15 AR2 AR5 AR8

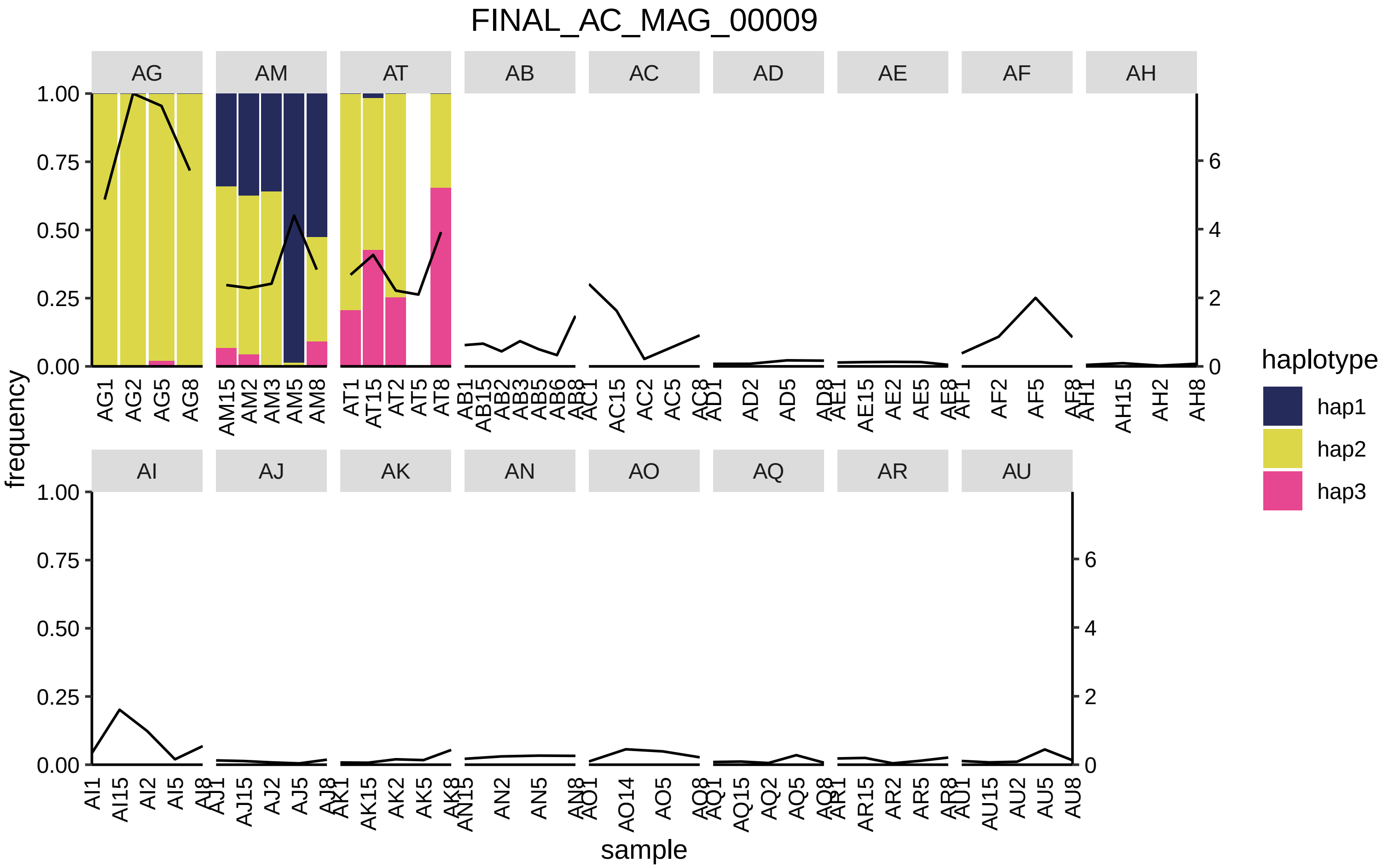
FINAL_AC_MAG_00009
AG
AM
AT
AB
AC
AD
AE
AF
AH
AI
AJ
AK
AN
AO
AQ
AR
AU
frequency
sample
haplotype
hap1
hap2
hap3
1.00
0.75
0.50
0.25
0.00
0
2
4
6
AG1 AG2 AG5 AG8
AM15 AM2 AM3 AM5 AM8
AT1 AT15 AT2 AT5 AT8
AB1 AB15 AB2 AB3 AB5 AB6 AB8
AC1 AC15 AC2 AC5 AC8
AD1 AD2 AD5 AD8
AE1 AE15 AE2 AE5 AE8
AF1 AF2 AF5 AF8
AH1 AH15 AH2 AH8
AI1 AI15 AI2 AI5 AI8
AJ1 AJ15 AJ2 AJ5 AJ8
AK1 AK15 AK2 AK5 AK8
AN15 AN2 AN5 AN8
AO1 AO14 AO5 AO8
AQ1 AQ15 AQ2 AQ5 AQ8
AR1 AR15 AR2 AR5 AR8
AU1 AU15 AU2 AU5 AU8

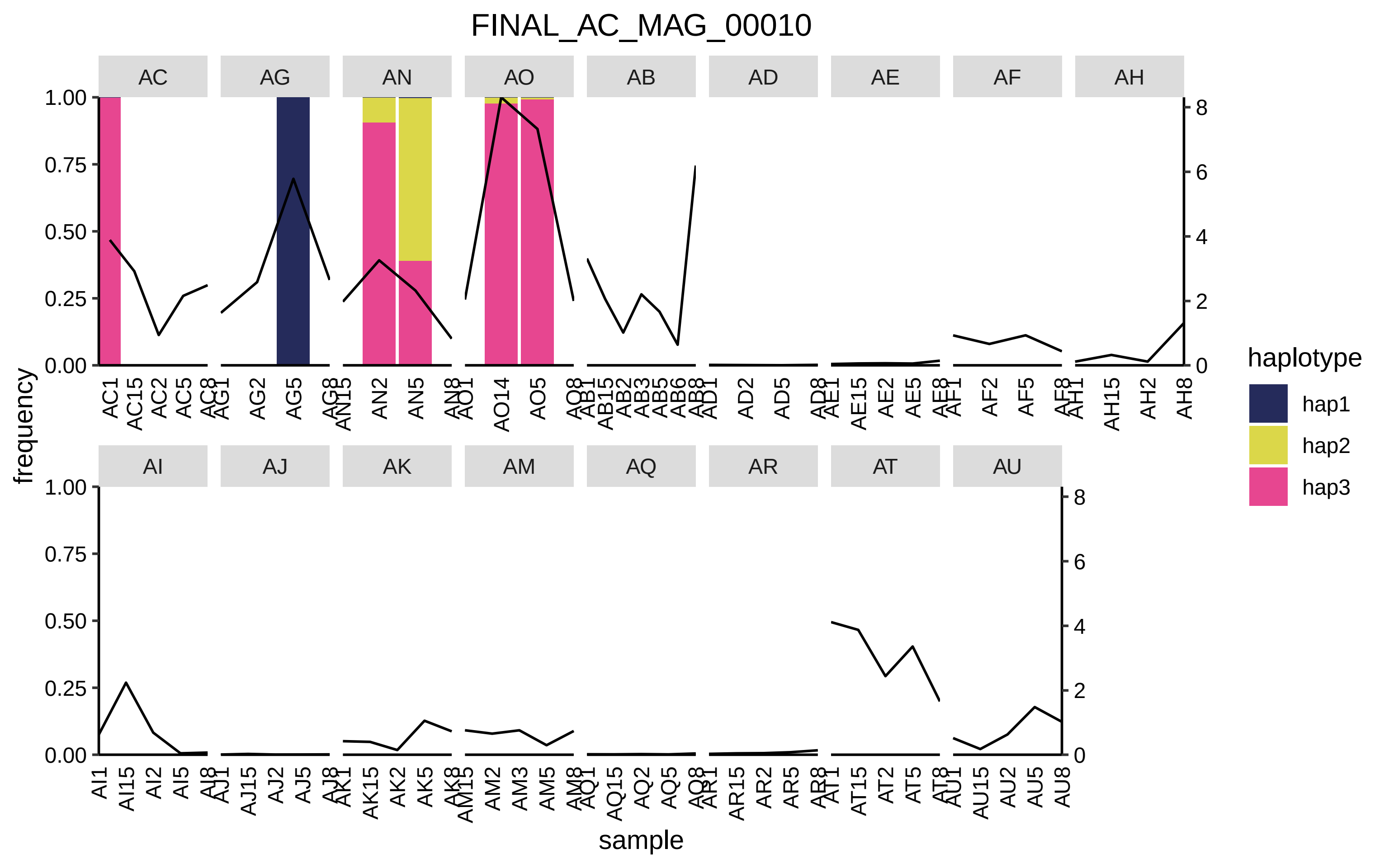
FINAL_AC_MAG_00010
AC
AG
AN
AO
AB
AD
AE
AF
AH
AI
AJ
AK
AM
AQ
AR
AT
AU
frequency
sample
haplotype
hap1
hap2
hap3
1.00
0.75
0.50
0.25
0.00
8
6
4
2
0
AC1
AC15
AC2
AC5
AC8
AG1
AG2
AG5
AG8
AN15
AN2
AN5
AN8
AO1
AO14
AO5
AO8
AB1
AB15
AB2
AB3
AB5
AB6
AB8
AD1
AD2
AD5
AD8
AE1
AE15
AE2
AE5
AE8
AF1
AF2
AF5
AF8
AH1
AH15
AH2
AH8
AI1
AI15
AI2
AI5
AI8
AJ1
AJ15
AJ2
AJ5
AJ8
AK1
AK15
AK2
AK5
AK8
AM15
AM2
AM3
AM5
AM8
AQ1
AQ15
AQ2
AQ5
AQ8
AR1
AR15
AR2
AR5
AR8
AT1
AT15
AT2
AT5
AT8
AU1
AU15
AU2
AU5
AU8

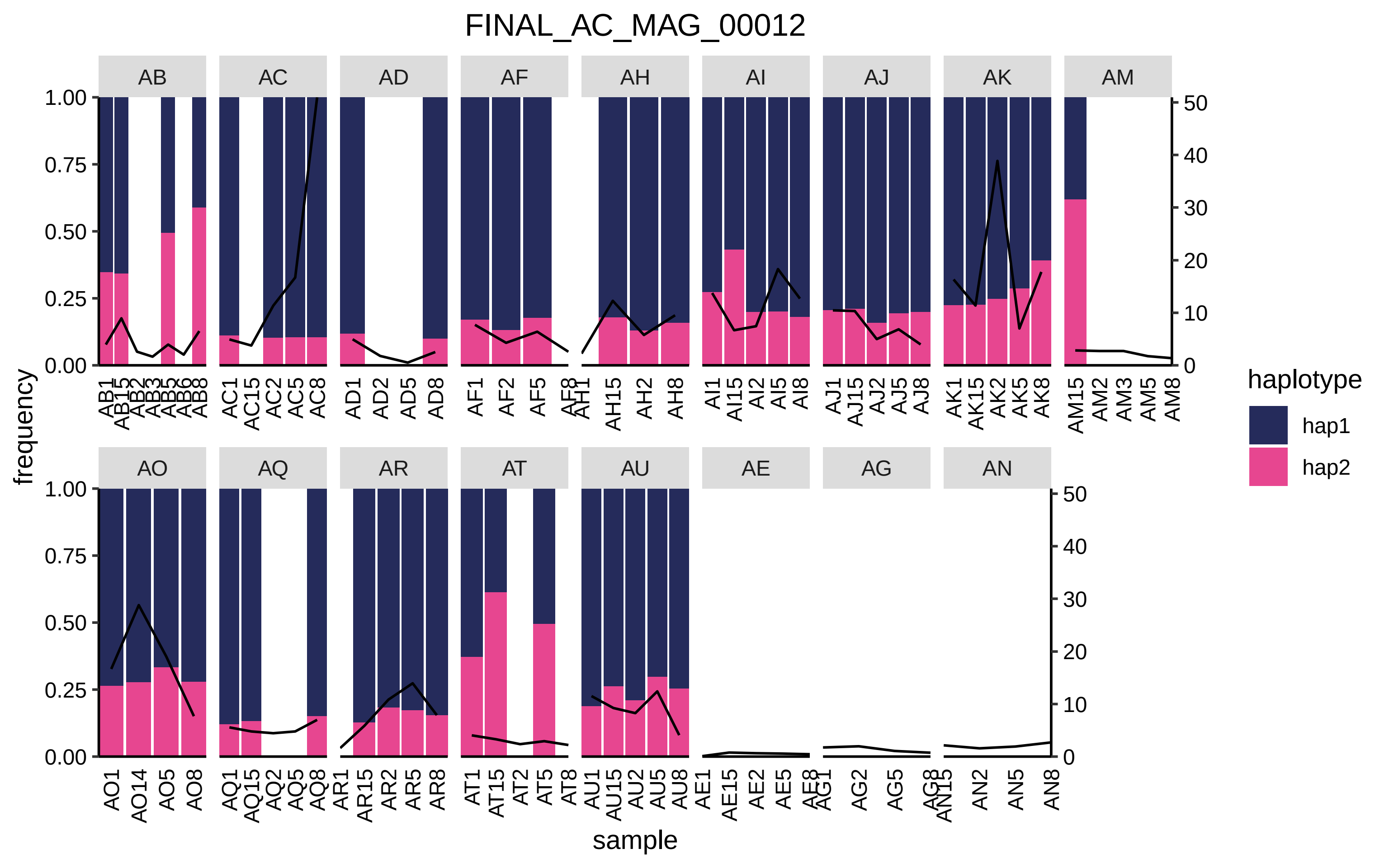
FINAL_AC_MAG_00012
AB
AC
AD
AF
AH
AI
AJ
AK
AM
AO
AQ
AR
AT
AU
AE
AG
AN
frequency
1.00
0.75
0.50
0.25
0.00
0
10
20
30
40
50
sample
AO1 AO14 AO5 AO8
AQ1 AQ15 AQ2 AQ5 AQ8
AR1 AR15 AR2 AR5 AR8
AT1 AT15 AT2 AT5 AT8
AU1 AU15 AU2 AU5 AU8
AN2 AN5 AN8
AM15 AM2 AM3 AM5 AM8
AK1 AK15 AK2 AK5 AK8
AJ1 AJ15 AJ2 AJ5 AJ8
AI1 AI15 AI2 AI5 AI8
AH15 AH2 AH8
AF1 AF2 AF5
AD1 AD2 AD5 AD8
AC1 AC15 AC2 AC5 AC8
AE1 AE15 AE2 AE5
AG2 AG5
haplotype
hap1
hap2

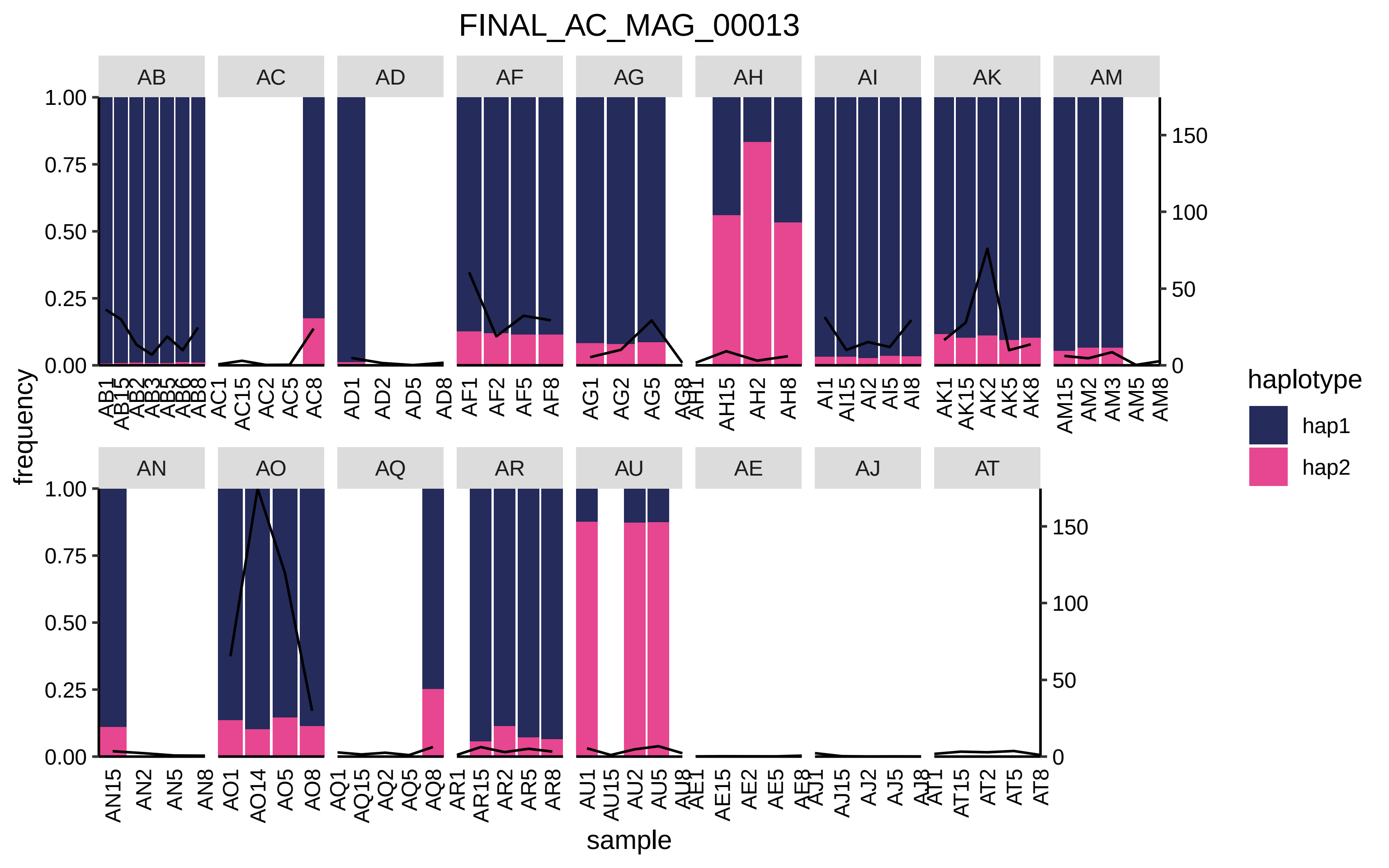

FINAL_AC_MAG_00013
AB
AC
AD
AF
AG
AH
AI
AK
AM
AN
AO
AQ
AR
AU
AE
AJ
AT
frequency
sample
haplotype
hap1
hap2

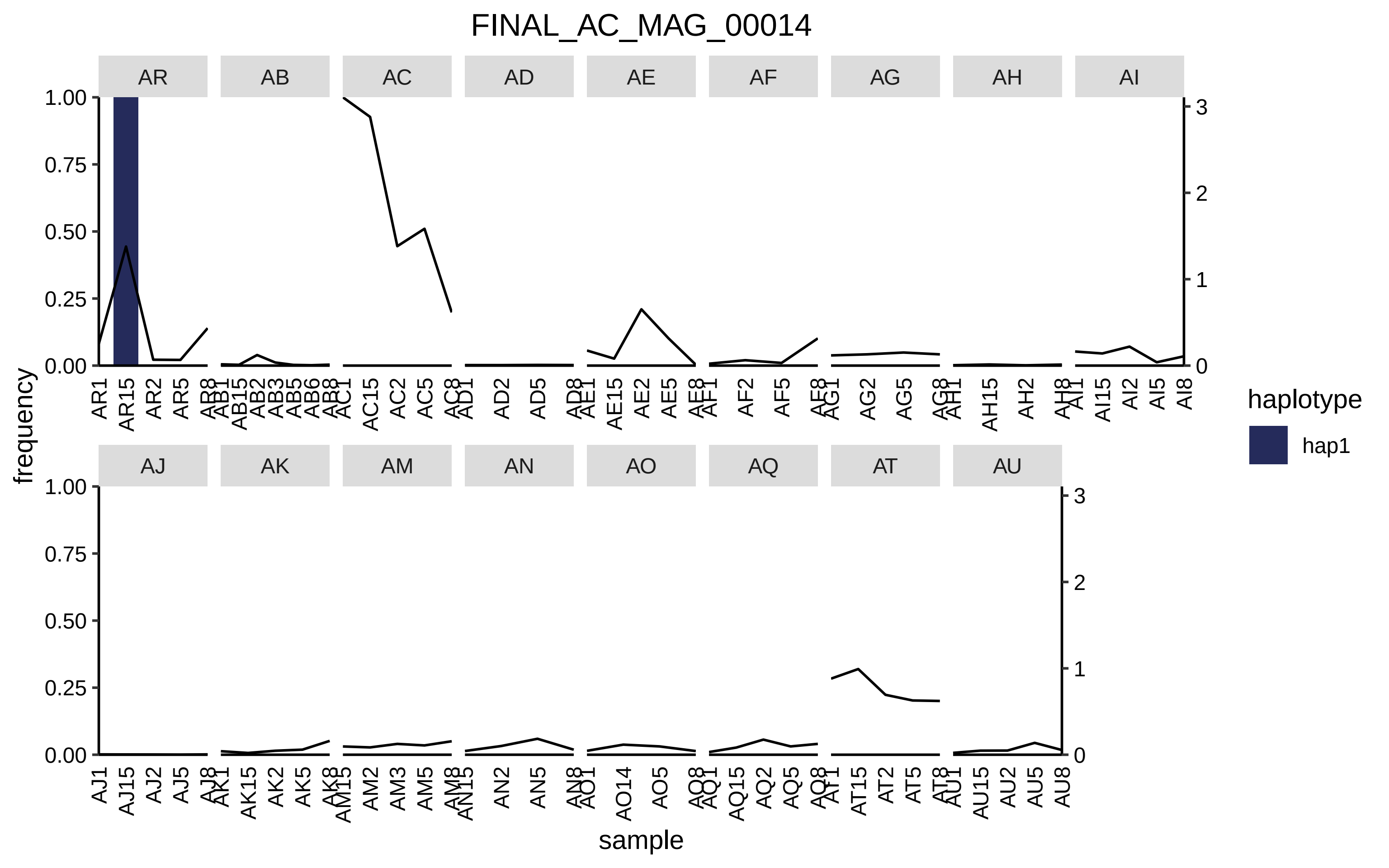
FINAL_AC_MAG_00014
AR
AB
AC
AD
AE
AF
AG
AH
AI
AJ
AK
AM
AN
AO
AQ
AT
AU
frequency
sample
haplotype
hap1

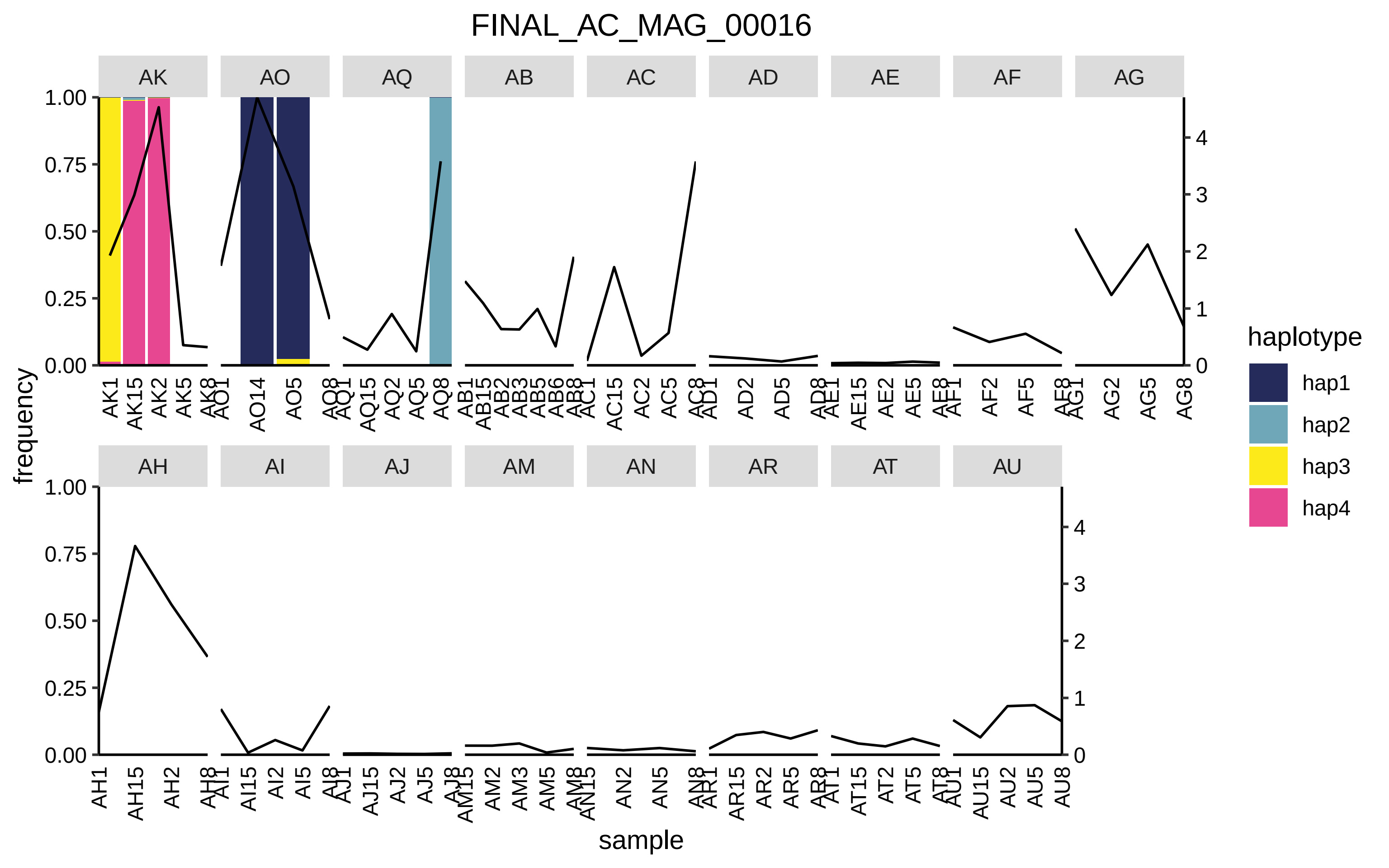

FINAL_AC_MAG_00016
AK
AO
AQ
AB
AC
AD
AE
AF
AG
AH
AI
AJ
AM
AN
AR
AT
AU
frequency
sample
haplotype
hap1
hap2
hap3
hap4
1.00
0.75
0.50
0.25
0.00
0
1
2
3
4
AK1 AK15 AK2 AK5 AK8
AO1 AO14 AO5 AO8
AQ1 AQ15 AQ2 AQ5 AQ8
AB1 AB15 AB2 AB3 AB5 AB6 AB8
AC1 AC15 AC2 AC5 AC8
AD1 AD2 AD5 AD8
AE1 AE15 AE2 AE5 AE8
AF1 AF2 AF5 AF8
AG1 AG2 AG5 AG8
AH1 AH15 AH2 AH8
AI1 AI15 AI2 AI5 AI8
AJ1 AJ15 AJ2 AJ5 AJ8
AM15 AM2 AM3 AM5 AM8
AN15 AN2 AN5 AN8
AR1 AR15 AR2 AR5 AR8
AT1 AT15 AT2 AT5 AT8
AU1 AU15 AU2 AU5 AU8

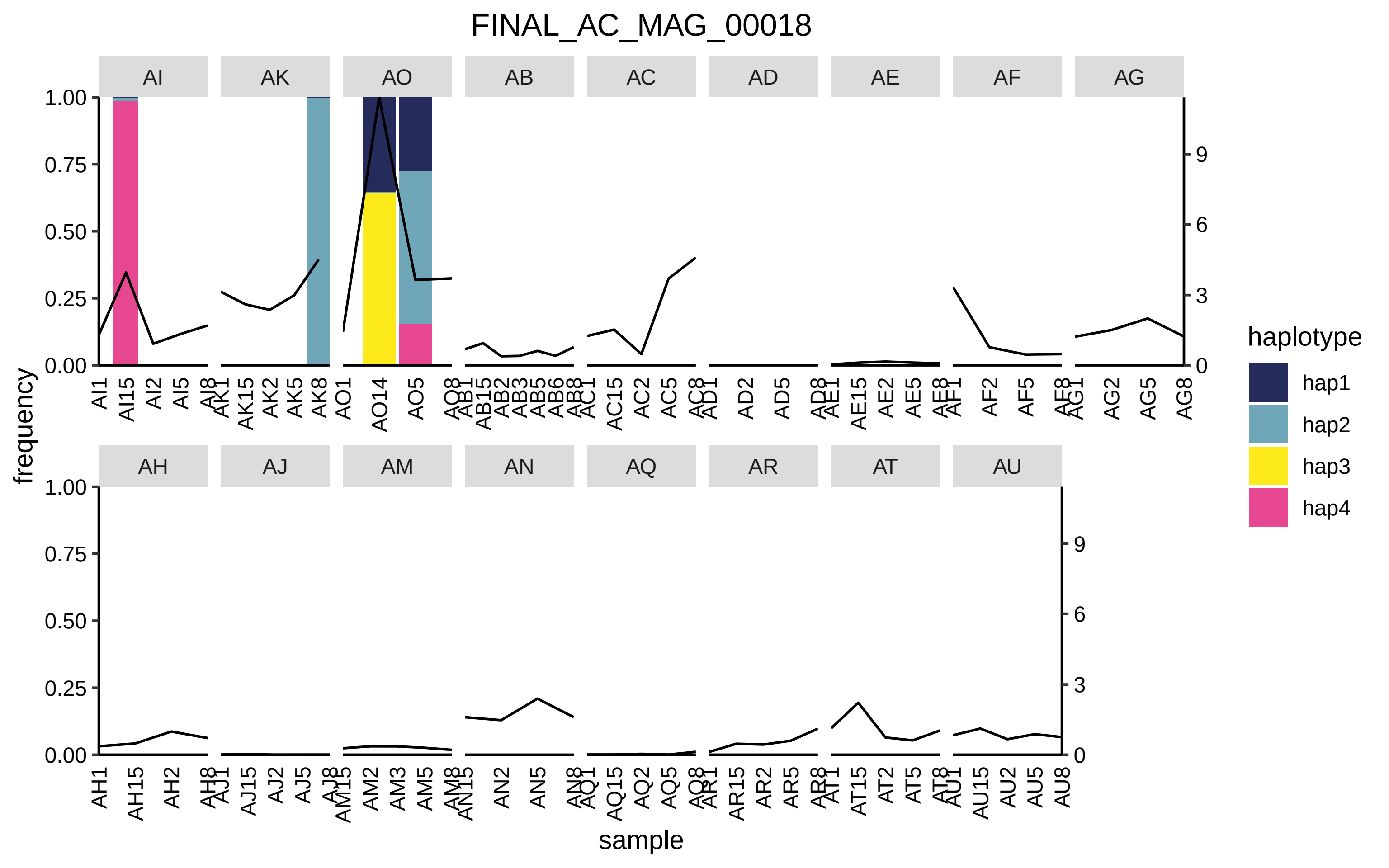

FINAL_AC_MAG_00018
AI
AK
AO
AB
AC
AD
AE
AF
AG
AH
AJ
AM
AN
AQ
AR
AT
AU
frequency
sample
haplotype
hap1
hap2
hap3
hap4

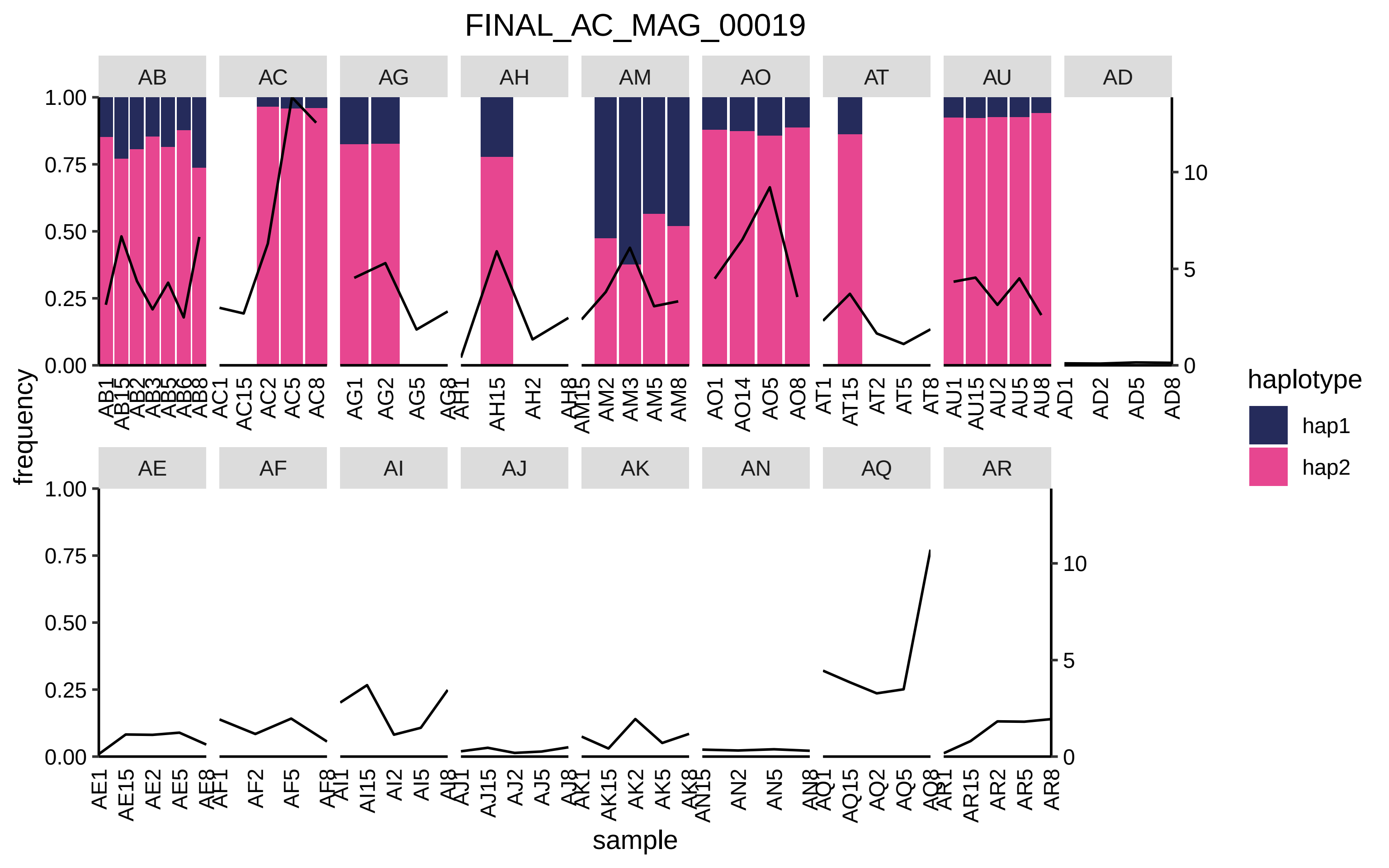
FINAL_AC_MAG_00019
AB
AC
AG
AH
AM
AO
AT
AU
AD
AE
AF
AI
AJ
AK
AN
AQ
AR
frequency
1.00
0.75
0.50
0.25
0.00
0
5
10
sample
haplotype
hap1
hap2
AB1 AB15 AB2 AB3 AB5 AB6 AB8
AC1 AC15 AC2 AC5 AC8
AG1 AG2 AG5 AG8
AH1 AH15 AH2 AH8
AM15 AM2 AM3 AM5 AM8
AO1 AO14 AO5 AO8
AT1 AT15 AT2 AT5 AT8
AU1 AU15 AU2 AU5 AU8
AD1 AD2 AD5 AD8
AE1 AE15 AE2 AE5 AE8
AF1 AF2 AF5 AF8
AI1 AI15 AI2 AI5 AI8
AJ1 AJ15 AJ2 AJ5 AJ8
AK1 AK15 AK2 AK5 AK8
AN15 AN2 AN5 AN8
AQ1 AQ15 AQ2 AQ5 AQ8
AR1 AR15 AR2 AR5 AR8

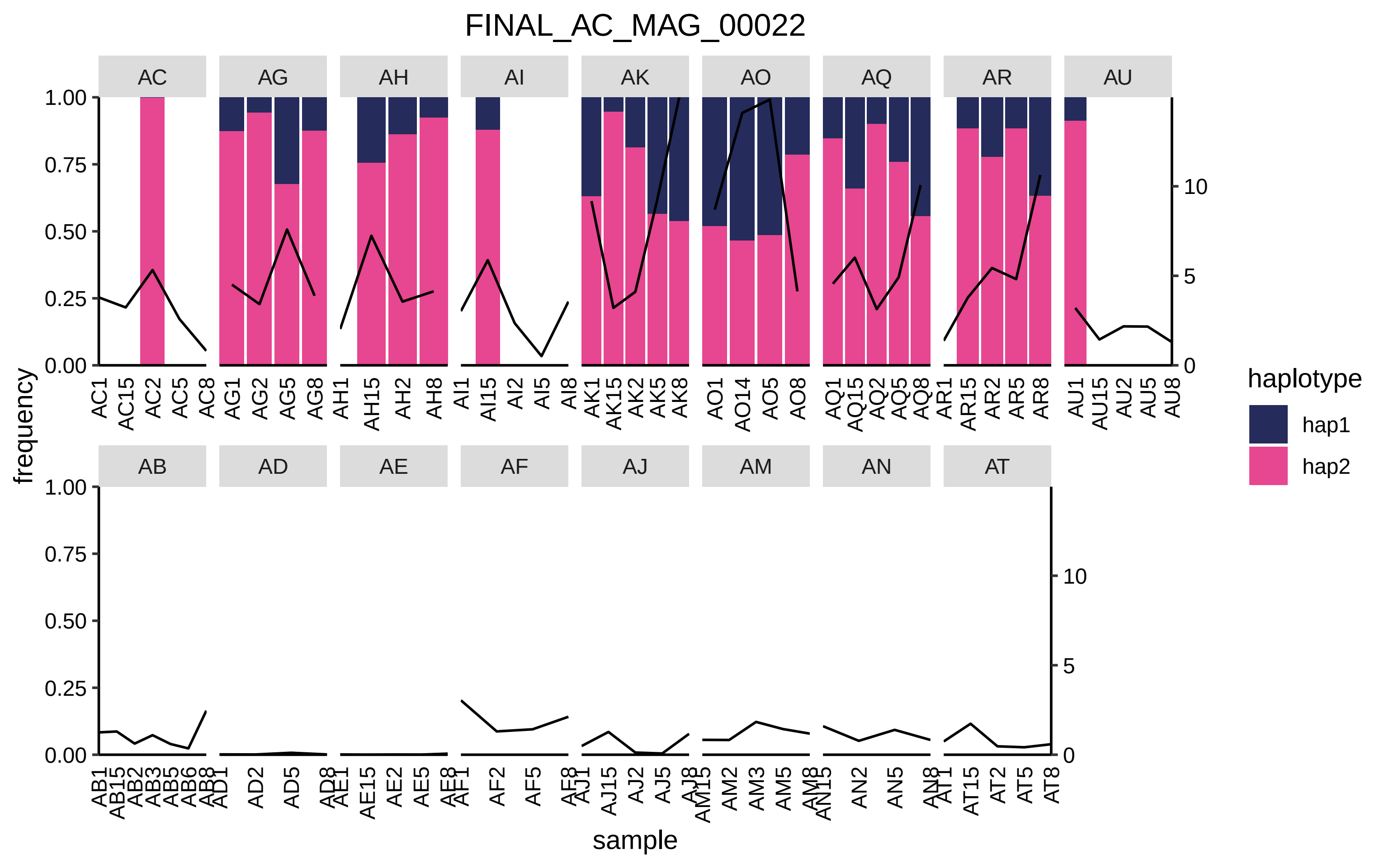

FINAL_AC_MAG_00022
AC
AG
AH
AI
AK
AO
AQ
AR
AU
AB
AD
AE
AF
AJ
AM
AN
AT
frequency
1.00
0.75
0.50
0.25
0.00
0
5
10
sample
haplotype
hap1
hap2
AC1 AC15 AC2 AC5 AC8 AG1 AG2 AG5 AG8 AH1 AH15 AH2 AH8 AI1 AI15 AI2 AI5 AI8 AK1 AK15 AK2 AK5 AK8 AO1 AO14 AO5 AO8 AQ1 AQ15 AQ2 AQ5 AQ8 AR1 AR15 AR2 AR5 AR8 AU1 AU15 AU2 AU5 AU8
AB1 AB15 AB2 AB3 AB5 AB6 AB8 AD1 AD2 AD5 AD8 AE1 AE15 AE2 AE5 AE8 AF1 AF2 AF5 AF8 AJ1 AJ15 AJ2 AJ5 AJ8 AM15 AM2 AM3 AM5 AM8 AN15 AN2 AN5 AN8 AT1 AT15 AT2 AT5 AT8

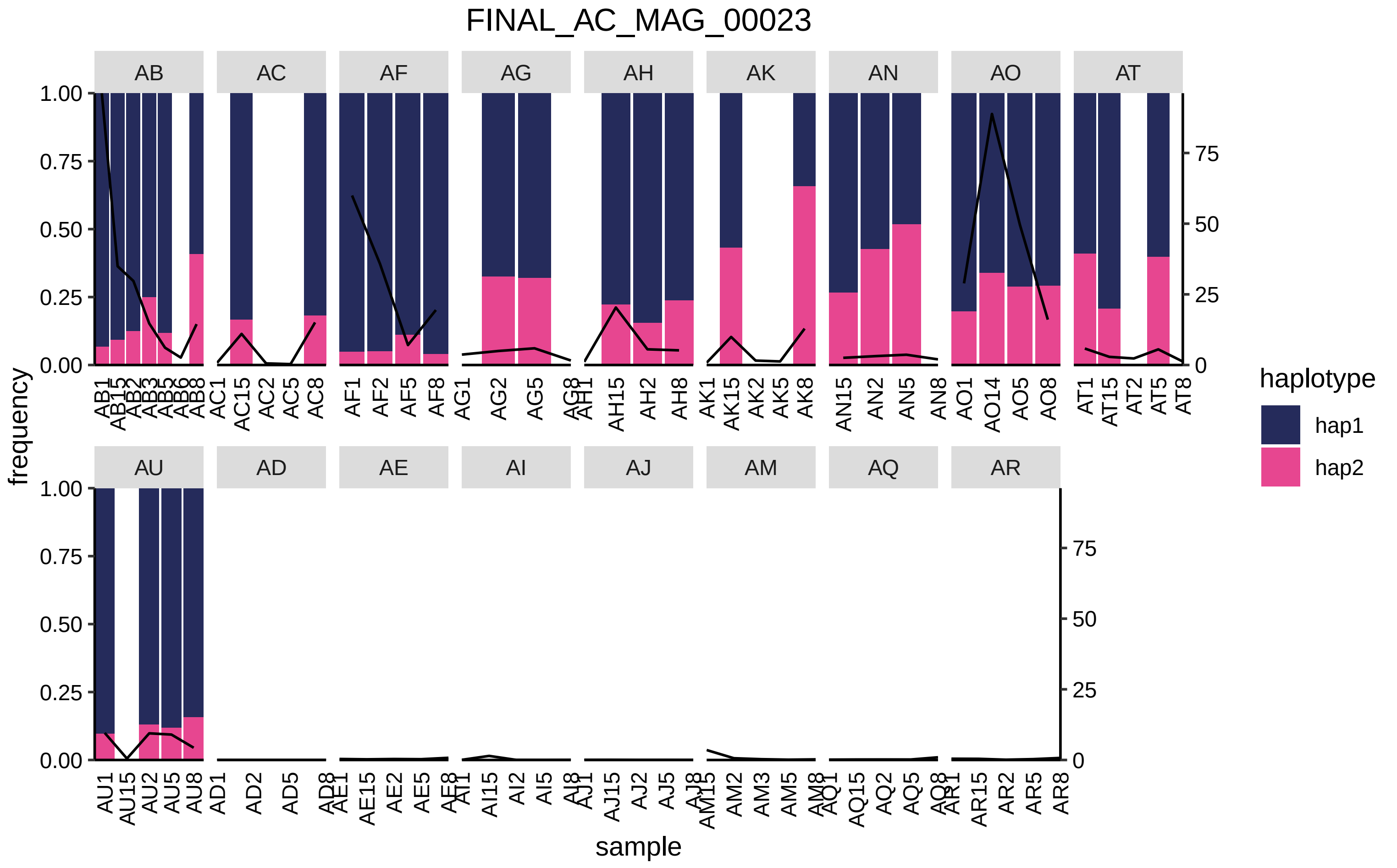
FINAL_AC_MAG_00023
AB
AC
AF
AG
AH
AK
AN
AO
AT
AU
AD
AE
AI
AJ
AM
AQ
AR
1.00
0.75
0.50
0.25
0.00
75
50
25
0
frequency
sample
haplotype
hap1
hap2
AB1 AB15 AB2 AB3 AB5 AB6 AB8
AC1 AC15 AC2 AC5 AC8
AF1 AF2 AF5 AF8
AG1 AG2 AG5 AG8
AH1 AH15 AH2 AH8
AK1 AK15 AK2 AK5 AK8
AN15 AN2 AN5 AN8
AO1 AO14 AO5 AO8
AT1 AT15 AT2 AT5 AT8
AU1 AU15 AU2 AU5 AU8
AD1 AD2 AD5 AD8
AE1 AE15 AE2 AE5 AE8
AI1 AI15 AI2 AI5 AI8
AJ1 AJ15 AJ2 AJ5 AJ8
AM15 AM2 AM3 AM5 AM8
AQ1 AQ15 AQ2 AQ5 AQ8
AR1 AR15 AR2 AR5 AR8

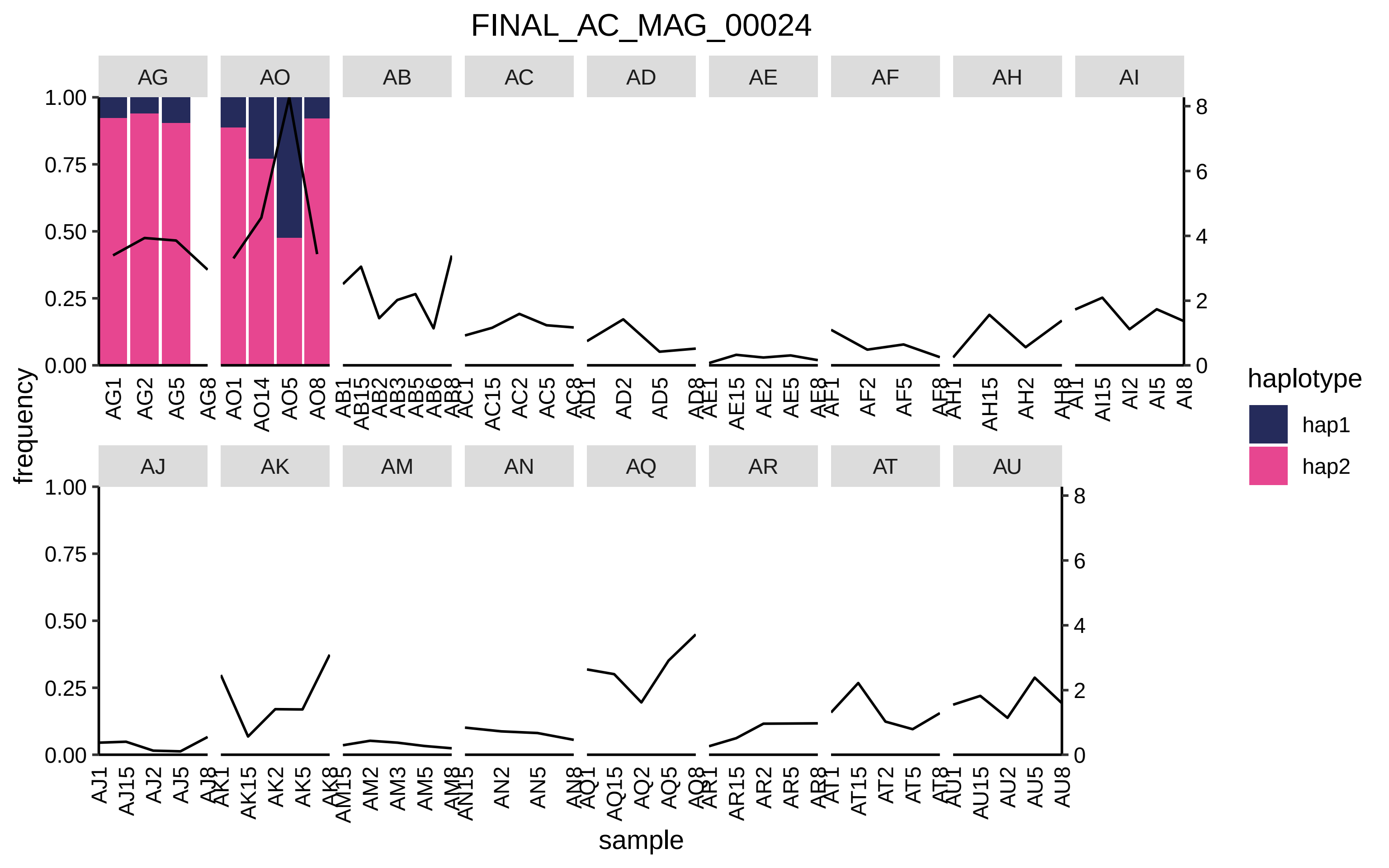

FINAL_AC_MAG_00024
AG
AO
AB
AC
AD
AE
AF
AH
AI
AJ
AK
AM
AN
AQ
AR
AT
AU
frequency
sample
haplotype
hap1
hap2
1.00
0.75
0.50
0.25
0.00
0
2
4
6
8

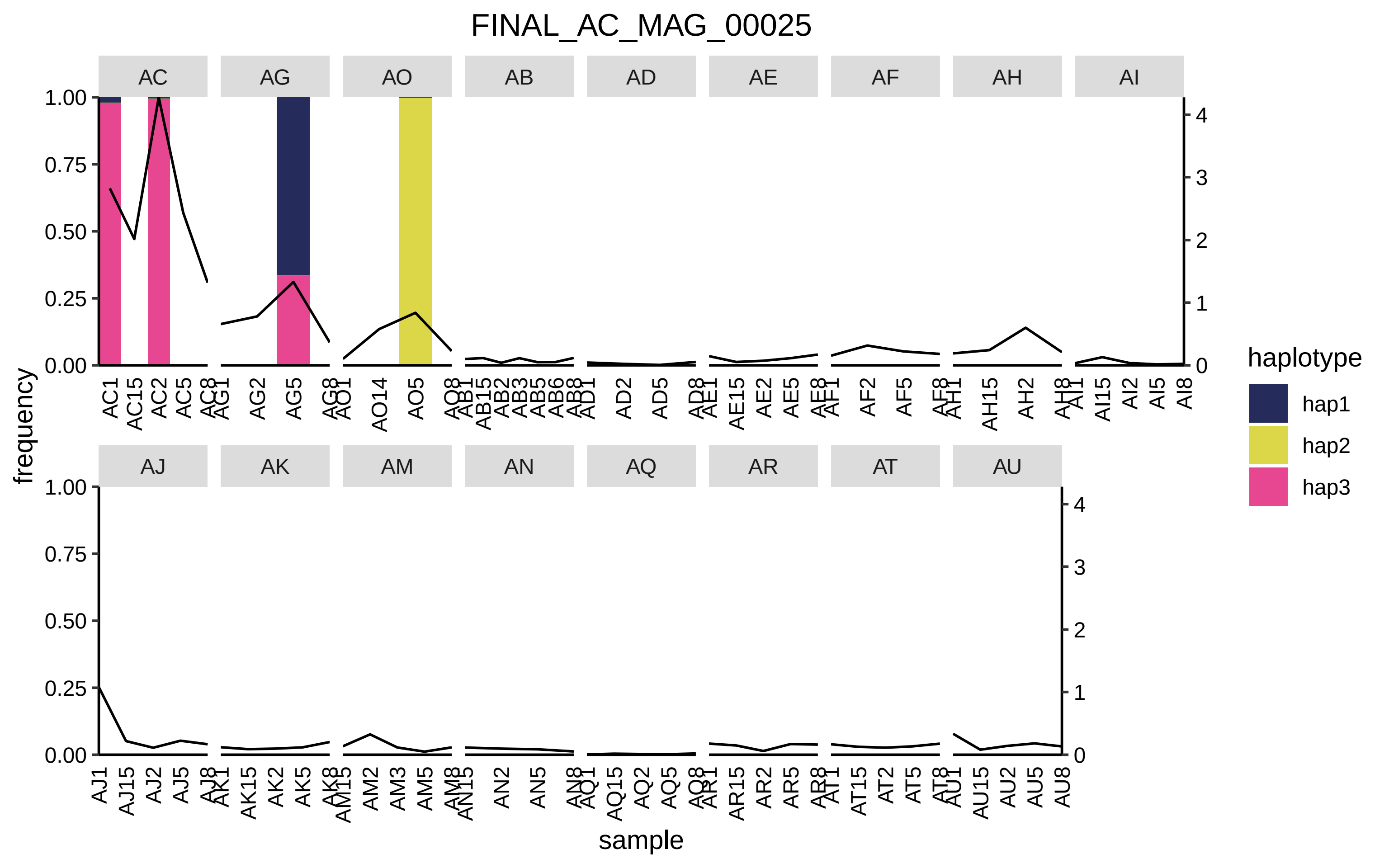
FINAL_AC_MAG_00025
AC
AG
AO
AB
AD
AE
AF
AH
AI
AJ
AK
AM
AN
AQ
AR
AT
AU
frequency
sample
haplotype
hap1
hap2
hap3
1.00
0.75
0.50
0.25
0.00
0
1
2
3
4

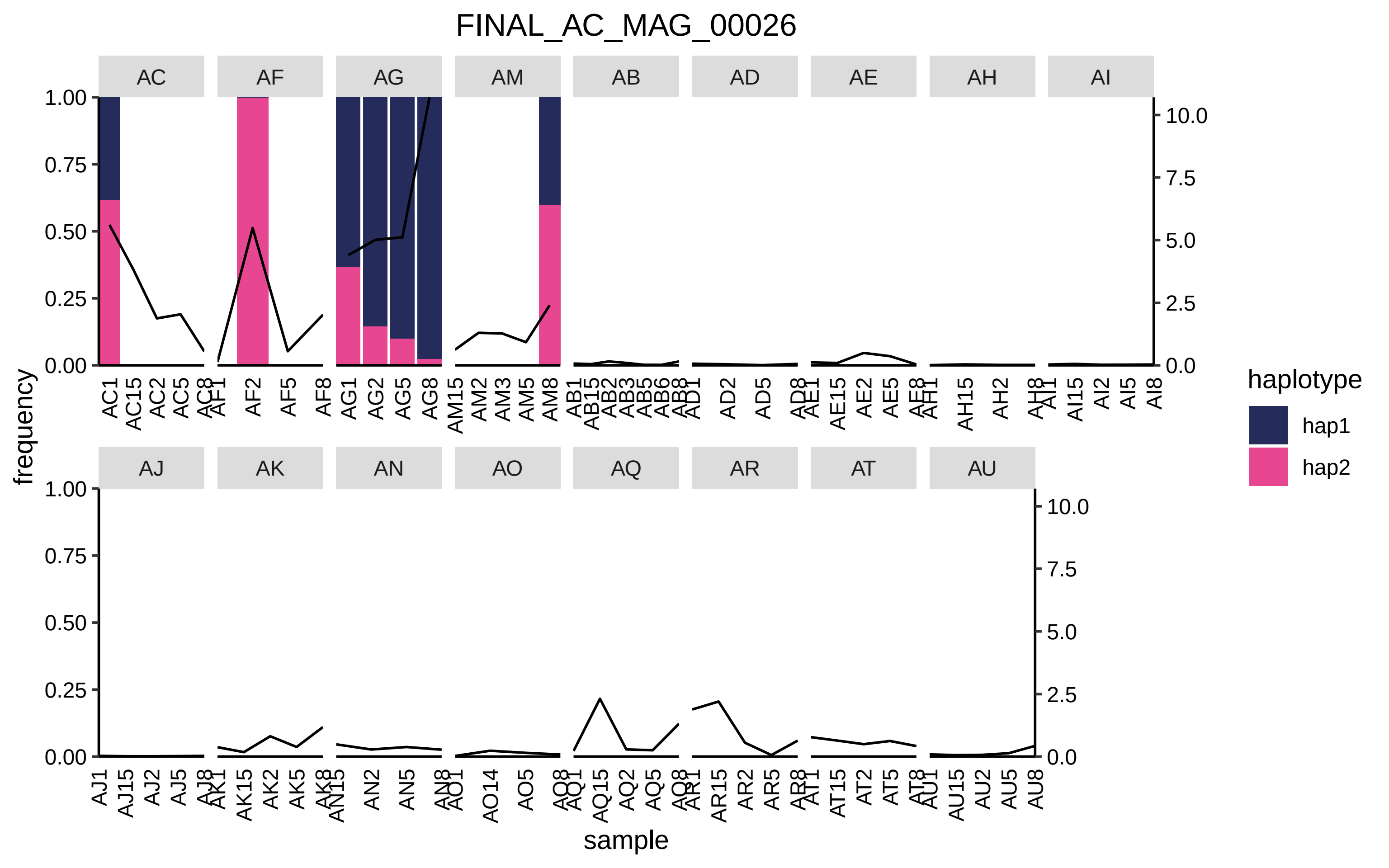

FINAL_AC_MAG_00026
AC
AF
AG
AM
AB
AD
AE
AH
AI
AJ
AK
AN
AO
AQ
AR
AT
AU
frequency
sample
haplotype
hap1
hap2
1.00
0.75
0.50
0.25
0.00
10.0
7.5
5.0
2.5
0.0

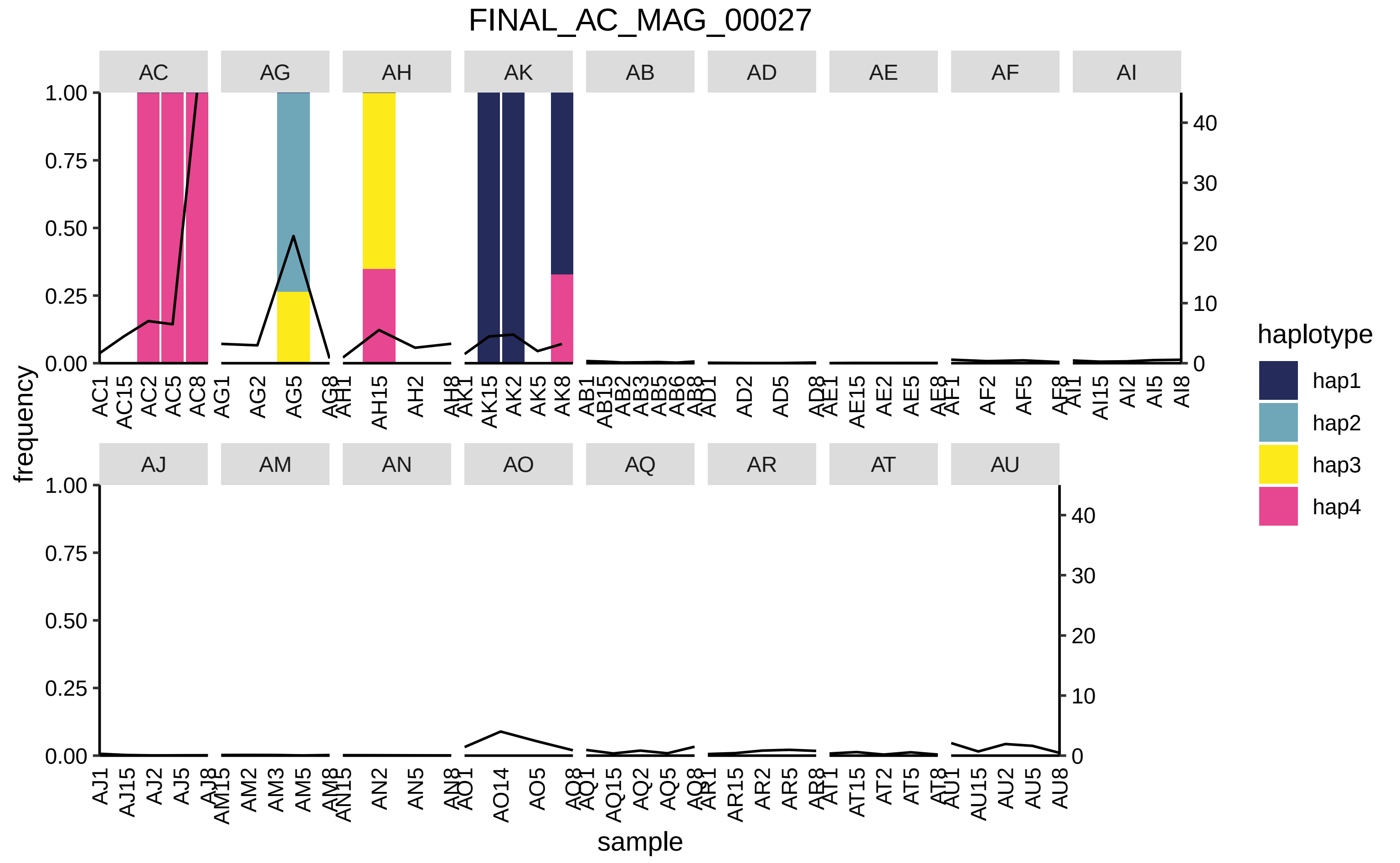

FINAL_AC_MAG_00027
AC
AG
AH
AK
AB
AD
AE
AF
AI
AJ
AM
AN
AO
AQ
AR
AT
AU
frequency
sample
haplotype
hap1
hap2
hap3
hap4

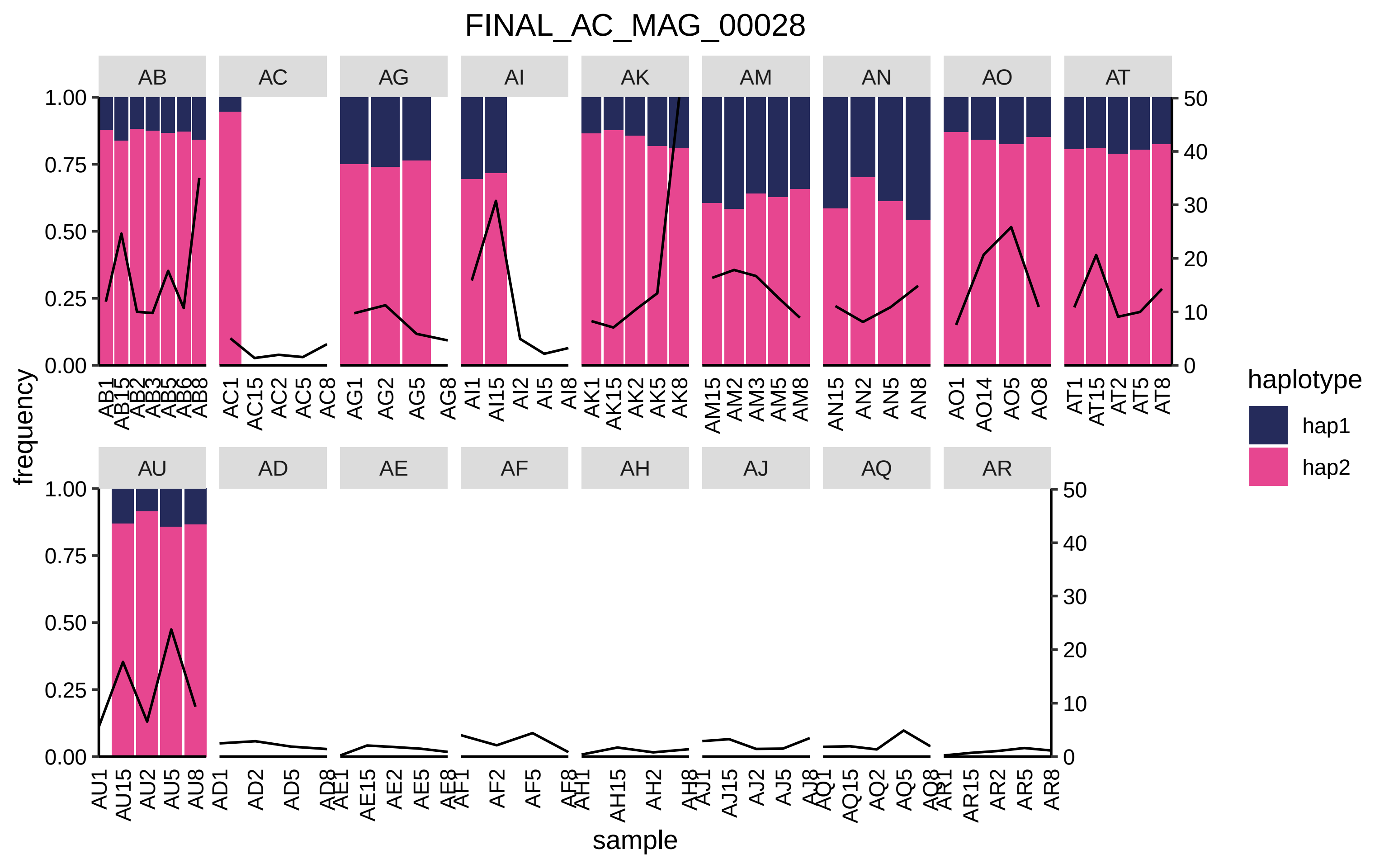
FINAL_AC_MAG_00028
AB
AC
AG
AI
AK
AM
AN
AO
AT
AU
AD
AE
AF
AH
AJ
AQ
AR
frequency
sample
haplotype
hap1
hap2

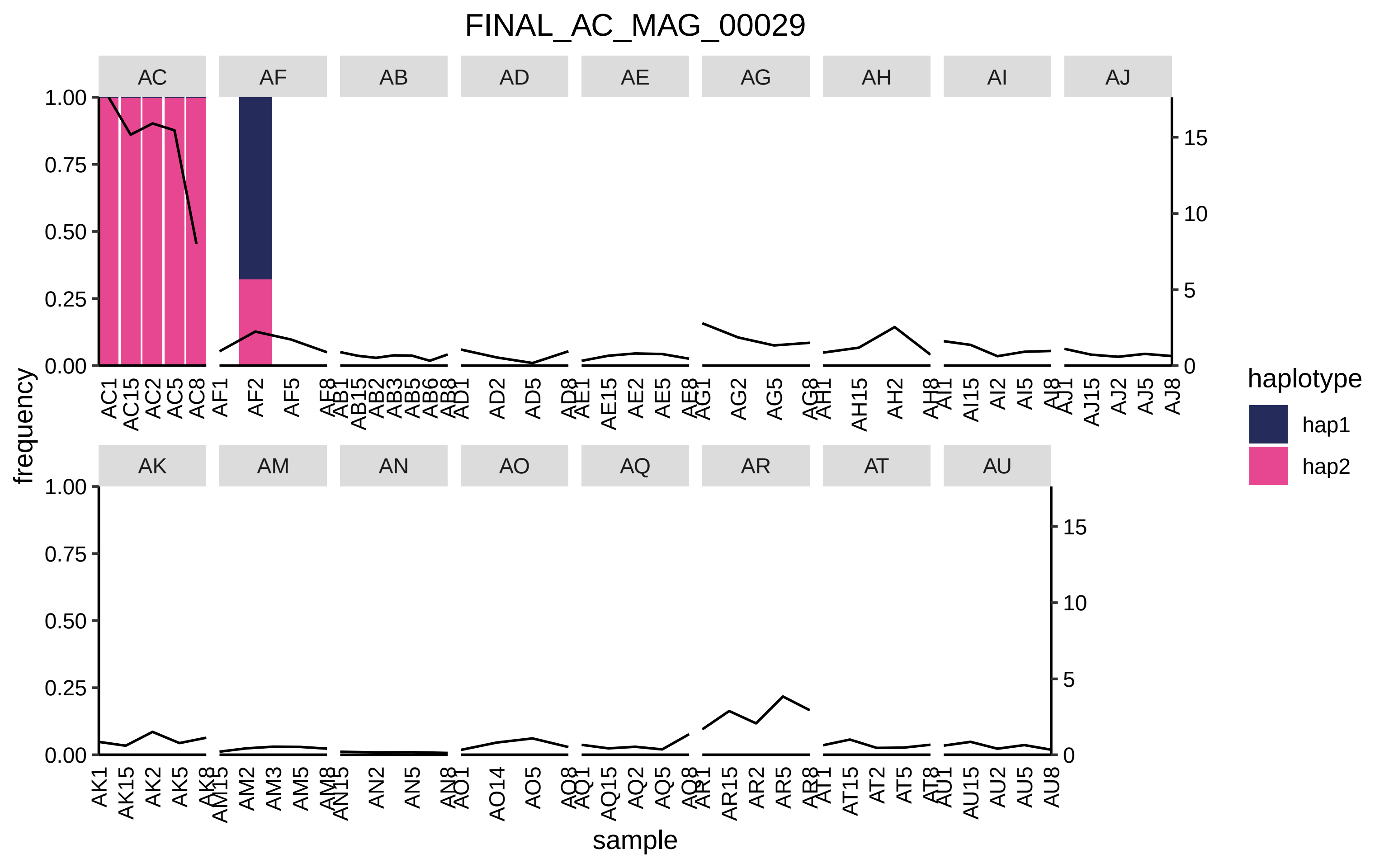

FINAL_AC_MAG_00029
AC
AF
AB
AD
AE
AG
AH
AI
AJ
AK
AM
AN
AO
AQ
AR
AT
AU
frequency
sample
haplotype
hap1
hap2

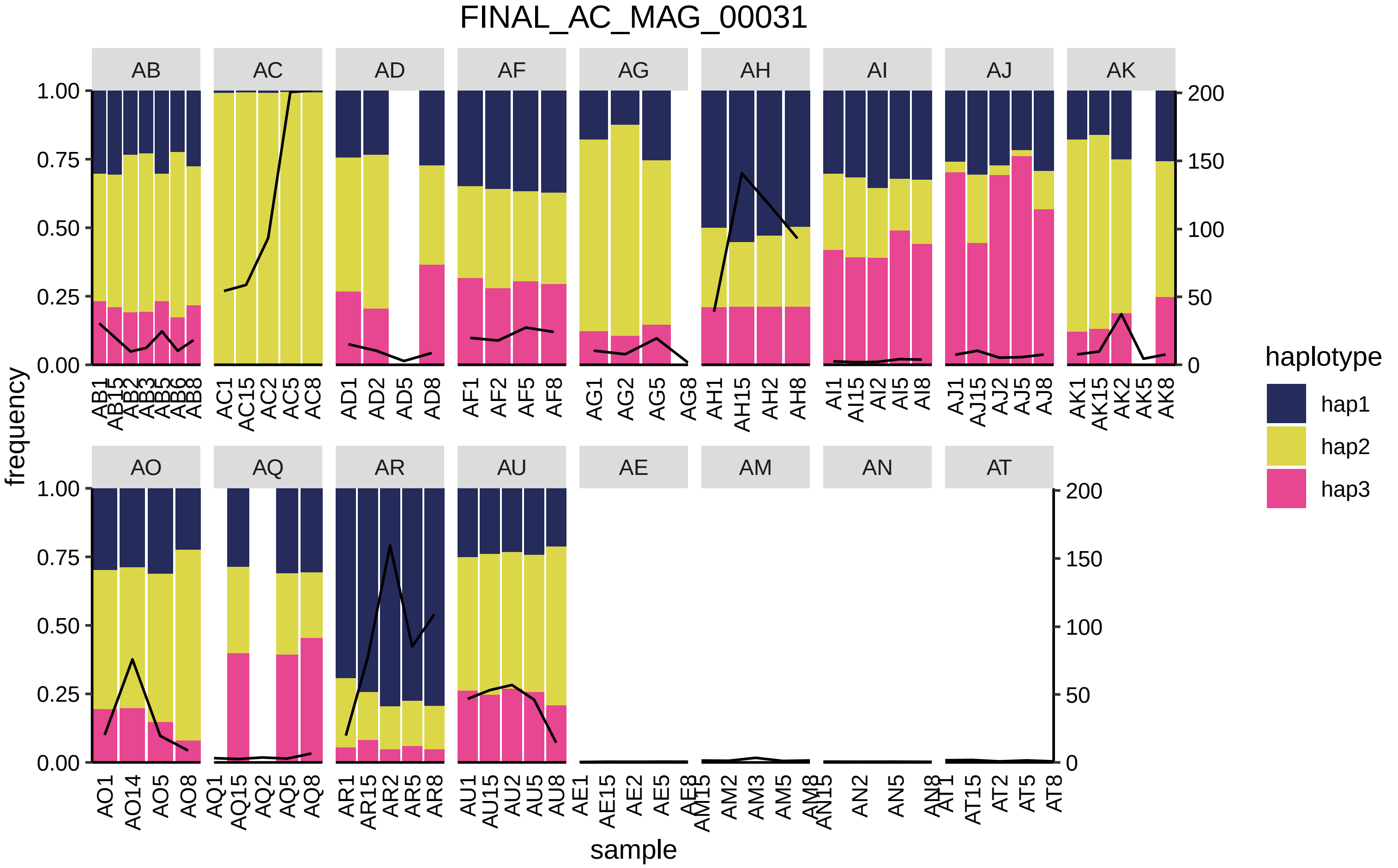

FINAL_AC_MAG_00031
AB
AC
AD
AF
AG
AH
AI
AJ
AK
AO
AQ
AR
AU
AE
AM
AN
AT
frequency
sample
haplotype
hap1
hap2
hap3

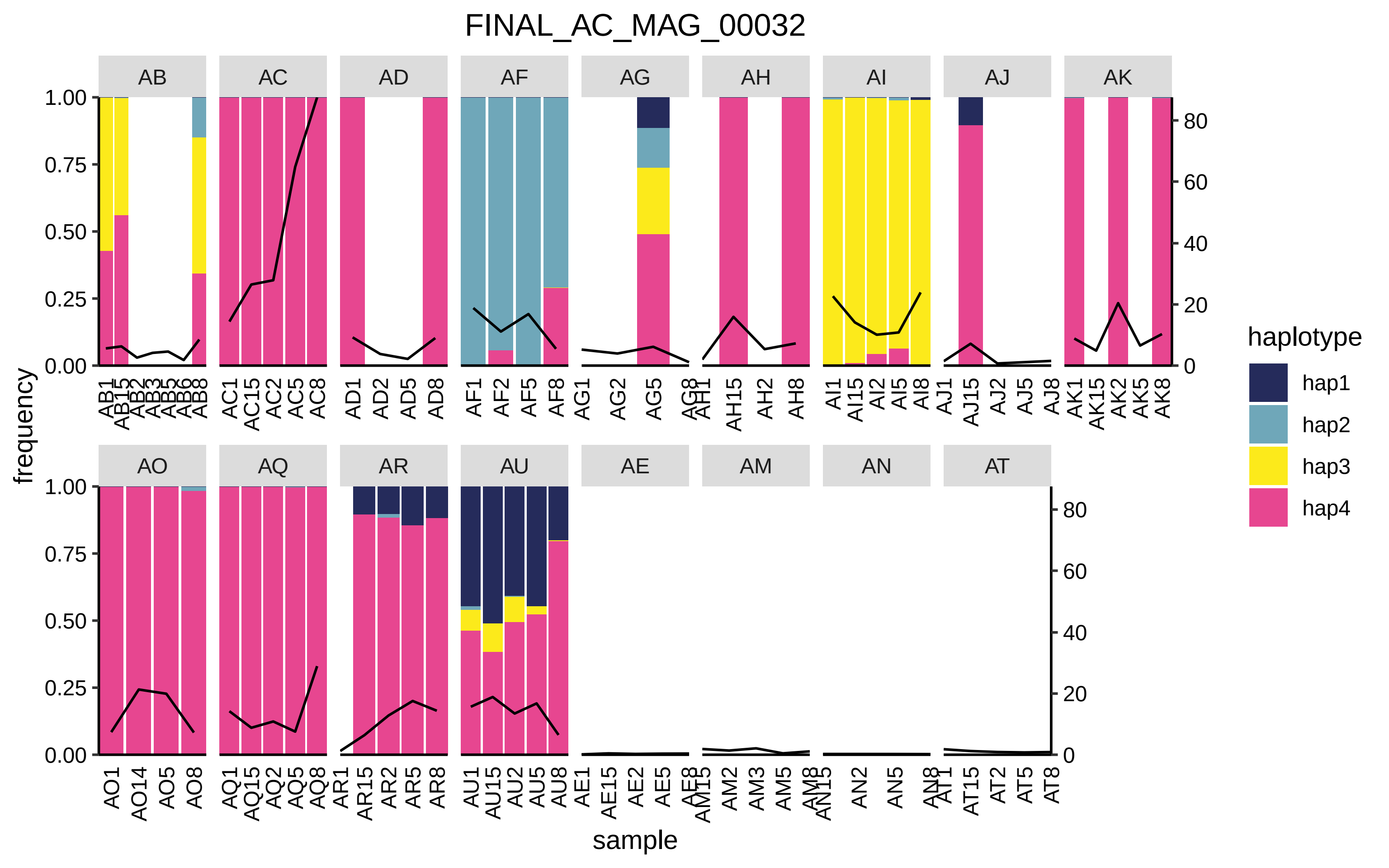

FINAL_AC_MAG_00032
AB
AC
AD
AF
AG
AH
AI
AJ
AK
AO
AQ
AR
AU
AE
AM
AN
AT
frequency
sample
haplotype
hap1
hap2
hap3
hap4

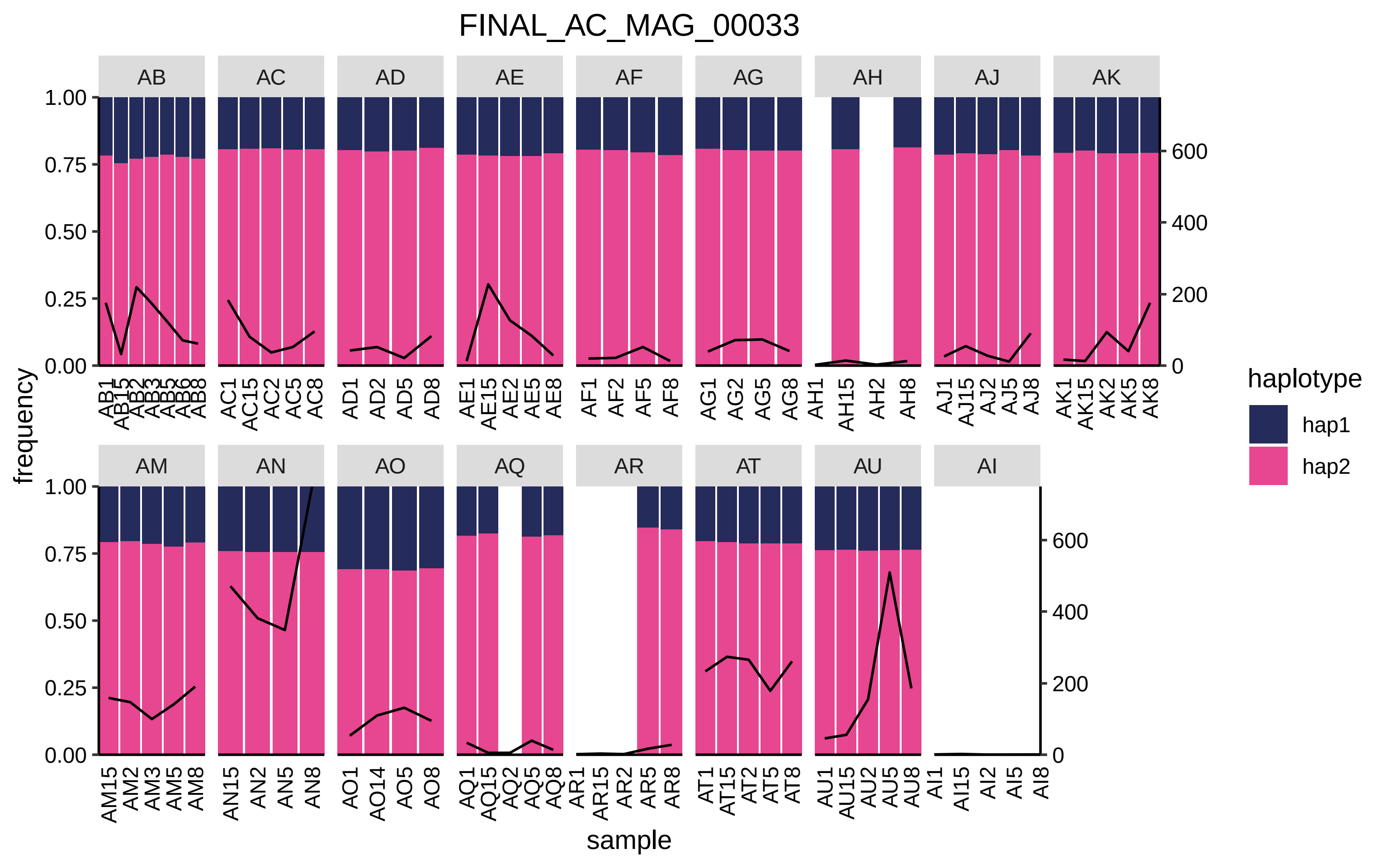
FINAL_AC_MAG_00033
AB
AC
AD
AE
AF
AG
AH
AJ
AK
AM
AN
AO
AQ
AR
AT
AU
AI
frequency
sample
1.00
0.75
0.50
0.25
0.00
600
400
200
0
AB1 AB15 AB2 AB3 AB5 AB6 AB8
AC1 AC15 AC2 AC5 AC8
AD1 AD2 AD5 AD8
AE1 AE15 AE2 AE5 AE8
AF1 AF2 AF5 AF8
AG1 AG2 AG5 AG8
AH1 AH15 AH2 AH8
AJ1 AJ15 AJ2 AJ5 AJ8
AK1 AK15 AK2 AK5 AK8
AM15 AM2 AM3 AM5 AM8
AN15 AN2 AN5 AN8
AO1 AO14 AO5 AO8
AQ1 AQ15 AQ2 AQ5 AQ8
AR1 AR15 AR2 AR5 AR8
AT1 AT15 AT2 AT5 AT8
AU1 AU15 AU2 AU5 AU8
AI1 AI15 AI2 AI5 AI8
haplotype
hap1
hap2

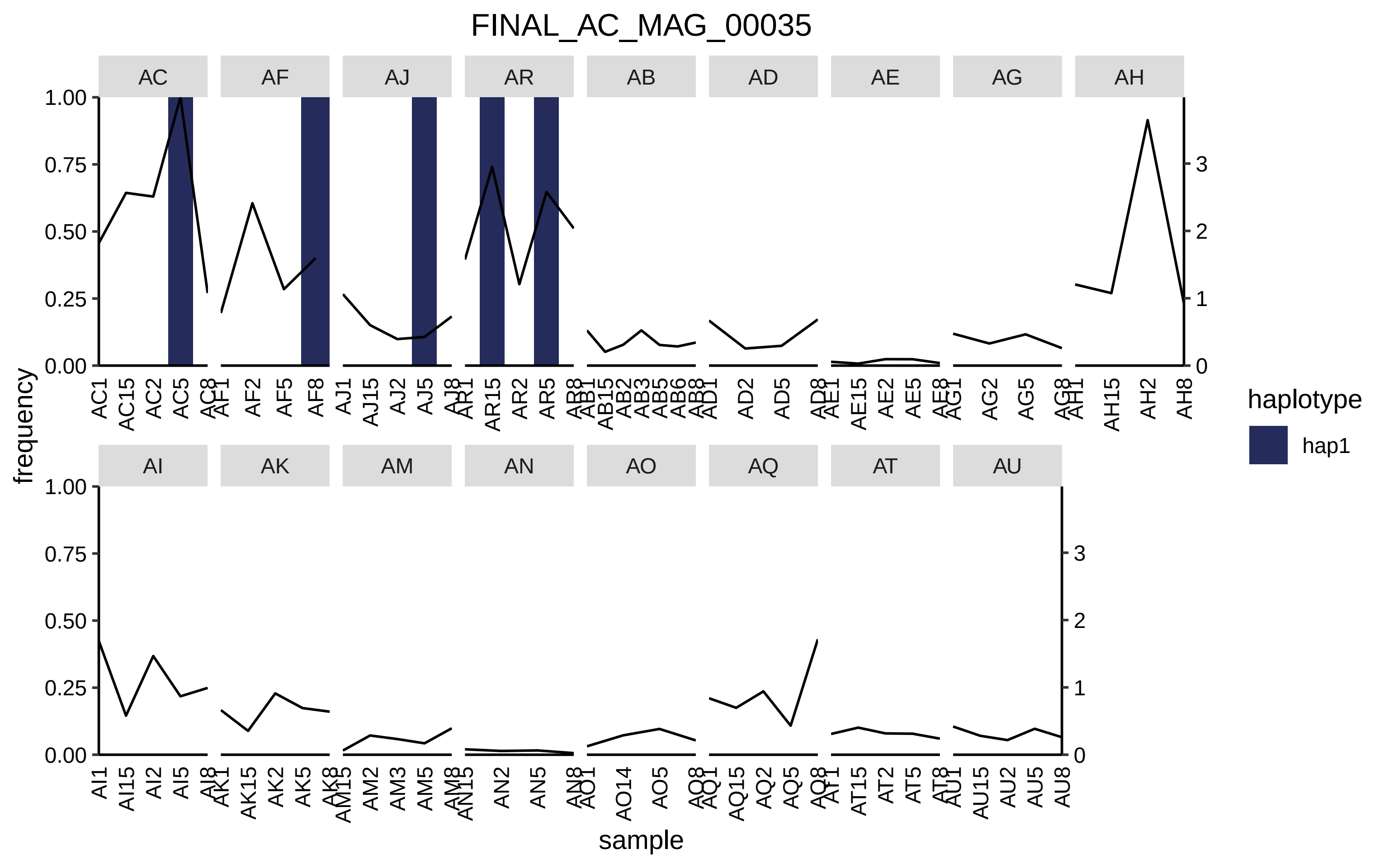
FINAL_AC_MAG_00035
AC
AF
AJ
AR
AB
AD
AE
AG
AH
AI
AK
AM
AN
AO
AQ
AT
AU
frequency
sample
haplotype
hap1
1.00
0.75
0.50
0.25
0.00
0
1
2
3

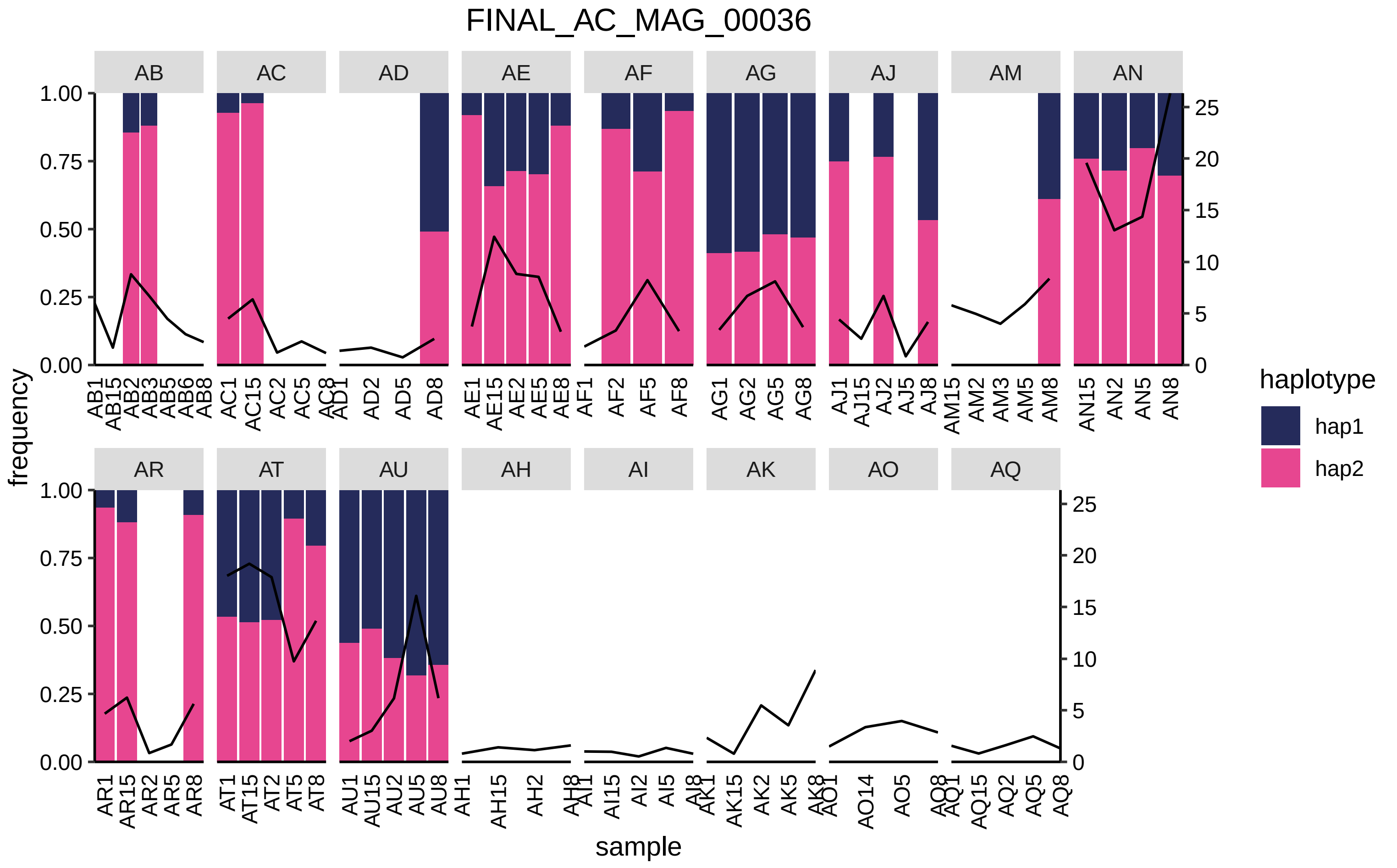

FINAL_AC_MAG_00036
AB
AC
AD
AE
AF
AG
AJ
AM
AN
AR
AT
AU
AH
AI
AK
AO
AQ
frequency
1.00
0.75
0.50
0.25
0.00
0
5
10
15
20
25
AB1 AB15 AB2 AB3 AB5 AB6 AB8
AC1 AC15 AC2 AC5 AC8
AD1 AD2 AD5 AD8
AE1 AE15 AE2 AE5 AE8
AF1 AF2 AF5 AF8
AG1 AG2 AG5 AG8
AJ1 AJ15 AJ2 AJ5 AJ8
AM15 AM2 AM3 AM5 AM8
AN15 AN2 AN5 AN8
AR1 AR15 AR2 AR5 AR8
AT1 AT15 AT2 AT5 AT8
AU1 AU15 AU2 AU5 AU8
AH1 AH15 AH2 AH8
AI1 AI15 AI2 AI5 AI8
AK1 AK15 AK2 AK5 AK8
AO1 AO14 AO5 AO8
AQ1 AQ15 AQ2 AQ5 AQ8
sample
haplotype
hap1
hap2

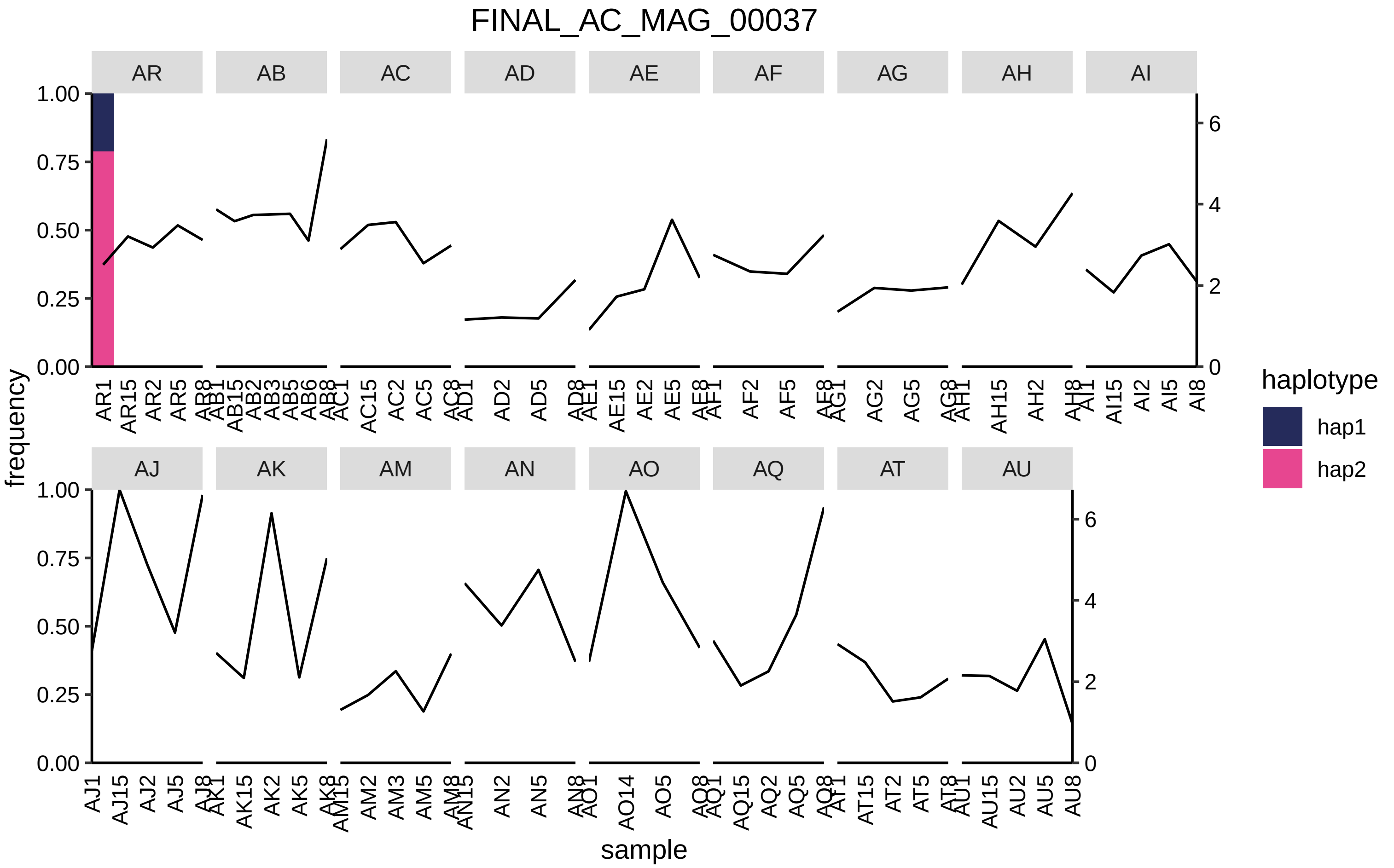
FINAL_AC_MAG_00037
AR
AB
AC
AD
AE
AF
AG
AH
AI
AJ
AK
AM
AN
AO
AQ
AT
AU
frequency
sample
haplotype
hap1
hap2
1.00
0.75
0.50
0.25
0.00
0
2
4
6

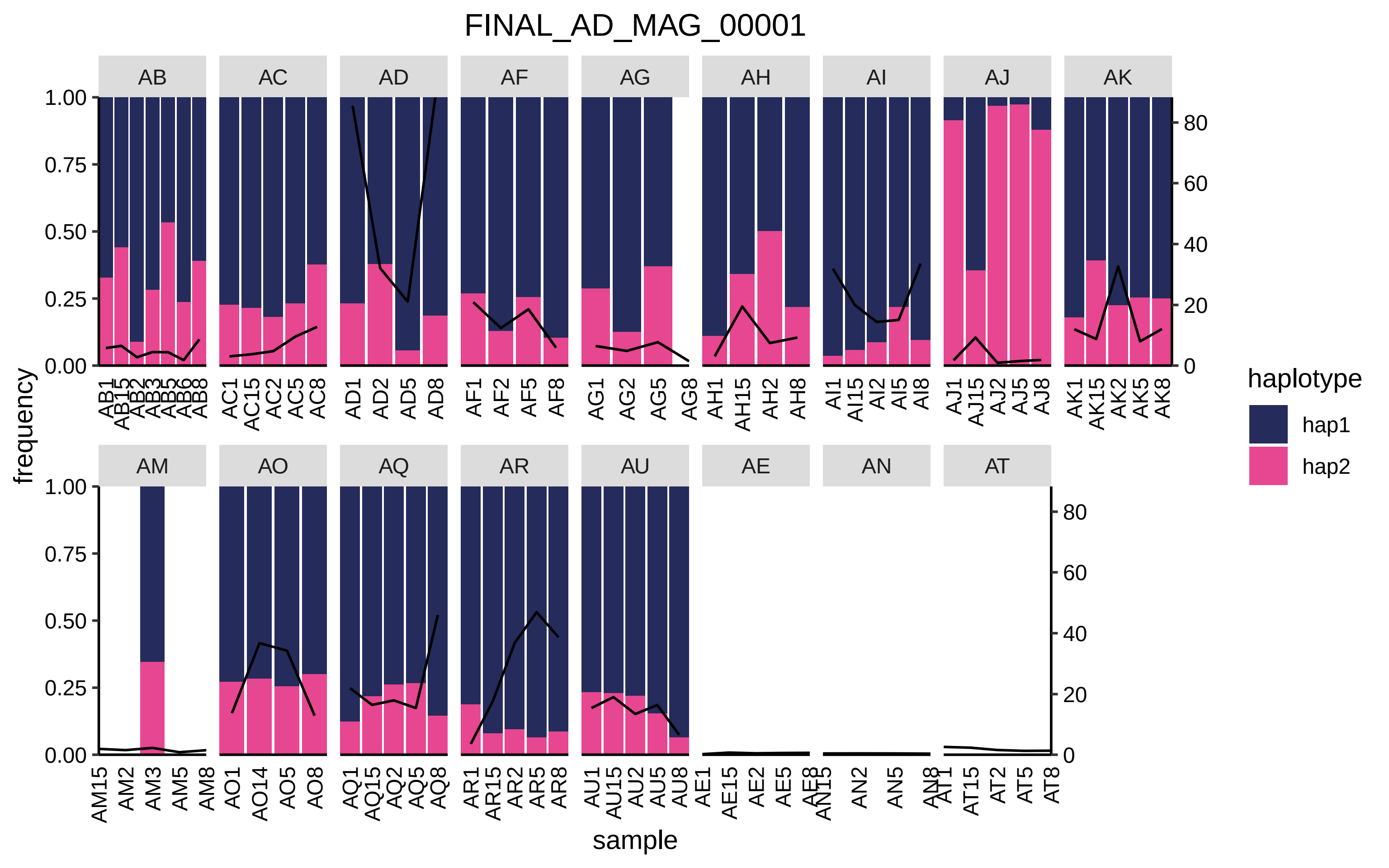

FINAL_AD_MAG_00001
AB
AC
AD
AF
AG
AH
AI
AJ
AK
AM
AO
AQ
AR
AU
AE
AN
AT
frequency
sample
haplotype
hap1
hap2
1.00
0.75
0.50
0.25
0.00
0
20
40
60
80
AB1 AB15 AB2 AB3 AB5 AB6 AB8
AC1 AC15 AC2 AC5 AC8
AD1 AD2 AD5 AD8
AF1 AF2 AF5 AF8
AG1 AG2 AG5 AG8
AH1 AH15 AH2 AH8
AI1 AI15 AI2 AI5 AI8
AJ1 AJ15 AJ2 AJ5 AJ8
AK1 AK15 AK2 AK5 AK8
AM15 AM2 AM3 AM5 AM8
AO1 AO14 AO5 AO8
AQ1 AQ15 AQ2 AQ5 AQ8
AR1 AR15 AR2 AR5 AR8
AU1 AU15 AU2 AU5 AU8
AE1 AE15 AE2 AE5 AE8
AN15 AN2 AN5 AN8
AT1 AT15 AT2 AT5 AT8

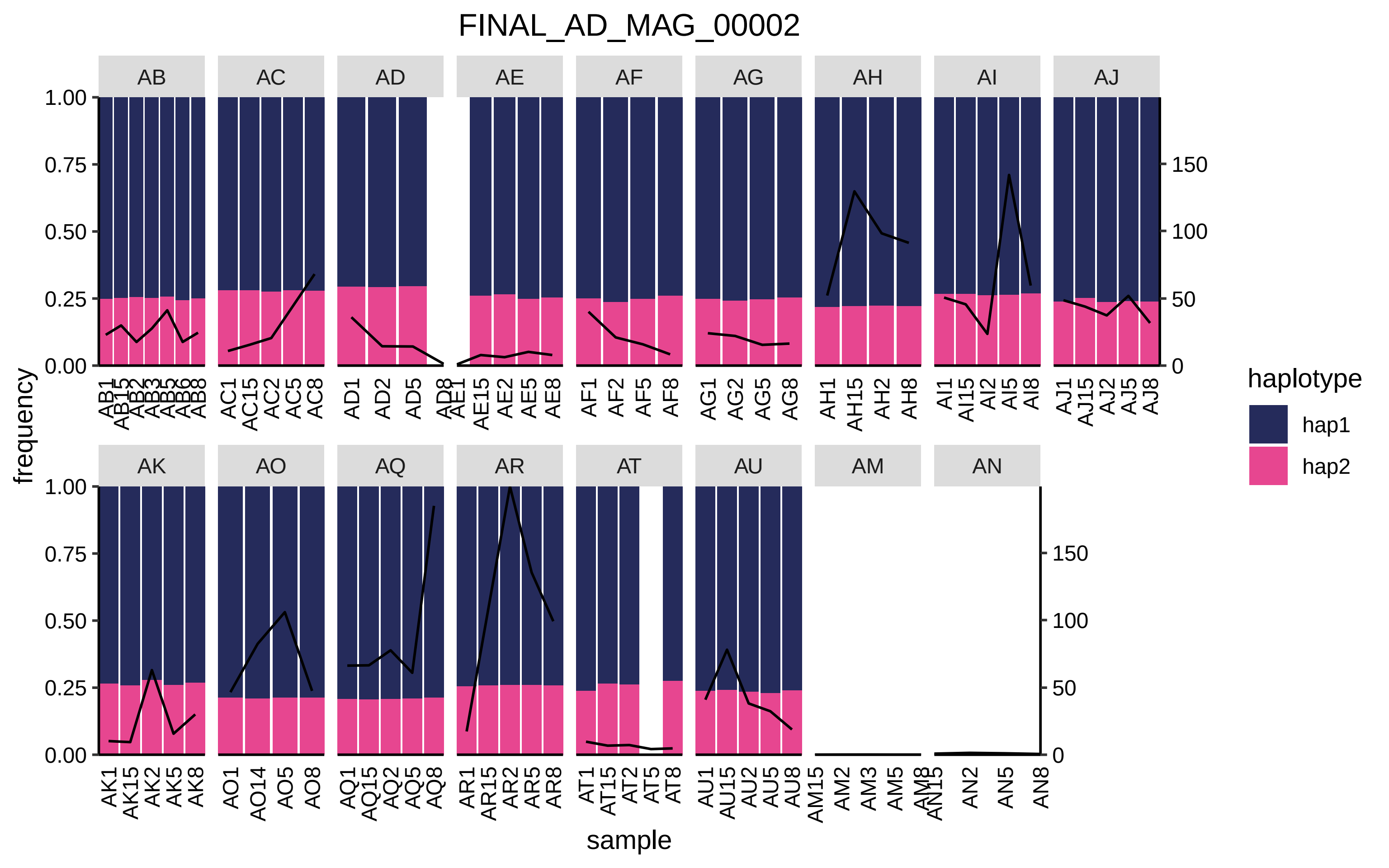

FINAL_AD_MAG_00002
AB
AC
AD
AE
AF
AG
AH
AI
AJ
AK
AO
AQ
AR
AT
AU
AM
AN
frequency
sample
haplotype
hap1
hap2

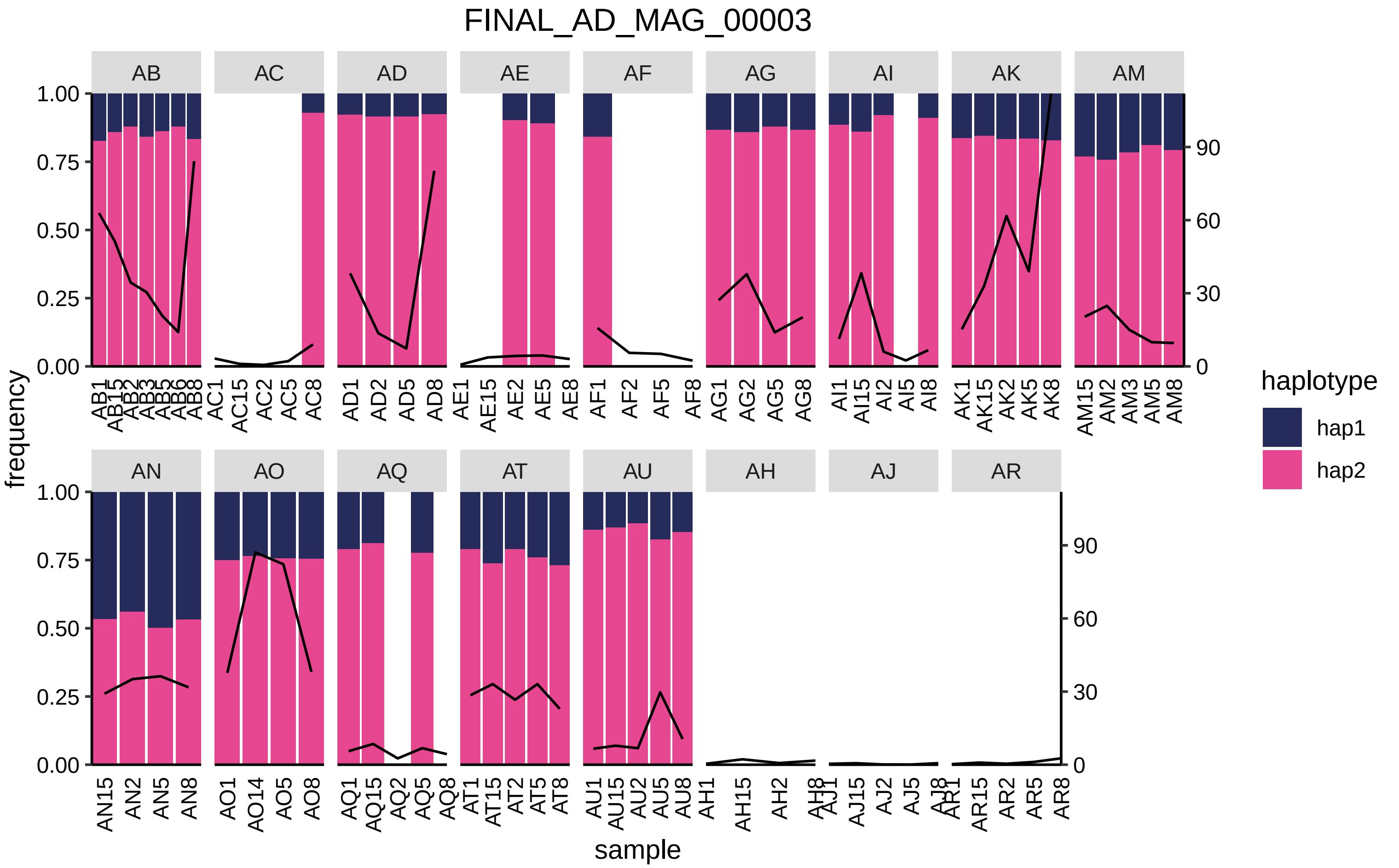
FINAL_AD_MAG_00003
AB
AC
AD
AE
AF
AG
AI
AK
AM
AN
AO
AQ
AT
AU
AH
AJ
AR
frequency
sample
haplotype
hap1
hap2
1.00
0.75
0.50
0.25
0.00
0
30
60
90
AB1 AB15 AB2 AB3 AB5 AB6 AB8
AC1 AC15 AC2 AC5 AC8
AD1 AD2 AD5 AD8
AE1 AE15 AE2 AE5 AE8
AF1 AF2 AF5 AF8
AG1 AG2 AG5 AG8
AI1 AI15 AI2 AI5 AI8
AK1 AK15 AK2 AK5 AK8
AM15 AM2 AM3 AM5 AM8
AN15 AN2 AN5 AN8
AO1 AO14 AO5 AO8
AQ1 AQ15 AQ2 AQ5 AQ8
AT1 AT15 AT2 AT5 AT8
AU1 AU15 AU2 AU5 AU8
AH1 AH15 AH2 AH8
AJ1 AJ15 AJ2 AJ5 AJ8
AR1 AR15 AR2 AR5 AR8

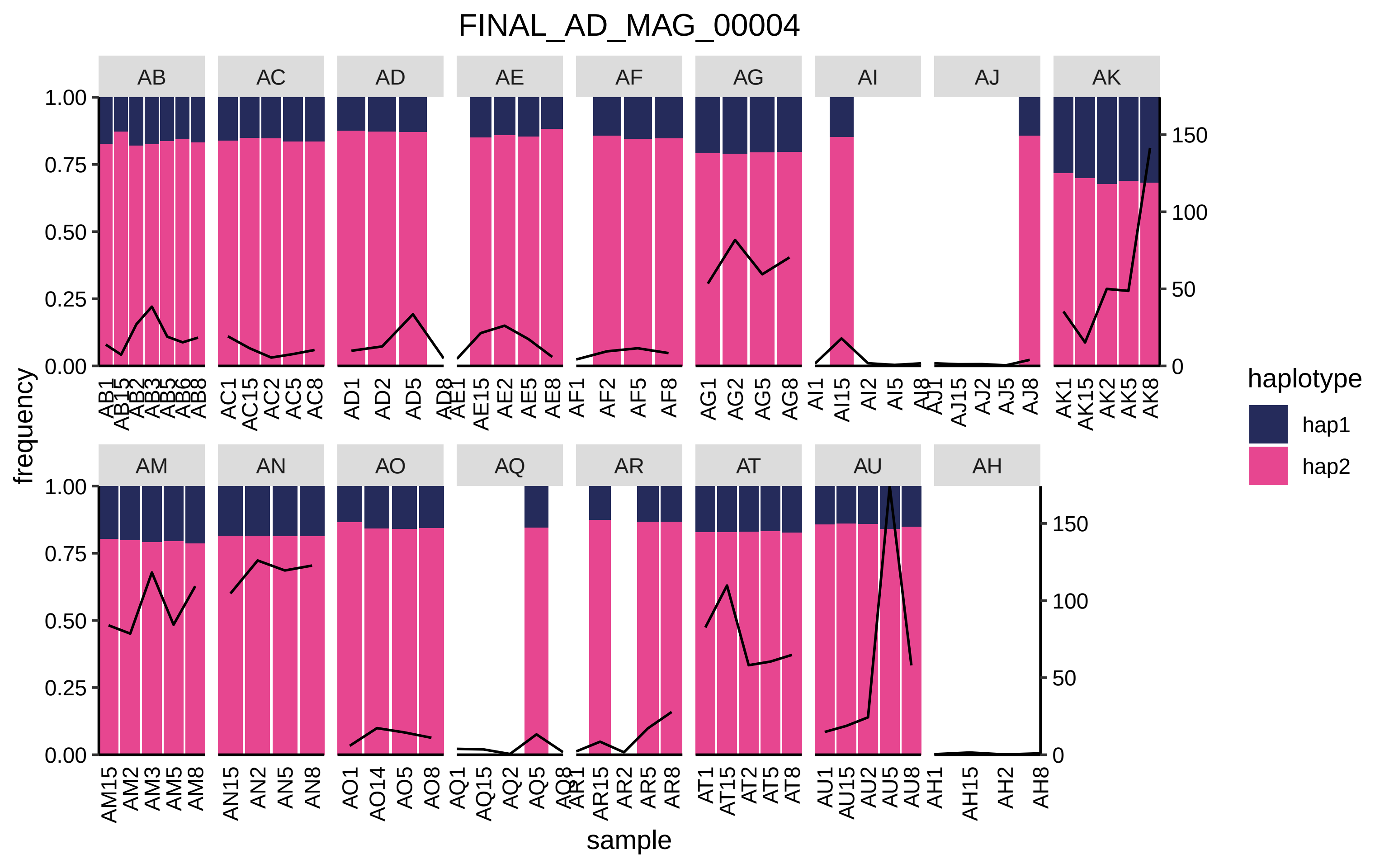

FINAL_AD_MAG_00004
AB
AC
AD
AE
AF
AG
AI
AJ
AK
AM
AN
AO
AQ
AR
AT
AU
AH
frequency
sample
haplotype
hap1
hap2
1.00
0.75
0.50
0.25
0.00
150
100
50
0
AB1 AB15 AB2 AB3 AB5 AB6 AB8
AC1 AC15 AC2 AC5 AC8
AD1 AD2 AD5 AD8
AE1 AE15 AE2 AE5 AE8
AF1 AF2 AF5 AF8
AG1 AG2 AG5 AG8
AI1 AI15 AI2 AI5 AI8
AJ1 AJ15 AJ2 AJ5 AJ8
AK1 AK15 AK2 AK5 AK8
AM15 AM2 AM3 AM5 AM8
AN15 AN2 AN5 AN8
AO1 AO14 AO5 AO8
AQ1 AQ15 AQ2 AQ5 AQ8
AR1 AR15 AR2 AR5 AR8
AT1 AT15 AT2 AT5 AT8
AU1 AU15 AU2 AU5 AU8
AH1 AH15 AH2 AH8

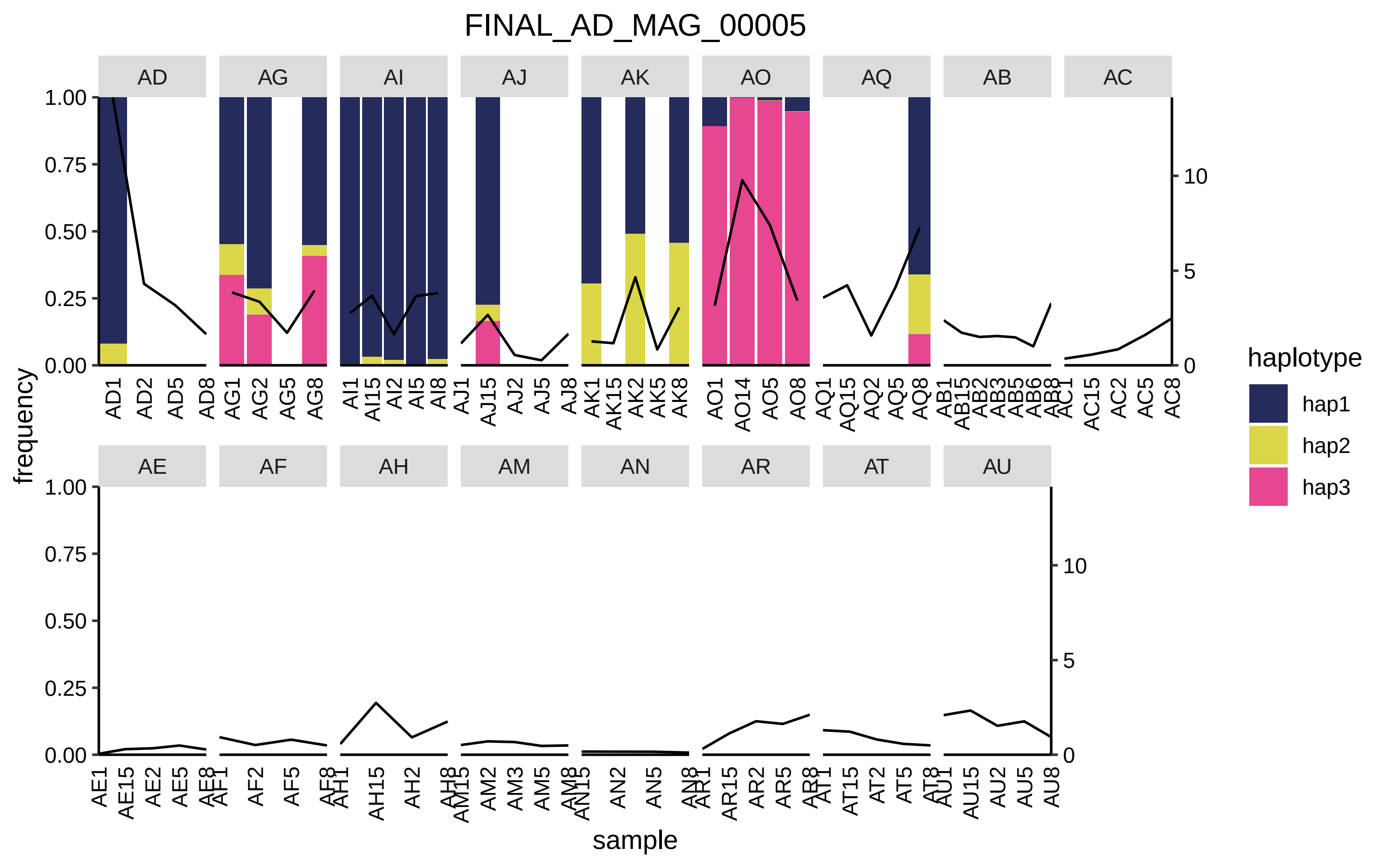

FINAL_AD_MAG_00005
AD
AG
AI
AJ
AK
AO
AQ
AB
AC
AE
AF
AH
AM
AN
AR
AT
AU
frequency
1.00
0.75
0.50
0.25
0.00
0
5
10
AD1 AD2 AD5 AD8
AG1 AG2 AG5 AG8
AI1 AI15 AI2 AI5 AI8
AJ1 AJ15 AJ2 AJ5 AJ8
AK1 AK15 AK2 AK5 AK8
AO1 AO14 AO5 AO8
AQ1 AQ15 AQ2 AQ5 AQ8
AB1 AB15 AB2 AB3 AB5 AB6 AB8
AC1 AC15 AC2 AC5 AC8
AE1 AE15 AE2 AE5 AE8
AF1 AF2 AF5 AF8
AH1 AH15 AH2 AH8
AM15 AM2 AM3 AM5 AM8
AN15 AN2 AN5 AN8
AR1 AR15 AR2 AR5 AR8
AT1 AT15 AT2 AT5 AT8
AU1 AU15 AU2 AU5 AU8
sample
haplotype
hap1
hap2
hap3

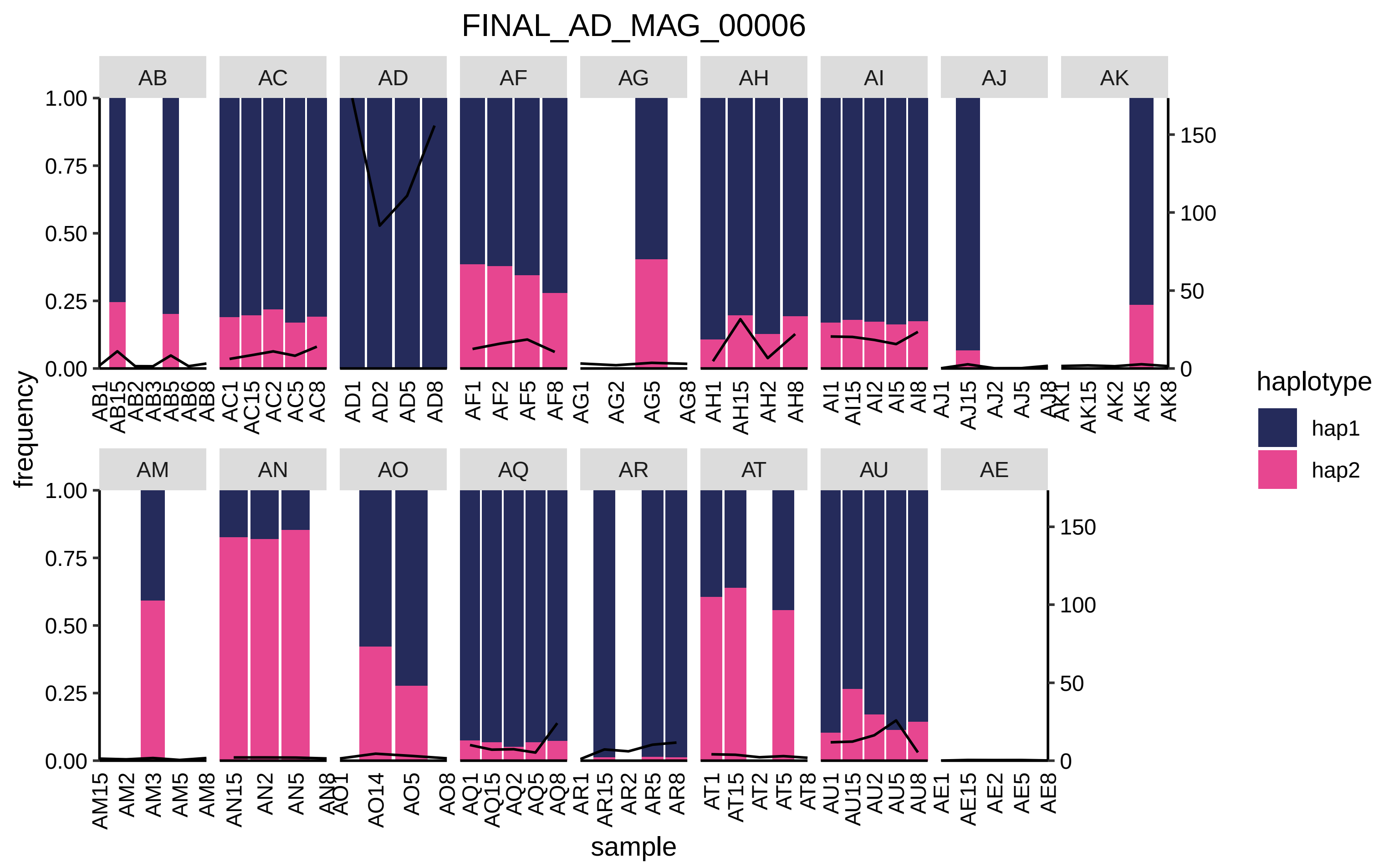

FINAL_AD_MAG_00006
AB
AC
AD
AF
AG
AH
AI
AJ
AK
AM
AN
AO
AQ
AR
AT
AU
AE
frequency
sample
haplotype
hap1
hap2

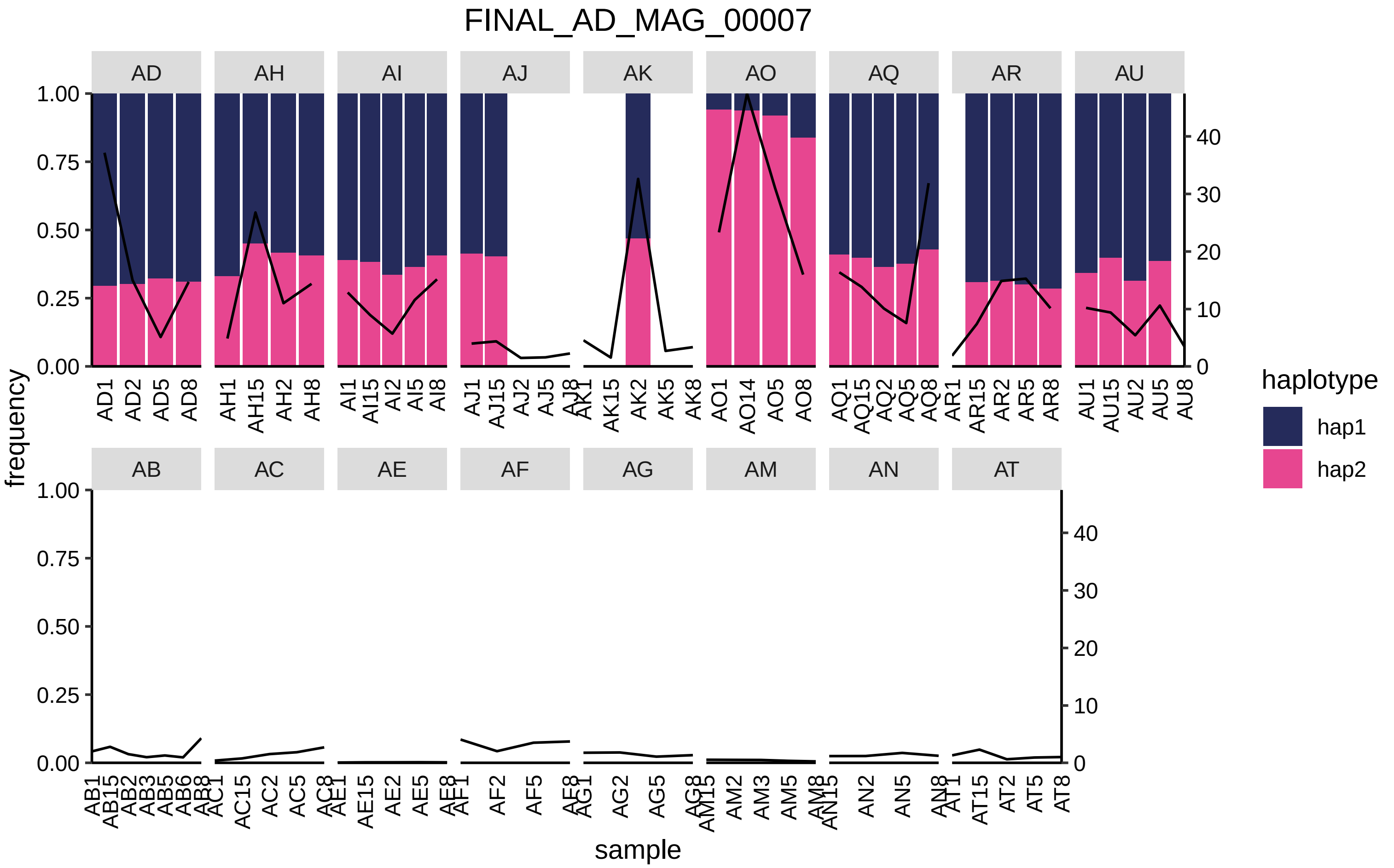
FINAL_AD_MAG_00007
AD
AH
AI
AJ
AK
AO
AQ
AR
AU
AB
AC
AE
AF
AG
AM
AN
AT
1.00
0.75
0.50
0.25
0.00
40
30
20
10
0
frequency
sample
haplotype
hap1
hap2
AD1
AD2
AD5
AD8
AH1
AH15
AH2
AH8
AI1
AI15
AI2
AI5
AI8
AJ1
AJ15
AJ2
AJ5
AJ8
AK1
AK15
AK2
AK5
AK8
AO1
AO14
AO5
AO8
AQ1
AQ15
AQ2
AQ5
AQ8
AR1
AR15
AR2
AR5
AR8
AU1
AU15
AU2
AU5
AU8
AB1
AB15
AB2
AB3
AB5
AB6
AB8
AC1
AC15
AC2
AC5
AC8
AE1
AE15
AE2
AE5
AE8
AF1
AF2
AF5
AF8
AG1
AG2
AG5
AG8
AM15
AM2
AM3
AM5
AM8
AN15
AN2
AN5
AN8
AT1
AT15
AT2
AT5
AT8

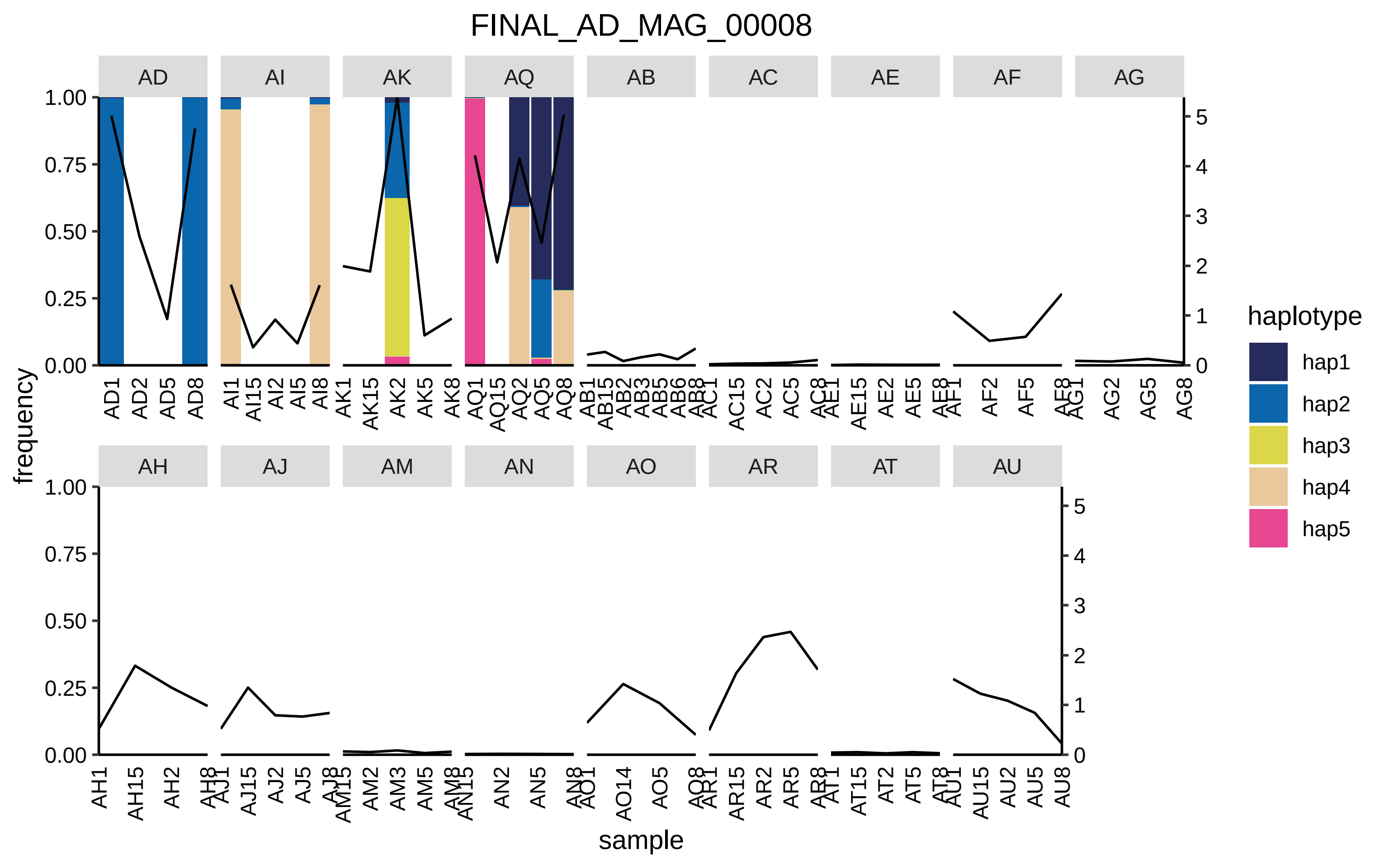

FINAL_AD_MAG_00008
AD
AI
AK
AQ
AB
AC
AE
AF
AG
AH
AJ
AM
AN
AO
AR
AT
AU
frequency
sample
haplotype
hap1
hap2
hap3
hap4
hap5

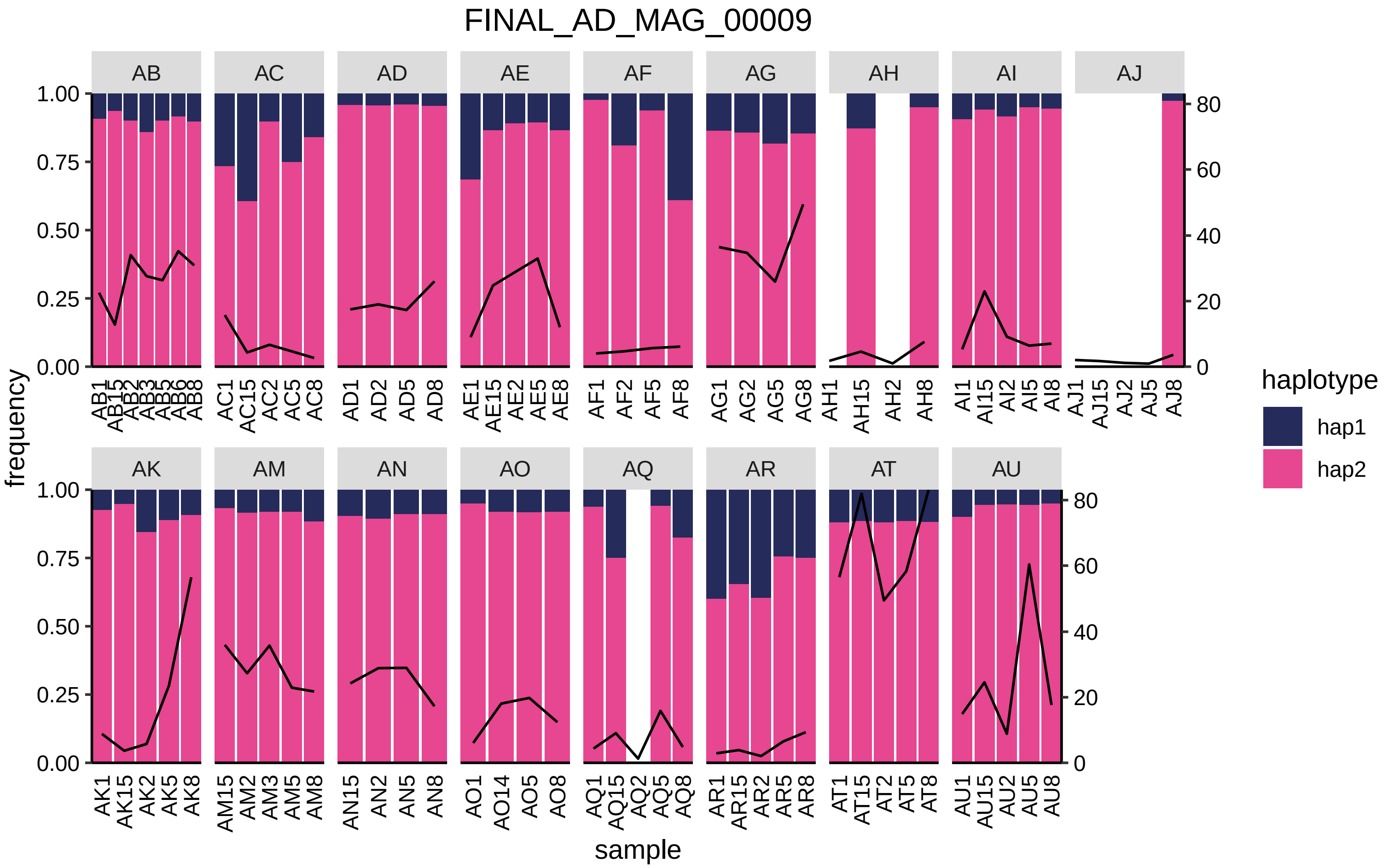

FINAL_AD_MAG_00009
AB
AC
AD
AE
AF
AG
AH
AI
AJ
AK
AM
AN
AO
AQ
AR
AT
AU
frequency
sample
haplotype
hap1
hap2
AB1 AB15 AB2 AB3 AB5 AB6 AB8
AC1 AC15 AC2 AC5 AC8
AD1 AD2 AD5 AD8
AE1 AE15 AE2 AE5 AE8
AF1 AF2 AF5 AF8
AG1 AG2 AG5 AG8
AH1 AH15 AH2 AH8
AI1 AI15 AI2 AI5 AI8
AJ1 AJ15 AJ2 AJ5 AJ8
AK1 AK15 AK2 AK5 AK8
AM15 AM2 AM3 AM5 AM8
AN15 AN2 AN5 AN8
AO1 AO14 AO5 AO8
AQ1 AQ15 AQ2 AQ5 AQ8
AR1 AR15 AR2 AR5 AR8
AT1 AT15 AT2 AT5 AT8
AU1 AU15 AU2 AU5 AU8
0.00 0.25 0.50 0.75 1.00
0 20 40 60 80

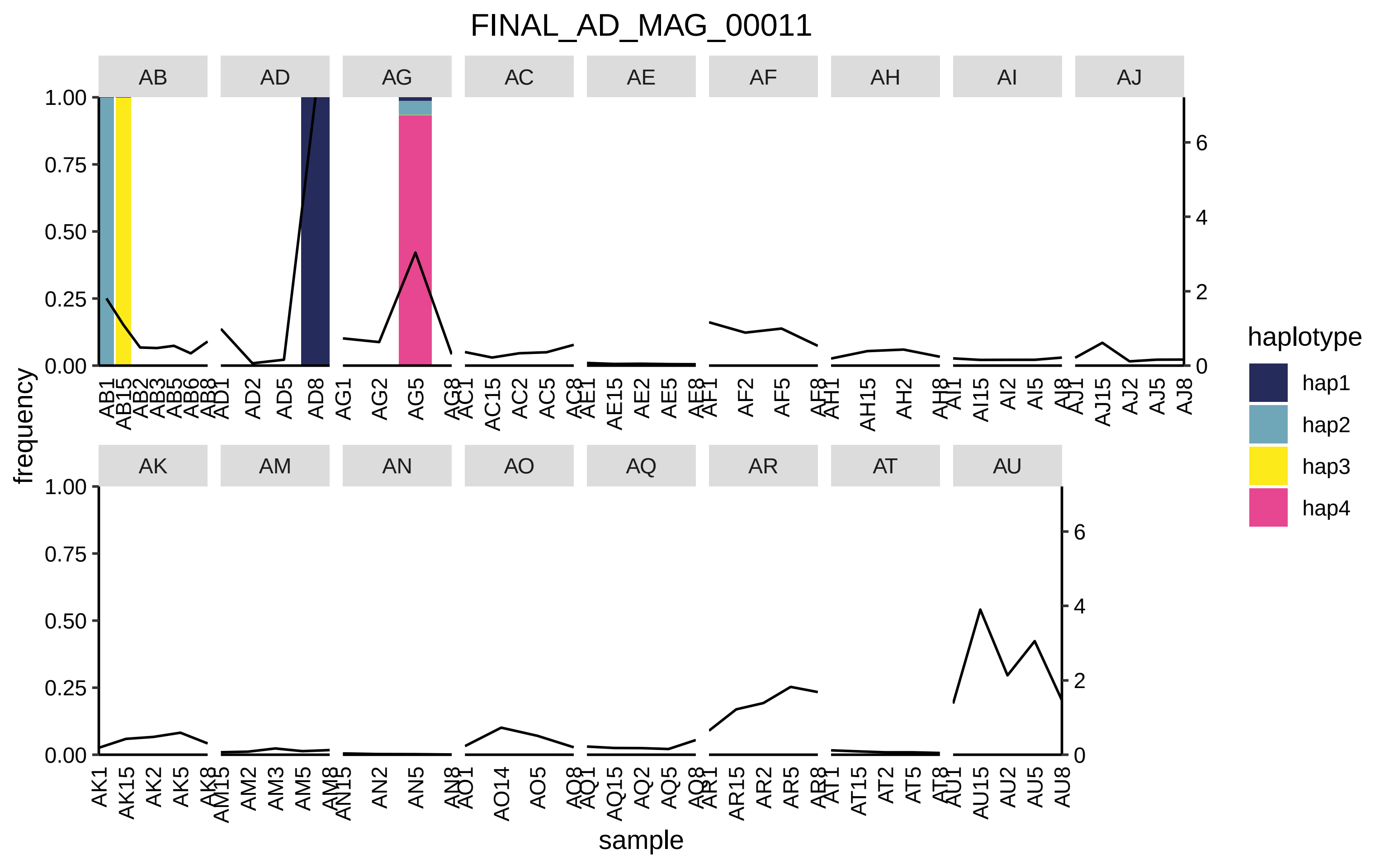

FINAL_AD_MAG_00011
AB
AD
AG
AC
AE
AF
AH
AI
AJ
AK
AM
AN
AO
AQ
AR
AT
AU
frequency
1.00
0.75
0.50
0.25
0.00
0
2
4
6
sample
haplotype
hap1
hap2
hap3
hap4

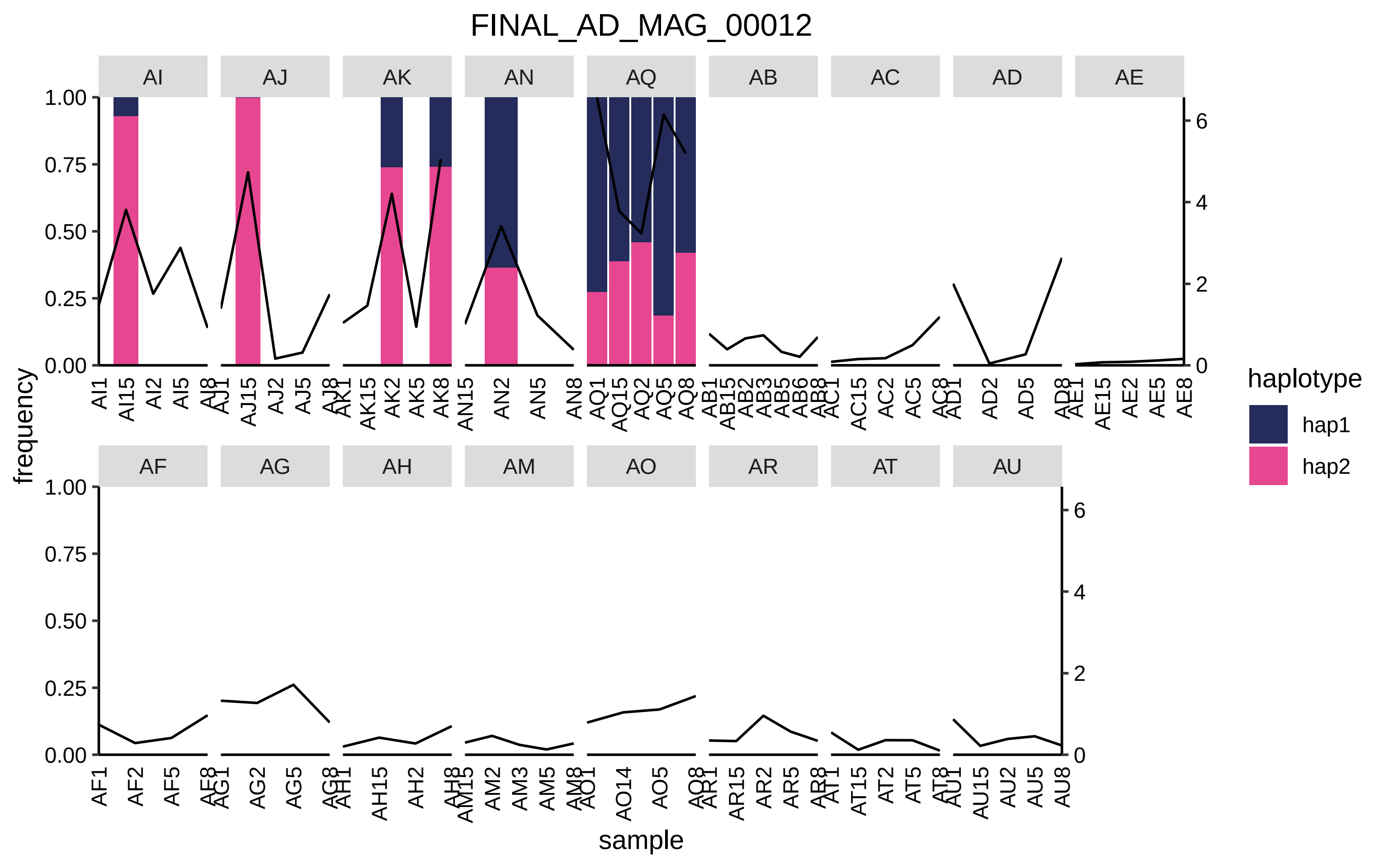

FINAL_AD_MAG_00012
AI
AJ
AK
AN
AQ
AB
AC
AD
AE
AF
AG
AH
AM
AO
AR
AT
AU
frequency
sample
haplotype
hap1
hap2

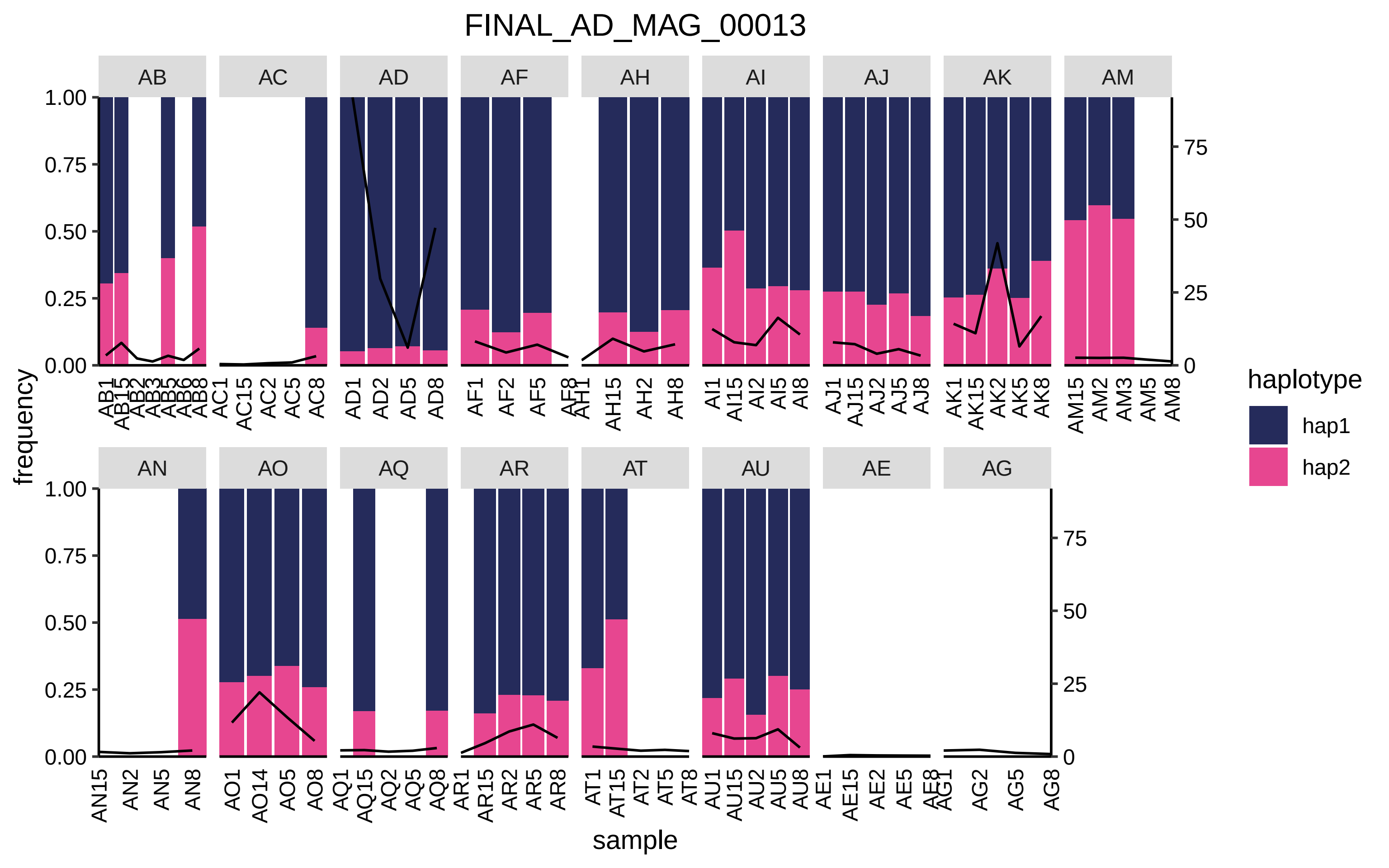
FINAL_AD_MAG_00013
AB
AC
AD
AF
AH
AI
AJ
AK
AM
AN
AO
AQ
AR
AT
AU
AE
AG
1.00
0.75
0.50
0.25
0.00
75
50
25
0
frequency
sample
AB1 AB15 AB2 AB3 AB5 AB6 AB8
AC1 AC15 AC2 AC5 AC8
AD1 AD2 AD5 AD8
AF1 AF2 AF5 AF8
AH1 AH15 AH2 AH8
AI1 AI15 AI2 AI5 AI8
AJ1 AJ15 AJ2 AJ5 AJ8
AK1 AK15 AK2 AK5 AK8
AM15 AM2 AM3 AM5 AM8
AN15 AN2 AN5 AN8
AO1 AO14 AO5 AO8
AQ1 AQ15 AQ2 AQ5 AQ8
AR1 AR15 AR2 AR5 AR8
AT1 AT15 AT2 AT5 AT8
AU1 AU15 AU2 AU5 AU8
AE1 AE15 AE2 AE5 AE8
AG1 AG2 AG5 AG8
haplotype
hap1
hap2

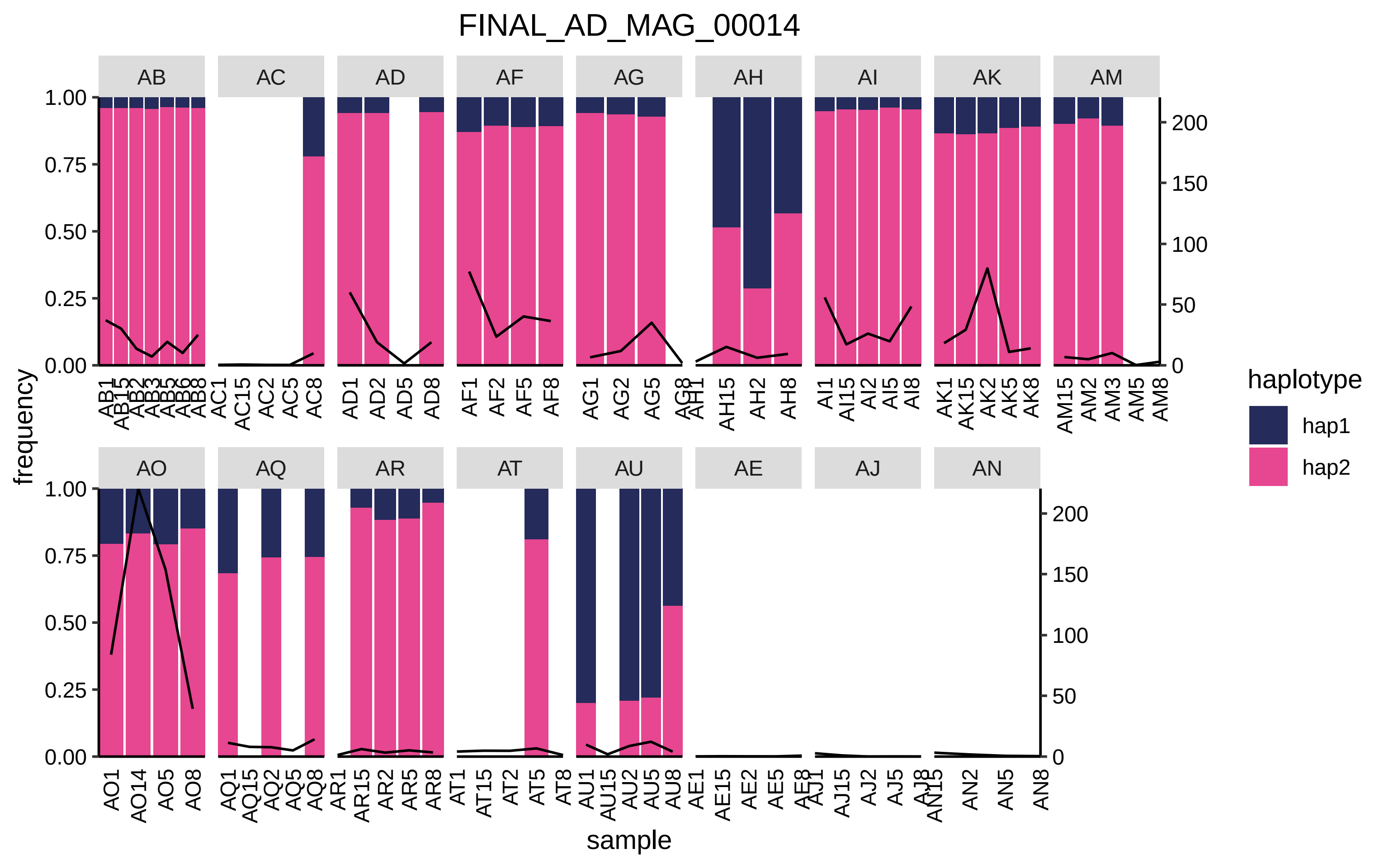

FINAL_AD_MAG_00014
AB
AC
AD
AF
AG
AH
AI
AK
AM
AO
AQ
AR
AT
AU
AE
AJ
AN
frequency
sample
haplotype
hap1
hap2

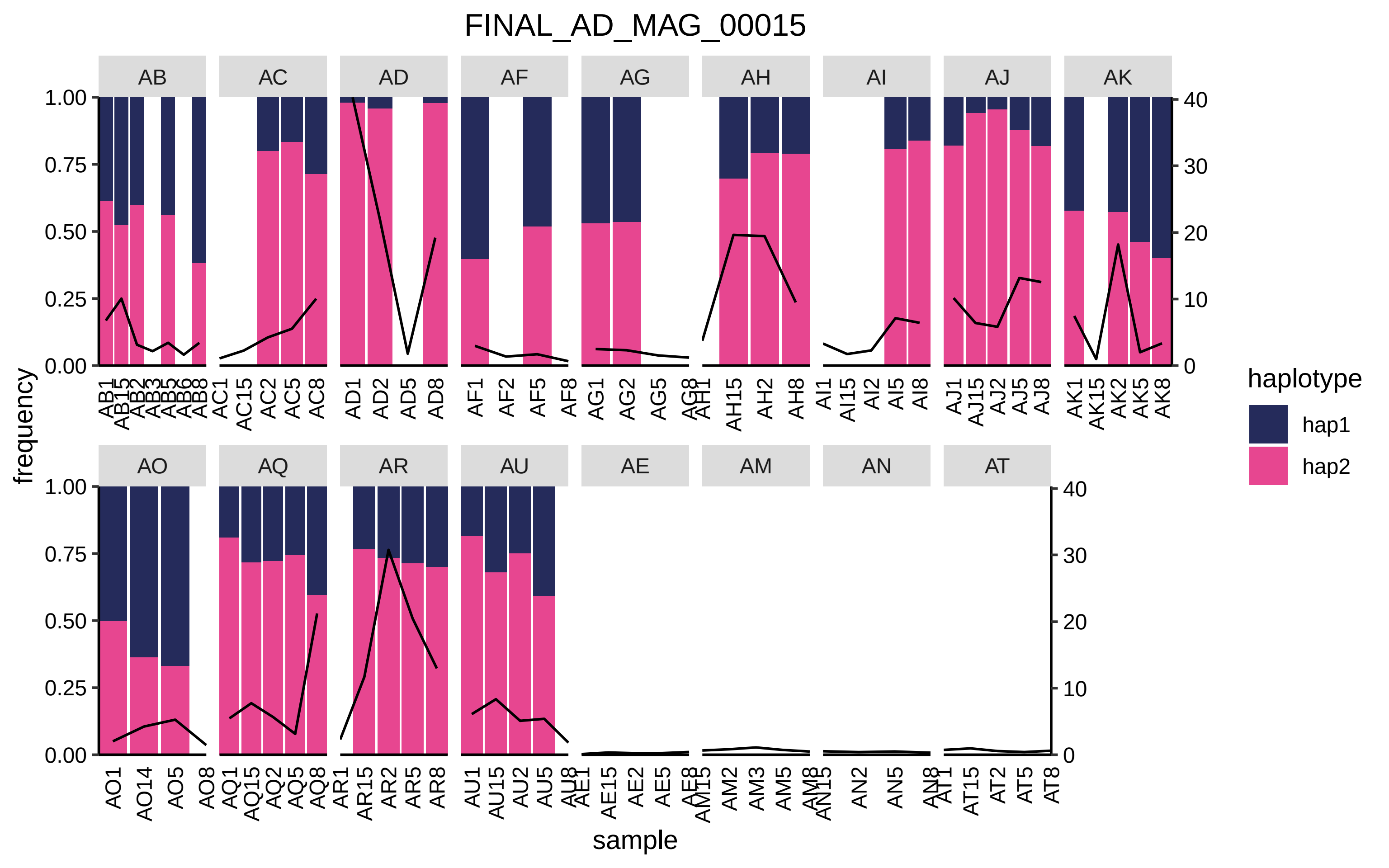

FINAL_AD_MAG_00015
AB
AC
AD
AF
AG
AH
AI
AJ
AK
AO
AQ
AR
AU
AE
AM
AN
AT
frequency
sample
haplotype
hap1
hap2

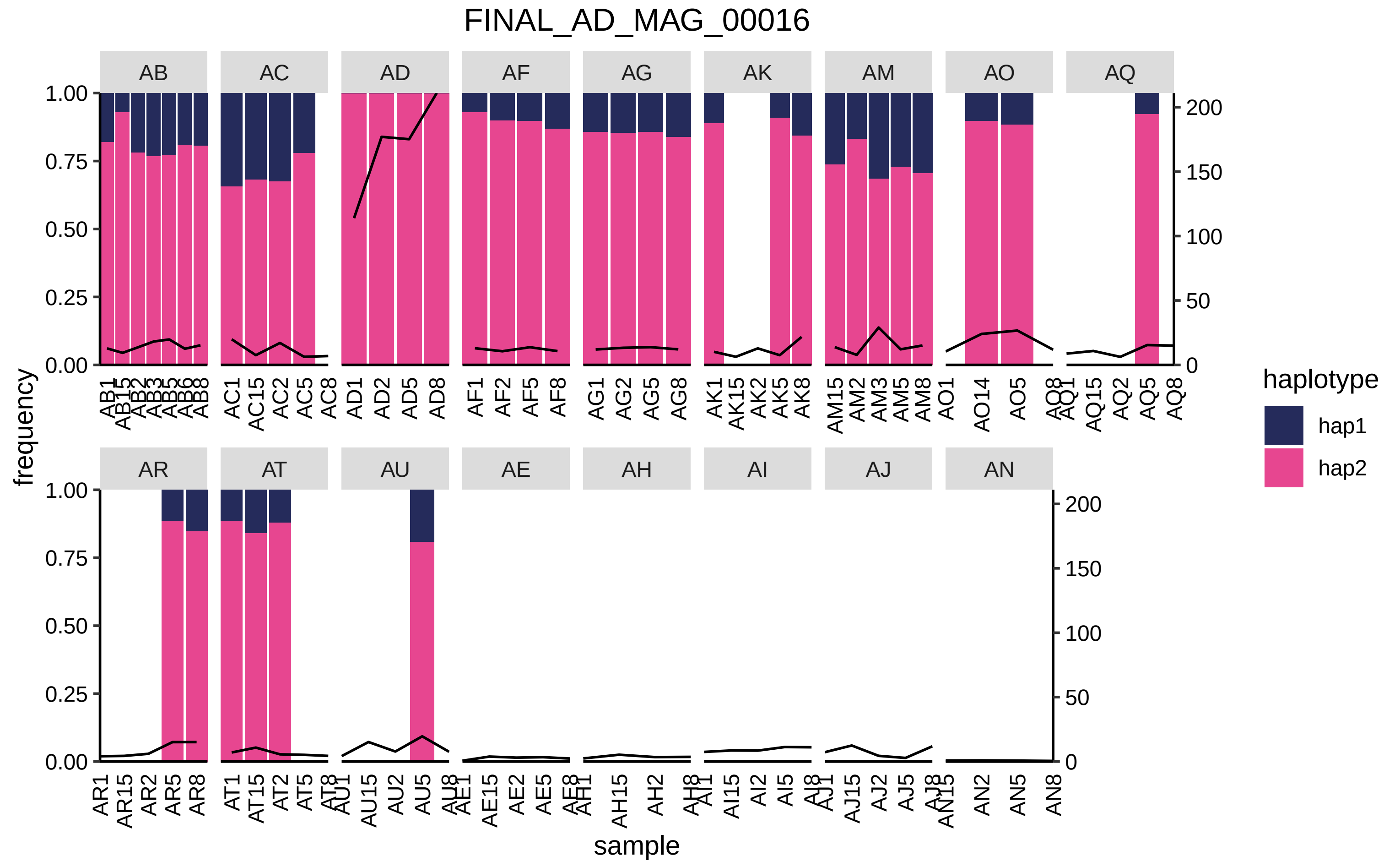

FINAL_AD_MAG_00016
AB
AC
AD
AF
AG
AK
AM
AO
AQ
AR
AT
AU
AE
AH
AI
AJ
AN
frequency
sample
haplotype
hap1
hap2
1.00
0.75
0.50
0.25
0.00
200
150
100
50
0
AB1 AB15 AB2 AB3 AB5 AB6 AB8
AC1 AC15 AC2 AC5 AC8
AD1 AD2 AD5 AD8
AF1 AF2 AF5 AF8
AG1 AG2 AG5 AG8
AK1 AK15 AK2 AK5 AK8
AM15 AM2 AM3 AM5 AM8
AO1 AO14 AO5 AO8
AQ1 AQ15 AQ2 AQ5 AQ8
AR1 AR15 AR2 AR5 AR8
AT1 AT15 AT2 AT5 AT8
AU1 AU15 AU2 AU5 AU8
AE1 AE15 AE2 AE5 AE8
AH1 AH15 AH2 AH8
AI1 AI15 AI2 AI5 AI8
AJ1 AJ15 AJ2 AJ5 AJ8
AN15 AN2 AN5 AN8

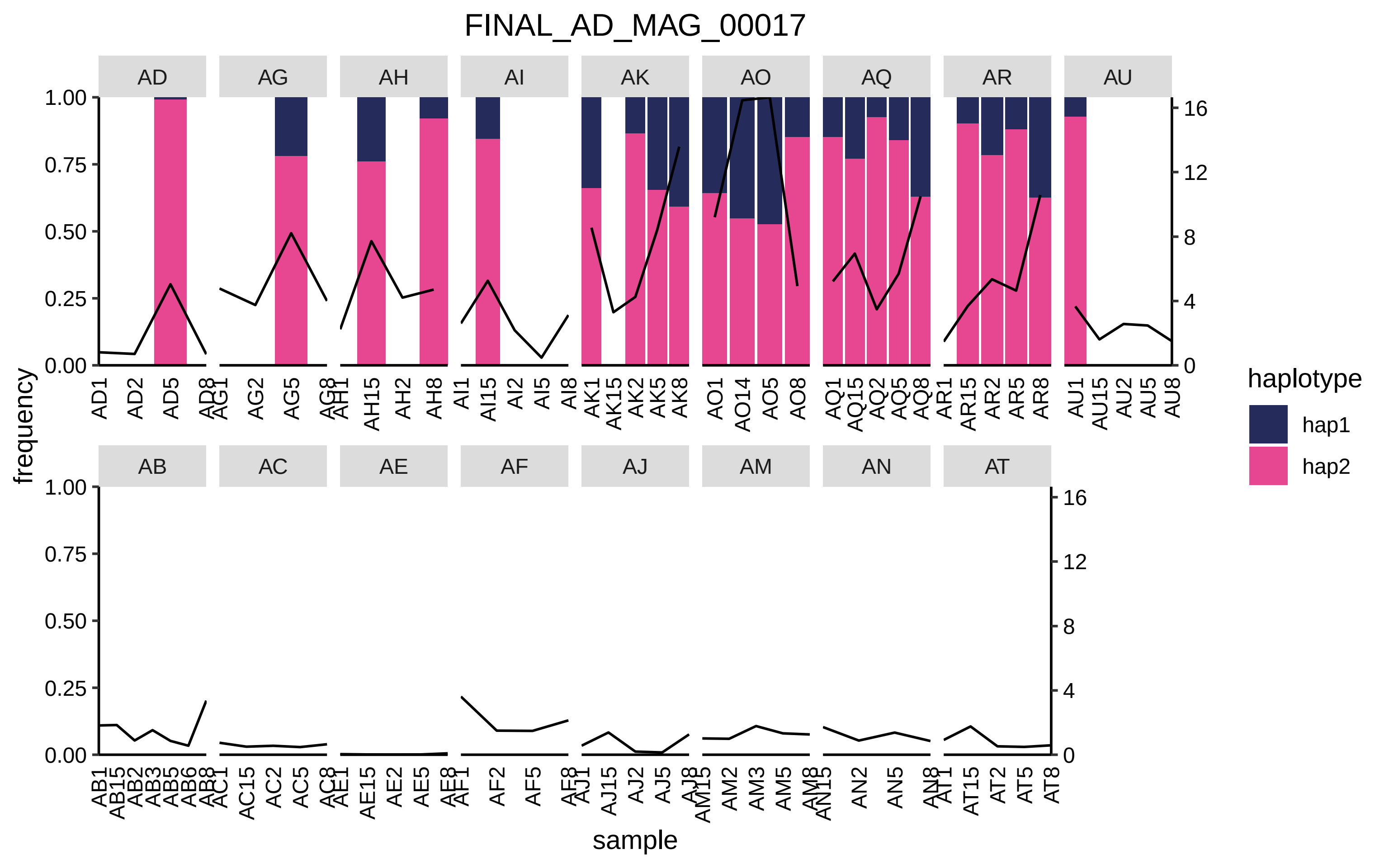
FINAL_AD_MAG_00017
AD
AG
AH
AI
AK
AO
AQ
AR
AU
AB
AC
AE
AF
AJ
AM
AN
AT
frequency
sample
haplotype
hap1
hap2

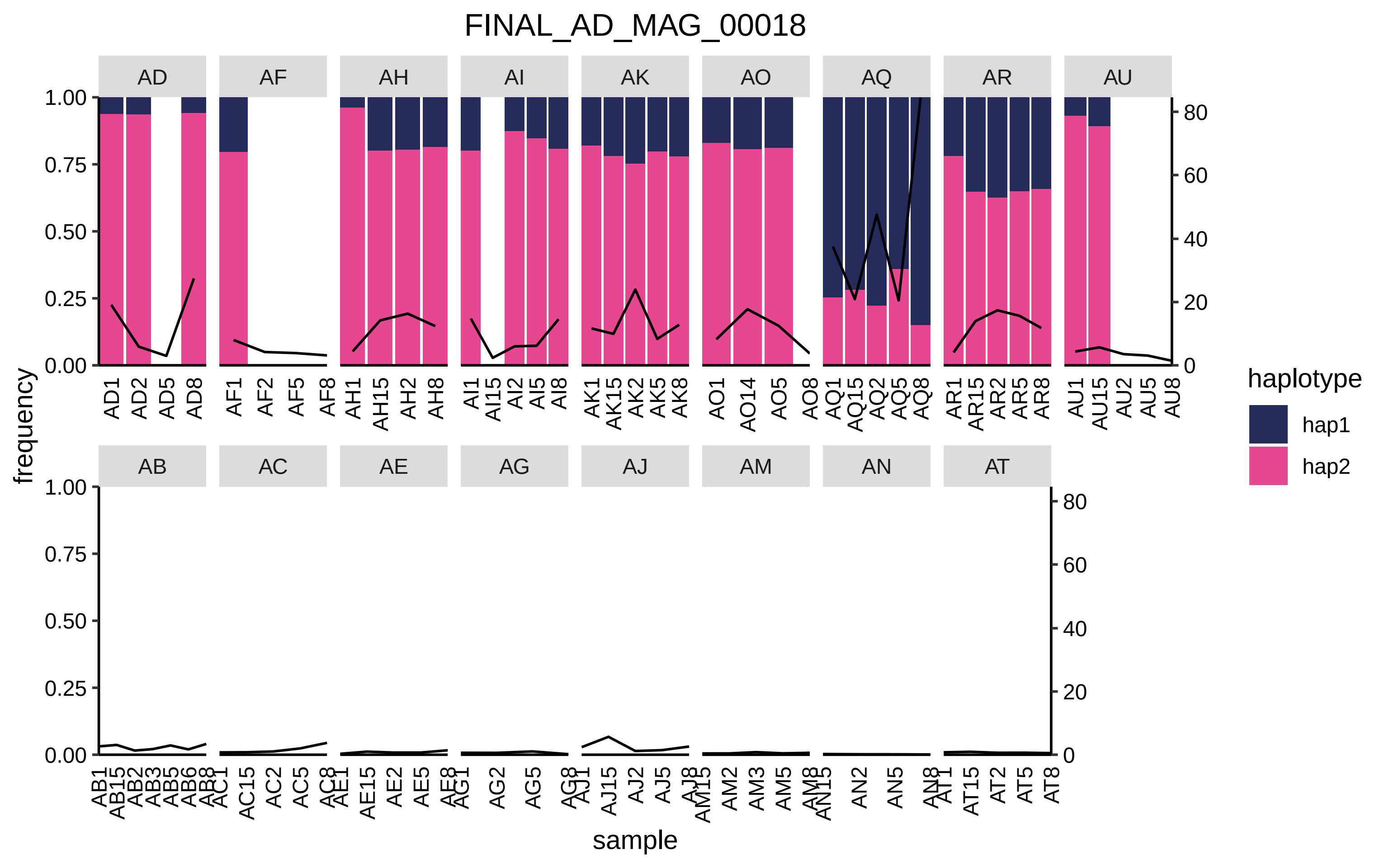

FINAL_AD_MAG_00018
AD
AF
AH
AI
AK
AO
AQ
AR
AU
AB
AC
AE
AG
AJ
AM
AN
AT
frequency
sample
haplotype
hap1
hap2

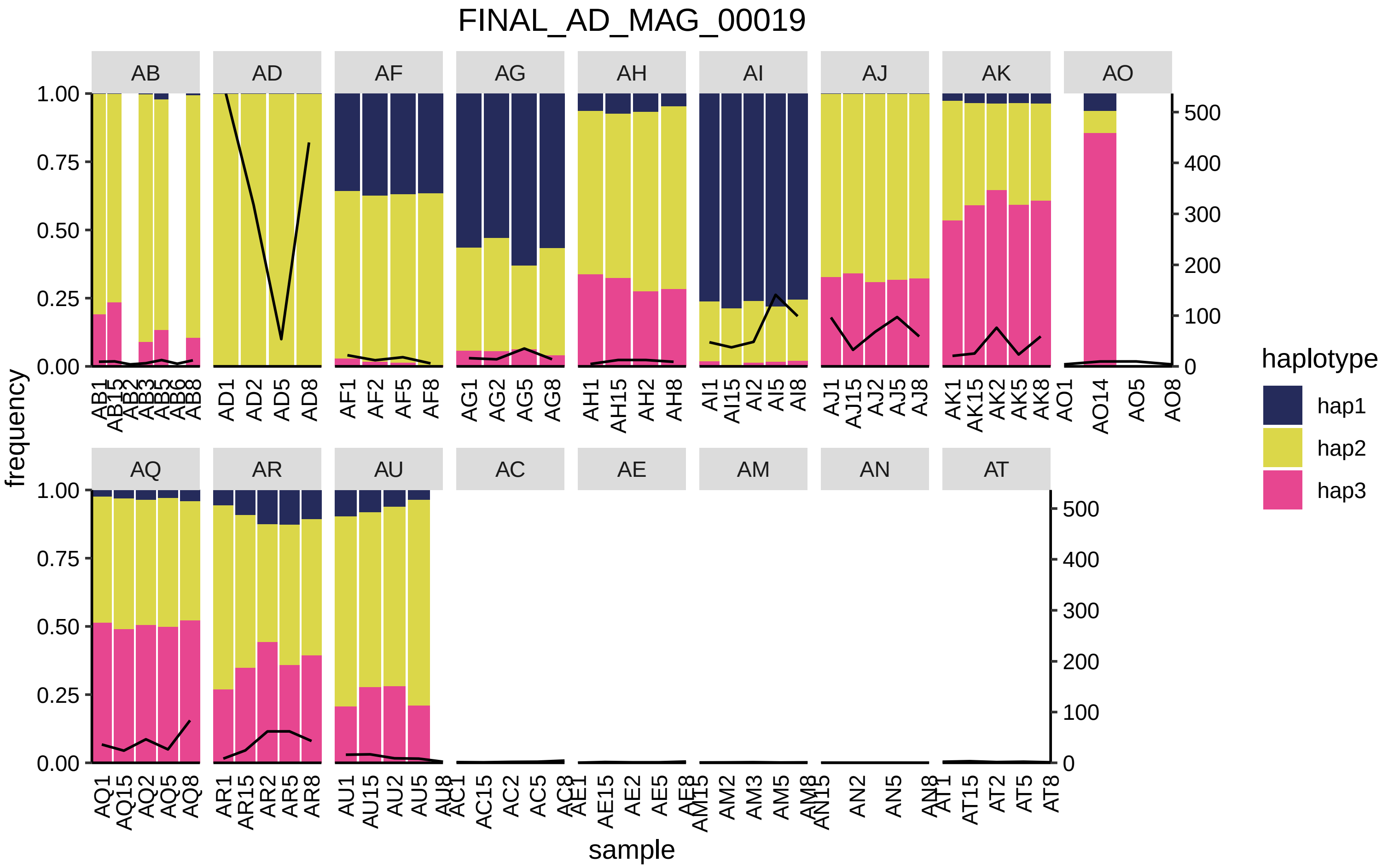

FINAL_AD_MAG_00019
AB
AD
AF
AG
AH
AI
AJ
AK
AO
AQ
AR
AU
AC
AE
AM
AN
AT
frequency
sample
haplotype
hap1
hap2
hap3
1.00
0.75
0.50
0.25
0.00
500
400
300
200
100
0

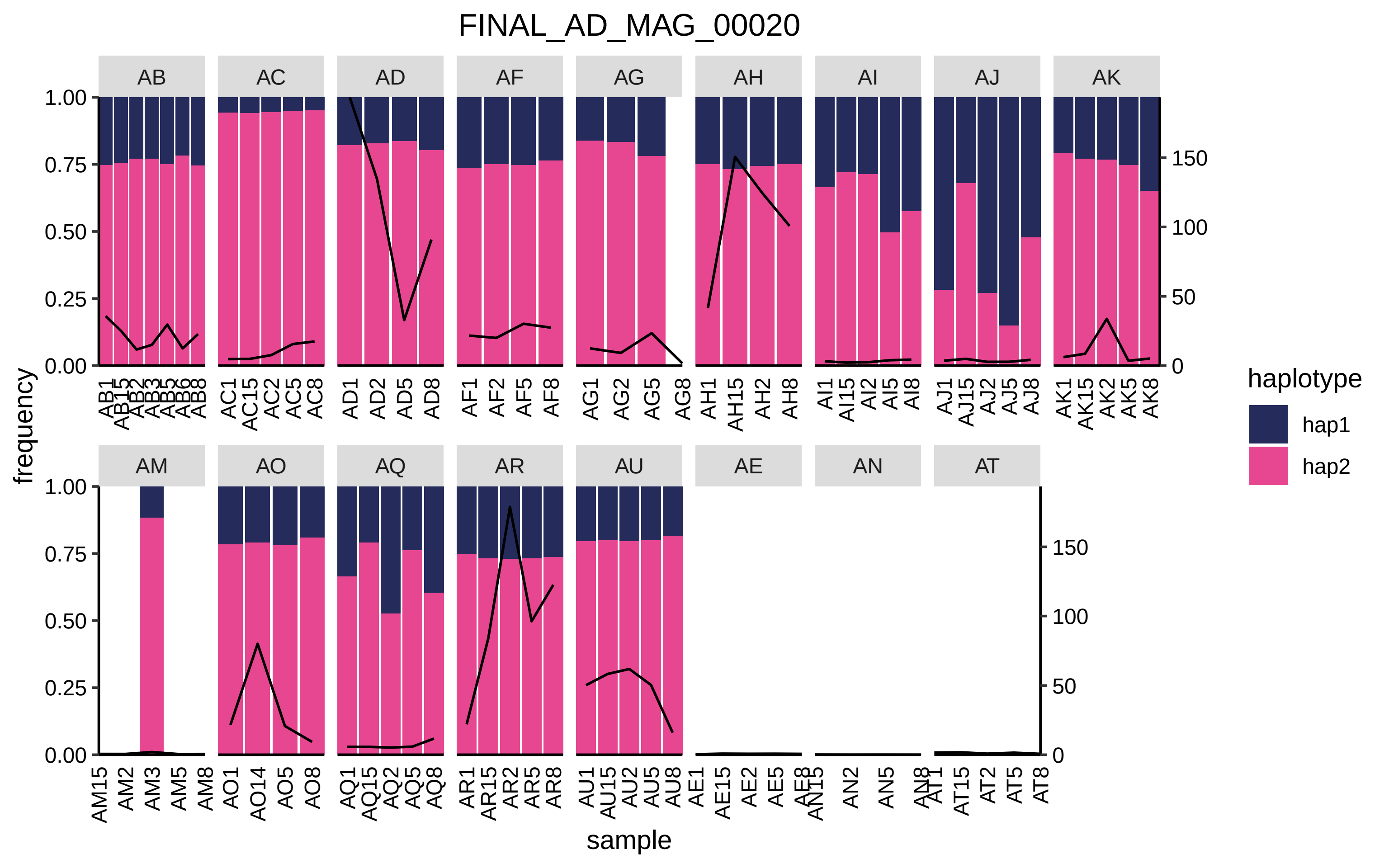

FINAL_AD_MAG_00020
AB
AC
AD
AF
AG
AH
AI
AJ
AK
AM
AO
AQ
AR
AU
AE
AN
AT
frequency
sample
haplotype
hap1
hap2

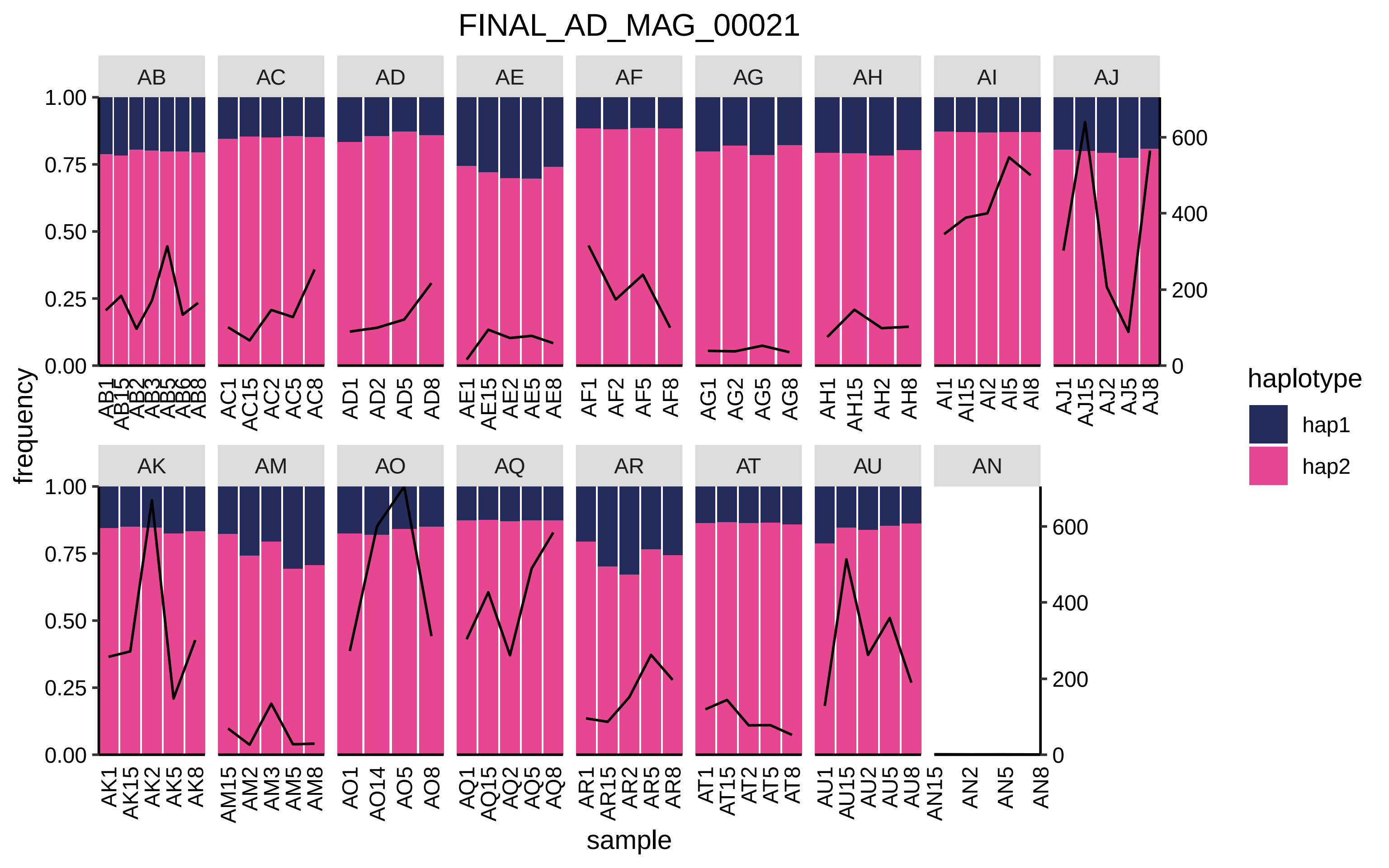

FINAL_AD_MAG_00021
AB
AC
AD
AE
AF
AG
AH
AI
AJ
AK
AM
AO
AQ
AR
AT
AU
AN
frequency
sample
1.00
0.75
0.50
0.25
0.00
0
200
400
600
AB1 AB15 AB2 AB3 AB5 AB6 AB8
AC1 AC15 AC2 AC5 AC8
AD1 AD2 AD5 AD8
AE1 AE15 AE2 AE5 AE8
AF1 AF2 AF5 AF8
AG1 AG2 AG5 AG8
AH1 AH15 AH2 AH8
AI1 AI15 AI2 AI5 AI8
AJ1 AJ15 AJ2 AJ5 AJ8
AK1 AK15 AK2 AK5 AK8
AM15 AM2 AM3 AM5 AM8
AO1 AO14 AO5 AO8
AQ1 AQ15 AQ2 AQ5 AQ8
AR1 AR15 AR2 AR5 AR8
AT1 AT15 AT2 AT5 AT8
AU1 AU15 AU2 AU5 AU8
AN15 AN2 AN5 AN8
haplotype
hap1
hap2

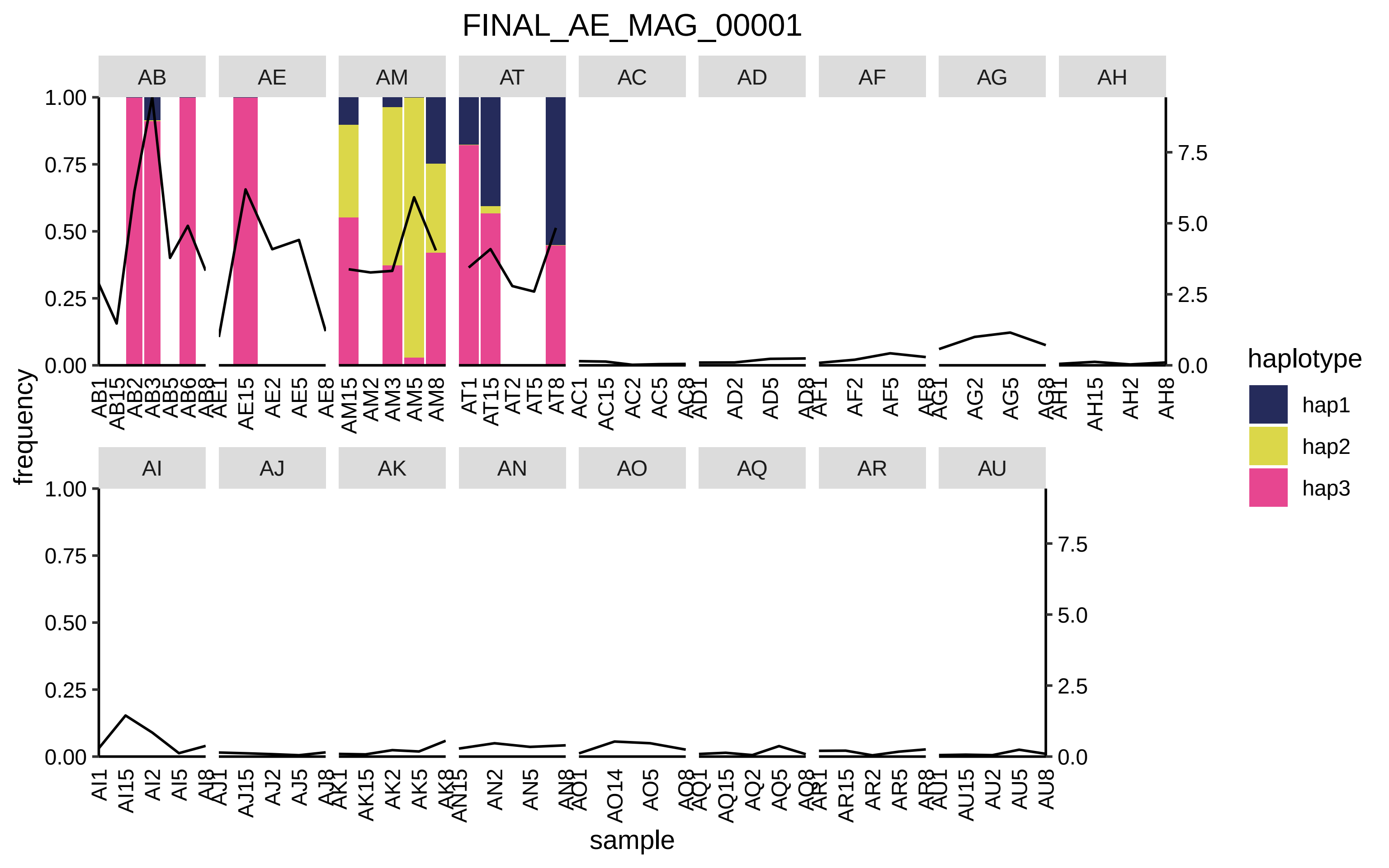

FINAL_AE_MAG_00001
AB
AE
AM
AT
AC
AD
AF
AG
AH
AI
AJ
AK
AN
AO
AQ
AR
AU
frequency
sample
1.00
0.75
0.50
0.25
0.00
7.5
5.0
2.5
0.0
AB1 AB15 AB2 AB3 AB5 AB6 AB8
AE1 AE15 AE2 AE5 AE8
AM15 AM2 AM3 AM5 AM8
AT1 AT15 AT2 AT5 AT8
AC1 AC15 AC2 AC5 AC8
AD1 AD2 AD5 AD8
AF1 AF2 AF5 AF8
AG1 AG2 AG5 AG8
AH1 AH15 AH2 AH8
AI1 AI15 AI2 AI5 AI8
AJ1 AJ15 AJ2 AJ5 AJ8
AK1 AK15 AK2 AK5 AK8
AN15 AN2 AN5 AN8
AO1 AO14 AO5 AO8
AQ1 AQ15 AQ2 AQ5 AQ8
AR1 AR15 AR2 AR5 AR8
AU1 AU15 AU2 AU5 AU8
haplotype
hap1
hap2
hap3

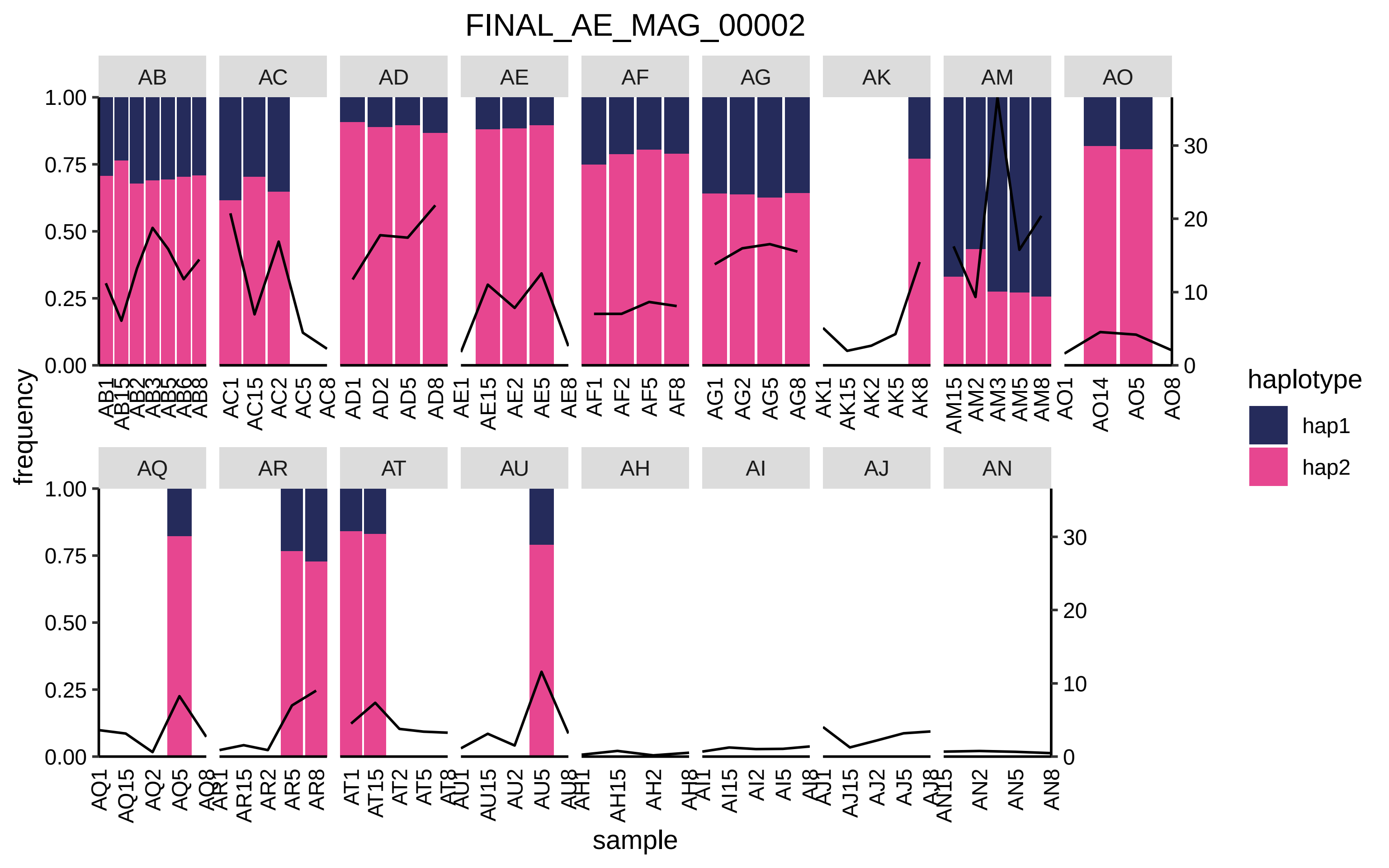
FINAL_AE_MAG_00002
AB
AC
AD
AE
AF
AG
AK
AM
AO
AQ
AR
AT
AU
AH
AI
AJ
AN
frequency
1.00
0.75
0.50
0.25
0.00
0
10
20
30
AB1 AB15 AB2 AB3 AB5 AB6 AB8
AC1 AC15 AC2 AC5 AC8
AD1 AD2 AD5 AD8
AE1 AE15 AE2 AE5 AE8
AF1 AF2 AF5 AF8
AG1 AG2 AG5 AG8
AK1 AK15 AK2 AK5 AK8
AM15 AM2 AM3 AM5 AM8
AO1 AO14 AO5 AO8
AQ1 AQ15 AQ2 AQ5 AQ8
AR1 AR15 AR2 AR5 AR8
AT1 AT15 AT2 AT5 AT8
AU1 AU15 AU2 AU5 AU8
AH1 AH15 AH2 AH8
AI1 AI15 AI2 AI5 AI8
AJ1 AJ15 AJ2 AJ5 AJ8
AN15 AN2 AN5 AN8
sample
haplotype
hap1
hap2

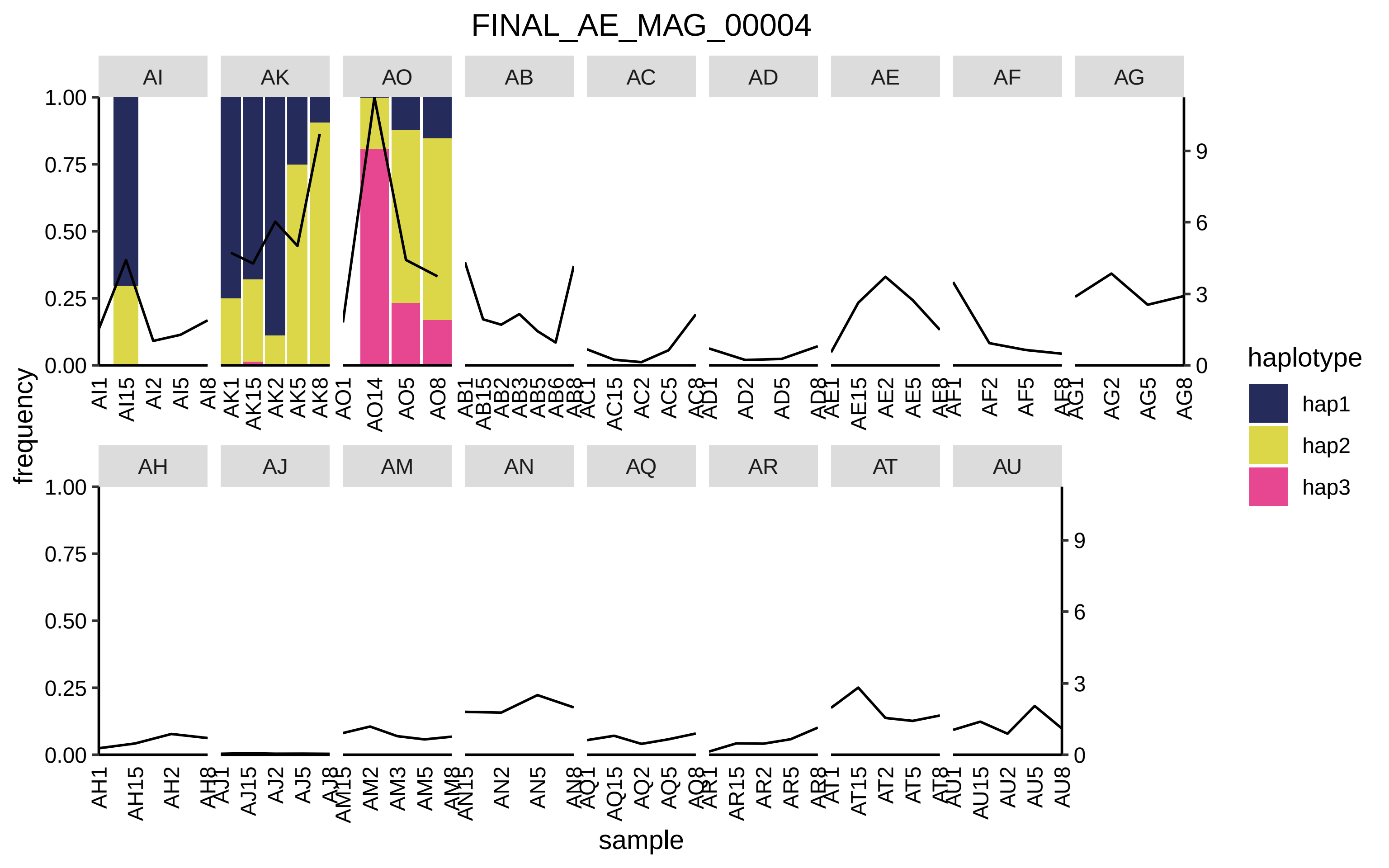
FINAL_AE_MAG_00004
AI
AK
AO
AB
AC
AD
AE
AF
AG
AH
AJ
AM
AN
AQ
AR
AT
AU
1.00
0.75
0.50
0.25
0.00
9
6
3
0
frequency
sample
AI1
AI15
AI2
AI5
AI8
AK1
AK15
AK2
AK5
AK8
AO1
AO14
AO5
AO8
AB1
AB15
AB2
AB3
AB5
AB6
AB8
AC1
AC15
AC2
AC5
AC8
AD1
AD2
AD5
AD8
AE1
AE15
AE2
AE5
AE8
AF1
AF2
AF5
AF8
AG1
AG2
AG5
AG8
AH1
AH15
AH2
AH8
AJ1
AJ15
AJ2
AJ5
AJ8
AM15
AM2
AM3
AM5
AM8
AN15
AN2
AN5
AN8
AQ1
AQ15
AQ2
AQ5
AQ8
AR1
AR15
AR2
AR5
AR8
AT1
AT15
AT2
AT5
AT8
AU1
AU15
AU2
AU5
AU8
haplotype
hap1
hap2
hap3

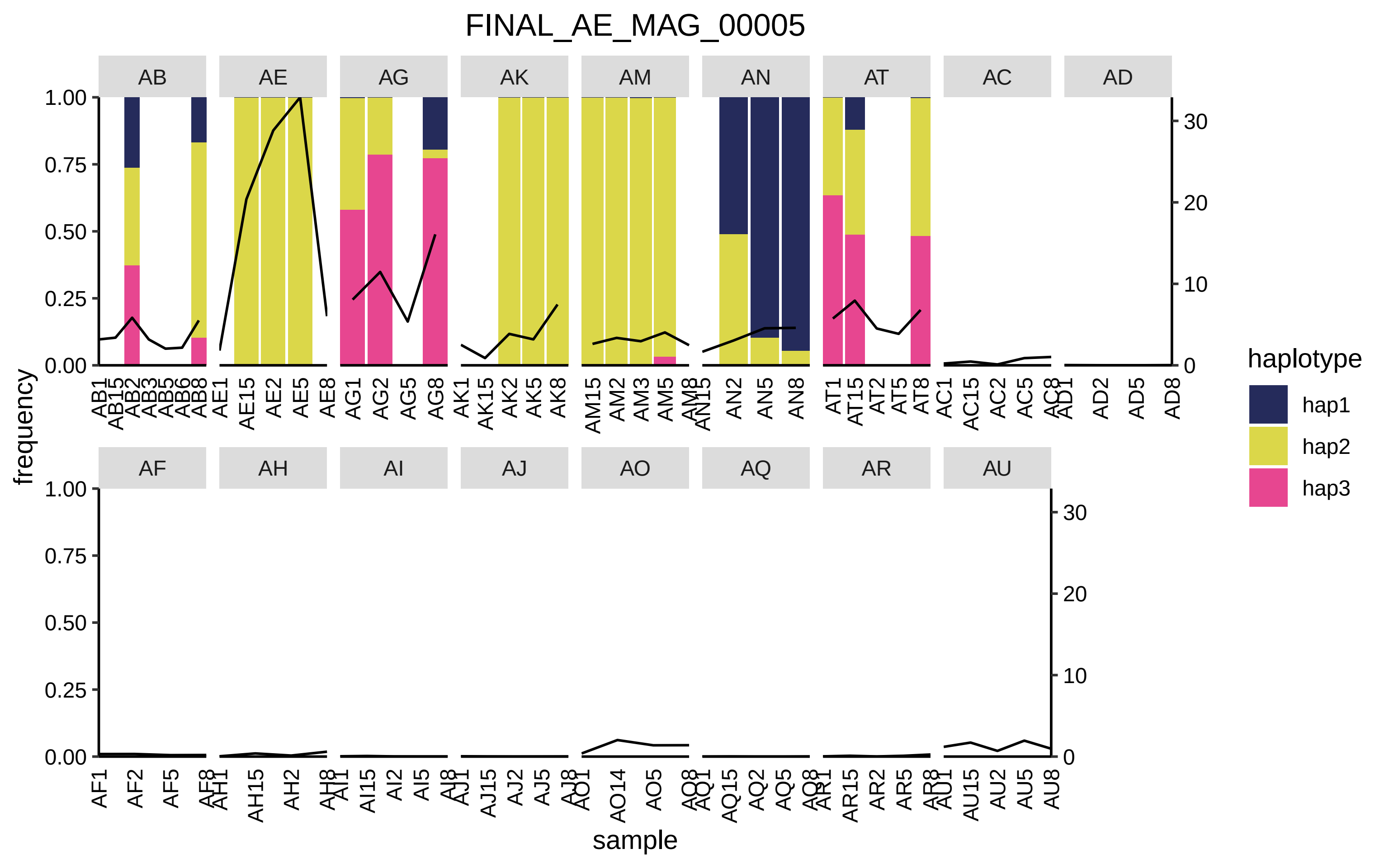

FINAL_AE_MAG_00005
AB
AE
AG
AK
AM
AN
AT
AC
AD
AF
AH
AI
AJ
AO
AQ
AR
AU
frequency
sample
haplotype
hap1
hap2
hap3
1.00
0.75
0.50
0.25
0.00
30
20
10
0
AB1
AB15
AB2
AB3
AB5
AB6
AB8
AE1
AE15
AE2
AE5
AE8
AG1
AG2
AG5
AG8
AK1
AK15
AK2
AK5
AK8
AM15
AM2
AM3
AM5
AM8
AN15
AN2
AN5
AN8
AT1
AT15
AT2
AT5
AT8
AC1
AC15
AC2
AC5
AC8
AD1
AD2
AD5
AD8
AF1
AF2
AF5
AF8
AH1
AH15
AH2
AH8
AI1
AI15
AI2
AI5
AI8
AJ1
AJ15
AJ2
AJ5
AJ8
AO1
AO14
AO5
AO8
AQ1
AQ15
AQ2
AQ5
AQ8
AR1
AR15
AR2
AR5
AR8
AU1
AU15
AU2
AU5
AU8

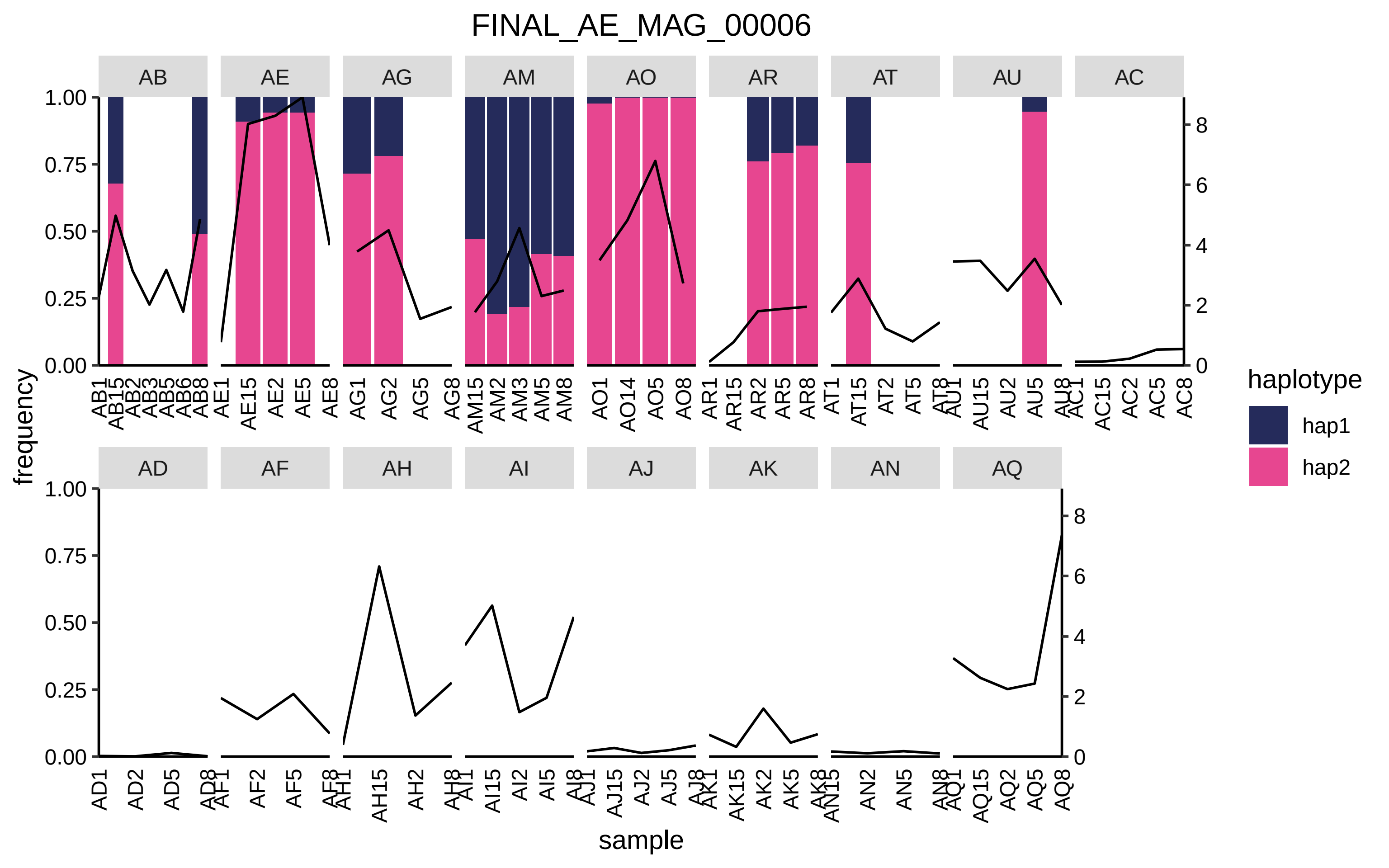

FINAL_AE_MAG_00006
AB
AE
AG
AM
AO
AR
AT
AU
AC
AD
AF
AH
AI
AJ
AK
AN
AQ
frequency
sample
haplotype
hap1
hap2
1.00
0.75
0.50
0.25
0.00
0
2
4
6
8
AB1 AB15 AB2 AB3 AB5 AB6 AB8
AE1 AE15 AE2 AE5 AE8
AG1 AG2 AG5 AG8
AM15 AM2 AM3 AM5 AM8
AO1 AO14 AO5 AO8
AR1 AR15 AR2 AR5 AR8
AT1 AT15 AT2 AT5 AT8
AU1 AU15 AU2 AU5 AU8
AC1 AC15 AC2 AC5 AC8
AD1 AD2 AD5 AD8
AF1 AF2 AF5 AF8
AH1 AH15 AH2 AH8
AI1 AI15 AI2 AI5 AI8
AJ1 AJ15 AJ2 AJ5 AJ8
AK1 AK15 AK2 AK5 AK8
AN15 AN2 AN5 AN8
AQ1 AQ15 AQ2 AQ5 AQ8

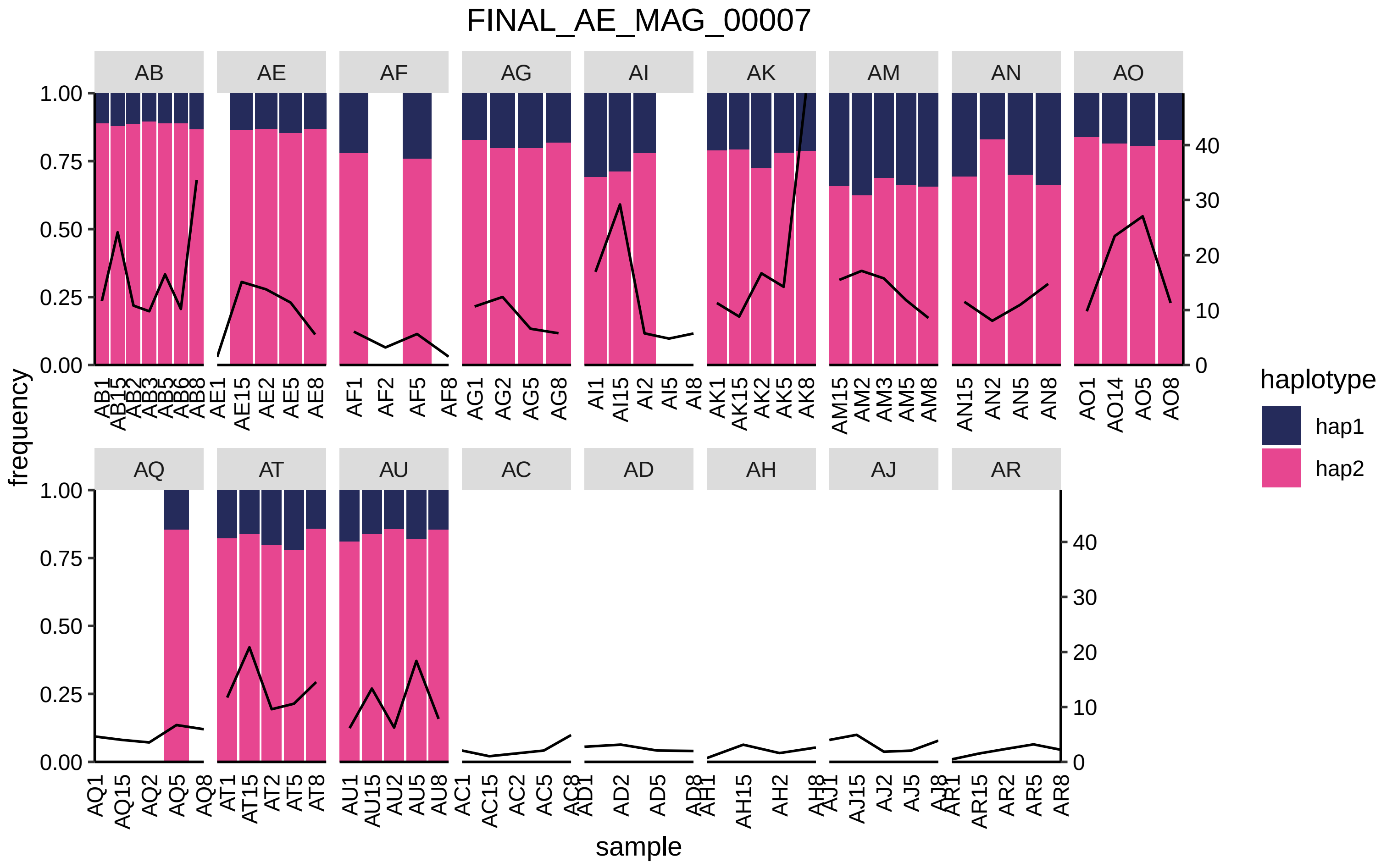

FINAL_AE_MAG_00007
AB
AE
AF
AG
AI
AK
AM
AN
AO
AQ
AT
AU
AC
AD
AH
AJ
AR
frequency
sample
1.00
0.75
0.50
0.25
0.00
40
30
20
10
0
haplotype
hap1
hap2

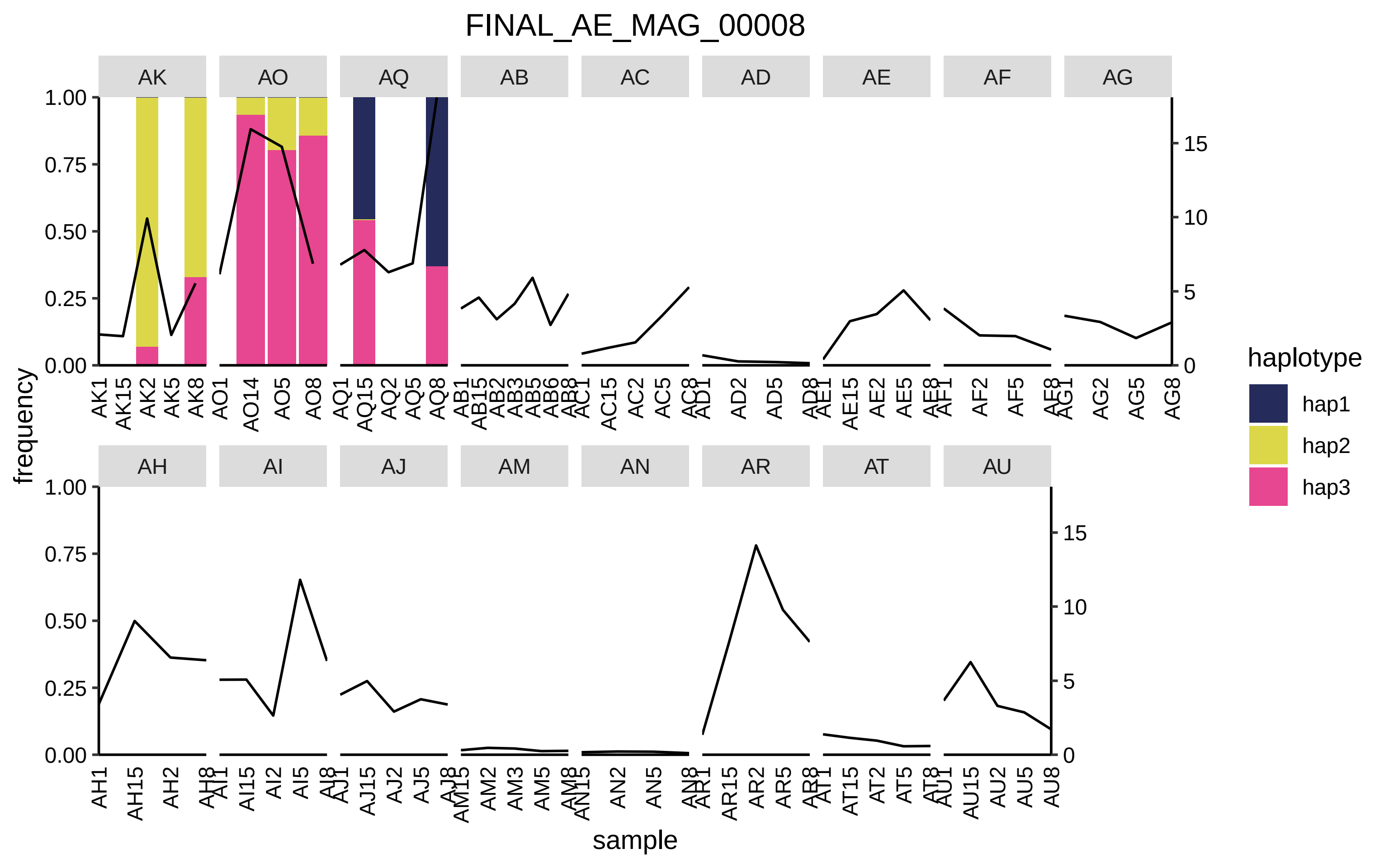

FINAL_AE_MAG_00008
AK
AO
AQ
AB
AC
AD
AE
AF
AG
AH
AI
AJ
AM
AN
AR
AT
AU
frequency
sample
haplotype
hap1
hap2
hap3
1.00
0.75
0.50
0.25
0.00
15
10
5
0

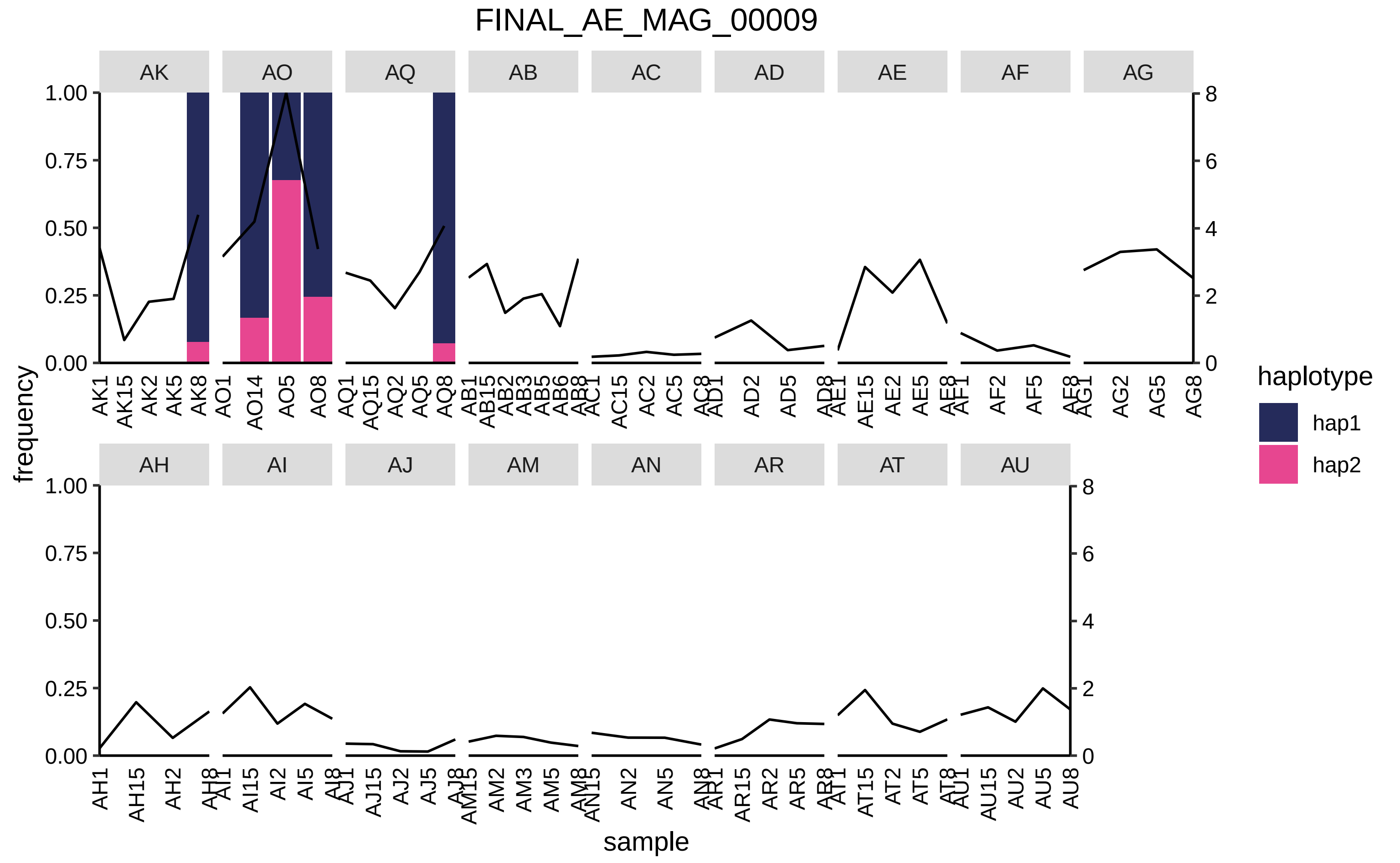

FINAL_AE_MAG_00009
AK
AO
AQ
AB
AC
AD
AE
AF
AG
AH
AI
AJ
AM
AN
AR
AT
AU
frequency
sample
haplotype
hap1
hap2
1.00
0.75
0.50
0.25
0.00
8
6
4
2
0
AK1 AK15 AK2 AK5 AK8
AO1 AO14 AO5 AO8
AQ1 AQ15 AQ2 AQ5 AQ8
AB1 AB15 AB2 AB3 AB5 AB6 AB8
AC1 AC15 AC2 AC5 AC8
AD1 AD2 AD5 AD8
AE1 AE15 AE2 AE5 AE8
AF1 AF2 AF5 AF8
AG1 AG2 AG5 AG8
AH1 AH15 AH2 AH8
AI1 AI15 AI2 AI5 AI8
AJ1 AJ15 AJ2 AJ5 AJ8
AM15 AM2 AM3 AM5 AM8
AN15 AN2 AN5 AN8
AR1 AR15 AR2 AR5 AR8
AT1 AT15 AT2 AT5 AT8
AU1 AU15 AU2 AU5 AU8

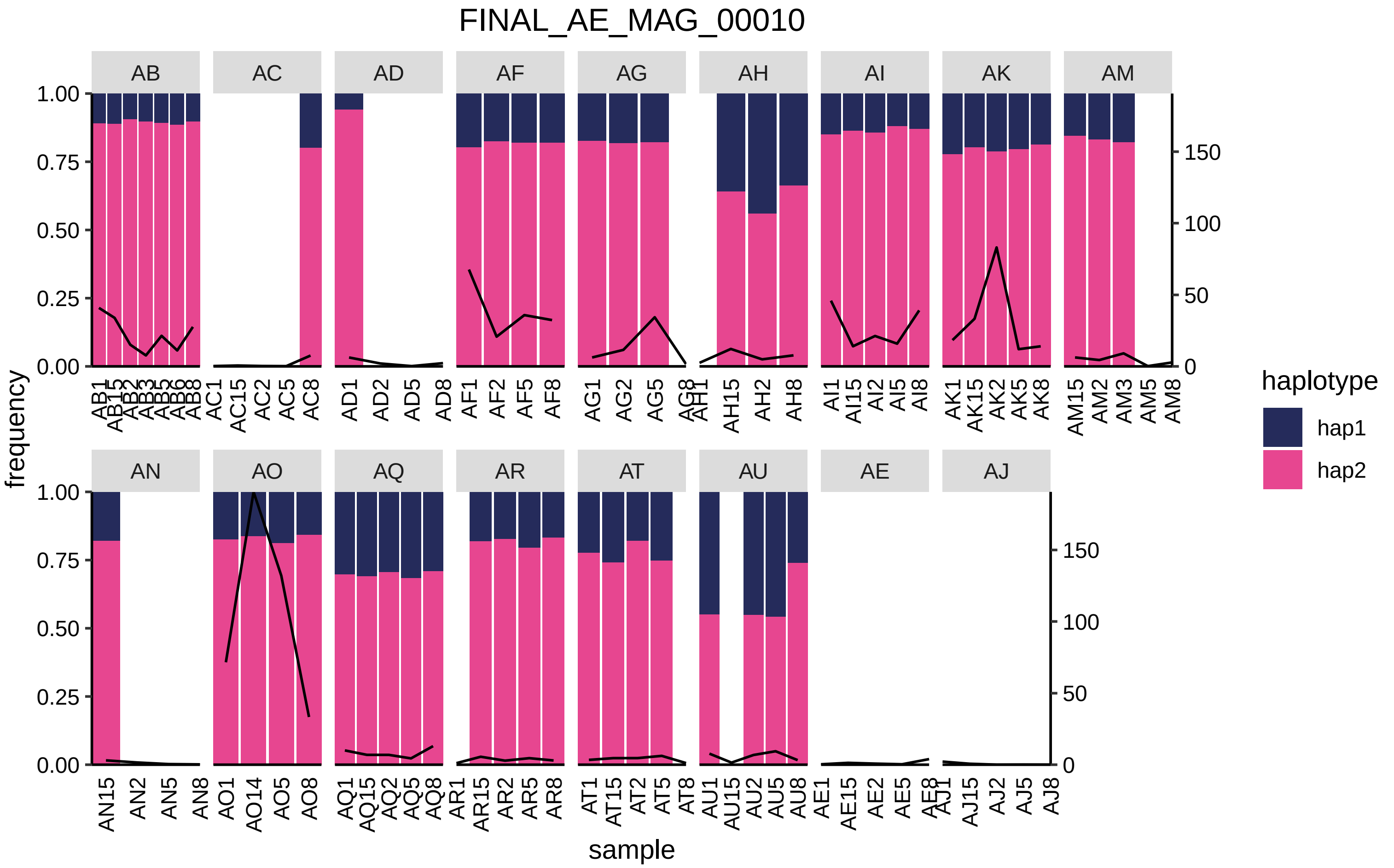

FINAL_AE_MAG_00010
AB
AC
AD
AF
AG
AH
AI
AK
AM
AN
AO
AQ
AR
AT
AU
AE
AJ
frequency
sample
haplotype
hap1
hap2

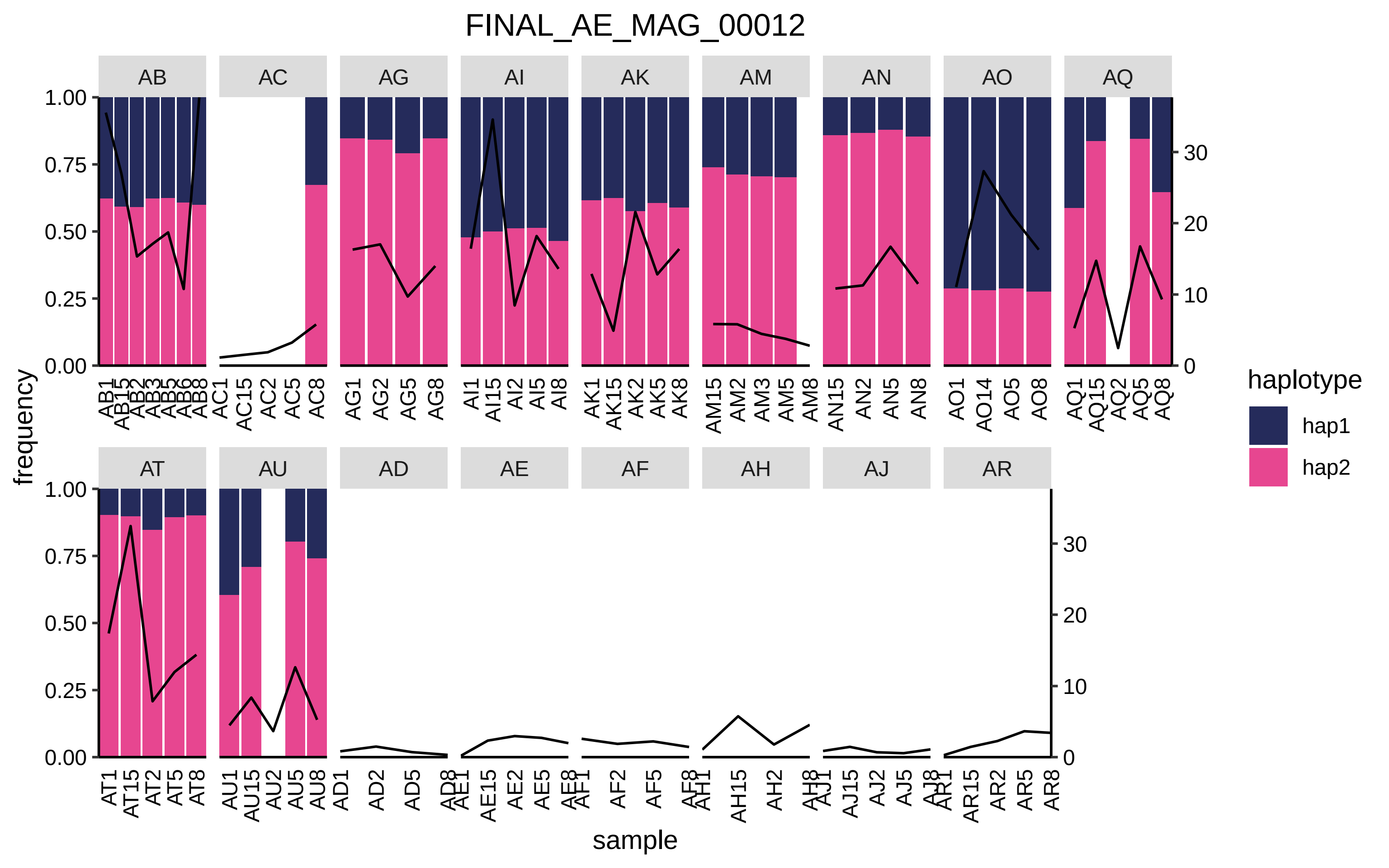

FINAL_AE_MAG_00012
AB
AC
AG
AI
AK
AM
AN
AO
AQ
AT
AU
AD
AE
AF
AH
AJ
AR
frequency
sample
haplotype
hap1
hap2
1.00
0.75
0.50
0.25
0.00
30
20
10
0
AB1 AB15 AB2 AB3 AB5 AB6 AB8
AC1 AC15 AC2 AC5 AC8
AG1 AG2 AG5 AG8
AI1 AI15 AI2 AI5 AI8
AK1 AK15 AK2 AK5 AK8
AM15 AM2 AM3 AM5 AM8
AN15 AN2 AN5 AN8
AO1 AO14 AO5 AO8
AQ1 AQ15 AQ2 AQ5 AQ8
AT1 AT15 AT2 AT5 AT8
AU1 AU15 AU2 AU5 AU8
AD1 AD2 AD5 AD8
AE1 AE15 AE2 AE5 AE8
AF1 AF2 AF5 AF8
AH1 AH15 AH2 AH8
AJ1 AJ15 AJ2 AJ5 AJ8
AR1 AR15 AR2 AR5 AR8

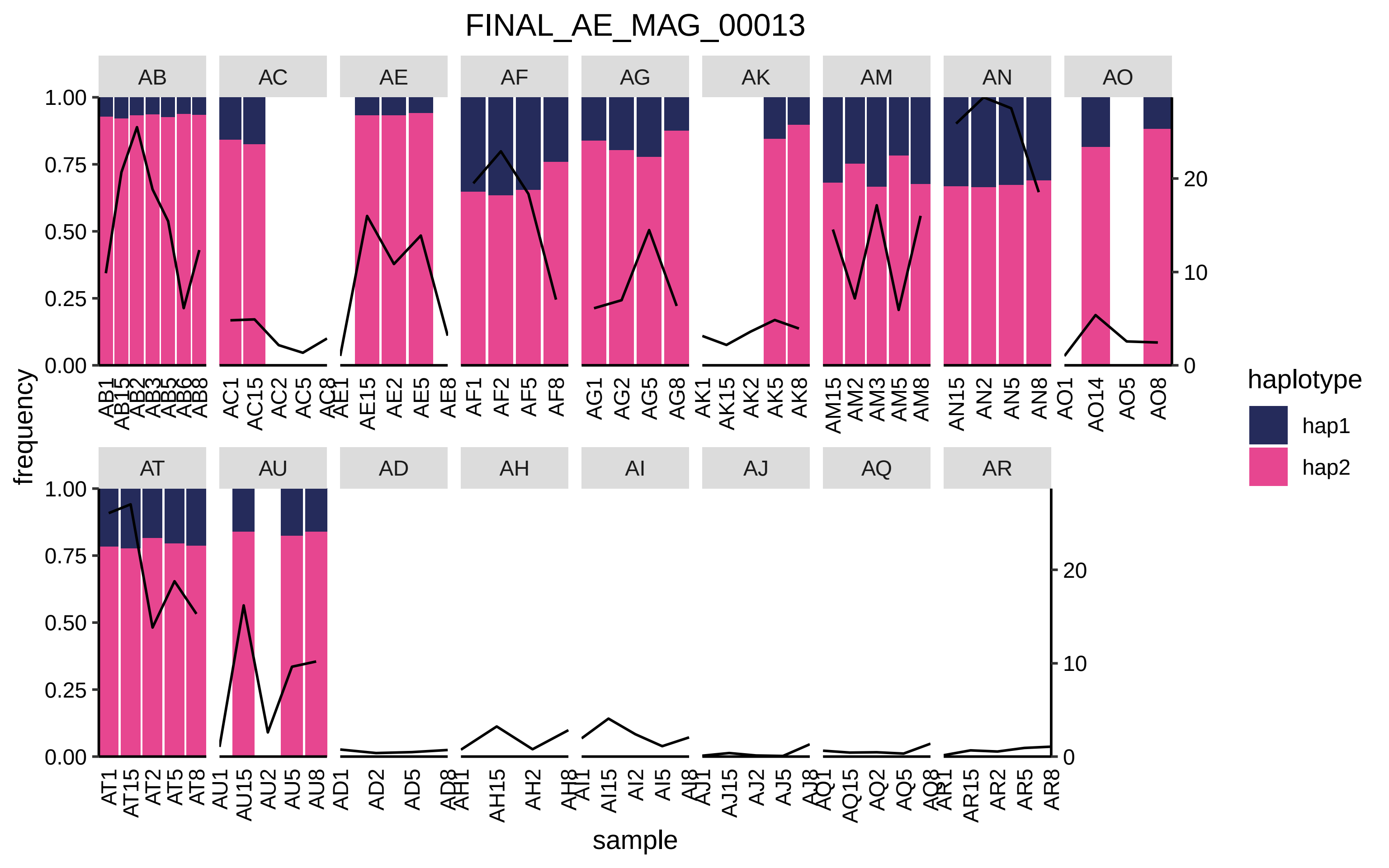
FINAL_AE_MAG_00013
AB
AC
AE
AF
AG
AK
AM
AN
AO
AT
AU
AD
AH
AI
AJ
AQ
AR
frequency
sample
haplotype
hap1
hap2
1.00
0.75
0.50
0.25
0.00
20
10
0

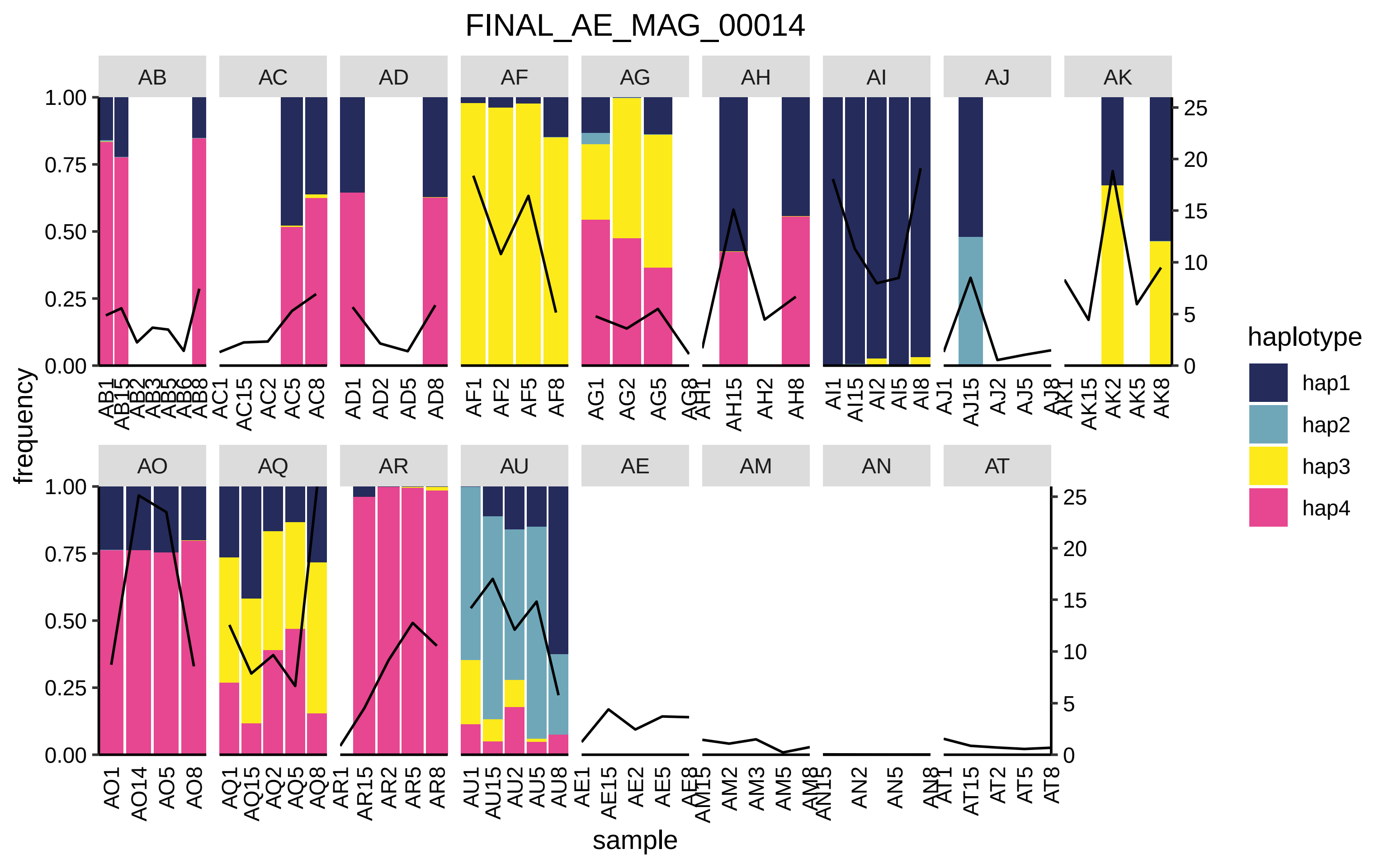

FINAL_AE_MAG_00014
AB
AC
AD
AF
AG
AH
AI
AJ
AK
AO
AQ
AR
AU
AE
AM
AN
AT
frequency
sample
haplotype
hap1
hap2
hap3
hap4

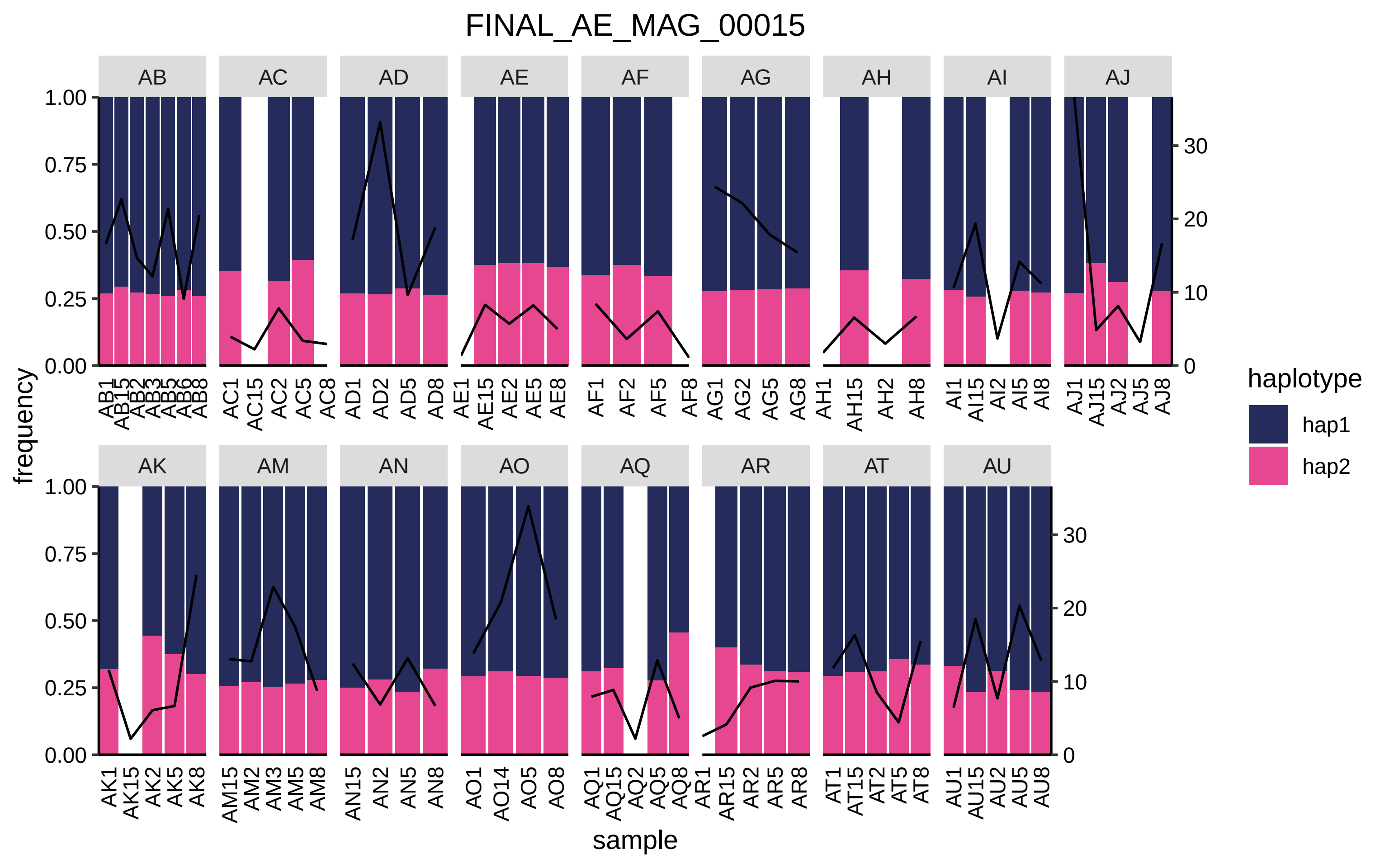

FINAL_AE_MAG_00015
AB
AC
AD
AE
AF
AG
AH
AI
AJ
AK
AM
AN
AO
AQ
AR
AT
AU
frequency
sample
haplotype
hap1
hap2
0.00
0.25
0.50
0.75
1.00
0
10
20
30
AB1 AB15 AB2 AB3 AB5 AB6 AB8
AC1 AC15 AC2 AC5 AC8
AD1 AD2 AD5 AD8
AE1 AE15 AE2 AE5 AE8
AF1 AF2 AF5 AF8
AG1 AG2 AG5 AG8
AH1 AH15 AH2 AH8
AI1 AI15 AI2 AI5 AI8
AJ1 AJ15 AJ2 AJ5 AJ8
AK1 AK15 AK2 AK5 AK8
AM15 AM2 AM3 AM5 AM8
AN15 AN2 AN5 AN8
AO1 AO14 AO5 AO8
AQ1 AQ15 AQ2 AQ5 AQ8
AR1 AR15 AR2 AR5 AR8
AT1 AT15 AT2 AT5 AT8
AU1 AU15 AU2 AU5 AU8

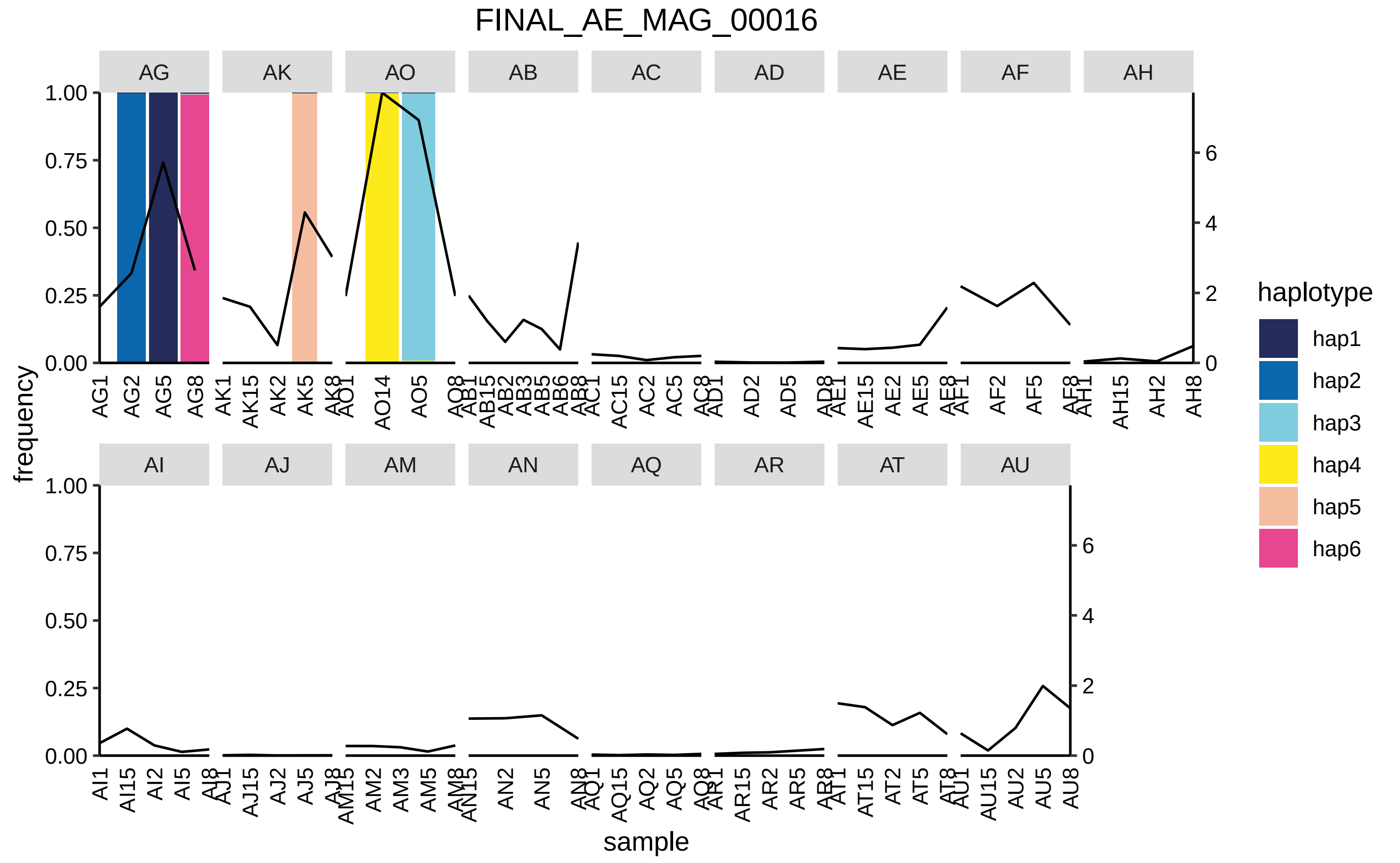
FINAL_AE_MAG_00016
AG
AK
AO
AB
AC
AD
AE
AF
AH
AI
AJ
AM
AN
AQ
AR
AT
AU
frequency
sample
haplotype
hap1
hap2
hap3
hap4
hap5
hap6

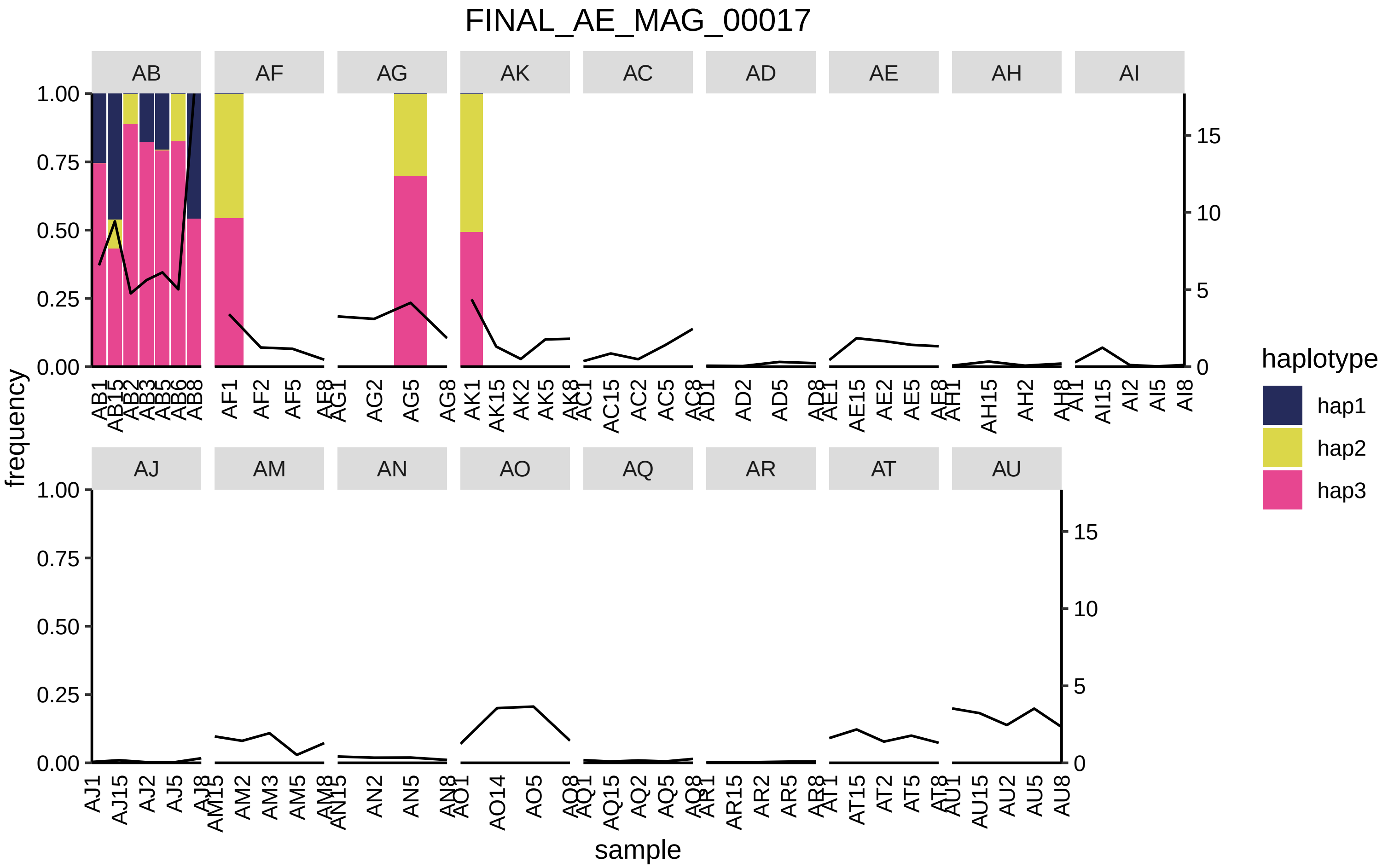
FINAL_AE_MAG_00017
AB
AF
AG
AK
AC
AD
AE
AH
AI
AJ
AM
AN
AO
AQ
AR
AT
AU
frequency
sample
haplotype
hap1
hap2
hap3

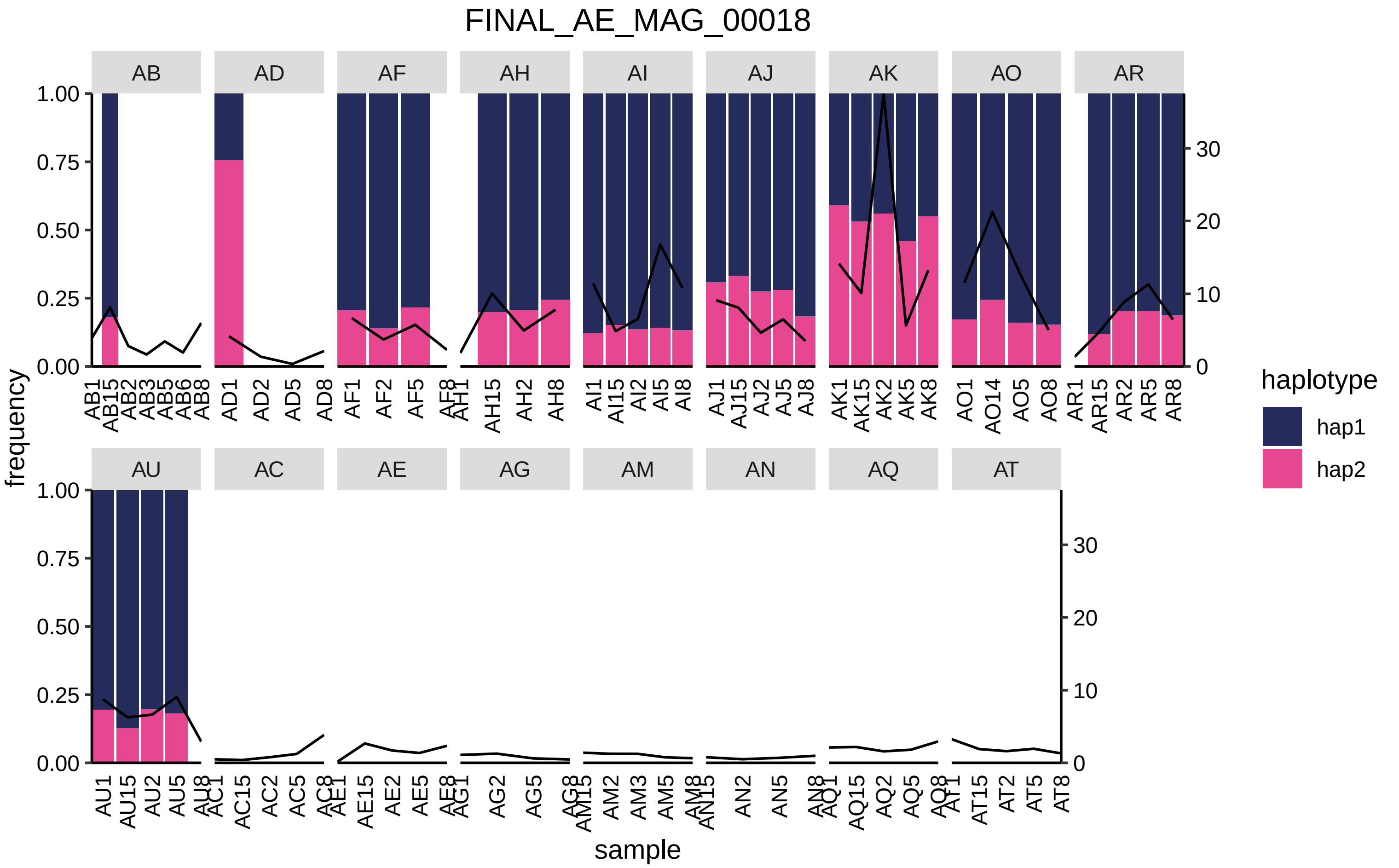

FINAL_AE_MAG_00018
AB
AD
AF
AH
AI
AJ
AK
AO
AR
AU
AC
AE
AG
AM
AN
AQ
AT
frequency
sample
haplotype
hap1
hap2
1.00
0.75
0.50
0.25
0.00
0
10
20
30
AB1 AB15 AB2 AB3 AB5 AB6 AB8
AD1 AD2 AD5 AD8
AF1 AF2 AF5 AF8
AH1 AH15 AH2 AH8
AI1 AI15 AI2 AI5 AI8
AJ1 AJ15 AJ2 AJ5 AJ8
AK1 AK15 AK2 AK5 AK8
AO1 AO14 AO5 AO8
AR1 AR15 AR2 AR5 AR8
AU1 AU15 AU2 AU5 AU8
AC1 AC15 AC2 AC5 AC8
AE1 AE15 AE2 AE5 AE8
AG1 AG2 AG5 AG8
AM15 AM2 AM3 AM5 AM8
AN15 AN2 AN5 AN8
AQ1 AQ15 AQ2 AQ5 AQ8
AT1 AT15 AT2 AT5 AT8

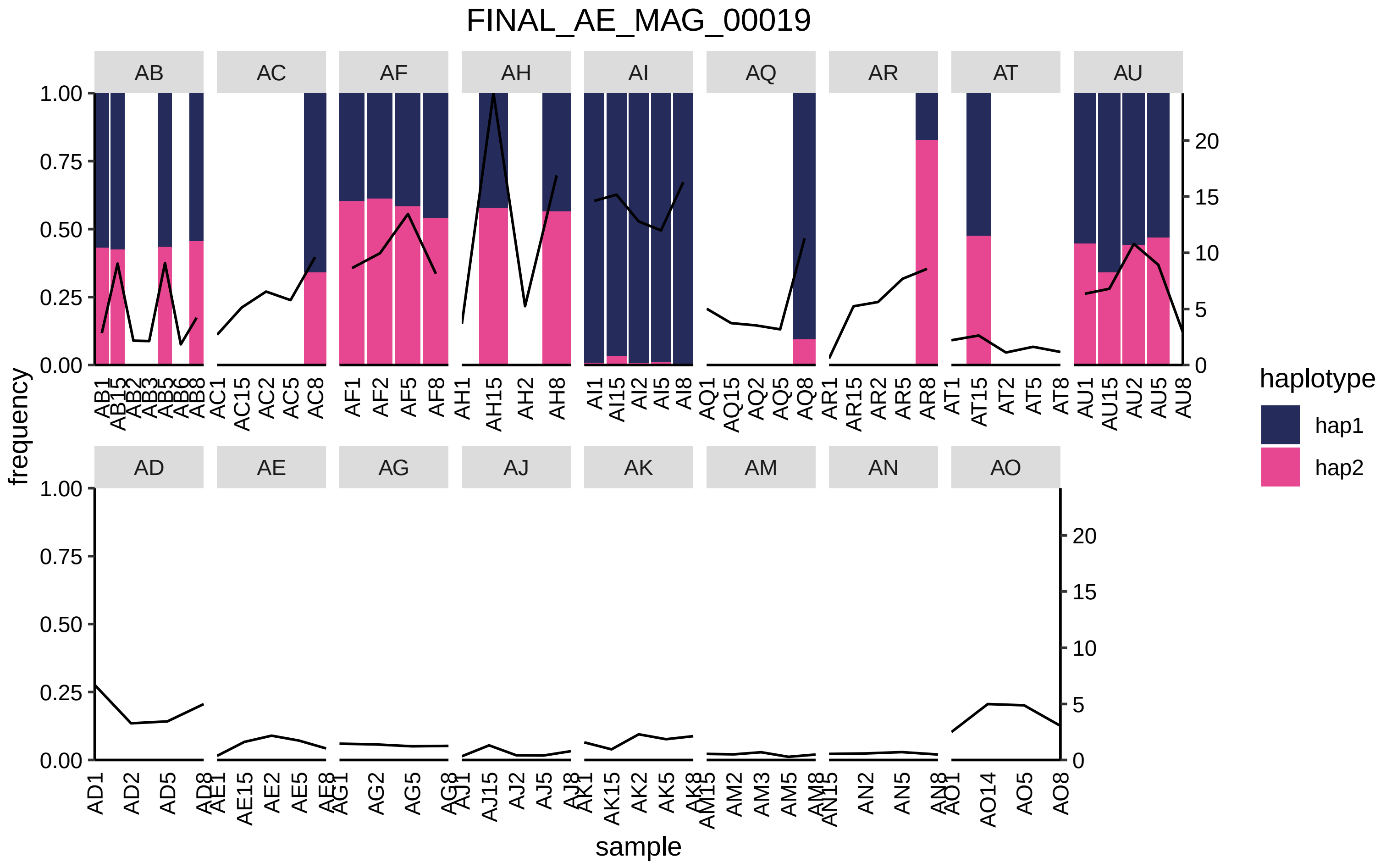

FINAL_AE_MAG_00019
AB
AC
AF
AH
AI
AQ
AR
AT
AU
AD
AE
AG
AJ
AK
AM
AN
AO
frequency
sample
haplotype
hap1
hap2

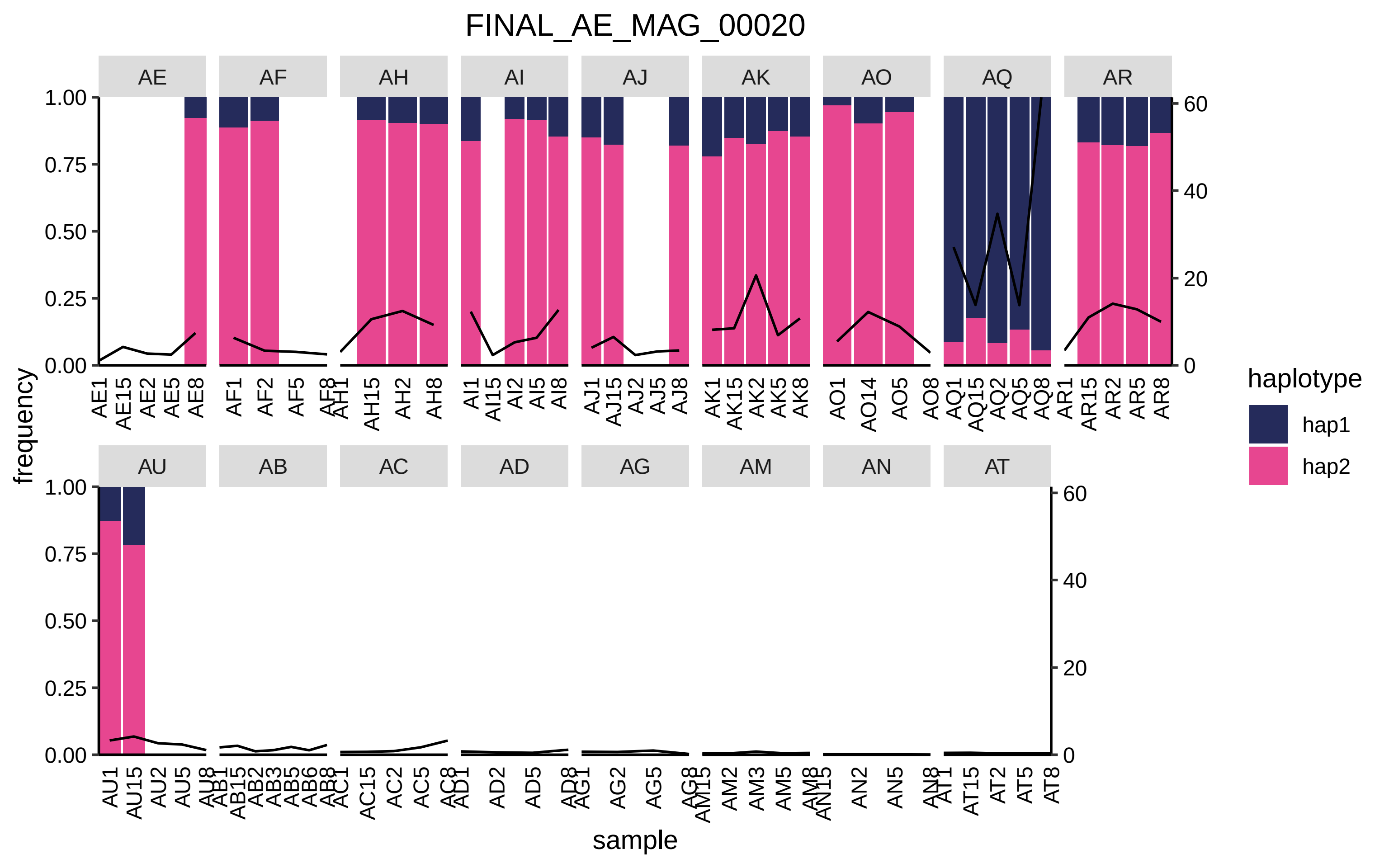
FINAL_AE_MAG_00020
AE
AF
AH
AI
AJ
AK
AO
AQ
AR
AU
AB
AC
AD
AG
AM
AN
AT
frequency
sample
haplotype
hap1
hap2
1.00
0.75
0.50
0.25
0.00
60
40
20
0

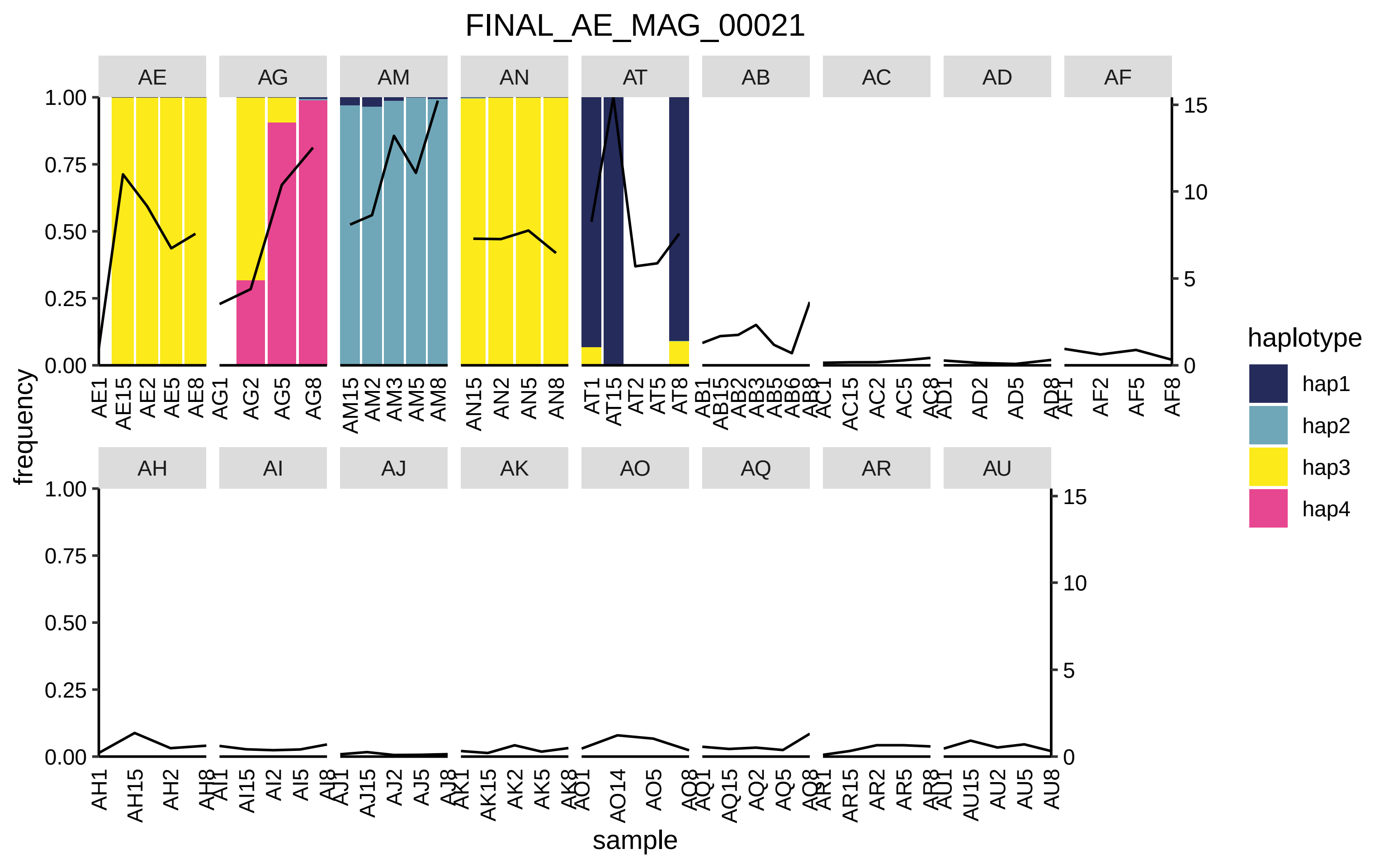
FINAL_AE_MAG_00021
AE
AG
AM
AN
AT
AB
AC
AD
AF
AH
AI
AJ
AK
AO
AQ
AR
AU
frequency
sample
1.00
0.75
0.50
0.25
0.00
15
10
5
0
haplotype
hap1
hap2
hap3
hap4
AE1 AE15 AE2 AE5 AE8
AG1 AG2 AG5 AG8
AM15 AM2 AM3 AM5 AM8
AN15 AN2 AN5 AN8
AT1 AT15 AT2 AT5 AT8
AB1 AB15 AB2 AB3 AB5 AB6 AB8
AC1 AC15 AC2 AC5 AC8
AD1 AD2 AD5 AD8
AF1 AF2 AF5 AF8
AH1 AH15 AH2 AH8
AI1 AI15 AI2 AI5 AI8
AJ1 AJ15 AJ2 AJ5 AJ8
AK1 AK15 AK2 AK5 AK8
AO1 AO14 AO5 AO8
AQ1 AQ15 AQ2 AQ5 AQ8
AR1 AR15 AR2 AR5 AR8
AU1 AU15 AU2 AU5 AU8

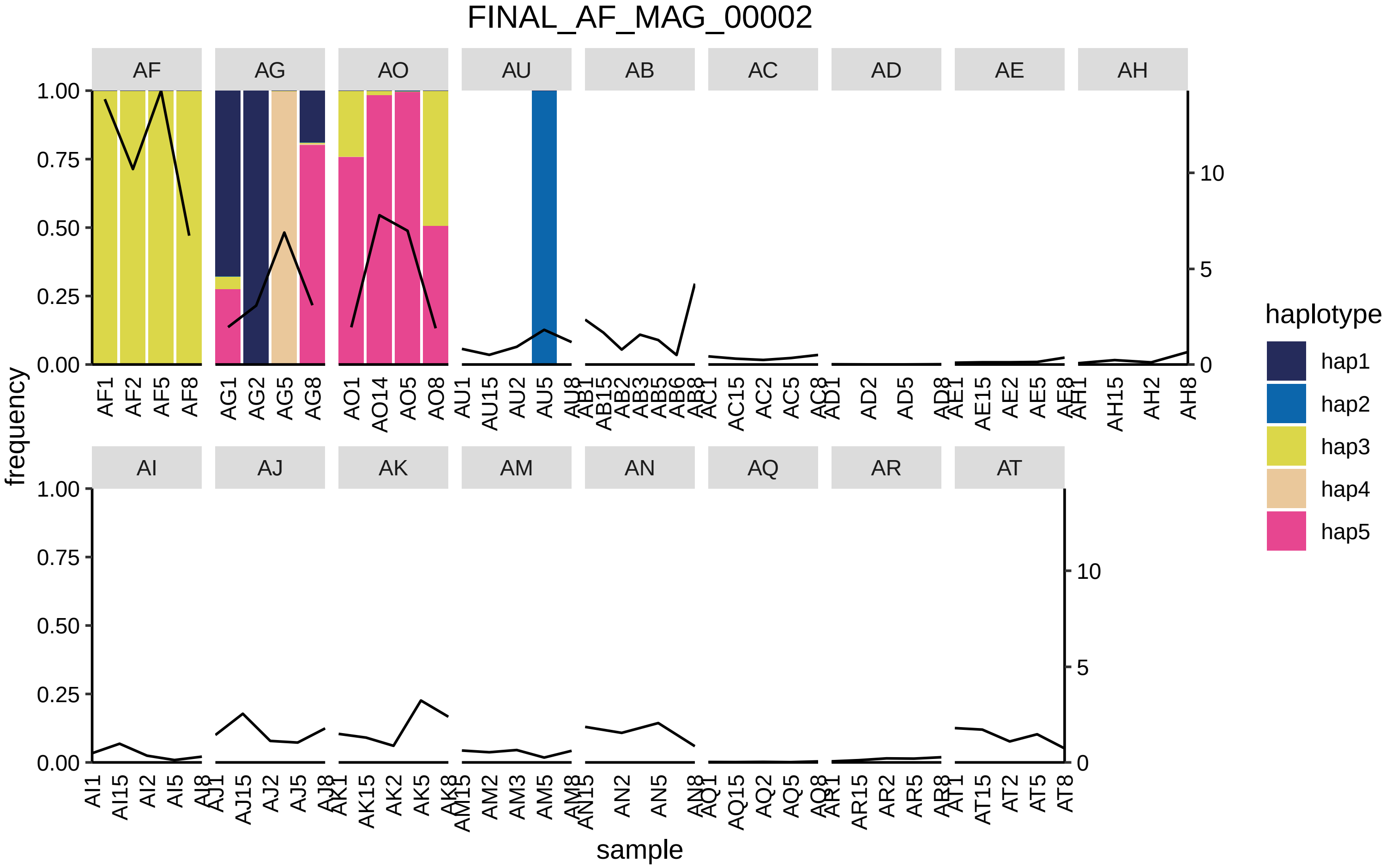
FINAL_AF_MAG_00002
AF
AG
AO
AU
AB
AC
AD
AE
AH
AI
AJ
AK
AM
AN
AQ
AR
AT
frequency
1.00
0.75
0.50
0.25
0.00
10
5
0
sample
AF1 AF2 AF5 AF8
AG1 AG2 AG5 AG8
AO1 AO14 AO5 AO8
AU1 AU15 AU2 AU5 AU8
AB1 AB15 AB2 AB3 AB5 AB6 AB8
AC1 AC15 AC2 AC5 AC8
AD1 AD2 AD5 AD8
AE1 AE15 AE2 AE5 AE8
AH1 AH15 AH2 AH8
AI1 AI15 AI2 AI5 AI8
AJ1 AJ15 AJ2 AJ5 AJ8
AK1 AK15 AK2 AK5 AK8
AM15 AM2 AM3 AM5 AM8
AN15 AN2 AN5 AN8
AQ1 AQ15 AQ2 AQ5 AQ8
AR1 AR15 AR2 AR5 AR8
AT1 AT15 AT2 AT5 AT8
haplotype
hap1
hap2
hap3
hap4
hap5

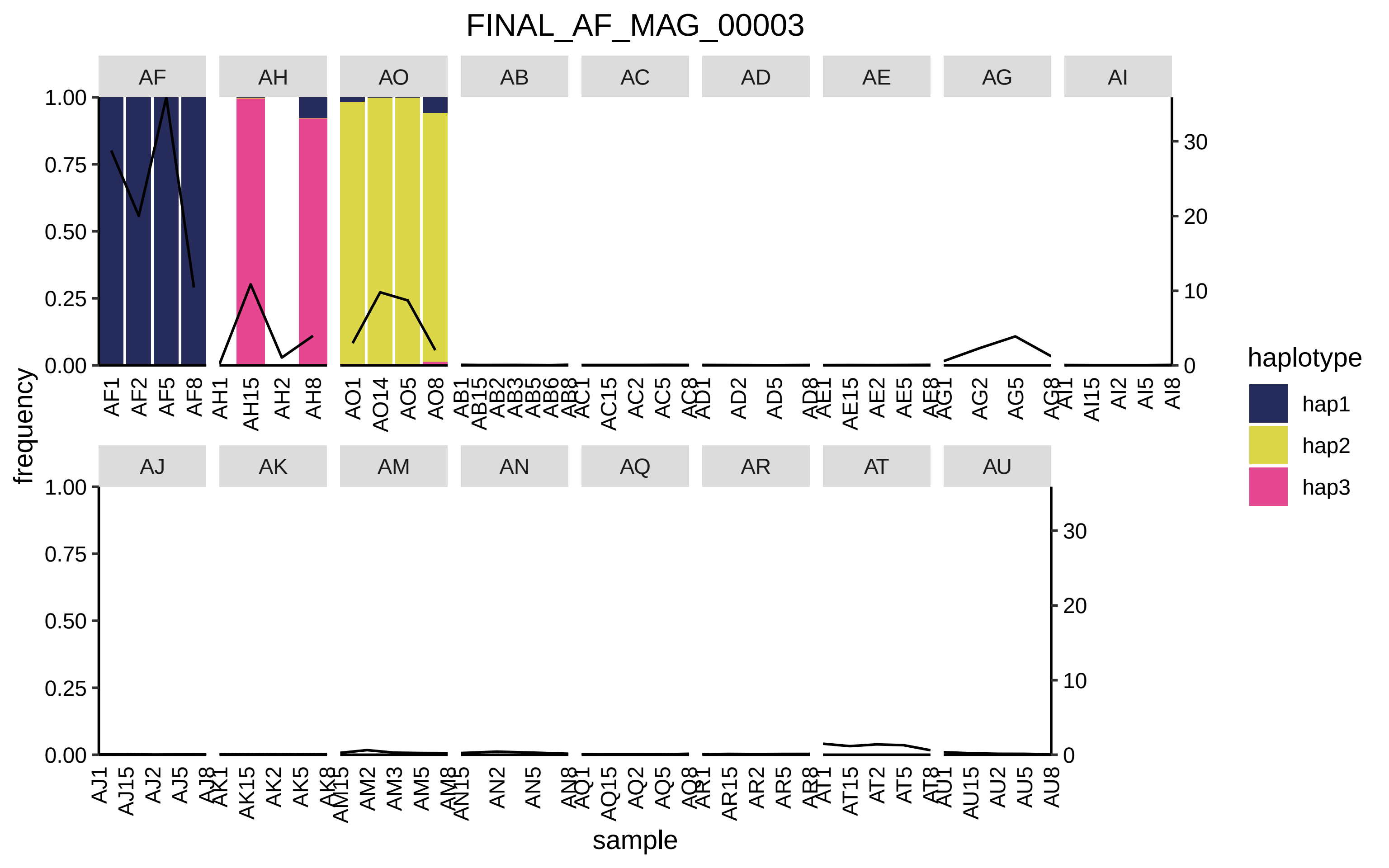
FINAL_AF_MAG_00003
AF
AH
AO
AB
AC
AD
AE
AG
AI
AJ
AK
AM
AN
AQ
AR
AT
AU
frequency
sample
haplotype
hap1
hap2
hap3
1.00
0.75
0.50
0.25
0.00
30
20
10
0

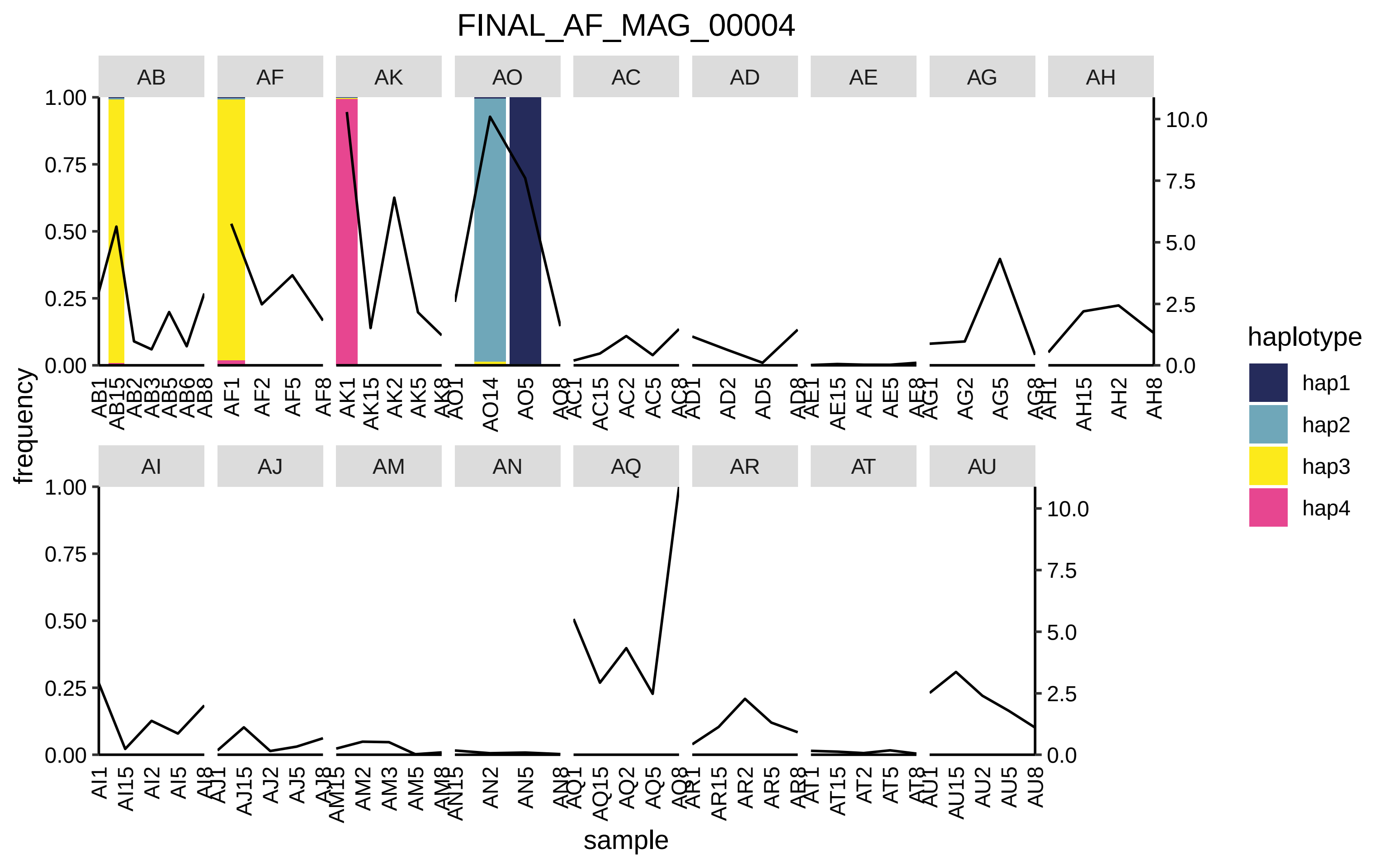

FINAL_AF_MAG_00004
AB
AF
AK
AO
AC
AD
AE
AG
AH
AI
AJ
AM
AN
AQ
AR
AT
AU
frequency
sample
haplotype
hap1
hap2
hap3
hap4
0.00
0.25
0.50
0.75
1.00
0.0
2.5
5.0
7.5
10.0

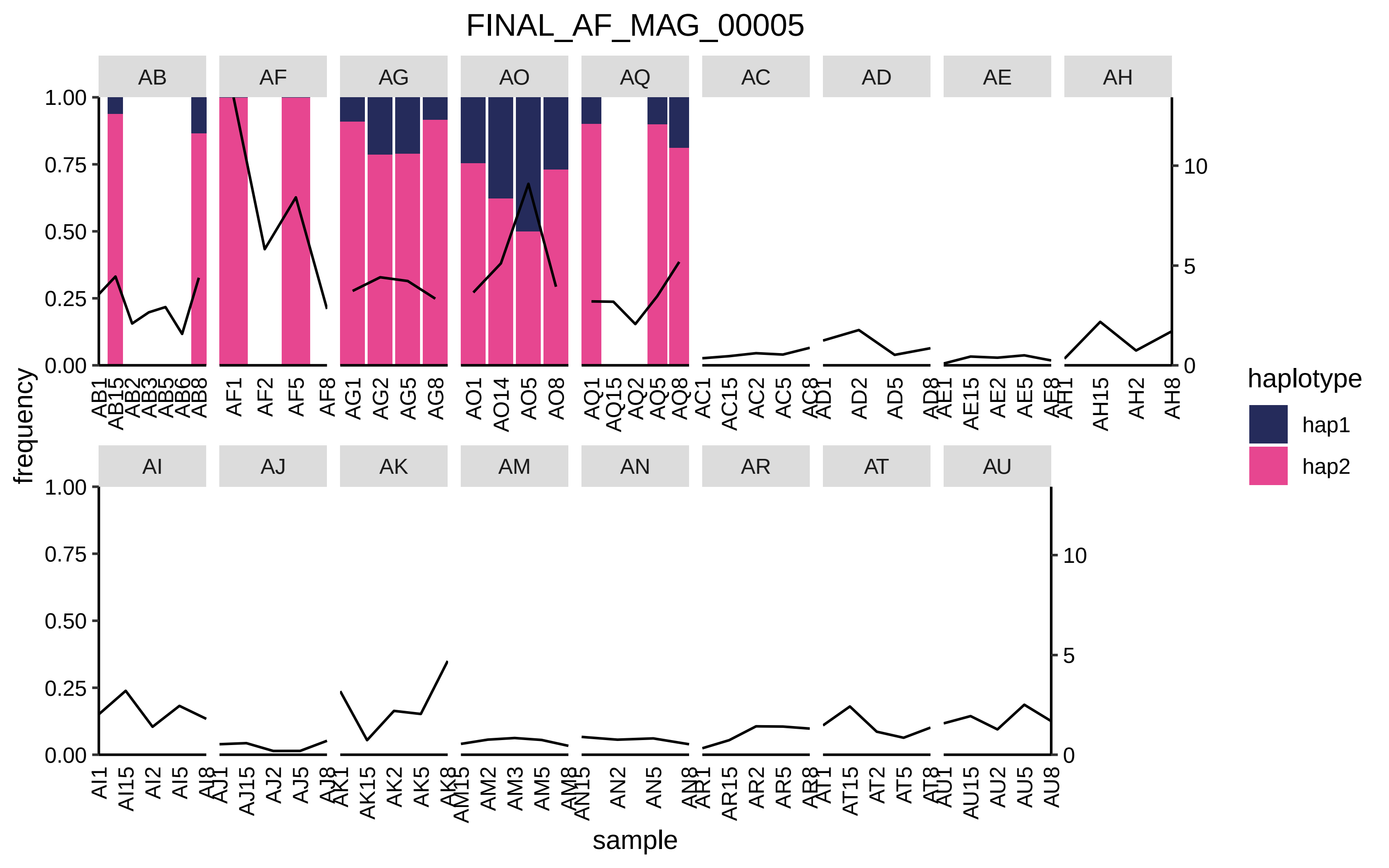

FINAL_AF_MAG_00005
AB
AF
AG
AO
AQ
AC
AD
AE
AH
AI
AJ
AK
AM
AN
AR
AT
AU
1.00
0.75
0.50
0.25
0.00
0
5
10
frequency
sample
haplotype
hap1
hap2
AB1 AB15 AB2 AB3 AB5 AB6 AB8
AF1 AF2 AF5 AF8
AG1 AG2 AG5 AG8
AO1 AO14 AO5 AO8
AQ1 AQ15 AQ2 AQ5 AQ8
AC1 AC15 AC2 AC5 AC8
AD1 AD2 AD5 AD8
AE1 AE15 AE2 AE5 AE8
AH1 AH15 AH2 AH8
AI1 AI15 AI2 AI5 AI8
AJ1 AJ15 AJ2 AJ5 AJ8
AK1 AK15 AK2 AK5 AK8
AM15 AM2 AM3 AM5 AM8
AN15 AN2 AN5 AN8
AR1 AR15 AR2 AR5 AR8
AT1 AT15 AT2 AT5 AT8
AU1 AU15 AU2 AU5 AU8

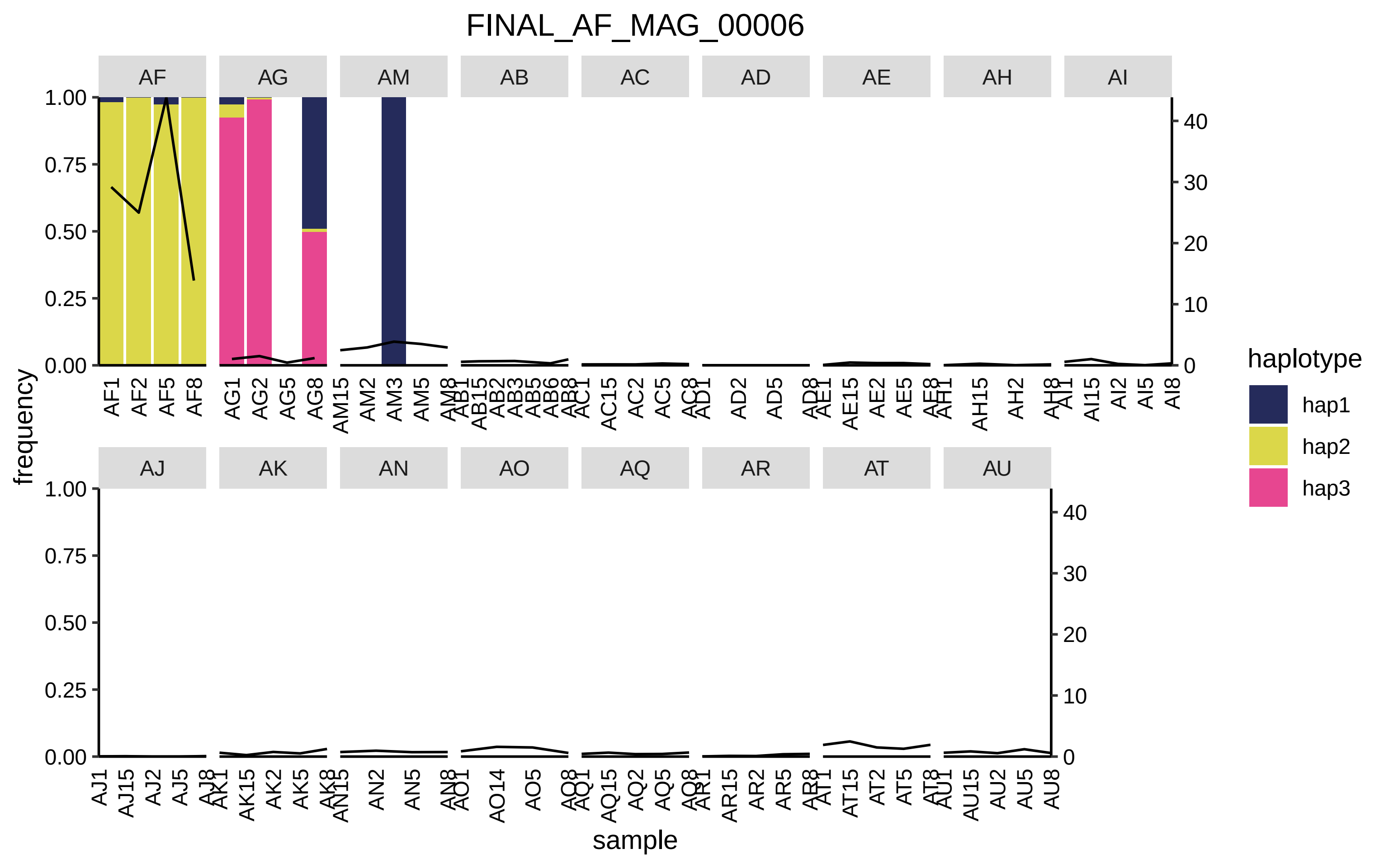

FINAL_AF_MAG_00006
AF
AG
AM
AB
AC
AD
AE
AH
AI
AJ
AK
AN
AO
AQ
AR
AT
AU
frequency
sample
haplotype
hap1
hap2
hap3
1.00
0.75
0.50
0.25
0.00
0
10
20
30
40
AF1
AF2
AF5
AF8
AG1
AG2
AG5
AG8
AM2
AM3
AM5
AD2
AD5
AH15
AH2
AI15
AI2
AI5
AI8
AO14
AO5
AN2
AN5
AU15
AU2
AU5
AU8

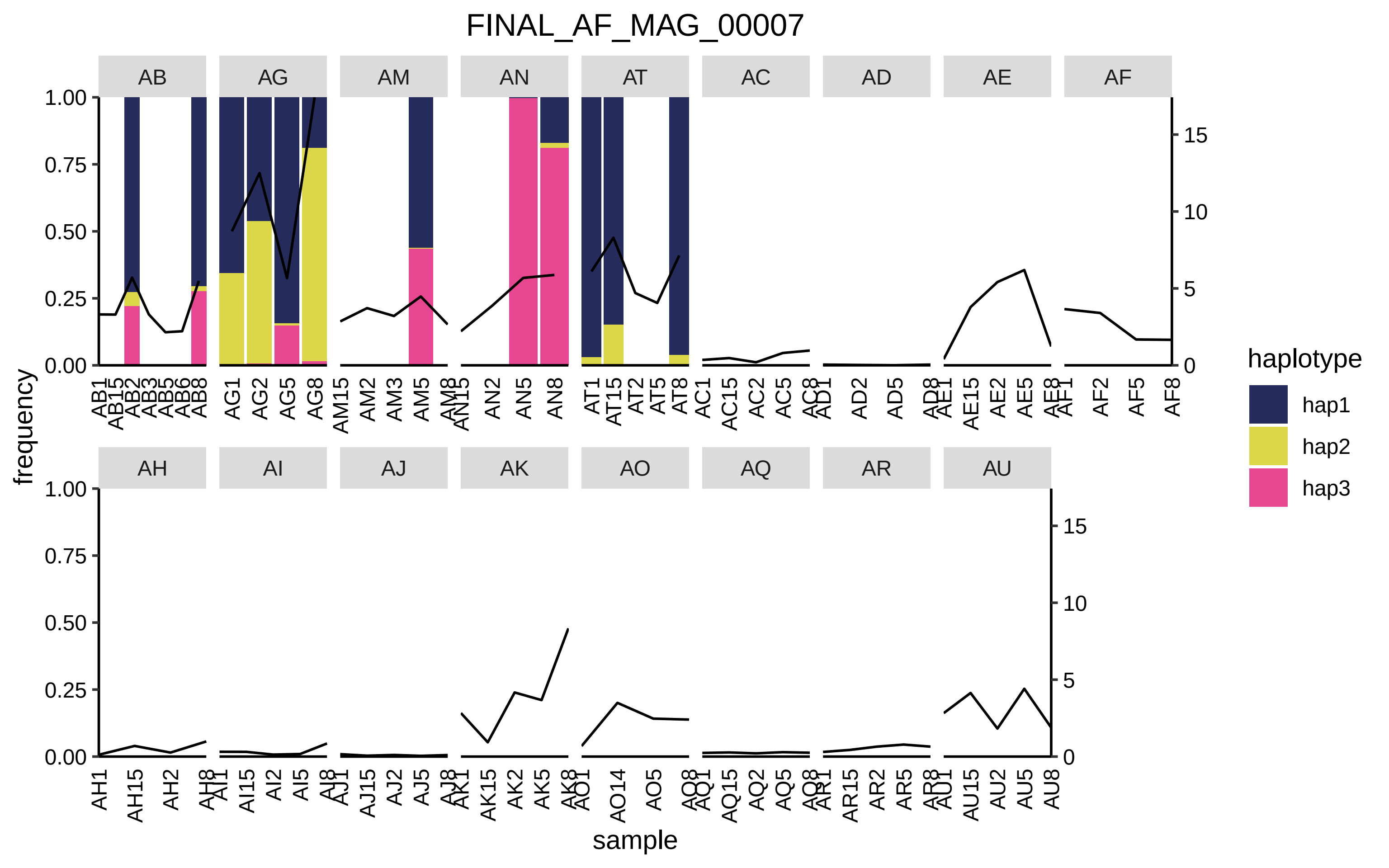

FINAL_AF_MAG_00007
AB
AG
AM
AN
AT
AC
AD
AE
AF
AH
AI
AJ
AK
AO
AQ
AR
AU
frequency
sample
1.00
0.75
0.50
0.25
0.00
0
5
10
15
AB1 AB15 AB2 AB3 AB5 AB6 AB8
AG1 AG2 AG5 AG8
AM15 AM2 AM3 AM5 AM8
AN15 AN2 AN5 AN8
AT1 AT15 AT2 AT5 AT8
AC1 AC15 AC2 AC5 AC8
AD1 AD2 AD5 AD8
AE1 AE15 AE2 AE5 AE8
AF1 AF2 AF5 AF8
AH1 AH15 AH2 AH8
AI1 AI15 AI2 AI5 AI8
AJ1 AJ15 AJ2 AJ5 AJ8
AK1 AK15 AK2 AK5 AK8
AO1 AO14 AO5 AO8
AQ1 AQ15 AQ2 AQ5 AQ8
AR1 AR15 AR2 AR5 AR8
AU1 AU15 AU2 AU5 AU8
haplotype
hap1
hap2
hap3

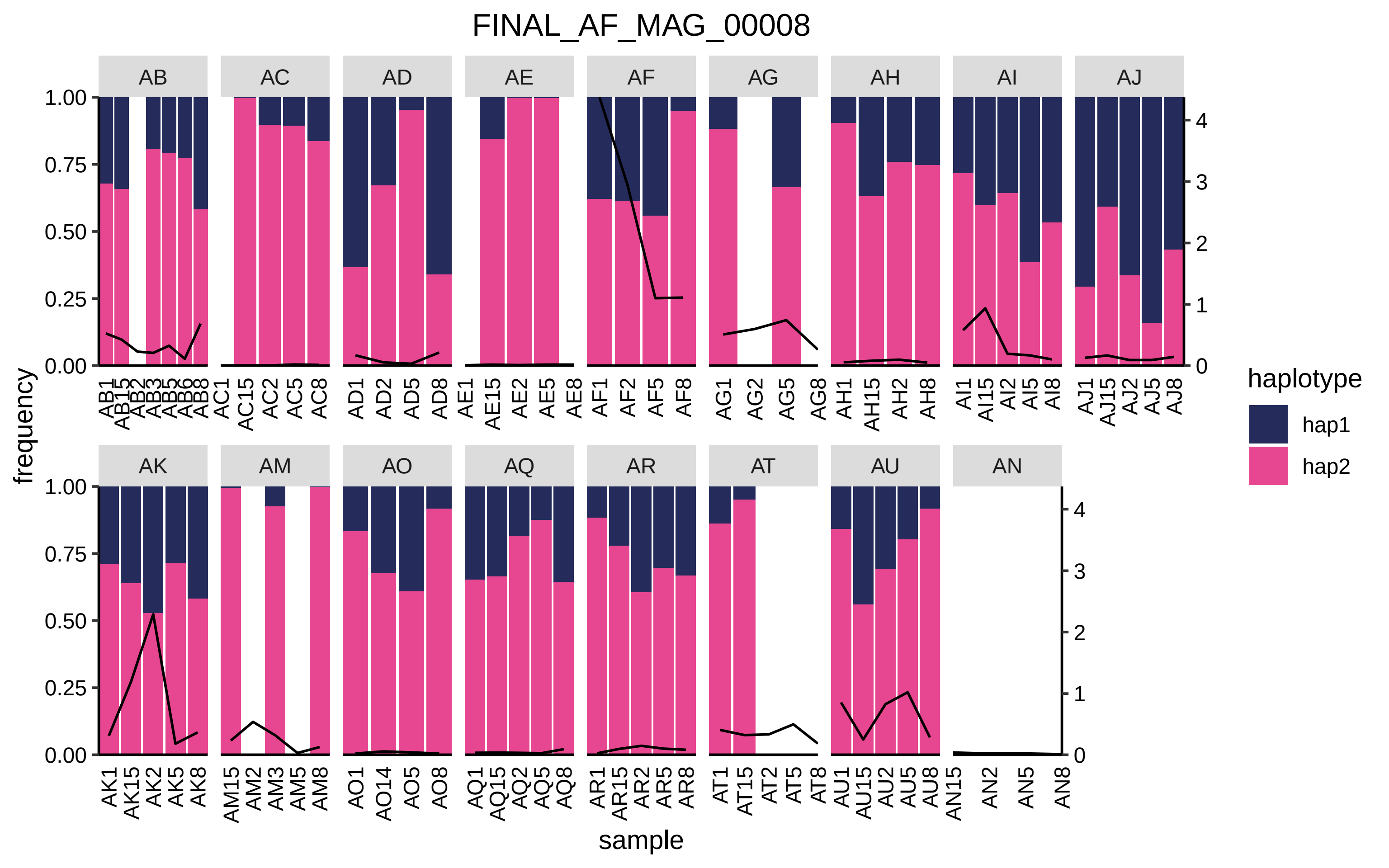

FINAL_AF_MAG_00008
AB
AC
AD
AE
AF
AG
AH
AI
AJ
AK
AM
AO
AQ
AR
AT
AU
AN
frequency
sample
haplotype
hap1
hap2

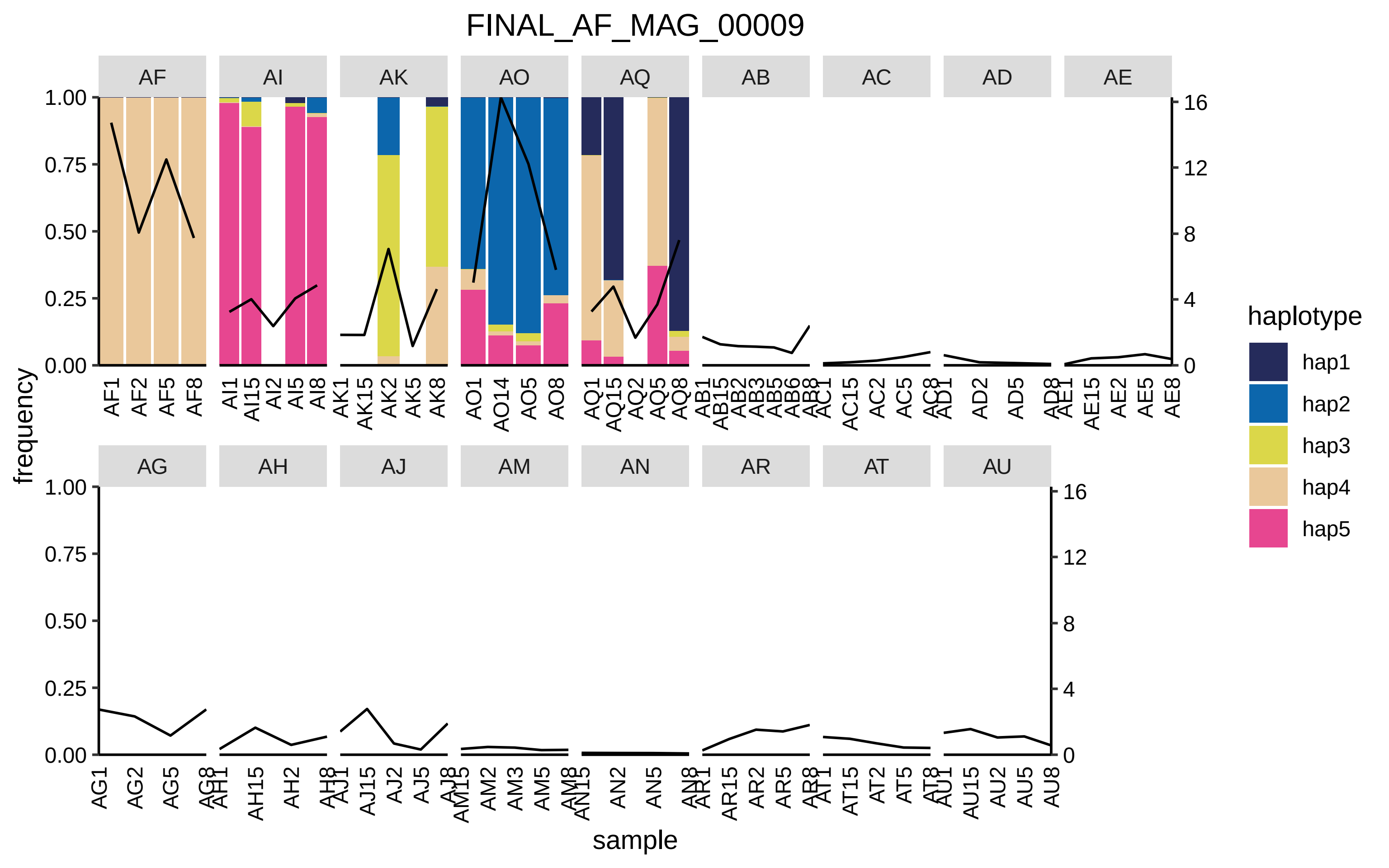

FINAL_AF_MAG_00009
AF
AI
AK
AO
AQ
AB
AC
AD
AE
AG
AH
AJ
AM
AN
AR
AT
AU
frequency
sample
haplotype
hap1
hap2
hap3
hap4
hap5

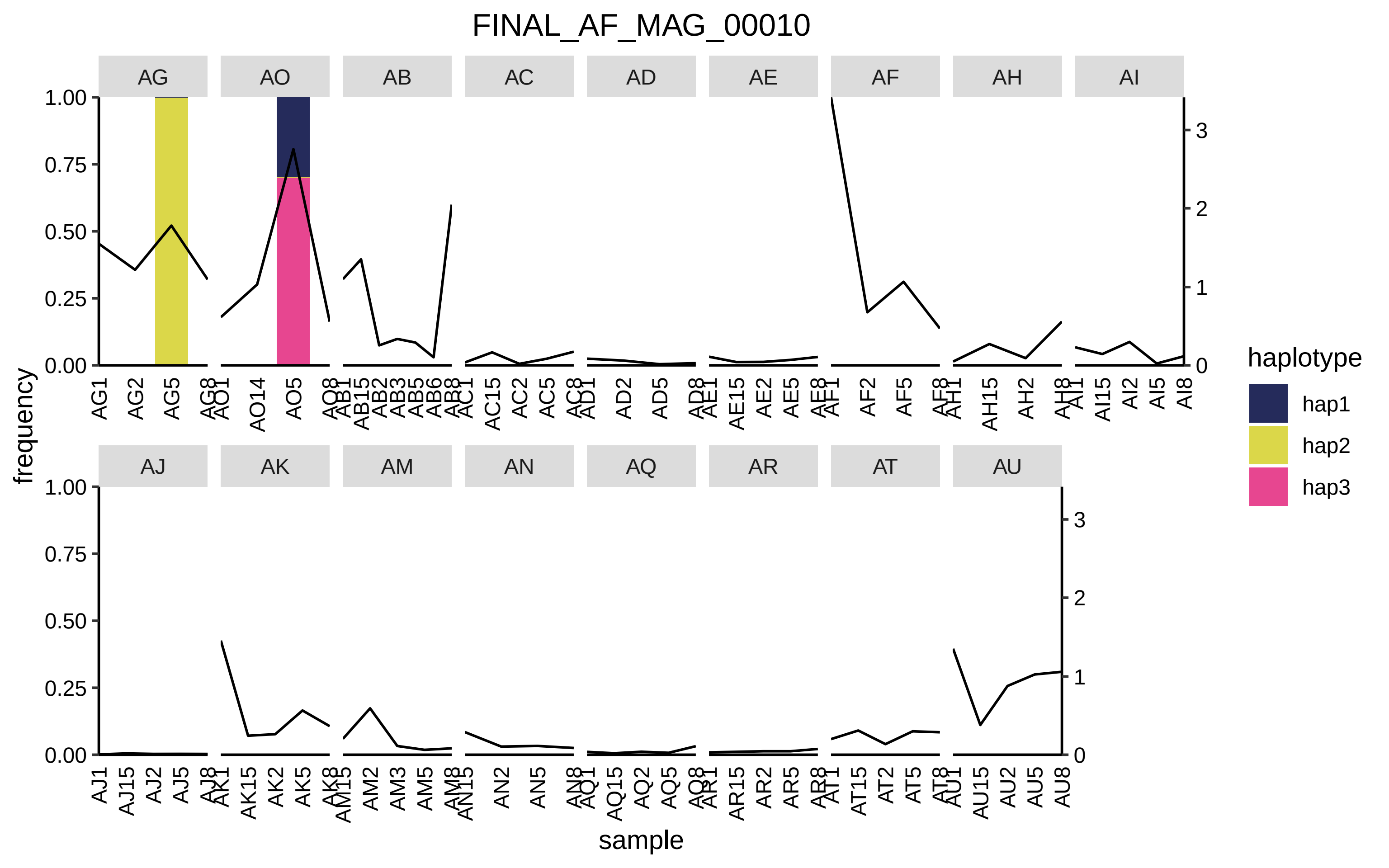

FINAL_AF_MAG_00010
AG
AO
AB
AC
AD
AE
AF
AH
AI
AJ
AK
AM
AN
AQ
AR
AT
AU
frequency
sample
haplotype
hap1
hap2
hap3
1.00
0.75
0.50
0.25
0.00
3
2
1
0

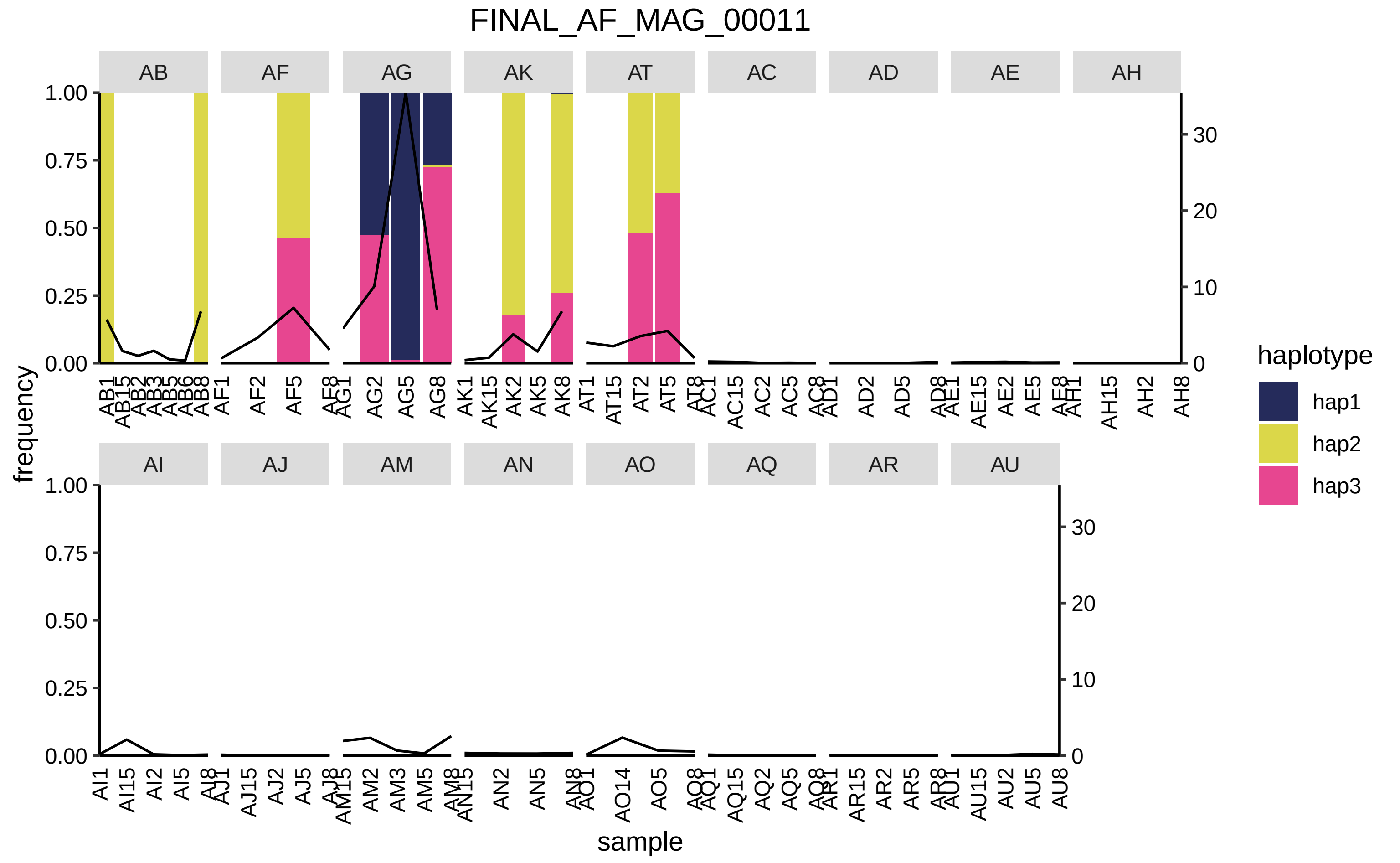

FINAL_AF_MAG_00011
AB
AF
AG
AK
AT
AC
AD
AE
AH
AI
AJ
AM
AN
AO
AQ
AR
AU
frequency
sample
haplotype
hap1
hap2
hap3

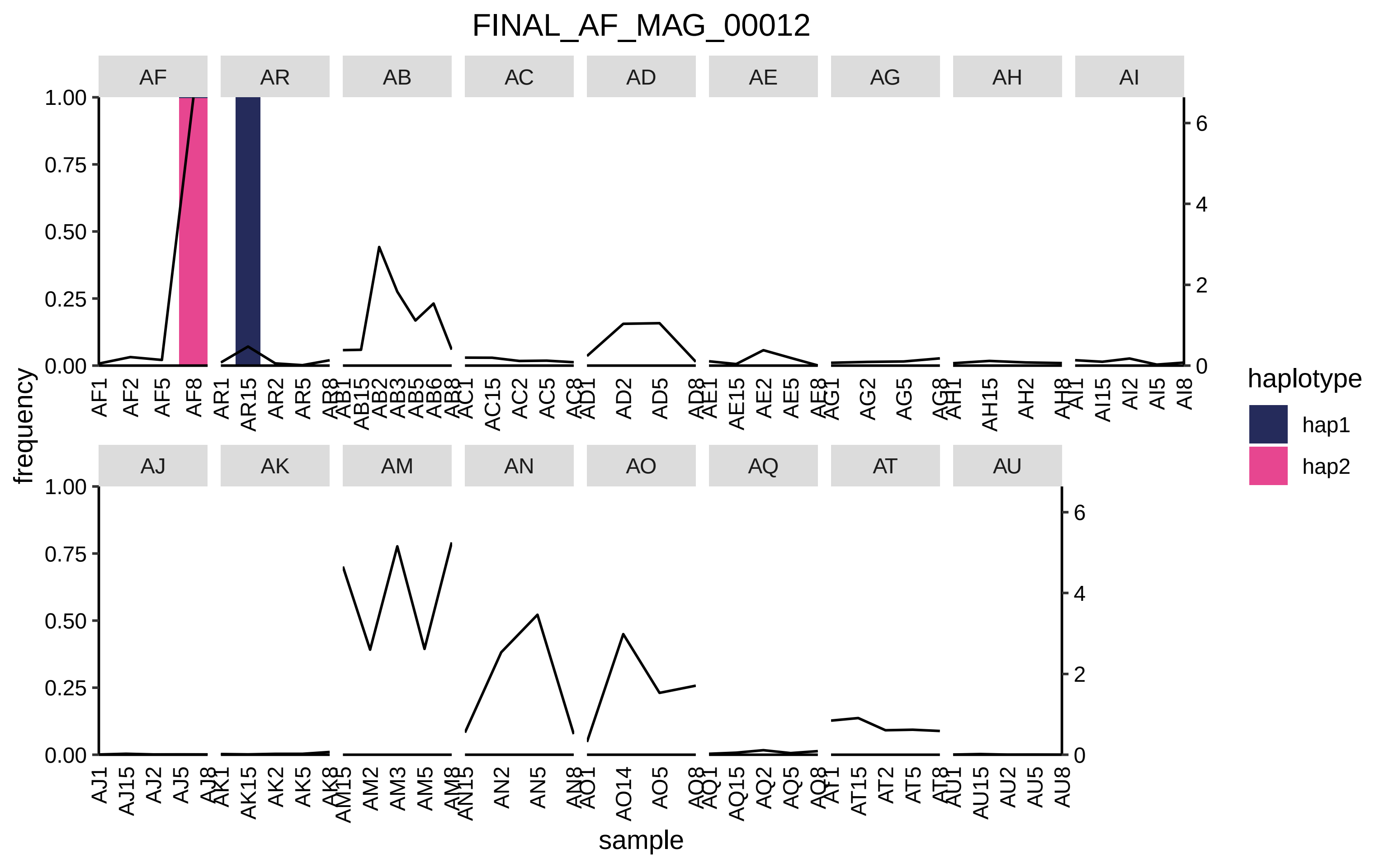
FINAL_AF_MAG_00012
AF
AR
AB
AC
AD
AE
AG
AH
AI
AJ
AK
AM
AN
AO
AQ
AT
AU
frequency
sample
haplotype
hap1
hap2
1.00
0.75
0.50
0.25
0.00
6
4
2
0

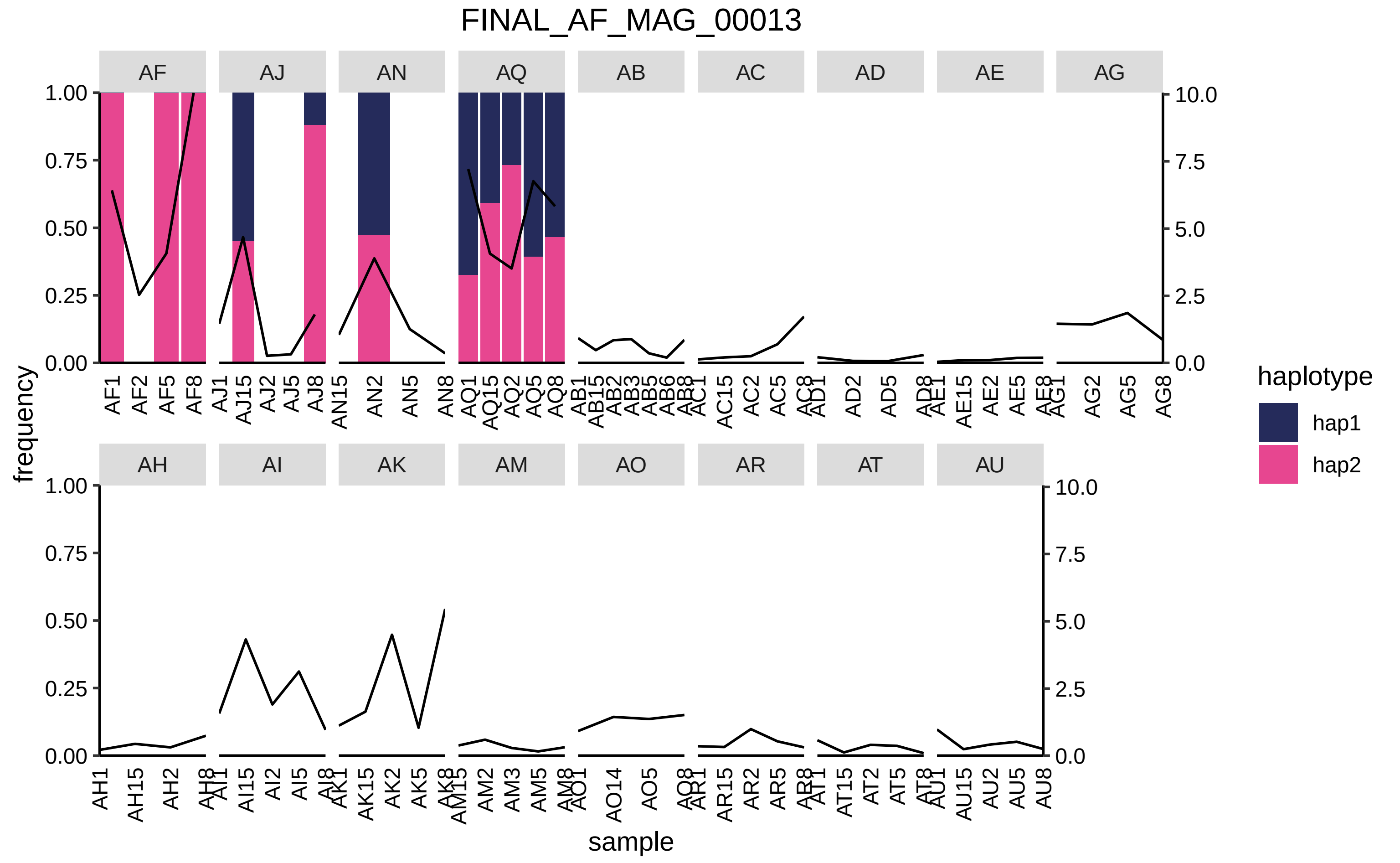

FINAL_AF_MAG_00013
AF
AJ
AN
AQ
AB
AC
AD
AE
AG
AH
AI
AK
AM
AO
AR
AT
AU
frequency
sample
haplotype
hap1
hap2
1.00
0.75
0.50
0.25
0.00
10.0
7.5
5.0
2.5
0.0
AF1
AF2
AF5
AF8
AJ1
AJ15
AJ2
AJ5
AJ8
AN15
AN2
AN5
AN8
AQ1
AQ15
AQ2
AQ5
AQ8
AB1
AB15
AB2
AB3
AB5
AB6
AB8
AC1
AC15
AC2
AC5
AC8
AD1
AD2
AD5
AD8
AE1
AE15
AE2
AE5
AE8
AG1
AG2
AG5
AG8
AH1
AH15
AH2
AH8
AI1
AI15
AI2
AI5
AI8
AK1
AK15
AK2
AK5
AK8
AM15
AM2
AM3
AM5
AM8
AO1
AO14
AO5
AO8
AR1
AR15
AR2
AR5
AR8
AT1
AT15
AT2
AT5
AT8
AU1
AU15
AU2
AU5
AU8

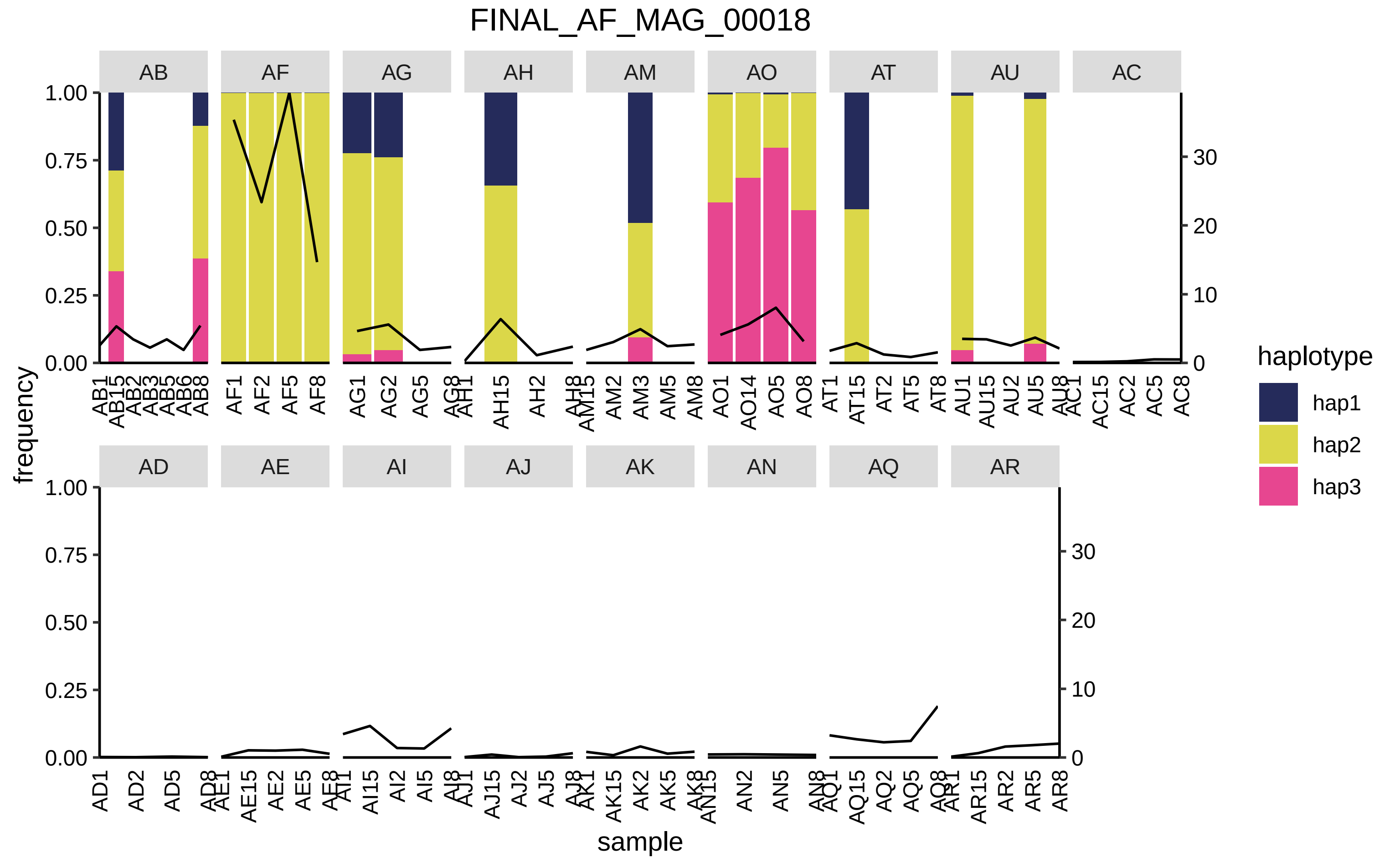
FINAL_AF_MAG_00018
AB
AF
AG
AH
AM
AO
AT
AU
AC
AD
AE
AI
AJ
AK
AN
AQ
AR
frequency
sample
haplotype
hap1
hap2
hap3

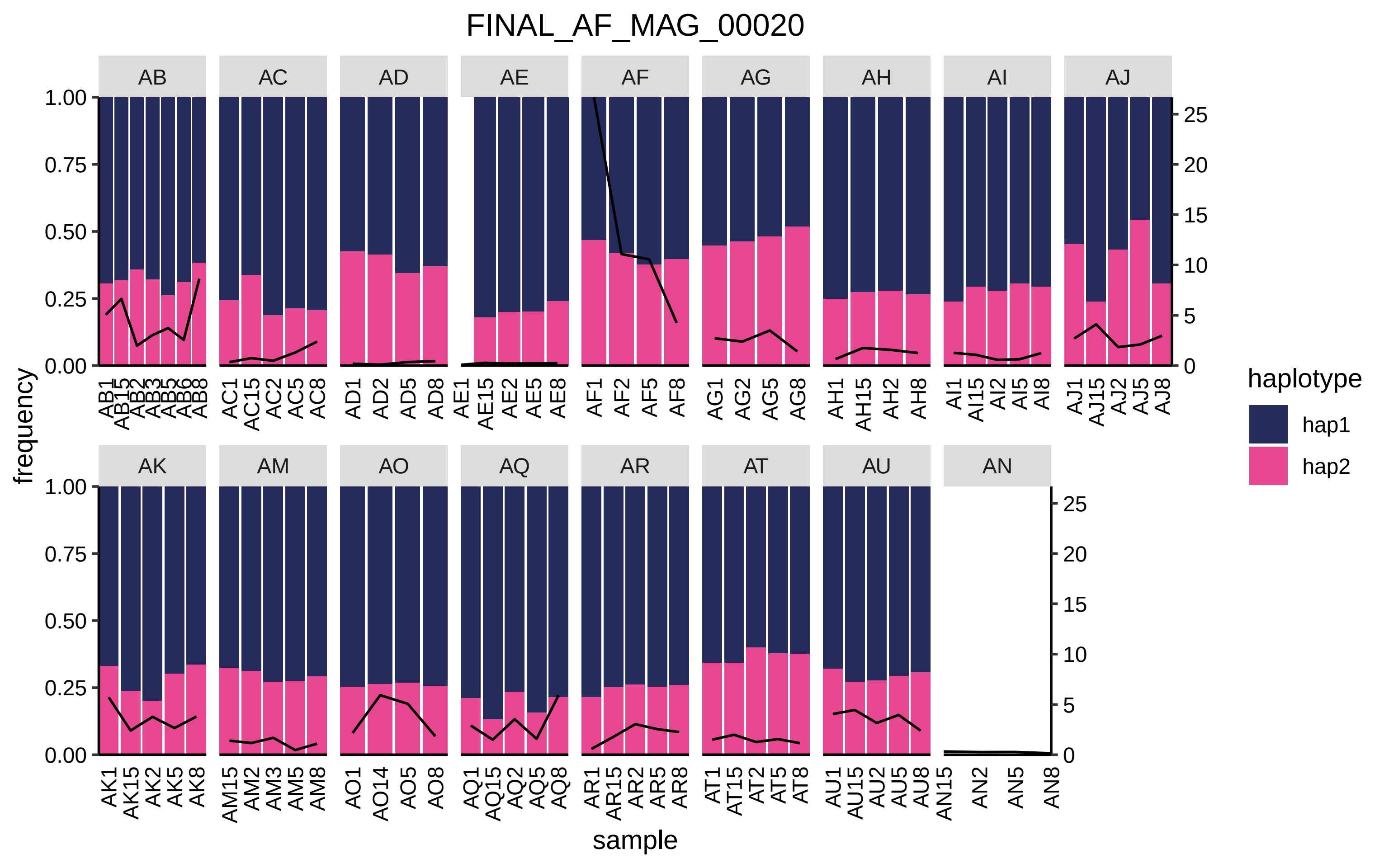

FINAL_AF_MAG_00020
AB
AC
AD
AE
AF
AG
AH
AI
AJ
AK
AM
AO
AQ
AR
AT
AU
AN
frequency
sample
haplotype
hap1
hap2
1.00
0.75
0.50
0.25
0.00
0
5
10
15
20
25
AB1 AB15 AB2 AB3 AB5 AB6 AB8
AC1 AC15 AC2 AC5 AC8
AD1 AD2 AD5 AD8
AE1 AE15 AE2 AE5 AE8
AF1 AF2 AF5 AF8
AG1 AG2 AG5 AG8
AH1 AH15 AH2 AH8
AI1 AI15 AI2 AI5 AI8
AJ1 AJ15 AJ2 AJ5 AJ8
AK1 AK15 AK2 AK5 AK8
AM15 AM2 AM3 AM5 AM8
AO1 AO14 AO5 AO8
AQ1 AQ15 AQ2 AQ5 AQ8
AR1 AR15 AR2 AR5 AR8
AT1 AT15 AT2 AT5 AT8
AU1 AU15 AU2 AU5 AU8
AN15 AN2 AN5 AN8

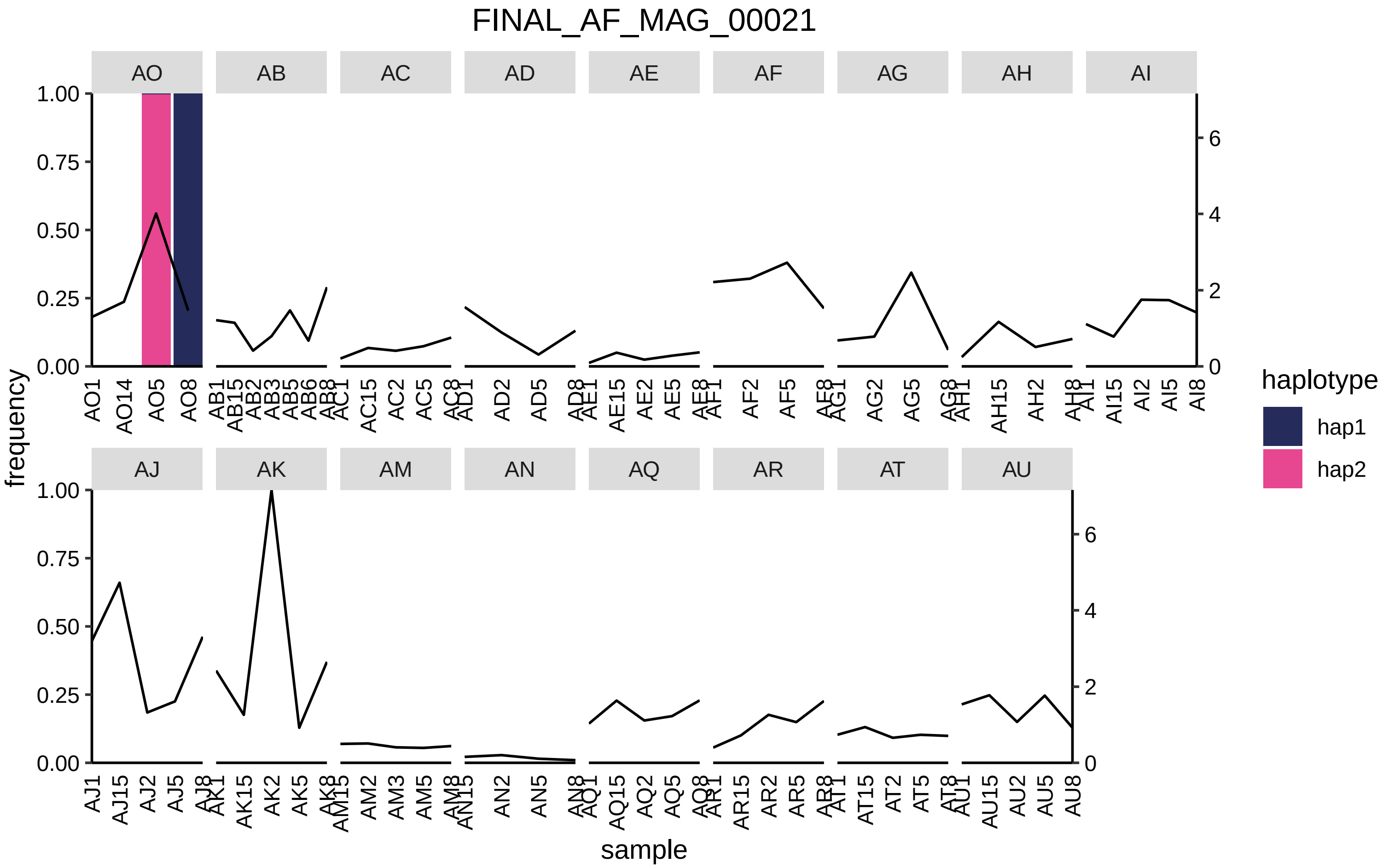

FINAL_AF_MAG_00021
AO
AB
AC
AD
AE
AF
AG
AH
AI
AJ
AK
AM
AN
AQ
AR
AT
AU
frequency
sample
haplotype
hap1
hap2

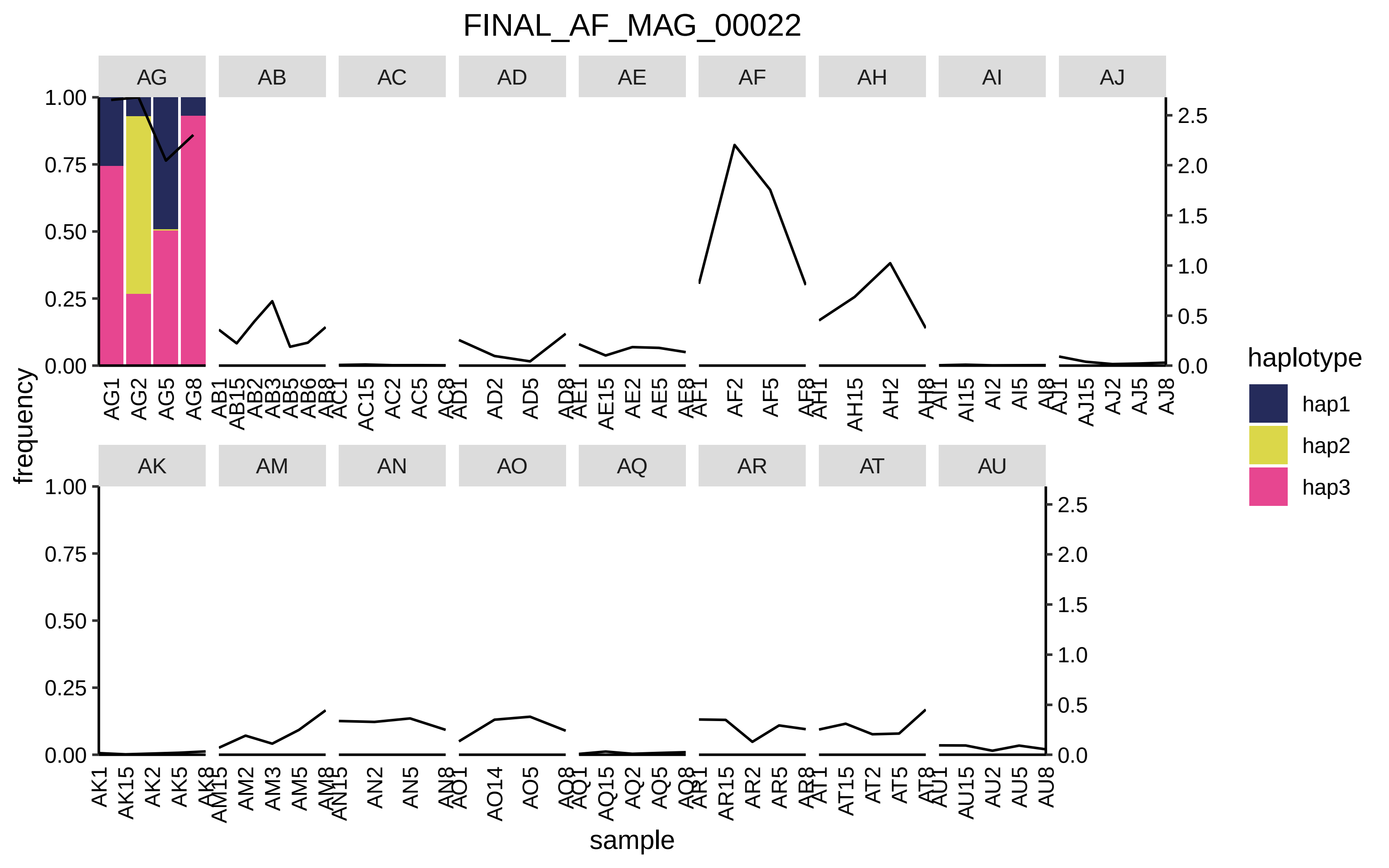

FINAL_AF_MAG_00022
AG
AB
AC
AD
AE
AF
AH
AI
AJ
AK
AM
AN
AO
AQ
AR
AT
AU
frequency
sample
1.00
0.75
0.50
0.25
0.00
2.5
2.0
1.5
1.0
0.5
0.0
haplotype
hap1
hap2
hap3

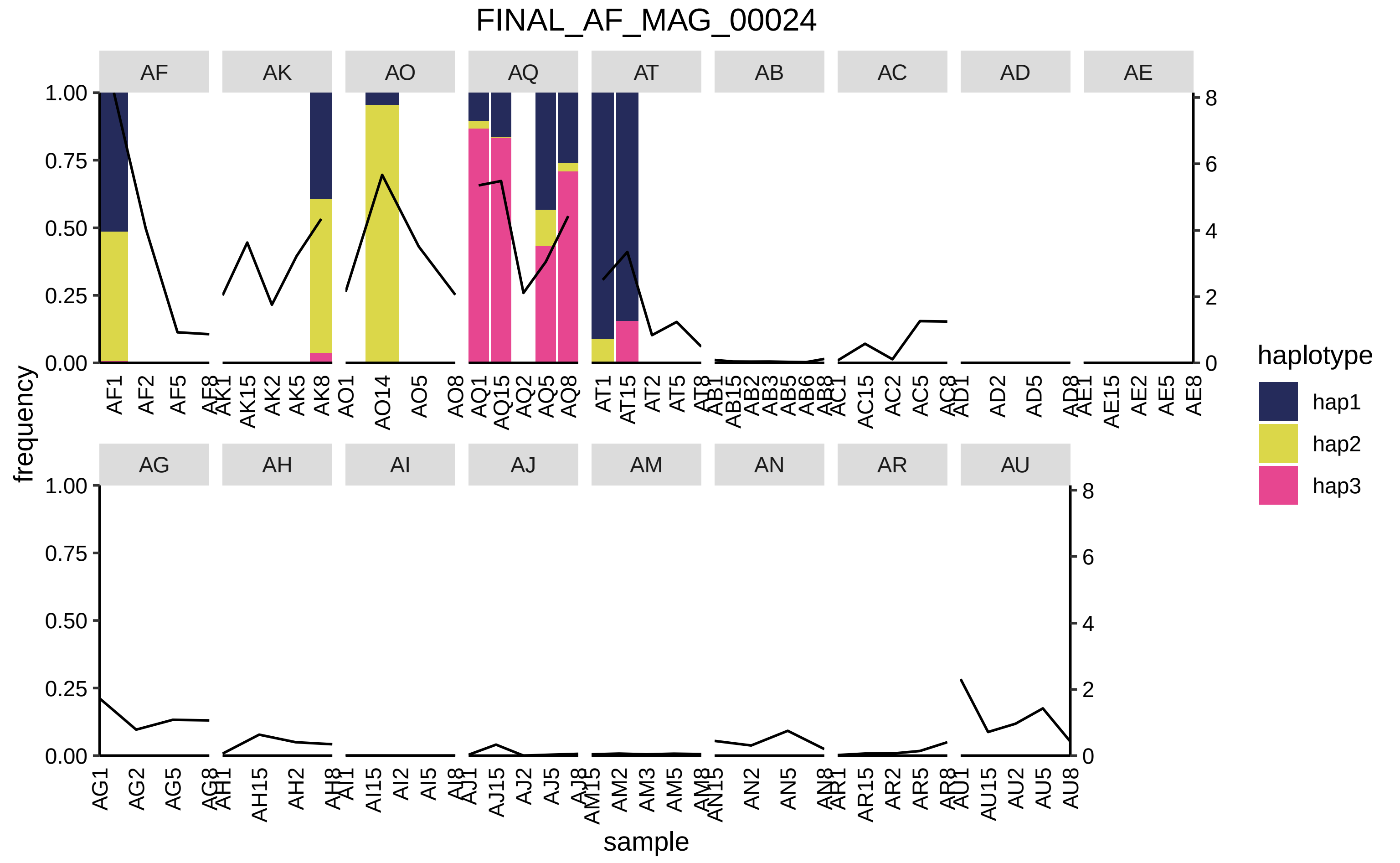

FINAL_AF_MAG_00024
AF
AK
AO
AQ
AT
AB
AC
AD
AE
AG
AH
AI
AJ
AM
AN
AR
AU
frequency
sample
haplotype
hap1
hap2
hap3
1.00
0.75
0.50
0.25
0.00
8
6
4
2
0
AF1 AF2 AF5 AF8 AK1 AK15 AK2 AK5 AK8 AO1 AO14 AO5 AO8 AQ1 AQ15 AQ2 AQ5 AQ8 AT1 AT15 AT2 AT5 AT8 AB1 AB15 AB2 AB3 AB5 AB6 AB8 AC1 AC15 AC2 AC5 AC8 AD1 AD2 AD5 AD8 AE1 AE15 AE2 AE5 AE8
AG1 AG2 AG5 AG8 AH1 AH15 AH2 AH8 AI1 AI15 AI2 AI5 AI8 AJ1 AJ15 AJ2 AJ5 AJ8 AM15 AM2 AM3 AM5 AM8 AN15 AN2 AN5 AN8 AR1 AR15 AR2 AR5 AR8 AU1 AU15 AU2 AU5 AU8

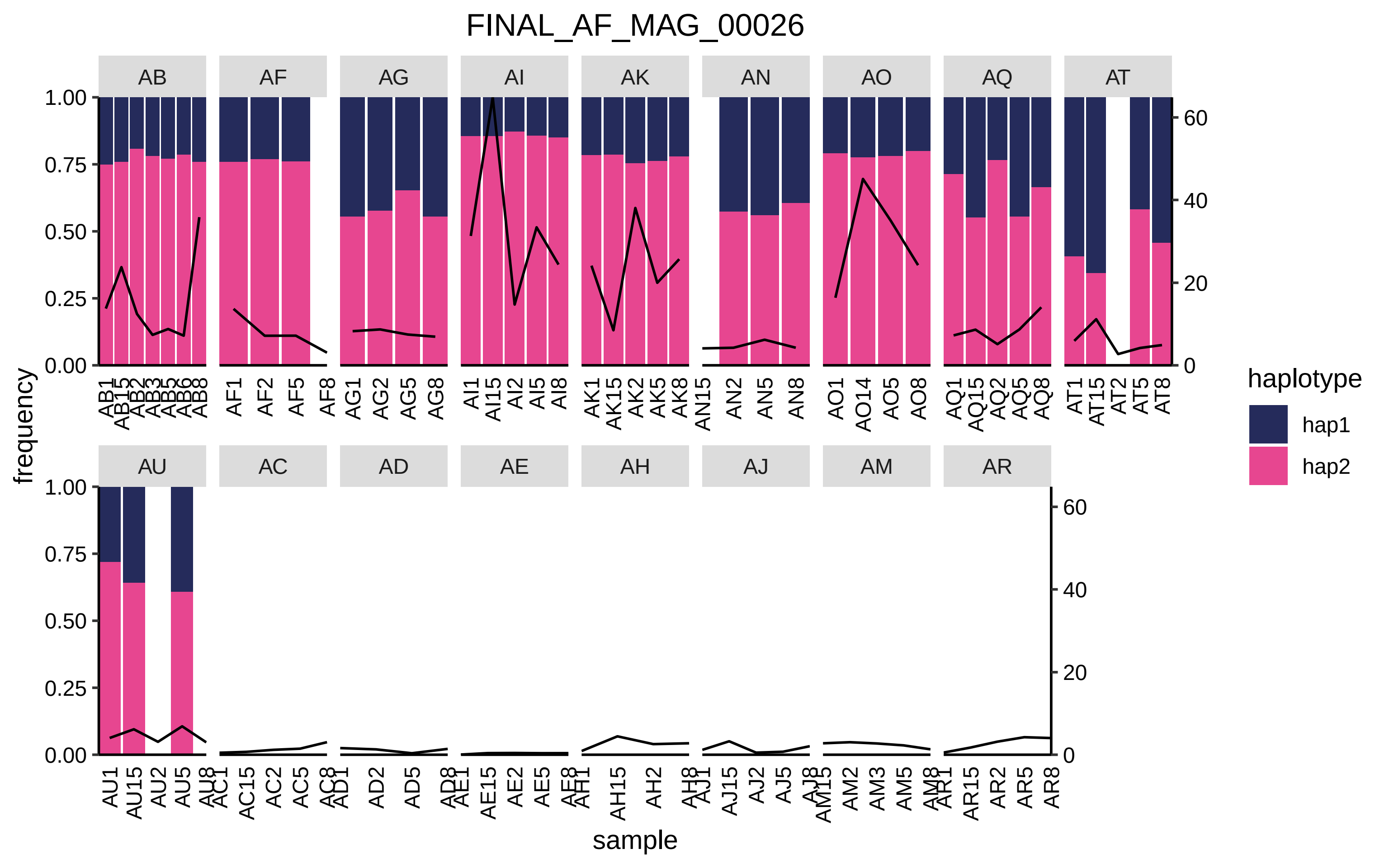
FINAL_AF_MAG_00026
AB
AF
AG
AI
AK
AN
AO
AQ
AT
AU
AC
AD
AE
AH
AJ
AM
AR
frequency
sample
haplotype
hap1
hap2
1.00
0.75
0.50
0.25
0.00
60
40
20
0

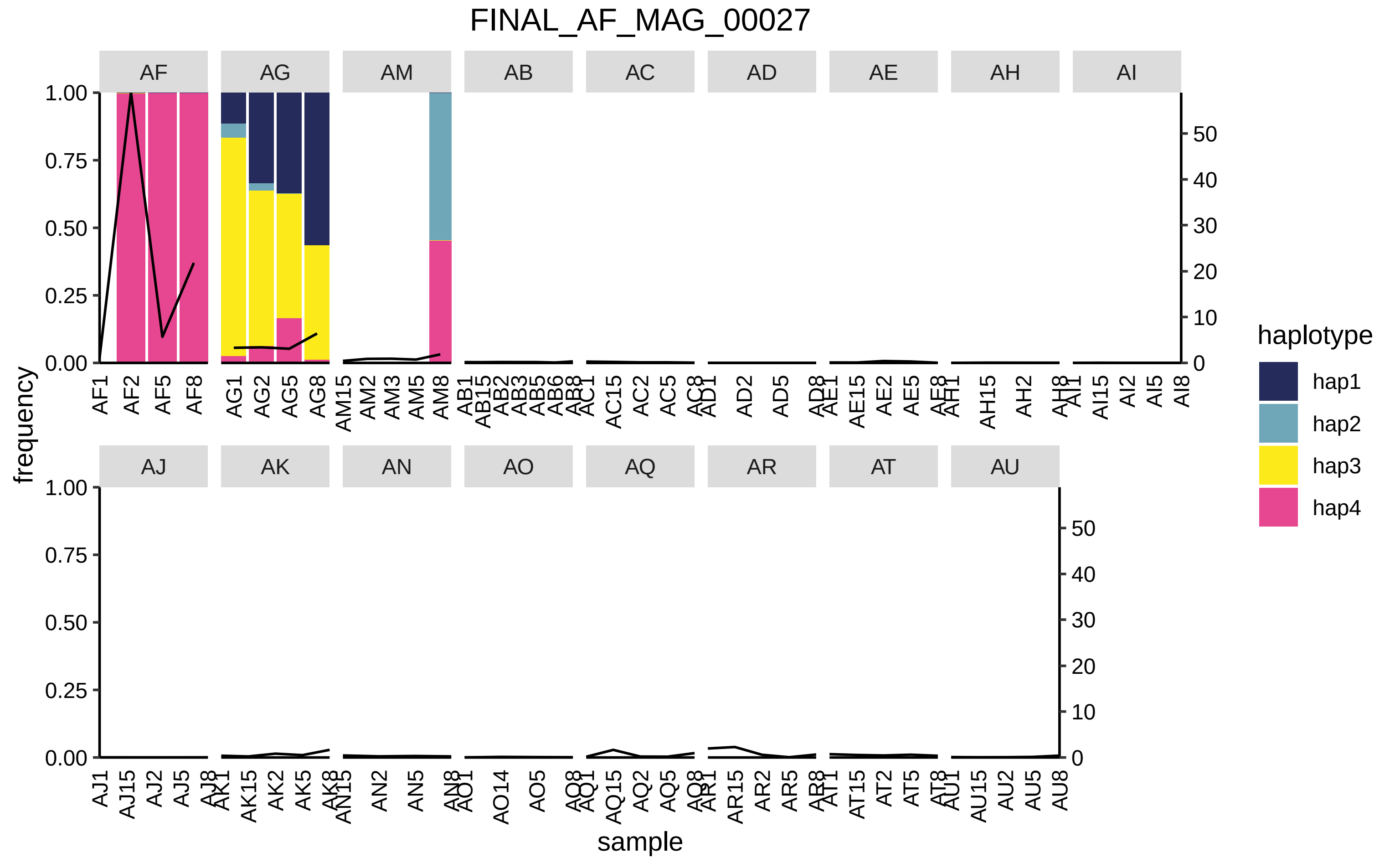

FINAL_AF_MAG_00027
AF
AG
AM
AB
AC
AD
AE
AH
AI
AJ
AK
AN
AO
AQ
AR
AT
AU
frequency
sample
haplotype
hap1
hap2
hap3
hap4

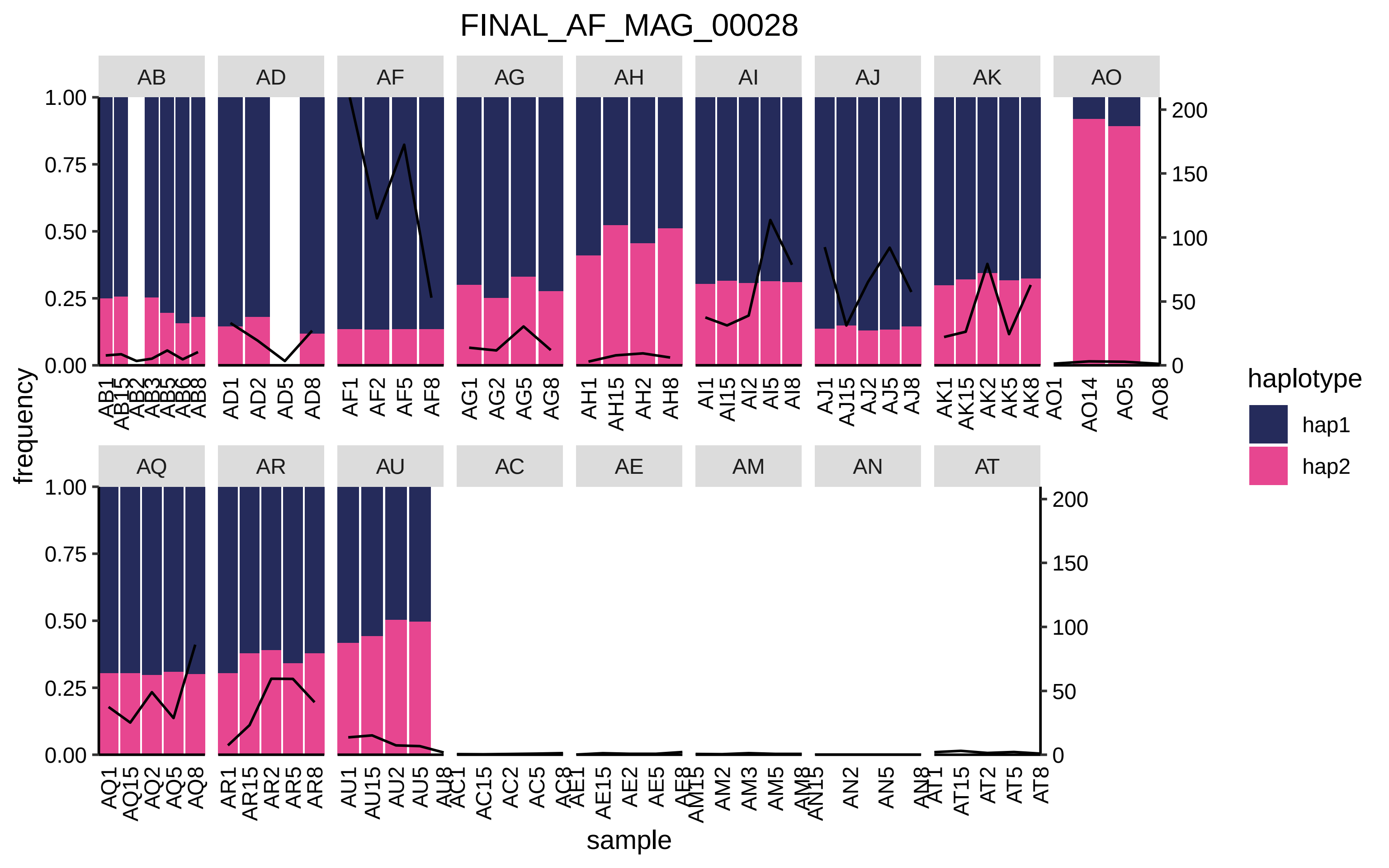
FINAL_AF_MAG_00028
AB
AD
AF
AG
AH
AI
AJ
AK
AO
AQ
AR
AU
AC
AE
AM
AN
AT
frequency
sample
haplotype
hap1
hap2
1.00
0.75
0.50
0.25
0.00
200
150
100
50
0

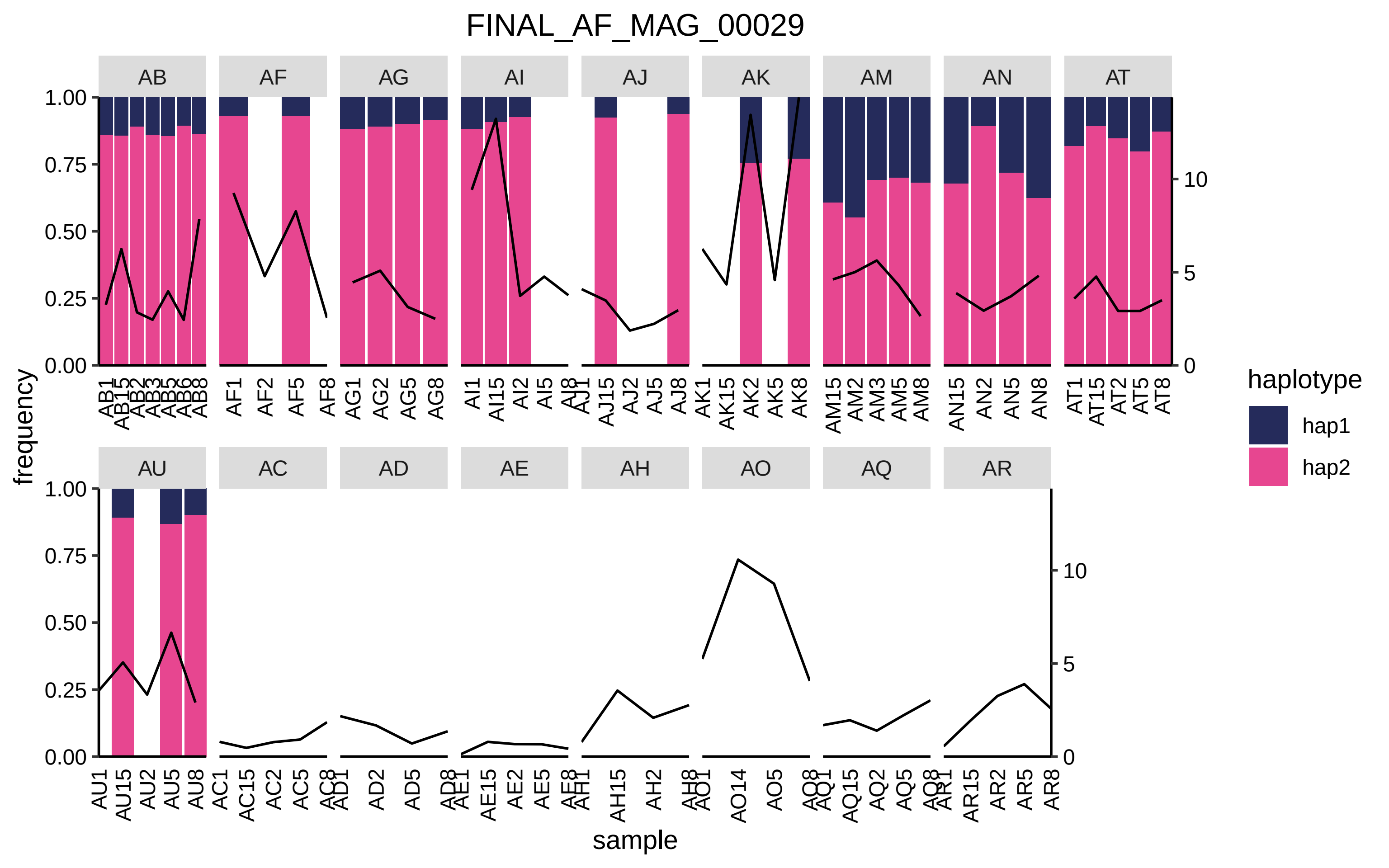

FINAL_AF_MAG_00029
AB
AF
AG
AI
AJ
AK
AM
AN
AT
AU
AC
AD
AE
AH
AO
AQ
AR
frequency
sample
haplotype
hap1
hap2

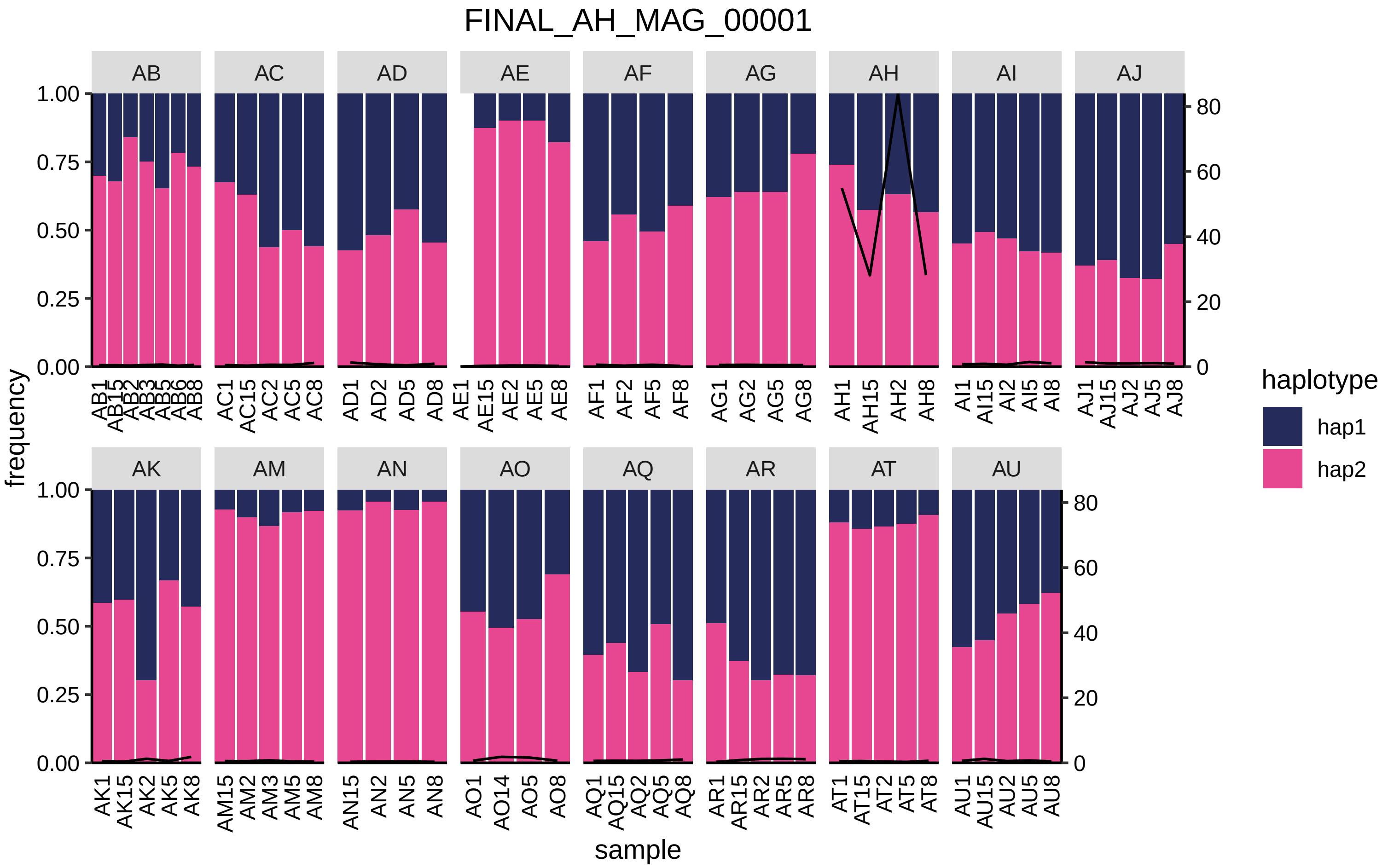

FINAL_AH_MAG_00001
AB
AC
AD
AE
AF
AG
AH
AI
AJ
AK
AM
AN
AO
AQ
AR
AT
AU
frequency
sample
haplotype
hap1
hap2
1.00
0.75
0.50
0.25
0.00
80
60
40
20
0
AB1 AB15 AB2 AB3 AB5 AB6 AB8
AC1 AC15 AC2 AC5 AC8
AD1 AD2 AD5 AD8
AE1 AE15 AE2 AE5 AE8
AF1 AF2 AF5 AF8
AG1 AG2 AG5 AG8
AH1 AH15 AH2 AH8
AI1 AI15 AI2 AI5 AI8
AJ1 AJ15 AJ2 AJ5 AJ8
AK1 AK15 AK2 AK5 AK8
AM15 AM2 AM3 AM5 AM8
AN15 AN2 AN5 AN8
AO1 AO14 AO5 AO8
AQ1 AQ15 AQ2 AQ5 AQ8
AR1 AR15 AR2 AR5 AR8
AT1 AT15 AT2 AT5 AT8
AU1 AU15 AU2 AU5 AU8

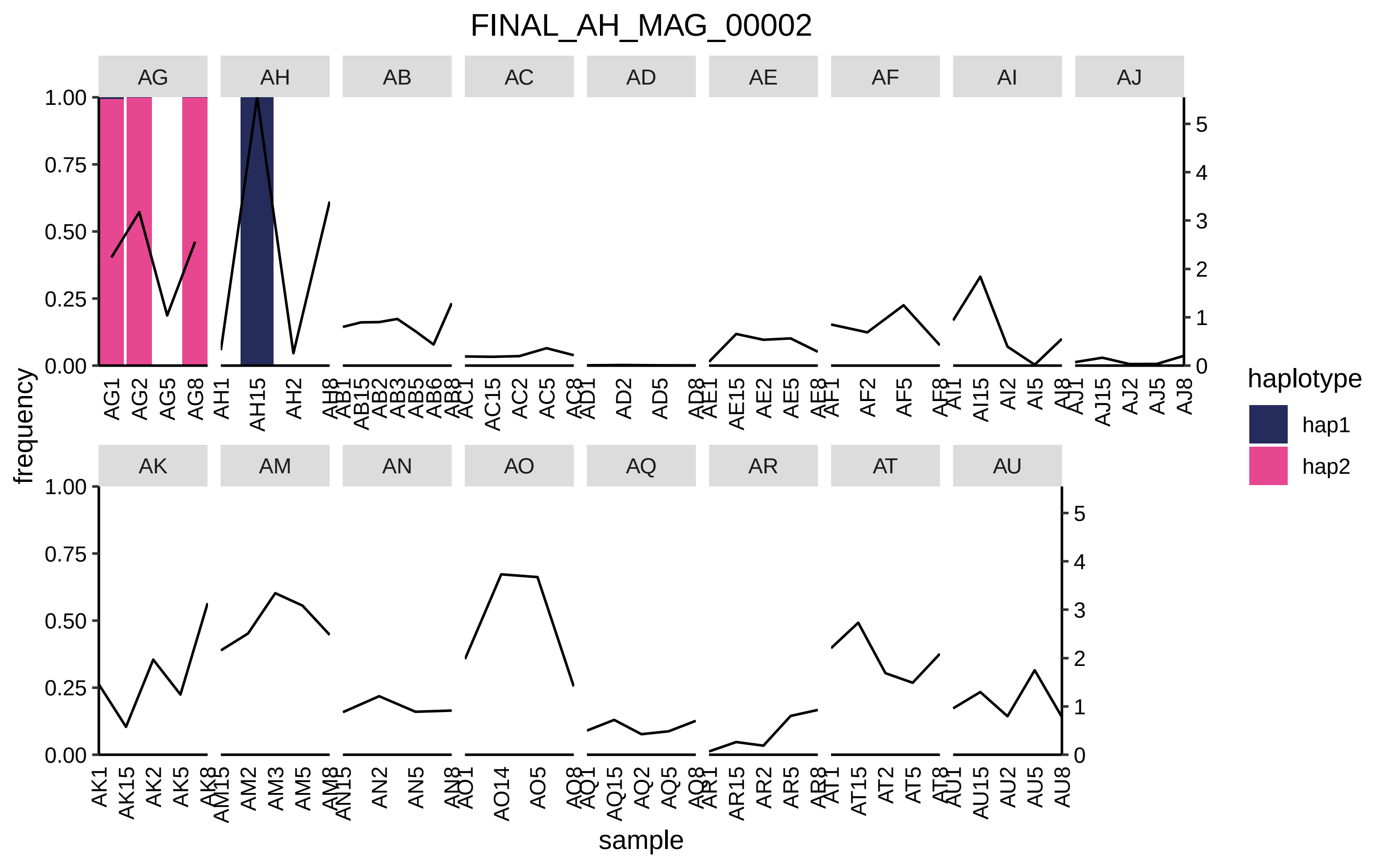

FINAL_AH_MAG_00002
AG
AH
AB
AC
AD
AE
AF
AI
AJ
AK
AM
AN
AO
AQ
AR
AT
AU
frequency
sample
haplotype
hap1
hap2
1.00
0.75
0.50
0.25
0.00
0
1
2
3
4
5

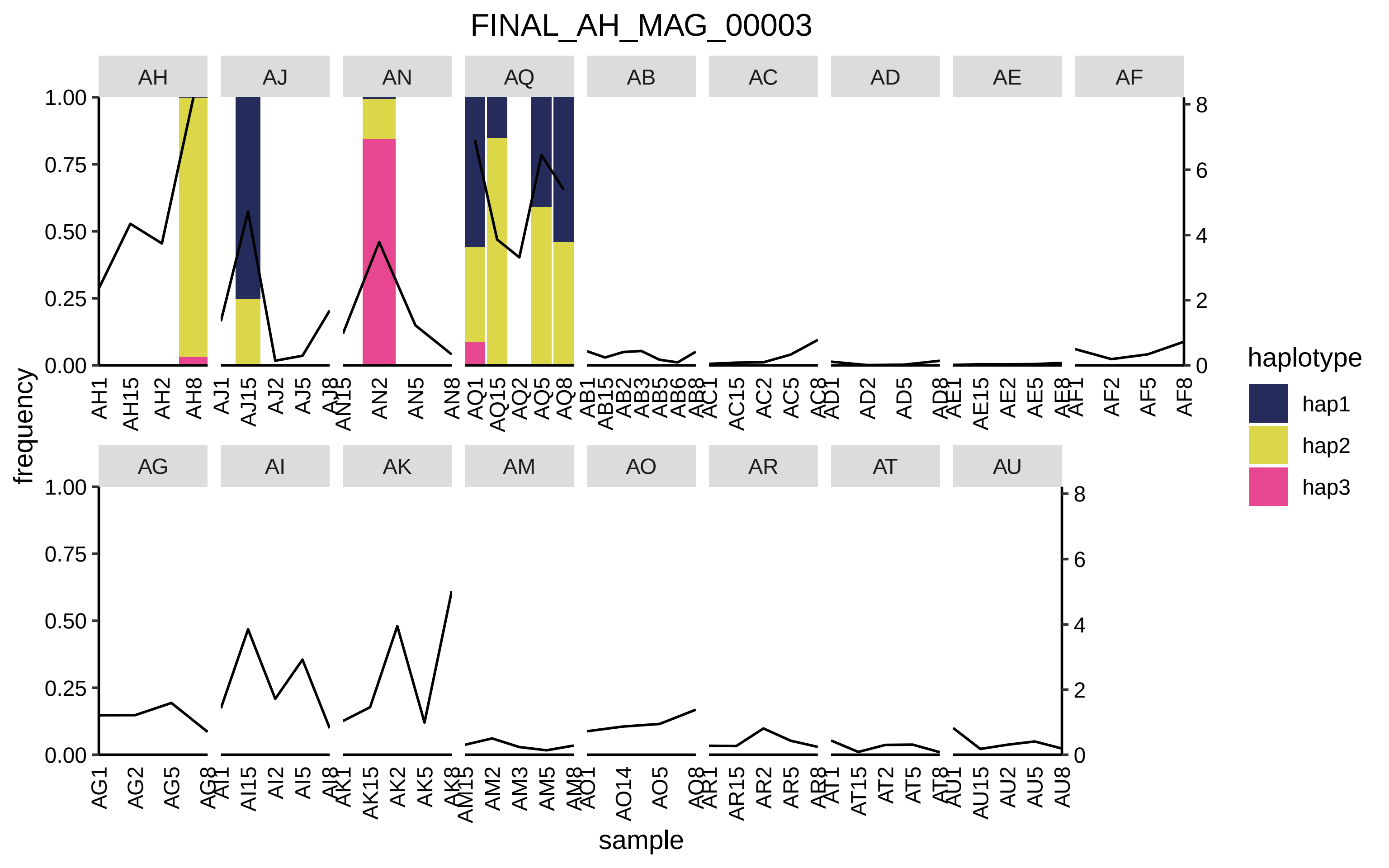

FINAL_AH_MAG_00003
AH
AJ
AN
AQ
AB
AC
AD
AE
AF
AG
AI
AK
AM
AO
AR
AT
AU
frequency
sample
haplotype
hap1
hap2
hap3

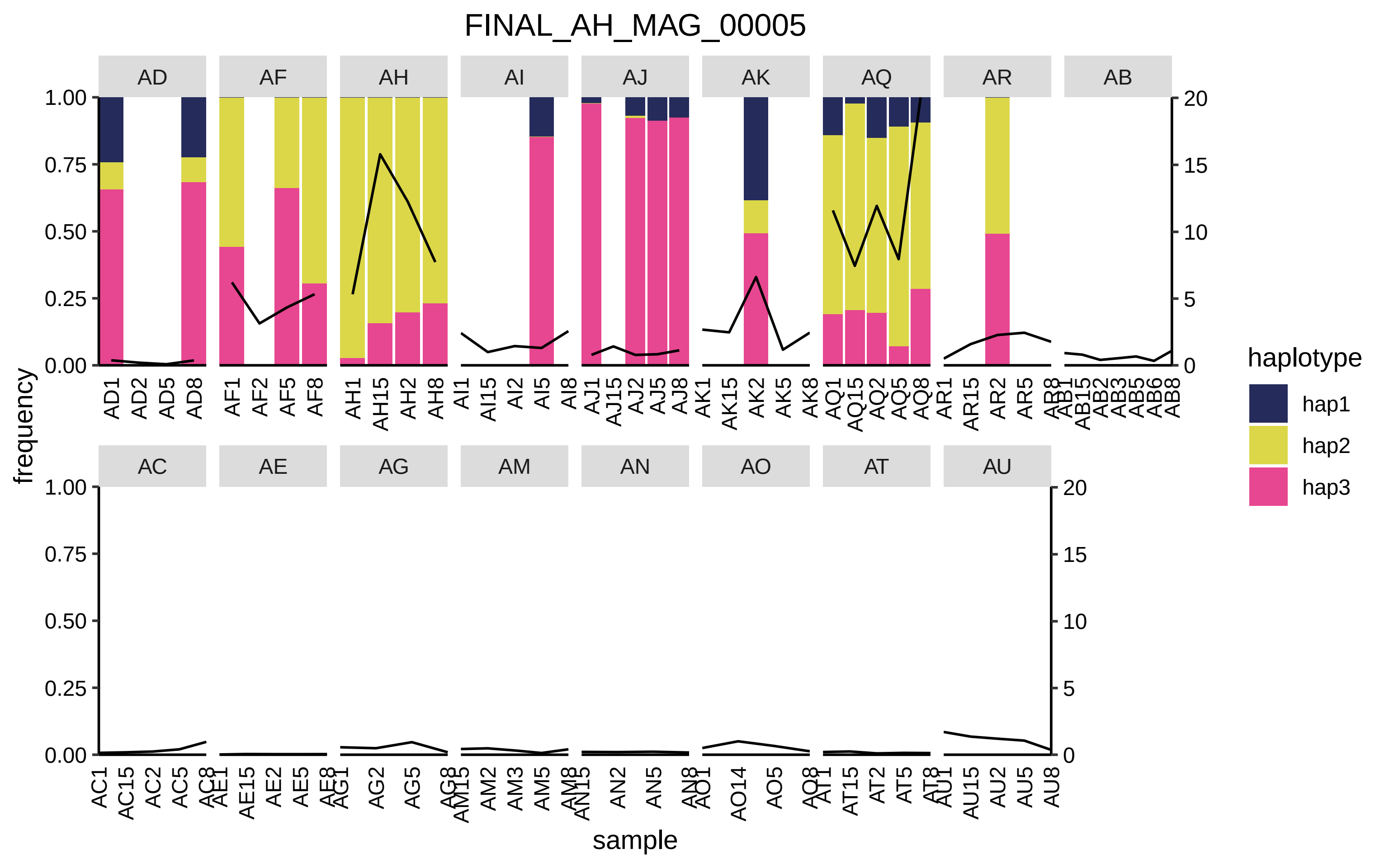
FINAL_AH_MAG_00005
AD
AF
AH
AI
AJ
AK
AQ
AR
AB
AC
AE
AG
AM
AN
AO
AT
AU
frequency
sample
haplotype
hap1
hap2
hap3
1.00
0.75
0.50
0.25
0.00
20
15
10
5
0
AD1 AD2 AD5 AD8
AF1 AF2 AF5 AF8
AH1 AH15 AH2 AH8
AI1 AI15 AI2 AI5 AI8
AJ1 AJ15 AJ2 AJ5 AJ8
AK1 AK15 AK2 AK5 AK8
AQ1 AQ15 AQ2 AQ5 AQ8
AR1 AR15 AR2 AR5 AR8
AB1 AB15 AB2 AB3 AB5 AB6 AB8
AC1 AC15 AC2 AC5 AC8
AE1 AE15 AE2 AE5 AE8
AG1 AG2 AG5 AG8
AM15 AM2 AM3 AM5 AM8
AN15 AN2 AN5 AN8
AO1 AO14 AO5 AO8
AT1 AT15 AT2 AT5 AT8
AU1 AU15 AU2 AU5 AU8

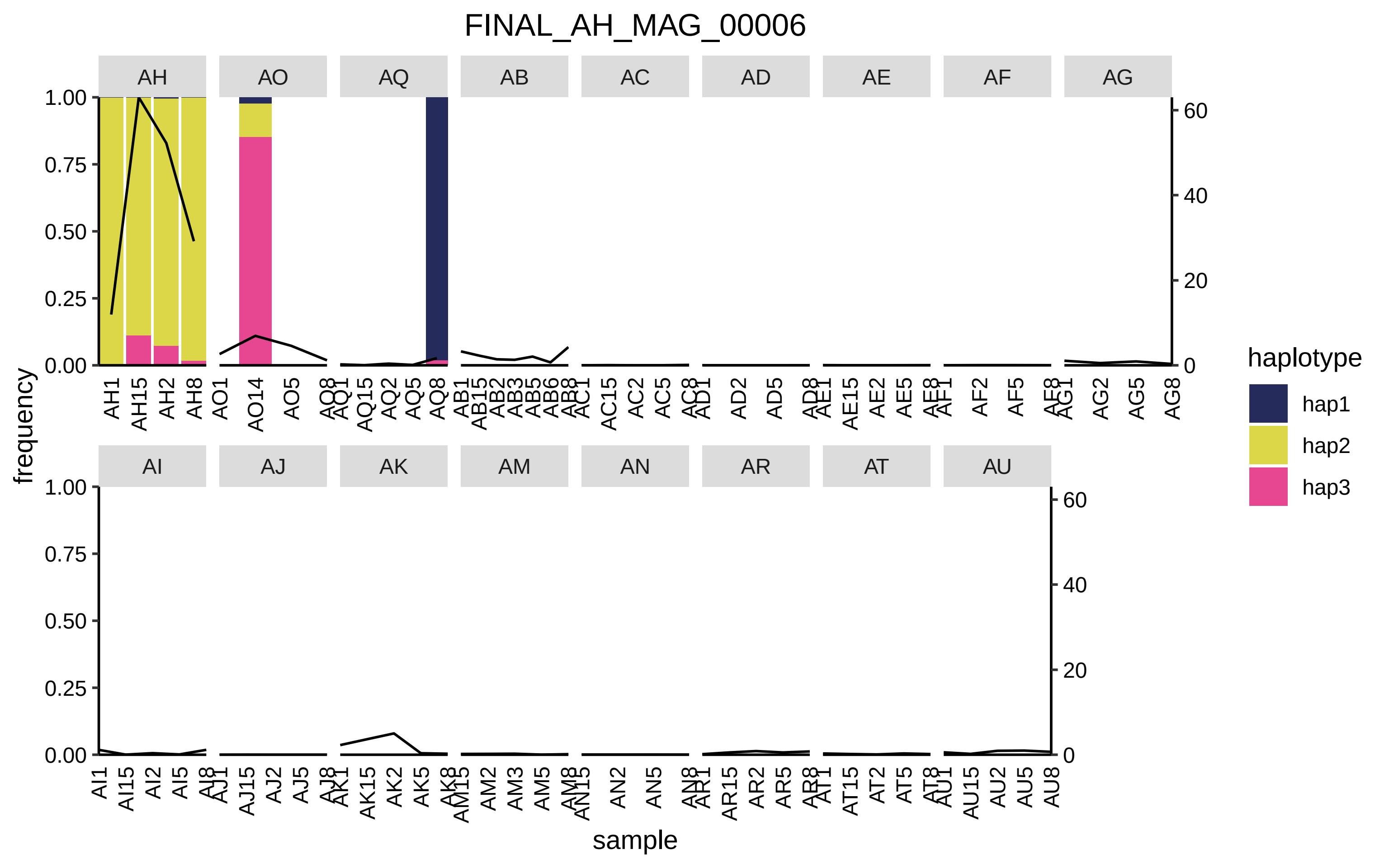

FINAL_AH_MAG_00006
AH
AO
AQ
AB
AC
AD
AE
AF
AG
AI
AJ
AK
AM
AN
AR
AT
AU
frequency
sample
haplotype
hap1
hap2
hap3

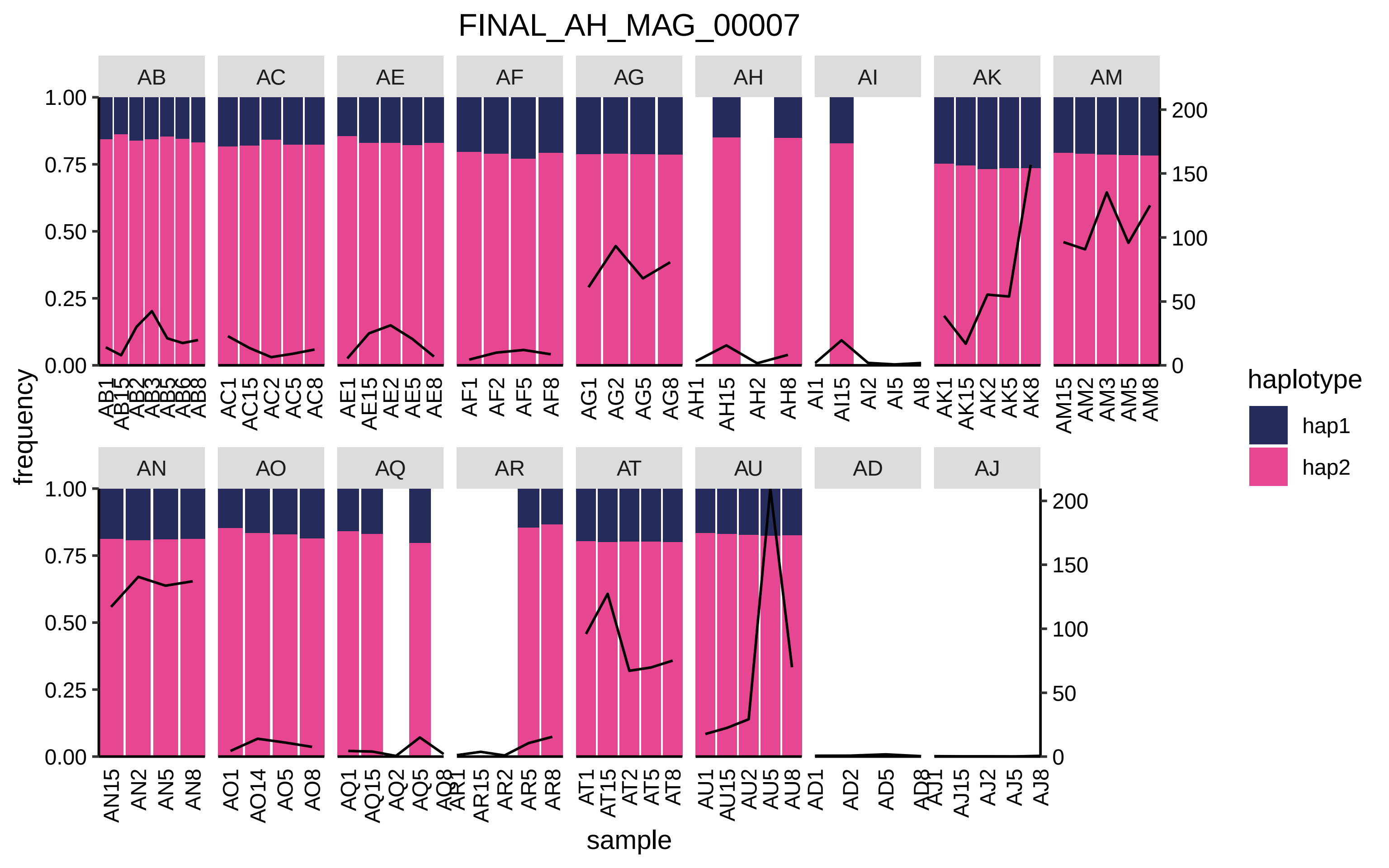

FINAL_AH_MAG_00007
AB
AC
AE
AF
AG
AH
AI
AK
AM
AN
AO
AQ
AR
AT
AU
AD
AJ
frequency
sample
haplotype
hap1
hap2

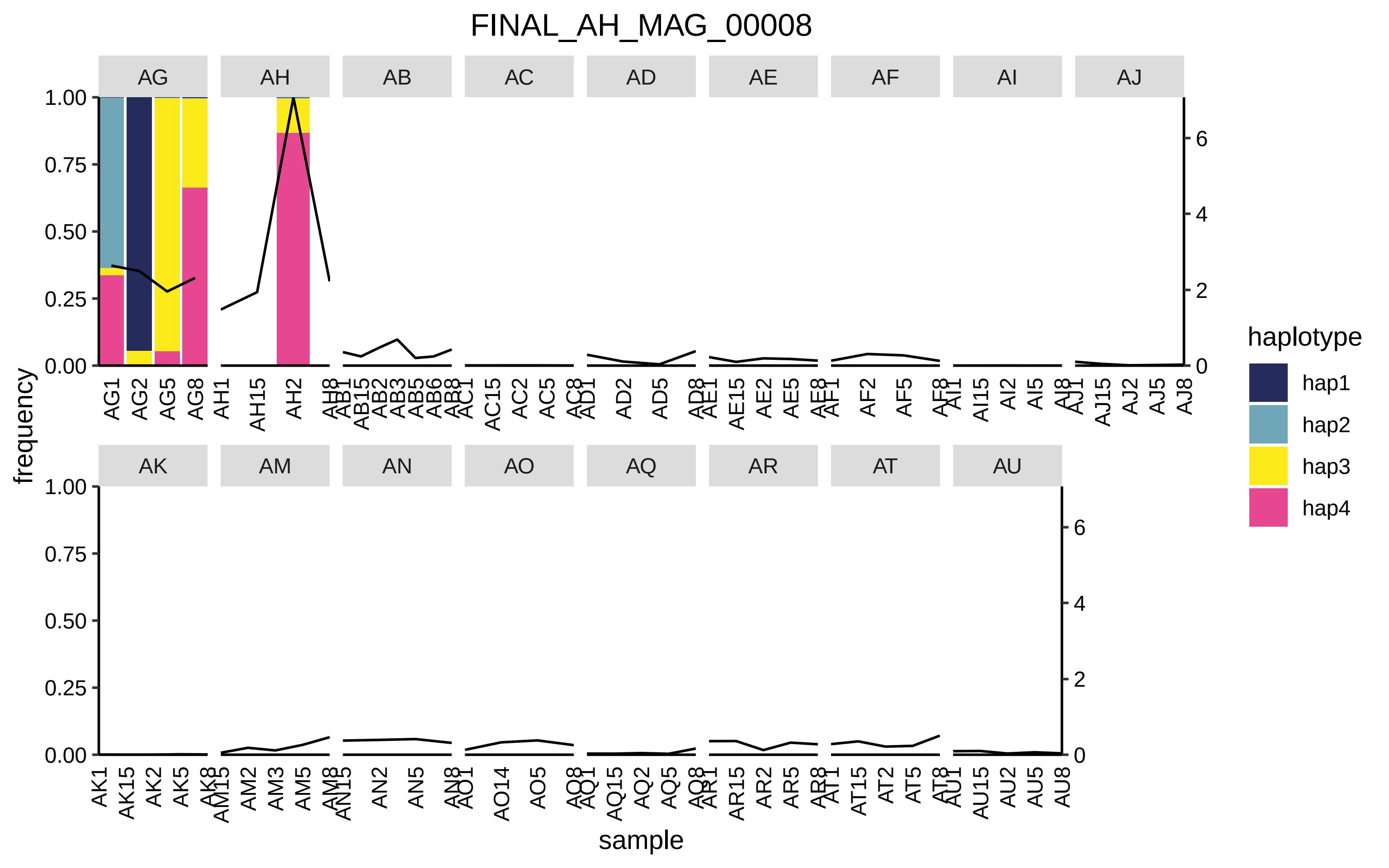
FINAL_AH_MAG_00008
AG
AH
AB
AC
AD
AE
AF
AI
AJ
AK
AM
AN
AO
AQ
AR
AT
AU
frequency
sample
haplotype
hap1
hap2
hap3
hap4
1.00
0.75
0.50
0.25
0.00
0
2
4
6
AG1 AG2 AG5 AG8
AH1 AH15 AH2 AH8
AB1 AB15 AB2 AB3 AB5 AB6 AB8
AC1 AC15 AC2 AC5 AC8
AD1 AD2 AD5 AD8
AE1 AE15 AE2 AE5 AE8
AF1 AF2 AF5 AF8
AI1 AI15 AI2 AI5 AI8
AJ1 AJ15 AJ2 AJ5 AJ8
AK1 AK15 AK2 AK5 AK8
AM15 AM2 AM3 AM5 AM8
AN15 AN2 AN5 AN8
AO1 AO14 AO5 AO8
AQ1 AQ15 AQ2 AQ5 AQ8
AR1 AR15 AR2 AR5 AR8
AT1 AT15 AT2 AT5 AT8
AU1 AU15 AU2 AU5 AU8

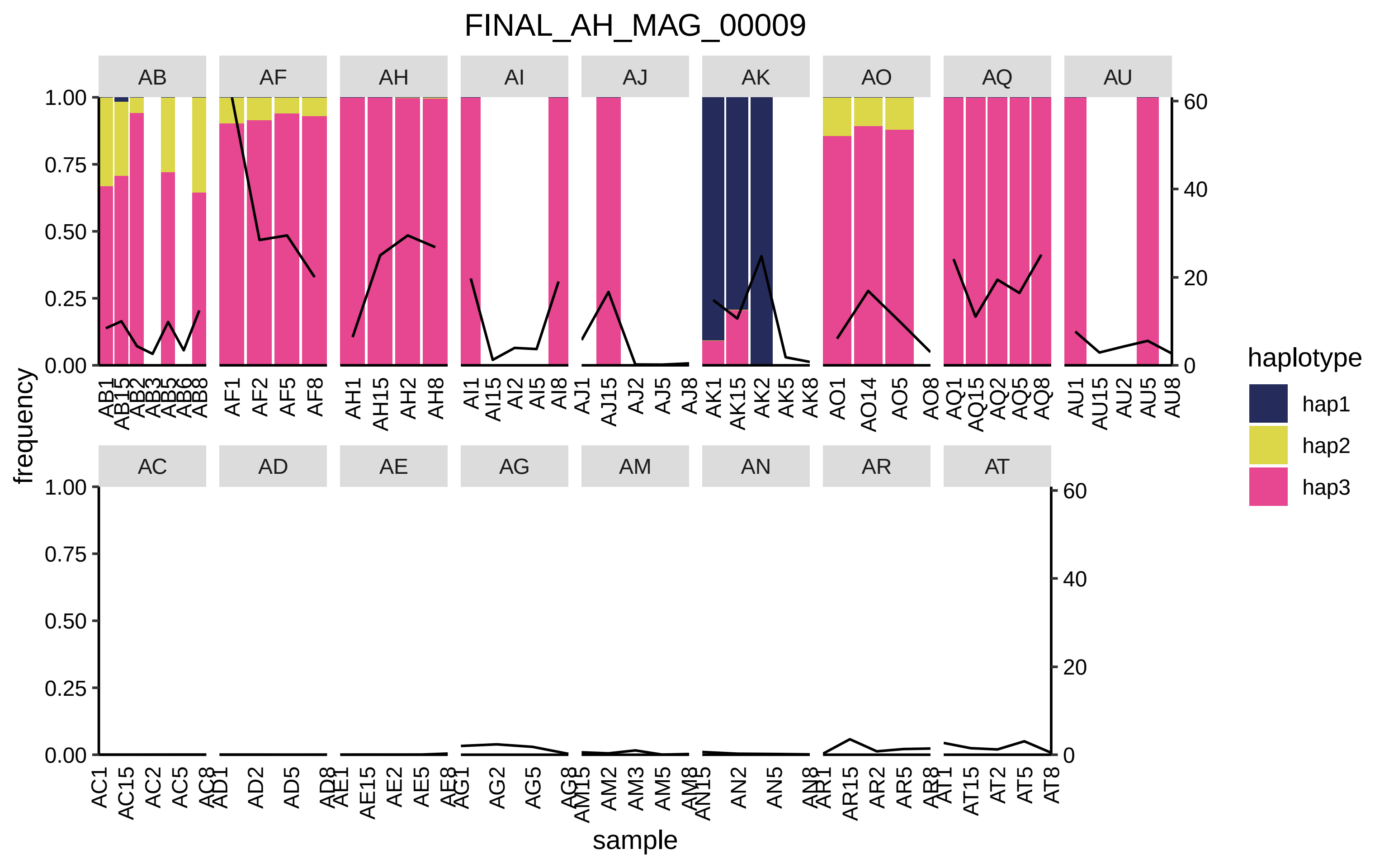

FINAL_AH_MAG_00009
AB
AF
AH
AI
AJ
AK
AO
AQ
AU
AC
AD
AE
AG
AM
AN
AR
AT
frequency
sample
1.00
0.75
0.50
0.25
0.00
60
40
20
0
haplotype
hap1
hap2
hap3
AB1 AB15 AB2 AB3 AB5 AB6 AB8
AF1 AF2 AF5 AF8
AH1 AH15 AH2 AH8
AI1 AI15 AI2 AI5 AI8
AJ1 AJ15 AJ2 AJ5 AJ8
AK1 AK15 AK2 AK5 AK8
AO1 AO14 AO5 AO8
AQ1 AQ15 AQ2 AQ5 AQ8
AU1 AU15 AU2 AU5 AU8
AC1 AC15 AC2 AC5 AC8
AD1 AD2 AD5 AD8
AE1 AE15 AE2 AE5 AE8
AG1 AG2 AG5 AG8
AM15 AM2 AM3 AM5 AM8
AN15 AN2 AN5 AN8
AR1 AR15 AR2 AR5 AR8
AT1 AT15 AT2 AT5 AT8

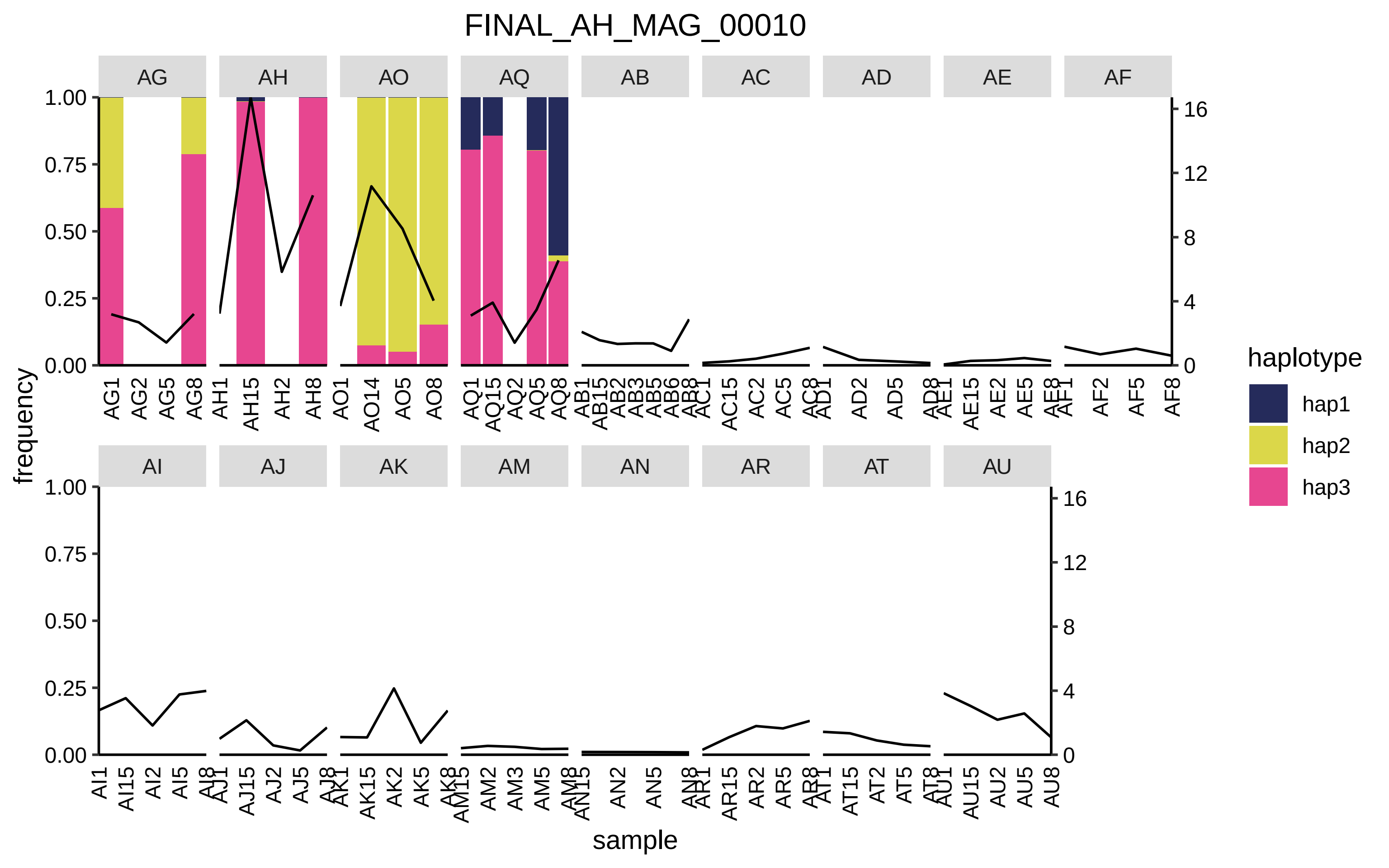
FINAL_AH_MAG_00010
AG
AH
AO
AQ
AB
AC
AD
AE
AF
AI
AJ
AK
AM
AN
AR
AT
AU
frequency
sample
haplotype
hap1
hap2
hap3

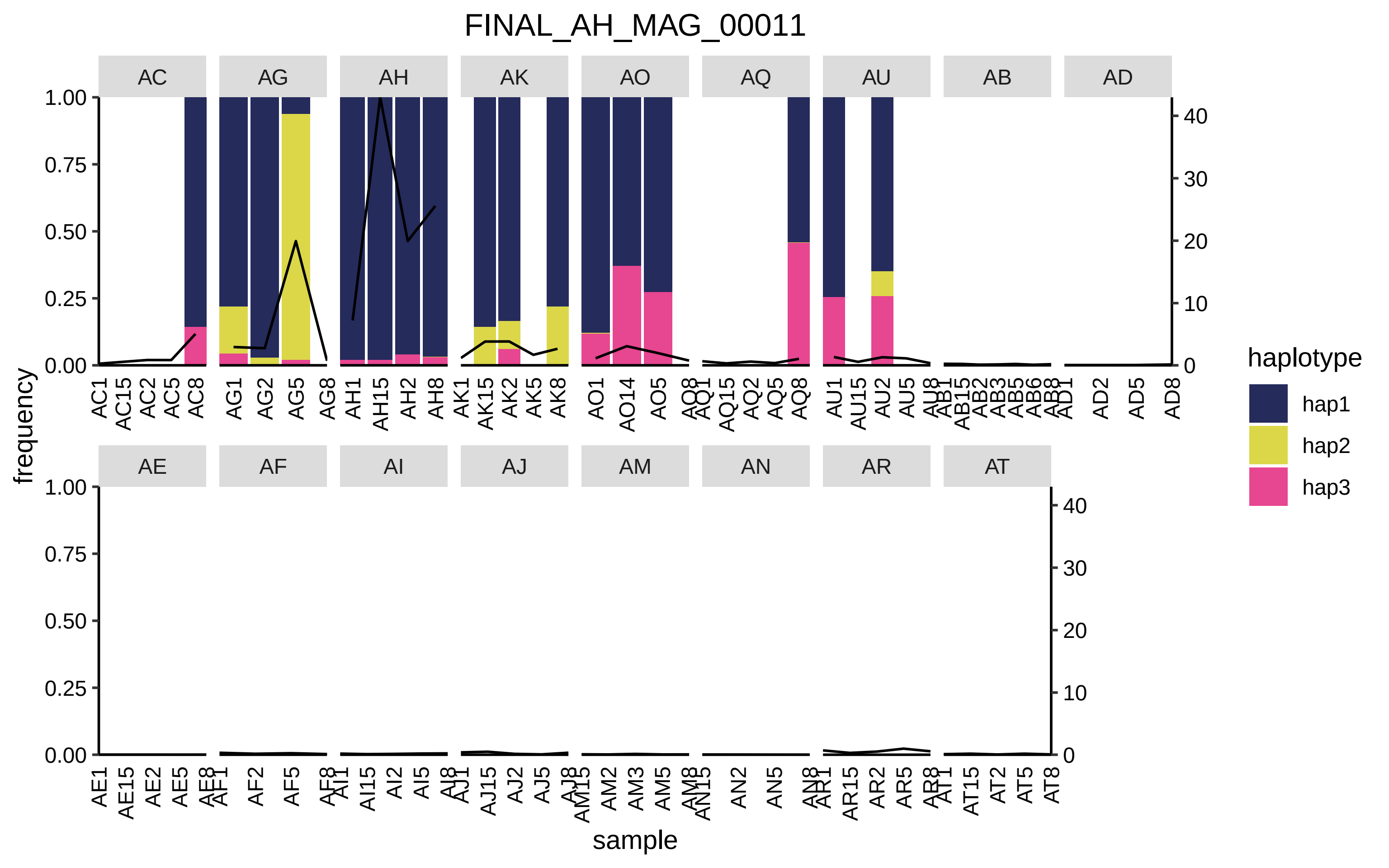

FINAL_AH_MAG_00011
AC
AG
AH
AK
AO
AQ
AU
AB
AD
AE
AF
AI
AJ
AM
AN
AR
AT
frequency
sample
haplotype
hap1
hap2
hap3

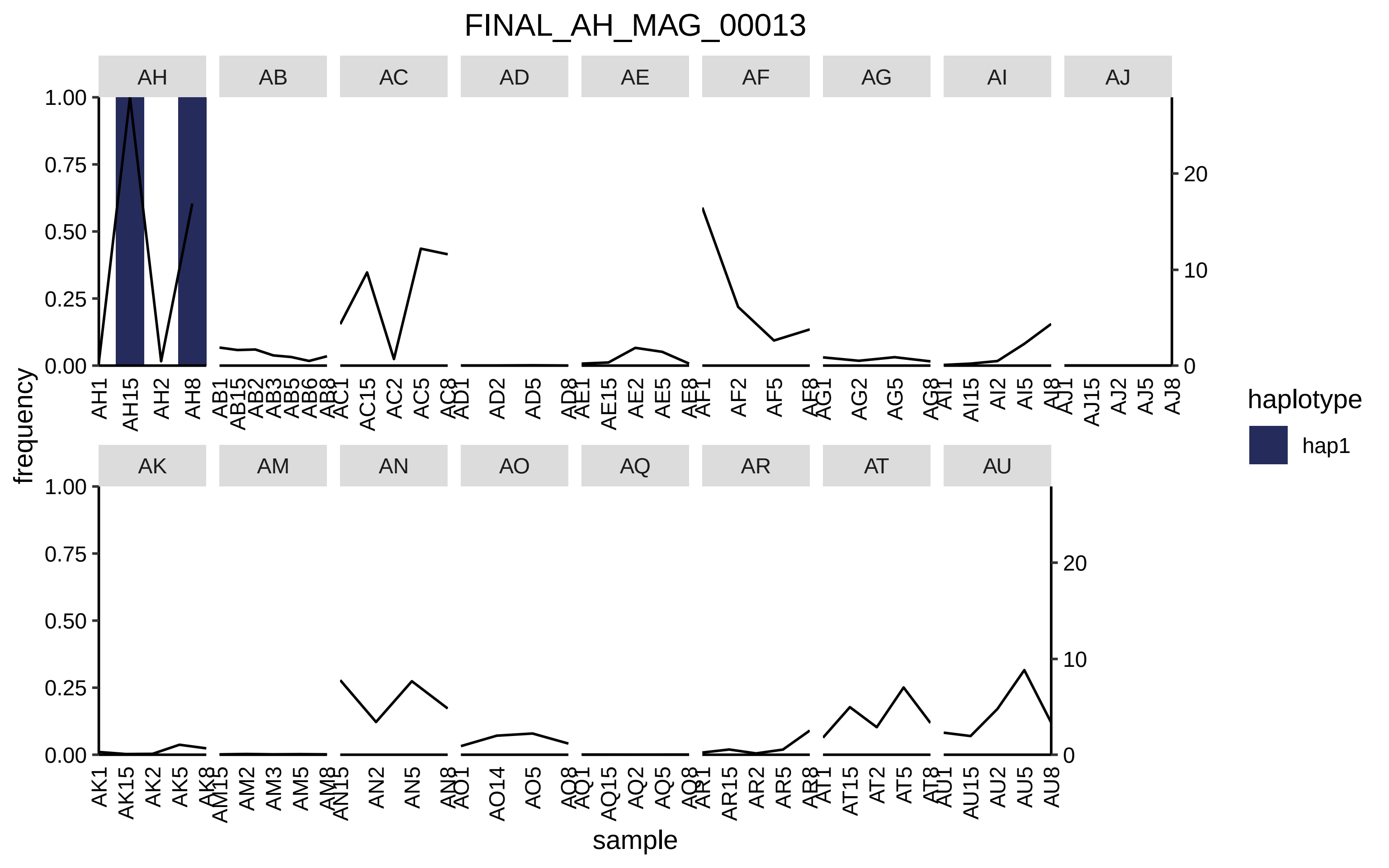
FINAL_AH_MAG_00013
AH
AB
AC
AD
AE
AF
AG
AI
AJ
AK
AM
AN
AO
AQ
AR
AT
AU
frequency
sample
haplotype
hap1
1.00
0.75
0.50
0.25
0.00
0
10
20

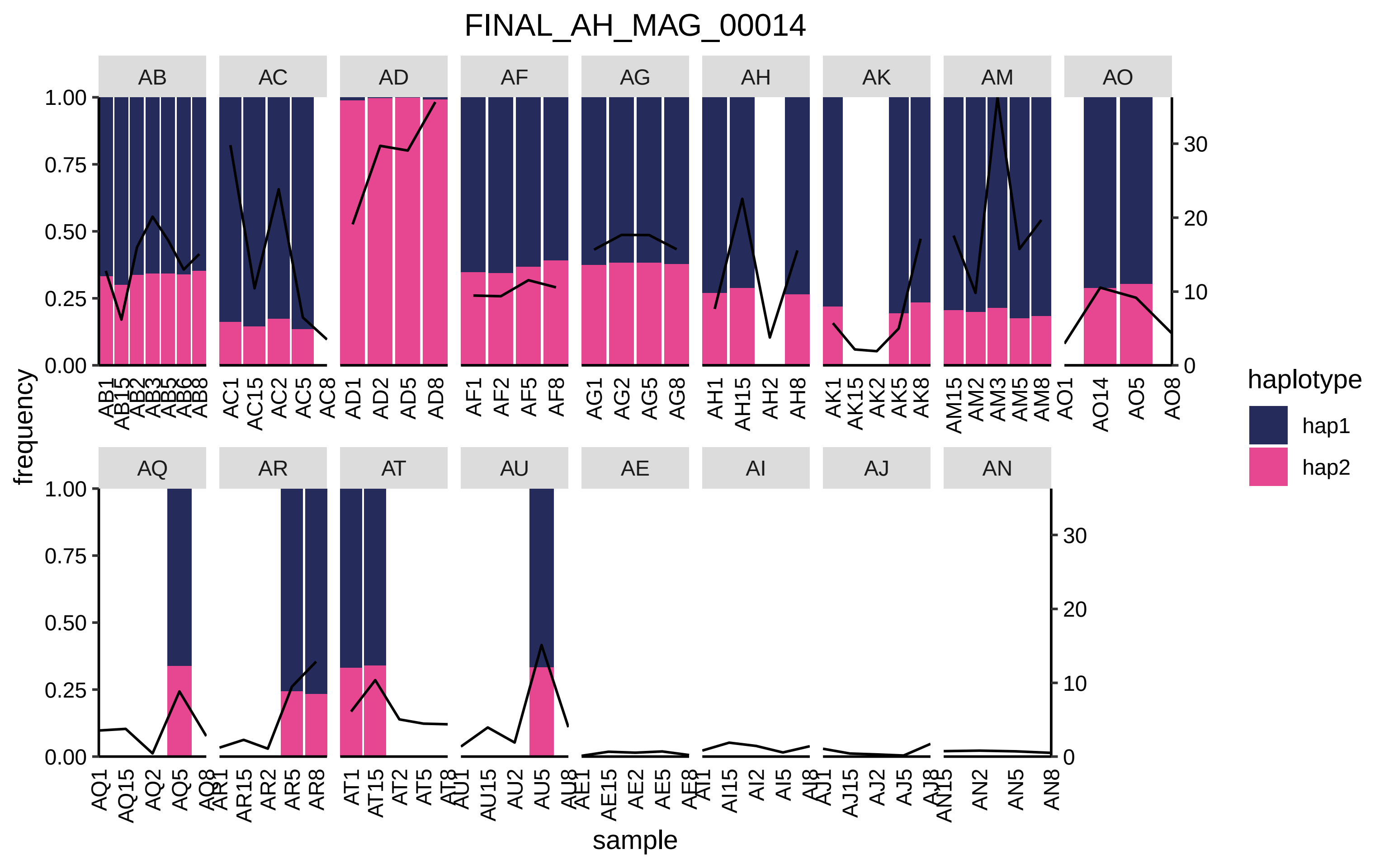

FINAL_AH_MAG_00014
AB
AC
AD
AF
AG
AH
AK
AM
AO
AQ
AR
AT
AU
AE
AI
AJ
AN
frequency
sample
haplotype
hap1
hap2

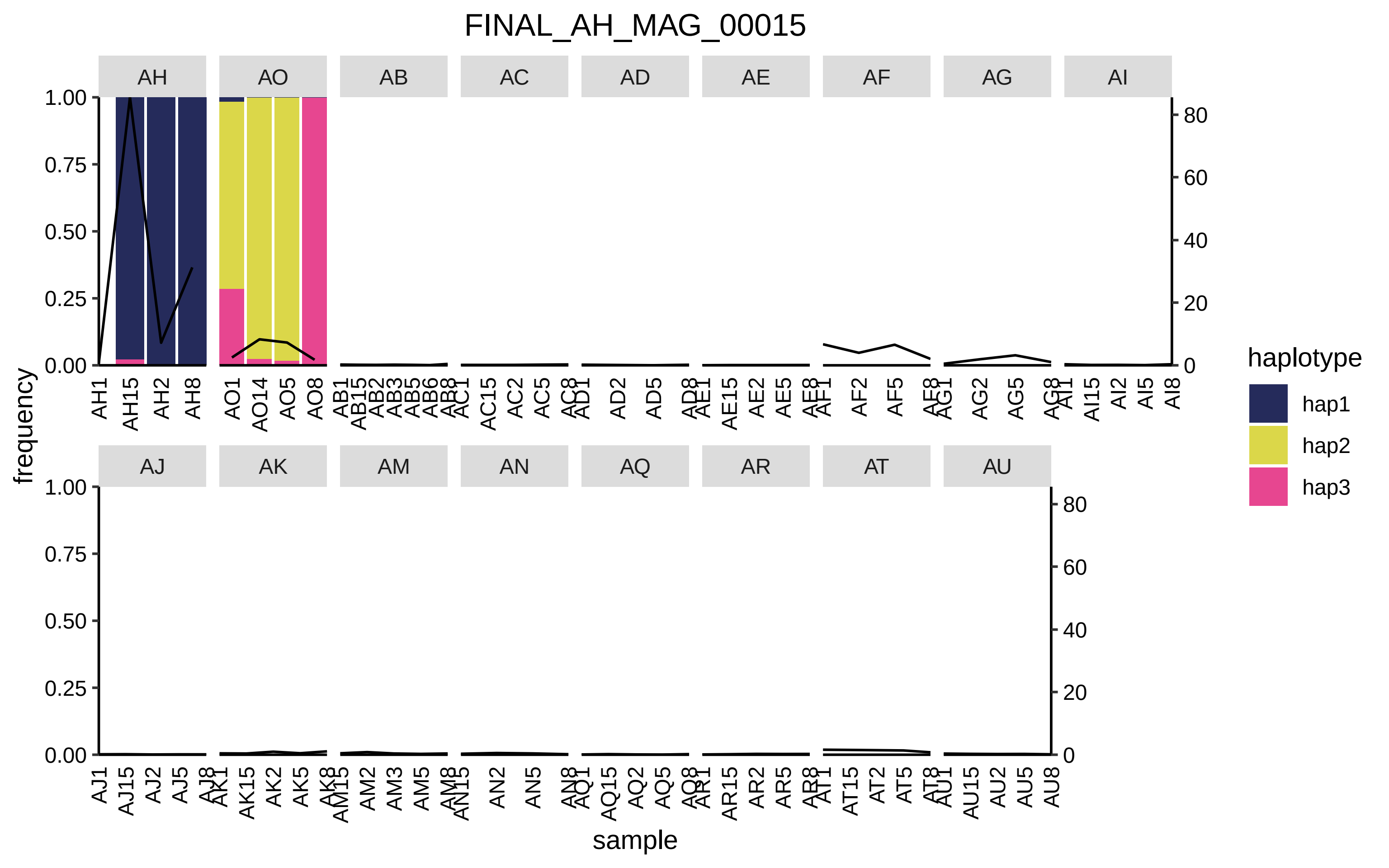

FINAL_AH_MAG_00015
AH
AO
AB
AC
AD
AE
AF
AG
AI
AJ
AK
AM
AN
AQ
AR
AT
AU
1.00
0.75
0.50
0.25
0.00
80
60
40
20
0
frequency
sample
AH1 AH15 AH2 AH8
AO1 AO14 AO5 AO8
AB1 AB15 AB2 AB3 AB5 AB6 AB8
AC1 AC15 AC2 AC5 AC8
AD1 AD2 AD5 AD8
AE1 AE15 AE2 AE5 AE8
AF1 AF2 AF5 AF8
AG1 AG2 AG5 AG8
AI1 AI15 AI2 AI5 AI8
AJ1 AJ15 AJ2 AJ5 AJ8
AK1 AK15 AK2 AK5 AK8
AM15 AM2 AM3 AM5 AM8
AN15 AN2 AN5 AN8
AQ1 AQ15 AQ2 AQ5 AQ8
AR1 AR15 AR2 AR5 AR8
AT1 AT15 AT2 AT5 AT8
AU1 AU15 AU2 AU5 AU8
haplotype
hap1
hap2
hap3

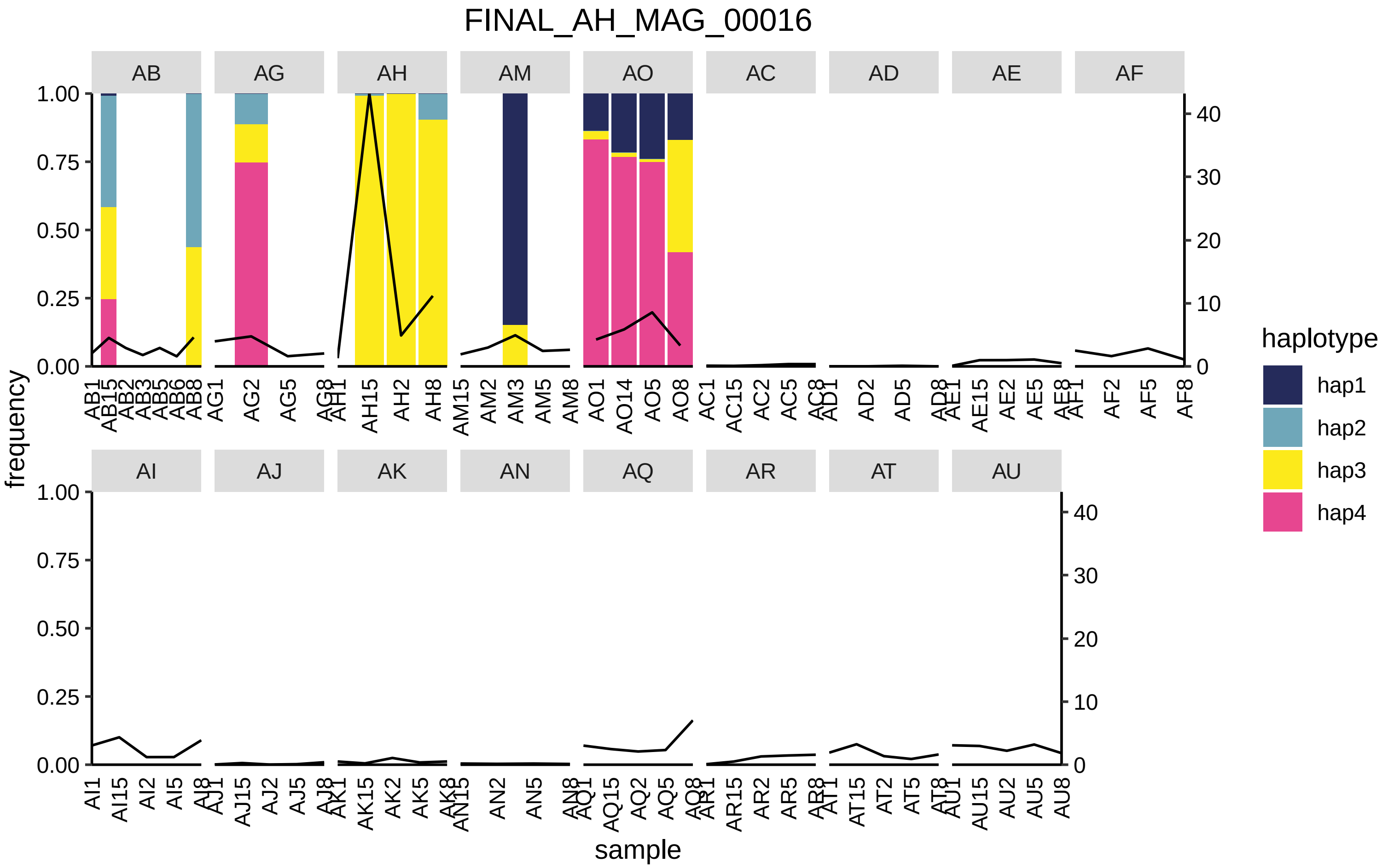
FINAL_AH_MAG_00016
AB
AG
AH
AM
AO
AC
AD
AE
AF
AI
AJ
AK
AN
AQ
AR
AT
AU
frequency
1.00
0.75
0.50
0.25
0.00
40
30
20
10
0
AB1 AB15 AB2 AB3 AB5 AB6 AB8
AG1 AG2 AG5 AG8
AH1 AH15 AH2 AH8
AM15 AM2 AM3 AM5 AM8
AO1 AO14 AO5 AO8
AC1 AC15 AC2 AC5 AC8
AD1 AD2 AD5 AD8
AE1 AE15 AE2 AE5 AE8
AF1 AF2 AF5 AF8
AI1 AI15 AI2 AI5 AI8
AJ1 AJ15 AJ2 AJ5 AJ8
AK1 AK15 AK2 AK5 AK8
AN15 AN2 AN5 AN8
AQ1 AQ15 AQ2 AQ5 AQ8
AR1 AR15 AR2 AR5 AR8
AT1 AT15 AT2 AT5 AT8
AU1 AU15 AU2 AU5 AU8
sample
haplotype
hap1
hap2
hap3
hap4

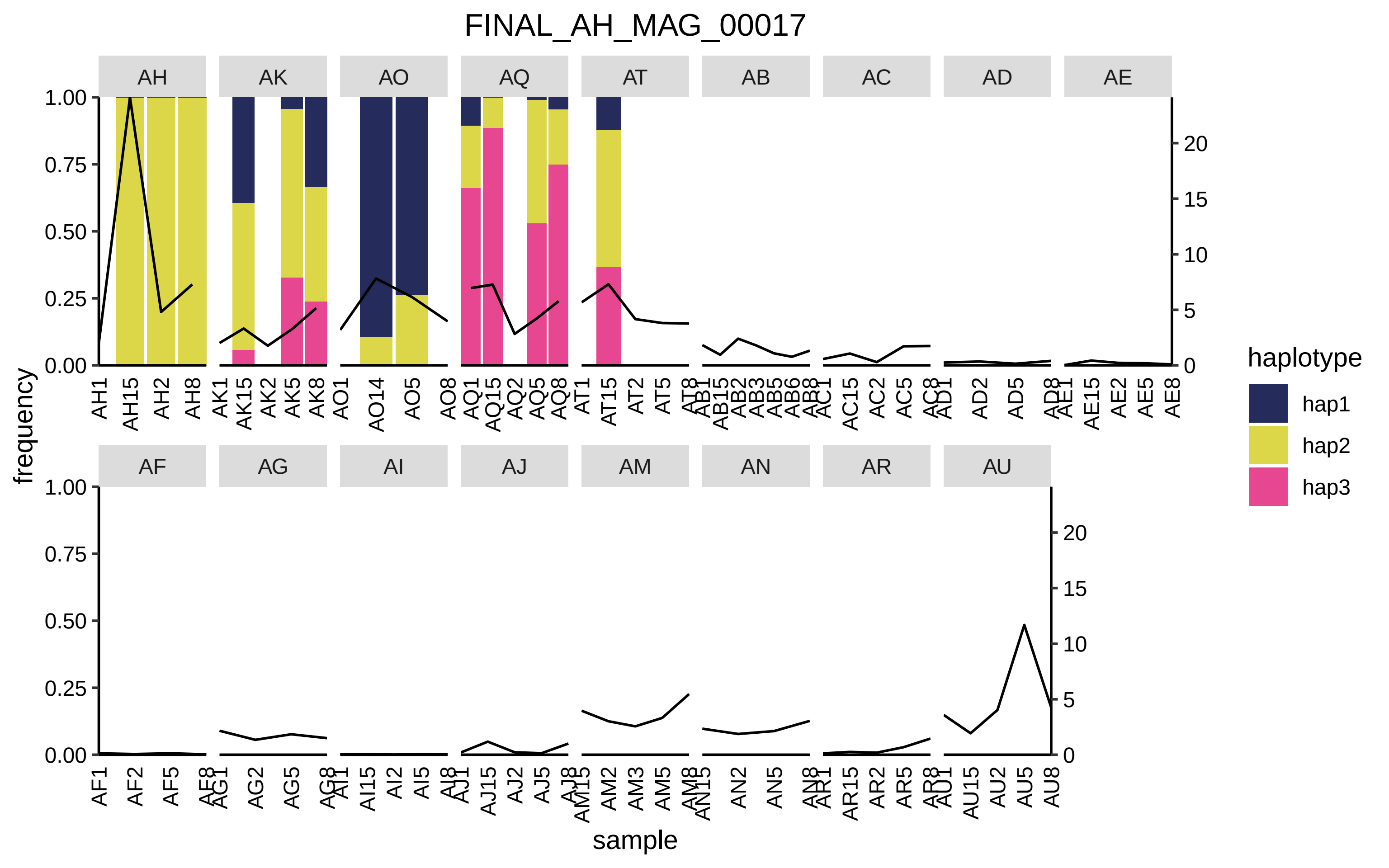

FINAL_AH_MAG_00017
AH
AK
AO
AQ
AT
AB
AC
AD
AE
AF
AG
AI
AJ
AM
AN
AR
AU
1.00
0.75
0.50
0.25
0.00
20
15
10
5
0
frequency
sample
haplotype
hap1
hap2
hap3

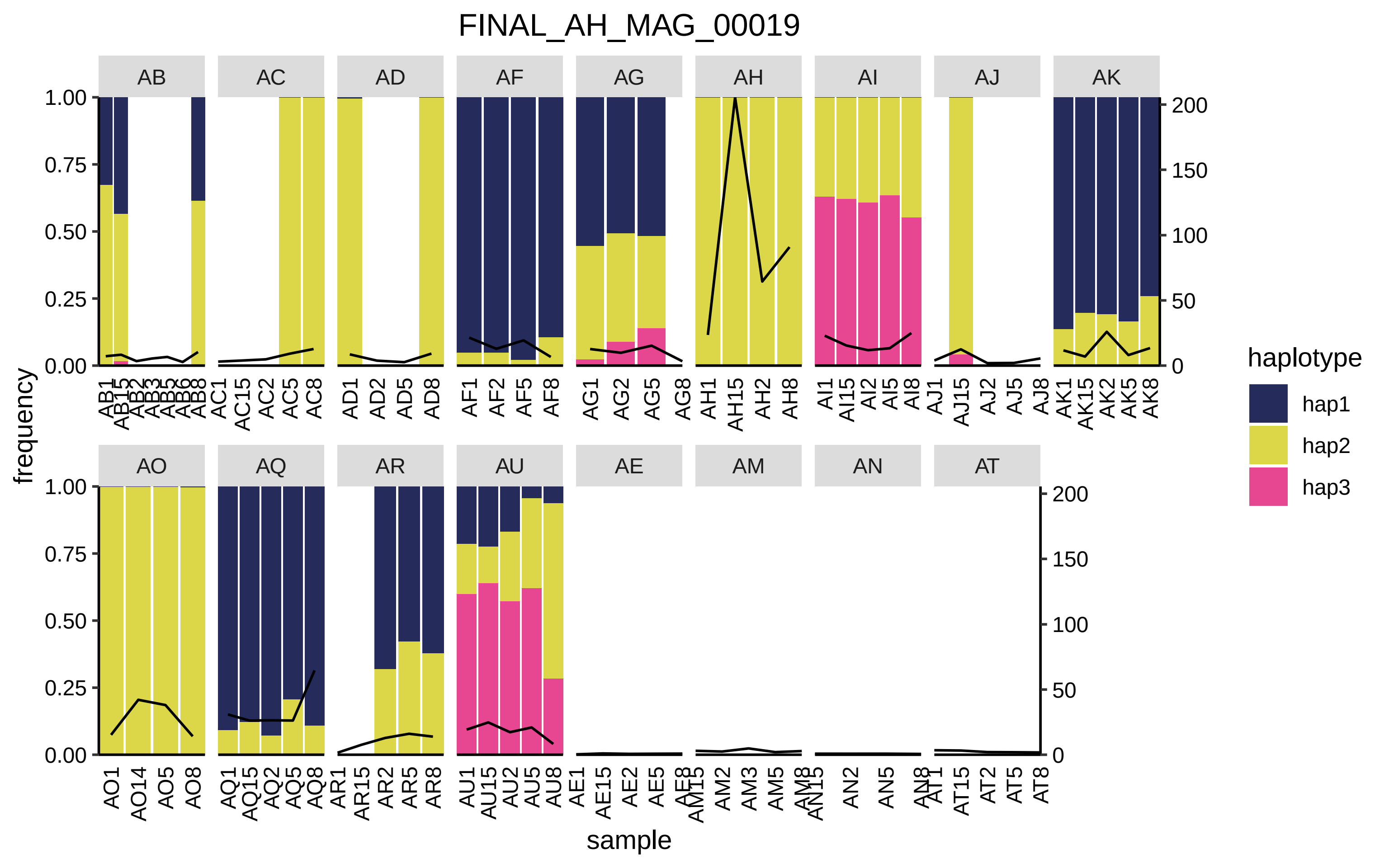

FINAL_AH_MAG_00019
AB
AC
AD
AF
AG
AH
AI
AJ
AK
AO
AQ
AR
AU
AE
AM
AN
AT
frequency
sample
haplotype
hap1
hap2
hap3

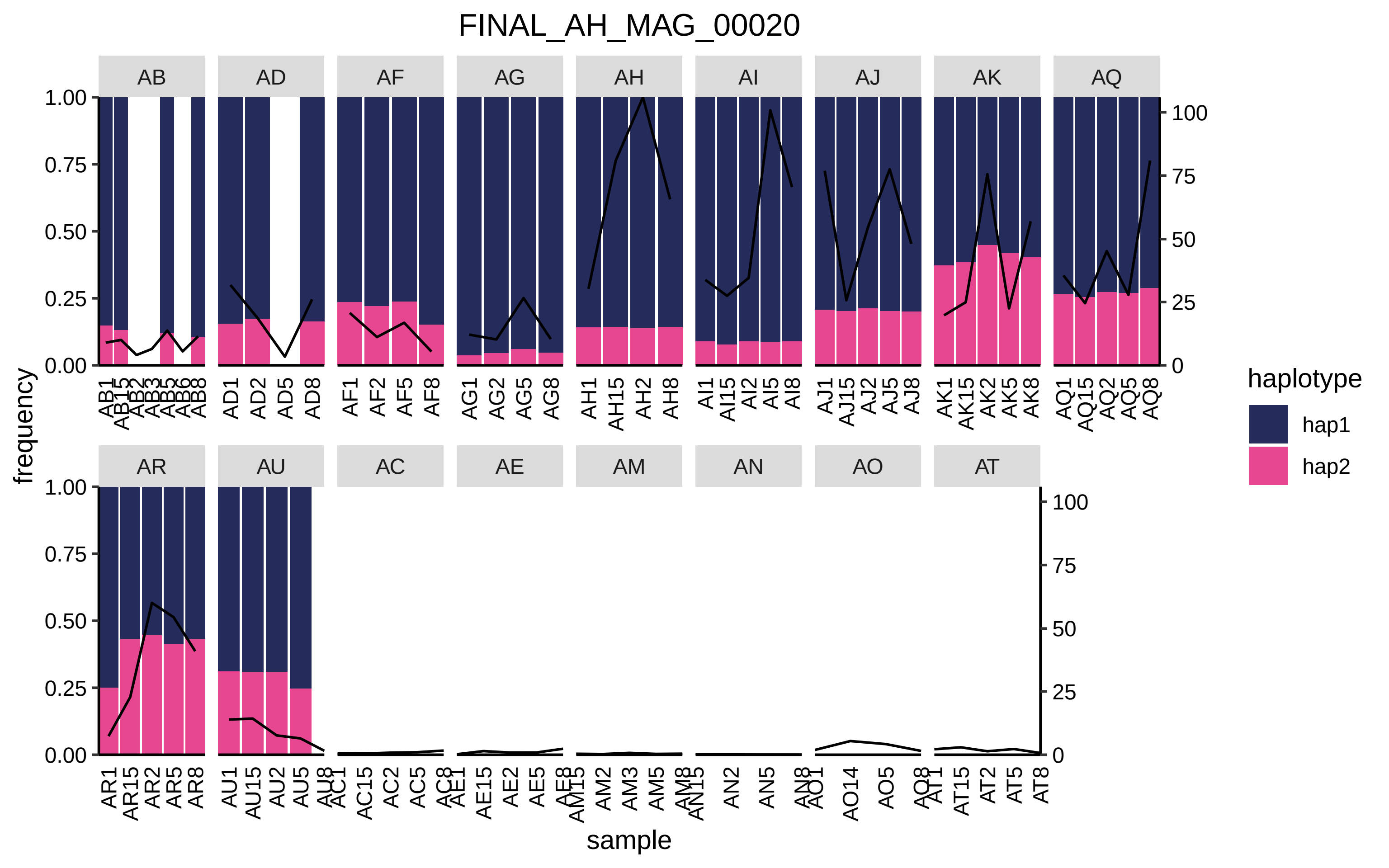

FINAL_AH_MAG_00020
AB
AD
AF
AG
AH
AI
AJ
AK
AQ
AR
AU
AC
AE
AM
AN
AO
AT
frequency
sample
haplotype
hap1
hap2
1.00
0.75
0.50
0.25
0.00
100
75
50
25
0
AD1
AD2
AD5
AD8
AF1
AF2
AF5
AF8
AG1
AG2
AG5
AG8
AH1
AH15
AH2
AH8
AI1
AI15
AI2
AI5
AI8
AJ1
AJ15
AJ2
AJ5
AJ8
AK1
AK15
AK2
AK5
AK8
AQ1
AQ15
AQ2
AQ5
AQ8
AR1
AR15
AR2
AR5
AR8
AU1
AU15
AU2
AU5
AC15
AC2
AC5
AE15
AE2
AE5
AM2
AM3
AM5
AN2
AN5
AO14
AO5
AT15
AT2
AT5
AT8

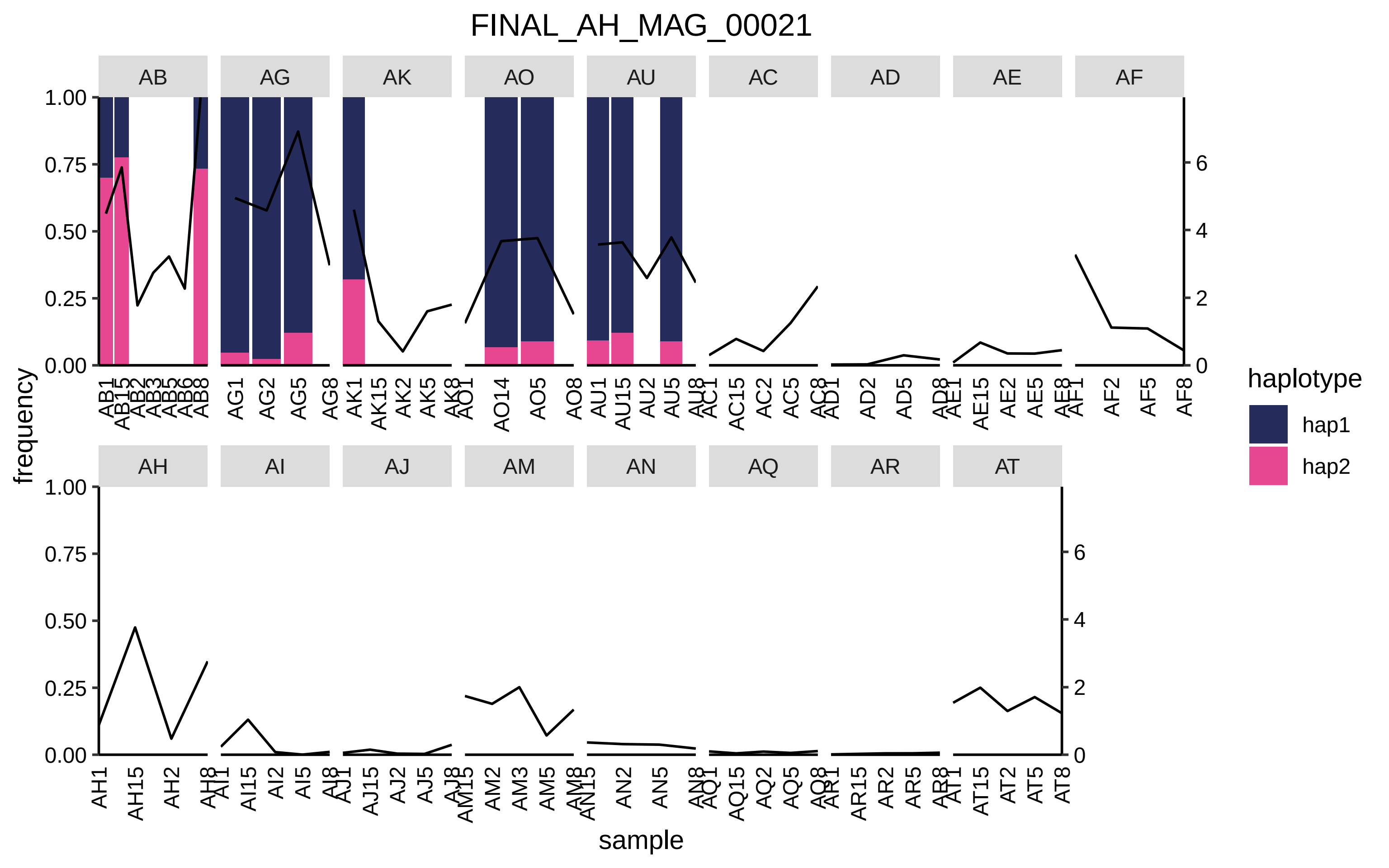

FINAL_AH_MAG_00021
AB
AG
AK
AO
AU
AC
AD
AE
AF
AH
AI
AJ
AM
AN
AQ
AR
AT
frequency
sample
haplotype
hap1
hap2
1.00
0.75
0.50
0.25
0.00
0
2
4
6
AB1 AB15 AB2 AB3 AB5 AB6 AB8
AG1 AG2 AG5 AG8
AK1 AK15 AK2 AK5 AK8
AO1 AO14 AO5 AO8
AU1 AU15 AU2 AU5 AU8
AC1 AC15 AC2 AC5 AC8
AD1 AD2 AD5 AD8
AE1 AE15 AE2 AE5 AE8
AF1 AF2 AF5 AF8
AH1 AH15 AH2 AH8
AI1 AI15 AI2 AI5 AI8
AJ1 AJ15 AJ2 AJ5 AJ8
AM15 AM2 AM3 AM5 AM8
AN15 AN2 AN5 AN8
AQ1 AQ15 AQ2 AQ5 AQ8
AR1 AR15 AR2 AR5 AR8
AT1 AT15 AT2 AT5 AT8

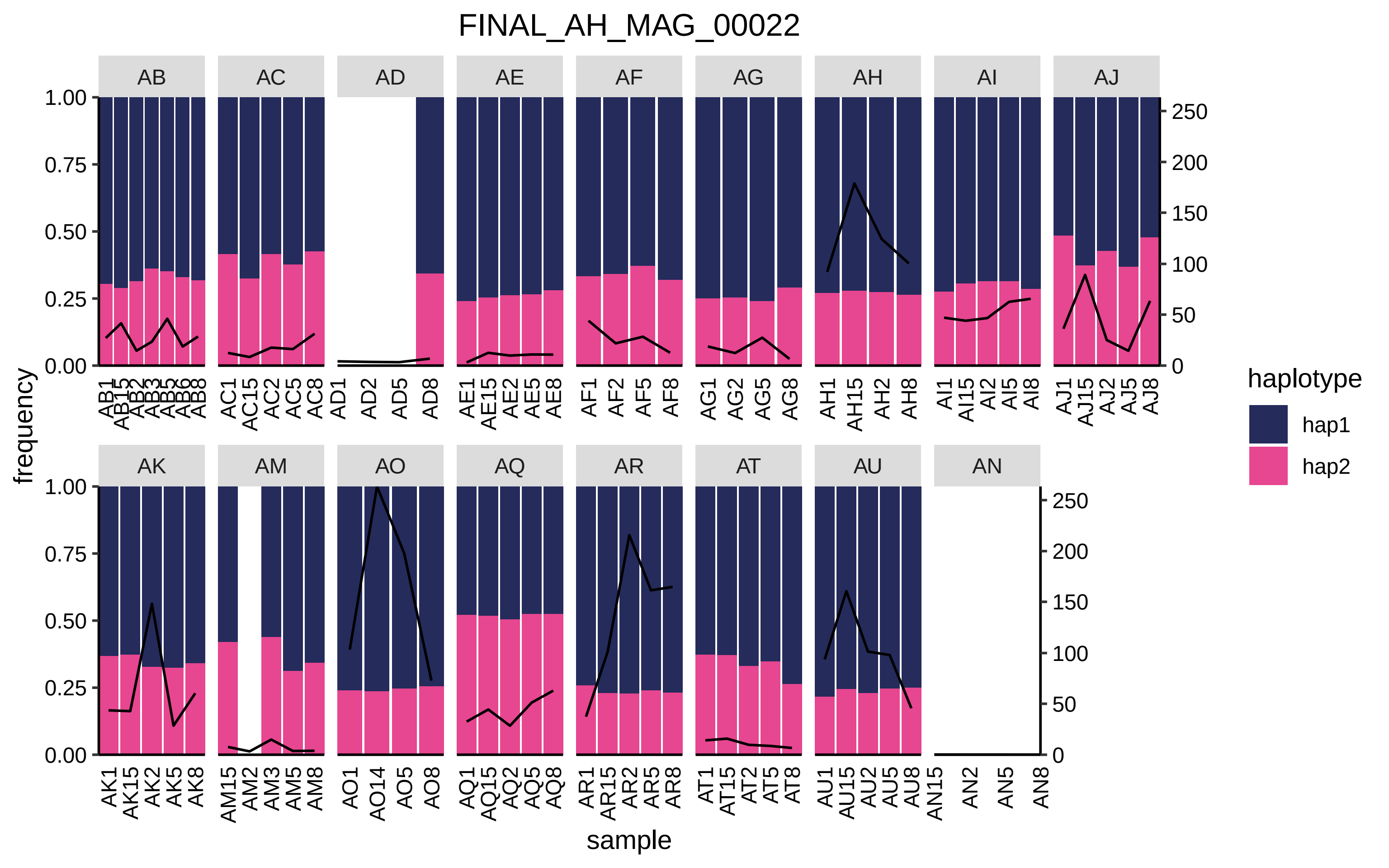

FINAL_AH_MAG_00022
AB
AC
AD
AE
AF
AG
AH
AI
AJ
AK
AM
AO
AQ
AR
AT
AU
AN
frequency
sample
haplotype
hap1
hap2

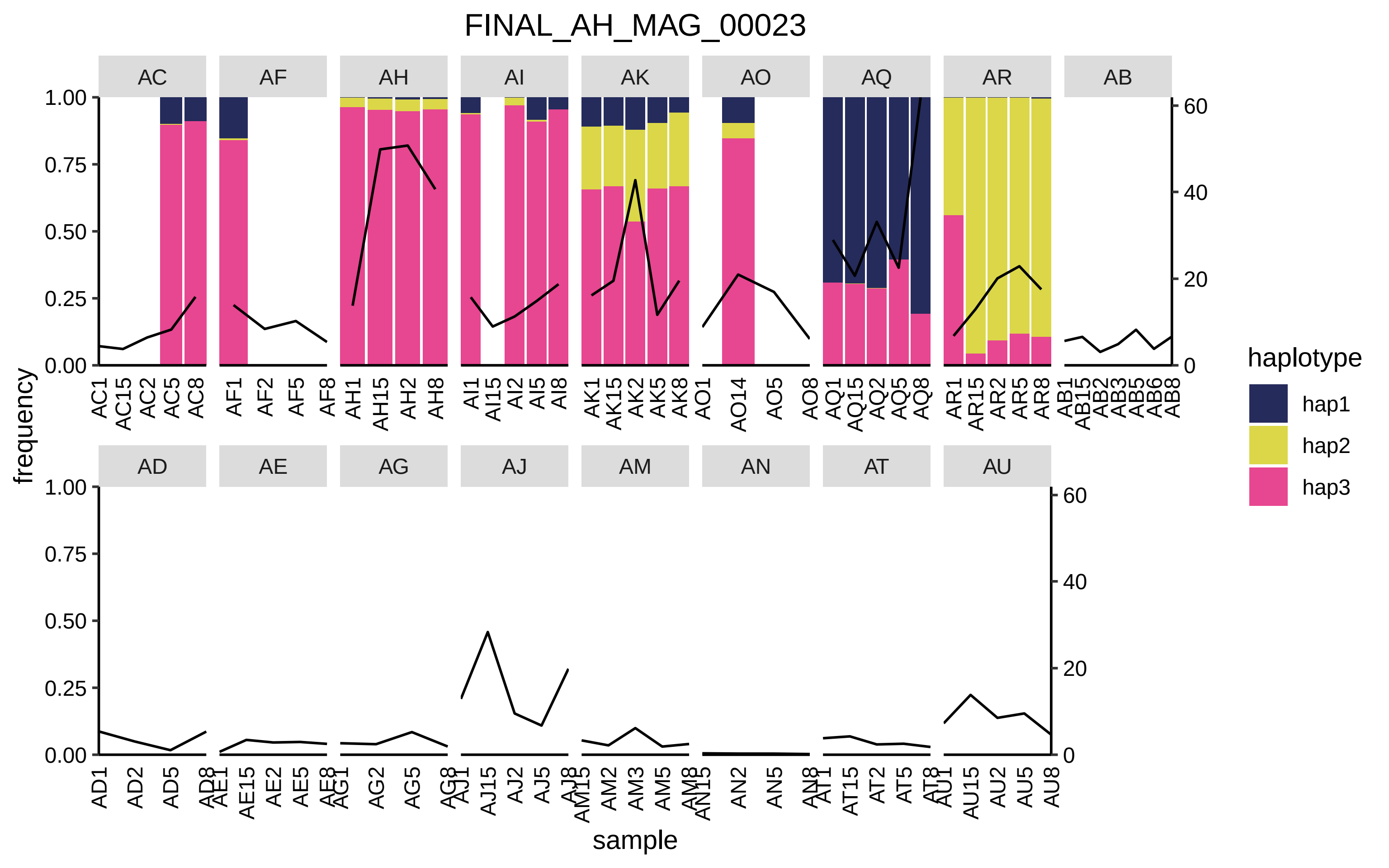

FINAL_AH_MAG_00023
AC
AF
AH
AI
AK
AO
AQ
AR
AB
AD
AE
AG
AJ
AM
AN
AT
AU
frequency
sample
haplotype
hap1
hap2
hap3

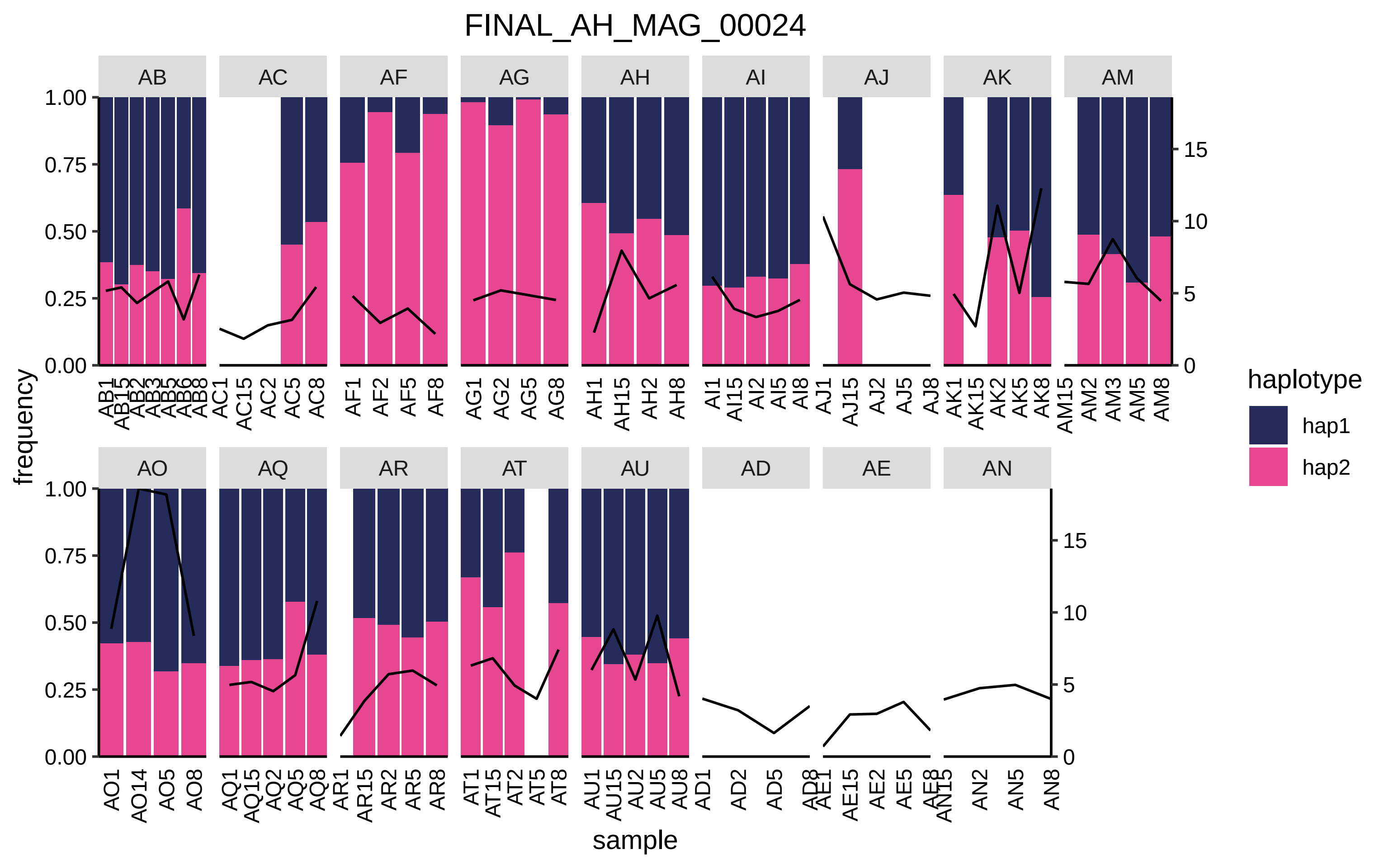

FINAL_AH_MAG_00024
AB
AC
AF
AG
AH
AI
AJ
AK
AM
AO
AQ
AR
AT
AU
AD
AE
AN
frequency
1.00
0.75
0.50
0.25
0.00
15
10
5
0
AB1 AB15 AB2 AB3 AB5 AB6 AB8
AC1 AC15 AC2 AC5 AC8
AF1 AF2 AF5 AF8
AG1 AG2 AG5 AG8
AH1 AH15 AH2 AH8
AI1 AI15 AI2 AI5 AI8
AJ1 AJ15 AJ2 AJ5 AJ8
AK1 AK15 AK2 AK5 AK8
AM15 AM2 AM3 AM5 AM8
AO1 AO14 AO5 AO8
AQ1 AQ15 AQ2 AQ5 AQ8
AR1 AR15 AR2 AR5 AR8
AT1 AT15 AT2 AT5 AT8
AU1 AU15 AU2 AU5 AU8
AD1 AD2 AD5 AD8
AE1 AE15 AE2 AE5 AE8
AN15 AN2 AN5 AN8
sample
haplotype
hap1
hap2

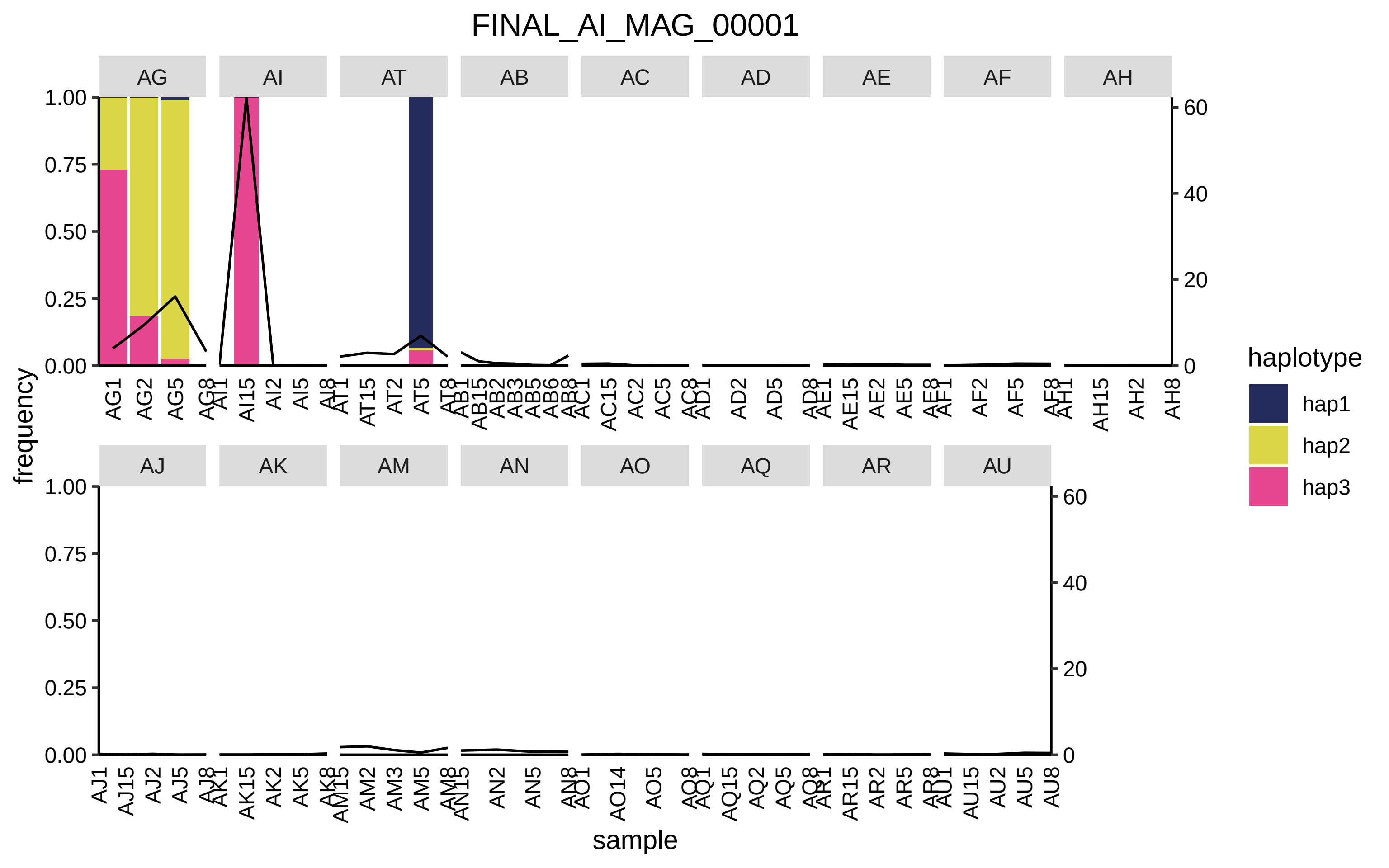

FINAL_AI_MAG_00001
AG
AI
AT
AB
AC
AD
AE
AF
AH
AJ
AK
AM
AN
AO
AQ
AR
AU
frequency
sample
haplotype
hap1
hap2
hap3

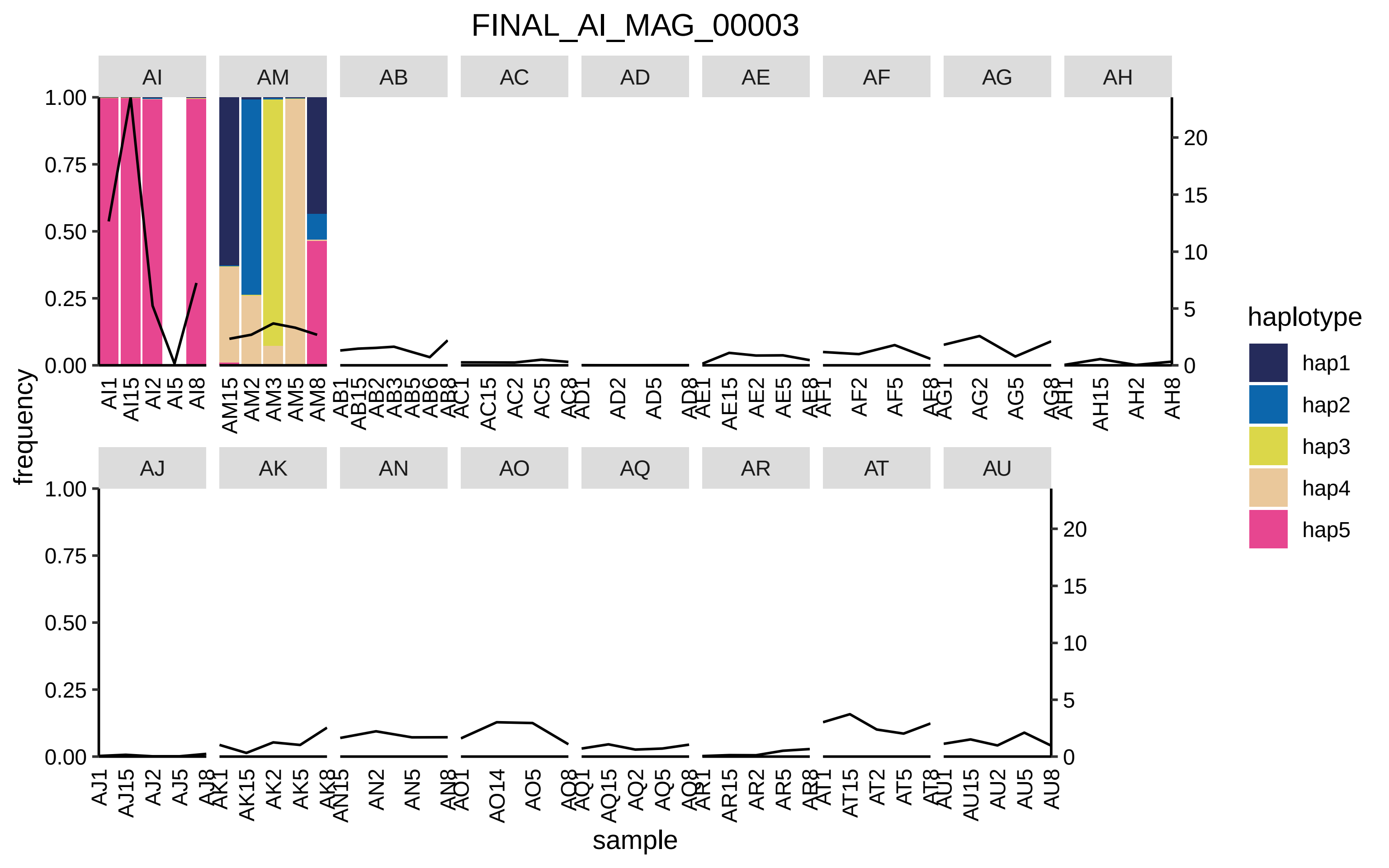
FINAL_AI_MAG_00003
AI
AM
AB
AC
AD
AE
AF
AG
AH
AJ
AK
AN
AO
AQ
AR
AT
AU
frequency
sample
haplotype
hap1
hap2
hap3
hap4
hap5

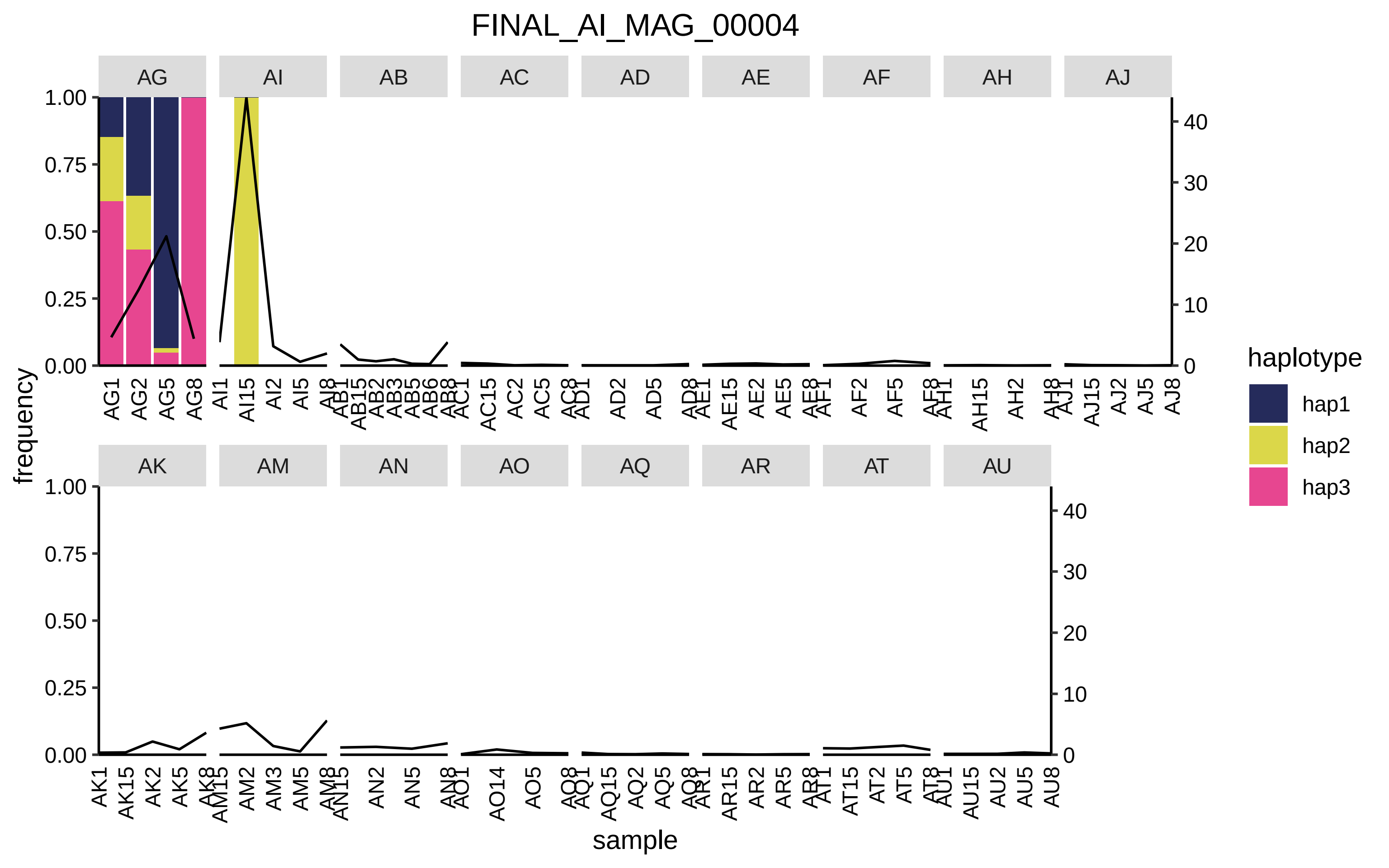

FINAL_AI_MAG_00004
AG
AI
AB
AC
AD
AE
AF
AH
AJ
AK
AM
AN
AO
AQ
AR
AT
AU
frequency
sample
haplotype
hap1
hap2
hap3
1.00
0.75
0.50
0.25
0.00
40
30
20
10
0
AG1 AG2 AG5 AG8 AI1 AI15 AI2 AI5 AI8 AB1 AB15 AB2 AB3 AB5 AB6 AB8 AC1 AC15 AC2 AC5 AC8 AD1 AD2 AD5 AD8 AE1 AE15 AE2 AE5 AE8 AF1 AF2 AF5 AF8 AH1 AH15 AH2 AH8 AJ1 AJ15 AJ2 AJ5 AJ8
AK1 AK15 AK2 AK5 AK8 AM15 AM2 AM3 AM5 AM8 AN15 AN2 AN5 AN8 AO1 AO14 AO5 AO8 AQ1 AQ15 AQ2 AQ5 AQ8 AR1 AR15 AR2 AR5 AR8 AT1 AT15 AT2 AT5 AT8 AU1 AU15 AU2 AU5 AU8

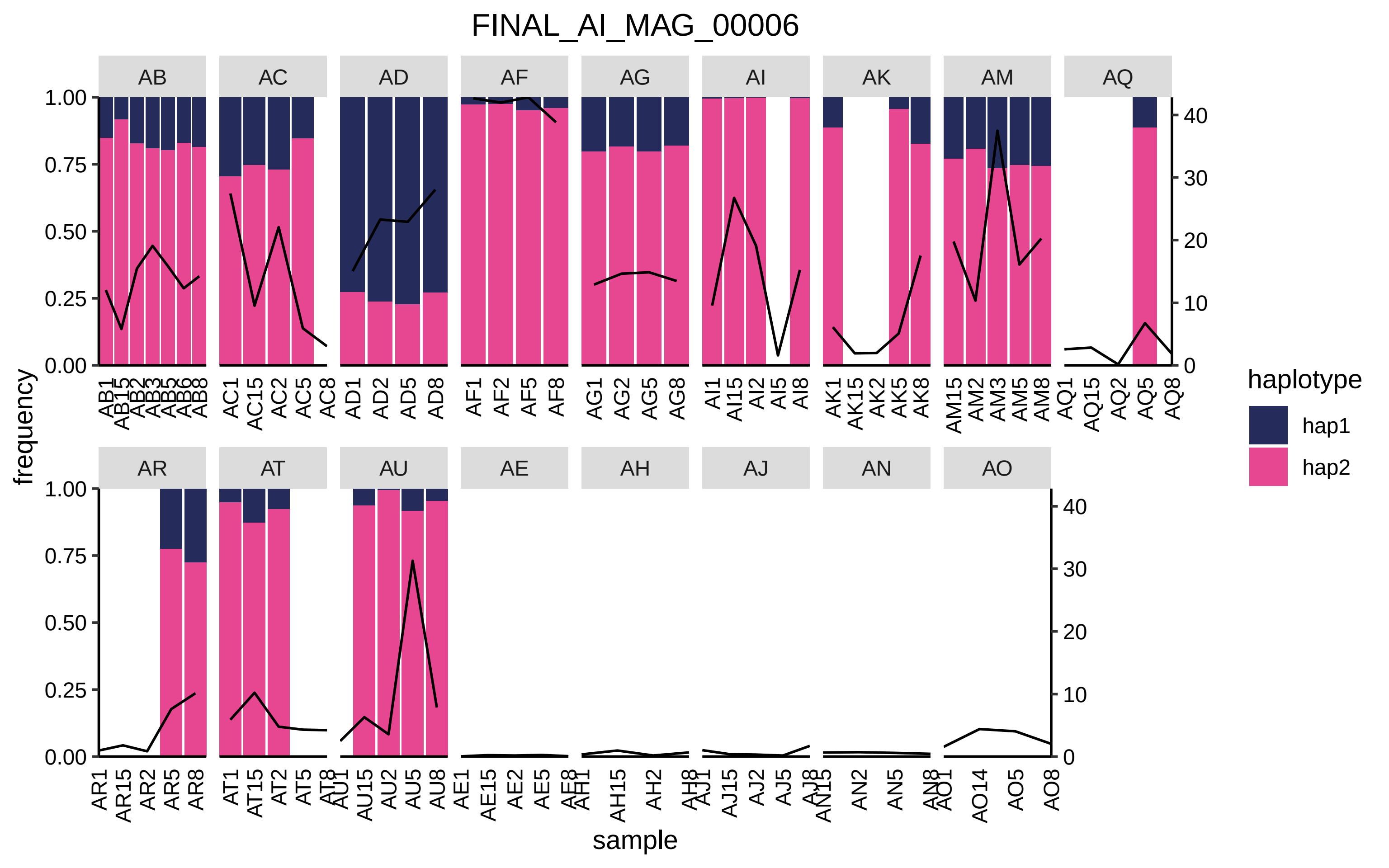
FINAL_AI_MAG_00006
AB
AC
AD
AF
AG
AI
AK
AM
AQ
AR
AT
AU
AE
AH
AJ
AN
AO
frequency
sample
haplotype
hap1
hap2
1.00
0.75
0.50
0.25
0.00
40
30
20
10
0
AB1 AB15 AB2 AB3 AB5 AB6 AB8
AC1 AC15 AC2 AC5 AC8
AD1 AD2 AD5 AD8
AF1 AF2 AF5 AF8
AG1 AG2 AG5 AG8
AI1 AI15 AI2 AI5 AI8
AK1 AK15 AK2 AK5 AK8
AM15 AM2 AM3 AM5 AM8
AQ1 AQ15 AQ2 AQ5 AQ8
AR1 AR15 AR2 AR5 AR8
AT1 AT15 AT2 AT5 AT8
AU1 AU15 AU2 AU5 AU8
AE1 AE15 AE2 AE5 AE8
AH1 AH15 AH2 AH8
AJ1 AJ15 AJ2 AJ5 AJ8
AN15 AN2 AN5 AN8
AO1 AO14 AO5 AO8

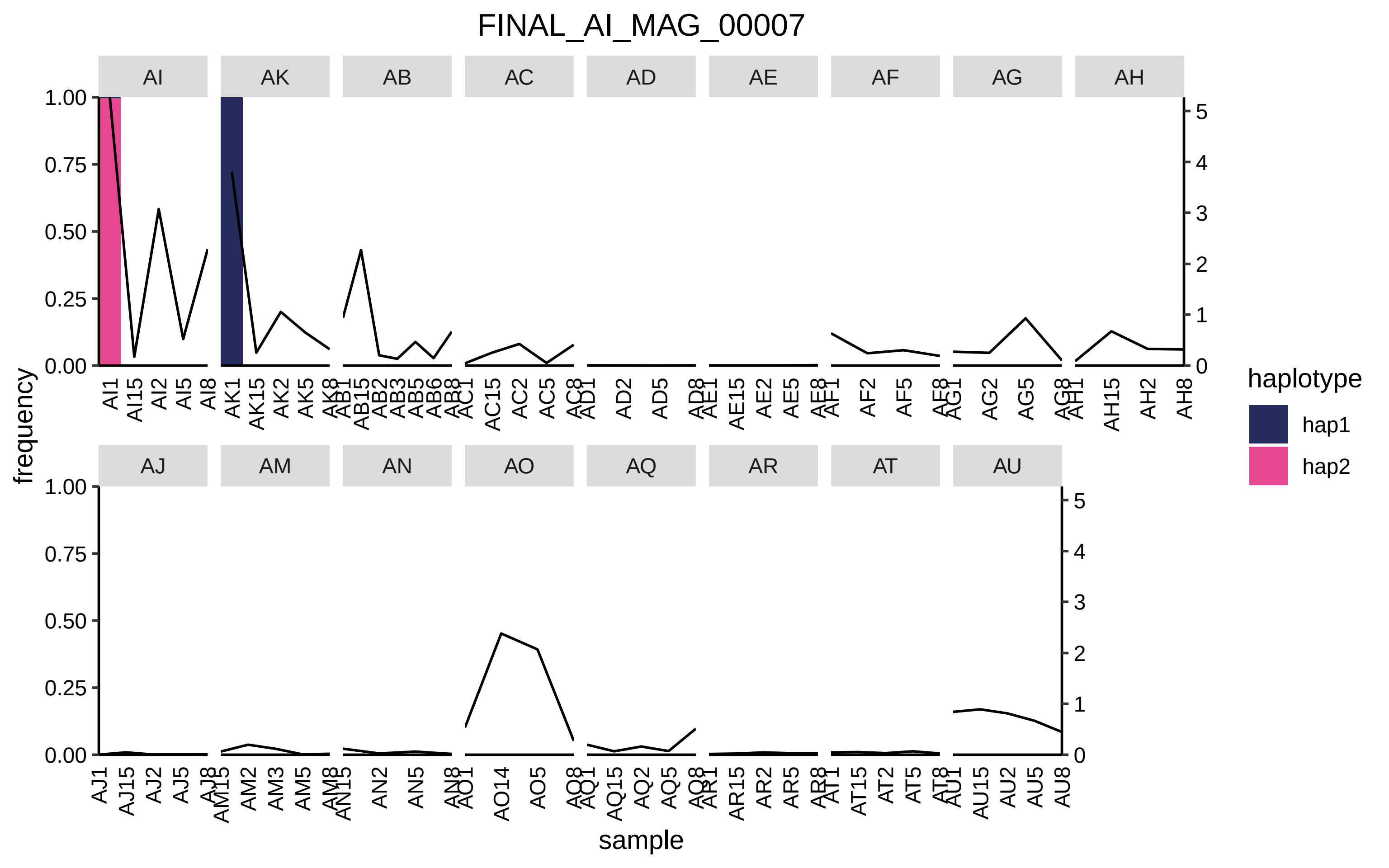

FINAL_AI_MAG_00007
AI
AK
AB
AC
AD
AE
AF
AG
AH
AJ
AM
AN
AO
AQ
AR
AT
AU
frequency
sample
haplotype
hap1
hap2

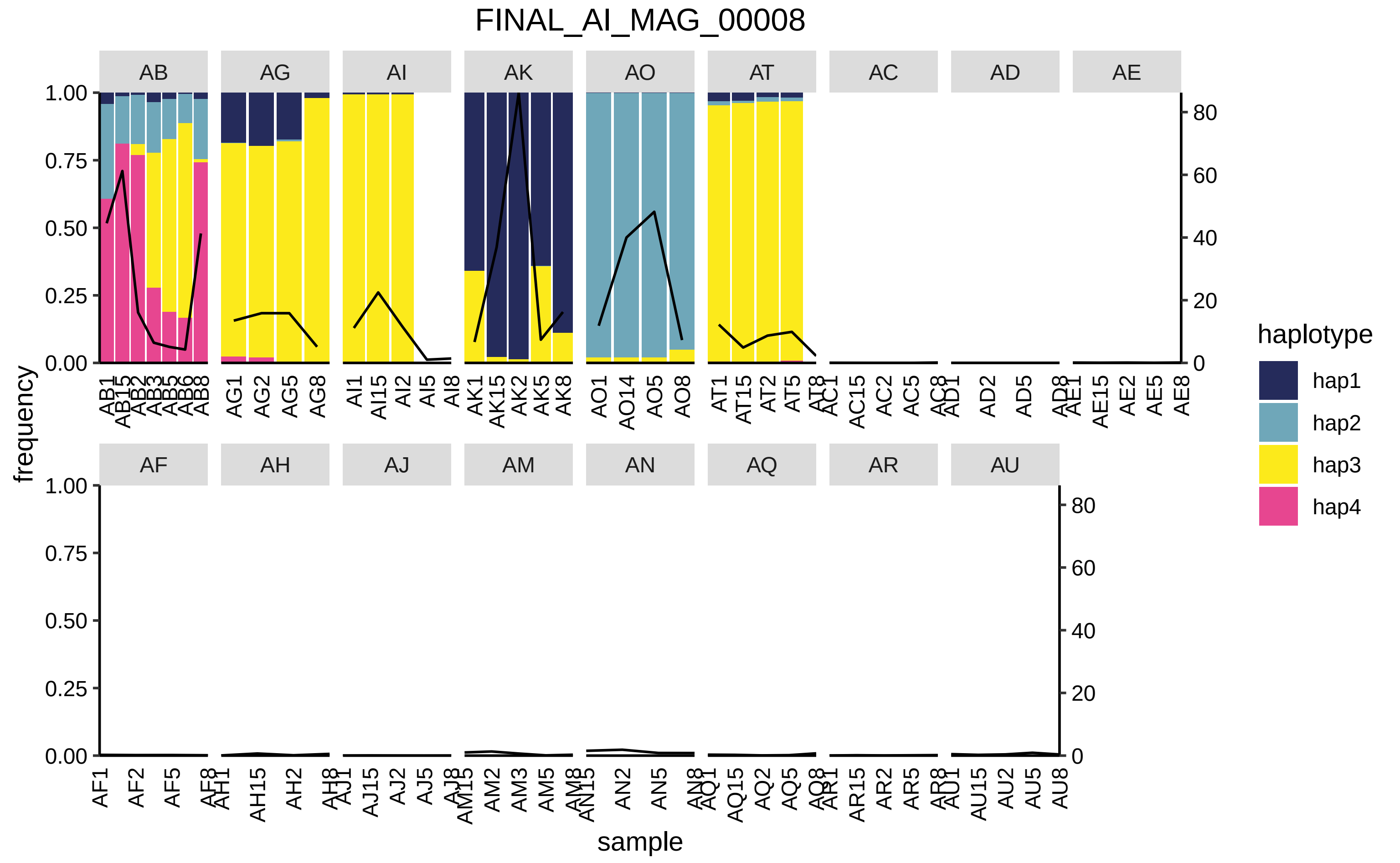
FINAL_AI_MAG_00008
AB
AG
AI
AK
AO
AT
AC
AD
AE
AF
AH
AJ
AM
AN
AQ
AR
AU
frequency
sample
haplotype
hap1
hap2
hap3
hap4

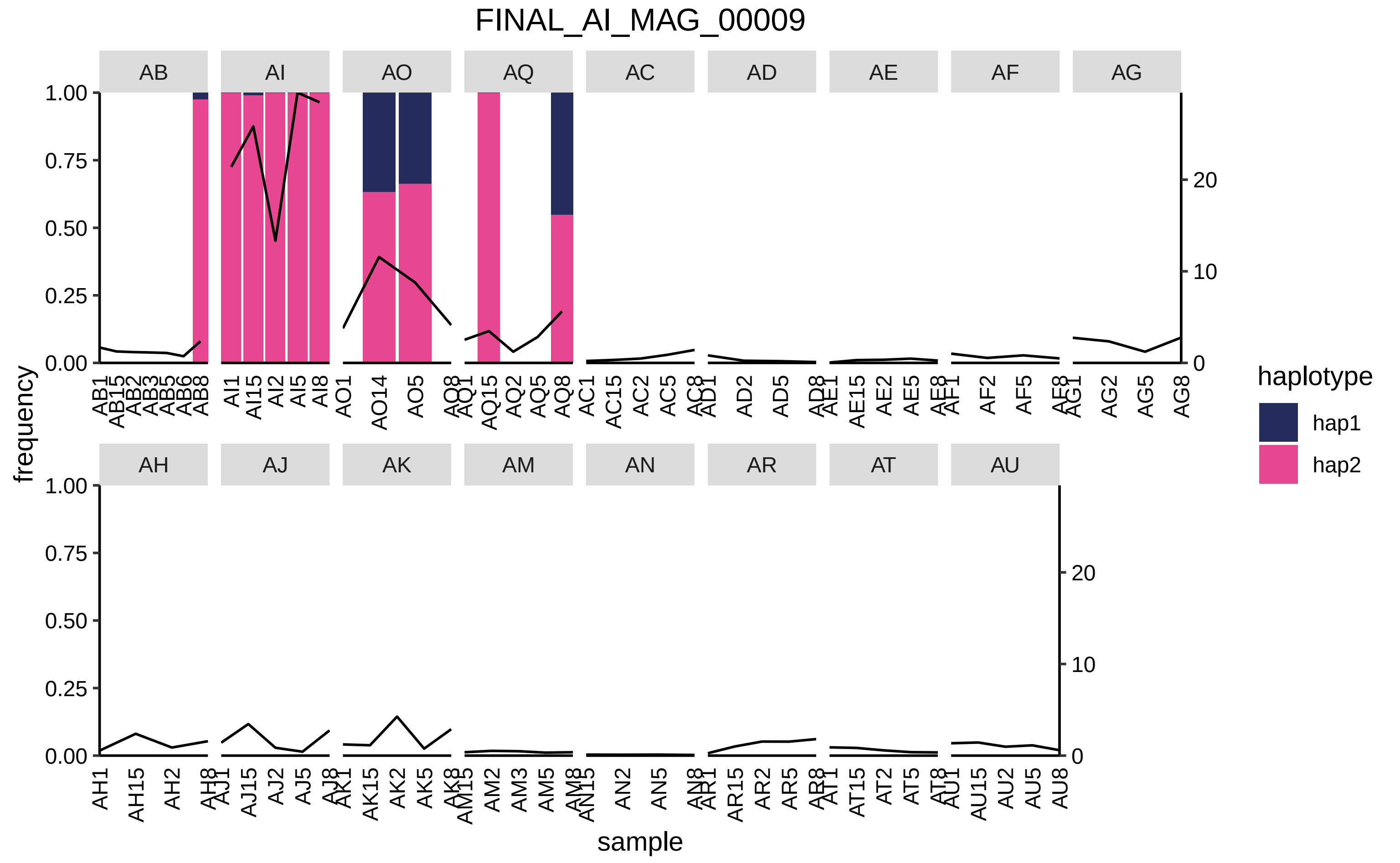
FINAL_AI_MAG_00009
AB
AI
AO
AQ
AC
AD
AE
AF
AG
AH
AJ
AK
AM
AN
AR
AT
AU
frequency
sample
haplotype
hap1
hap2
1.00
0.75
0.50
0.25
0.00
20
10
0

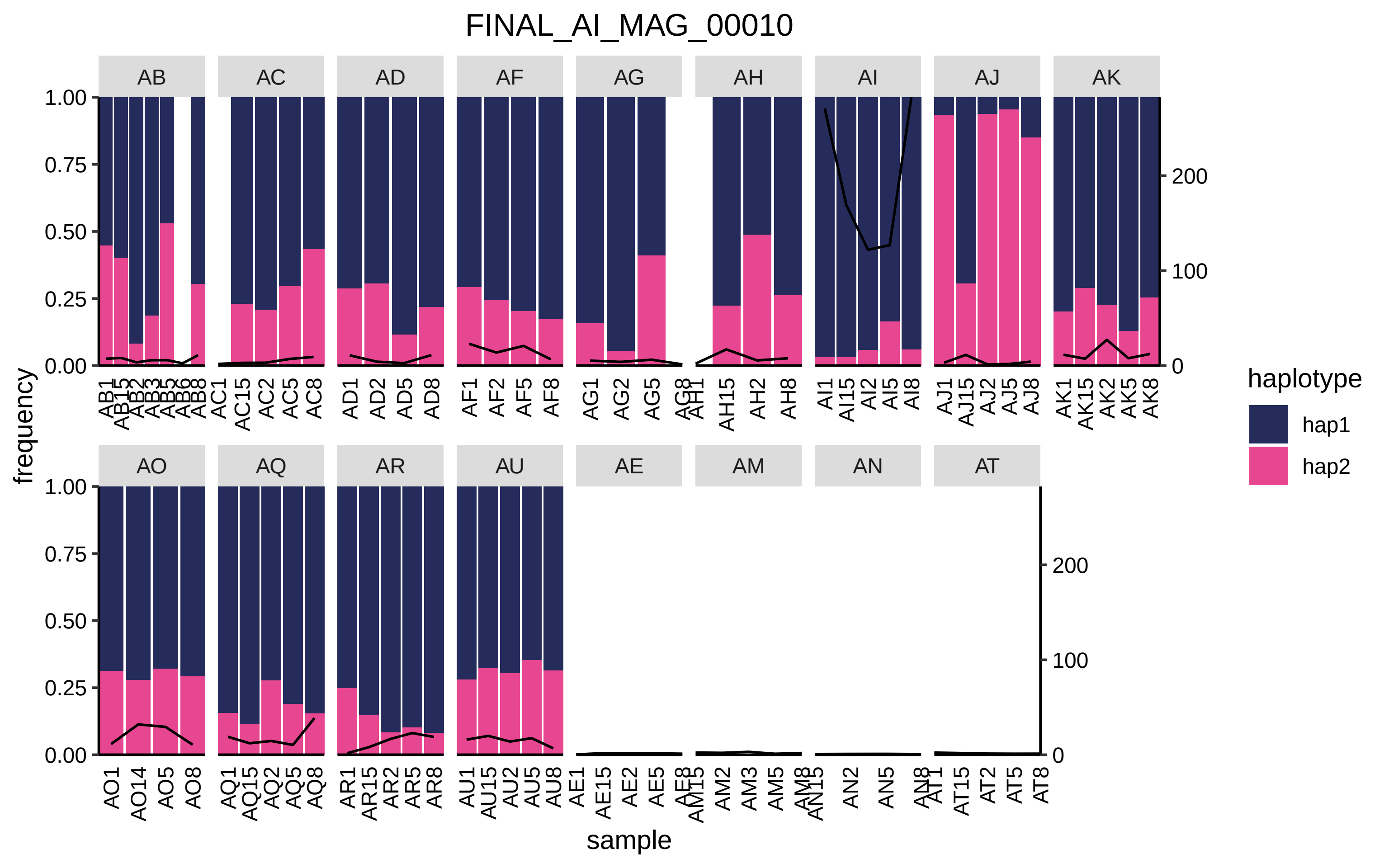

FINAL_AI_MAG_00010
AB
AC
AD
AF
AG
AH
AI
AJ
AK
AO
AQ
AR
AU
AE
AM
AN
AT
frequency
1.00
0.75
0.50
0.25
0.00
200
100
0
sample
haplotype
hap1
hap2
AB1 AB15 AB2 AB3 AB5 AB6 AB8
AC1 AC15 AC2 AC5 AC8
AD1 AD2 AD5 AD8
AF1 AF2 AF5 AF8
AG1 AG2 AG5 AG8
AH1 AH15 AH2 AH8
AI1 AI15 AI2 AI5 AI8
AJ1 AJ15 AJ2 AJ5 AJ8
AK1 AK15 AK2 AK5 AK8
AO1 AO14 AO5 AO8
AQ1 AQ15 AQ2 AQ5 AQ8
AR1 AR15 AR2 AR5 AR8
AU1 AU15 AU2 AU5 AU8
AE1 AE15 AE2 AE5 AE8
AM15 AM2 AM3 AM5 AM8
AN15 AN2 AN5 AN8
AT1 AT15 AT2 AT5 AT8

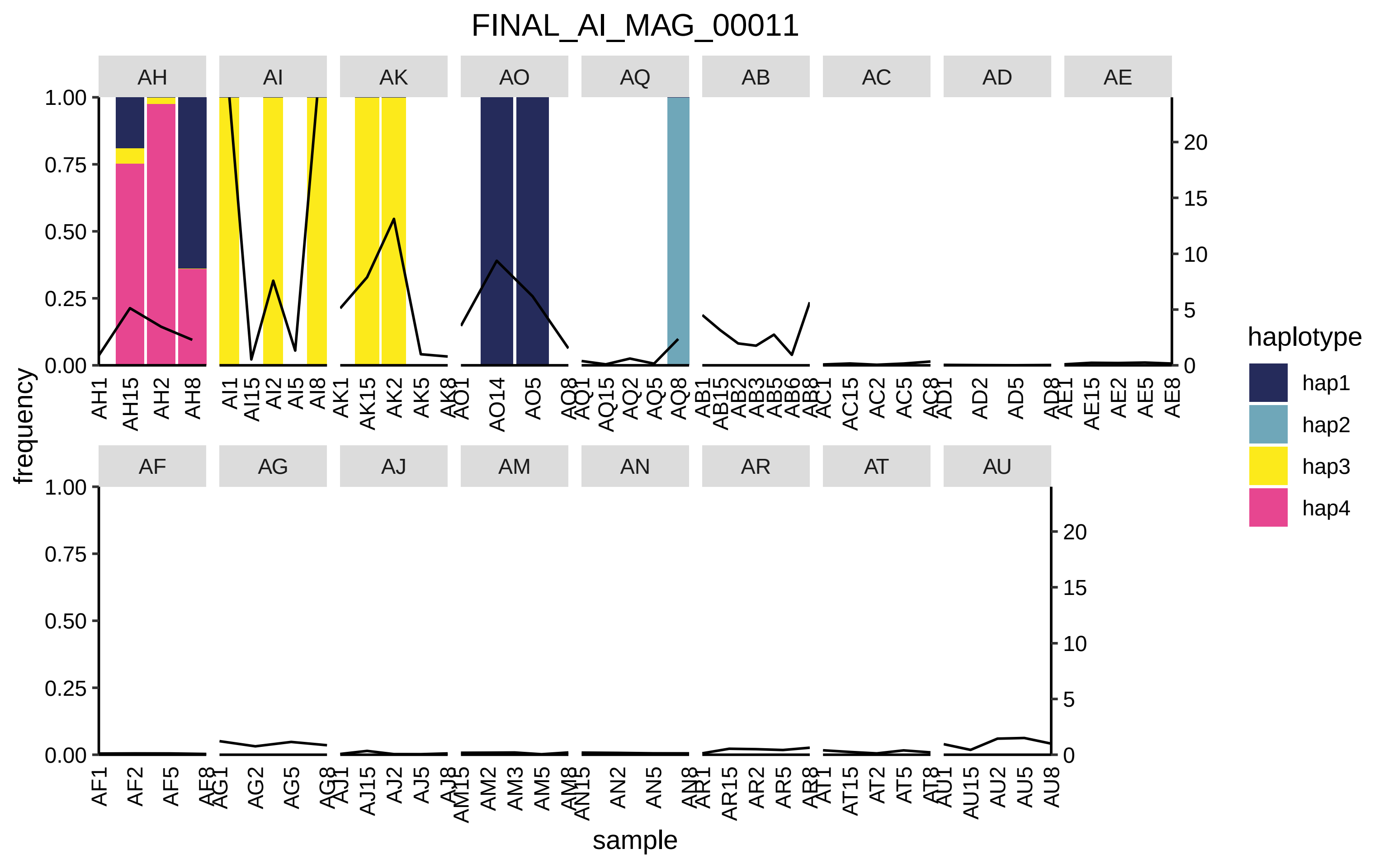
FINAL_AI_MAG_00011
AH
AI
AK
AO
AQ
AB
AC
AD
AE
AF
AG
AJ
AM
AN
AR
AT
AU
frequency
sample
haplotype
hap1
hap2
hap3
hap4

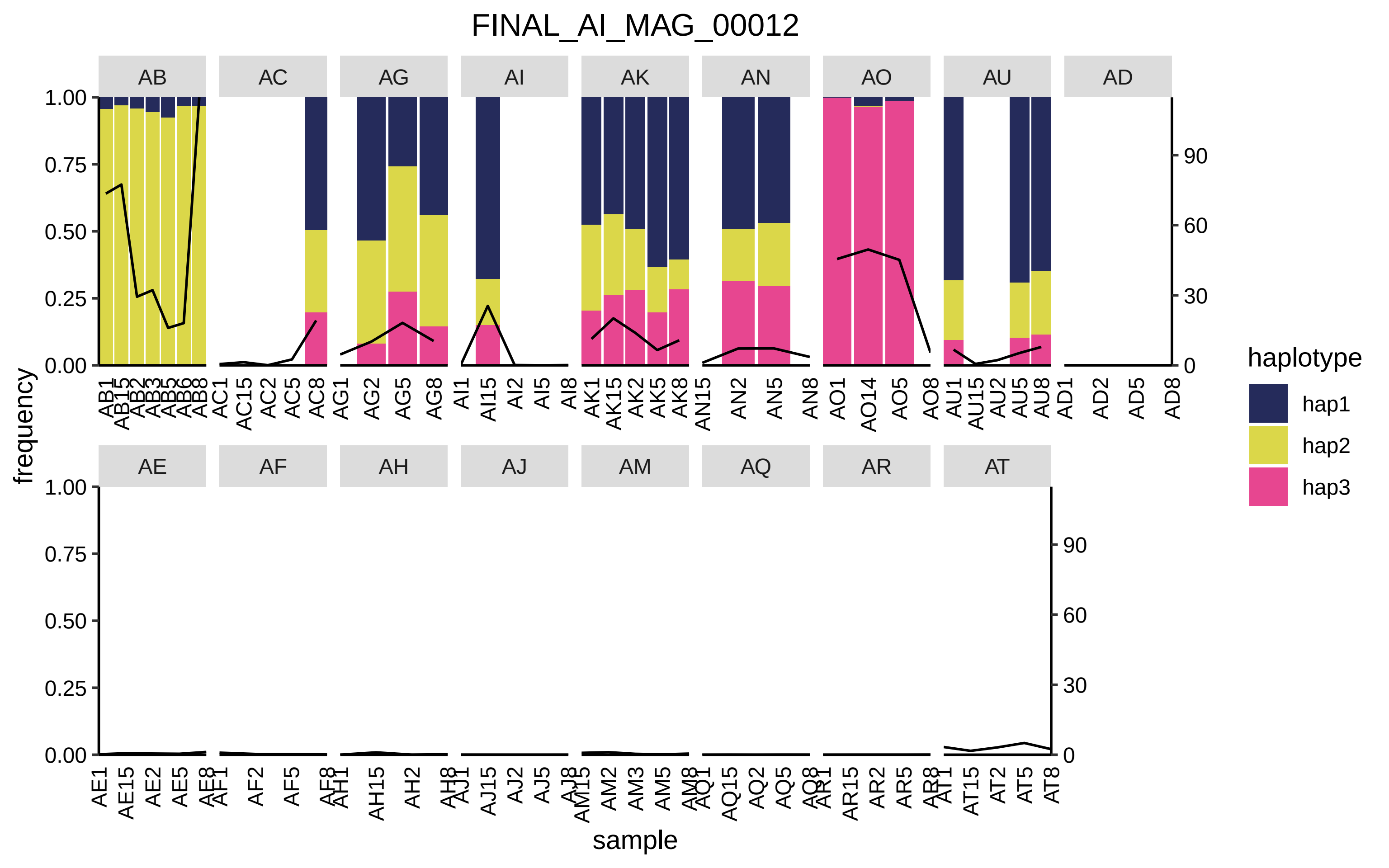

FINAL_AI_MAG_00012
AB
AC
AG
AI
AK
AN
AO
AU
AD
AE
AF
AH
AJ
AM
AQ
AR
AT
frequency
sample
haplotype
hap1
hap2
hap3

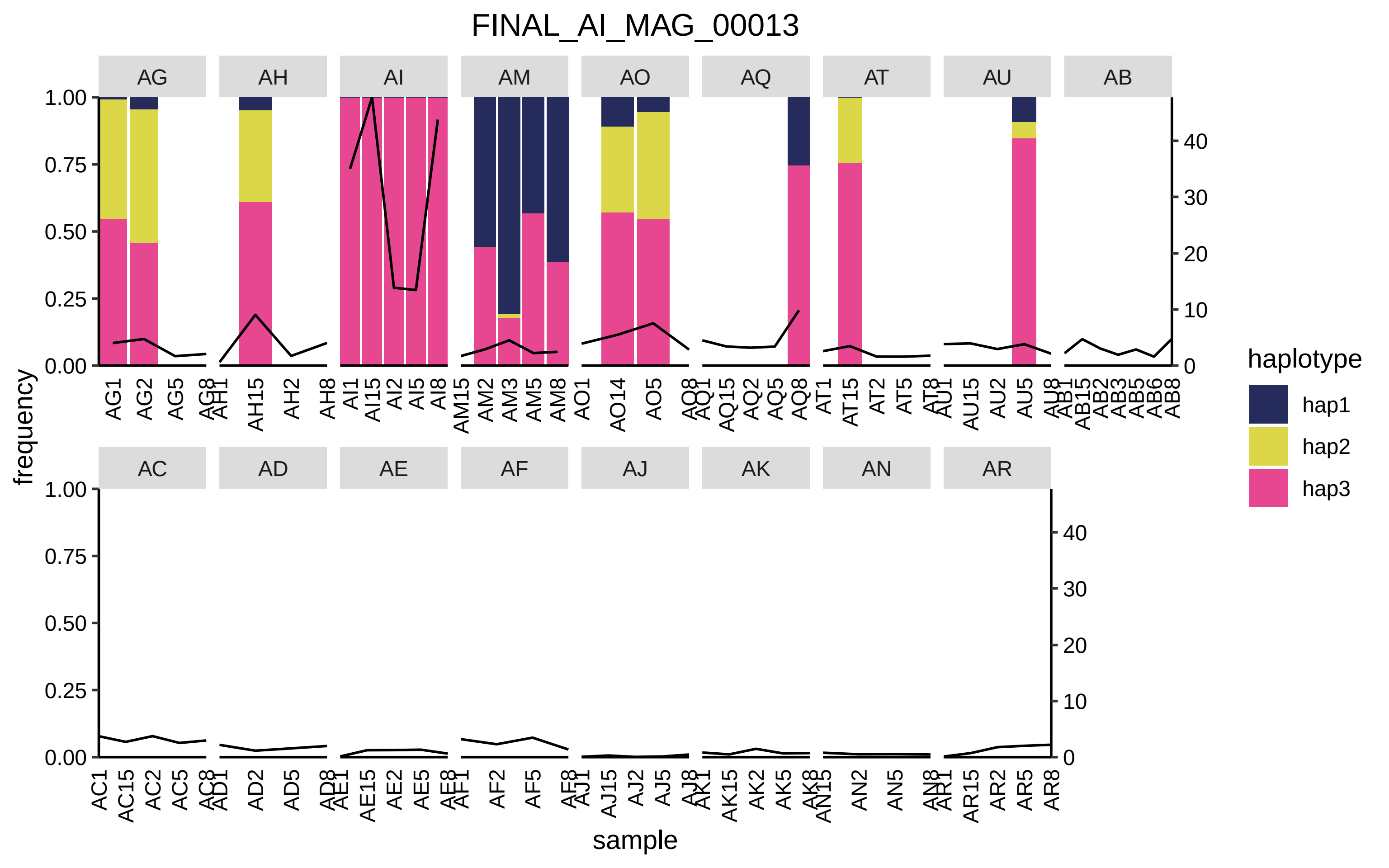
FINAL_AI_MAG_00013
AG
AH
AI
AM
AO
AQ
AT
AU
AB
AC
AD
AE
AF
AJ
AK
AN
AR
frequency
sample
haplotype
hap1
hap2
hap3

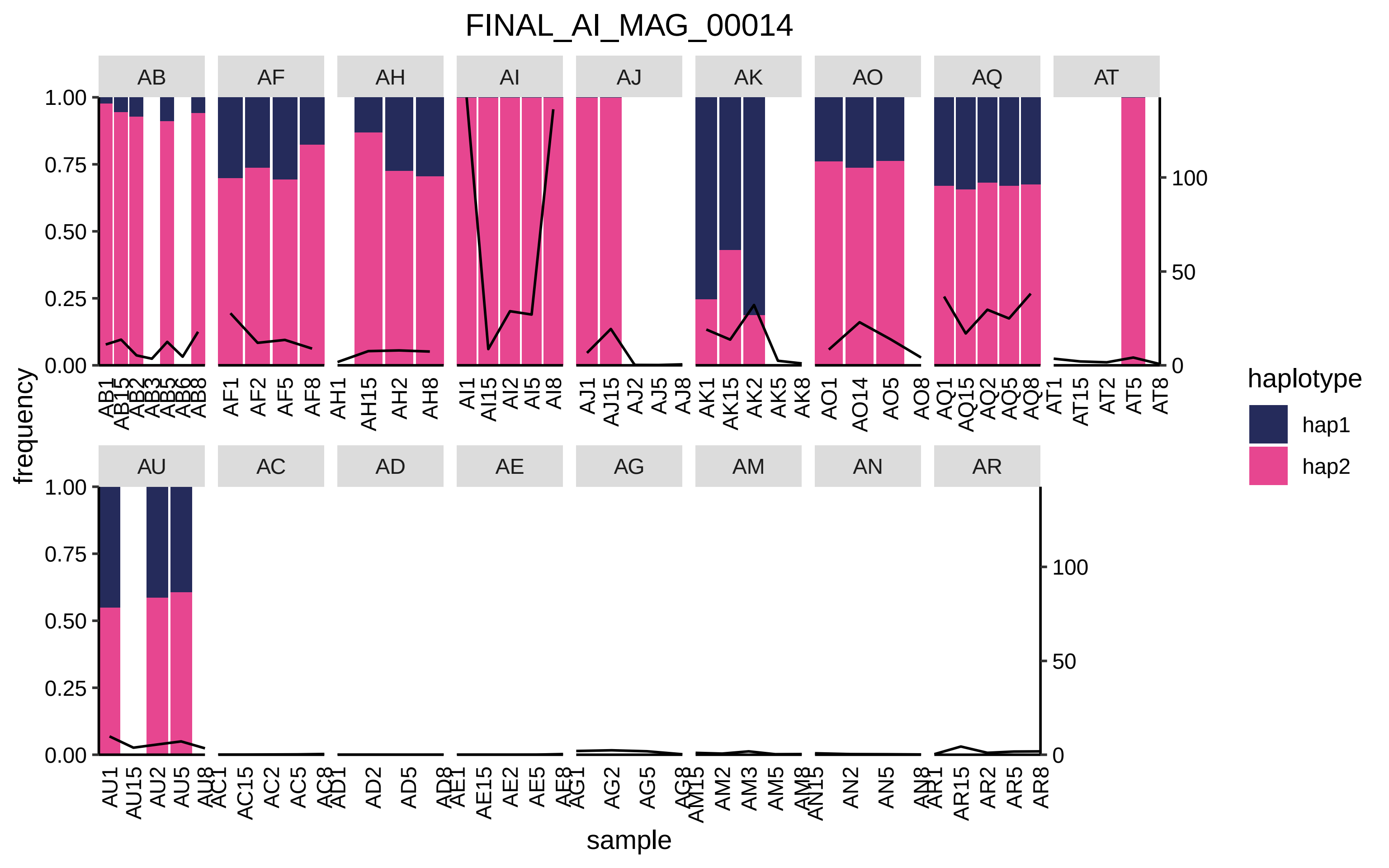

FINAL_AI_MAG_00014
AB
AF
AH
AI
AJ
AK
AO
AQ
AT
AU
AC
AD
AE
AG
AM
AN
AR
frequency
sample
haplotype
hap1
hap2

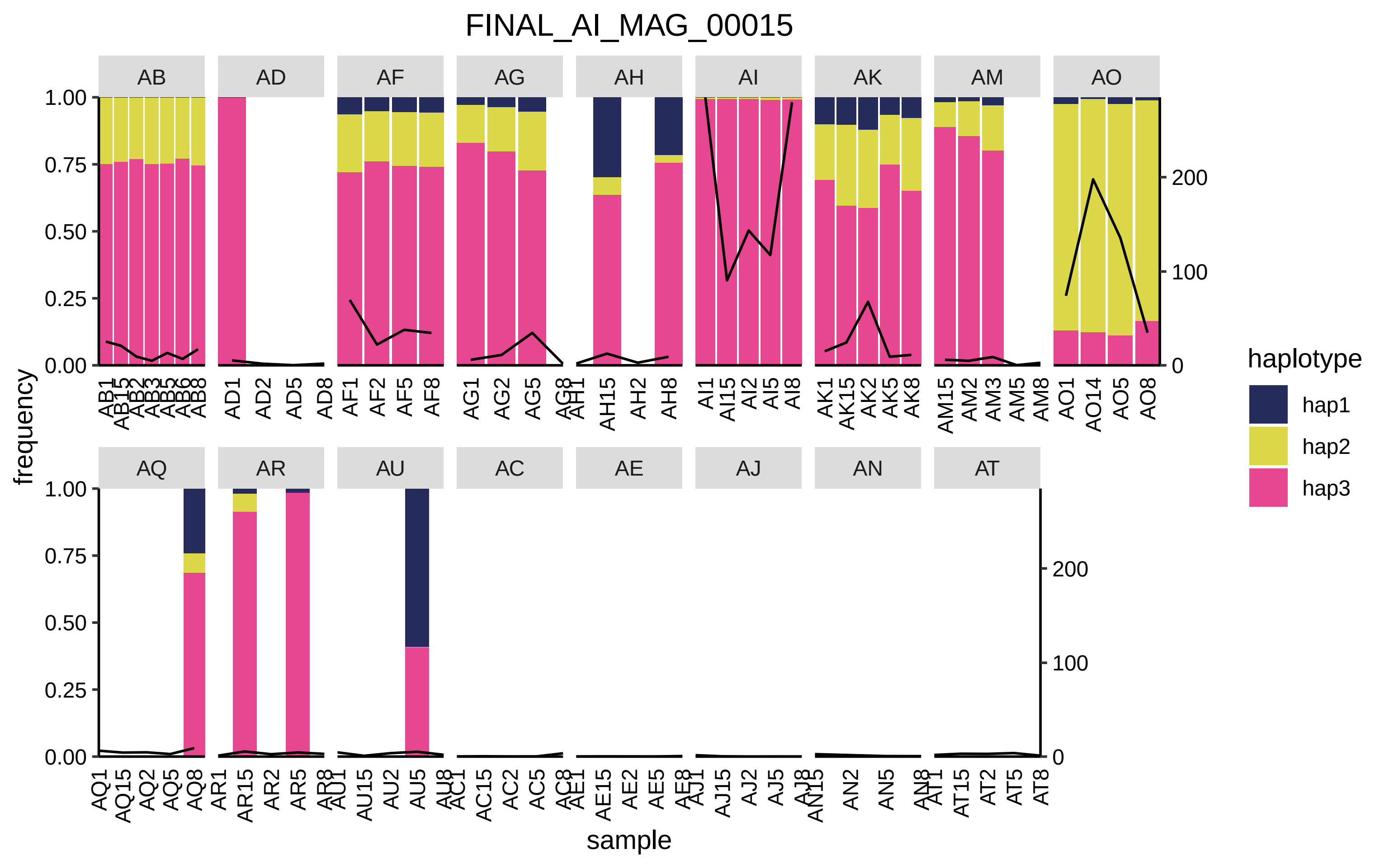
FINAL_AI_MAG_00015
AB
AD
AF
AG
AH
AI
AK
AM
AO
AQ
AR
AU
AC
AE
AJ
AN
AT
frequency
sample
haplotype
hap1
hap2
hap3
1.00
0.75
0.50
0.25
0.00
200
100
0

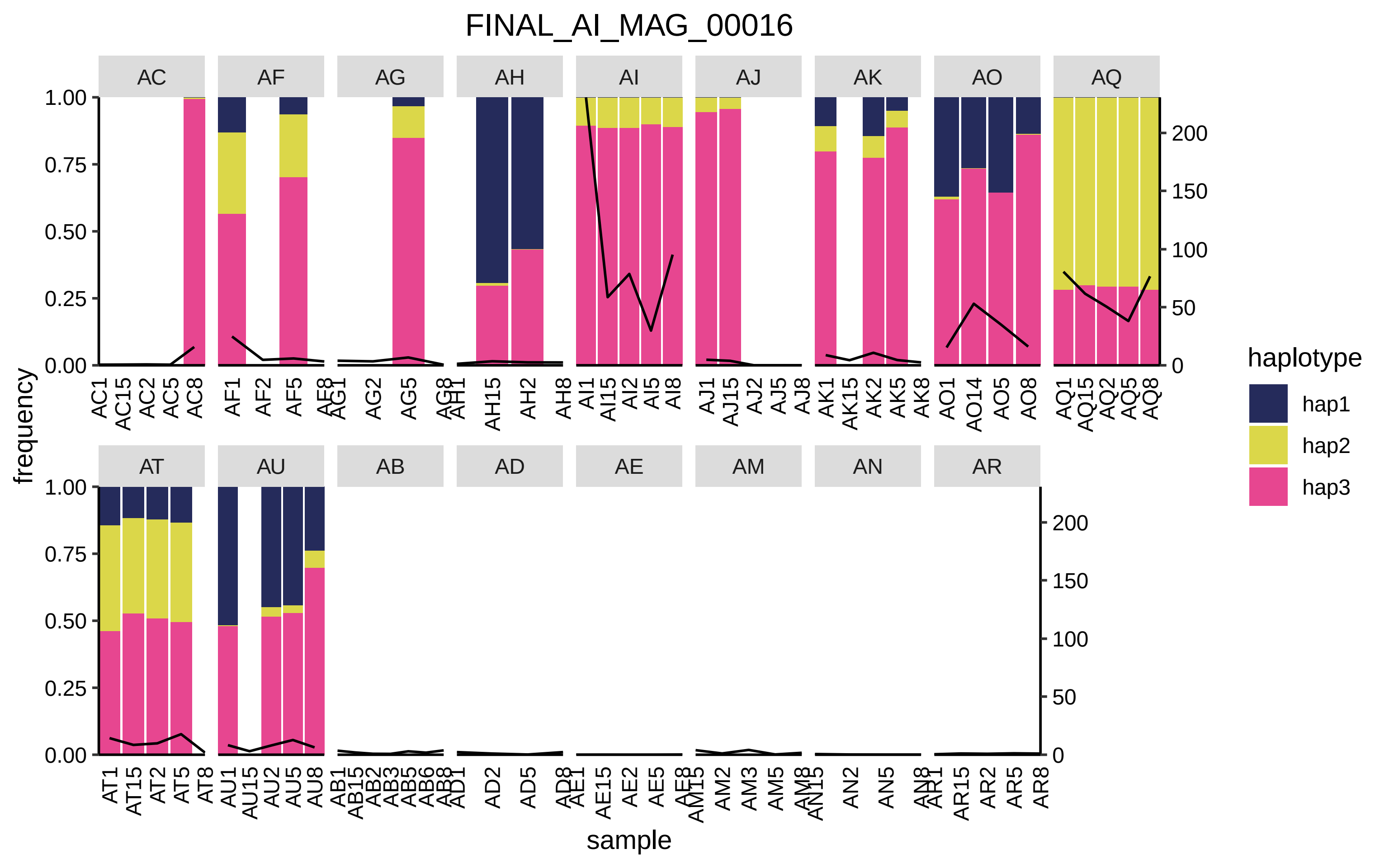

FINAL_AI_MAG_00016
AC
AF
AG
AH
AI
AJ
AK
AO
AQ
AT
AU
AB
AD
AE
AM
AN
AR
1.00
0.75
0.50
0.25
0.00
200
150
100
50
0
frequency
sample
haplotype
hap1
hap2
hap3

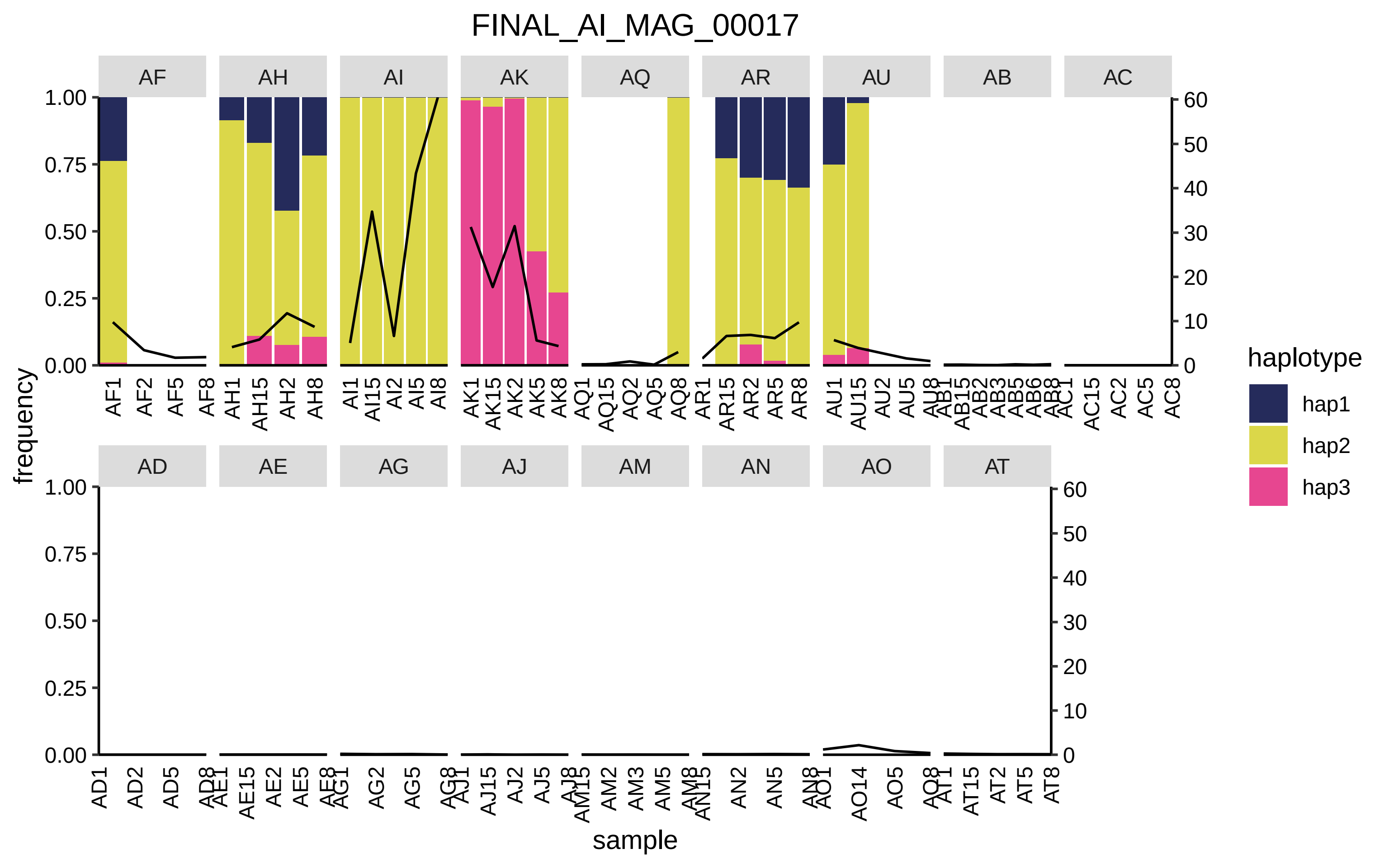

FINAL_AI_MAG_00017
AF
AH
AI
AK
AQ
AR
AU
AB
AC
AD
AE
AG
AJ
AM
AN
AO
AT
frequency
sample
haplotype
hap1
hap2
hap3

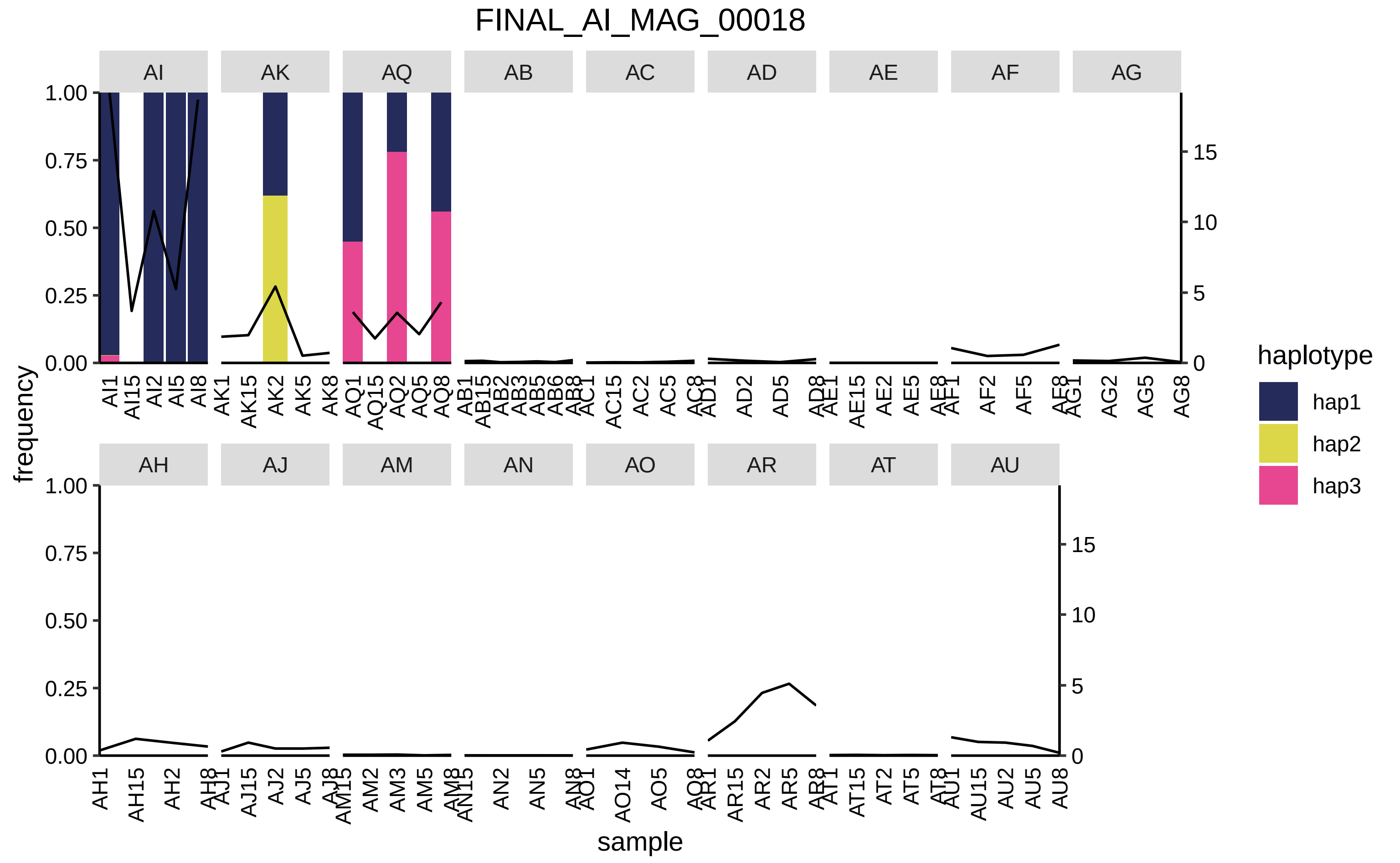
FINAL_AI_MAG_00018
frequency
sample
haplotype
hap1
hap2
hap3
AI
AK
AQ
AB
AC
AD
AE
AF
AG
AH
AJ
AM
AN
AO
AR
AT
AU

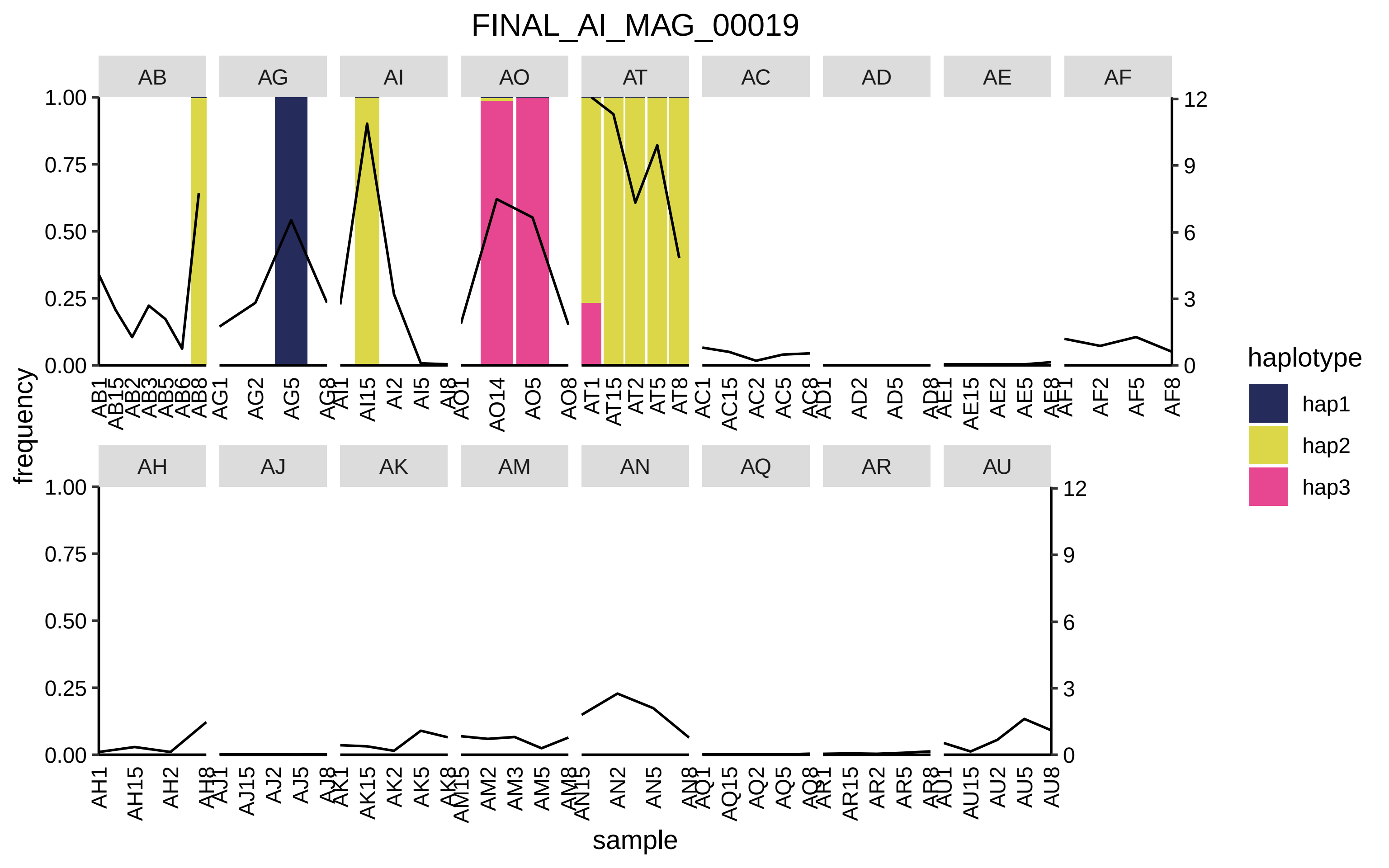

FINAL_AI_MAG_00019
AB
AG
AI
AO
AT
AC
AD
AE
AF
AH
AJ
AK
AM
AN
AQ
AR
AU
frequency
sample
1.00
0.75
0.50
0.25
0.00
12
9
6
3
0
haplotype
hap1
hap2
hap3

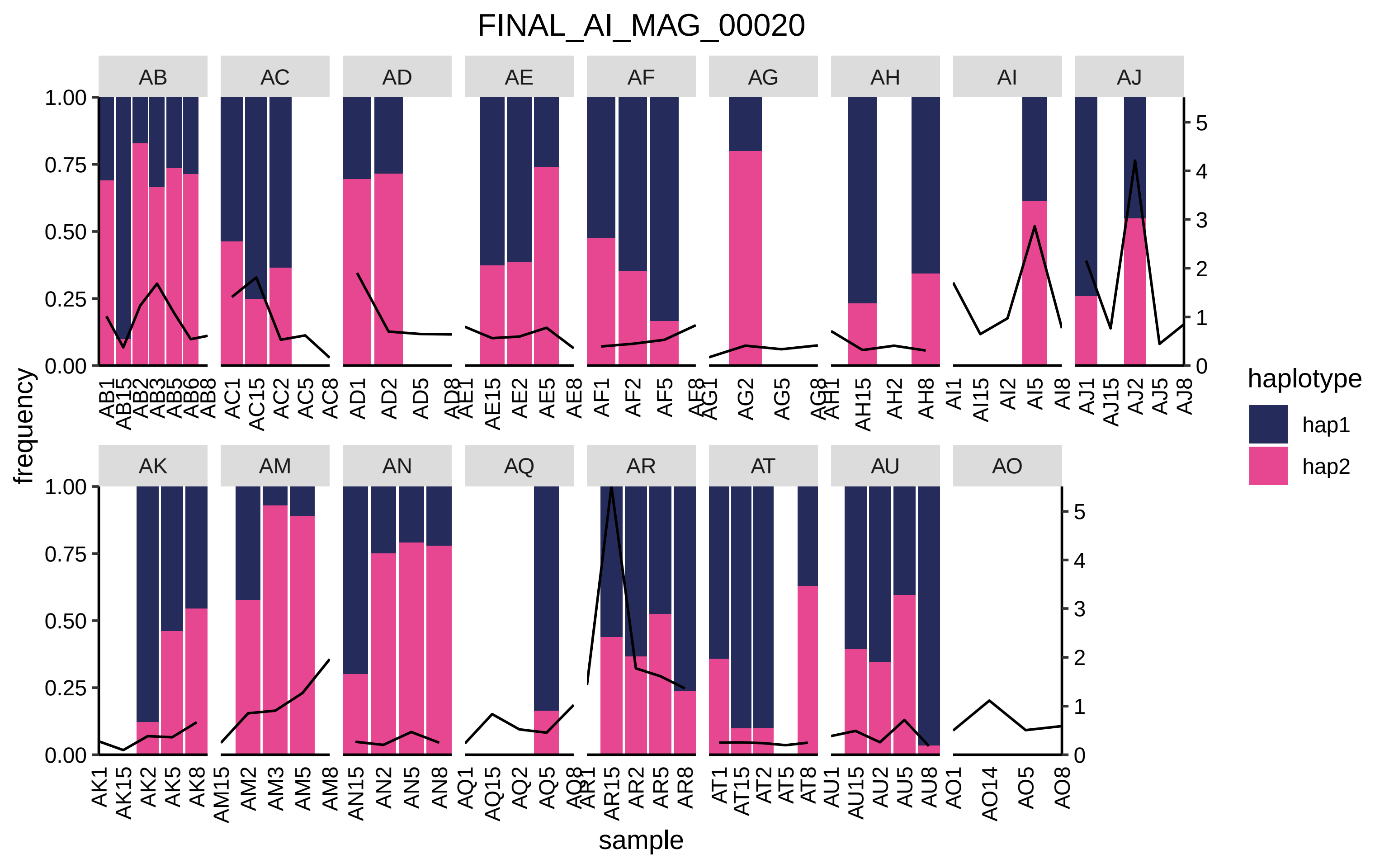

FINAL_AI_MAG_00020
AB
AC
AD
AE
AF
AG
AH
AI
AJ
AK
AM
AN
AQ
AR
AT
AU
AO
frequency
sample
haplotype
hap1
hap2
1.00
0.75
0.50
0.25
0.00
0
1
2
3
4
5
AB1 AB15 AB2 AB3 AB5 AB6 AB8
AC1 AC15 AC2 AC5 AC8
AD1 AD2 AD5 AD8
AE1 AE15 AE2 AE5 AE8
AF1 AF2 AF5 AF8
AG1 AG2 AG5 AG8
AH1 AH15 AH2 AH8
AI1 AI15 AI2 AI5 AI8
AJ1 AJ15 AJ2 AJ5 AJ8
AK1 AK15 AK2 AK5 AK8
AM15 AM2 AM3 AM5 AM8
AN15 AN2 AN5 AN8
AQ1 AQ15 AQ2 AQ5 AQ8
AR1 AR15 AR2 AR5 AR8
AT1 AT15 AT2 AT5 AT8
AU1 AU15 AU2 AU5 AU8
AO1 AO14 AO5 AO8

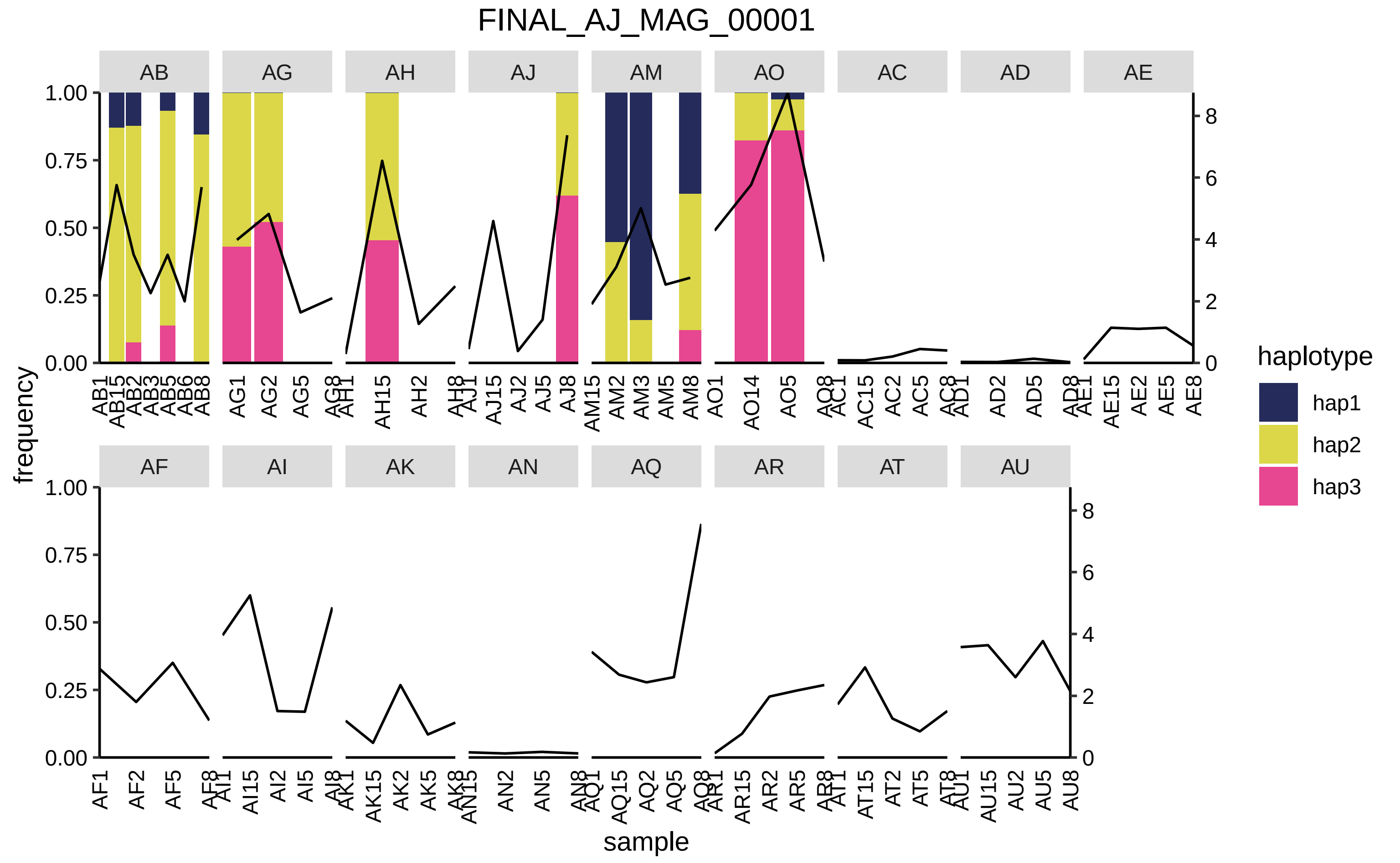

FINAL_AJ_MAG_00001
AB
AG
AH
AJ
AM
AO
AC
AD
AE
AF
AI
AK
AN
AQ
AR
AT
AU
frequency
sample
haplotype
hap1
hap2
hap3

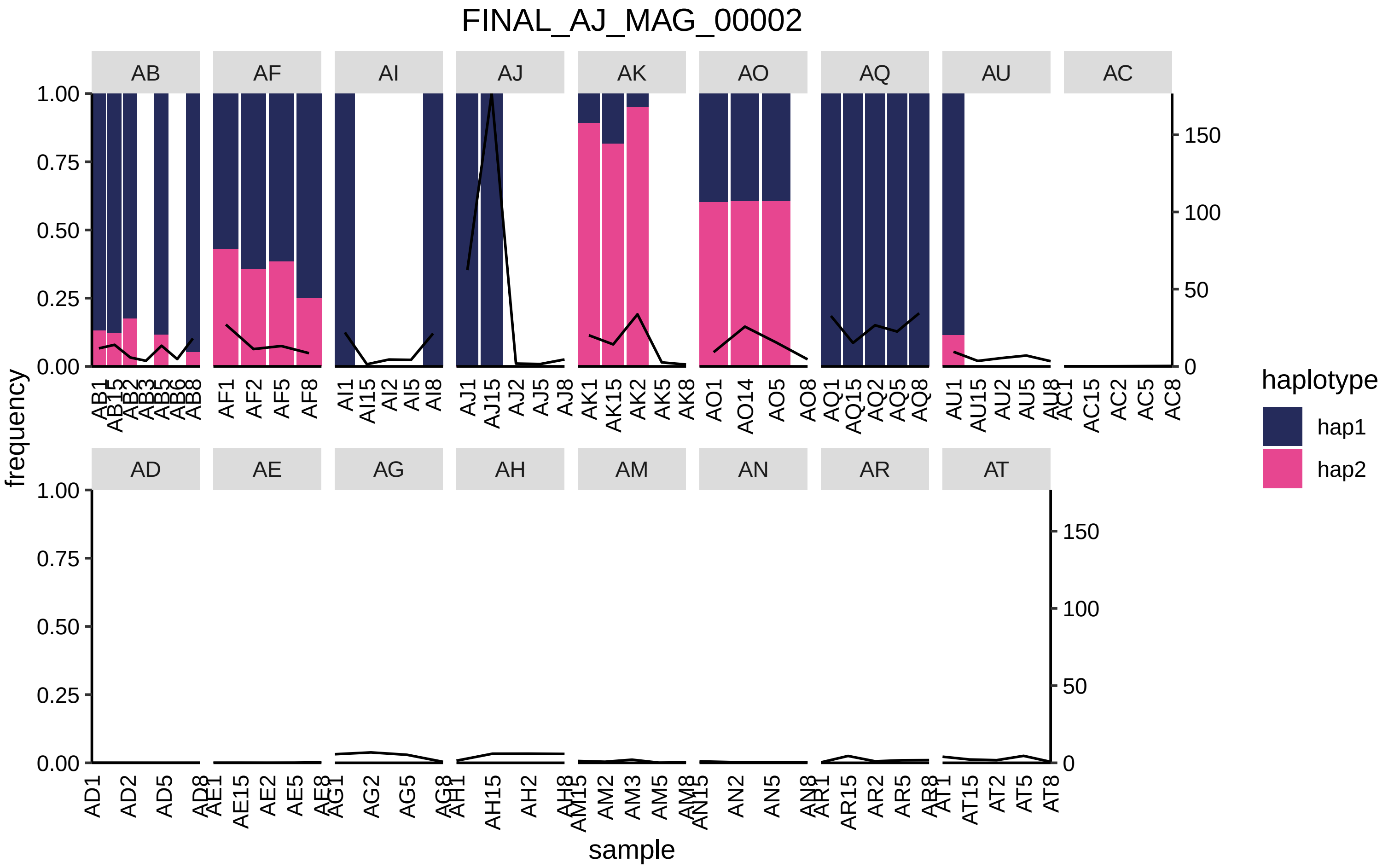
FINAL_AJ_MAG_00002
AB
AF
AI
AJ
AK
AO
AQ
AU
AC
AD
AE
AG
AH
AM
AN
AR
AT
frequency
sample
haplotype
hap1
hap2
1.00
0.75
0.50
0.25
0.00
150
100
50
0

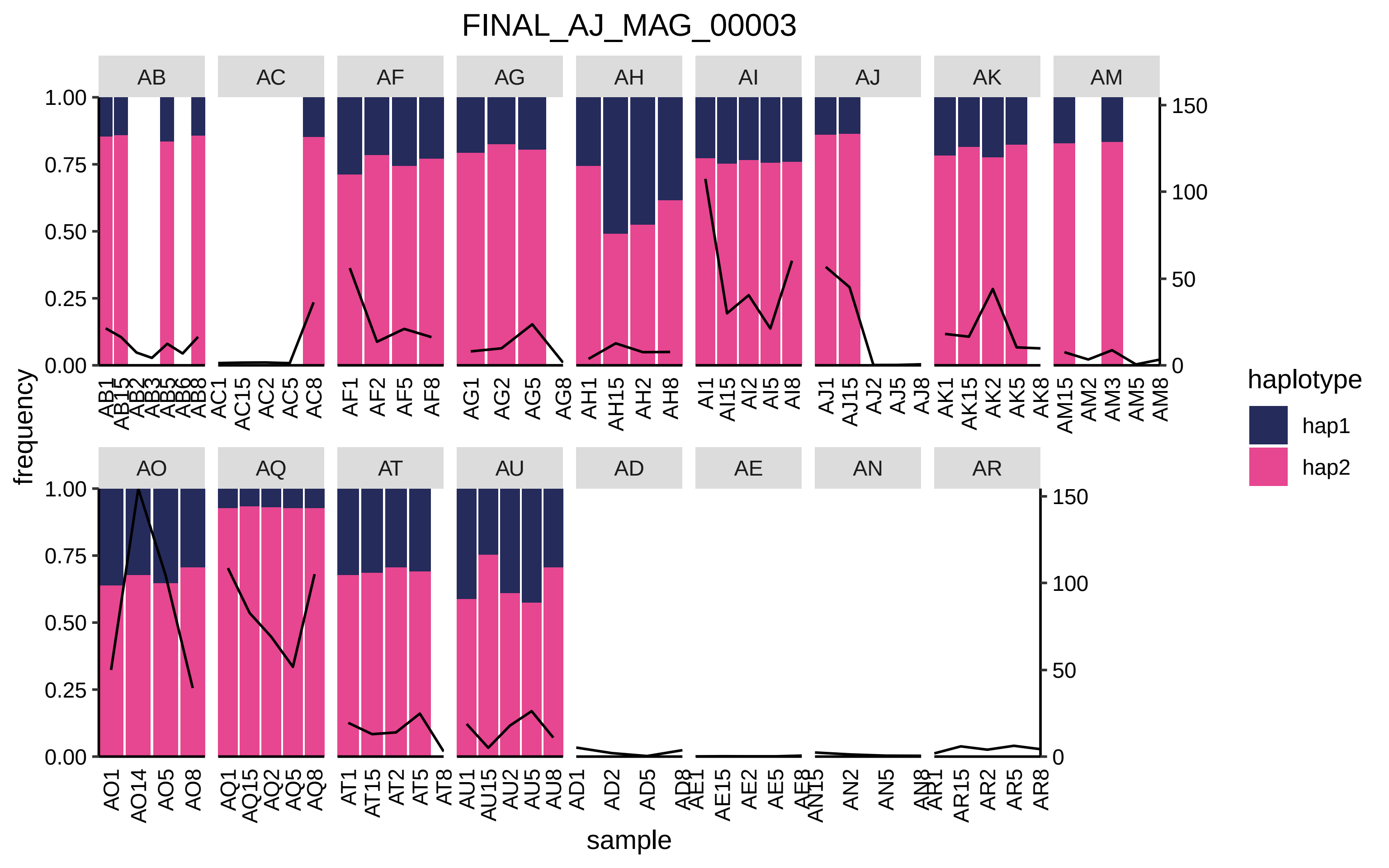

FINAL_AJ_MAG_00003
AB
AC
AF
AG
AH
AI
AJ
AK
AM
AO
AQ
AT
AU
AD
AE
AN
AR
frequency
sample
haplotype
hap1
hap2

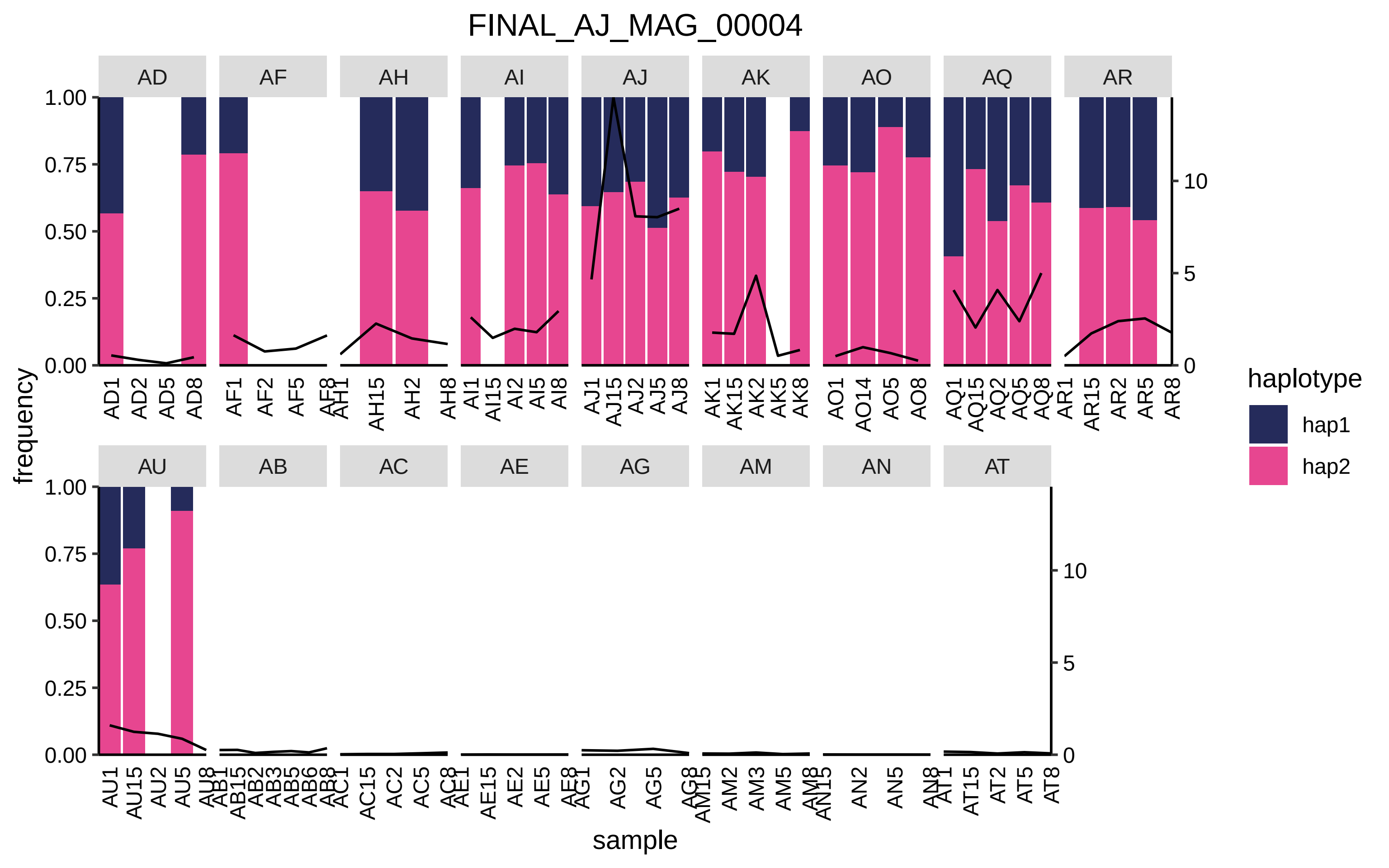

FINAL_AJ_MAG_00004
AD
AF
AH
AI
AJ
AK
AO
AQ
AR
AU
AB
AC
AE
AG
AM
AN
AT
frequency
sample
1.00
0.75
0.50
0.25
0.00
10
5
0
haplotype
hap1
hap2
AD1 AD2 AD5 AD8
AF1 AF2 AF5 AF8
AH1 AH15 AH2 AH8
AI1 AI15 AI2 AI5 AI8
AJ1 AJ15 AJ2 AJ5 AJ8
AK1 AK15 AK2 AK5 AK8
AO1 AO14 AO5 AO8
AQ1 AQ15 AQ2 AQ5 AQ8
AR1 AR15 AR2 AR5 AR8
AU1 AU15 AU2 AU5 AU8
AB1 AB15 AB2 AB3 AB5 AB6 AB8
AC1 AC15 AC2 AC5 AC8
AE1 AE15 AE2 AE5 AE8
AG1 AG2 AG5 AG8
AM15 AM2 AM3 AM5 AM8
AN15 AN2 AN5 AN8
AT1 AT15 AT2 AT5 AT8

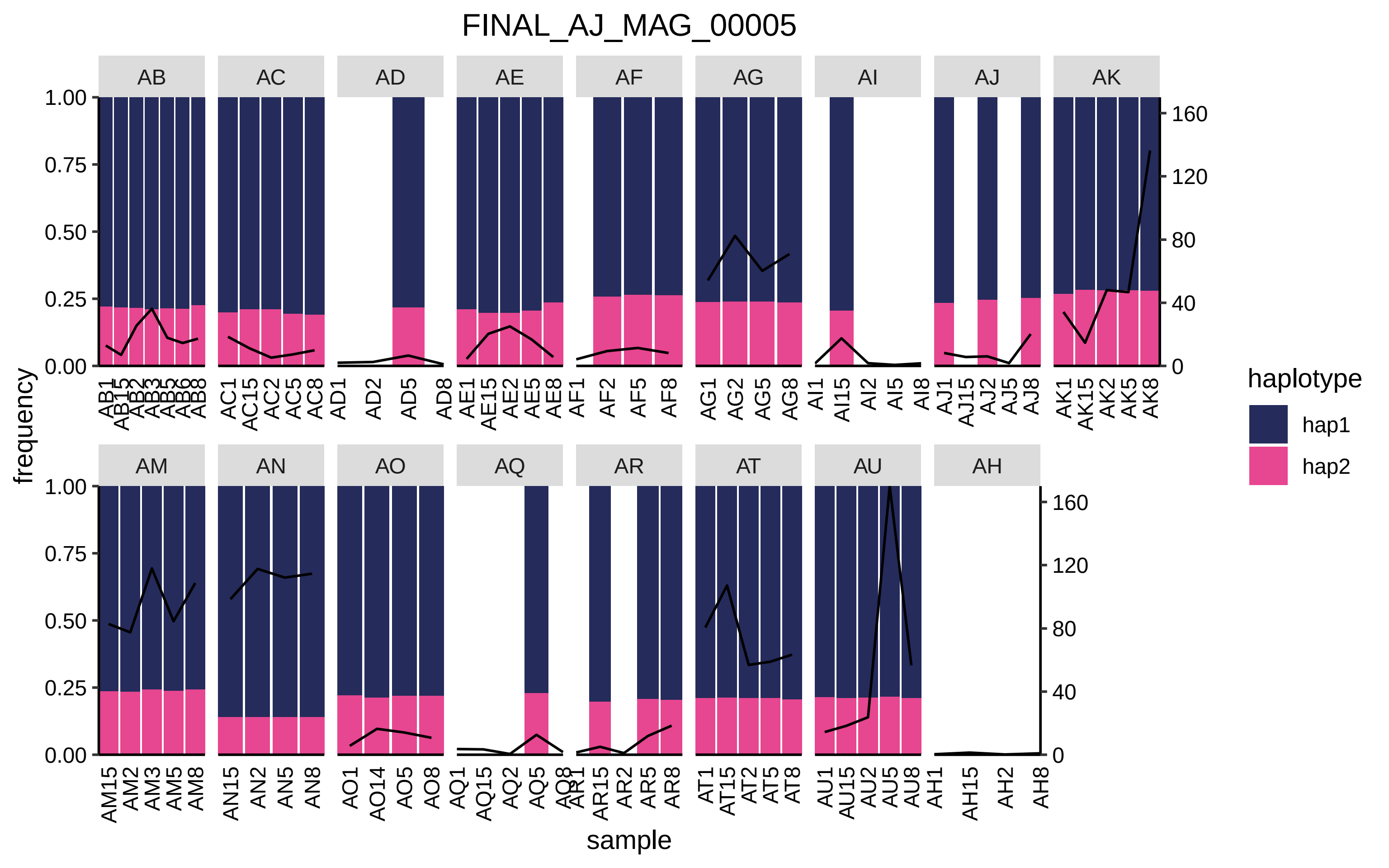
FINAL_AJ_MAG_00005
AB
AC
AD
AE
AF
AG
AI
AJ
AK
AM
AN
AO
AQ
AR
AT
AU
AH
frequency
1.00
0.75
0.50
0.25
0.00
160
120
80
40
0
sample
haplotype
hap1
hap2
AB1 AB15 AB2 AB3 AB5 AB6 AB8
AC1 AC15 AC2 AC5 AC8
AD1 AD2 AD5 AD8
AE1 AE15 AE2 AE5 AE8
AF1 AF2 AF5 AF8
AG1 AG2 AG5 AG8
AI1 AI15 AI2 AI5 AI8
AJ1 AJ15 AJ2 AJ5 AJ8
AK1 AK15 AK2 AK5 AK8
AM15 AM2 AM3 AM5 AM8
AN15 AN2 AN5 AN8
AO1 AO14 AO5 AO8
AQ1 AQ15 AQ2 AQ5 AQ8
AR1 AR15 AR2 AR5 AR8
AT1 AT15 AT2 AT5 AT8
AU1 AU15 AU2 AU5 AU8
AH1 AH15 AH2 AH8

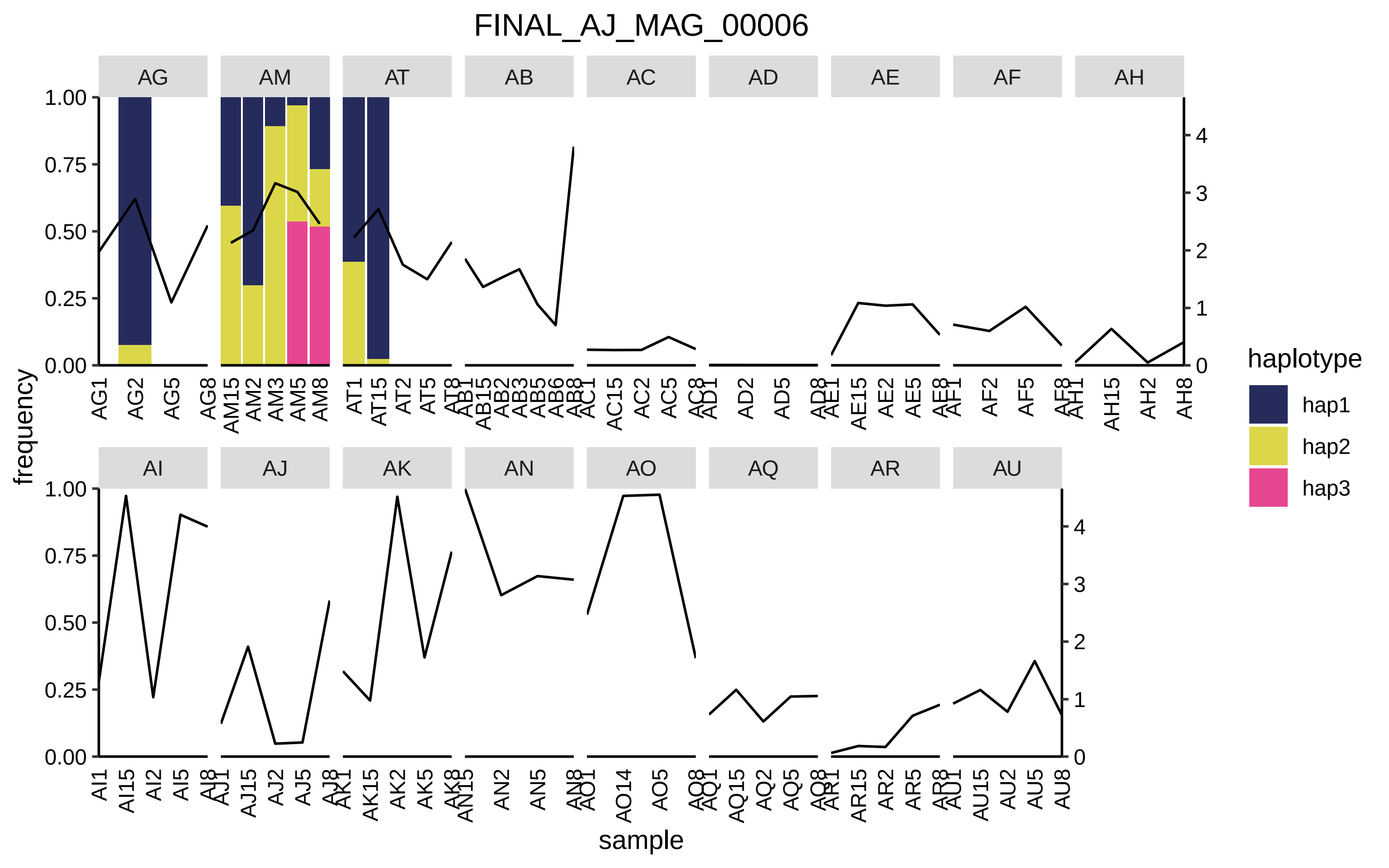
FINAL_AJ_MAG_00006
AG
AM
AT
AB
AC
AD
AE
AF
AH
AI
AJ
AK
AN
AO
AQ
AR
AU
frequency
sample
haplotype
hap1
hap2
hap3

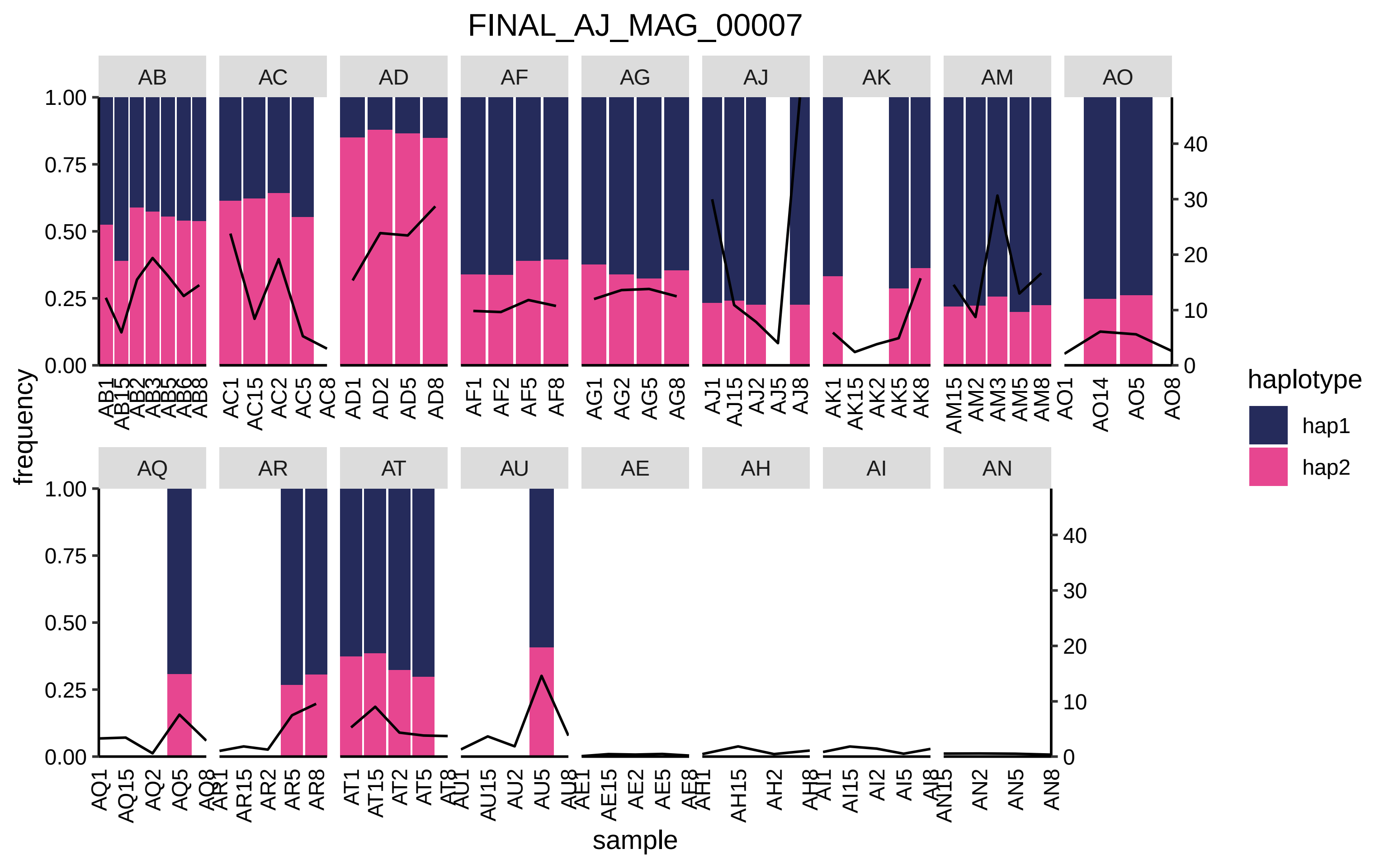

FINAL_AJ_MAG_00007
AB
AC
AD
AF
AG
AJ
AK
AM
AO
AQ
AR
AT
AU
AE
AH
AI
AN
frequency
sample
haplotype
hap1
hap2

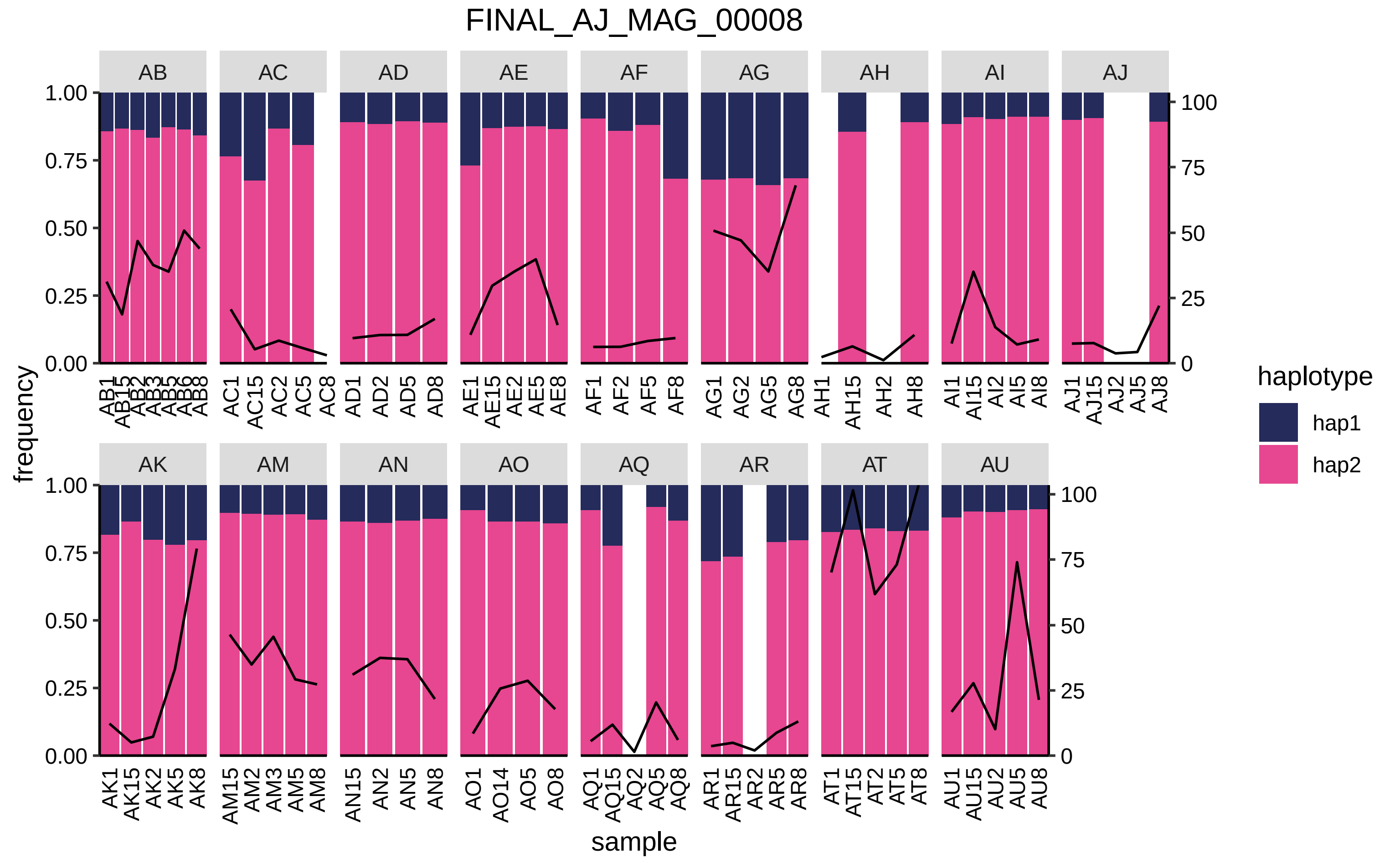
FINAL_AJ_MAG_00008
AB
AC
AD
AE
AF
AG
AH
AI
AJ
AK
AM
AN
AO
AQ
AR
AT
AU
frequency
sample
haplotype
hap1
hap2

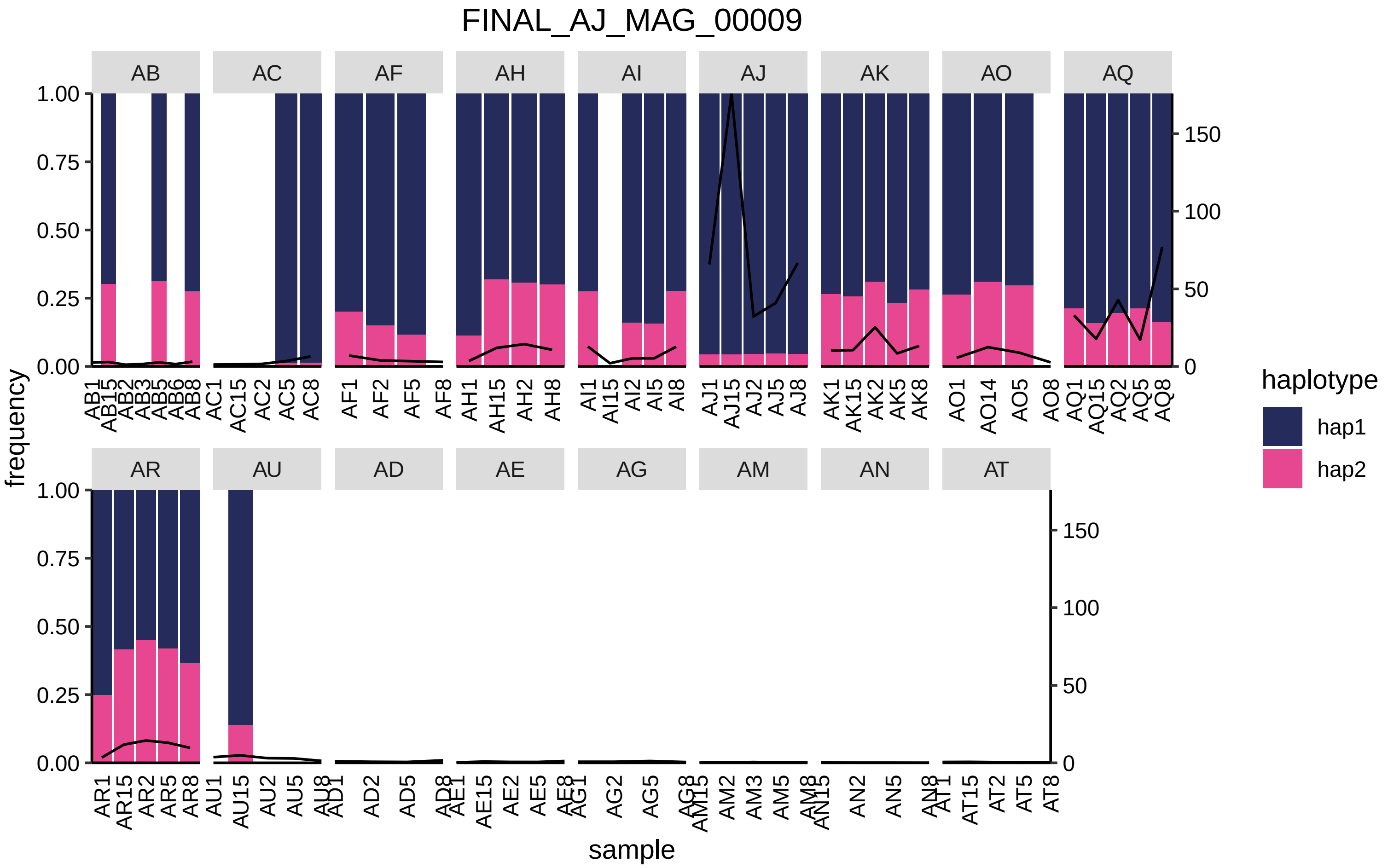
FINAL_AJ_MAG_00009
AB
AC
AF
AH
AI
AJ
AK
AO
AQ
AR
AU
AD
AE
AG
AM
AN
AT
frequency
sample
haplotype
hap1
hap2

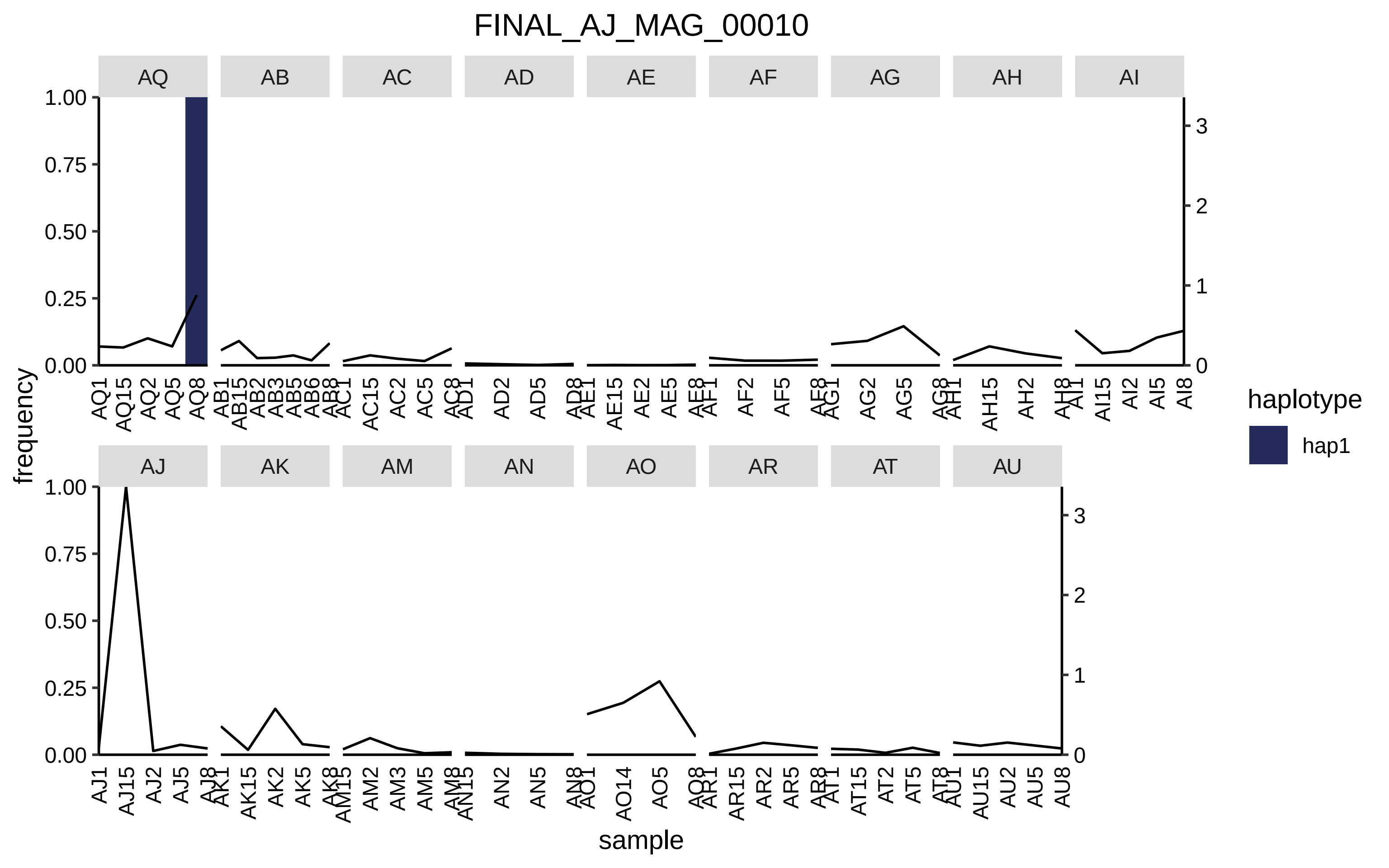
FINAL_AJ_MAG_00010
AQ
AB
AC
AD
AE
AF
AG
AH
AI
AJ
AK
AM
AN
AO
AR
AT
AU
frequency
sample
haplotype
hap1
1.00
0.75
0.50
0.25
0.00
3
2
1
0

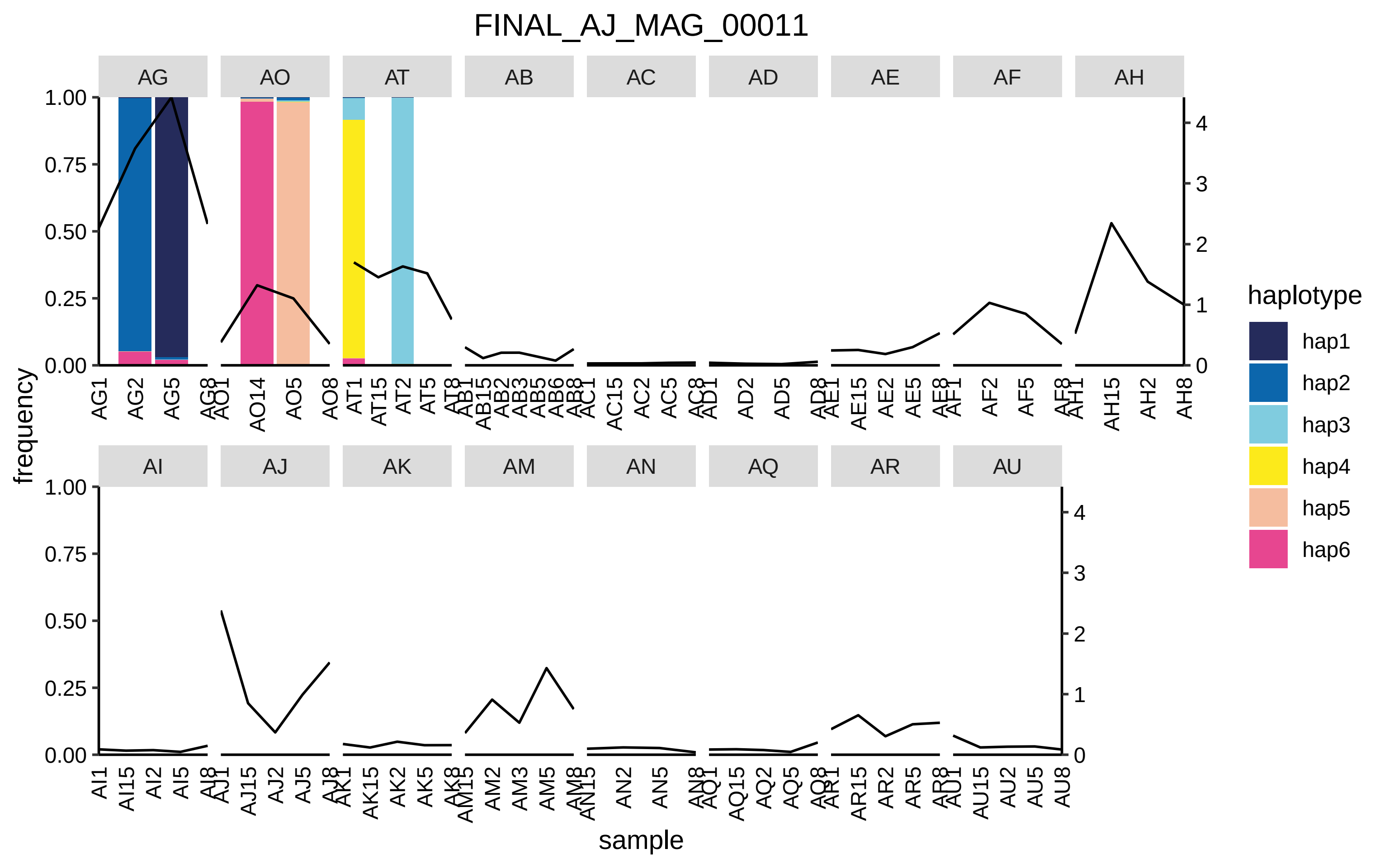
FINAL_AJ_MAG_00011
AG
AO
AT
AB
AC
AD
AE
AF
AH
AI
AJ
AK
AM
AN
AQ
AR
AU
frequency
sample
haplotype
hap1
hap2
hap3
hap4
hap5
hap6

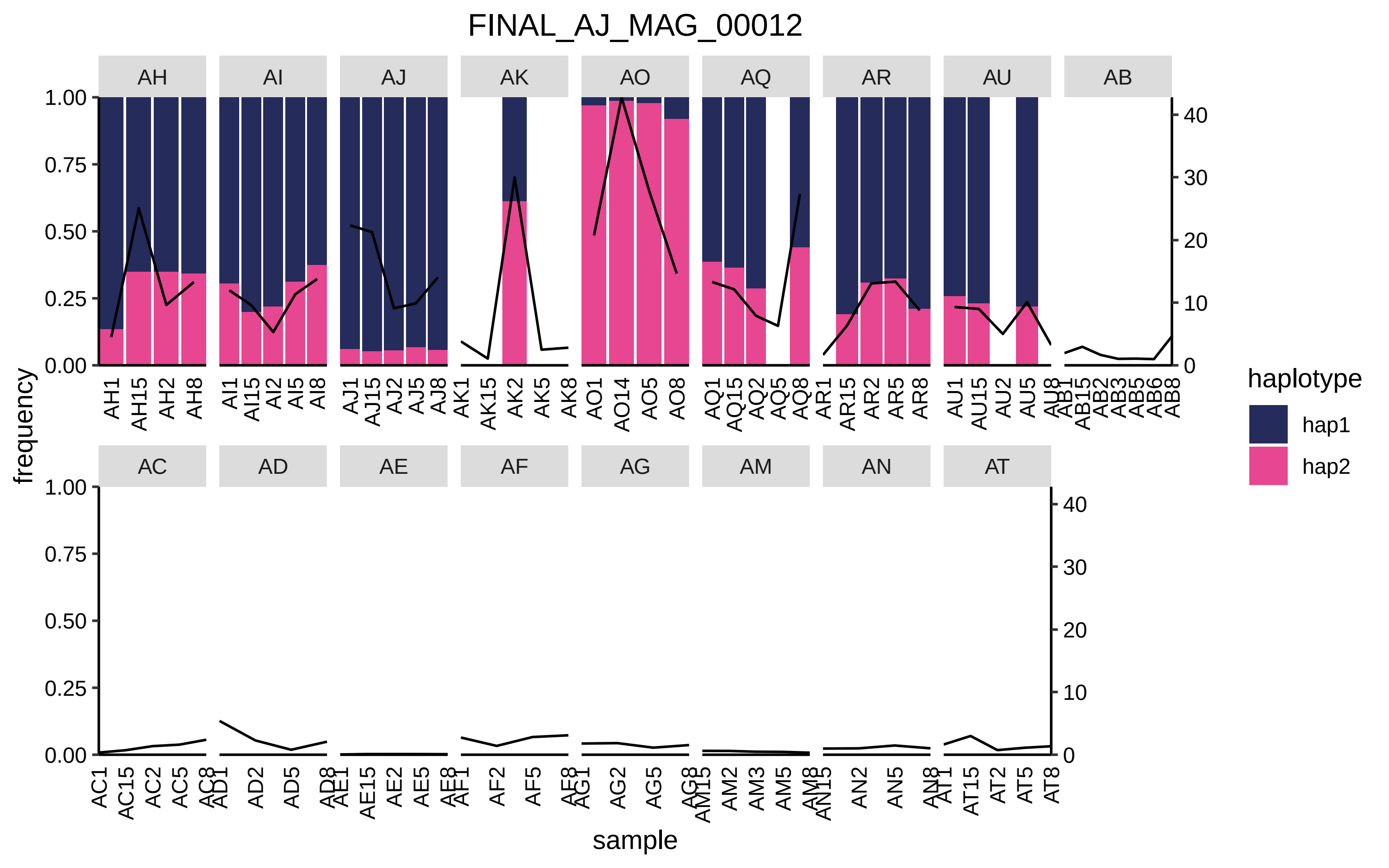

FINAL_AJ_MAG_00012
AH
AI
AJ
AK
AO
AQ
AR
AU
AB
AC
AD
AE
AF
AG
AM
AN
AT
frequency
sample
haplotype
hap1
hap2

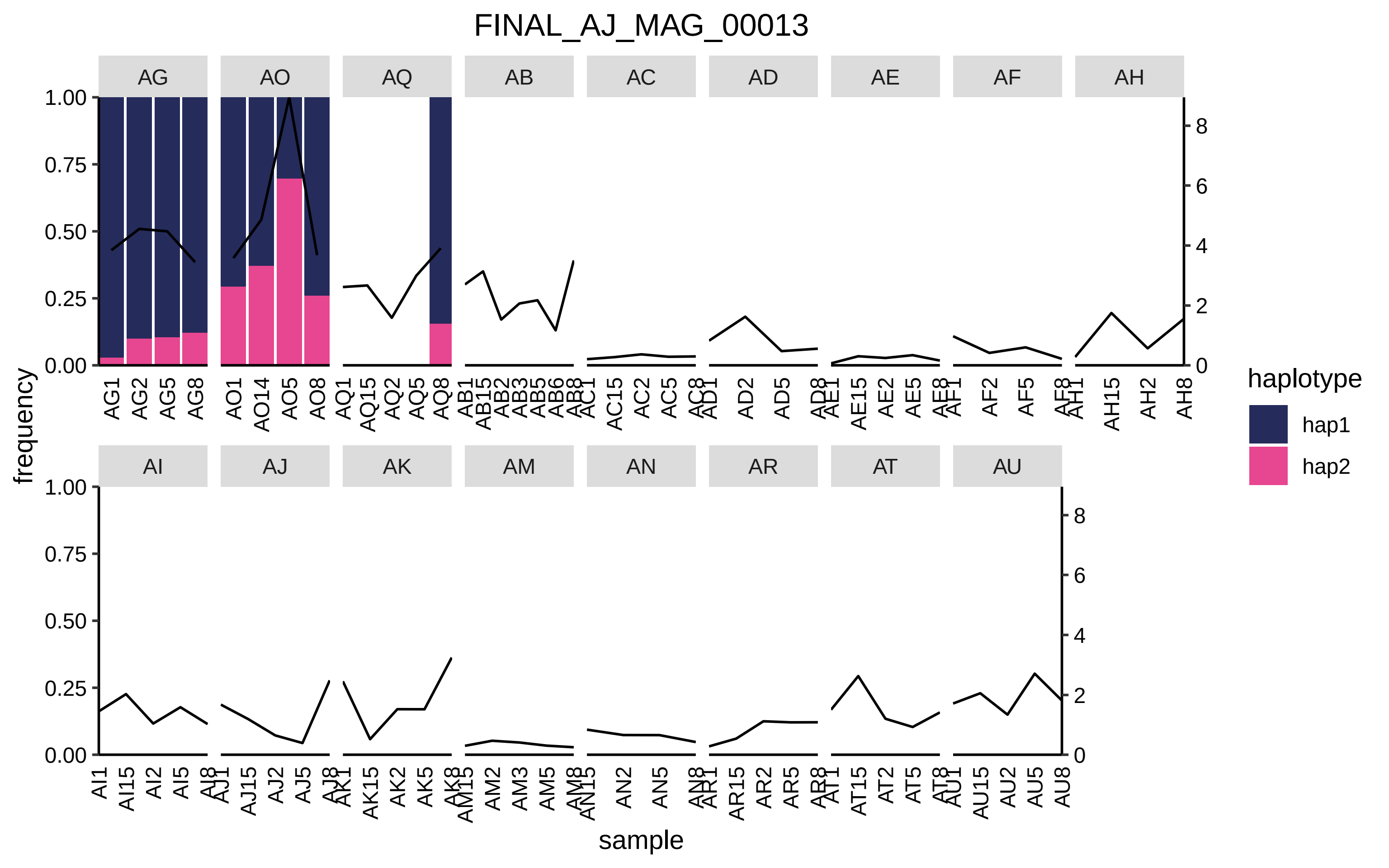
FINAL_AJ_MAG_00013
AG
AO
AQ
AB
AC
AD
AE
AF
AH
AI
AJ
AK
AM
AN
AR
AT
AU
frequency
sample
haplotype
hap1
hap2
1.00
0.75
0.50
0.25
0.00
8
6
4
2
0
AG1
AG2
AG5
AG8
AO1
AO14
AO5
AO8
AQ1
AQ15
AQ2
AQ5
AQ8
AB1
AB15
AB2
AB3
AB5
AB6
AB8
AC1
AC15
AC2
AC5
AC8
AD1
AD2
AD5
AD8
AE1
AE15
AE2
AE5
AE8
AF1
AF2
AF5
AF8
AH1
AH15
AH2
AH8
AI1
AI15
AI2
AI5
AI8
AJ1
AJ15
AJ2
AJ5
AJ8
AK1
AK15
AK2
AK5
AK8
AM15
AM2
AM3
AM5
AM8
AN15
AN2
AN5
AN8
AR1
AR15
AR2
AR5
AR8
AT1
AT15
AT2
AT5
AT8
AU1
AU15
AU2
AU5
AU8

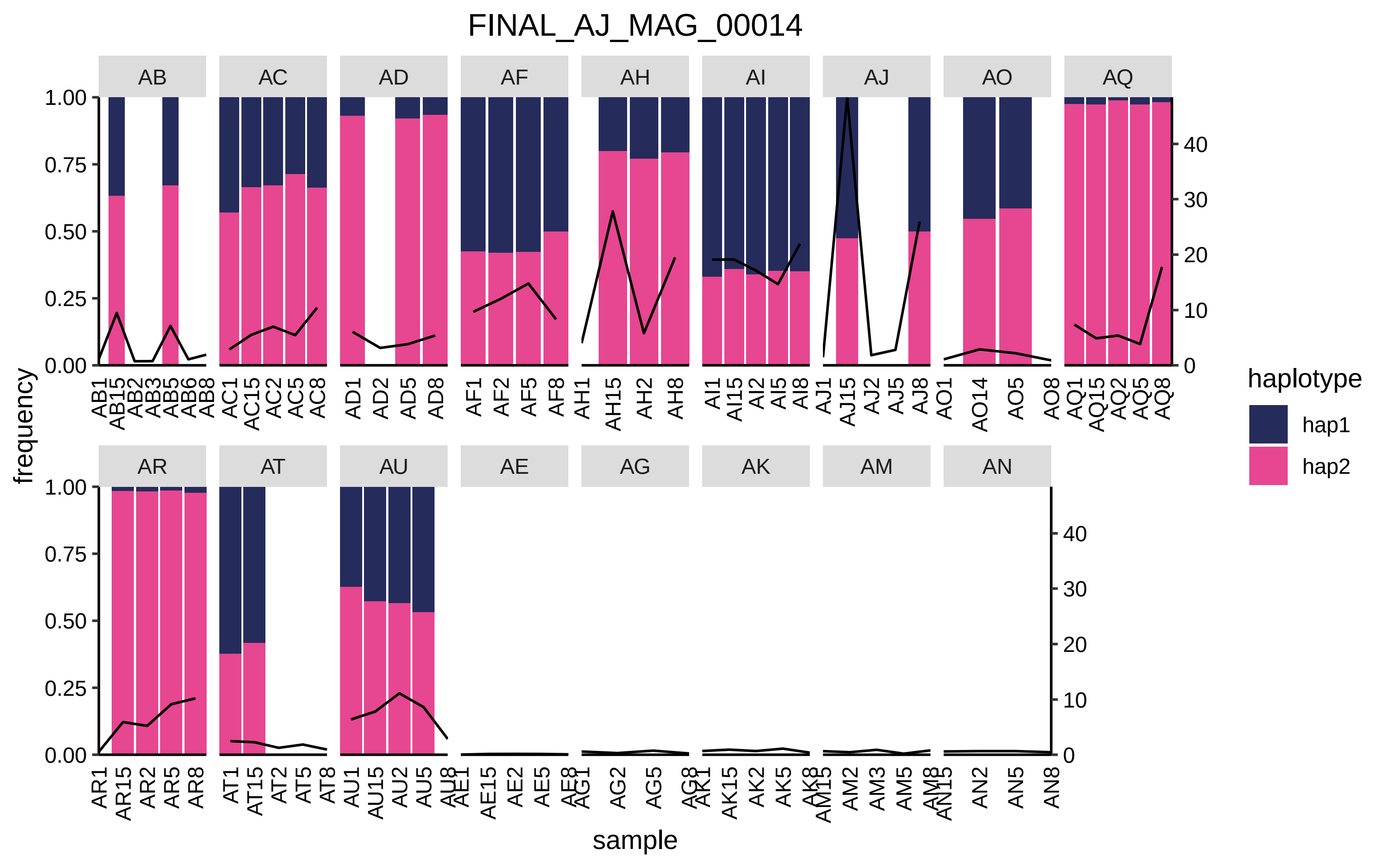
FINAL_AJ_MAG_00014
AB
AC
AD
AF
AH
AI
AJ
AO
AQ
AR
AT
AU
AE
AG
AK
AM
AN
frequency
sample
haplotype
hap1
hap2
1.00
0.75
0.50
0.25
0.00
40
30
20
10
0

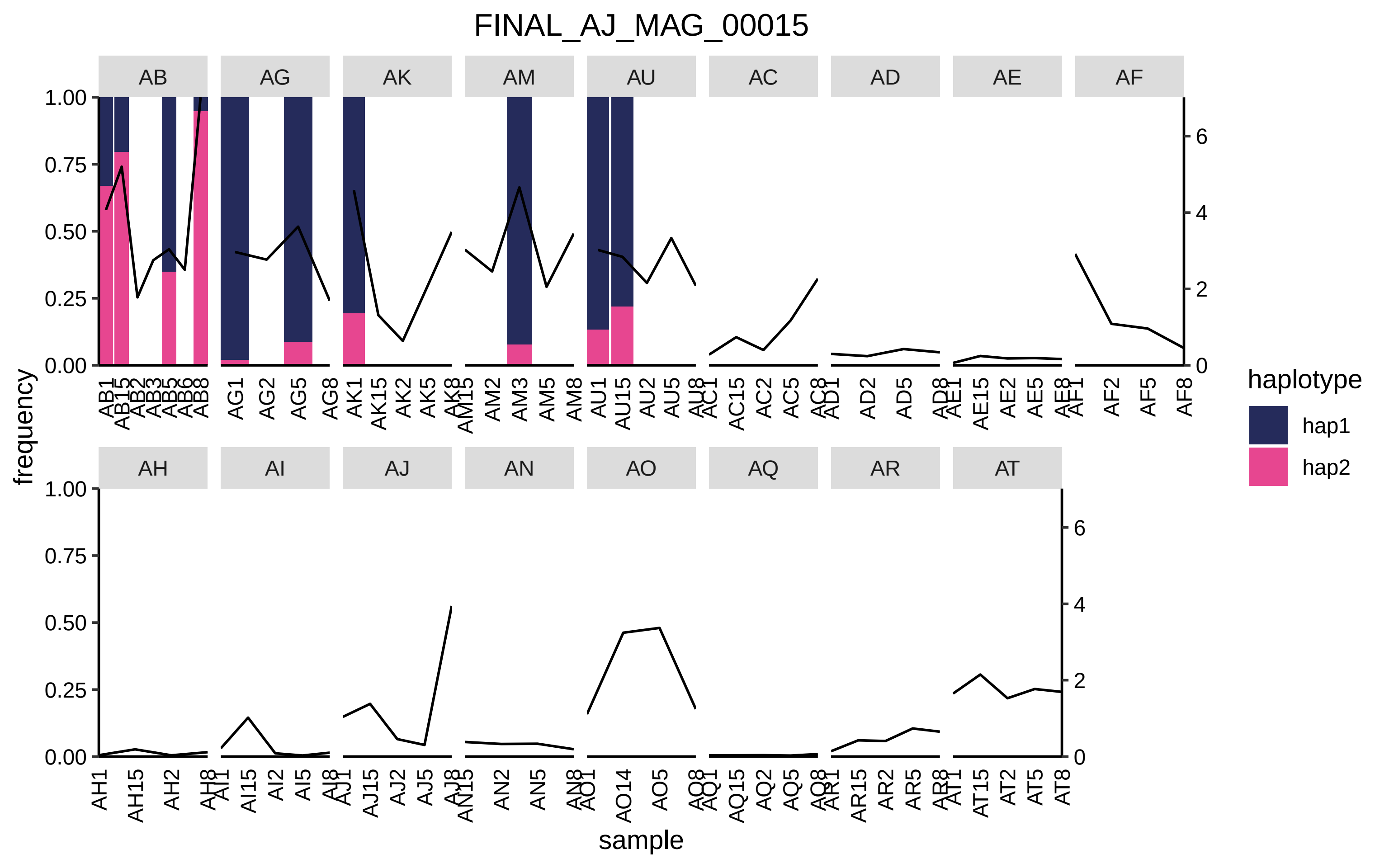
FINAL_AJ_MAG_00015
AB
AG
AK
AM
AU
AC
AD
AE
AF
AH
AI
AJ
AN
AO
AQ
AR
AT
frequency
sample
haplotype
hap1
hap2

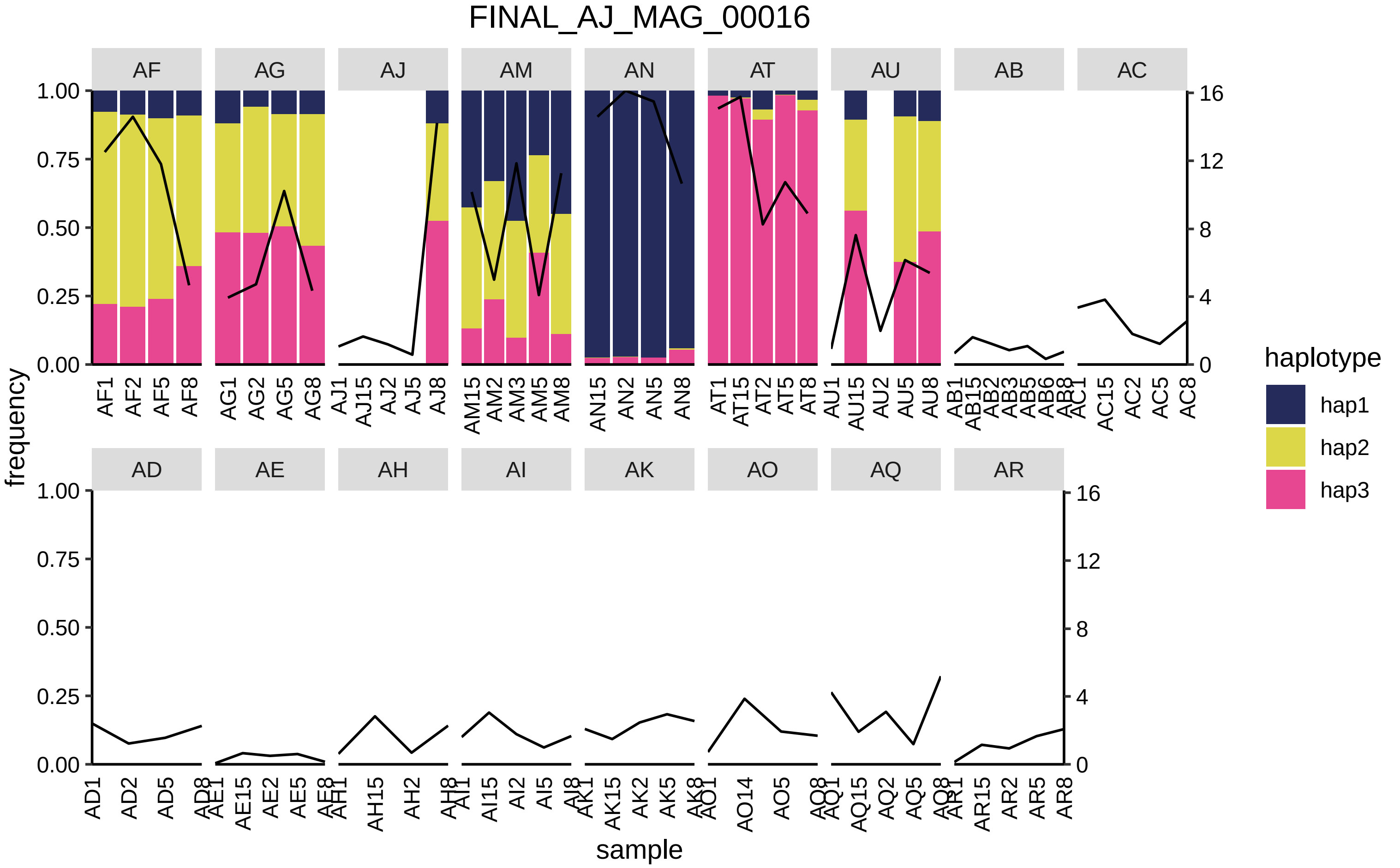

FINAL_AJ_MAG_00016
AF
AG
AJ
AM
AN
AT
AU
AB
AC
AD
AE
AH
AI
AK
AO
AQ
AR
frequency
sample
haplotype
hap1
hap2
hap3

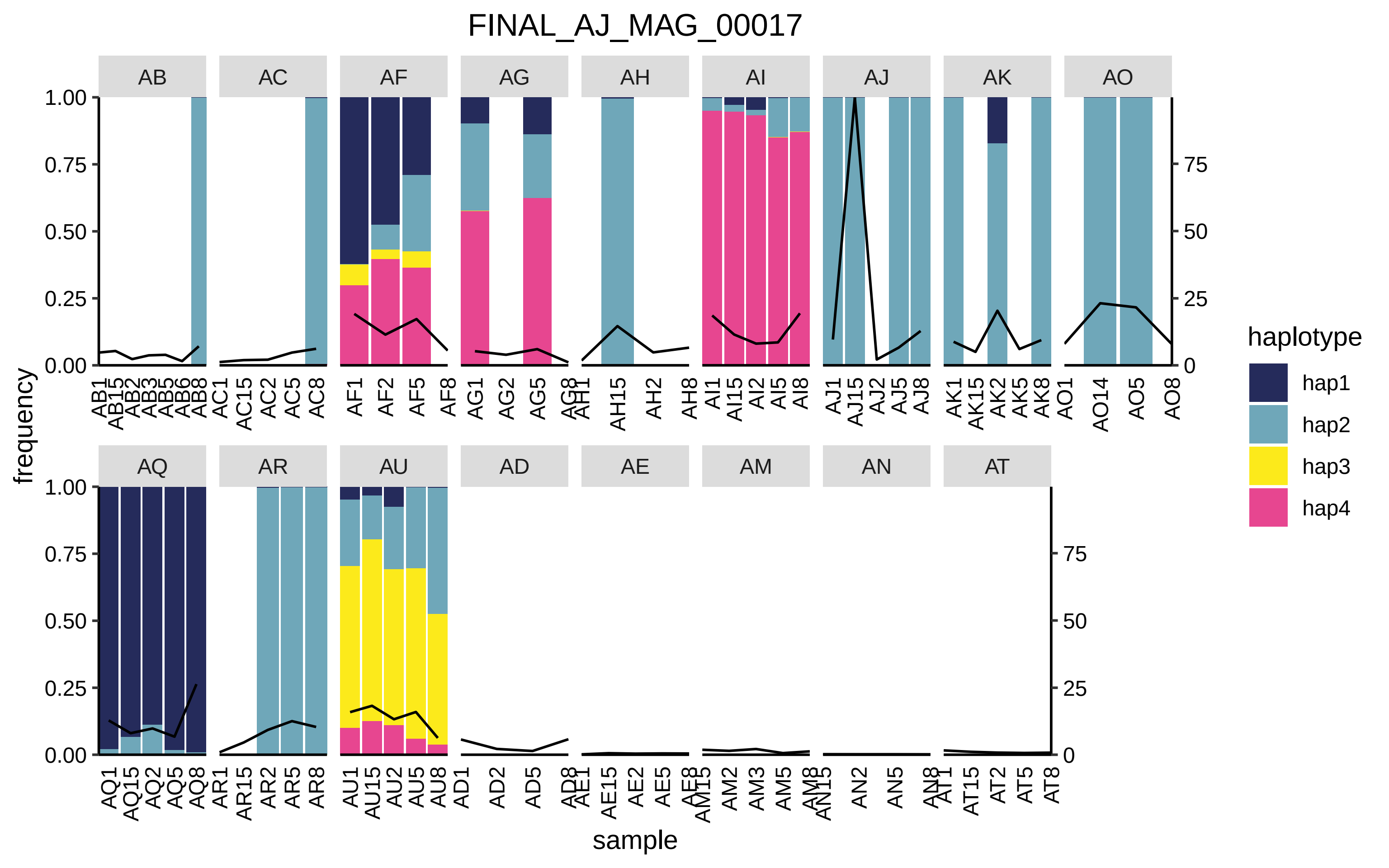

FINAL_AJ_MAG_00017
AB
AC
AF
AG
AH
AI
AJ
AK
AO
AQ
AR
AU
AD
AE
AM
AN
AT
frequency
sample
1.00
0.75
0.50
0.25
0.00
75
50
25
0
AF1 AF2 AF5 AF8
AG1 AG2 AG5
AH15 AH2 AH8
AI1 AI15 AI2 AI5 AI8
AJ1 AJ15 AJ2 AJ5 AJ8
AK1 AK15 AK2 AK5 AK8
AO1 AO14 AO5 AO8
AQ1 AQ15 AQ2 AQ5 AQ8
AR1 AR15 AR2 AR5 AR8
AU1 AU15 AU2 AU5 AU8
AD1 AD2 AD5
AE15 AE2 AE5
AM2 AM3 AM5
AN2 AN5
AT15 AT2 AT5 AT8
AC1 AC15 AC2 AC5 AC8
haplotype
hap1
hap2
hap3
hap4

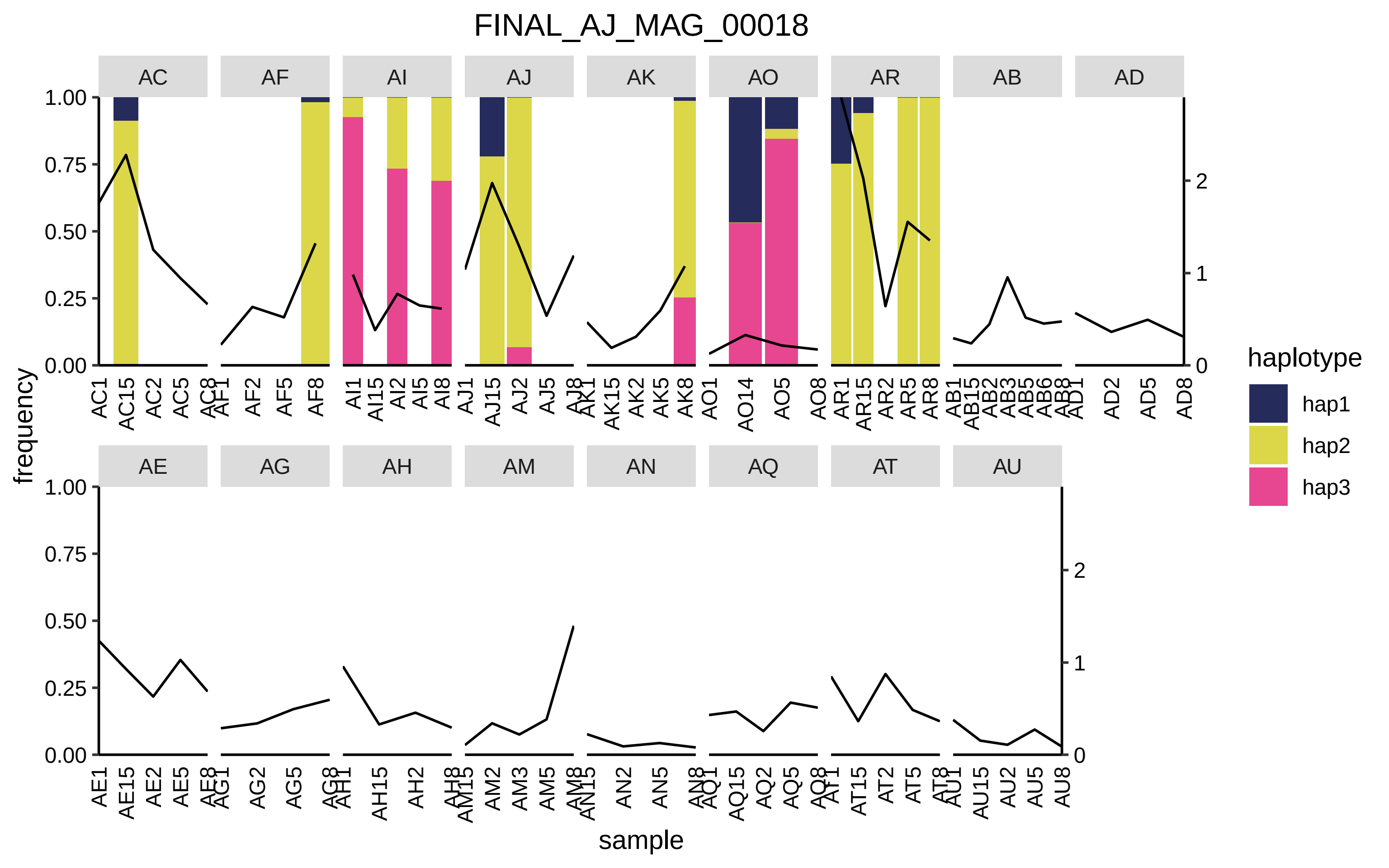
FINAL_AJ_MAG_00018
AC
AF
AI
AJ
AK
AO
AR
AB
AD
AE
AG
AH
AM
AN
AQ
AT
AU
frequency
sample
1.00
0.75
0.50
0.25
0.00
0
1
2
AC1 AC15 AC2 AC5 AC8
AF1 AF2 AF5 AF8
AI1 AI15 AI2 AI5 AI8
AJ1 AJ15 AJ2 AJ5 AJ8
AK1 AK15 AK2 AK5 AK8
AO1 AO14 AO5 AO8
AR1 AR15 AR2 AR5 AR8
AB1 AB15 AB2 AB3 AB5 AB6 AB8
AD1 AD2 AD5 AD8
AE1 AE15 AE2 AE5 AE8
AG1 AG2 AG5 AG8
AH1 AH15 AH2 AH8
AM15 AM2 AM3 AM5 AM8
AN15 AN2 AN5 AN8
AQ1 AQ15 AQ2 AQ5 AQ8
AT1 AT15 AT2 AT5 AT8
AU1 AU15 AU2 AU5 AU8
haplotype
hap1
hap2
hap3

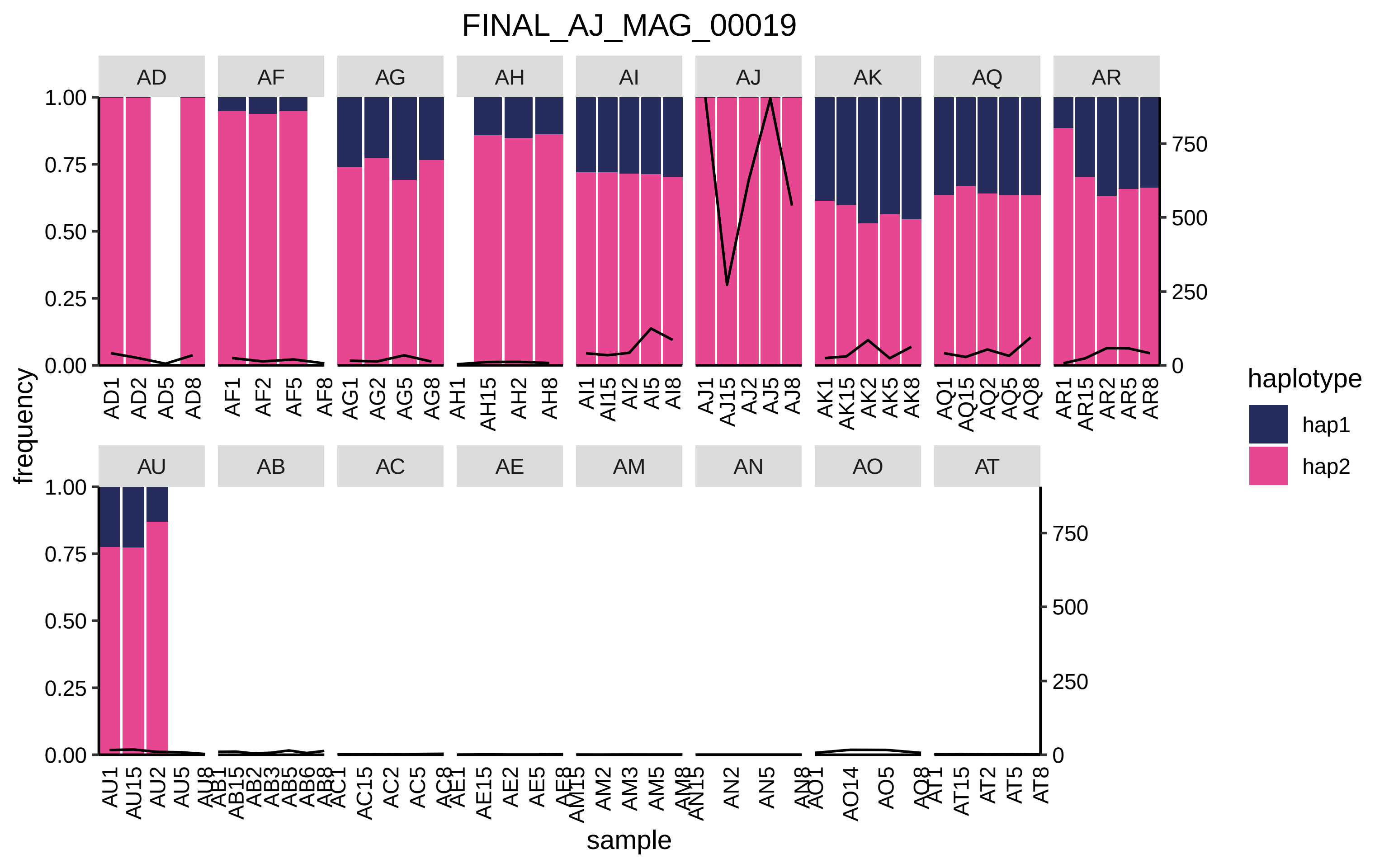

FINAL_AJ_MAG_00019
AD
AF
AG
AH
AI
AJ
AK
AQ
AR
AU
AB
AC
AE
AM
AN
AO
AT
frequency
sample
haplotype
hap1
hap2
1.00
0.75
0.50
0.25
0.00
750
500
250
0

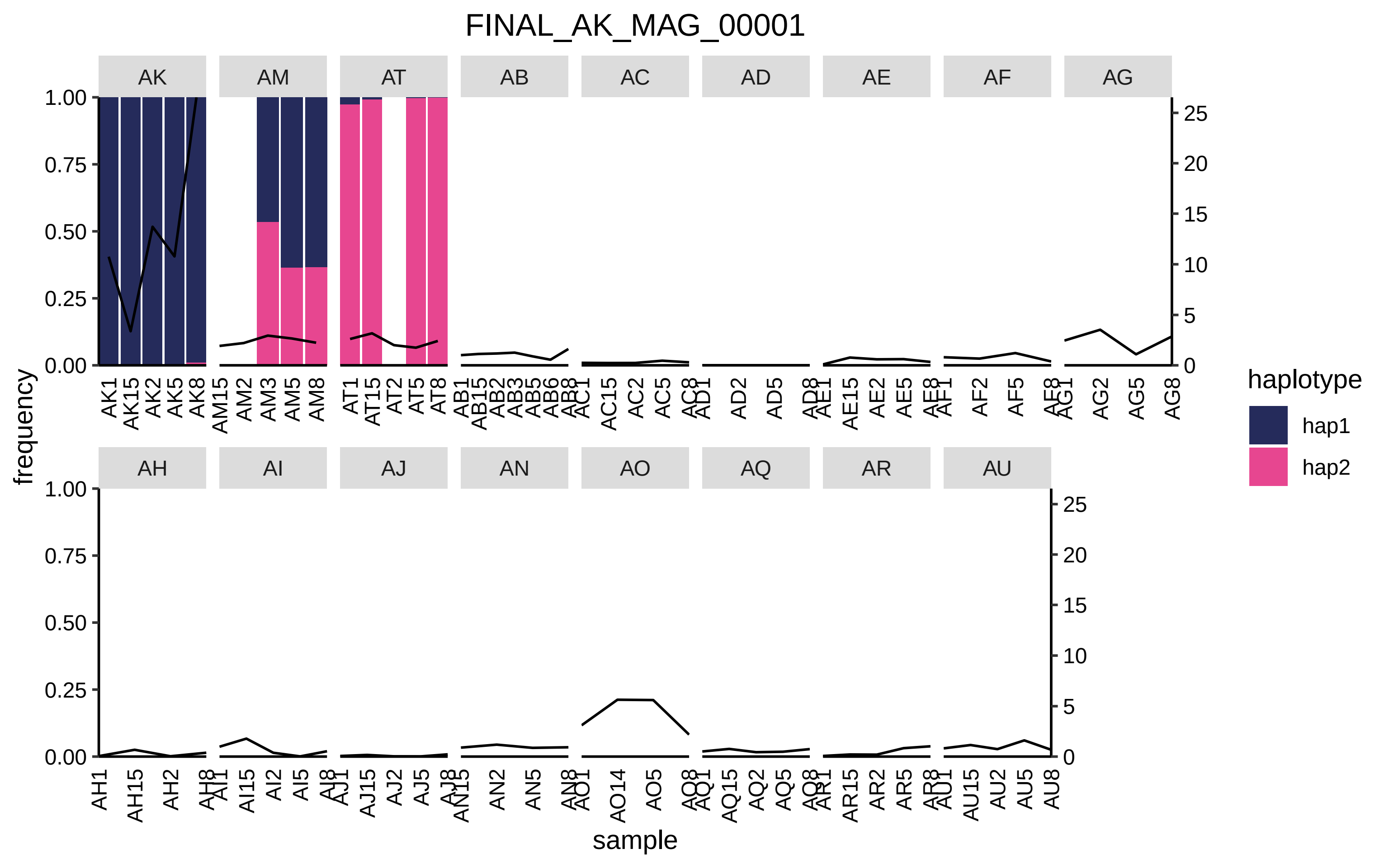
FINAL_AK_MAG_00001
AK
AM
AT
AB
AC
AD
AE
AF
AG
AH
AI
AJ
AN
AO
AQ
AR
AU
frequency
sample
haplotype
hap1
hap2

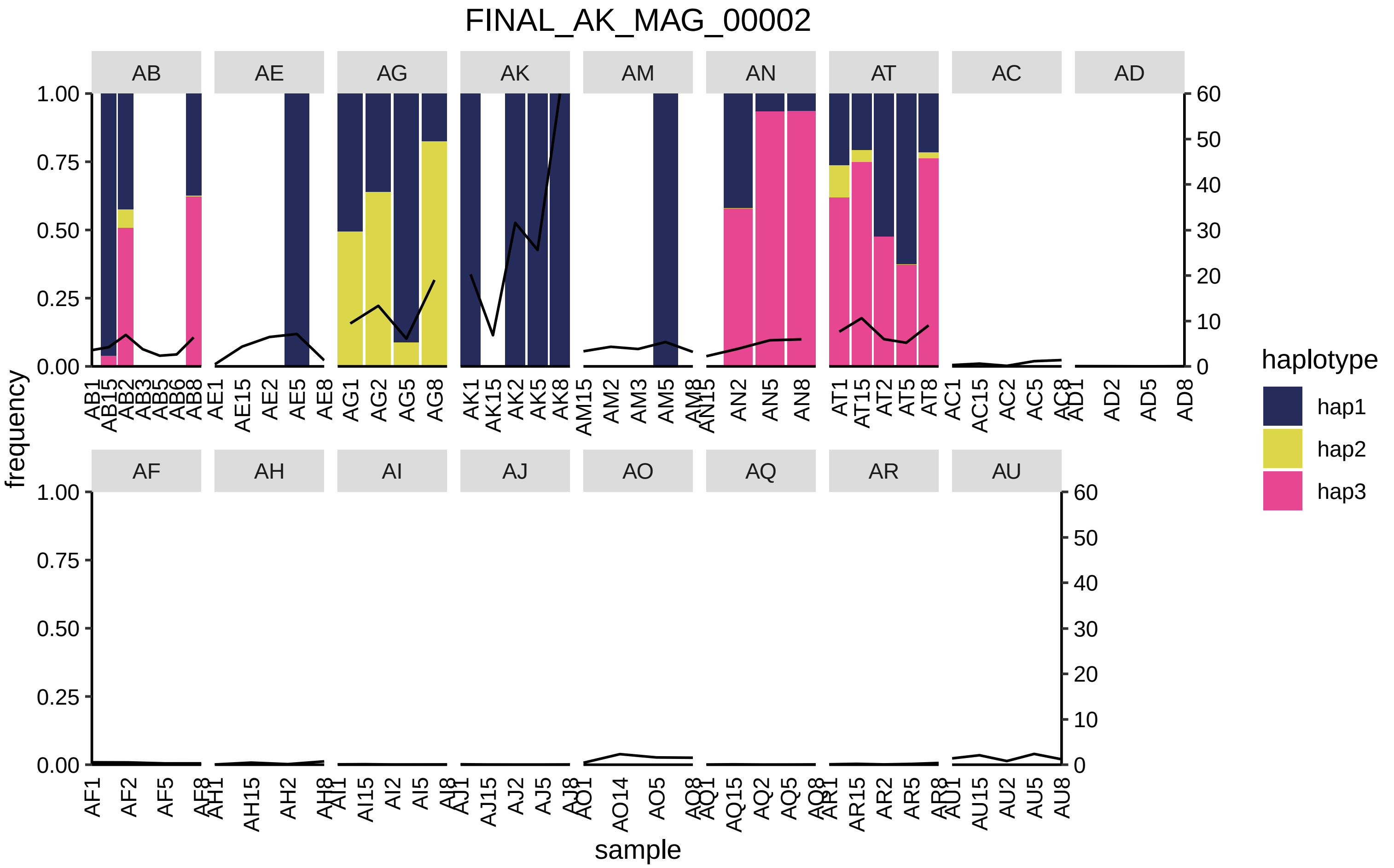
FINAL_AK_MAG_00002
AB
AE
AG
AK
AM
AN
AT
AC
AD
AF
AH
AI
AJ
AO
AQ
AR
AU
frequency
sample
haplotype
hap1
hap2
hap3

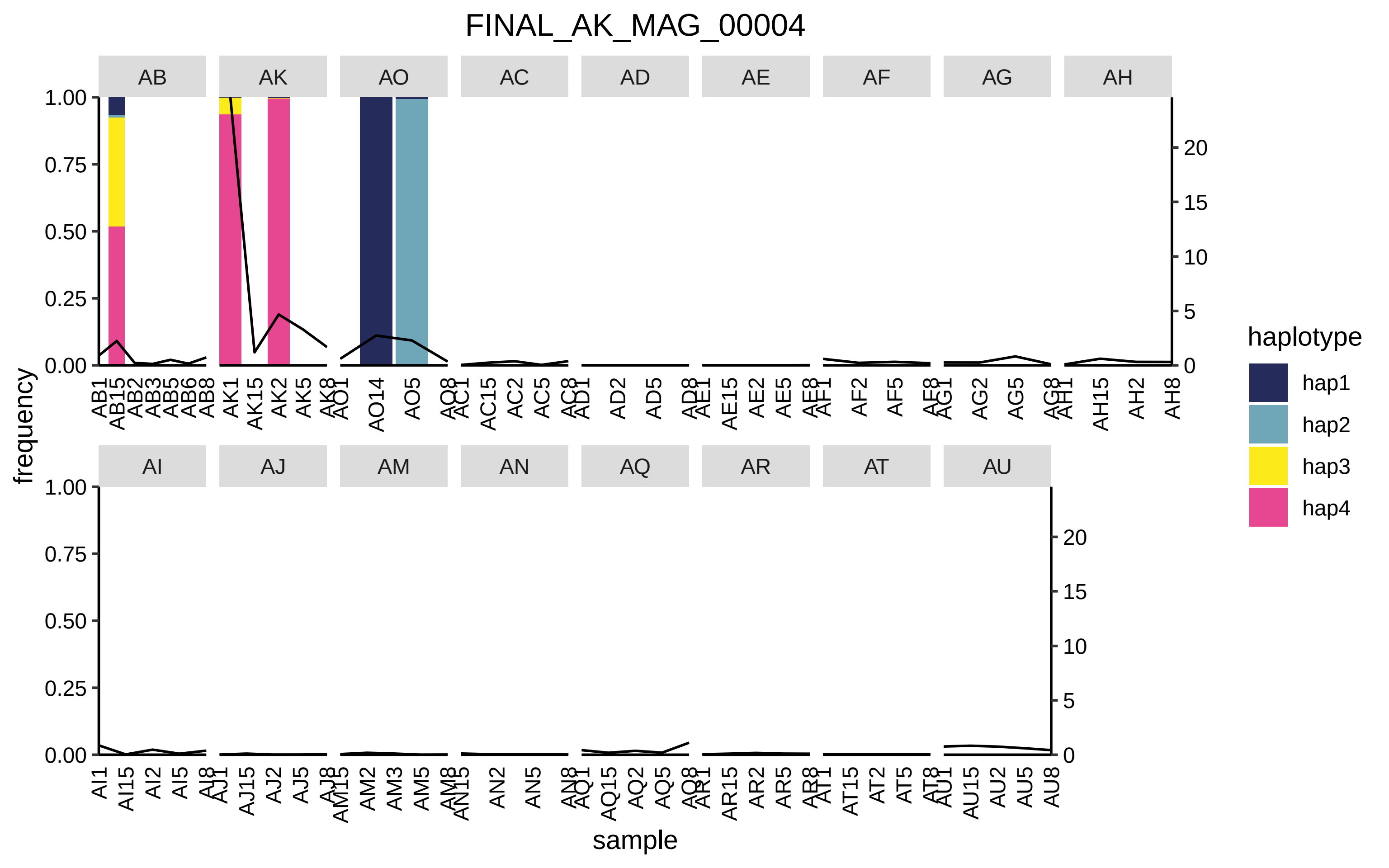

FINAL_AK_MAG_00004
AB
AK
AO
AC
AD
AE
AF
AG
AH
AI
AJ
AM
AN
AQ
AR
AT
AU
frequency
sample
haplotype
hap1
hap2
hap3
hap4
1.00
0.75
0.50
0.25
0.00
20
15
10
5
0
AB1 AB15 AB2 AB3 AB5 AB6 AB8
AK1 AK15 AK2 AK5 AK8
AO1 AO14 AO5 AO8
AC1 AC15 AC2 AC5 AC8
AD1 AD2 AD5 AD8
AE1 AE15 AE2 AE5 AE8
AF1 AF2 AF5 AF8
AG1 AG2 AG5 AG8
AH1 AH15 AH2 AH8
AI1 AI15 AI2 AI5 AI8
AJ1 AJ15 AJ2 AJ5 AJ8
AM15 AM2 AM3 AM5 AM8
AN15 AN2 AN5 AN8
AQ1 AQ15 AQ2 AQ5 AQ8
AR1 AR15 AR2 AR5 AR8
AT1 AT15 AT2 AT5 AT8
AU1 AU15 AU2 AU5 AU8

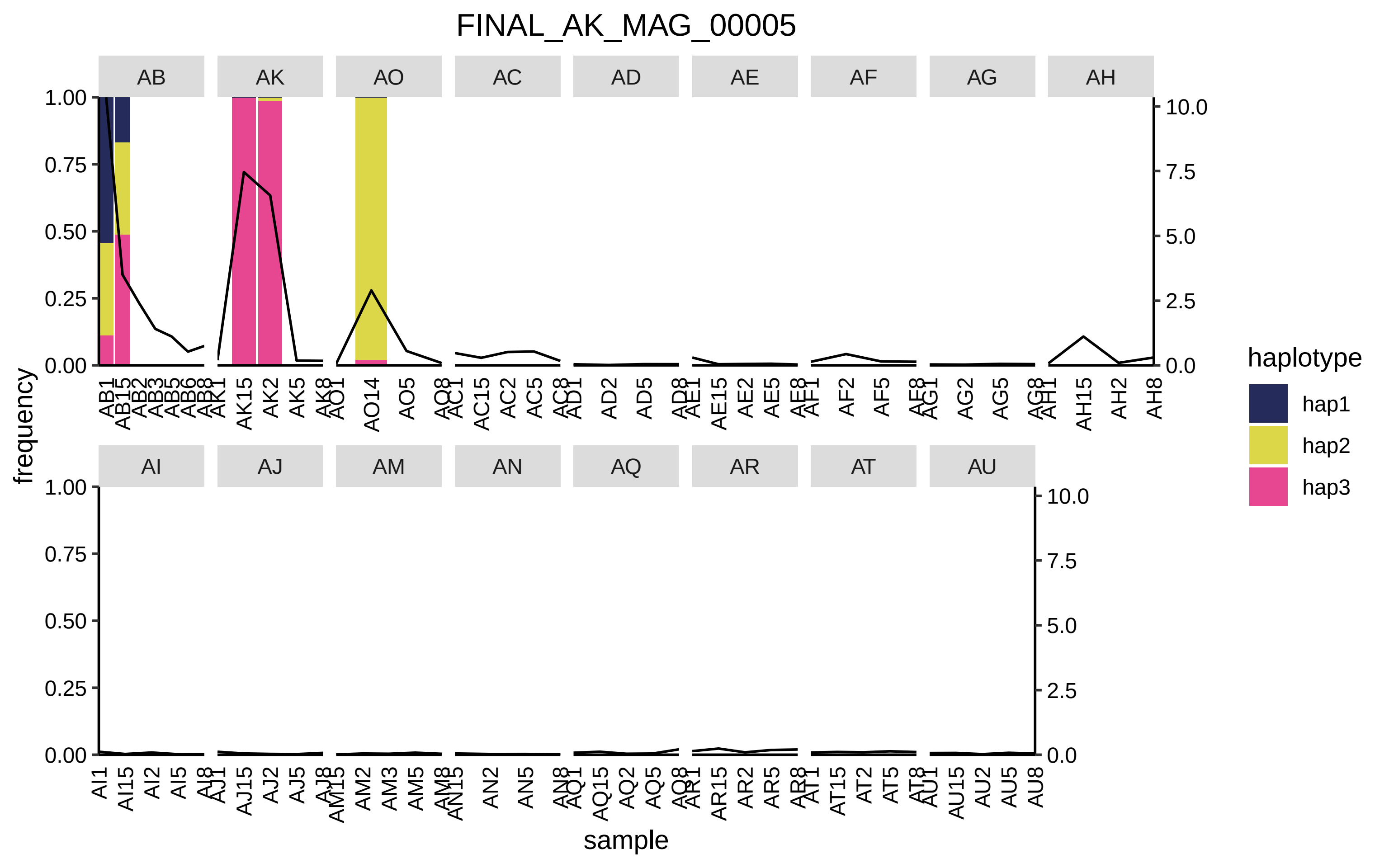
FINAL_AK_MAG_00005
AB
AK
AO
AC
AD
AE
AF
AG
AH
AI
AJ
AM
AN
AQ
AR
AT
AU
frequency
sample
haplotype
hap1
hap2
hap3

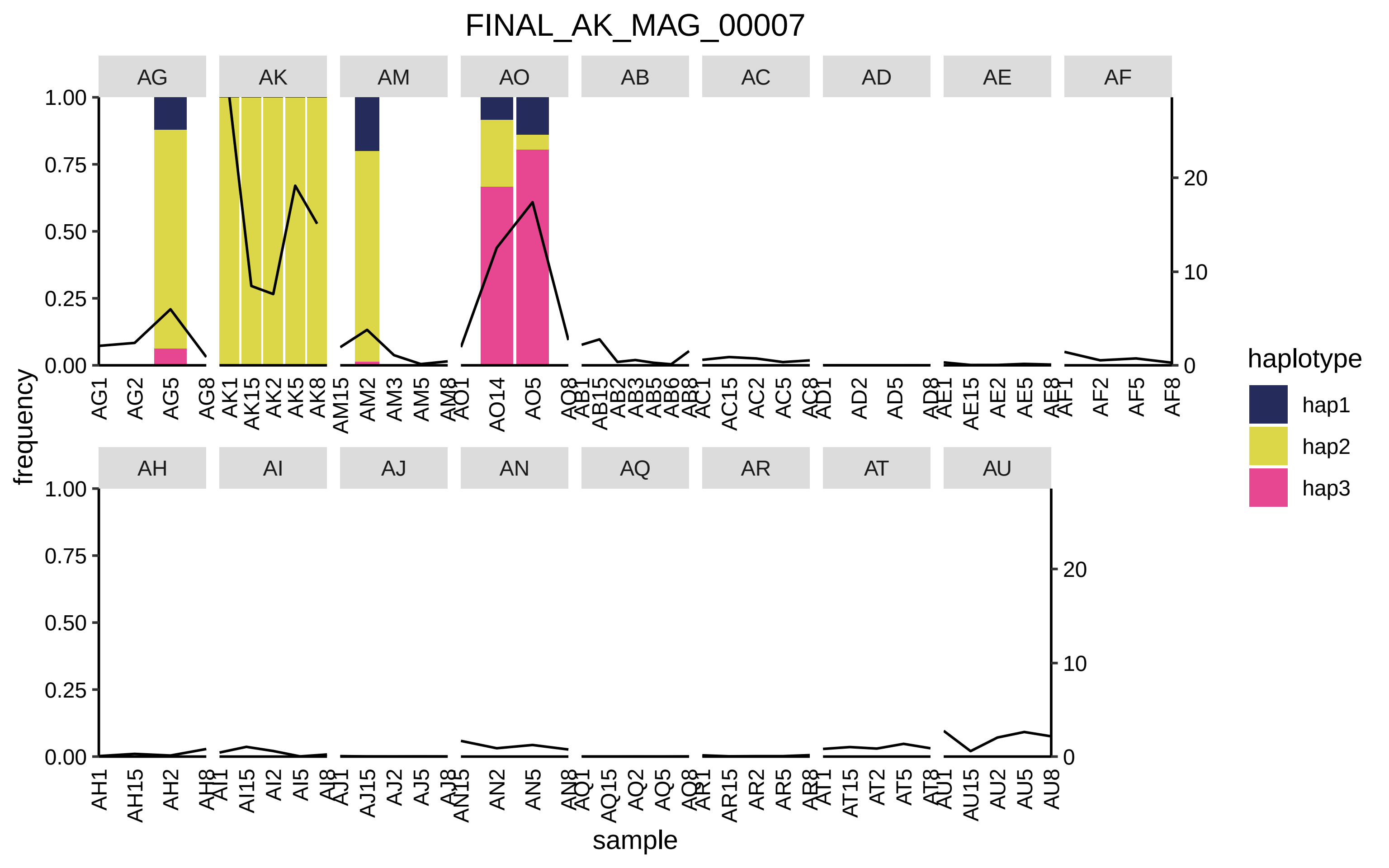
FINAL_AK_MAG_00007
AG
AK
AM
AO
AB
AC
AD
AE
AF
AH
AI
AJ
AN
AQ
AR
AT
AU
1.00
0.75
0.50
0.25
0.00
20
10
0
frequency
sample
AG1
AG2
AG5
AG8
AK1
AK15
AK2
AK5
AK8
AM15
AM2
AM3
AM5
AM8
AO1
AO14
AO5
AO8
AB1
AB15
AB2
AB3
AB5
AB6
AB8
AC1
AC15
AC2
AC5
AC8
AD1
AD2
AD5
AD8
AE1
AE15
AE2
AE5
AE8
AF1
AF2
AF5
AF8
AH1
AH15
AH2
AH8
AI1
AI15
AI2
AI5
AI8
AJ1
AJ15
AJ2
AJ5
AJ8
AN15
AN2
AN5
AN8
AQ1
AQ15
AQ2
AQ5
AQ8
AR1
AR15
AR2
AR5
AR8
AT1
AT15
AT2
AT5
AT8
AU1
AU15
AU2
AU5
AU8
haplotype
hap1
hap2
hap3

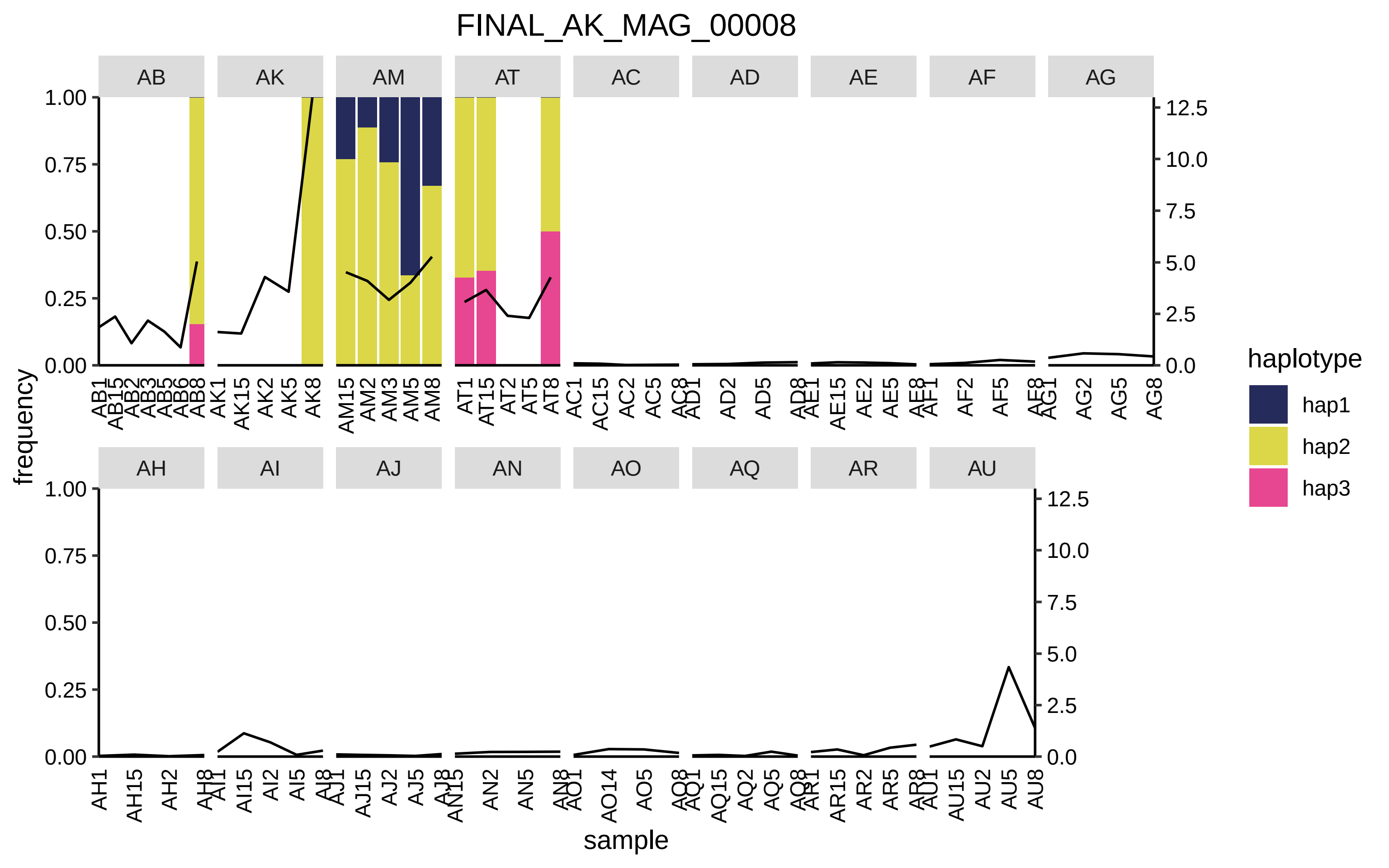
FINAL_AK_MAG_00008
AB
AK
AM
AT
AC
AD
AE
AF
AG
AH
AI
AJ
AN
AO
AQ
AR
AU
frequency
sample
haplotype
hap1
hap2
hap3

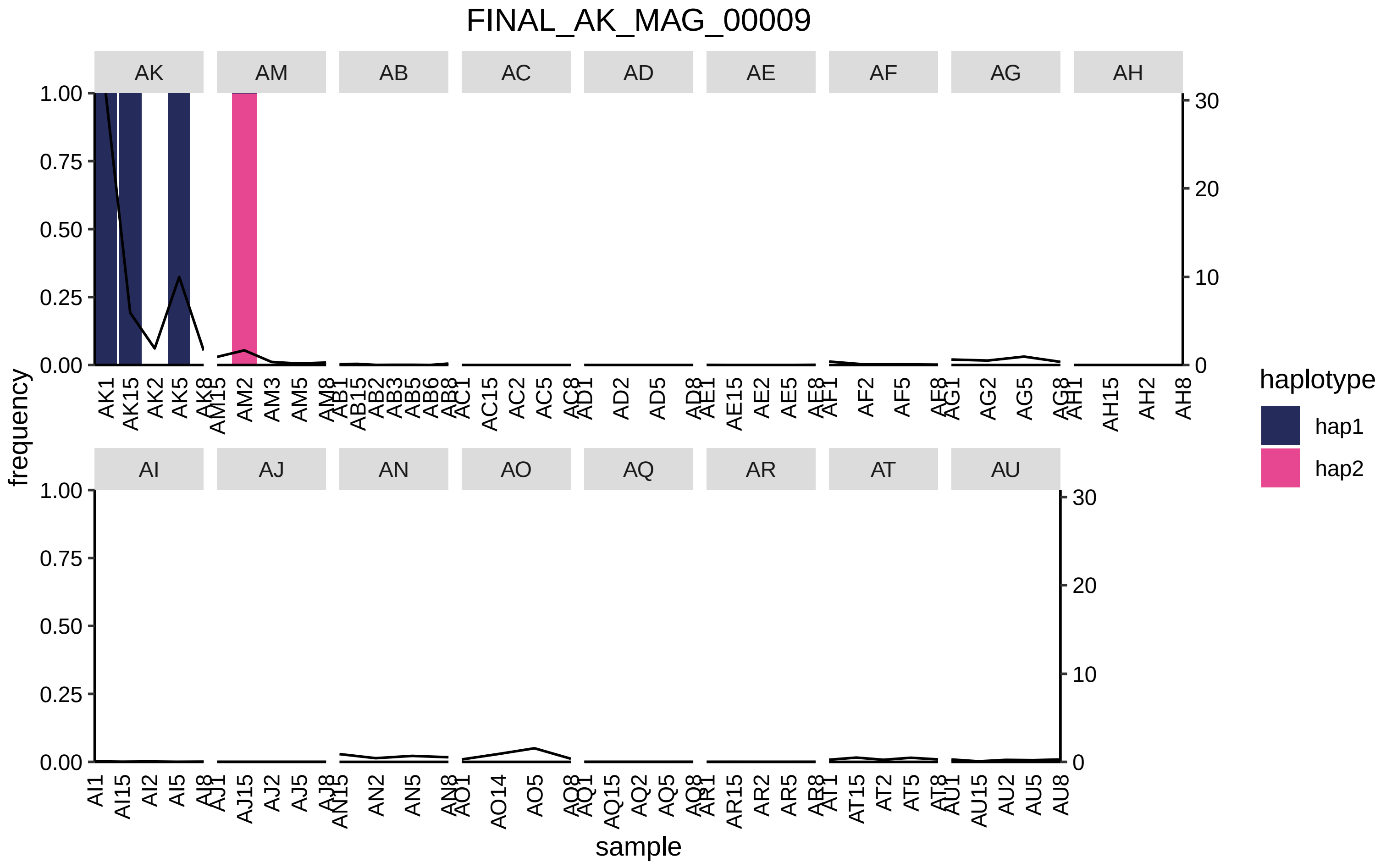

FINAL_AK_MAG_00009
AK
AM
AB
AC
AD
AE
AF
AG
AH
AI
AJ
AN
AO
AQ
AR
AT
AU
frequency
sample
haplotype
hap1
hap2
1.00
0.75
0.50
0.25
0.00
30
20
10
0

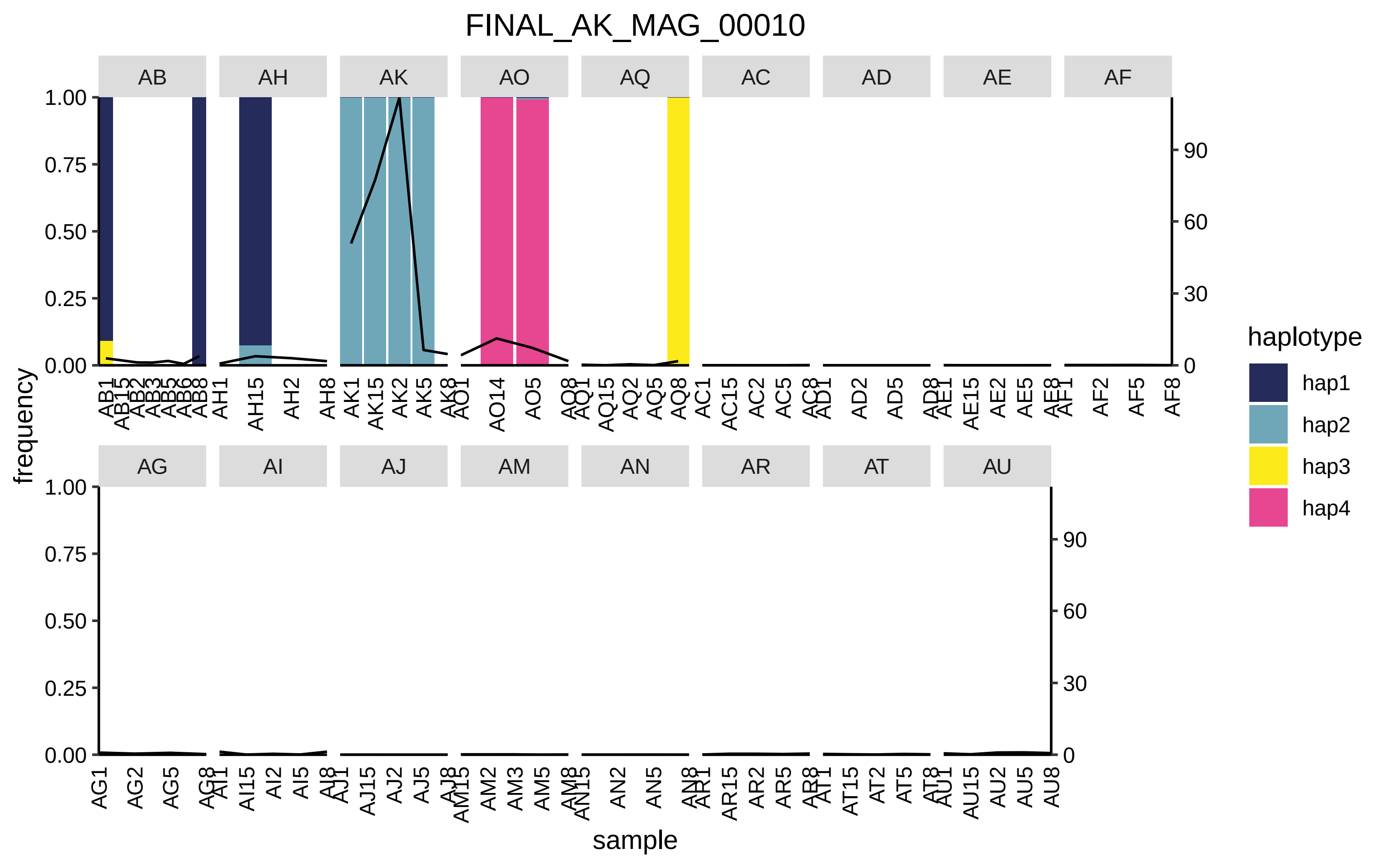
FINAL_AK_MAG_00010
AB
AH
AK
AO
AQ
AC
AD
AE
AF
AG
AI
AJ
AM
AN
AR
AT
AU
frequency
sample
haplotype
hap1
hap2
hap3
hap4
1.00
0.75
0.50
0.25
0.00
90
60
30
0

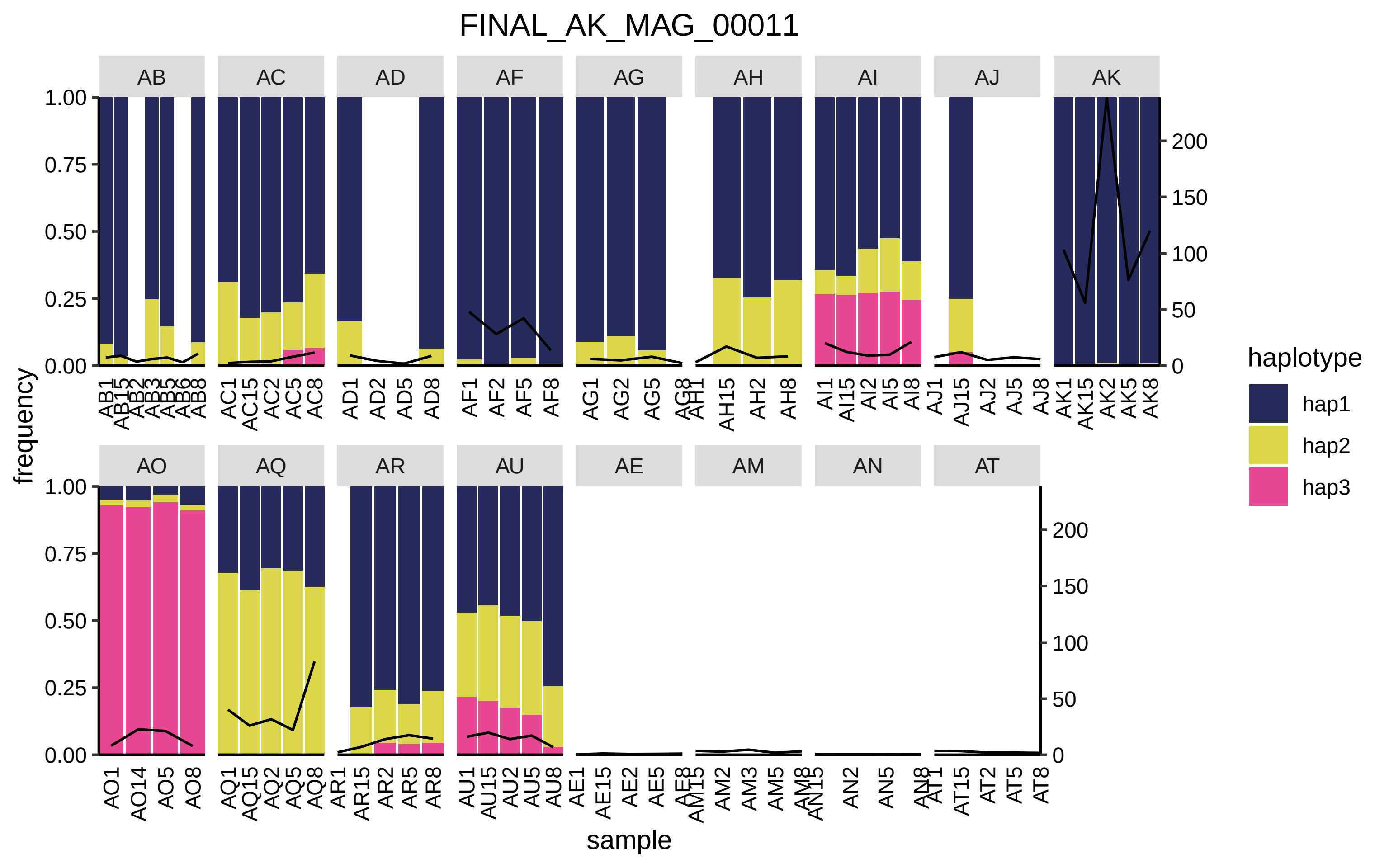

FINAL_AK_MAG_00011
AB
AC
AD
AF
AG
AH
AI
AJ
AK
AO
AQ
AR
AU
AE
AM
AN
AT
frequency
sample
1.00
0.75
0.50
0.25
0.00
200
150
100
50
0
haplotype
hap1
hap2
hap3

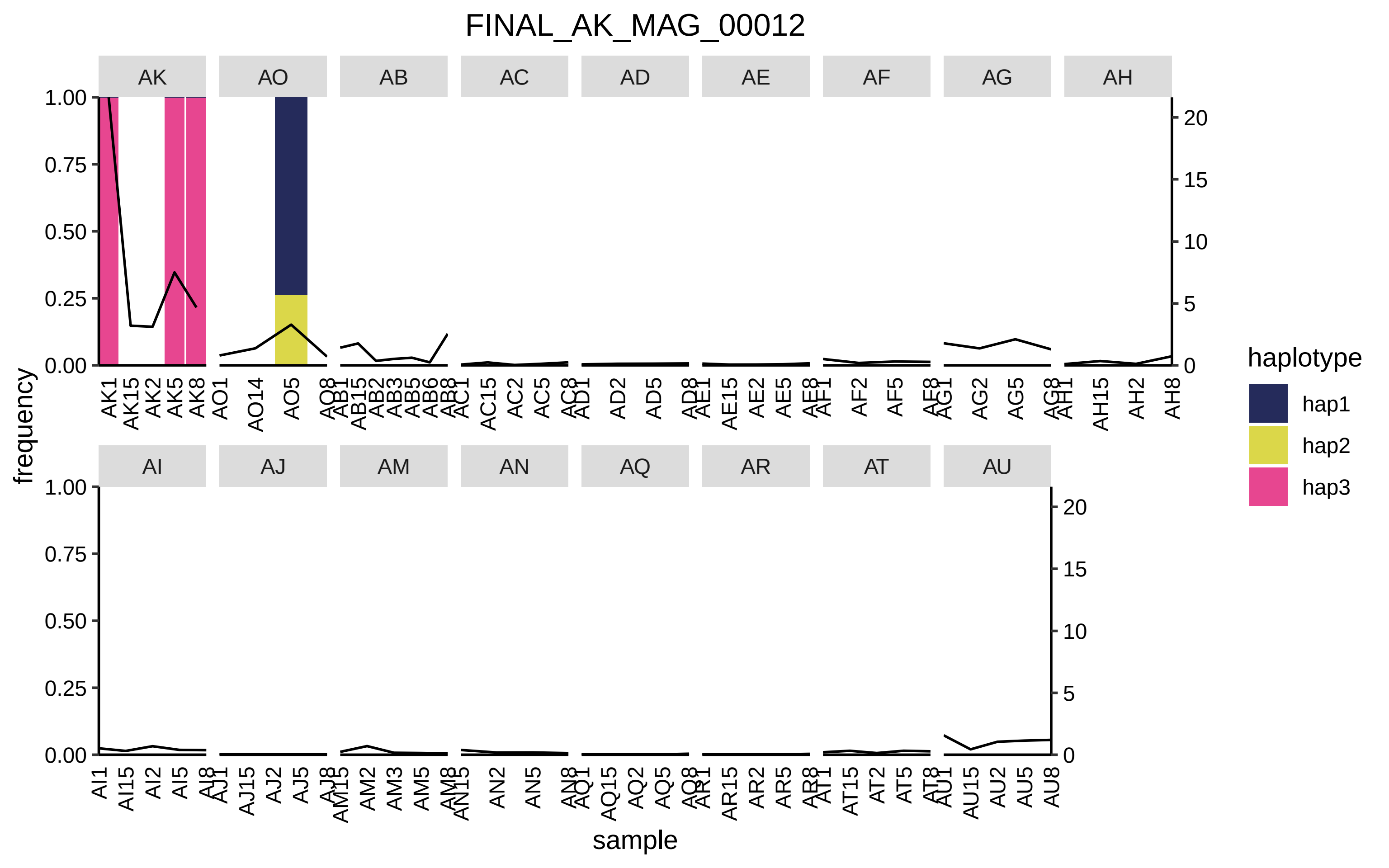
FINAL_AK_MAG_00012
AK
AO
AB
AC
AD
AE
AF
AG
AH
AI
AJ
AM
AN
AQ
AR
AT
AU
frequency
sample
haplotype
hap1
hap2
hap3

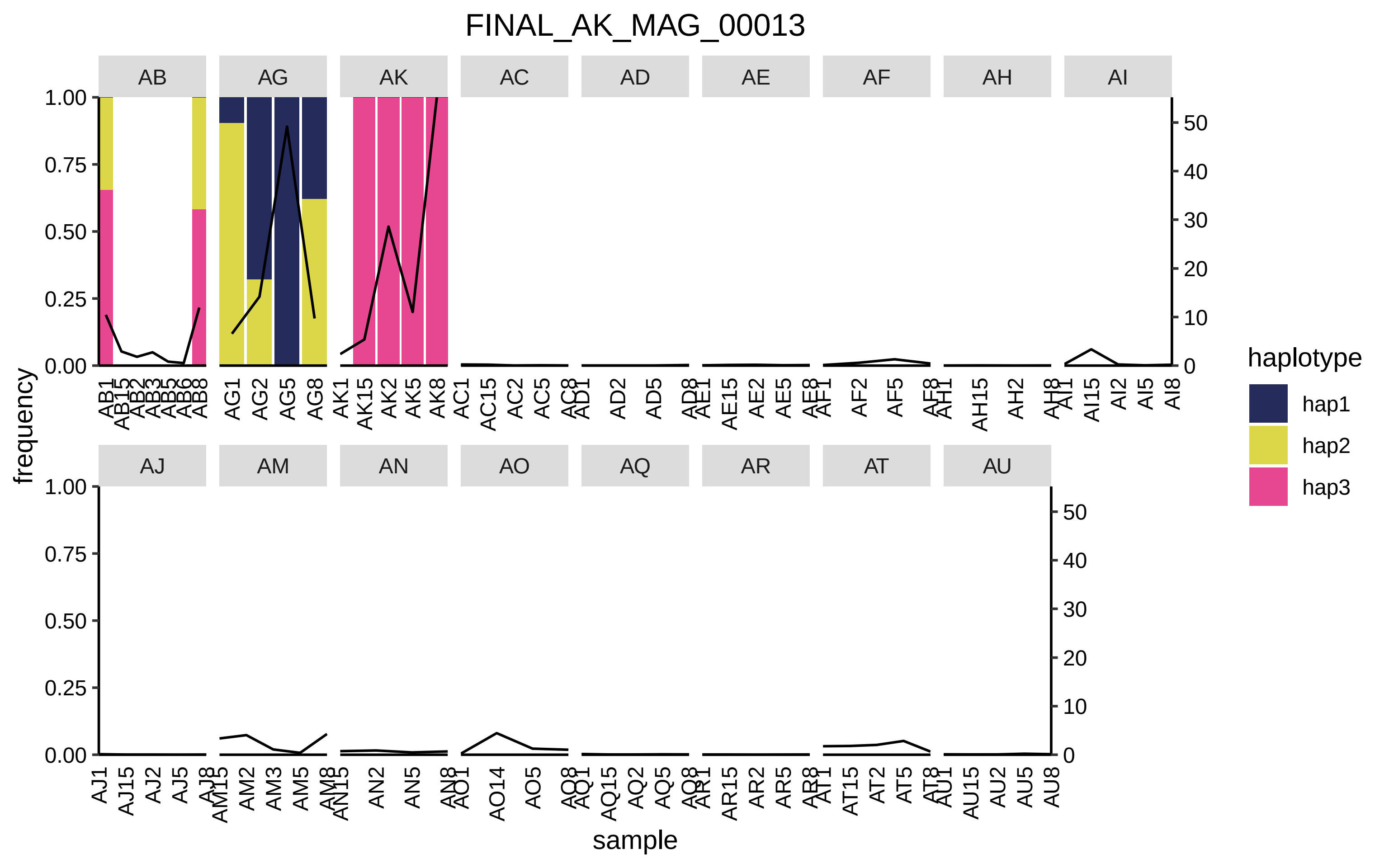

FINAL_AK_MAG_00013
AB
AG
AK
AC
AD
AE
AF
AH
AI
AJ
AM
AN
AO
AQ
AR
AT
AU
frequency
sample
haplotype
hap1
hap2
hap3

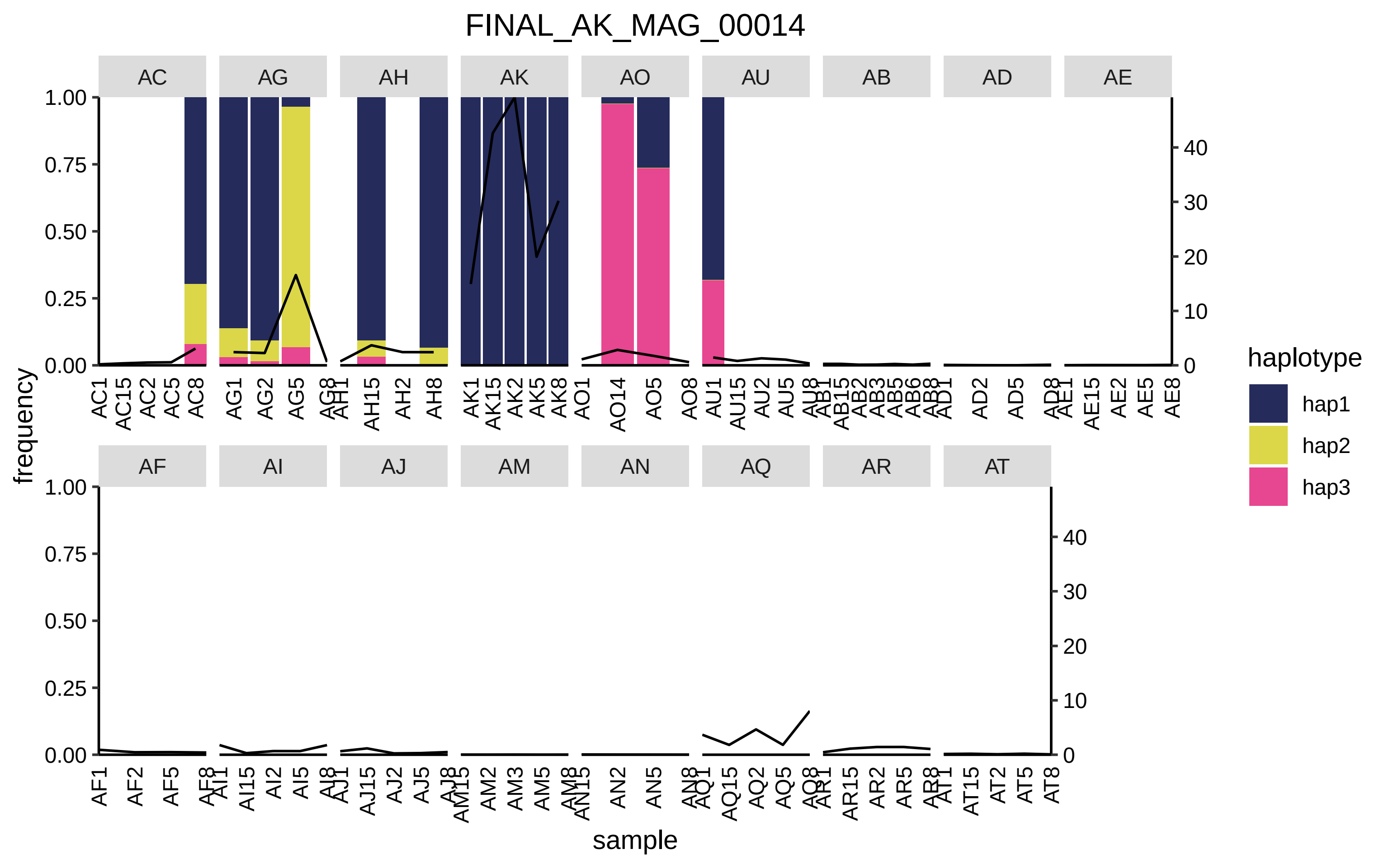
FINAL_AK_MAG_00014
AC
AG
AH
AK
AO
AU
AB
AD
AE
AF
AI
AJ
AM
AN
AQ
AR
AT
frequency
sample
haplotype
hap1
hap2
hap3

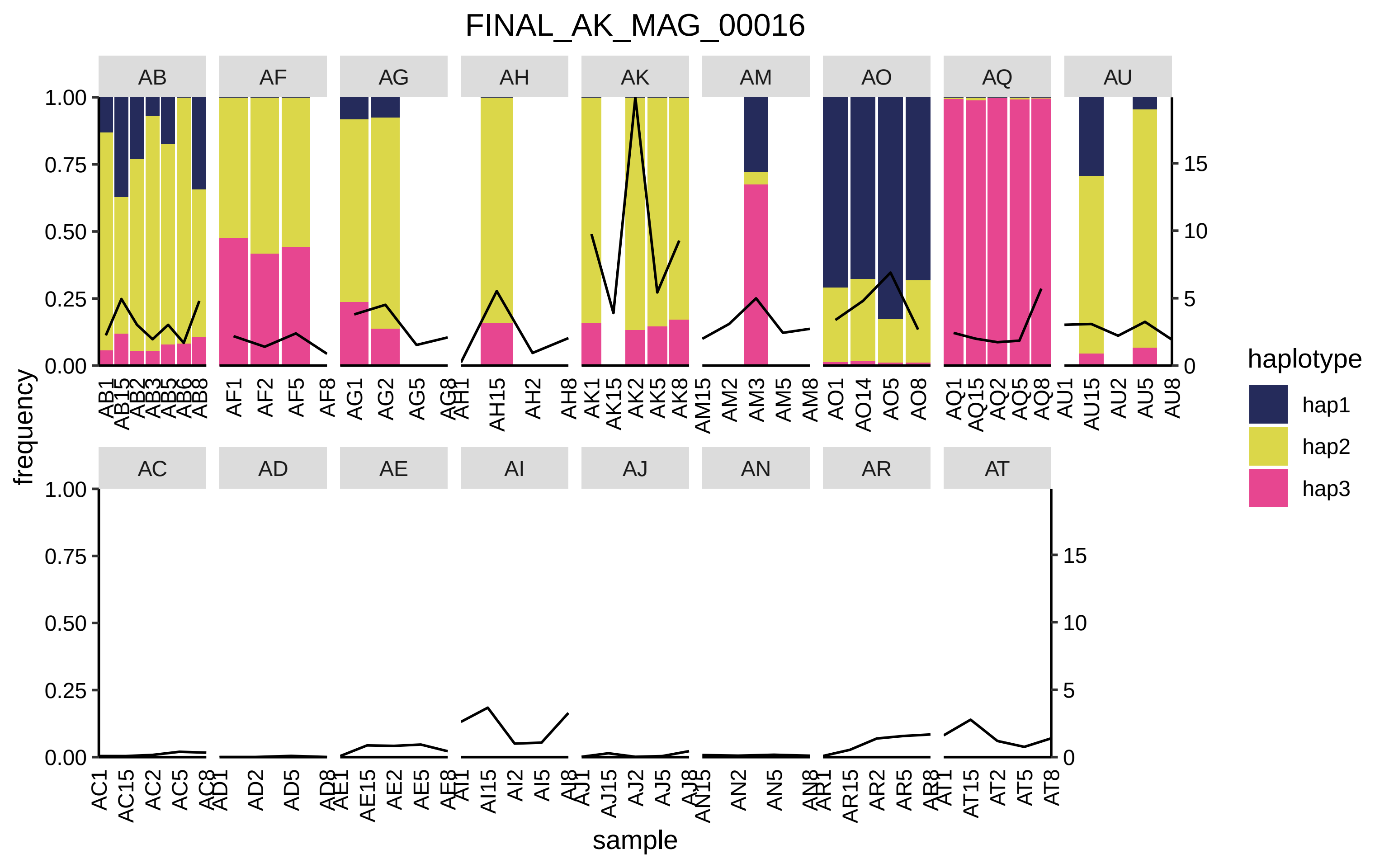

FINAL_AK_MAG_00016
AB
AF
AG
AH
AK
AM
AO
AQ
AU
AC
AD
AE
AI
AJ
AN
AR
AT
frequency
sample
haplotype
hap1
hap2
hap3

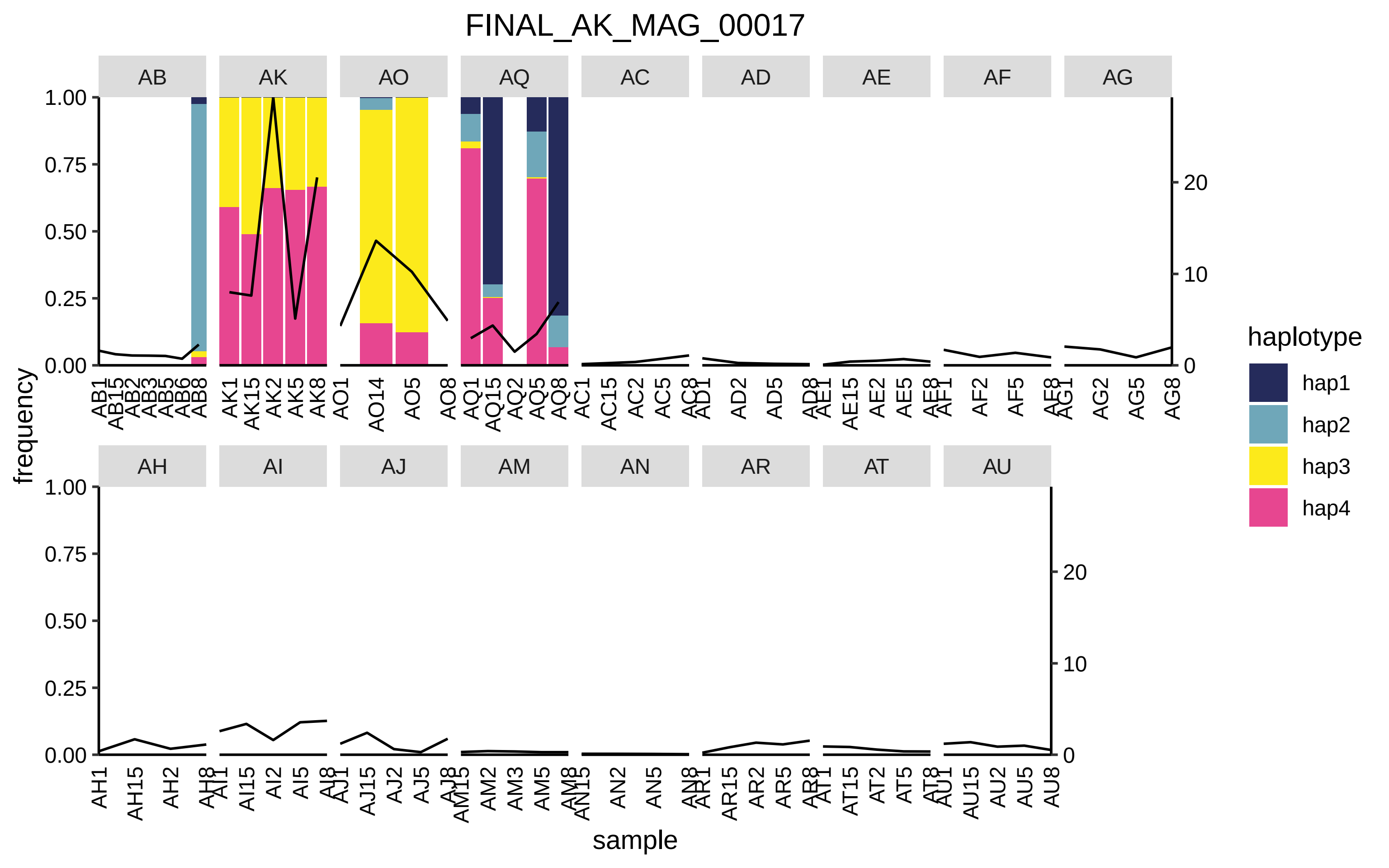
FINAL_AK_MAG_00017
AB
AK
AO
AQ
AC
AD
AE
AF
AG
AH
AI
AJ
AM
AN
AR
AT
AU
frequency
sample
haplotype
hap1
hap2
hap3
hap4
1.00
0.75
0.50
0.25
0.00
20
10
0

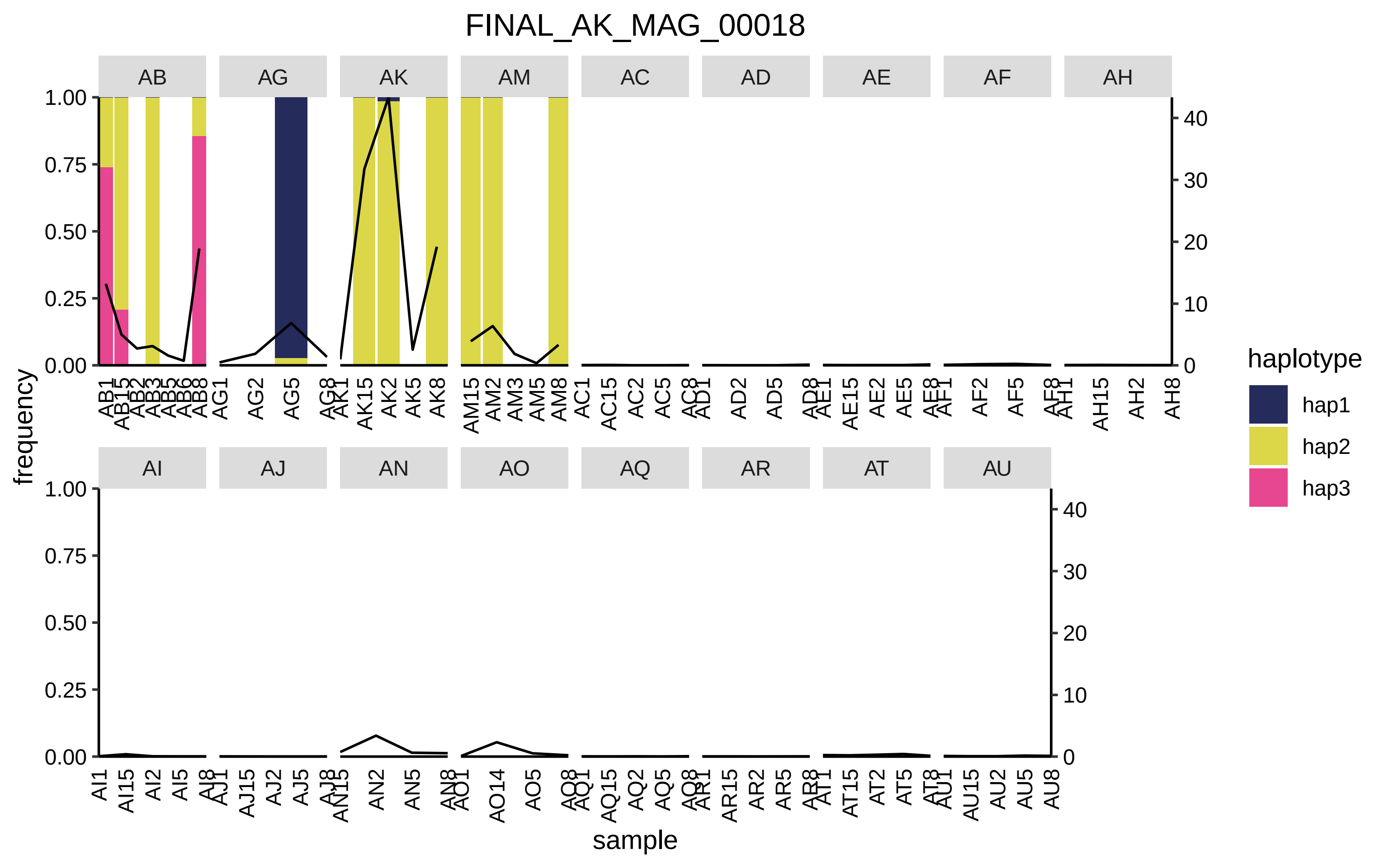
FINAL_AK_MAG_00018
AB
AG
AK
AM
AC
AD
AE
AF
AH
AI
AJ
AN
AO
AQ
AR
AT
AU
frequency
sample
haplotype
hap1
hap2
hap3

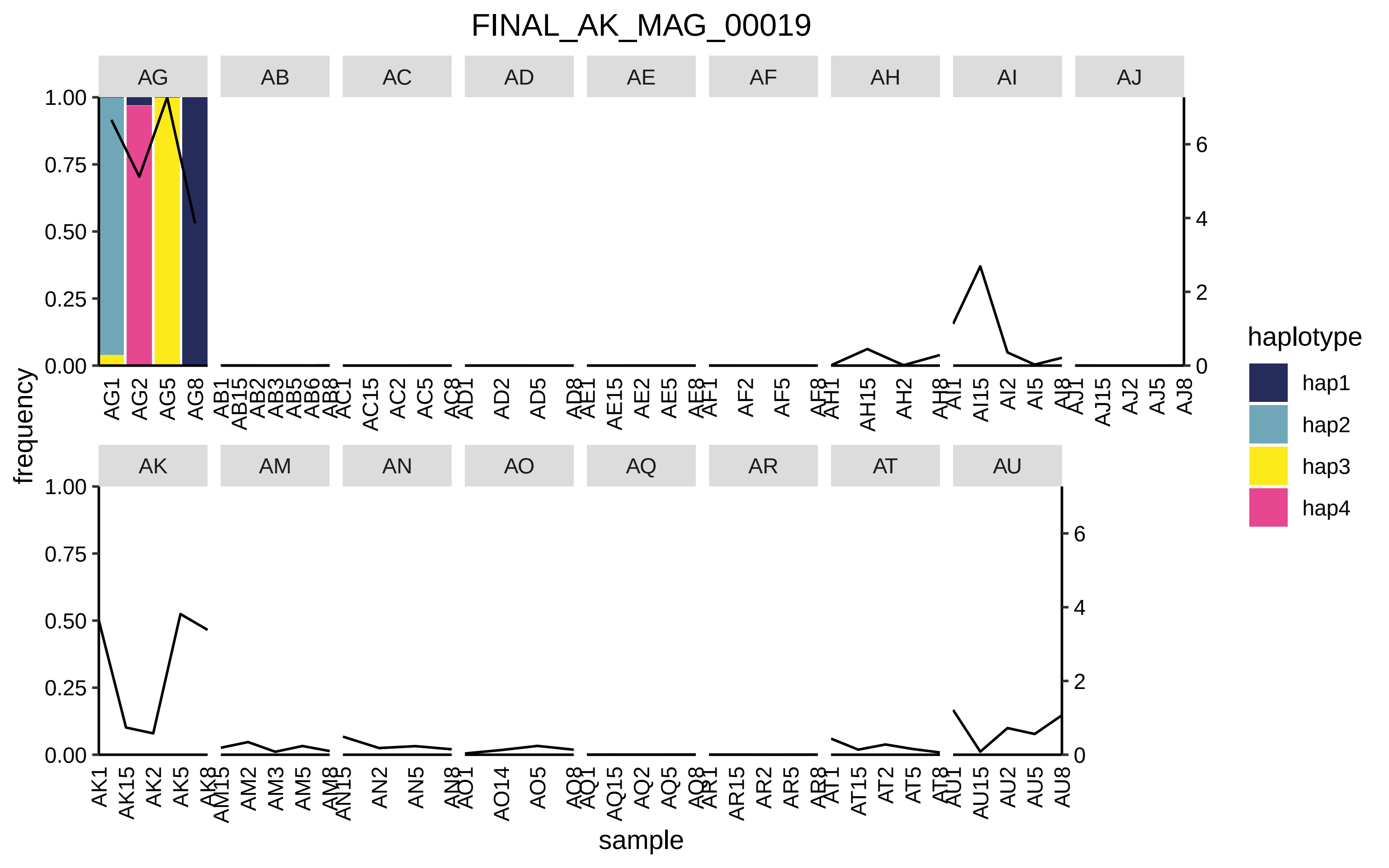

FINAL_AK_MAG_00019
AG
AB
AC
AD
AE
AF
AH
AI
AJ
AK
AM
AN
AO
AQ
AR
AT
AU
frequency
sample
haplotype
hap1
hap2
hap3
hap4

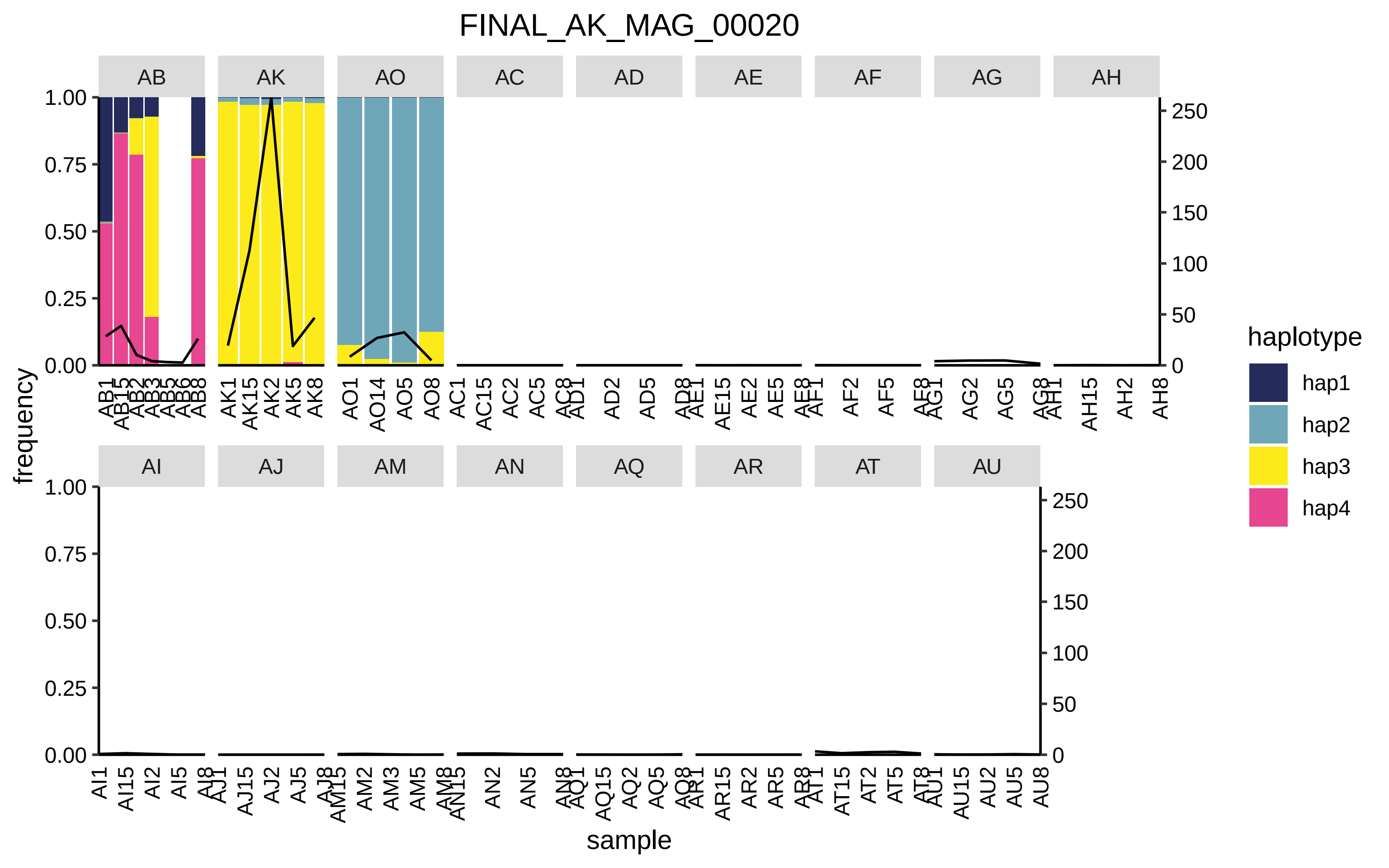
FINAL_AK_MAG_00020
AB
AK
AO
AC
AD
AE
AF
AG
AH
AI
AJ
AM
AN
AQ
AR
AT
AU
frequency
sample
haplotype
hap1
hap2
hap3
hap4
1.00
0.75
0.50
0.25
0.00
250
200
150
100
50
0

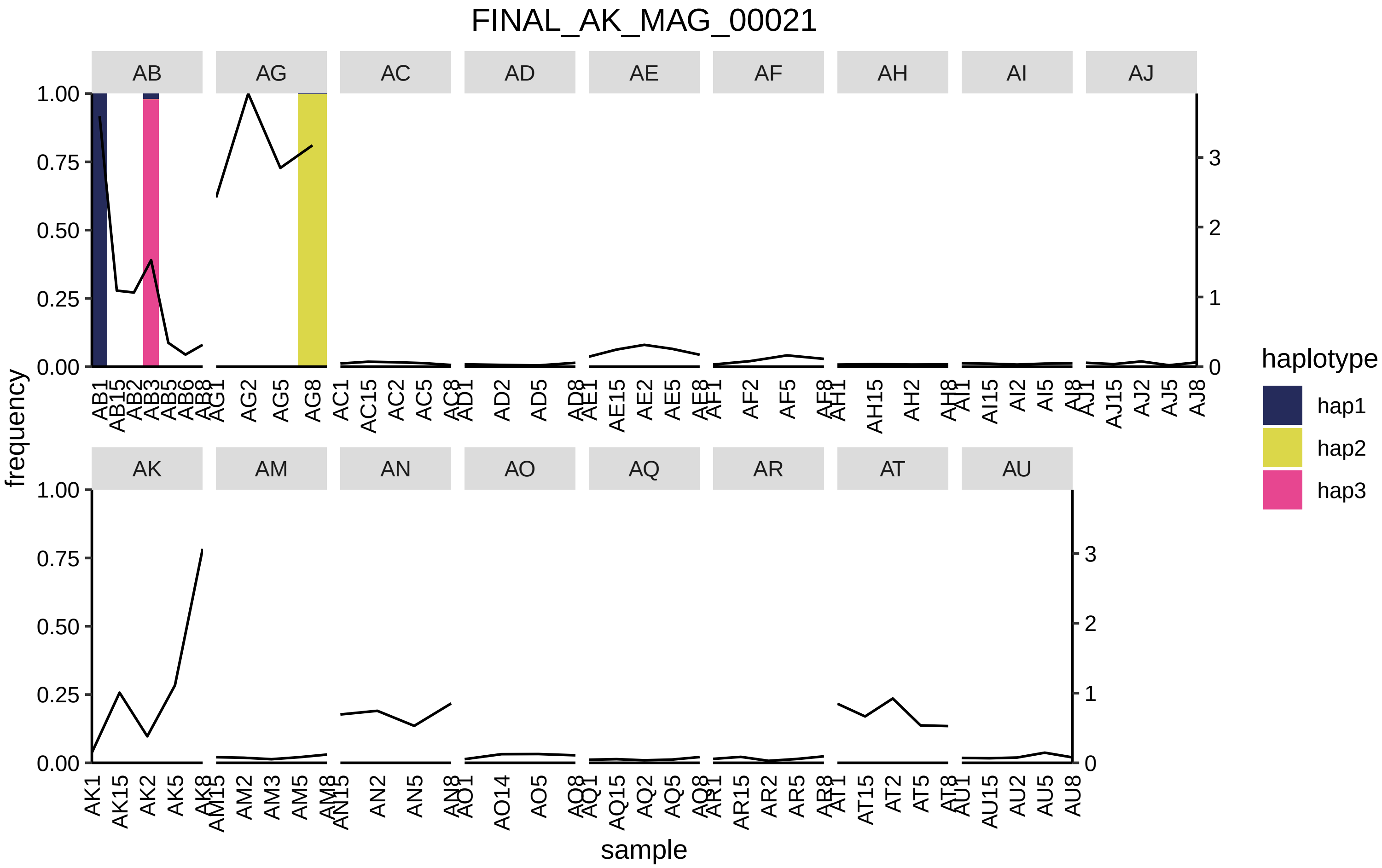

FINAL_AK_MAG_00021
AB
AG
AC
AD
AE
AF
AH
AI
AJ
AK
AM
AN
AO
AQ
AR
AT
AU
frequency
sample
haplotype
hap1
hap2
hap3

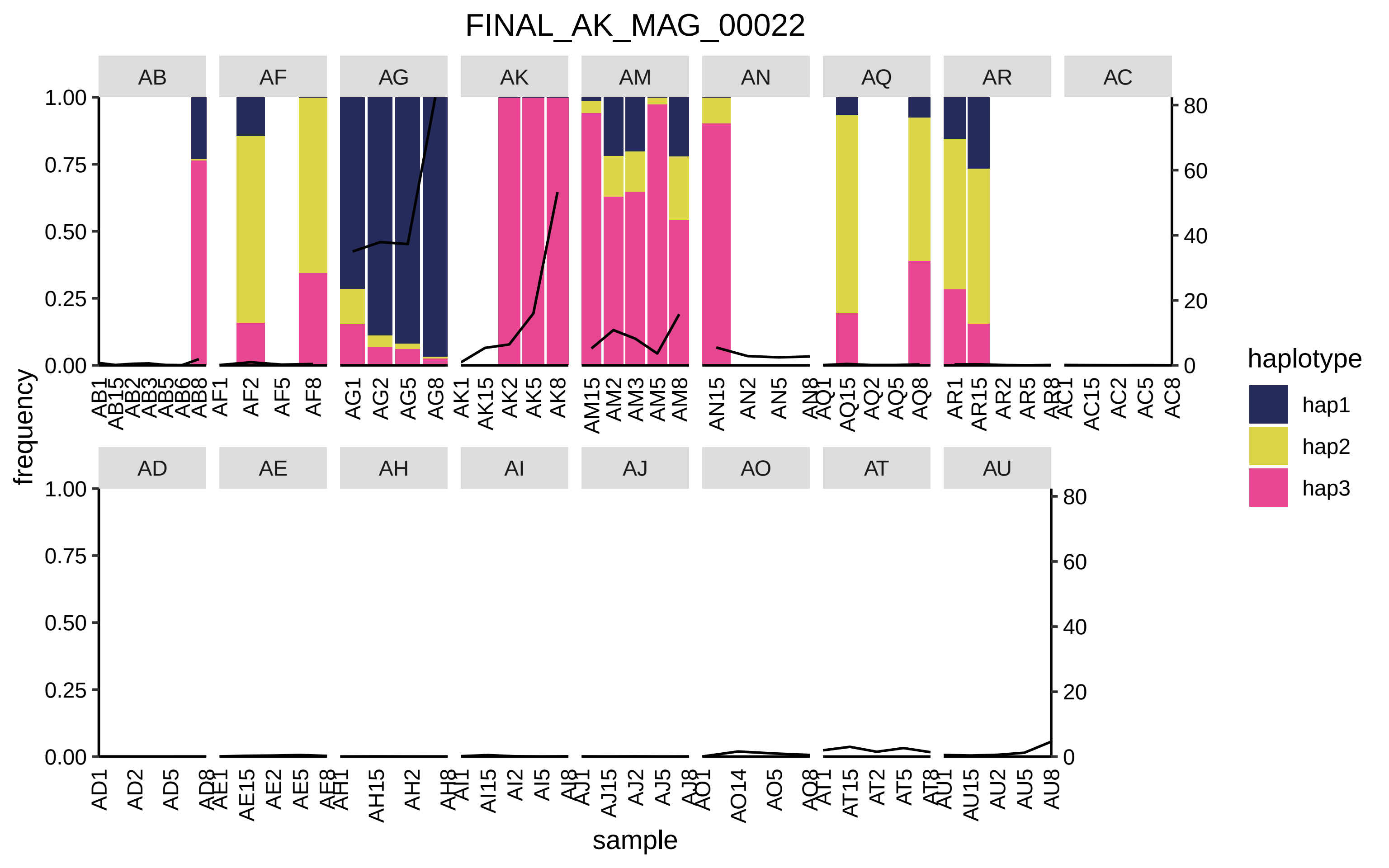
FINAL_AK_MAG_00022
AB
AF
AG
AK
AM
AN
AQ
AR
AC
AD
AE
AH
AI
AJ
AO
AT
AU
frequency
sample
haplotype
hap1
hap2
hap3

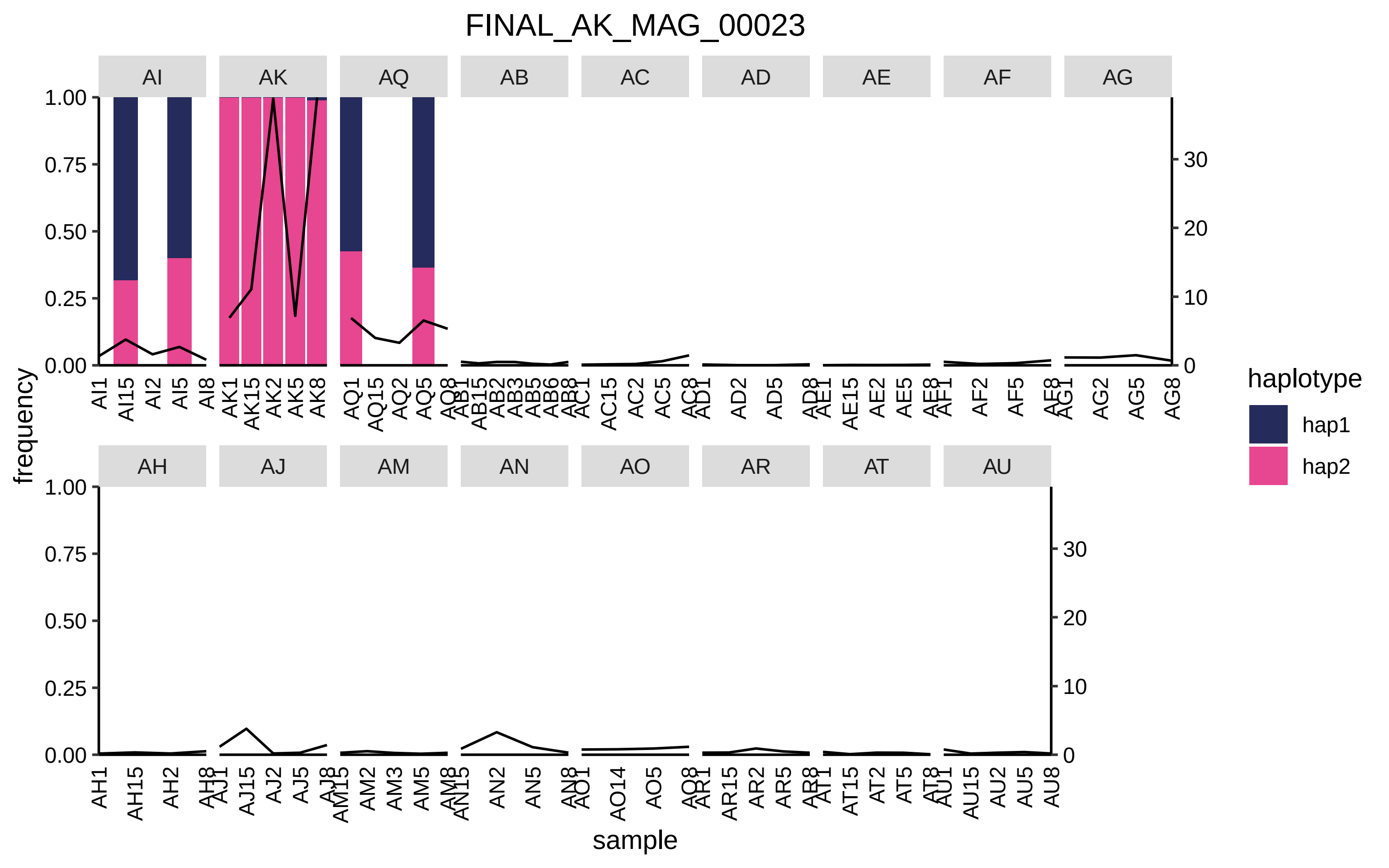

FINAL_AK_MAG_00023
AI
AK
AQ
AB
AC
AD
AE
AF
AG
AH
AJ
AM
AN
AO
AR
AT
AU
frequency
sample
haplotype
hap1
hap2
1.00
0.75
0.50
0.25
0.00
30
20
10
0
AI1 AI15 AI2 AI5 AI8
AK1 AK15 AK2 AK5 AK8
AQ1 AQ15 AQ2 AQ5 AQ8
AB1 AB15 AB2 AB3 AB5 AB6 AB8
AC1 AC15 AC2 AC5 AC8
AD1 AD2 AD5 AD8
AE1 AE15 AE2 AE5 AE8
AF1 AF2 AF5 AF8
AG1 AG2 AG5 AG8
AH1 AH15 AH2 AH8
AJ1 AJ15 AJ2 AJ5 AJ8
AM15 AM2 AM3 AM5 AM8
AN15 AN2 AN5 AN8
AO1 AO14 AO5 AO8
AR1 AR15 AR2 AR5 AR8
AT1 AT15 AT2 AT5 AT8
AU1 AU15 AU2 AU5 AU8

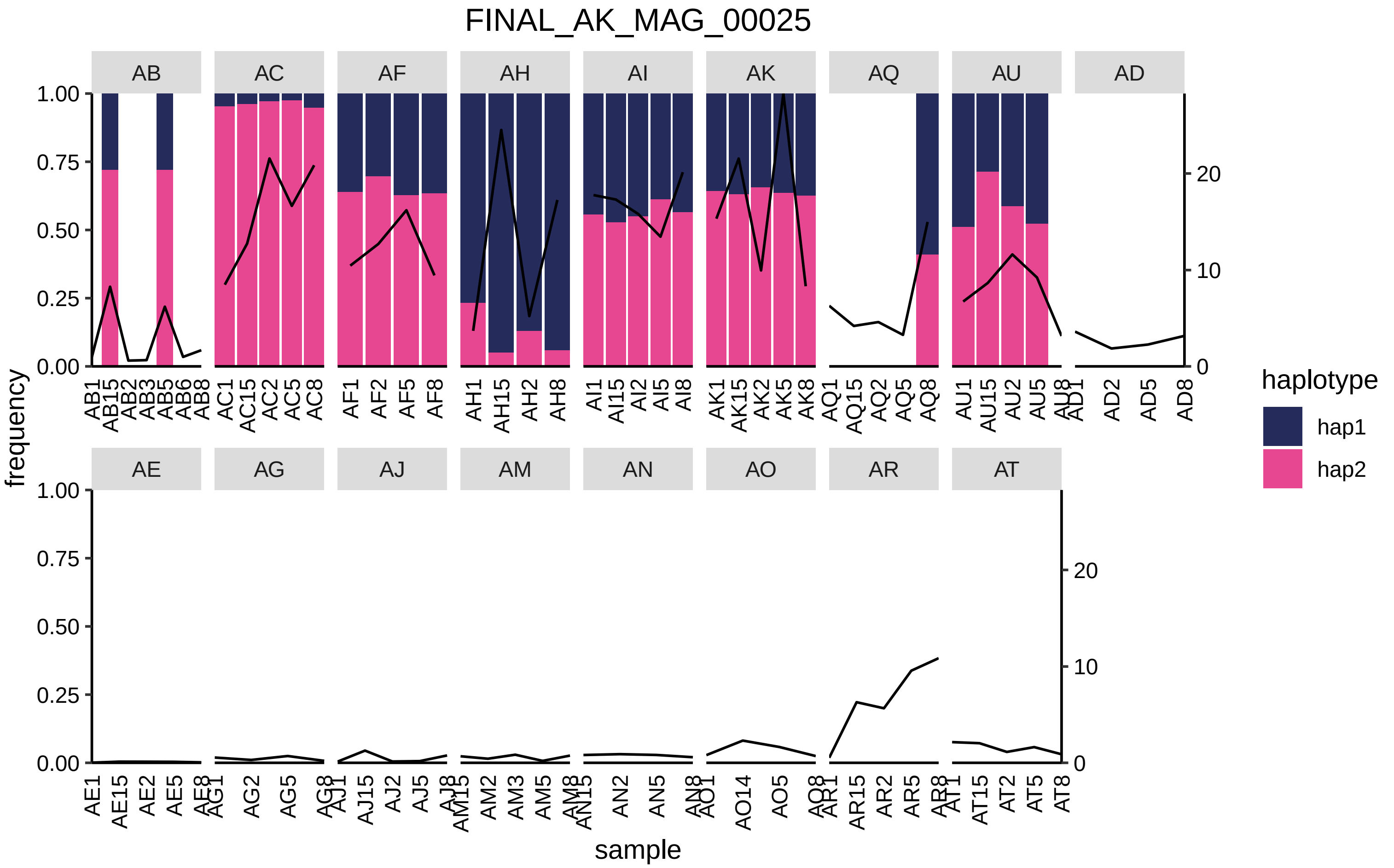

FINAL_AK_MAG_00025
AB
AC
AF
AH
AI
AK
AQ
AU
AD
AE
AG
AJ
AM
AN
AO
AR
AT
frequency
sample
haplotype
hap1
hap2

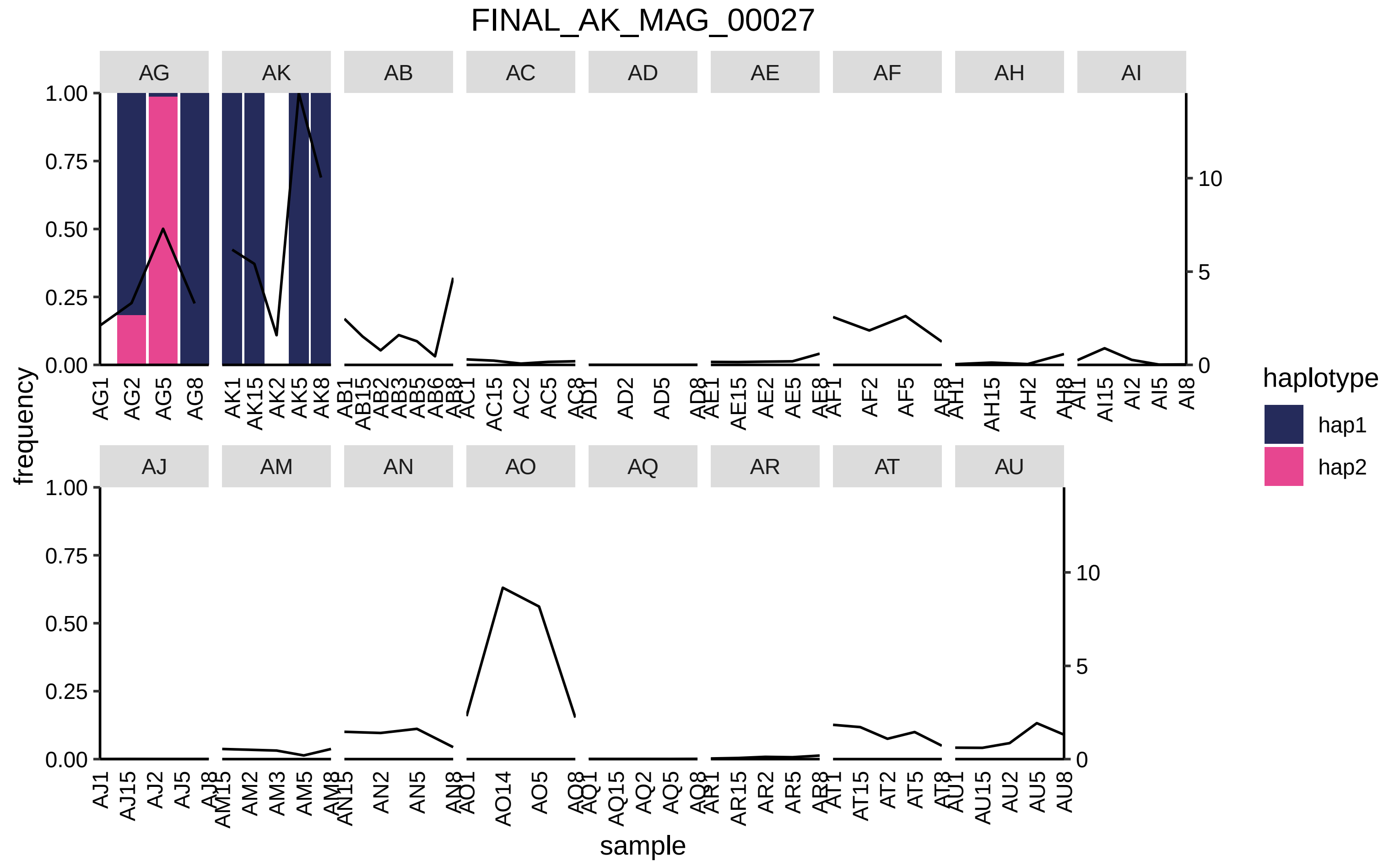
FINAL_AK_MAG_00027
AG
AK
AB
AC
AD
AE
AF
AH
AI
AJ
AM
AN
AO
AQ
AR
AT
AU
frequency
sample
haplotype
hap1
hap2
1.00
0.75
0.50
0.25
0.00
0
5
10

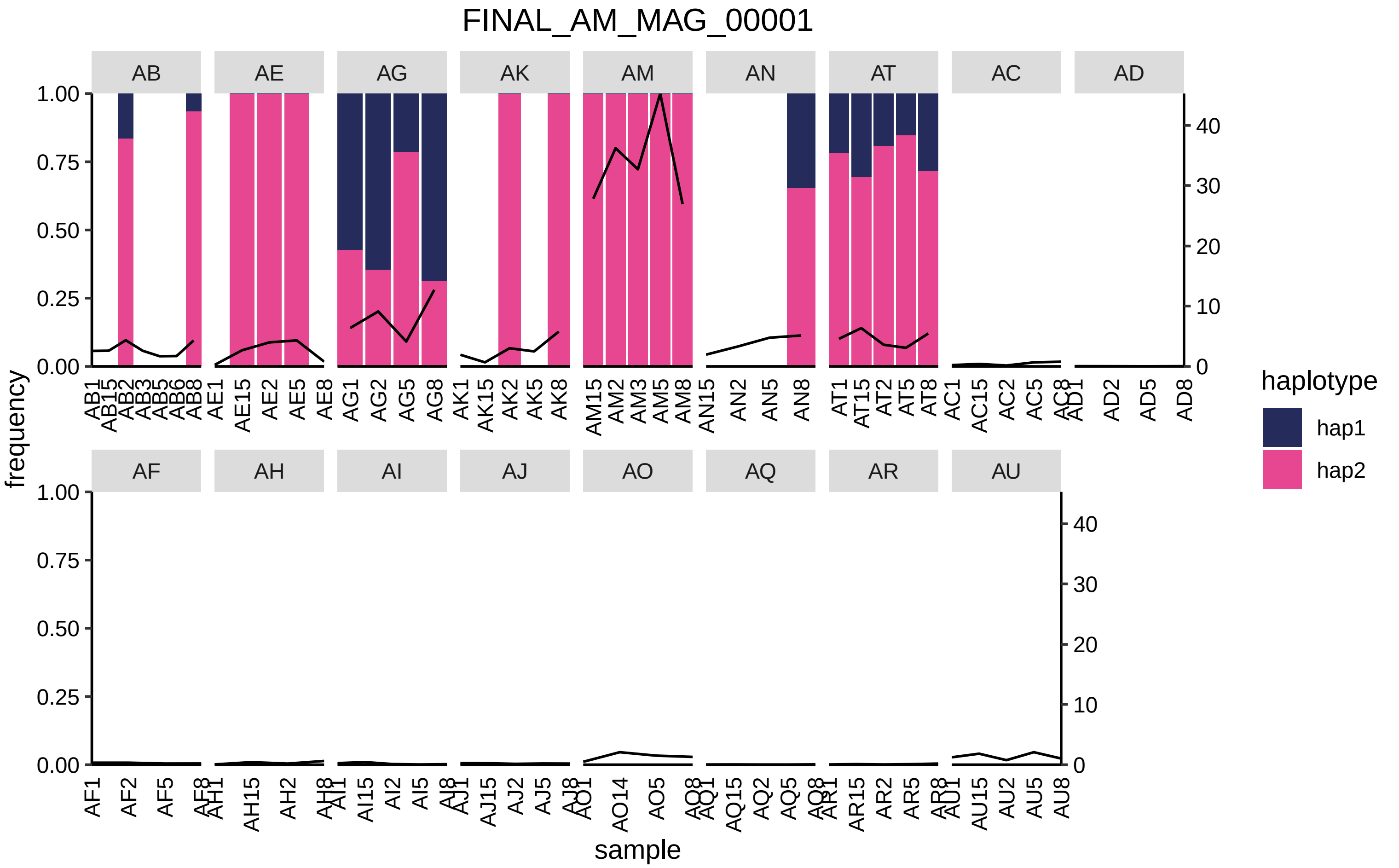
FINAL_AM_MAG_00001
AB
AE
AG
AK
AM
AN
AT
AC
AD
AF
AH
AI
AJ
AO
AQ
AR
AU
frequency
sample
haplotype
hap1
hap2

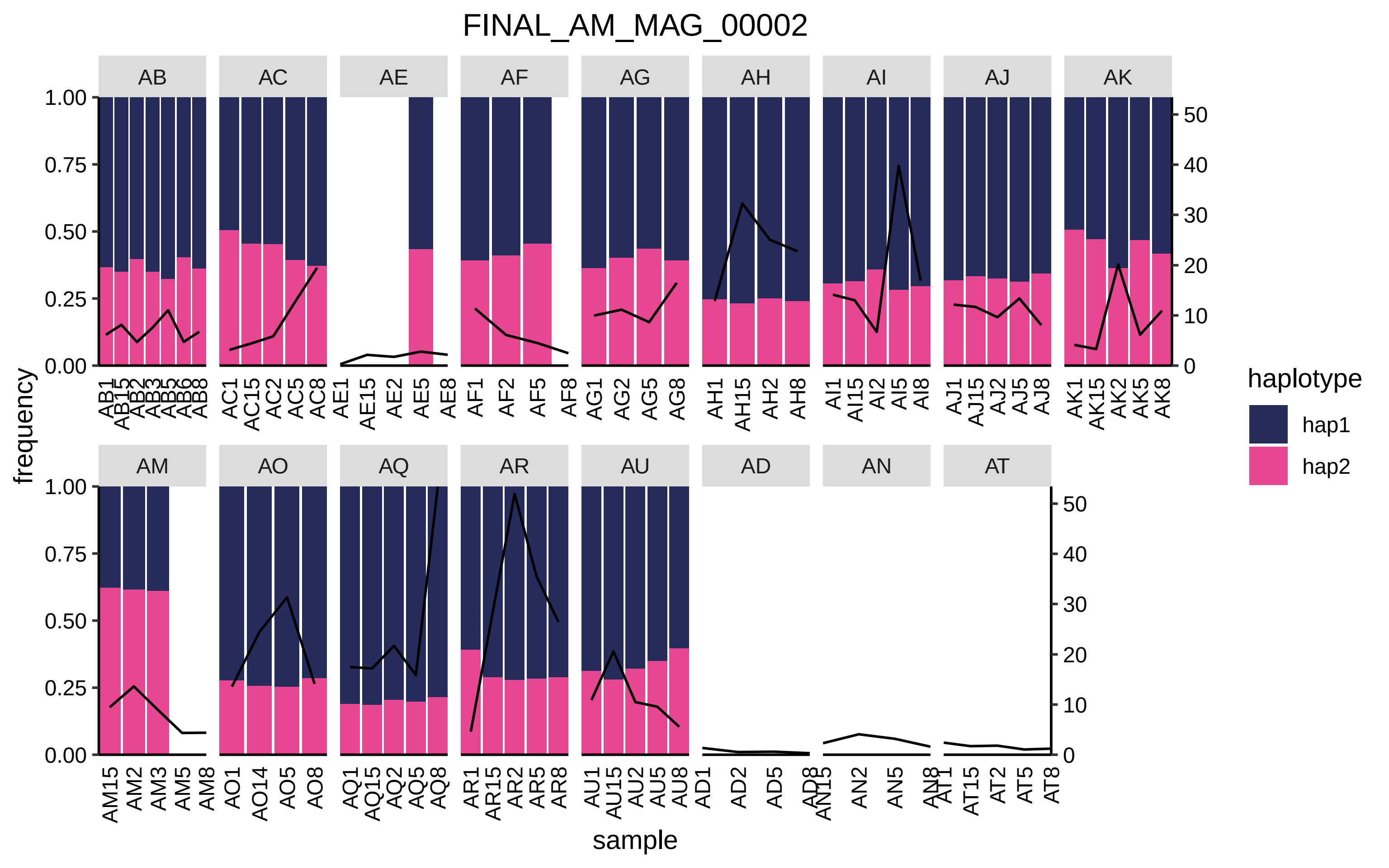

FINAL_AM_MAG_00002
AB
AC
AE
AF
AG
AH
AI
AJ
AK
AM
AO
AQ
AR
AU
AD
AN
AT
frequency
sample
1.00
0.75
0.50
0.25
0.00
50
40
30
20
10
0
AB1 AB15 AB2 AB3 AB5 AB6 AB8
AC1 AC15 AC2 AC5 AC8
AE1 AE15 AE2 AE5 AE8
AF1 AF2 AF5 AF8
AG1 AG2 AG5 AG8
AH1 AH15 AH2 AH8
AI1 AI15 AI2 AI5 AI8
AJ1 AJ15 AJ2 AJ5 AJ8
AK1 AK15 AK2 AK5 AK8
AM15 AM2 AM3 AM5 AM8
AO1 AO14 AO5 AO8
AQ1 AQ15 AQ2 AQ5 AQ8
AR1 AR15 AR2 AR5 AR8
AU1 AU15 AU2 AU5 AU8
AD1 AD2 AD5 AD8
AN15 AN2 AN5 AN8
AT1 AT15 AT2 AT5 AT8
haplotype
hap1
hap2

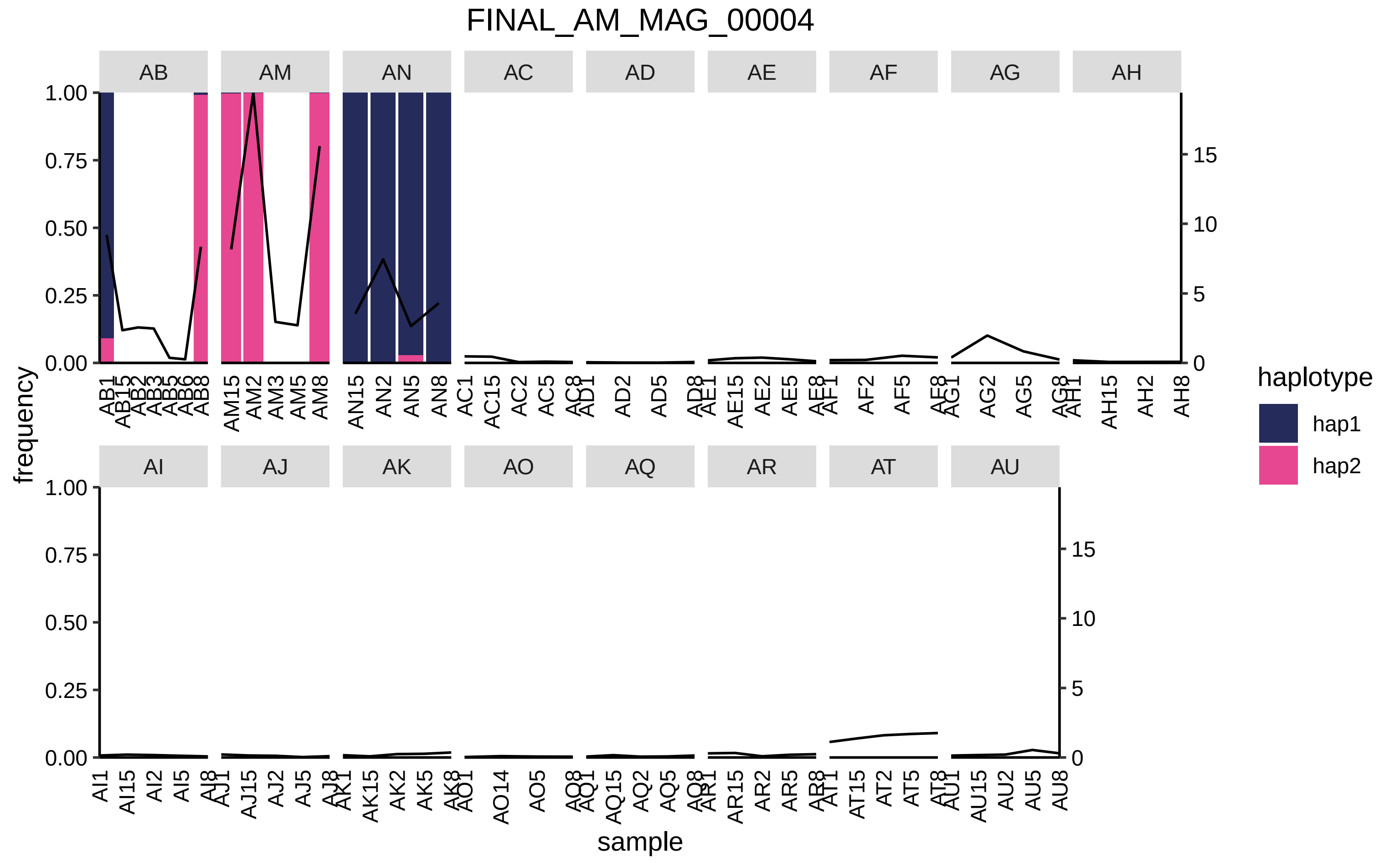

FINAL_AM_MAG_00004
AB
AM
AN
AC
AD
AE
AF
AG
AH
AI
AJ
AK
AO
AQ
AR
AT
AU
frequency
sample
haplotype
hap1
hap2
1.00
0.75
0.50
0.25
0.00
15
10
5
0

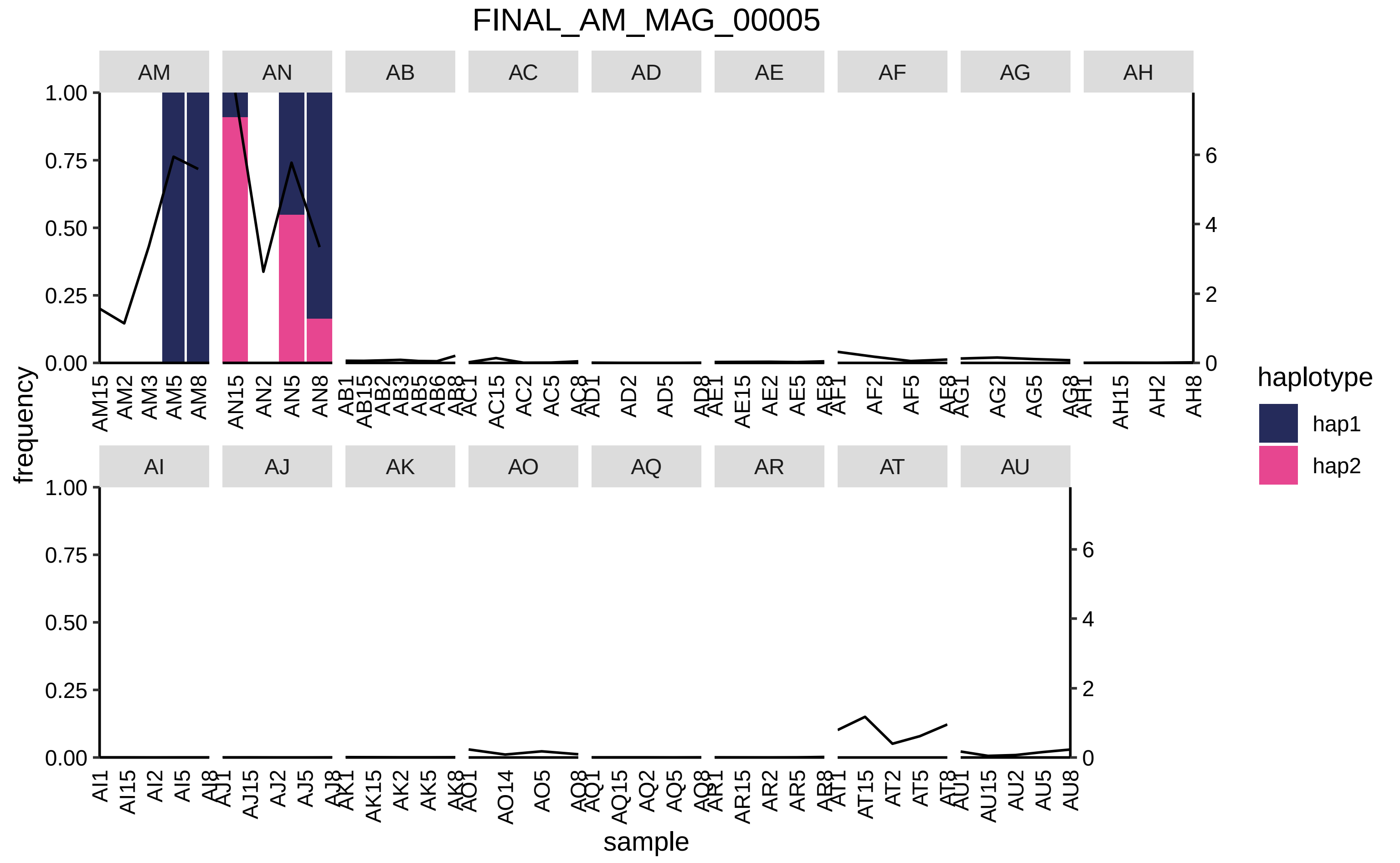

FINAL_AM_MAG_00005
AM
AN
AB
AC
AD
AE
AF
AG
AH
AI
AJ
AK
AO
AQ
AR
AT
AU
frequency
sample
haplotype
hap1
hap2

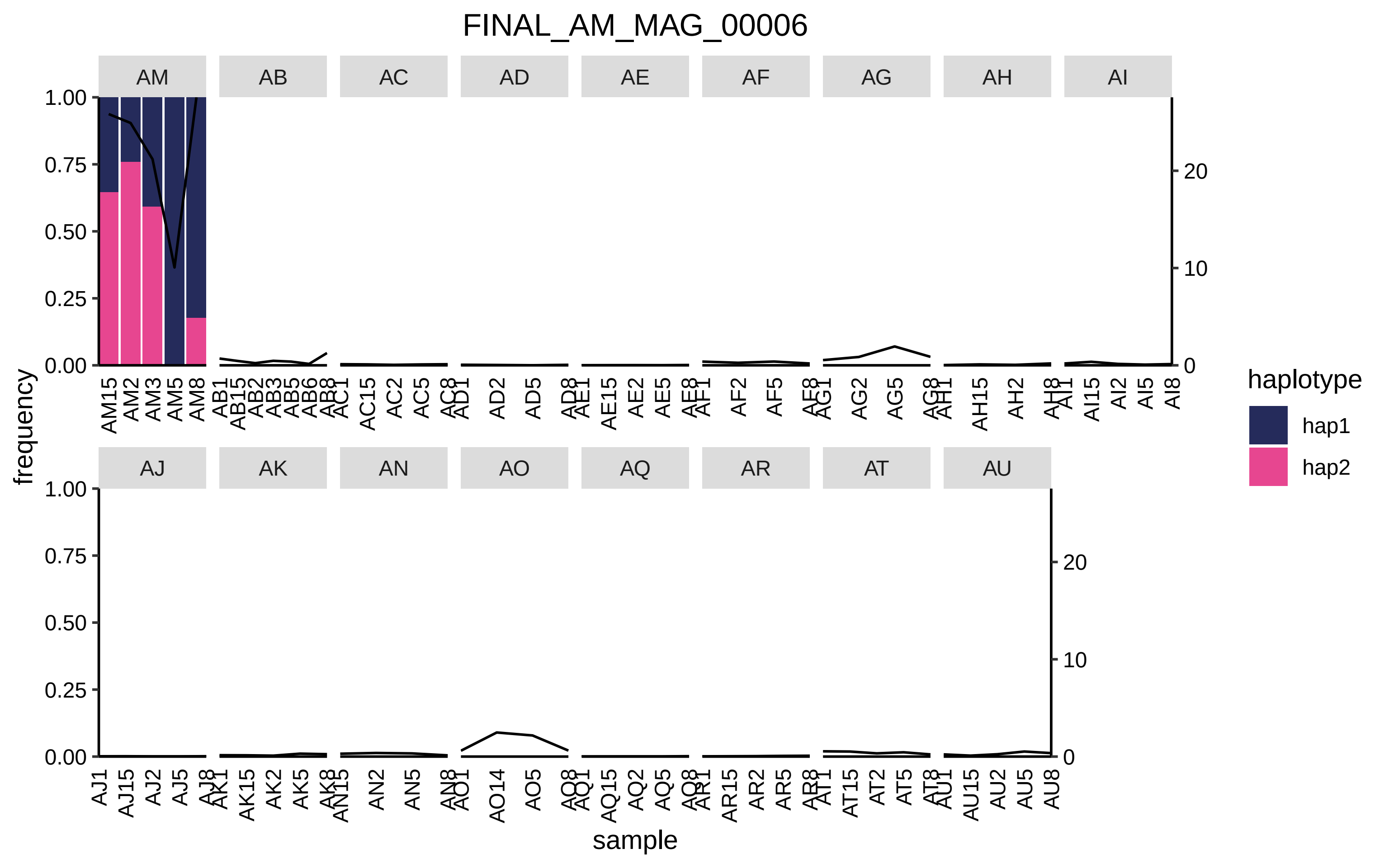
FINAL_AM_MAG_00006
AM
AB
AC
AD
AE
AF
AG
AH
AI
AJ
AK
AN
AO
AQ
AR
AT
AU
frequency
sample
haplotype
hap1
hap2
1.00
0.75
0.50
0.25
0.00
20
10
0
AM15 AM2 AM3 AM5 AM8
AB1 AB15 AB2 AB3 AB5 AB6 AB8
AC1 AC15 AC2 AC5 AC8
AD1 AD2 AD5 AD8
AE1 AE15 AE2 AE5 AE8
AF1 AF2 AF5 AF8
AG1 AG2 AG5 AG8
AH1 AH15 AH2 AH8
AI1 AI15 AI2 AI5 AI8
AJ1 AJ15 AJ2 AJ5 AJ8
AK1 AK15 AK2 AK5 AK8
AN15 AN2 AN5 AN8
AO1 AO14 AO5 AO8
AQ1 AQ15 AQ2 AQ5 AQ8
AR1 AR15 AR2 AR5 AR8
AT1 AT15 AT2 AT5 AT8
AU1 AU15 AU2 AU5 AU8

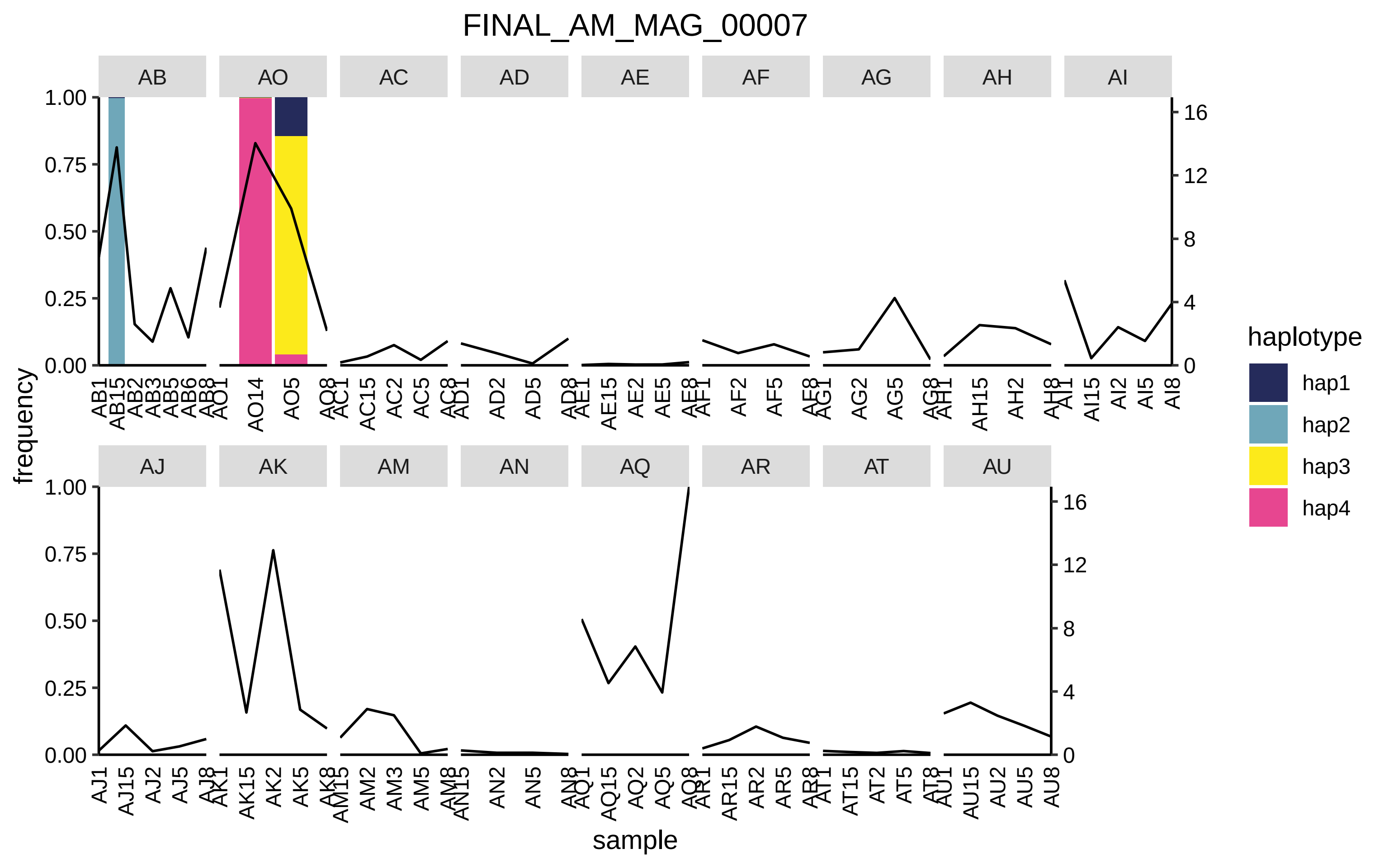

FINAL_AM_MAG_00007
AB
AO
AC
AD
AE
AF
AG
AH
AI
AJ
AK
AM
AN
AQ
AR
AT
AU
frequency
sample
haplotype
hap1
hap2
hap3
hap4

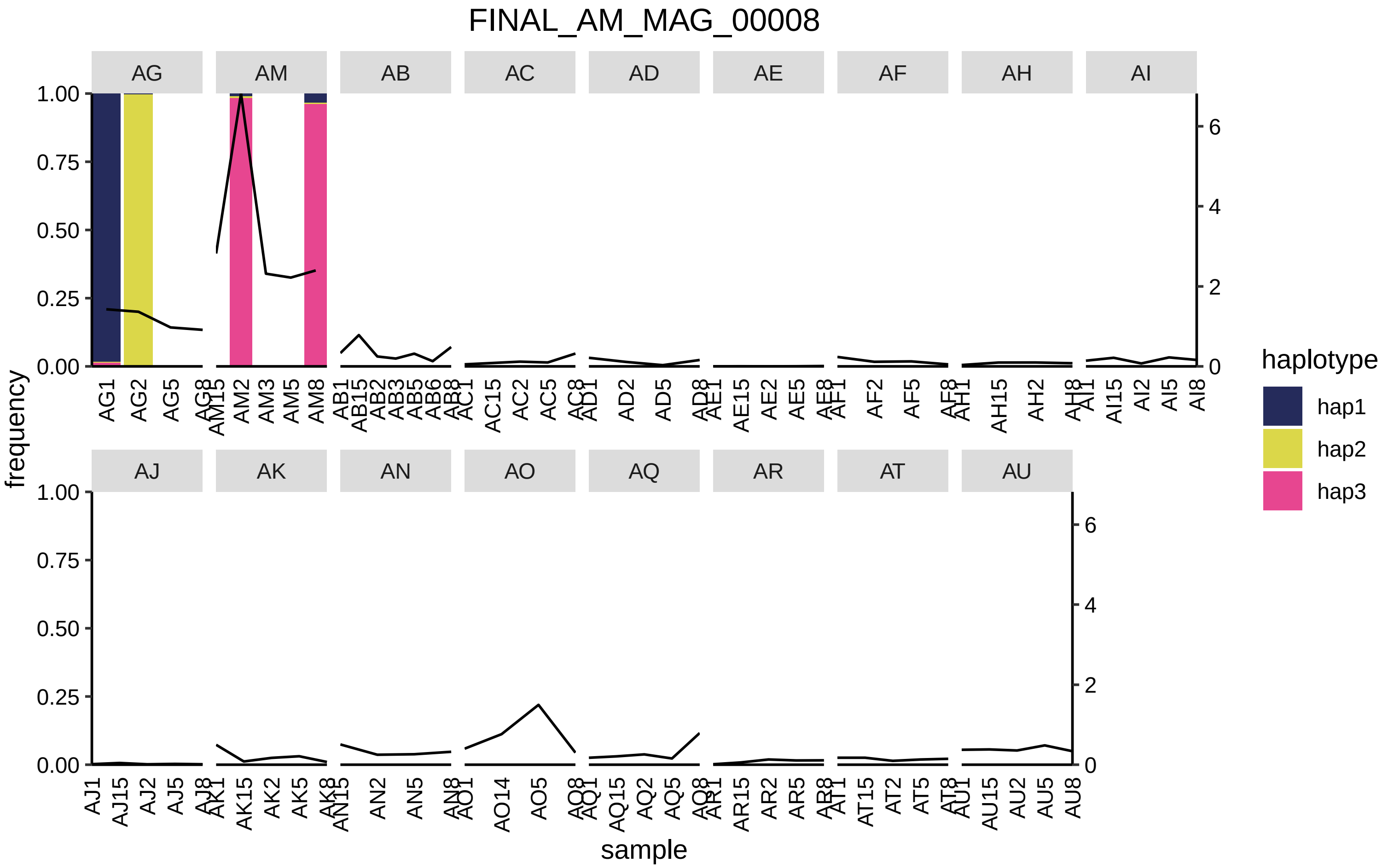
FINAL_AM_MAG_00008
AG
AM
AB
AC
AD
AE
AF
AH
AI
AJ
AK
AN
AO
AQ
AR
AT
AU
frequency
sample
haplotype
hap1
hap2
hap3
1.00
0.75
0.50
0.25
0.00
0
2
4
6

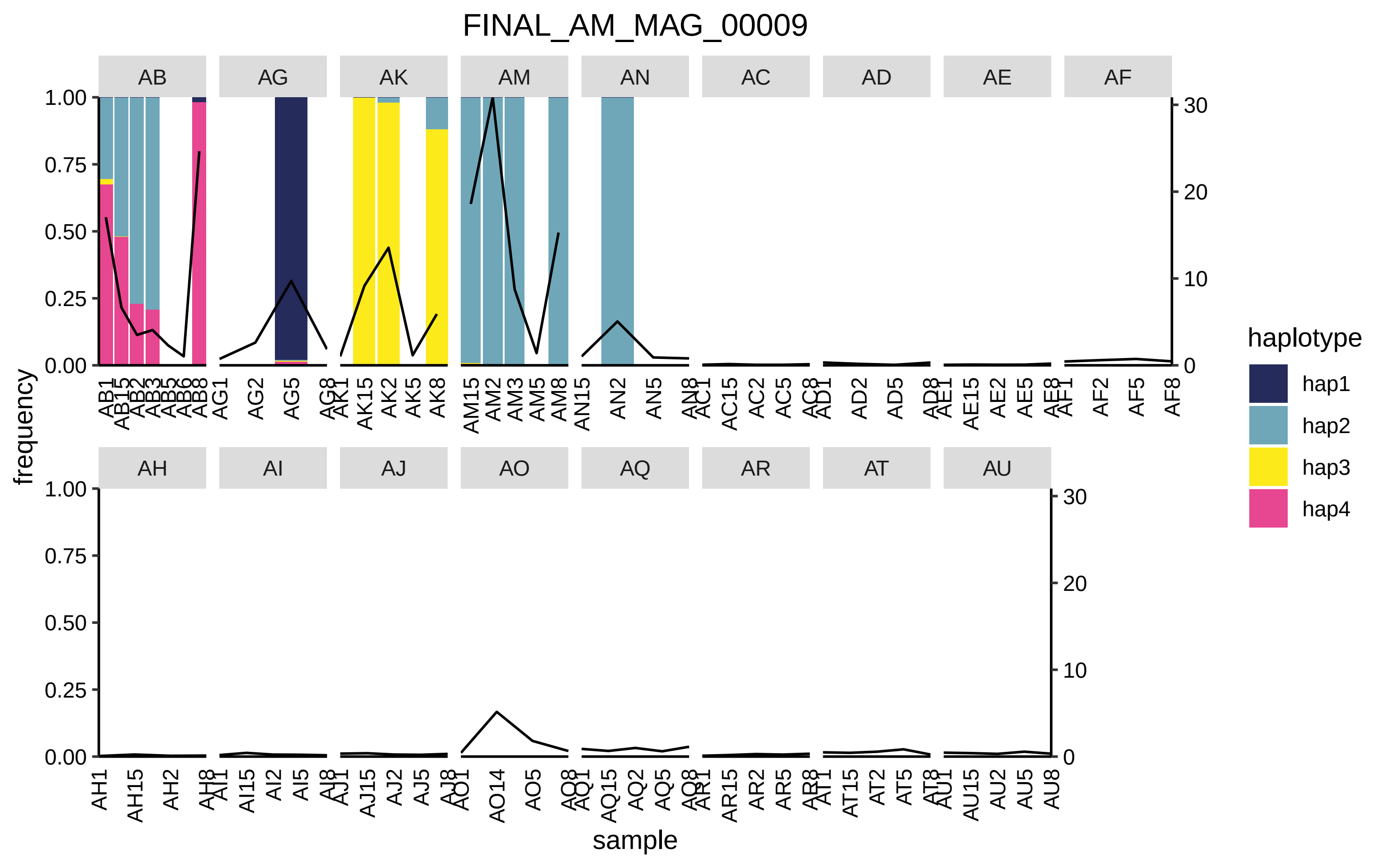
FINAL_AM_MAG_00009
AB
AG
AK
AM
AN
AC
AD
AE
AF
AH
AI
AJ
AO
AQ
AR
AT
AU
frequency
sample
haplotype
hap1
hap2
hap3
hap4
1.00
0.75
0.50
0.25
0.00
30
20
10
0
AB1 AB15 AB2 AB3 AB5 AB6 AB8 AG1 AG2 AG5 AG8 AK1 AK15 AK2 AK5 AK8 AM15 AM2 AM3 AM5 AM8 AN15 AN2 AN5 AN8 AC1 AC15 AC2 AC5 AC8 AD1 AD2 AD5 AD8 AE1 AE15 AE2 AE5 AE8 AF1 AF2 AF5 AF8
AH1 AH15 AH2 AH8 AI1 AI15 AI2 AI5 AI8 AJ1 AJ15 AJ2 AJ5 AJ8 AO1 AO14 AO5 AO8 AQ1 AQ15 AQ2 AQ5 AQ8 AR1 AR15 AR2 AR5 AR8 AT1 AT15 AT2 AT5 AT8 AU1 AU15 AU2 AU5 AU8

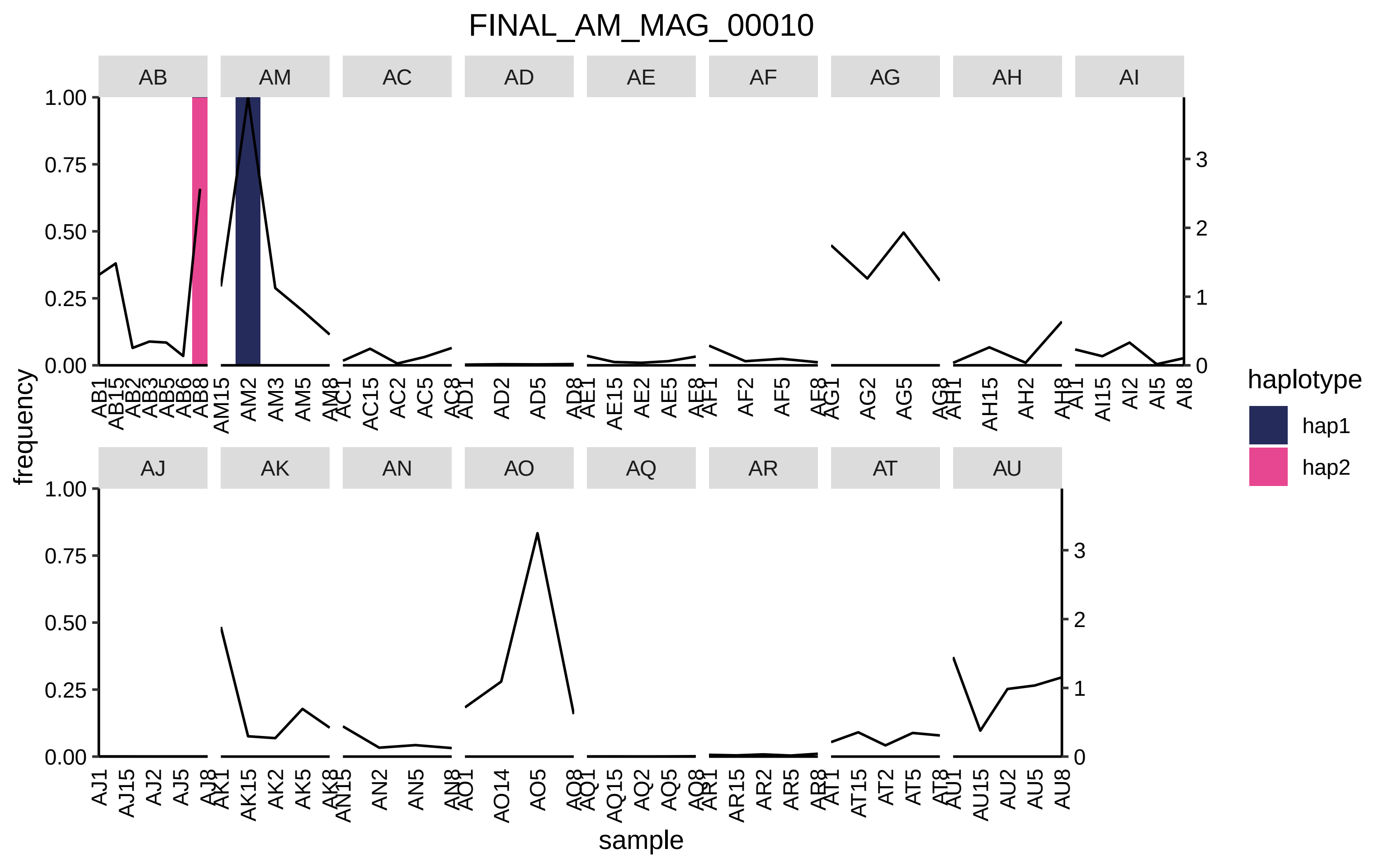

FINAL_AM_MAG_00010
AB
AM
AC
AD
AE
AF
AG
AH
AI
AJ
AK
AN
AO
AQ
AR
AT
AU
frequency
sample
haplotype
hap1
hap2

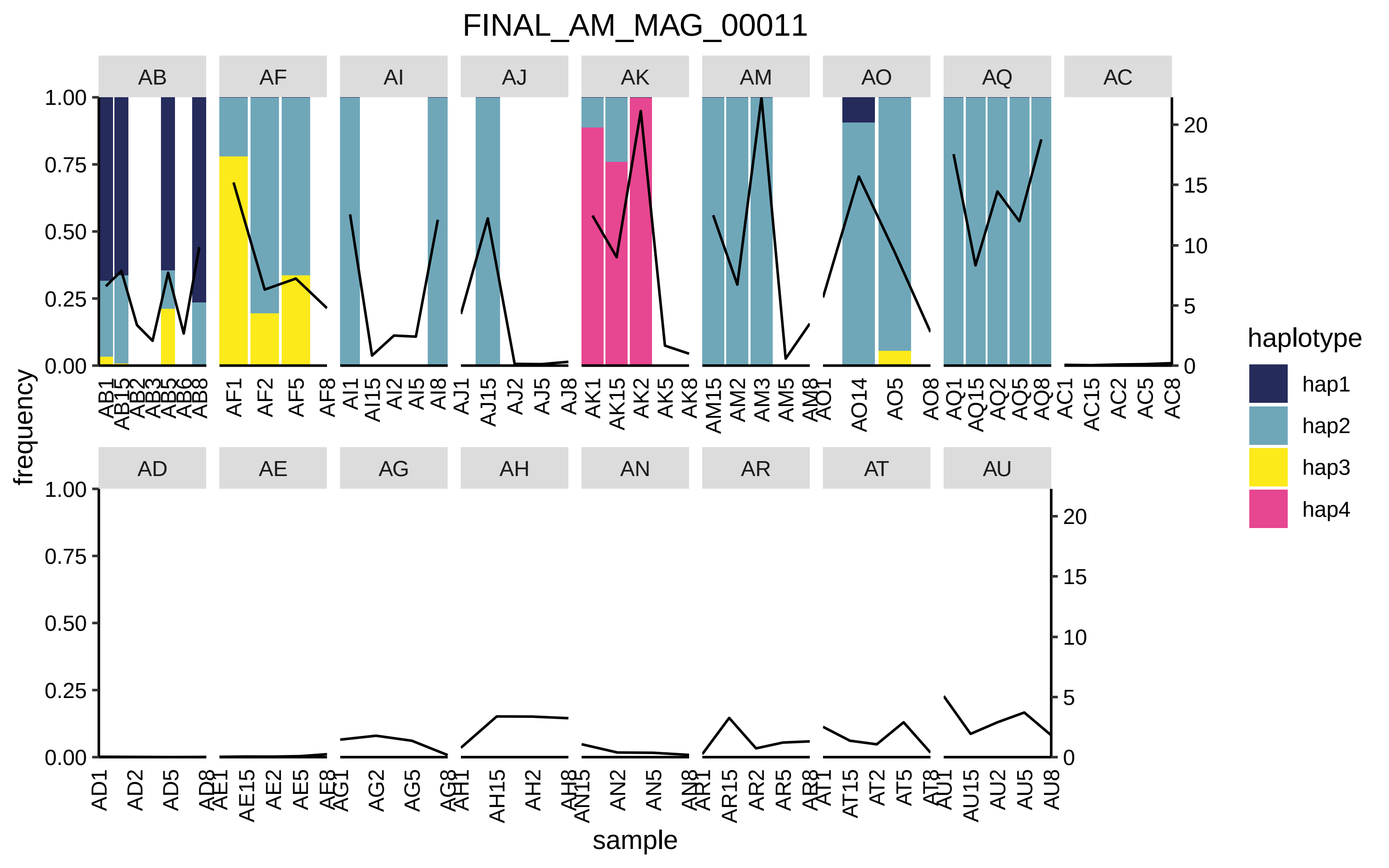

FINAL_AM_MAG_00011
AB
AF
AI
AJ
AK
AM
AO
AQ
AC
AD
AE
AG
AH
AN
AR
AT
AU
frequency
sample
haplotype
hap1
hap2
hap3
hap4

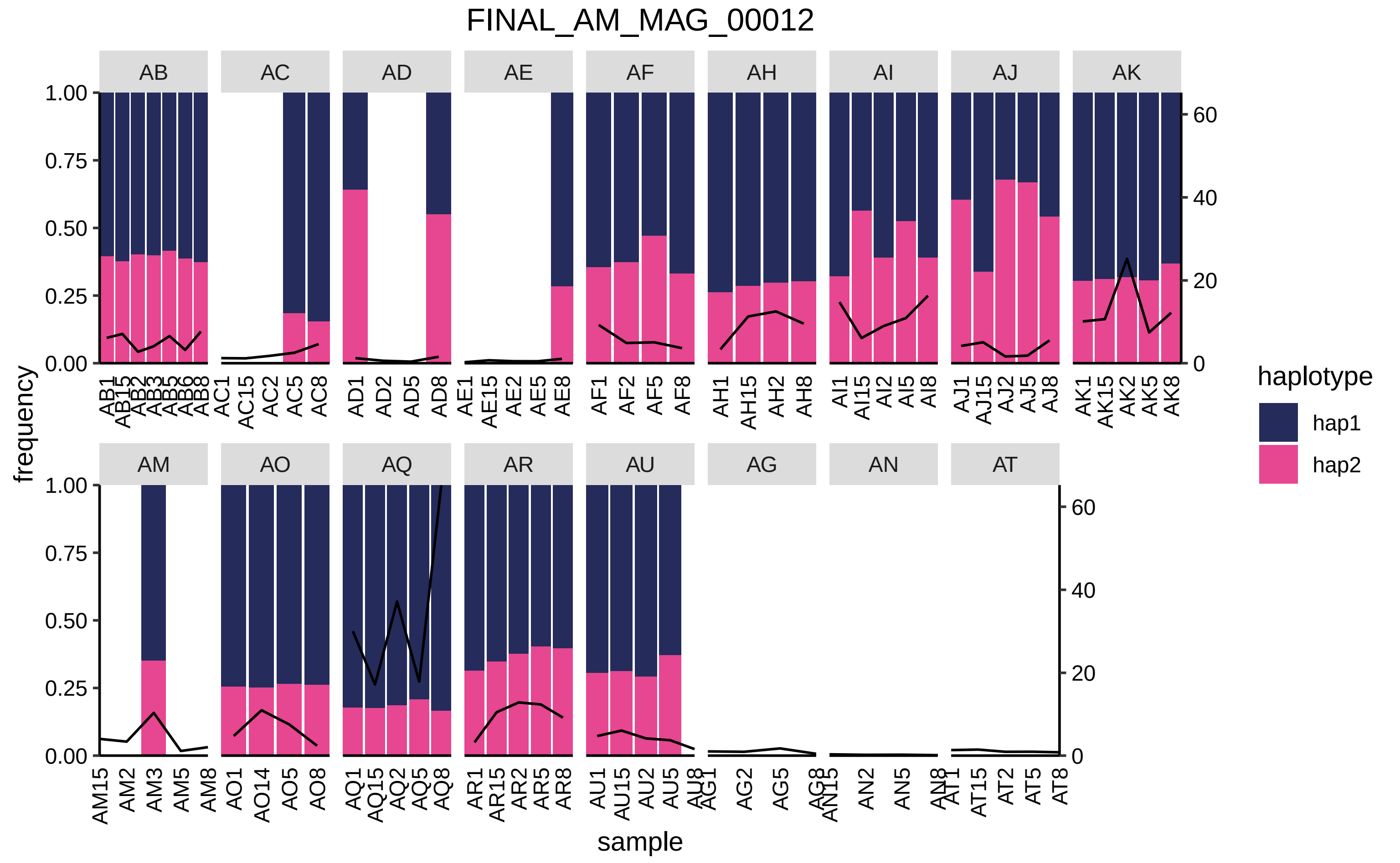

FINAL_AM_MAG_00012
AB
AC
AD
AE
AF
AH
AI
AJ
AK
AM
AO
AQ
AR
AU
AG
AN
AT
frequency
sample
1.00
0.75
0.50
0.25
0.00
60
40
20
0
AB1 AB15 AB2 AB3 AB5 AB6 AB8
AC1 AC15 AC2 AC5 AC8
AD1 AD2 AD5 AD8
AE1 AE15 AE2 AE5 AE8
AF1 AF2 AF5 AF8
AH1 AH15 AH2 AH8
AI1 AI15 AI2 AI5 AI8
AJ1 AJ15 AJ2 AJ5 AJ8
AK1 AK15 AK2 AK5 AK8
AM15 AM2 AM3 AM5 AM8
AO1 AO14 AO5 AO8
AQ1 AQ15 AQ2 AQ5 AQ8
AR1 AR15 AR2 AR5 AR8
AU1 AU15 AU2 AU5 AU8
AG1 AG2 AG5 AG8
AN15 AN2 AN5 AN8
AT1 AT15 AT2 AT5 AT8
haplotype
hap1
hap2

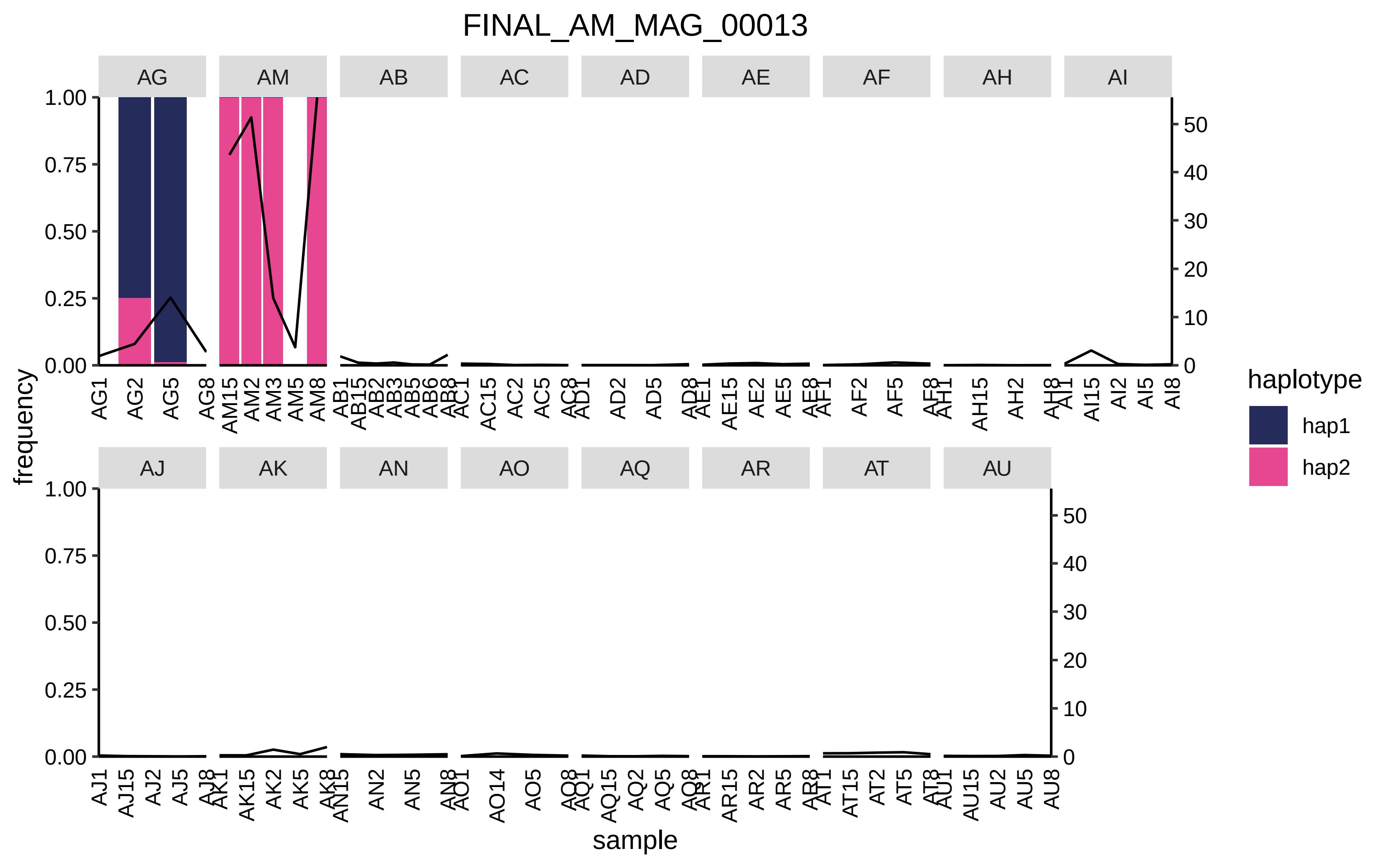

FINAL_AM_MAG_00013
AG
AM
AB
AC
AD
AE
AF
AH
AI
AJ
AK
AN
AO
AQ
AR
AT
AU
frequency
sample
haplotype
hap1
hap2

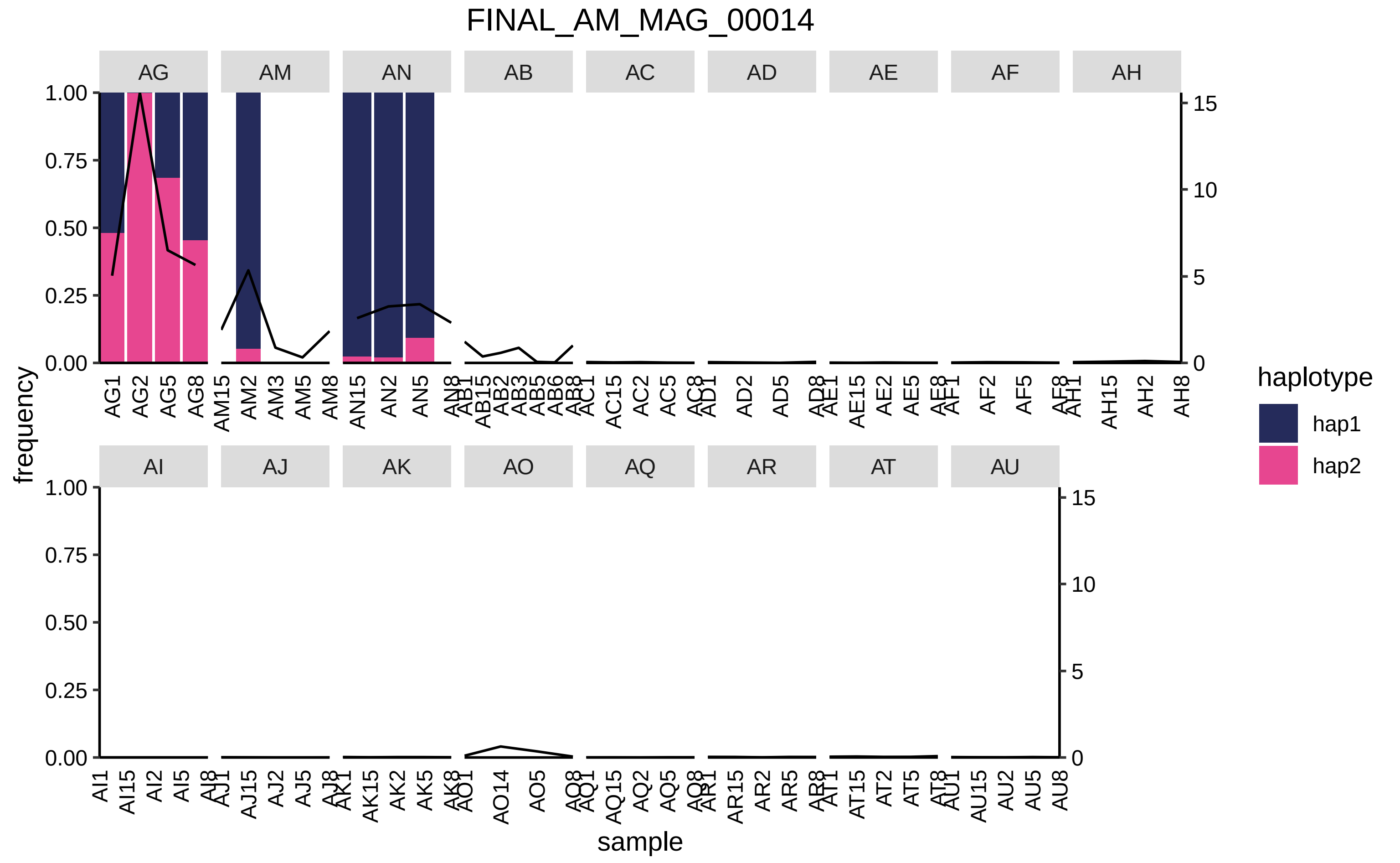
FINAL_AM_MAG_00014
AG
AM
AN
AB
AC
AD
AE
AF
AH
AI
AJ
AK
AO
AQ
AR
AT
AU
frequency
sample
haplotype
hap1
hap2
1.00
0.75
0.50
0.25
0.00
15
10
5
0

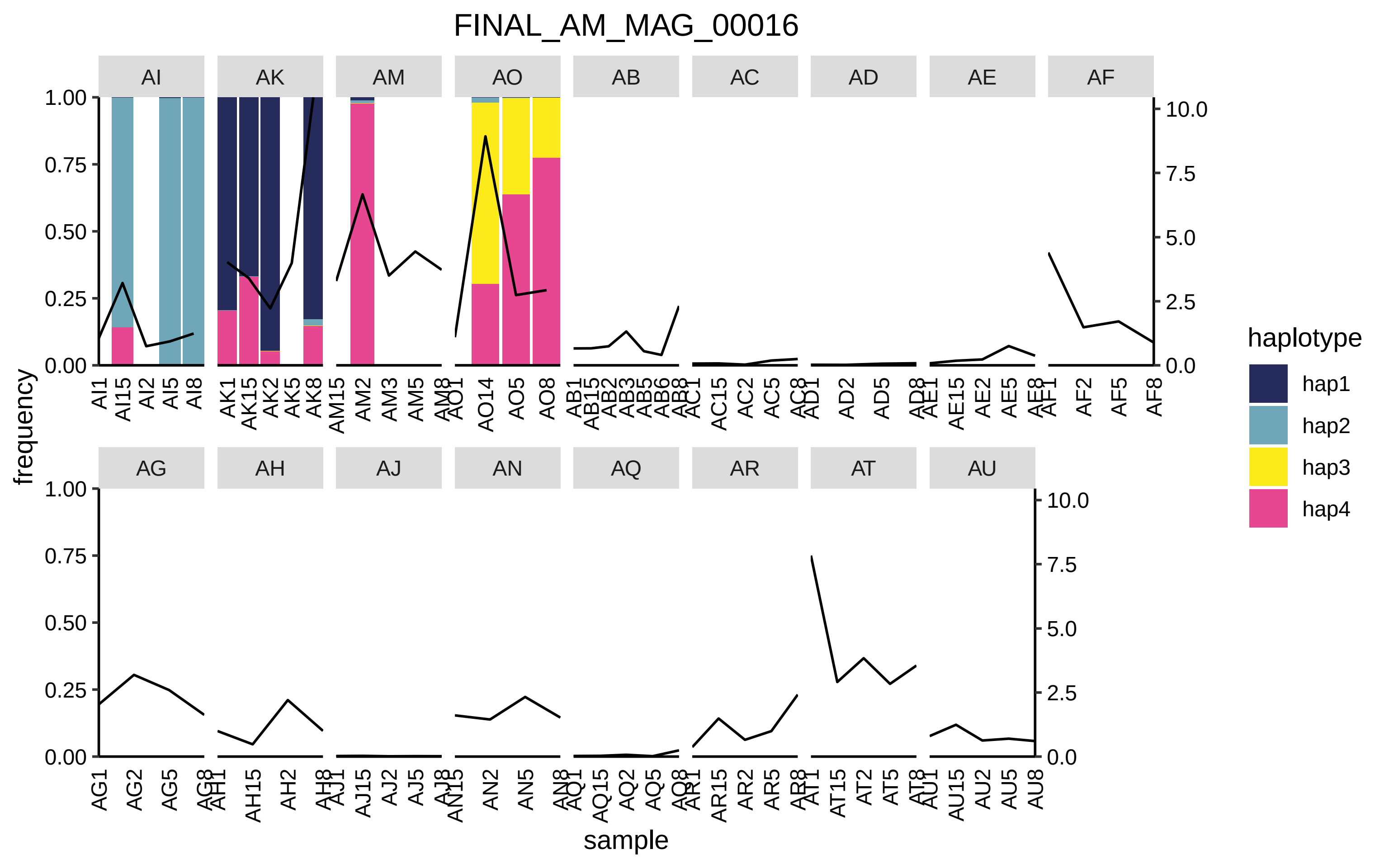

FINAL_AM_MAG_00016
AI
AK
AM
AO
AB
AC
AD
AE
AF
AG
AH
AJ
AN
AQ
AR
AT
AU
frequency
sample
haplotype
hap1
hap2
hap3
hap4

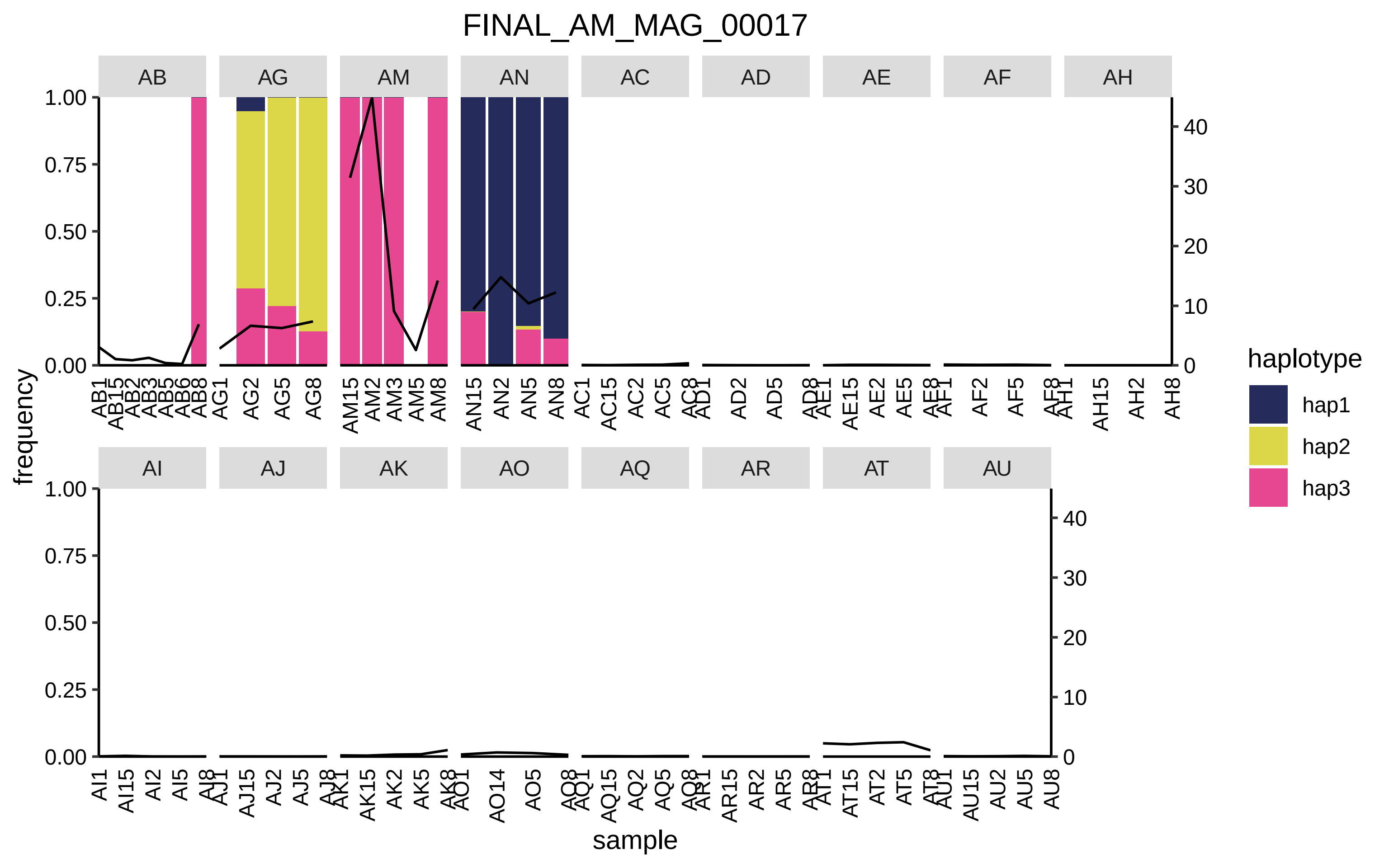

FINAL_AM_MAG_00017
AB
AG
AM
AN
AC
AD
AE
AF
AH
AI
AJ
AK
AO
AQ
AR
AT
AU
frequency
sample
1.00
0.75
0.50
0.25
0.00
40
30
20
10
0
AB1 AB15 AB2 AB3 AB5 AB6 AB8
AG1 AG2 AG5 AG8
AM15 AM2 AM3 AM5 AM8
AN15 AN2 AN5 AN8
AC1 AC15 AC2 AC5 AC8
AD1 AD2 AD5 AD8
AE1 AE15 AE2 AE5 AE8
AF1 AF2 AF5 AF8
AH1 AH15 AH2 AH8
AI1 AI15 AI2 AI5 AI8
AJ1 AJ15 AJ2 AJ5 AJ8
AK1 AK15 AK2 AK5 AK8
AO1 AO14 AO5 AO8
AQ1 AQ15 AQ2 AQ5 AQ8
AR1 AR15 AR2 AR5 AR8
AT1 AT15 AT2 AT5 AT8
AU1 AU15 AU2 AU5 AU8
haplotype
hap1
hap2
hap3

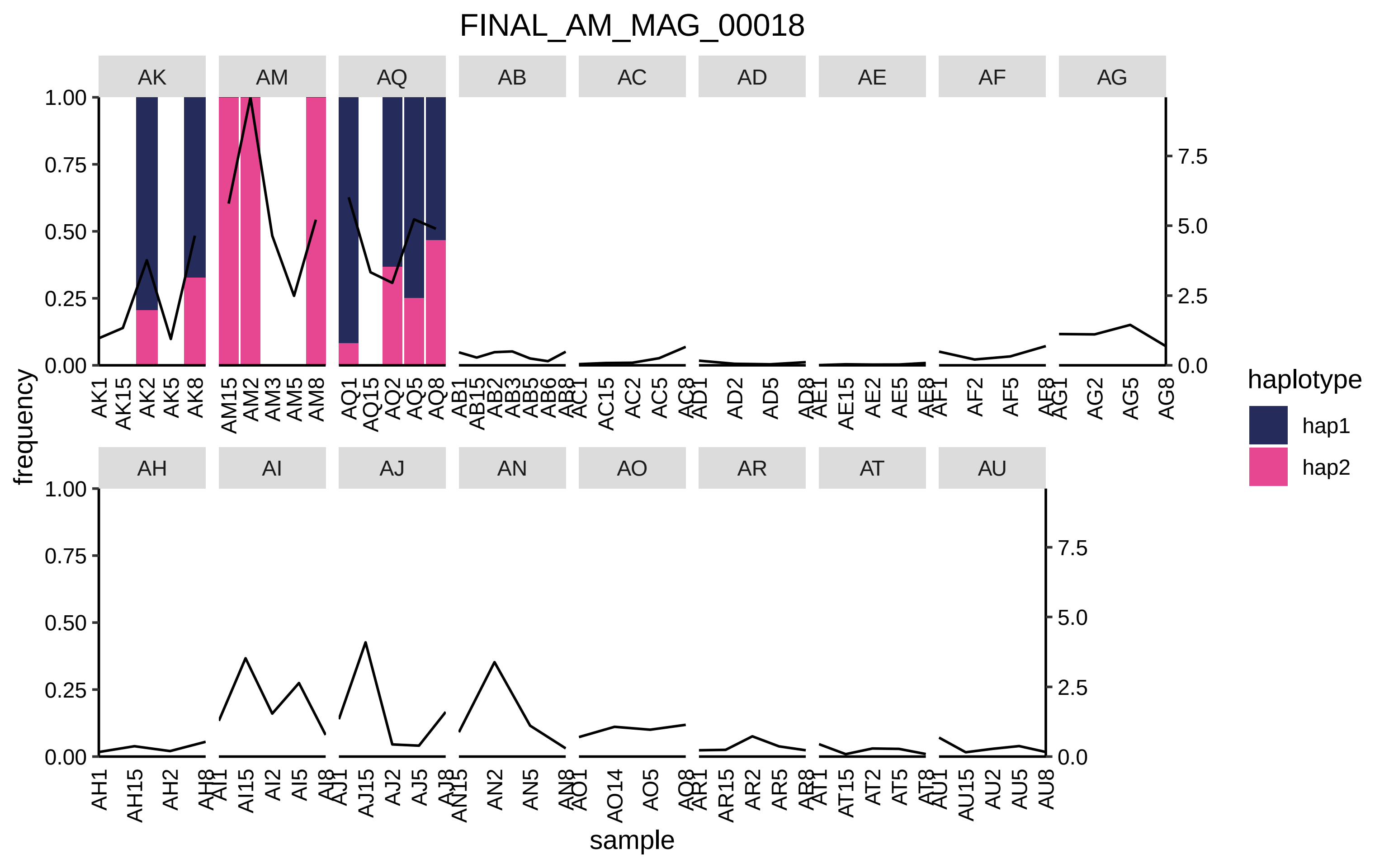
FINAL_AM_MAG_00018
AK
AM
AQ
AB
AC
AD
AE
AF
AG
AH
AI
AJ
AN
AO
AR
AT
AU
frequency
sample
haplotype
hap1
hap2
1.00
0.75
0.50
0.25
0.00
7.5
5.0
2.5
0.0
AK1 AK15 AK2 AK5 AK8
AM15 AM2 AM3 AM5 AM8
AQ1 AQ15 AQ2 AQ5 AQ8
AB1 AB15 AB2 AB3 AB5 AB6 AB8
AC1 AC15 AC2 AC5 AC8
AD1 AD2 AD5 AD8
AE1 AE15 AE2 AE5 AE8
AF1 AF2 AF5 AF8
AG1 AG2 AG5 AG8
AH1 AH15 AH2 AH8
AI1 AI15 AI2 AI5 AI8
AJ1 AJ15 AJ2 AJ5 AJ8
AN15 AN2 AN5 AN8
AO1 AO14 AO5 AO8
AR1 AR15 AR2 AR5 AR8
AT1 AT15 AT2 AT5 AT8
AU1 AU15 AU2 AU5 AU8

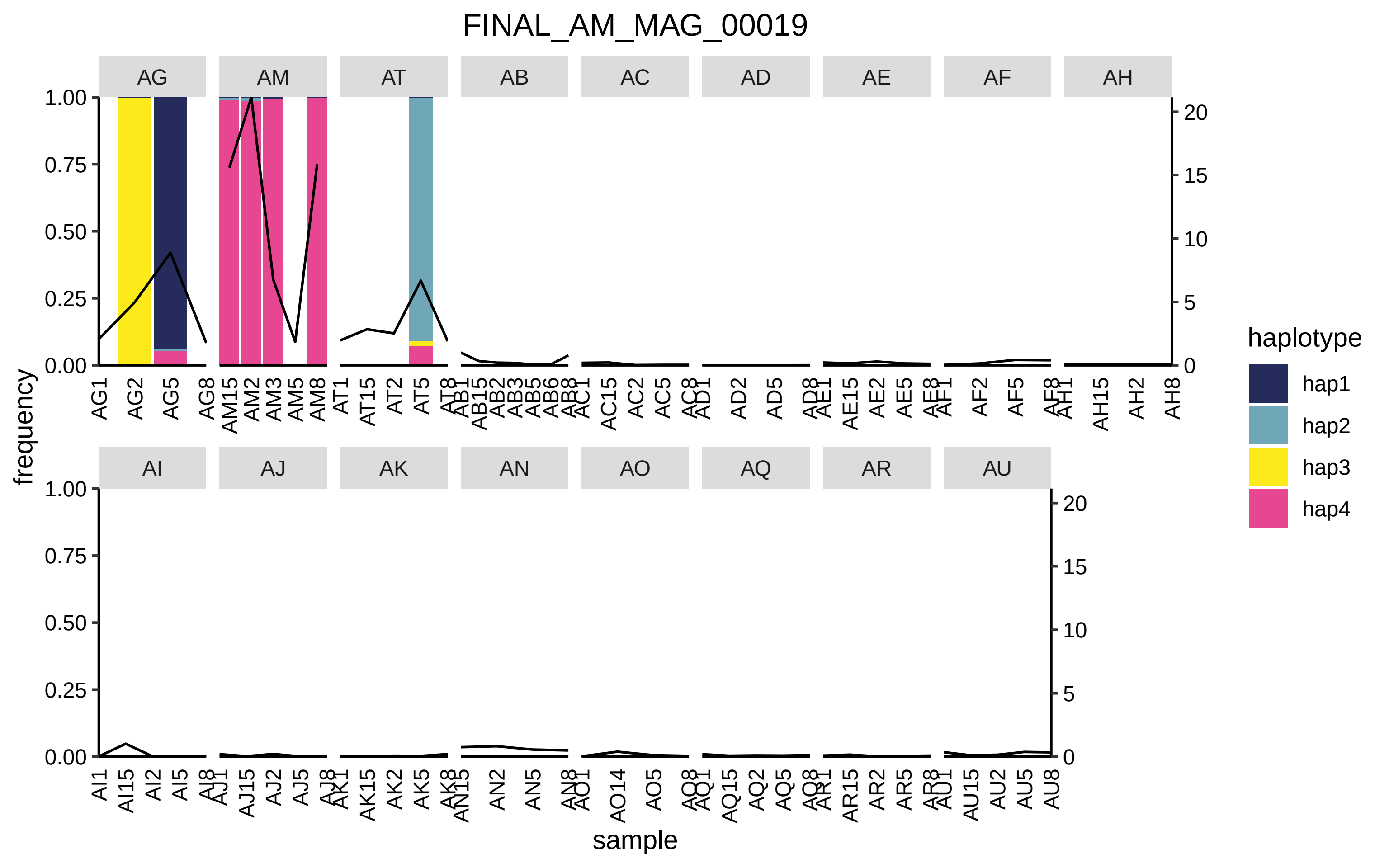

FINAL_AM_MAG_00019
AG
AM
AT
AB
AC
AD
AE
AF
AH
AI
AJ
AK
AN
AO
AQ
AR
AU
frequency
sample
haplotype
hap1
hap2
hap3
hap4
1.00
0.75
0.50
0.25
0.00
20
15
10
5
0
AG1 AG2 AG5 AG8
AM15 AM2 AM3 AM5 AM8
AT1 AT15 AT2 AT5 AT8
AB1 AB15 AB2 AB3 AB5 AB6 AB8
AC1 AC15 AC2 AC5 AC8
AD1 AD2 AD5 AD8
AE1 AE15 AE2 AE5 AE8
AF1 AF2 AF5 AF8
AH1 AH15 AH2 AH8
AI1 AI15 AI2 AI5 AI8
AJ1 AJ15 AJ2 AJ5 AJ8
AK1 AK15 AK2 AK5 AK8
AN15 AN2 AN5 AN8
AO1 AO14 AO5 AO8
AQ1 AQ15 AQ2 AQ5 AQ8
AR1 AR15 AR2 AR5 AR8
AU1 AU15 AU2 AU5 AU8

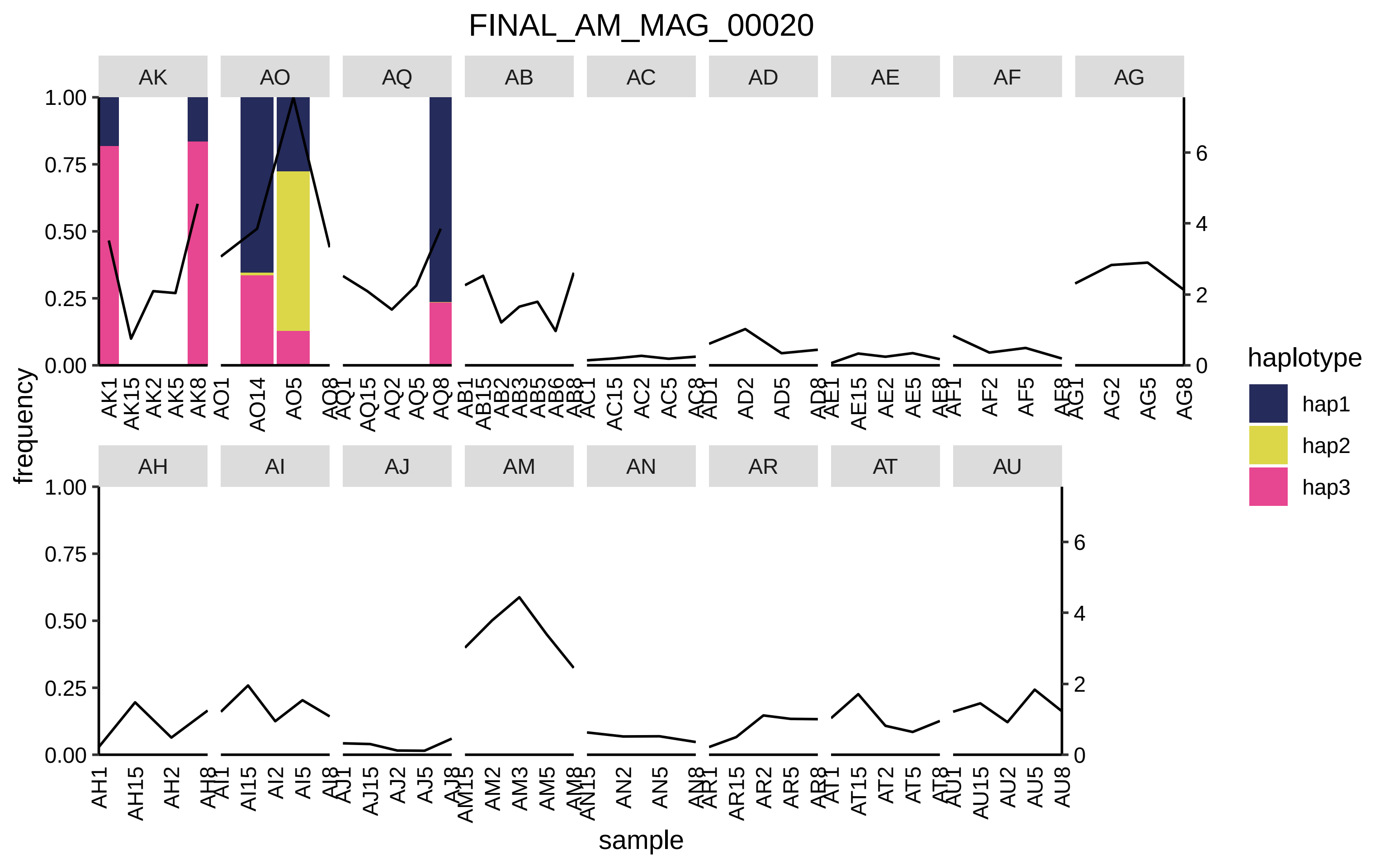
FINAL_AM_MAG_00020
AK
AO
AQ
AB
AC
AD
AE
AF
AG
AH
AI
AJ
AM
AN
AR
AT
AU
frequency
sample
haplotype
hap1
hap2
hap3
1.00
0.75
0.50
0.25
0.00
0
2
4
6

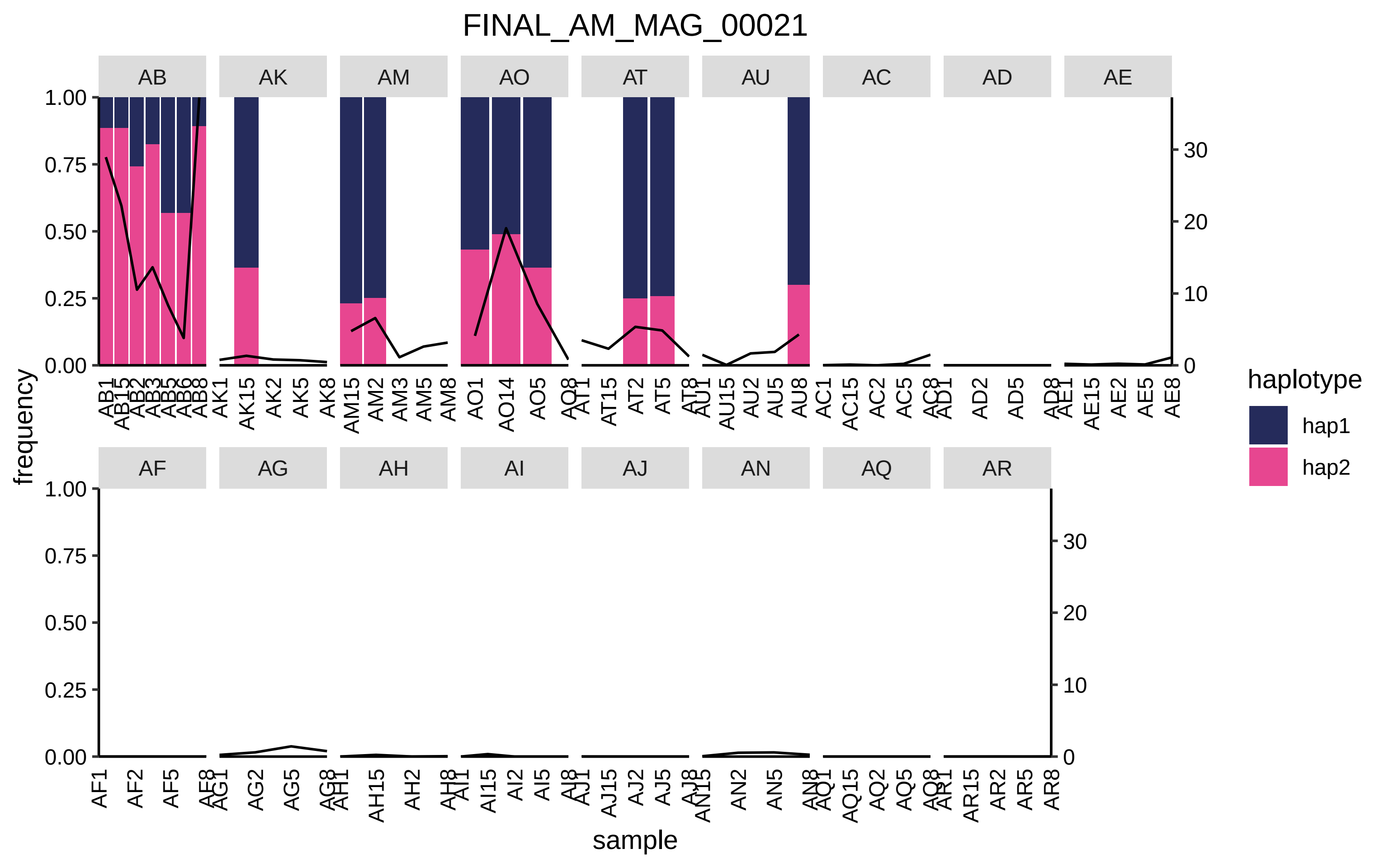
FINAL_AM_MAG_00021
AB
AK
AM
AO
AT
AU
AC
AD
AE
AF
AG
AH
AI
AJ
AN
AQ
AR
frequency
sample
haplotype
hap1
hap2
1.00
0.75
0.50
0.25
0.00
30
20
10
0

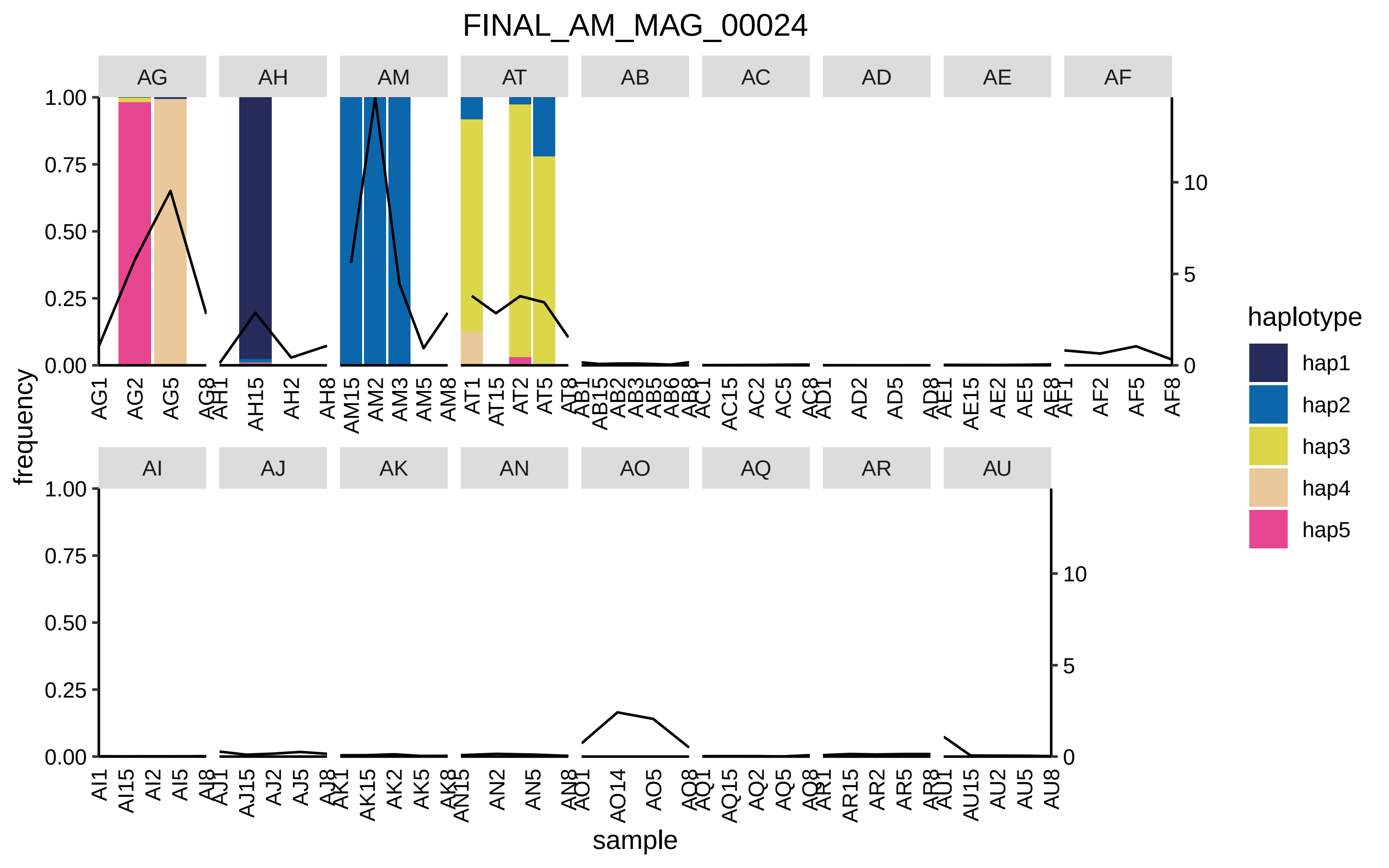

FINAL_AM_MAG_00024
AG
AH
AM
AT
AB
AC
AD
AE
AF
AI
AJ
AK
AN
AO
AQ
AR
AU
frequency
sample
haplotype
hap1
hap2
hap3
hap4
hap5

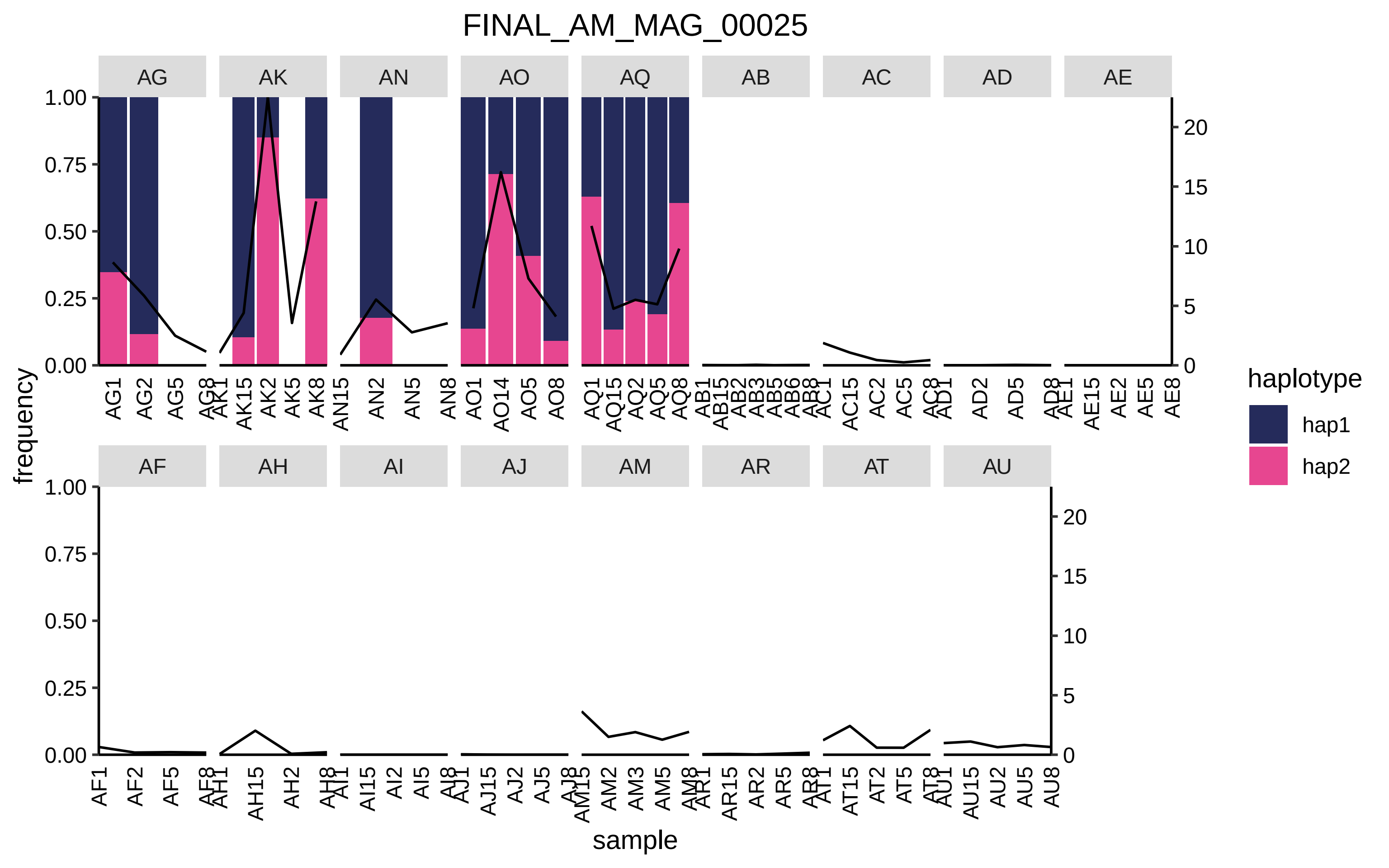

FINAL_AM_MAG_00025
AG
AK
AN
AO
AQ
AB
AC
AD
AE
AF
AH
AI
AJ
AM
AR
AT
AU
frequency
sample
haplotype
hap1
hap2

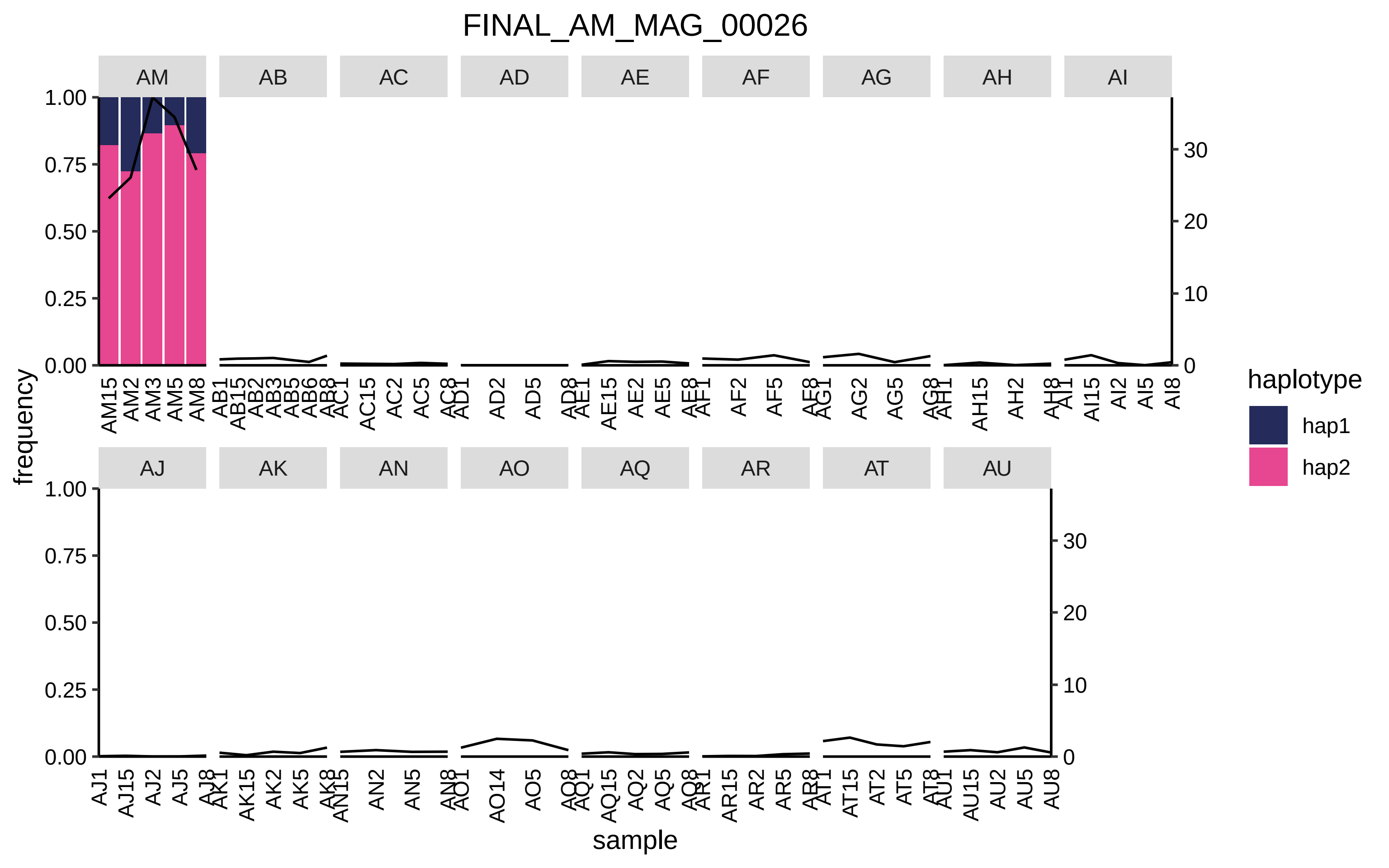

FINAL_AM_MAG_00026
AM
AB
AC
AD
AE
AF
AG
AH
AI
AJ
AK
AN
AO
AQ
AR
AT
AU
frequency
sample
haplotype
hap1
hap2

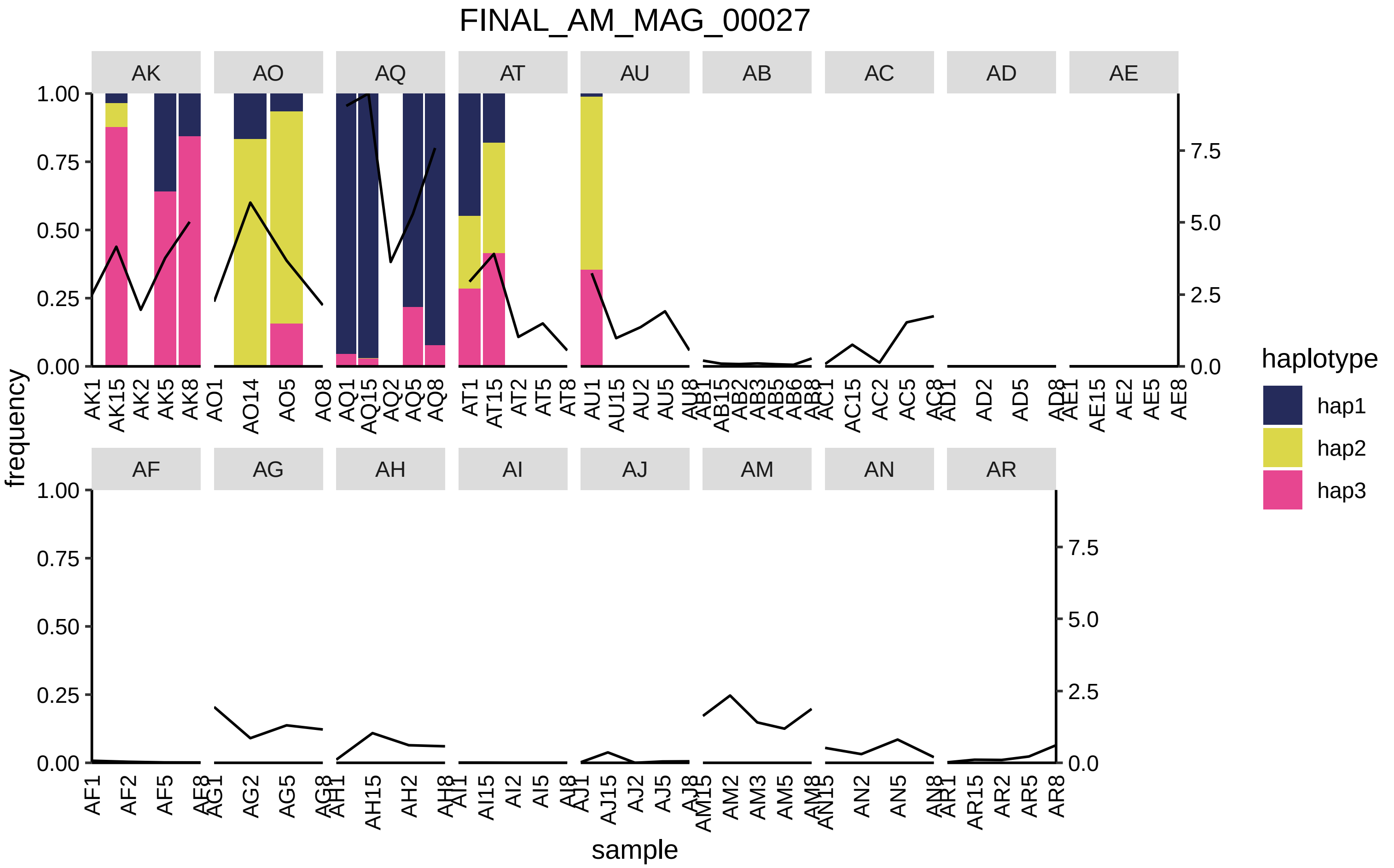
FINAL_AM_MAG_00027
AK
AO
AQ
AT
AU
AB
AC
AD
AE
AF
AG
AH
AI
AJ
AM
AN
AR
frequency
sample
haplotype
hap1
hap2
hap3
1.00
0.75
0.50
0.25
0.00
7.5
5.0
2.5
0.0
AK1
AK15
AK2
AK5
AK8
AO1
AO14
AO5
AO8
AQ1
AQ15
AQ2
AQ5
AQ8
AT1
AT15
AT2
AT5
AT8
AU1
AU15
AU2
AU5
AU8
AB1
AB15
AB2
AB3
AB5
AB6
AB8
AC1
AC15
AC2
AC5
AC8
AD1
AD2
AD5
AD8
AE1
AE15
AE2
AE5
AE8
AF1
AF2
AF5
AF8
AG1
AG2
AG5
AG8
AH1
AH15
AH2
AH8
AI1
AI15
AI2
AI5
AI8
AJ1
AJ15
AJ2
AJ5
AJ8
AM15
AM2
AM3
AM5
AM8
AN15
AN2
AN5
AN8
AR1
AR15
AR2
AR5
AR8

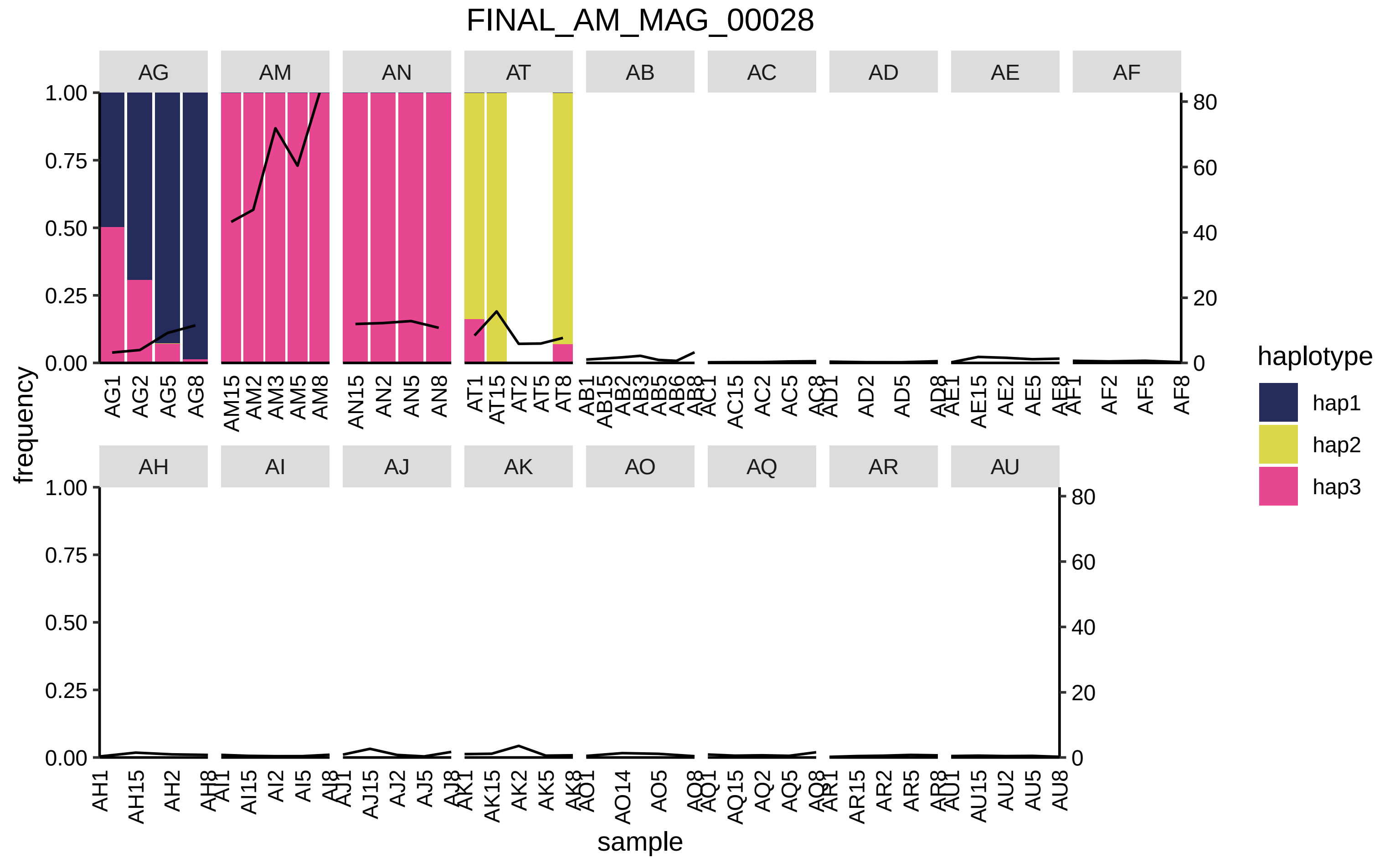

FINAL_AM_MAG_00028
AG
AM
AN
AT
AB
AC
AD
AE
AF
AH
AI
AJ
AK
AO
AQ
AR
AU
frequency
sample
haplotype
hap1
hap2
hap3

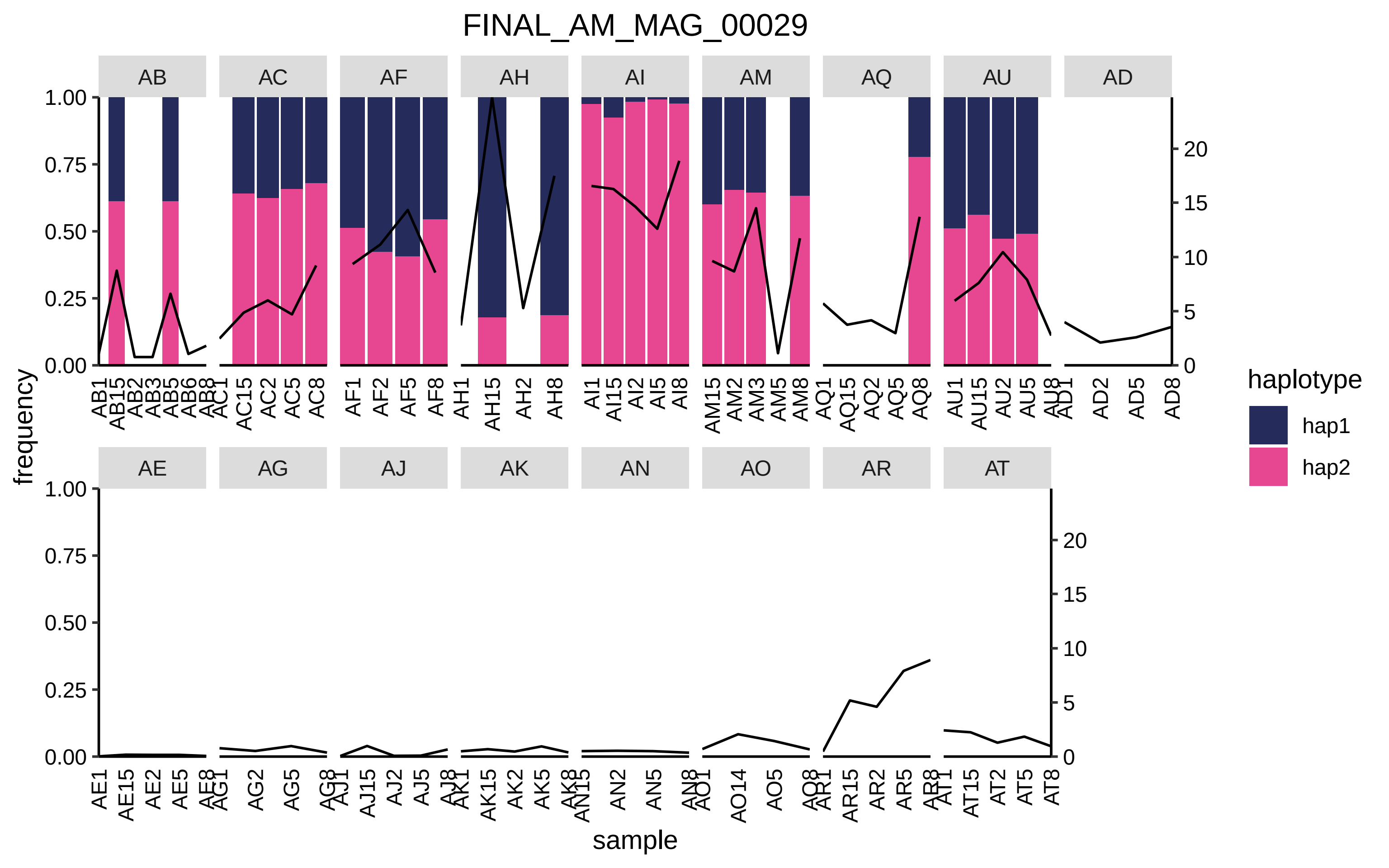

FINAL_AM_MAG_00029
AB
AC
AF
AH
AI
AM
AQ
AU
AD
AE
AG
AJ
AK
AN
AO
AR
AT
frequency
sample
haplotype
hap1
hap2
1.00
0.75
0.50
0.25
0.00
20
15
10
5
0

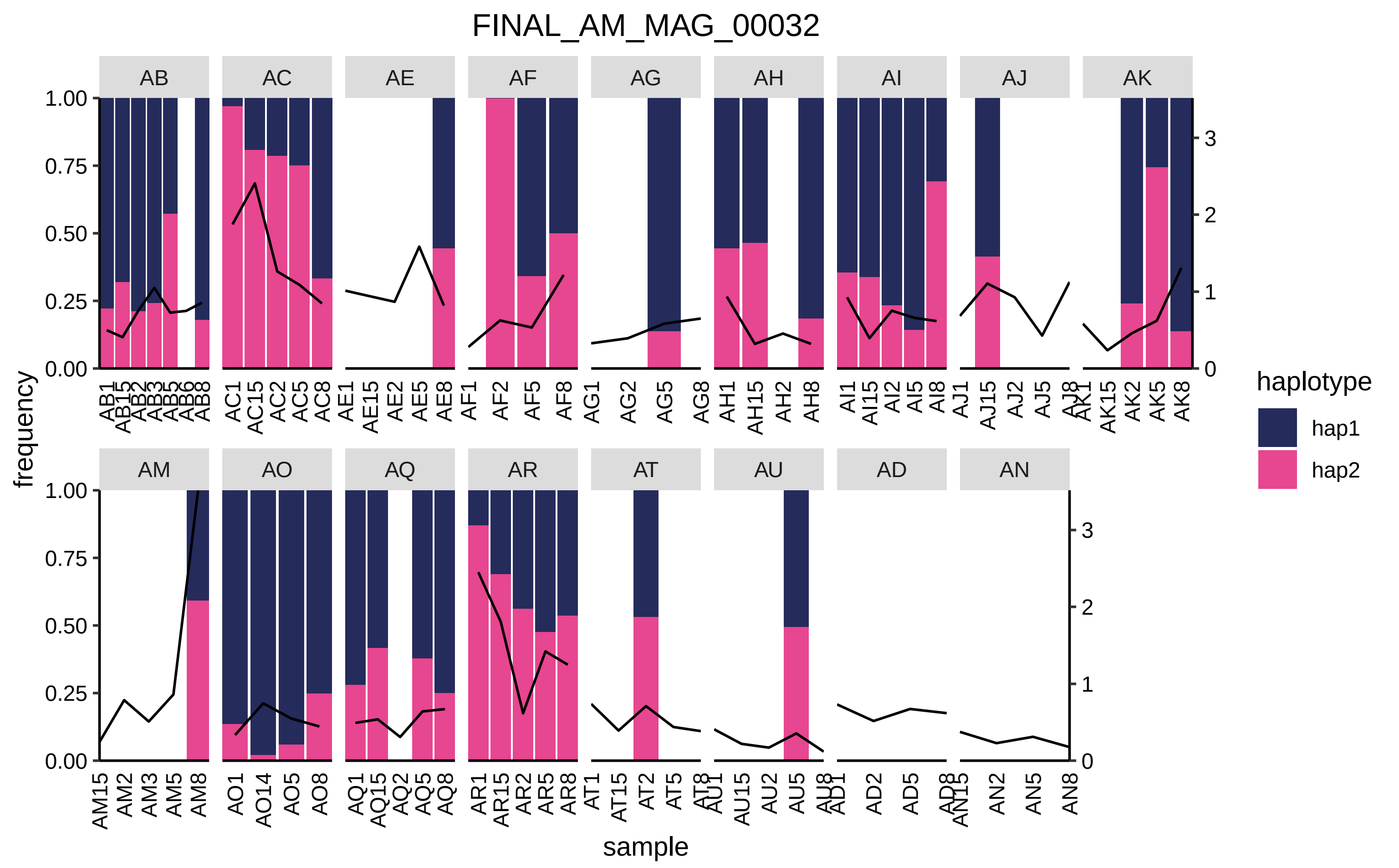

FINAL_AM_MAG_00032
AB
AC
AE
AF
AG
AH
AI
AJ
AK
AM
AO
AQ
AR
AT
AU
AD
AN
frequency
sample
haplotype
hap1
hap2
1.00
0.75
0.50
0.25
0.00
0
1
2
3
AB1
AB15
AB2
AB3
AB5
AB6
AB8
AC1
AC15
AC2
AC5
AC8
AE1
AE15
AE2
AE5
AE8
AF1
AF2
AF5
AF8
AG1
AG2
AG5
AG8
AH1
AH15
AH2
AH8
AI1
AI15
AI2
AI5
AI8
AJ1
AJ15
AJ2
AJ5
AJ8
AK1
AK15
AK2
AK5
AK8
AM15
AM2
AM3
AM5
AM8
AO1
AO14
AO5
AO8
AQ1
AQ15
AQ2
AQ5
AQ8
AR1
AR15
AR2
AR5
AR8
AT1
AT15
AT2
AT5
AT8
AU1
AU15
AU2
AU5
AU8
AD1
AD2
AD5
AD8
AN15
AN2
AN5
AN8

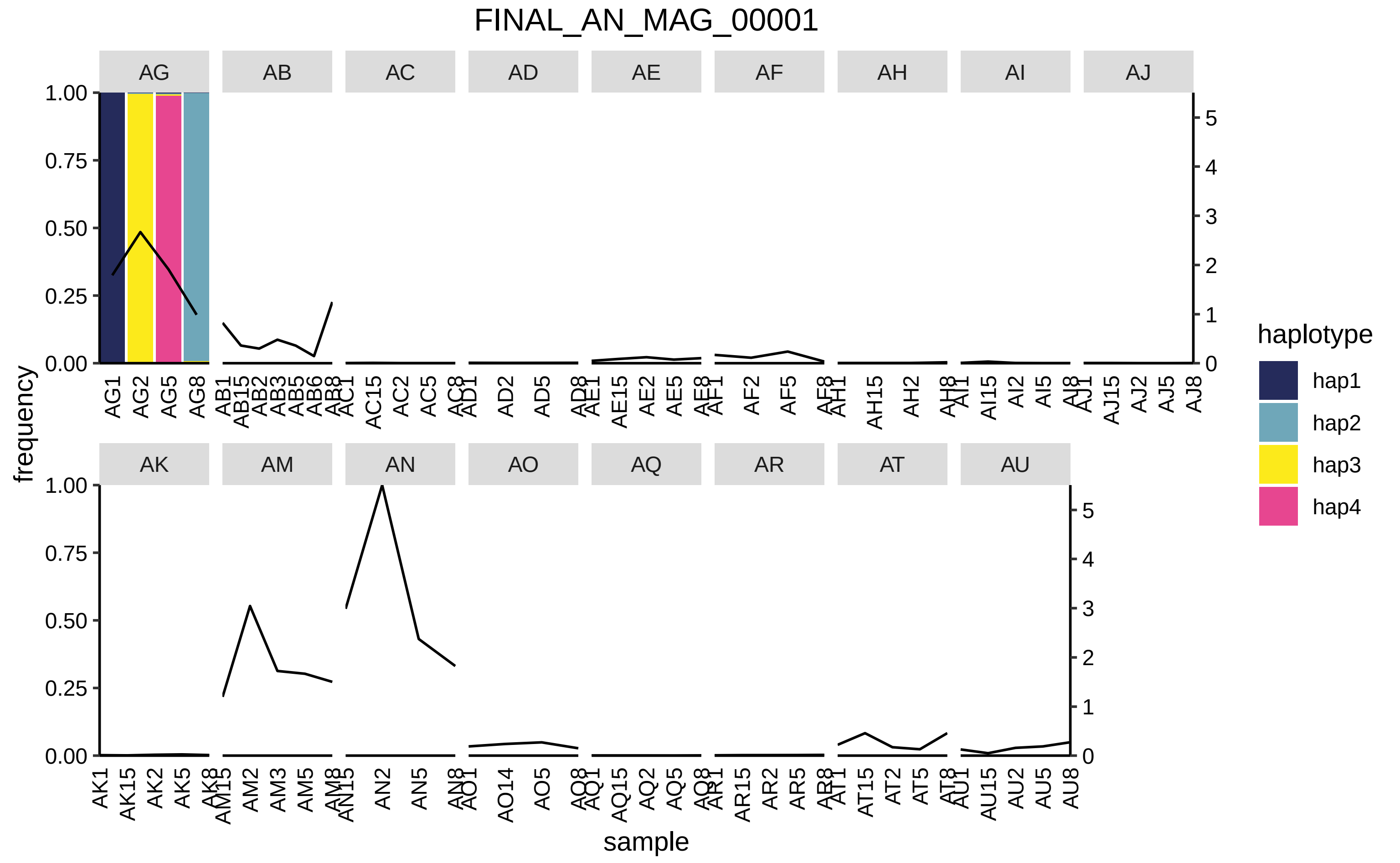

FINAL_AN_MAG_00001
AG
AB
AC
AD
AE
AF
AH
AI
AJ
AK
AM
AN
AO
AQ
AR
AT
AU
frequency
sample
haplotype
hap1
hap2
hap3
hap4

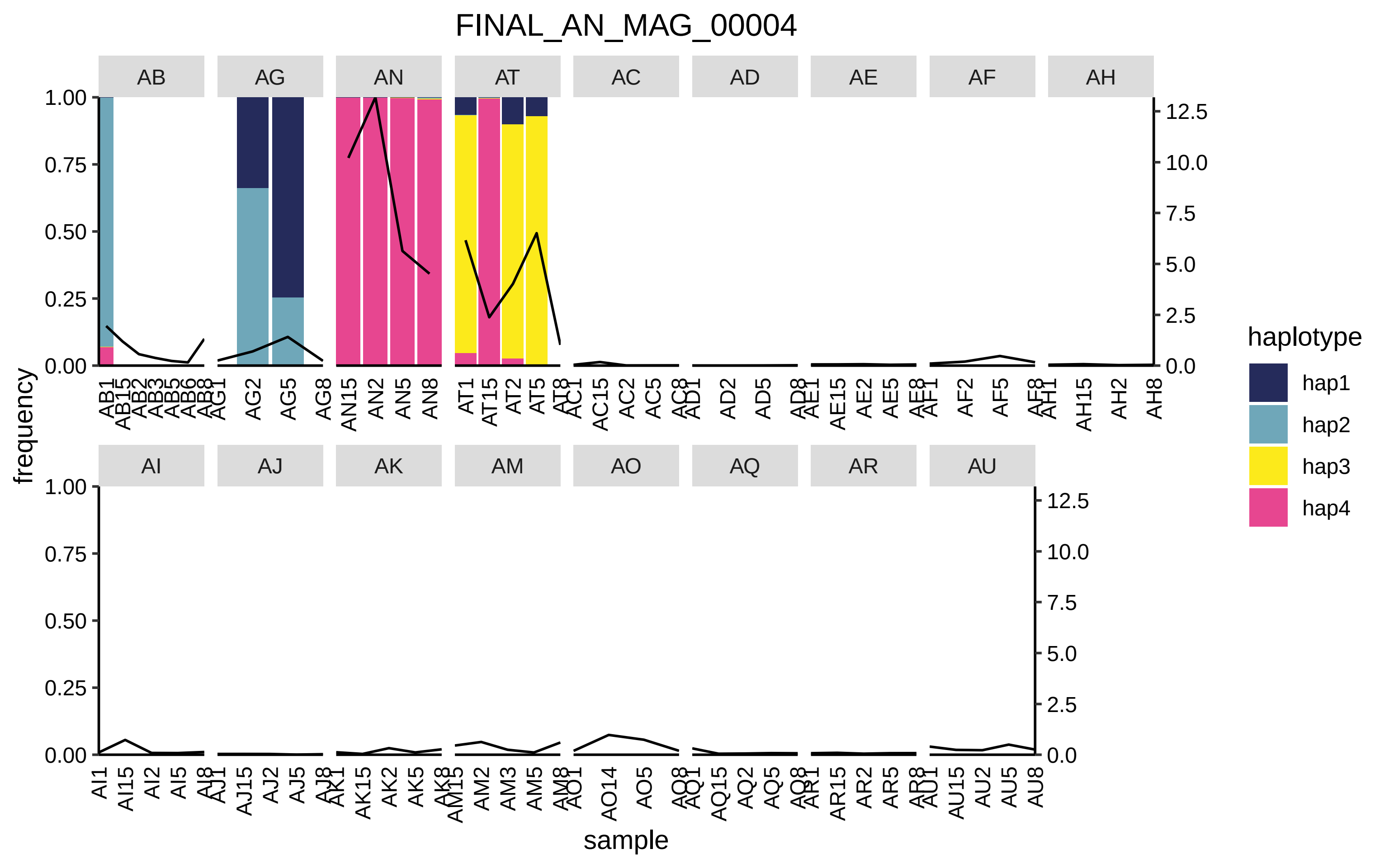

FINAL_AN_MAG_00004
AB
AG
AN
AT
AC
AD
AE
AF
AH
AI
AJ
AK
AM
AO
AQ
AR
AU
frequency
sample
1.00
0.75
0.50
0.25
0.00
12.5
10.0
7.5
5.0
2.5
0.0
haplotype
hap1
hap2
hap3
hap4

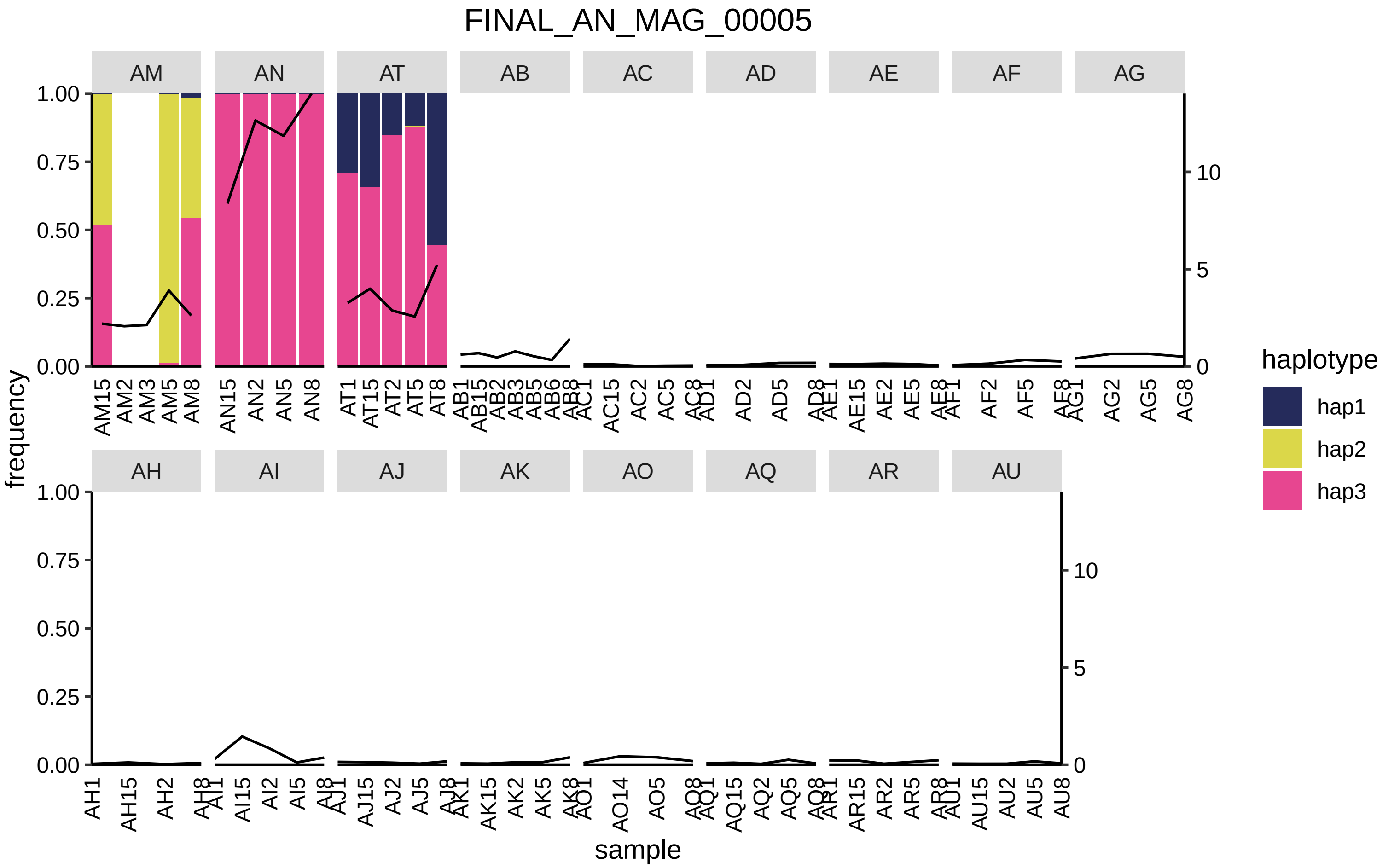

FINAL_AN_MAG_00005
AM
AN
AT
AB
AC
AD
AE
AF
AG
AH
AI
AJ
AK
AO
AQ
AR
AU
frequency
sample
haplotype
hap1
hap2
hap3

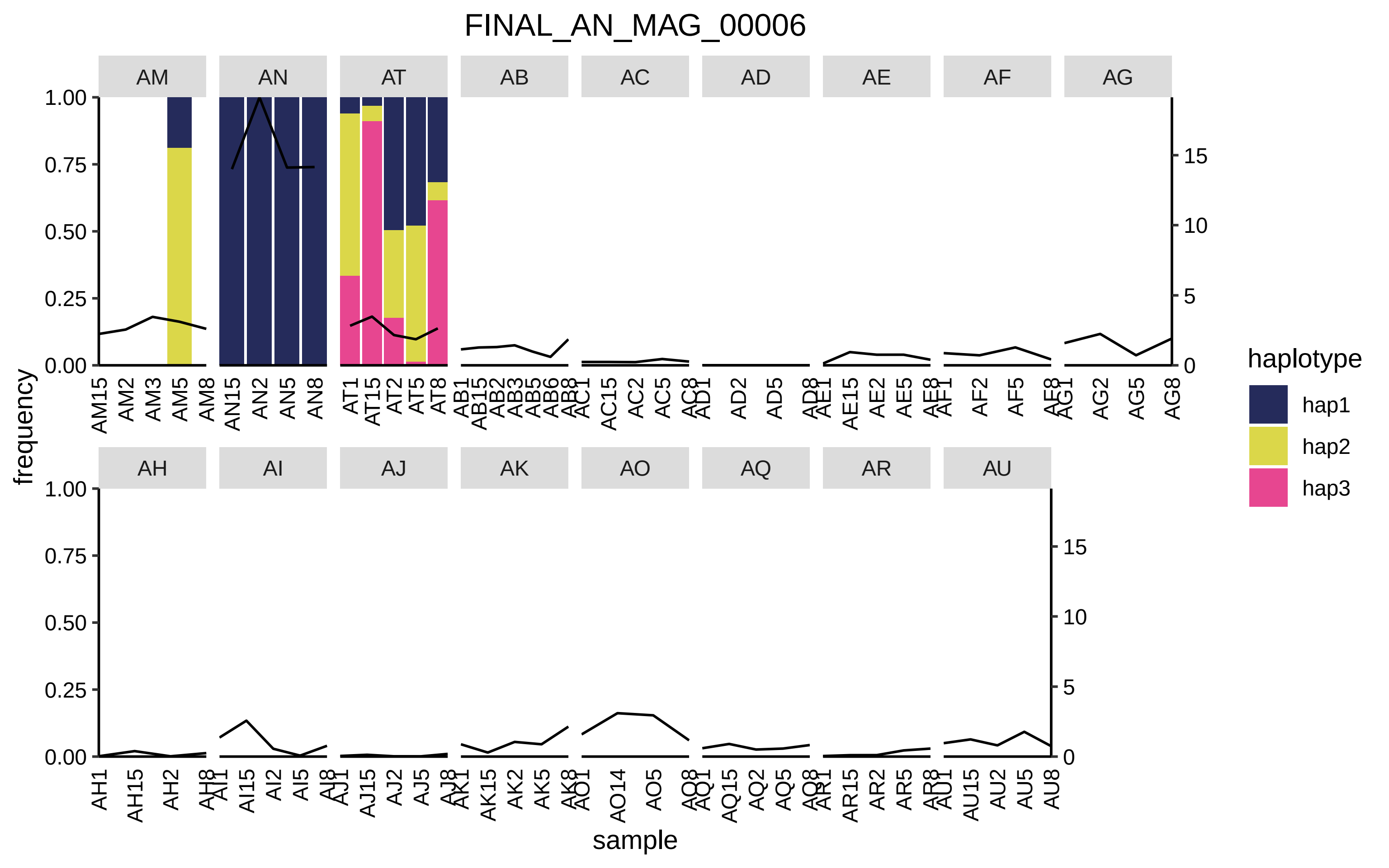
FINAL_AN_MAG_00006
AM
AN
AT
AB
AC
AD
AE
AF
AG
AH
AI
AJ
AK
AO
AQ
AR
AU
frequency
sample
haplotype
hap1
hap2
hap3

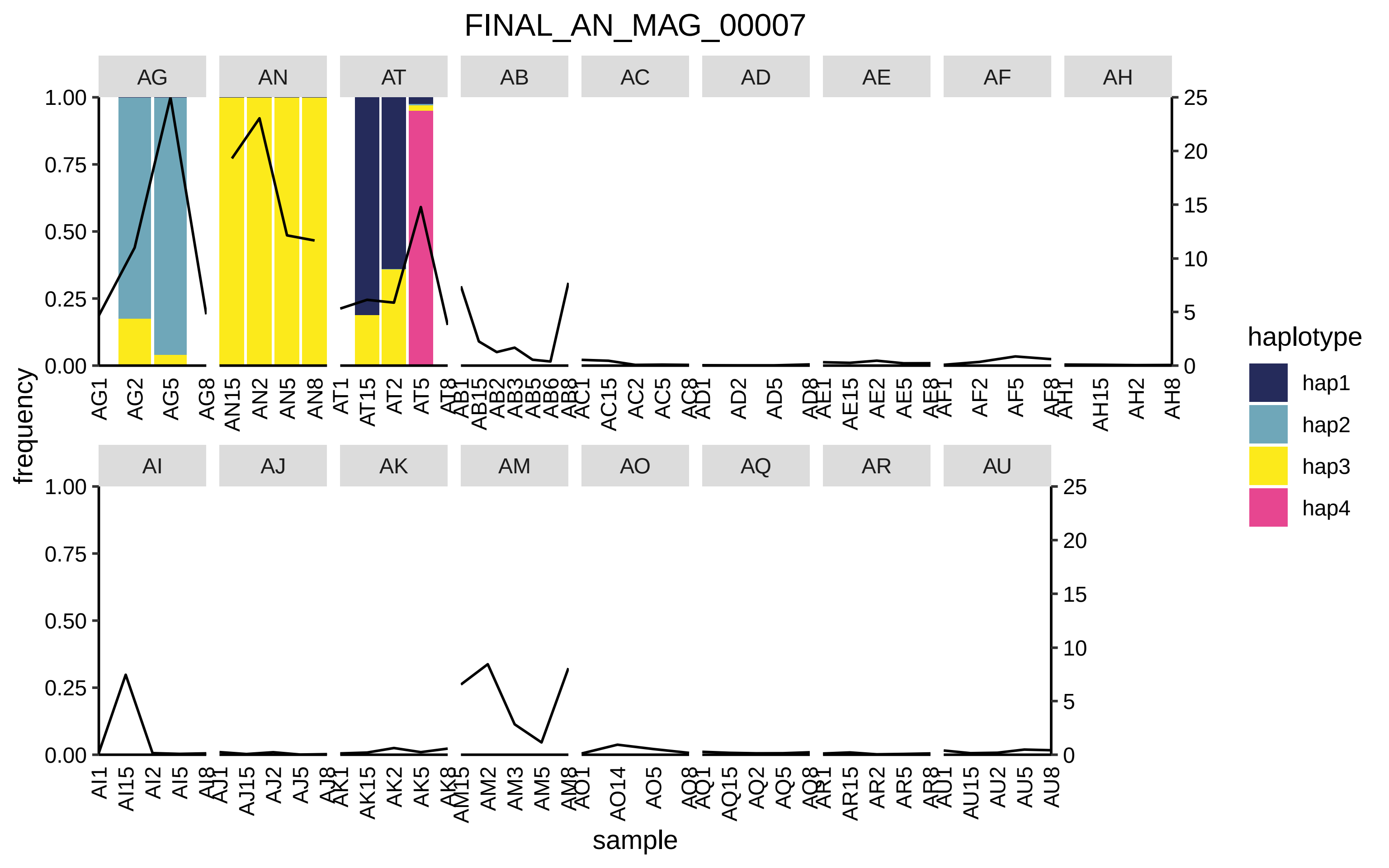
FINAL_AN_MAG_00007
AG
AN
AT
AB
AC
AD
AE
AF
AH
AI
AJ
AK
AM
AO
AQ
AR
AU
frequency
sample
haplotype
hap1
hap2
hap3
hap4

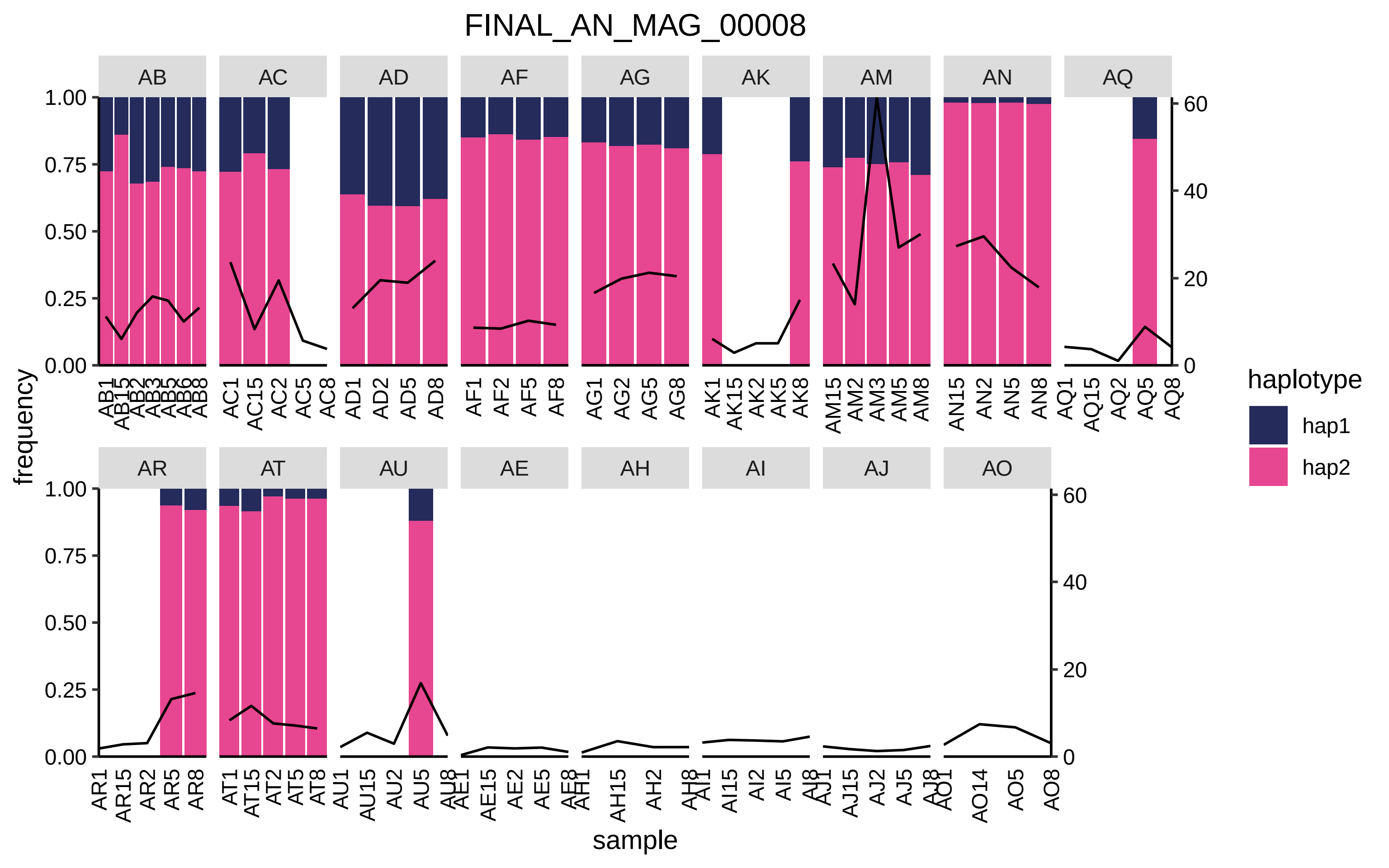

FINAL_AN_MAG_00008
AB
AC
AD
AF
AG
AK
AM
AN
AQ
AR
AT
AU
AE
AH
AI
AJ
AO
frequency
sample
1.00
0.75
0.50
0.25
0.00
60
40
20
0
haplotype
hap1
hap2
AB1 AB15 AB2 AB3 AB5 AB6 AB8
AC1 AC15 AC2 AC5 AC8
AD1 AD2 AD5 AD8
AF1 AF2 AF5 AF8
AG1 AG2 AG5 AG8
AK1 AK15 AK2 AK5 AK8
AM15 AM2 AM3 AM5 AM8
AN15 AN2 AN5 AN8
AQ1 AQ15 AQ2 AQ5 AQ8
AR1 AR15 AR2 AR5 AR8
AT1 AT15 AT2 AT5 AT8
AU1 AU15 AU2 AU5 AU8
AE1 AE15 AE2 AE5 AE8
AH1 AH15 AH2 AH8
AI1 AI15 AI2 AI5 AI8
AJ1 AJ15 AJ2 AJ5 AJ8
AO1 AO14 AO5 AO8

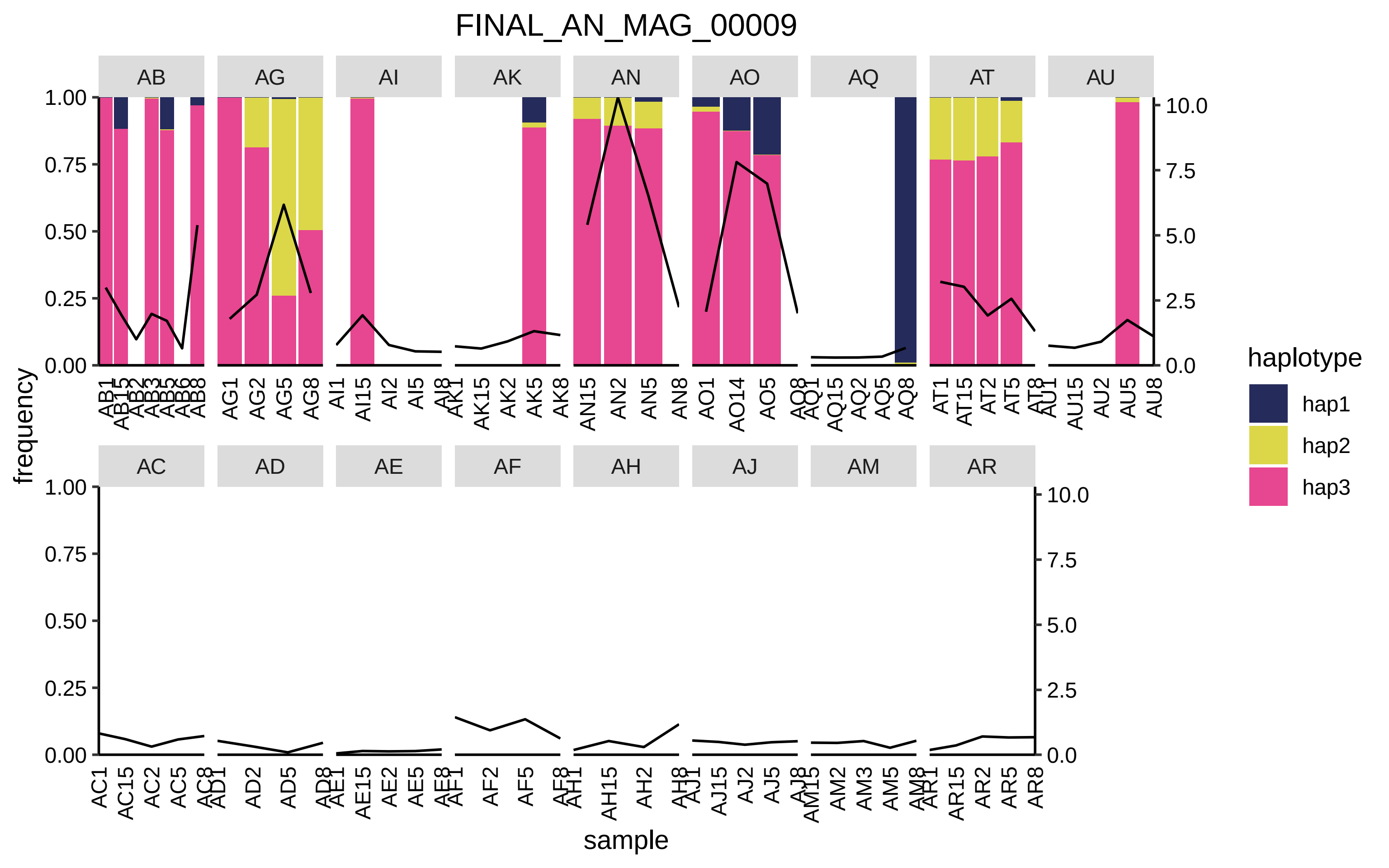

FINAL_AN_MAG_00009
AB
AG
AI
AK
AN
AO
AQ
AT
AU
AC
AD
AE
AF
AH
AJ
AM
AR
frequency
sample
haplotype
hap1
hap2
hap3
1.00
0.75
0.50
0.25
0.00
10.0
7.5
5.0
2.5
0.0

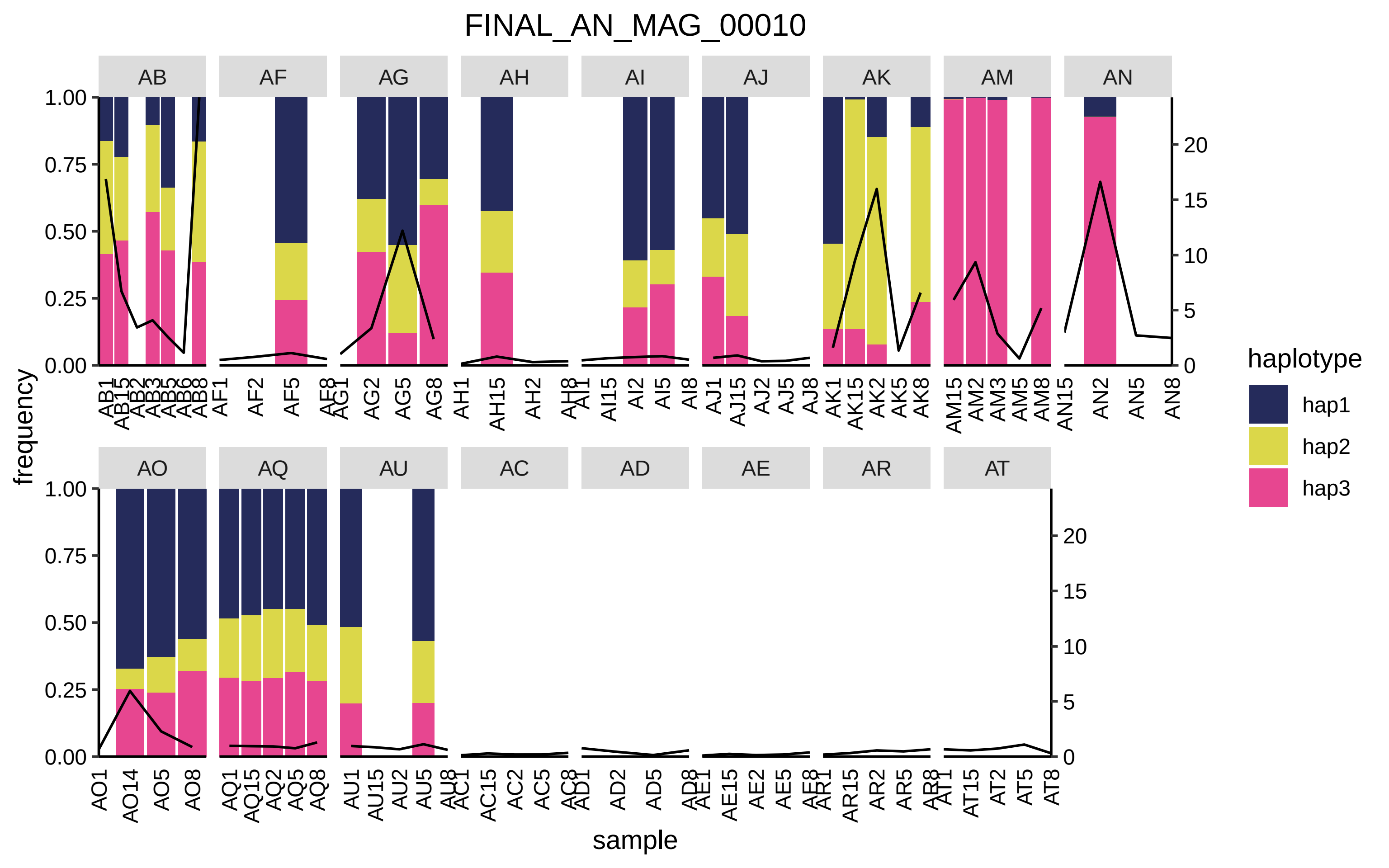
FINAL_AN_MAG_00010
AB
AF
AG
AH
AI
AJ
AK
AM
AN
AO
AQ
AU
AC
AD
AE
AR
AT
frequency
sample
haplotype
hap1
hap2
hap3

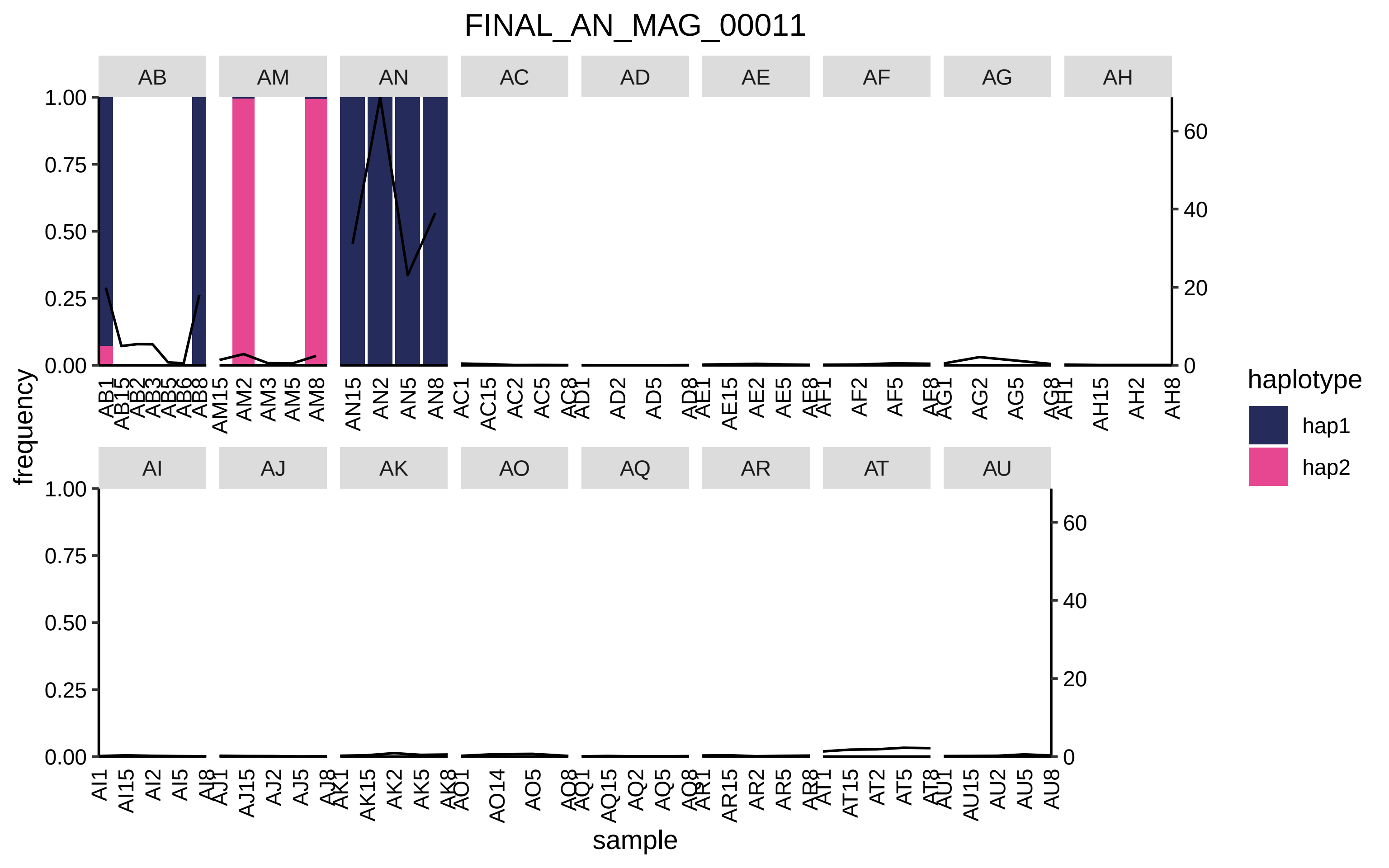
FINAL_AN_MAG_00011
AB
AM
AN
AC
AD
AE
AF
AG
AH
AI
AJ
AK
AO
AQ
AR
AT
AU
frequency
sample
haplotype
hap1
hap2
1.00
0.75
0.50
0.25
0.00
0
20
40
60

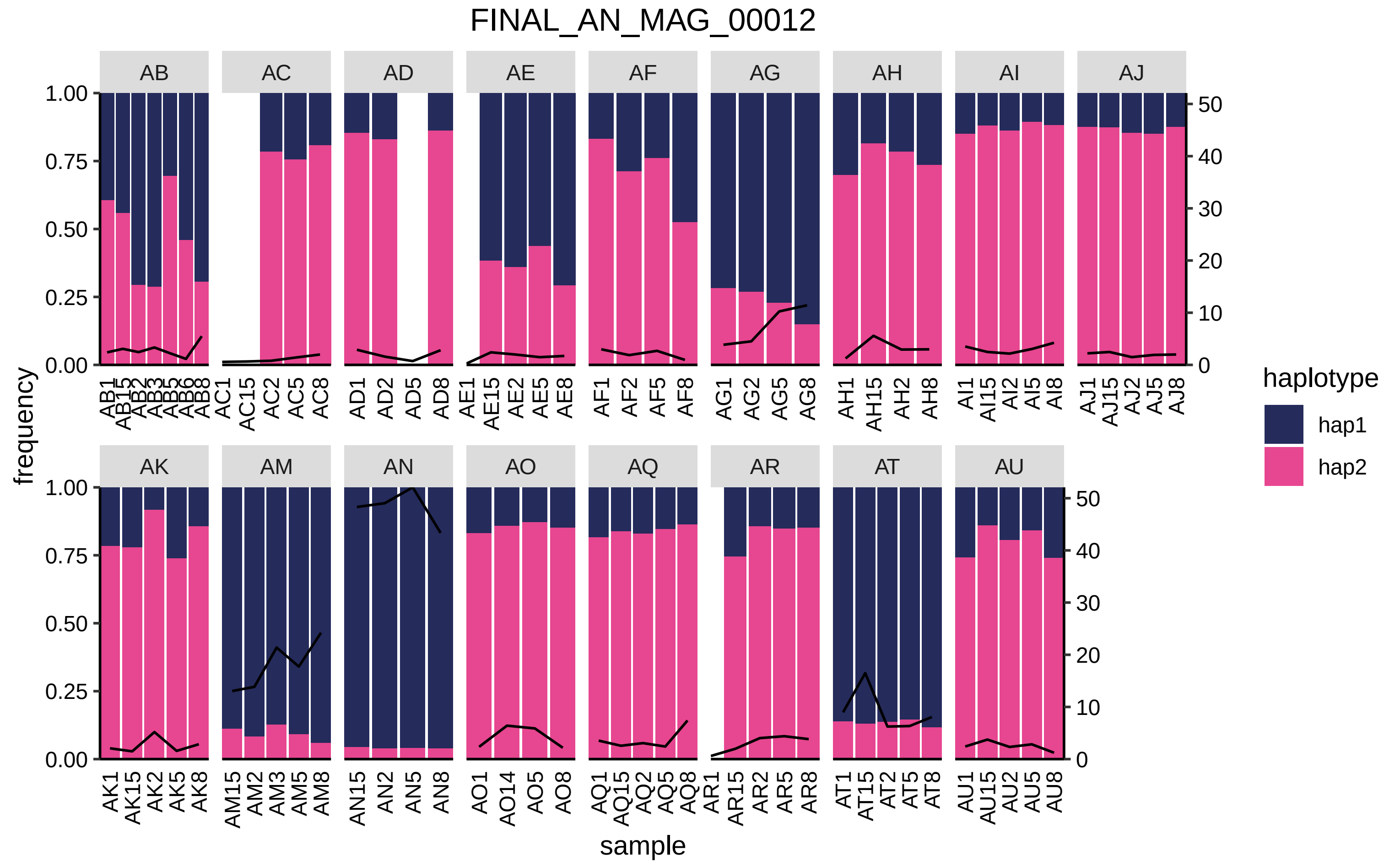

FINAL_AN_MAG_00012
frequency
sample
haplotype
hap1
hap2
AB
AC
AD
AE
AF
AG
AH
AI
AJ
AK
AM
AN
AO
AQ
AR
AT
AU
0.00
0.25
0.50
0.75
1.00
0
10
20
30
40
50
AB1 AB15 AB2 AB3 AB5 AB6 AB8
AC1 AC15 AC2 AC5 AC8
AD1 AD2 AD5 AD8
AE1 AE15 AE2 AE5 AE8
AF1 AF2 AF5 AF8
AG1 AG2 AG5 AG8
AH1 AH15 AH2 AH8
AI1 AI15 AI2 AI5 AI8
AJ1 AJ15 AJ2 AJ5 AJ8
AK1 AK15 AK2 AK5 AK8
AM15 AM2 AM3 AM5 AM8
AN15 AN2 AN5 AN8
AO1 AO14 AO5 AO8
AQ1 AQ15 AQ2 AQ5 AQ8
AR1 AR15 AR2 AR5 AR8
AT1 AT15 AT2 AT5 AT8
AU1 AU15 AU2 AU5 AU8

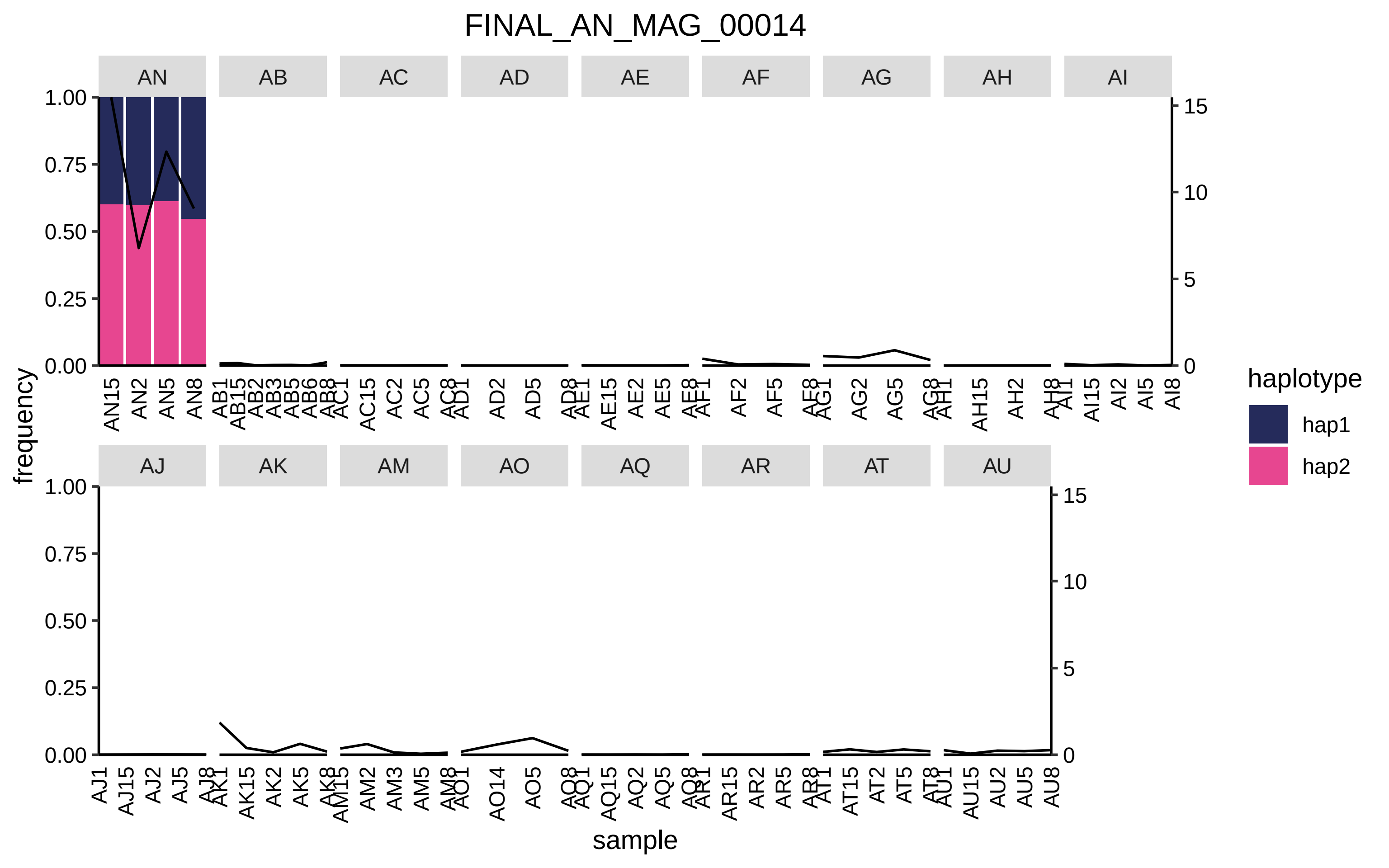

FINAL_AN_MAG_00014
AN
AB
AC
AD
AE
AF
AG
AH
AI
AJ
AK
AM
AO
AQ
AR
AT
AU
frequency
sample
haplotype
hap1
hap2

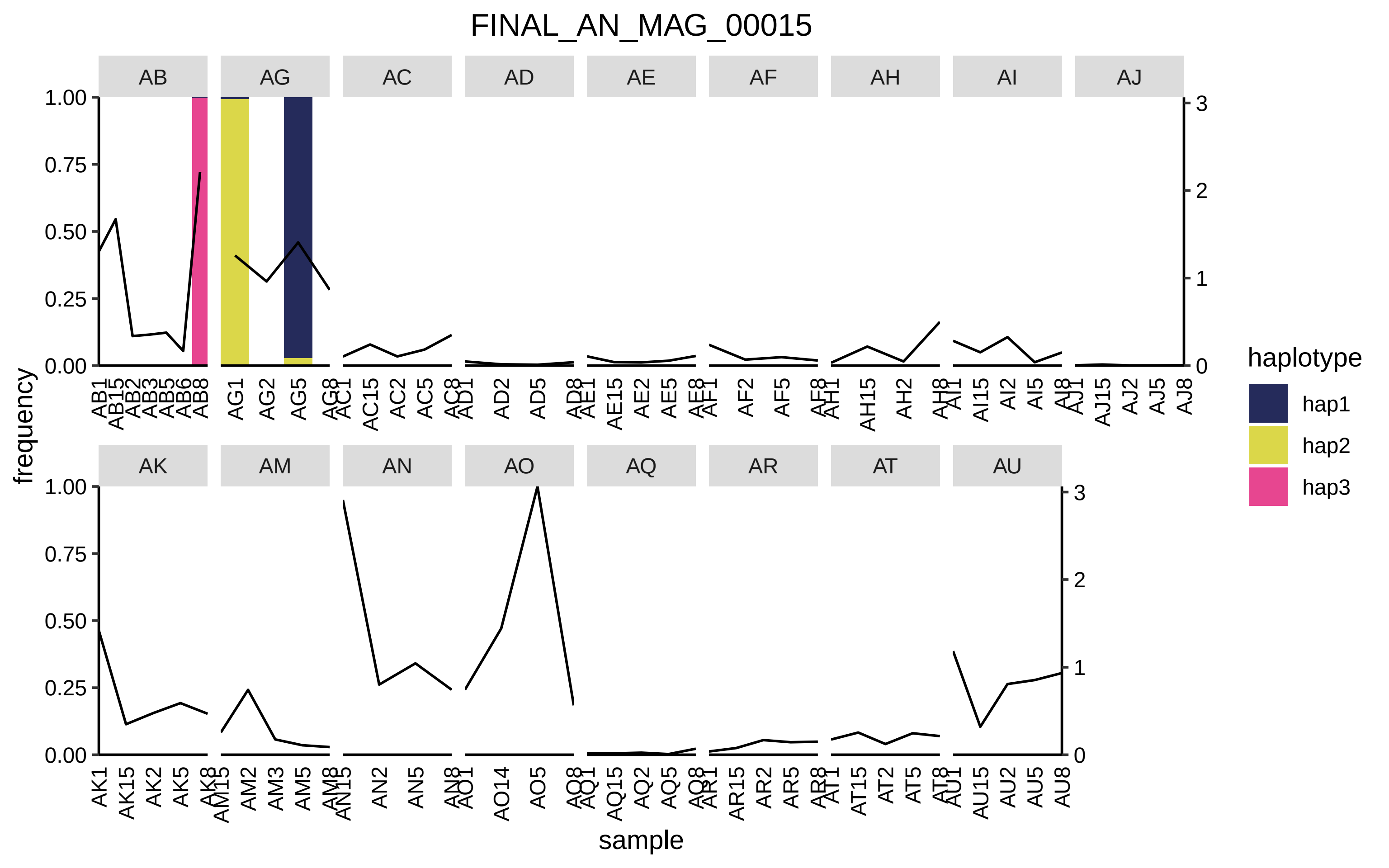
FINAL_AN_MAG_00015
AB
AG
AC
AD
AE
AF
AH
AI
AJ
AK
AM
AN
AO
AQ
AR
AT
AU
frequency
sample
haplotype
hap1
hap2
hap3
1.00
0.75
0.50
0.25
0.00
3
2
1
0

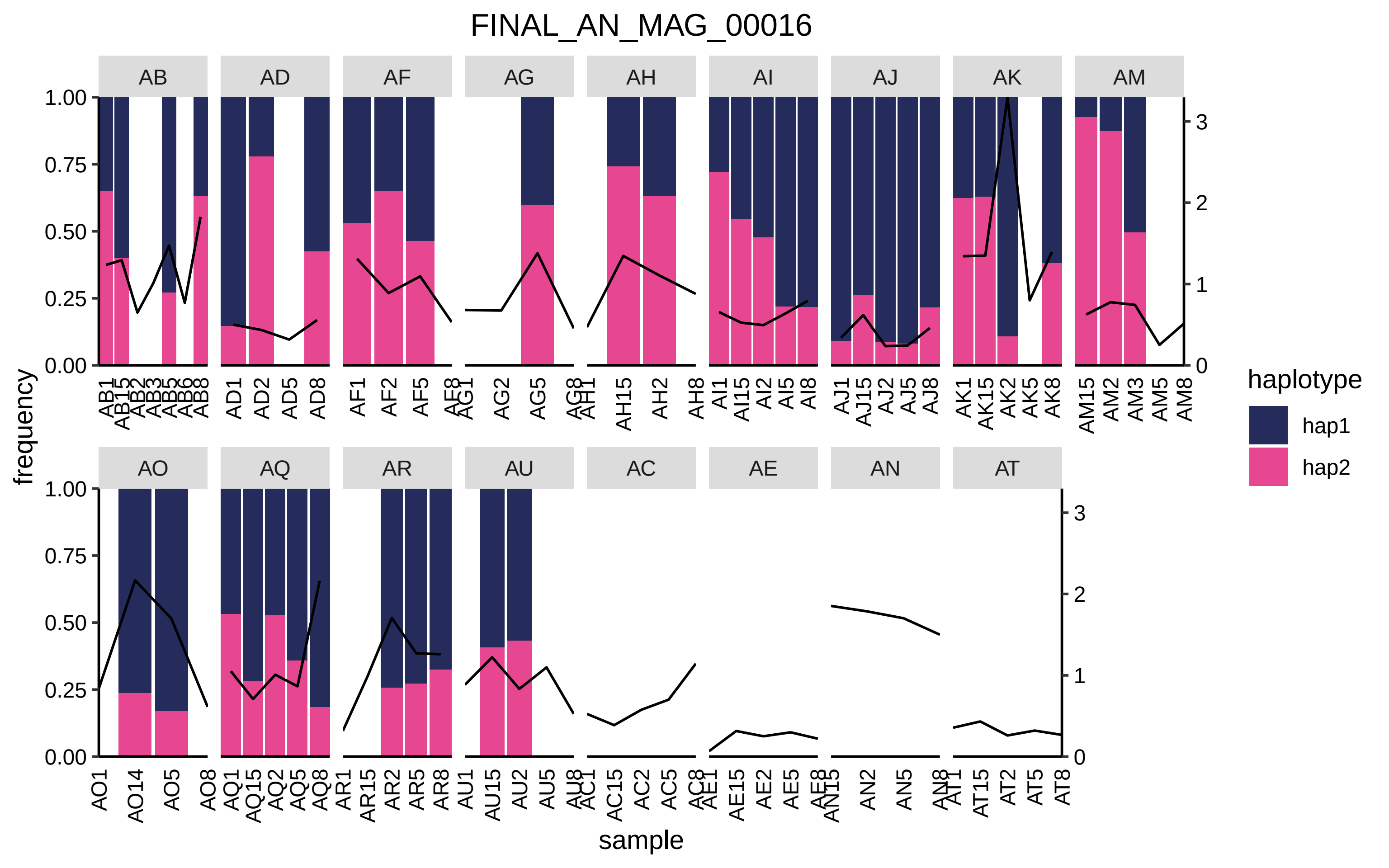

FINAL_AN_MAG_00016
AB
AD
AF
AG
AH
AI
AJ
AK
AM
AO
AQ
AR
AU
AC
AE
AN
AT
1.00
0.75
0.50
0.25
0.00
0
1
2
3
frequency
sample
haplotype
hap1
hap2
AB1 AB15 AB2 AB3 AB5 AB6 AB8
AD1 AD2 AD5 AD8
AF1 AF2 AF5 AF8
AG1 AG2 AG5 AG8
AH1 AH15 AH2 AH8
AI1 AI15 AI2 AI5 AI8
AJ1 AJ15 AJ2 AJ5 AJ8
AK1 AK15 AK2 AK5 AK8
AM15 AM2 AM3 AM5 AM8
AO1 AO14 AO5 AO8
AQ1 AQ15 AQ2 AQ5 AQ8
AR1 AR15 AR2 AR5 AR8
AU1 AU15 AU2 AU5 AU8
AC1 AC15 AC2 AC5 AC8
AE1 AE15 AE2 AE5 AE8
AN15 AN2 AN5 AN8
AT1 AT15 AT2 AT5 AT8

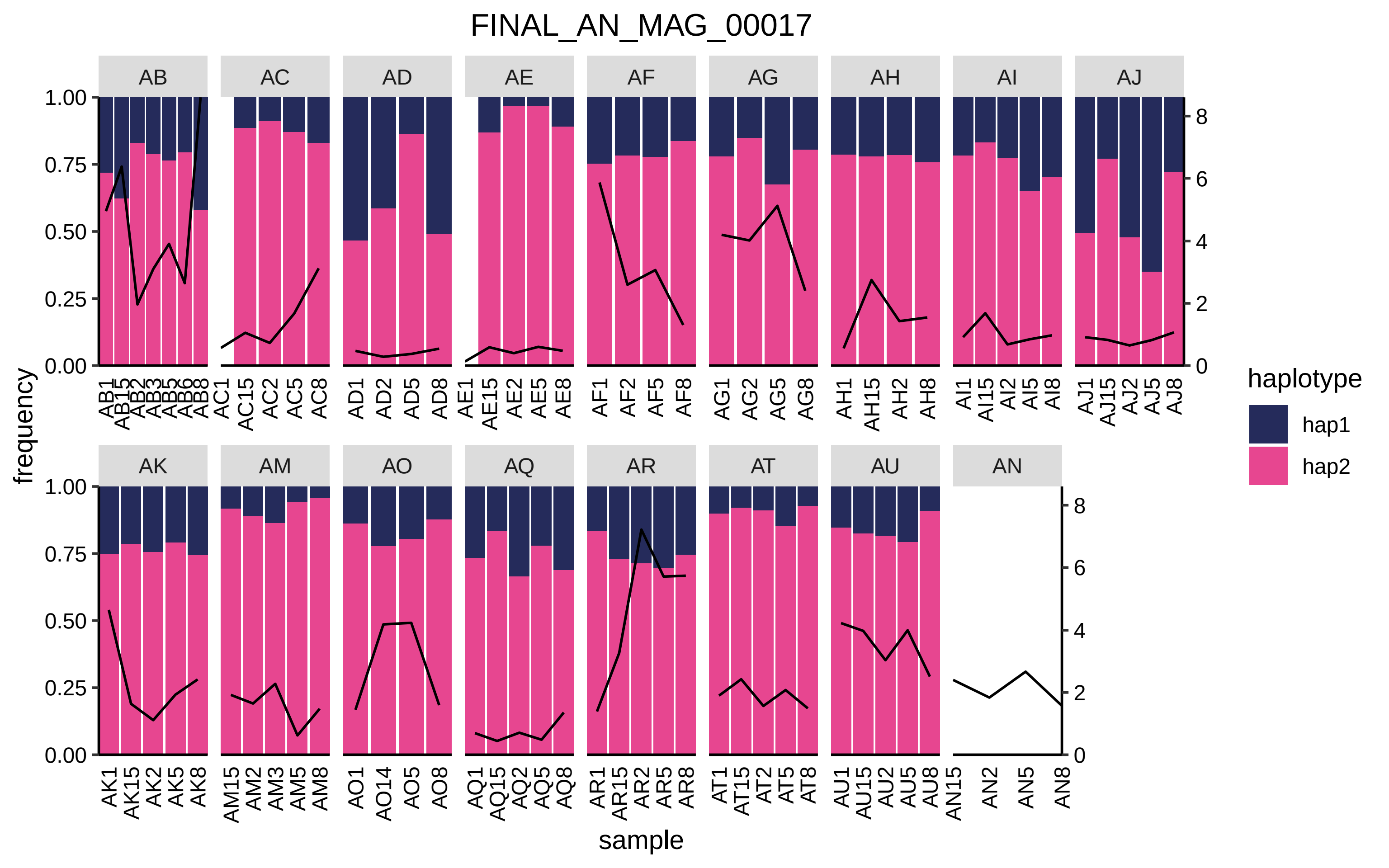
FINAL_AN_MAG_00017
AB
AC
AD
AE
AF
AG
AH
AI
AJ
AK
AM
AO
AQ
AR
AT
AU
AN
frequency
1.00
0.75
0.50
0.25
0.00
8
6
4
2
0
AB1 AB15 AB2 AB3 AB5 AB6 AB8
AC1 AC15 AC2 AC5 AC8
AD1 AD2 AD5 AD8
AE1 AE15 AE2 AE5 AE8
AF1 AF2 AF5 AF8
AG1 AG2 AG5 AG8
AH1 AH15 AH2 AH8
AI1 AI15 AI2 AI5 AI8
AJ1 AJ15 AJ2 AJ5 AJ8
AK1 AK15 AK2 AK5 AK8
AM15 AM2 AM3 AM5 AM8
AO1 AO14 AO5 AO8
AQ1 AQ15 AQ2 AQ5 AQ8
AR1 AR15 AR2 AR5 AR8
AT1 AT15 AT2 AT5 AT8
AU1 AU15 AU2 AU5 AU8
AN15 AN2 AN5 AN8
sample
haplotype
hap1
hap2

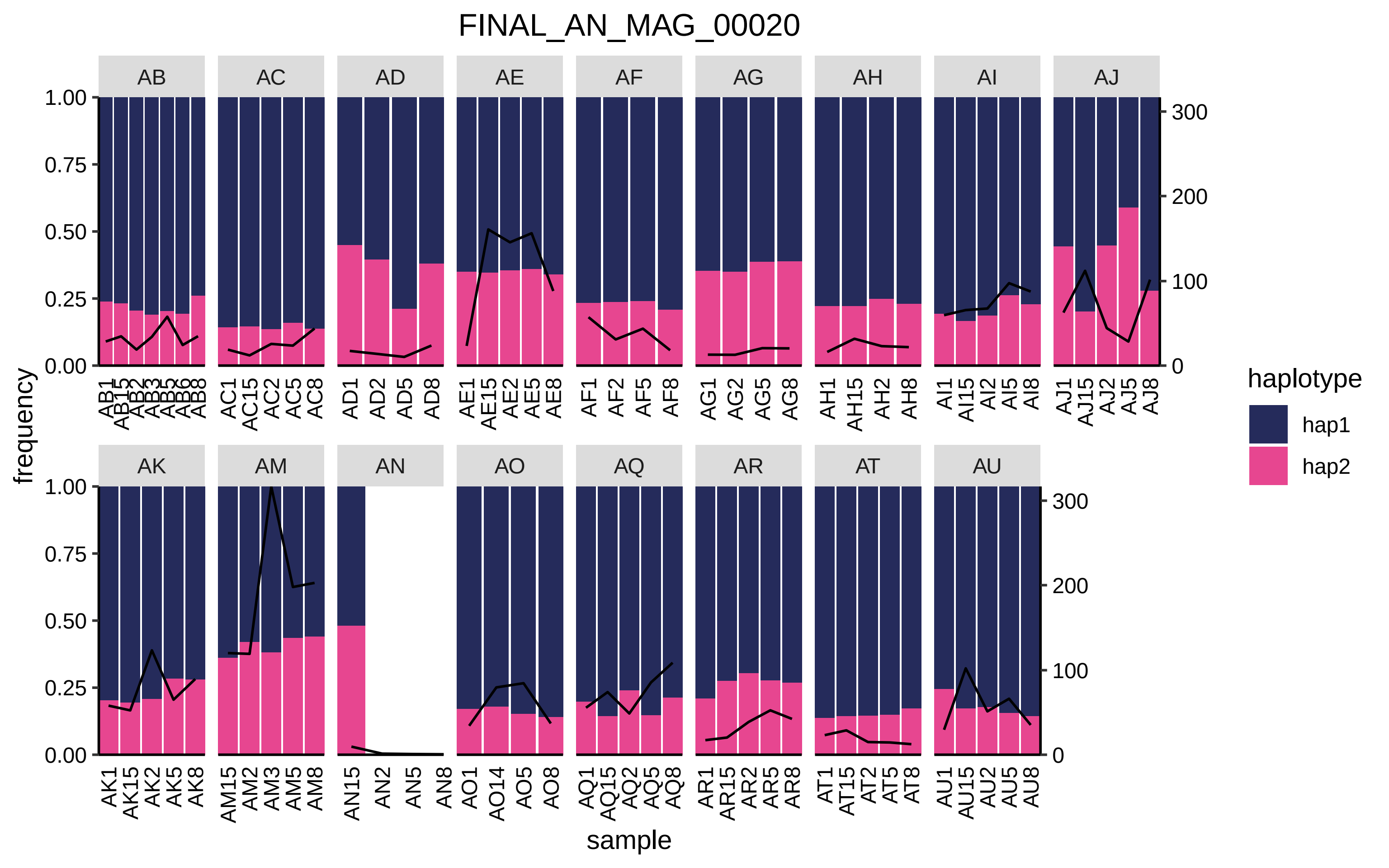
FINAL_AN_MAG_00020
AB
AC
AD
AE
AF
AG
AH
AI
AJ
AK
AM
AN
AO
AQ
AR
AT
AU
frequency
sample
haplotype
hap1
hap2
1.00
0.75
0.50
0.25
0.00
300
200
100
0
AB1
AB15
AB2
AB3
AB5
AB6
AB8
AC1
AC15
AC2
AC5
AC8
AD1
AD2
AD5
AD8
AE1
AE15
AE2
AE5
AE8
AF1
AF2
AF5
AF8
AG1
AG2
AG5
AG8
AH1
AH15
AH2
AH8
AI1
AI15
AI2
AI5
AI8
AJ1
AJ15
AJ2
AJ5
AJ8
AK1
AK15
AK2
AK5
AK8
AM15
AM2
AM3
AM5
AM8
AN15
AN2
AN5
AN8
AO1
AO14
AO5
AO8
AQ1
AQ15
AQ2
AQ5
AQ8
AR1
AR15
AR2
AR5
AR8
AT1
AT15
AT2
AT5
AT8
AU1
AU15
AU2
AU5
AU8

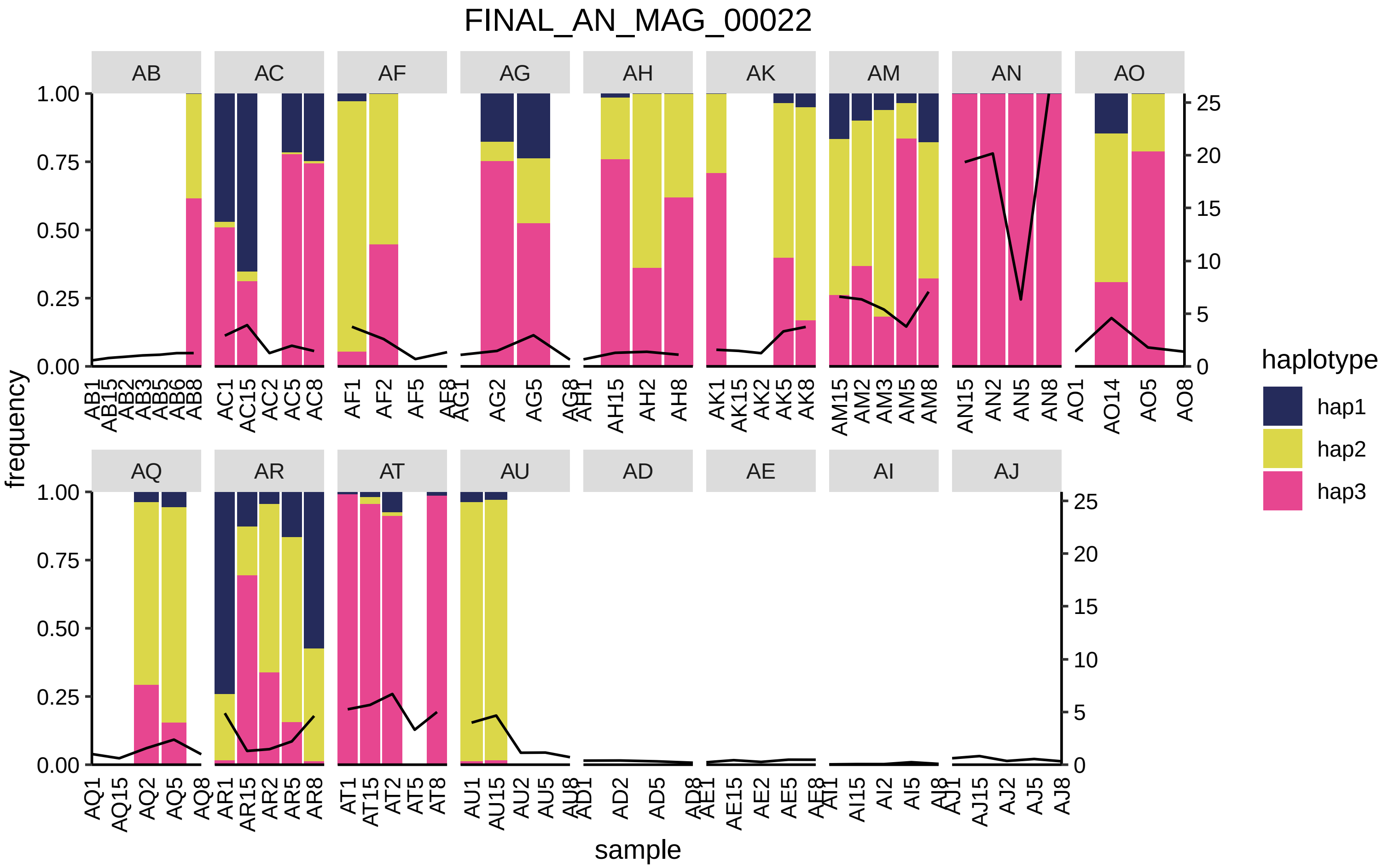

FINAL_AN_MAG_00022
AB
AC
AF
AG
AH
AK
AM
AN
AO
AQ
AR
AT
AU
AD
AE
AI
AJ
frequency
sample
haplotype
hap1
hap2
hap3

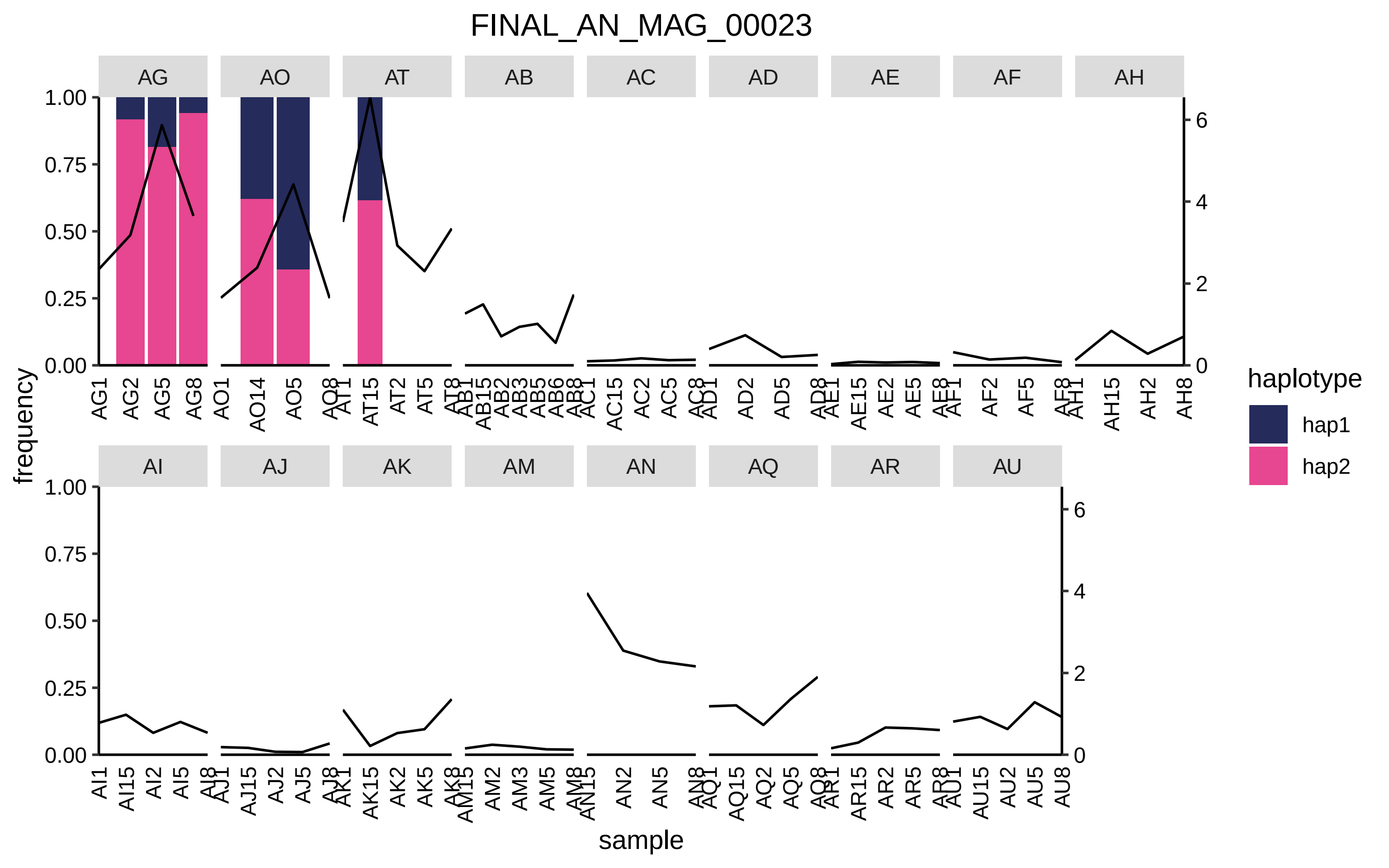

FINAL_AN_MAG_00023
AG
AO
AT
AB
AC
AD
AE
AF
AH
AI
AJ
AK
AM
AN
AQ
AR
AU
frequency
sample
haplotype
hap1
hap2
1.00
0.75
0.50
0.25
0.00
0
2
4
6

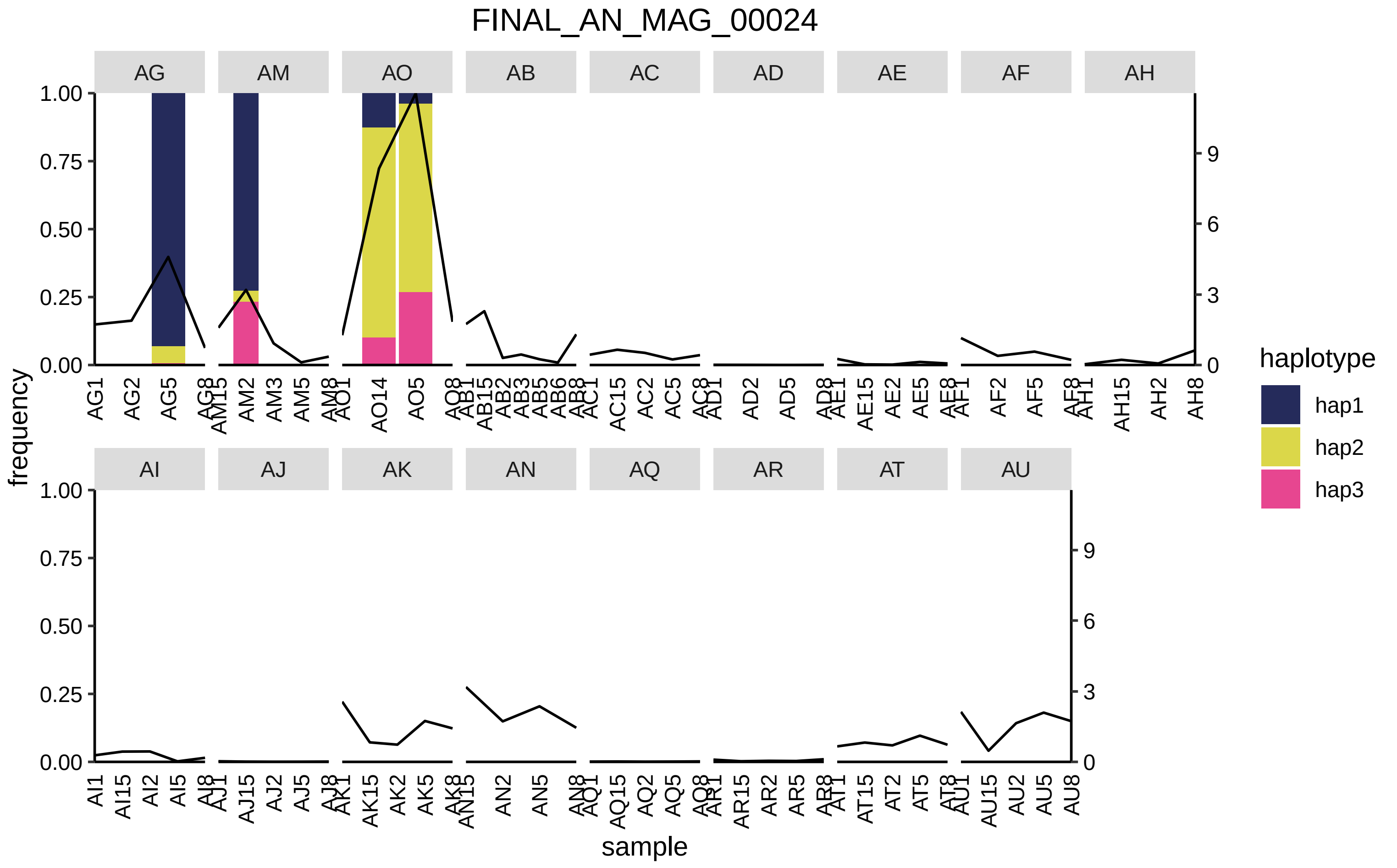

FINAL_AN_MAG_00024
frequency
sample
haplotype
hap1
hap2
hap3
AG
AM
AO
AB
AC
AD
AE
AF
AH
AI
AJ
AK
AN
AQ
AR
AT
AU

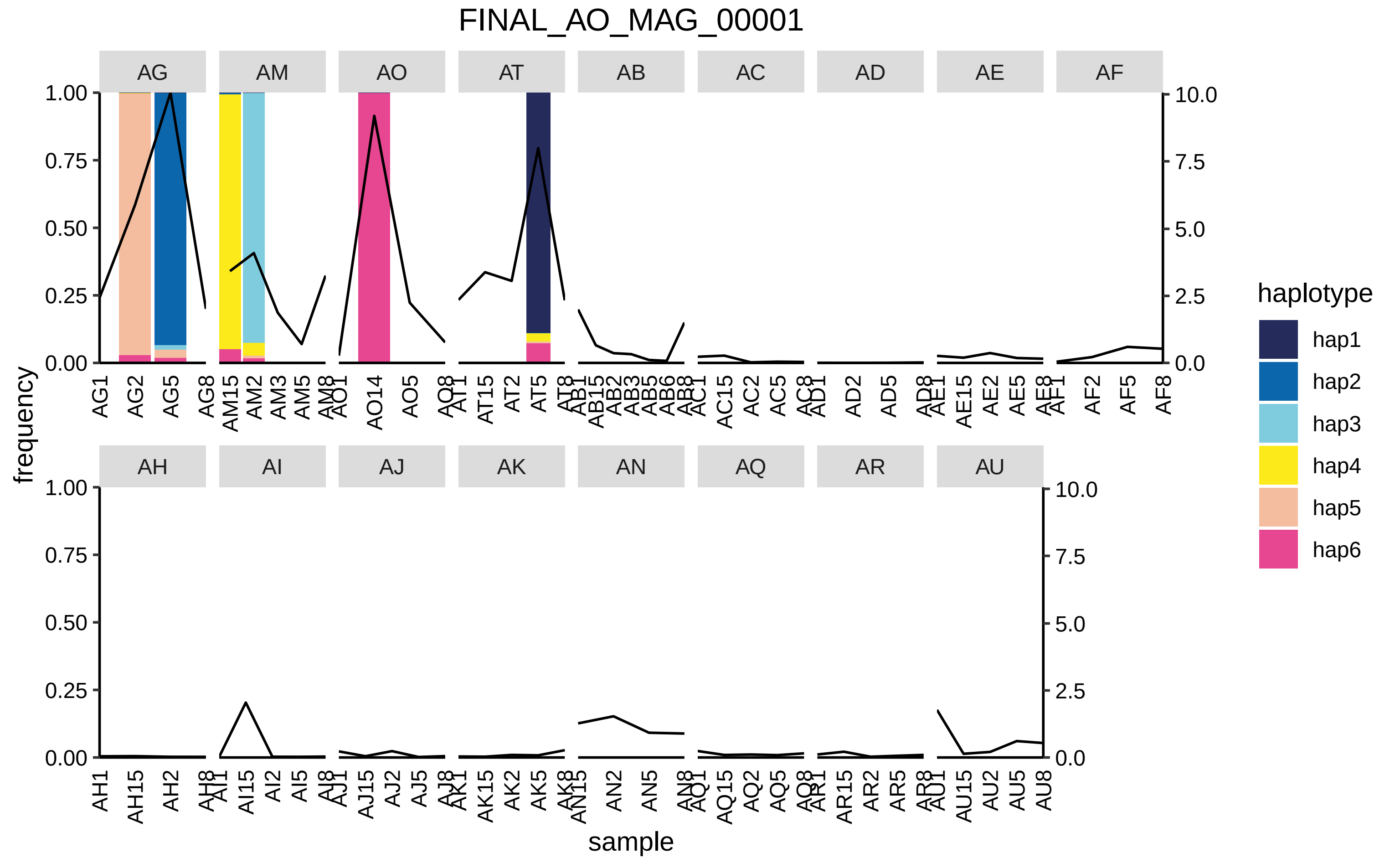
FINAL_AO_MAG_00001
AG
AM
AO
AT
AB
AC
AD
AE
AF
AH
AI
AJ
AK
AN
AQ
AR
AU
frequency
sample
haplotype
hap1
hap2
hap3
hap4
hap5
hap6
1.00
0.75
0.50
0.25
0.00
10.0
7.5
5.0
2.5
0.0

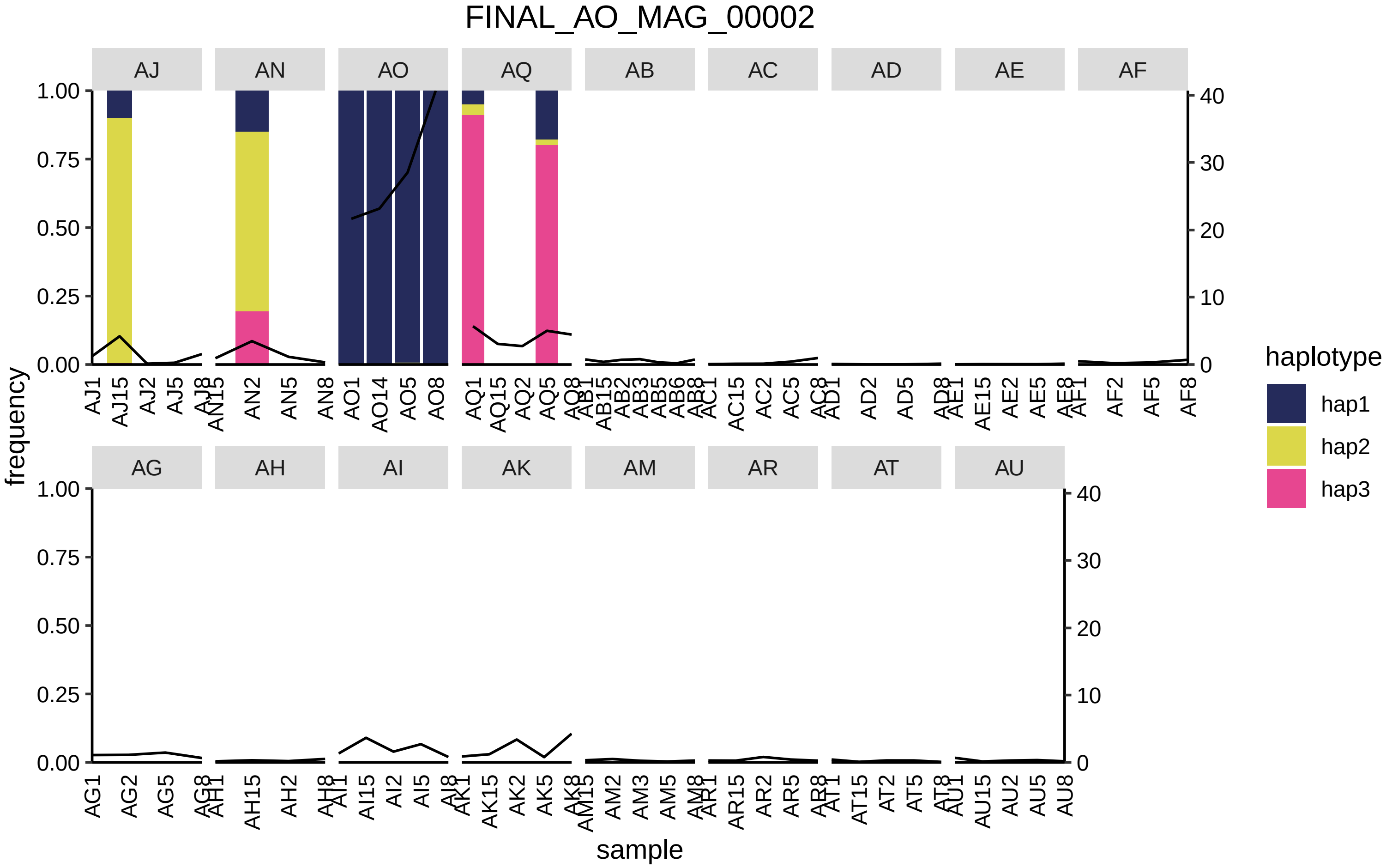
FINAL_AO_MAG_00002
AJ
AN
AO
AQ
AB
AC
AD
AE
AF
AG
AH
AI
AK
AM
AR
AT
AU
frequency
sample
haplotype
hap1
hap2
hap3
1.00
0.75
0.50
0.25
0.00
40
30
20
10
0
AJ1 AJ15 AJ2 AJ5 AJ8
AN15 AN2 AN5 AN8
AO1 AO14 AO5 AO8
AQ1 AQ15 AQ2 AQ5 AQ8
AB1 AB15 AB2 AB3 AB5 AB6 AB8
AC1 AC15 AC2 AC5 AC8
AD1 AD2 AD5 AD8
AE1 AE15 AE2 AE5 AE8
AF1 AF2 AF5 AF8
AG1 AG2 AG5 AG8
AH1 AH15 AH2 AH8
AI1 AI15 AI2 AI5 AI8
AK1 AK15 AK2 AK5 AK8
AM15 AM2 AM3 AM5 AM8
AR1 AR15 AR2 AR5 AR8
AT1 AT15 AT2 AT5 AT8
AU1 AU15 AU2 AU5 AU8

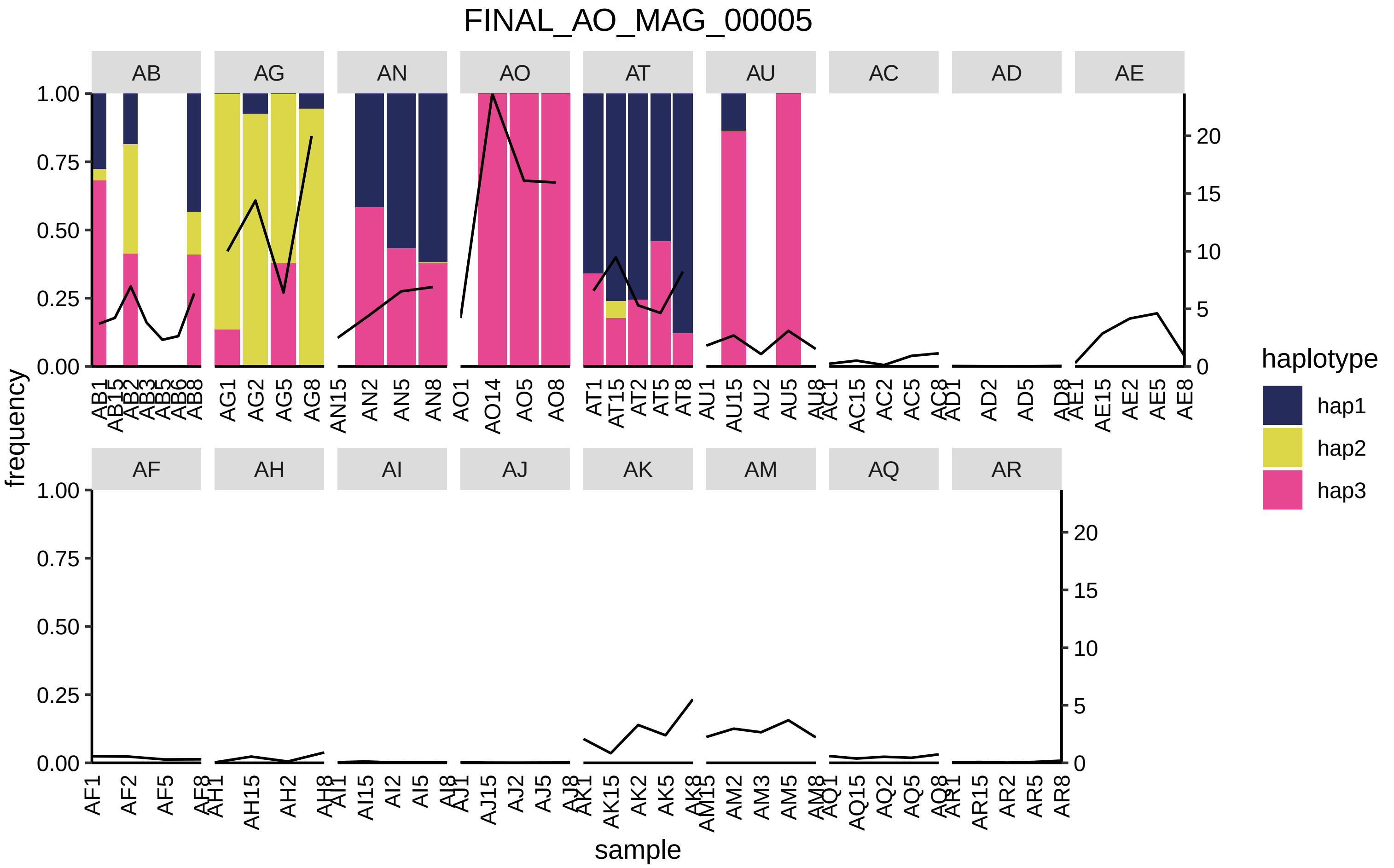

FINAL_AO_MAG_00005
AB
AG
AN
AO
AT
AU
AC
AD
AE
AF
AH
AI
AJ
AK
AM
AQ
AR
frequency
sample
haplotype
hap1
hap2
hap3

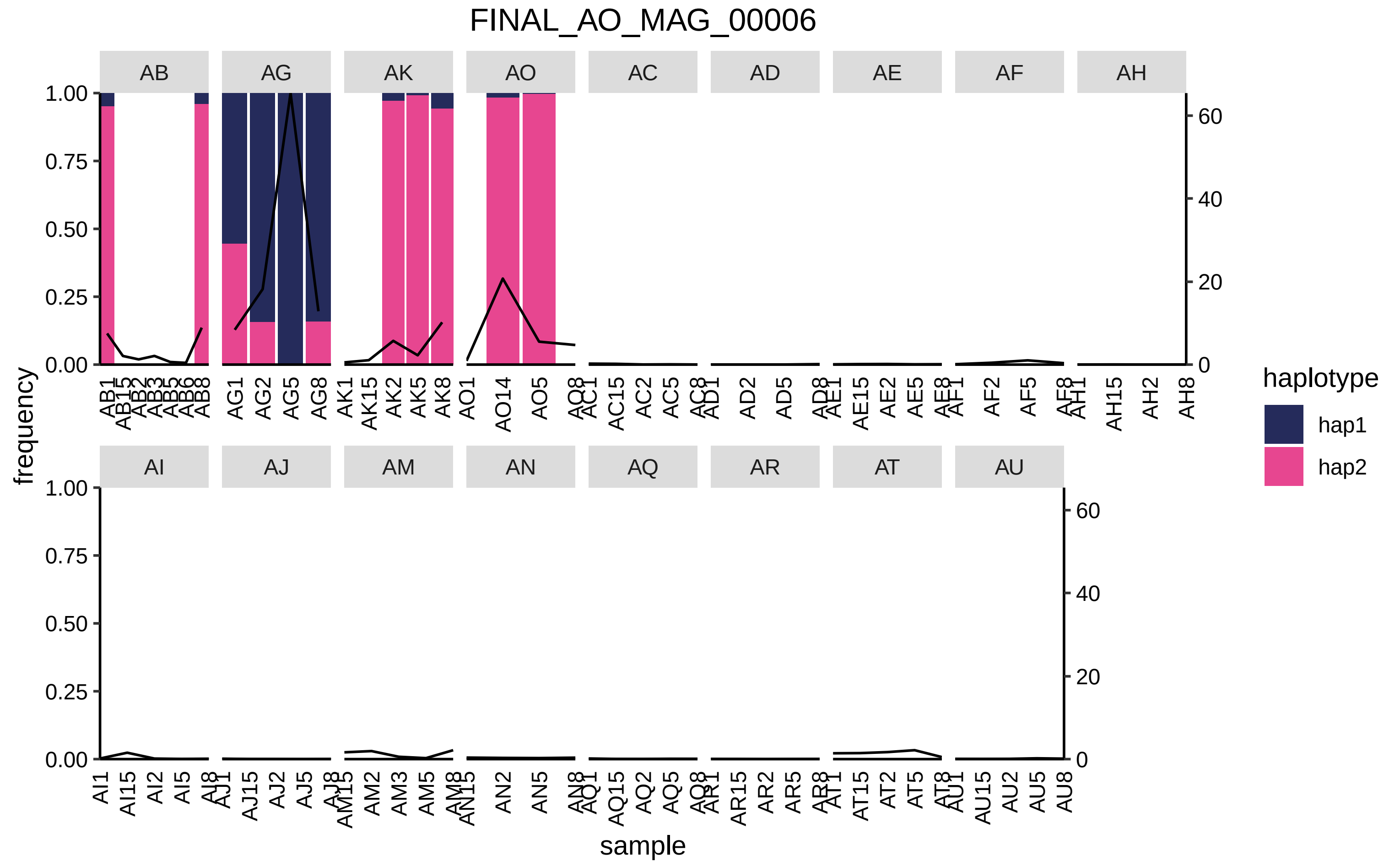
FINAL_AO_MAG_00006
AB
AG
AK
AO
AC
AD
AE
AF
AH
AI
AJ
AM
AN
AQ
AR
AT
AU
frequency
sample
haplotype
hap1
hap2

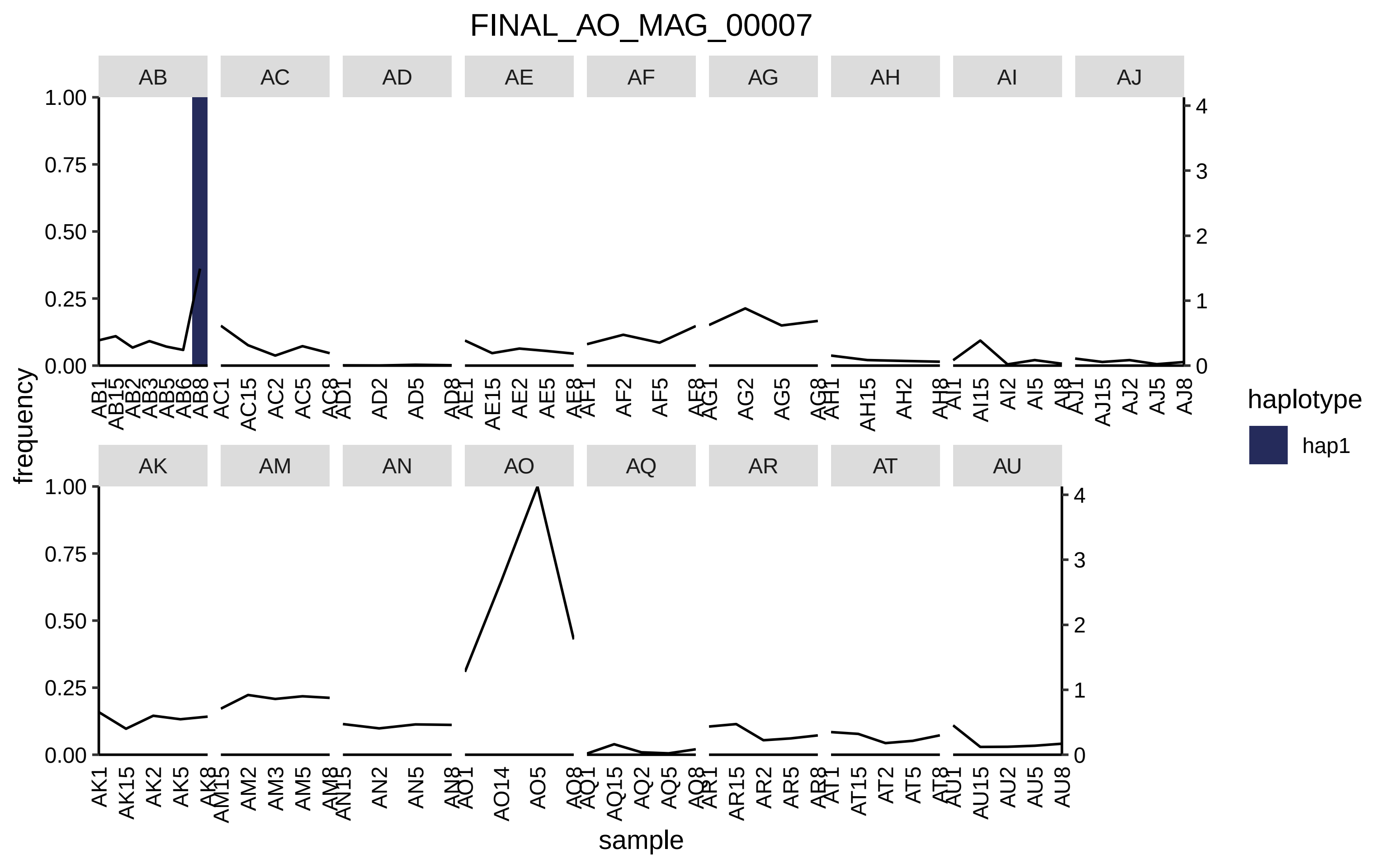

FINAL_AO_MAG_00007
AB
AC
AD
AE
AF
AG
AH
AI
AJ
AK
AM
AN
AO
AQ
AR
AT
AU
frequency
sample
haplotype
hap1

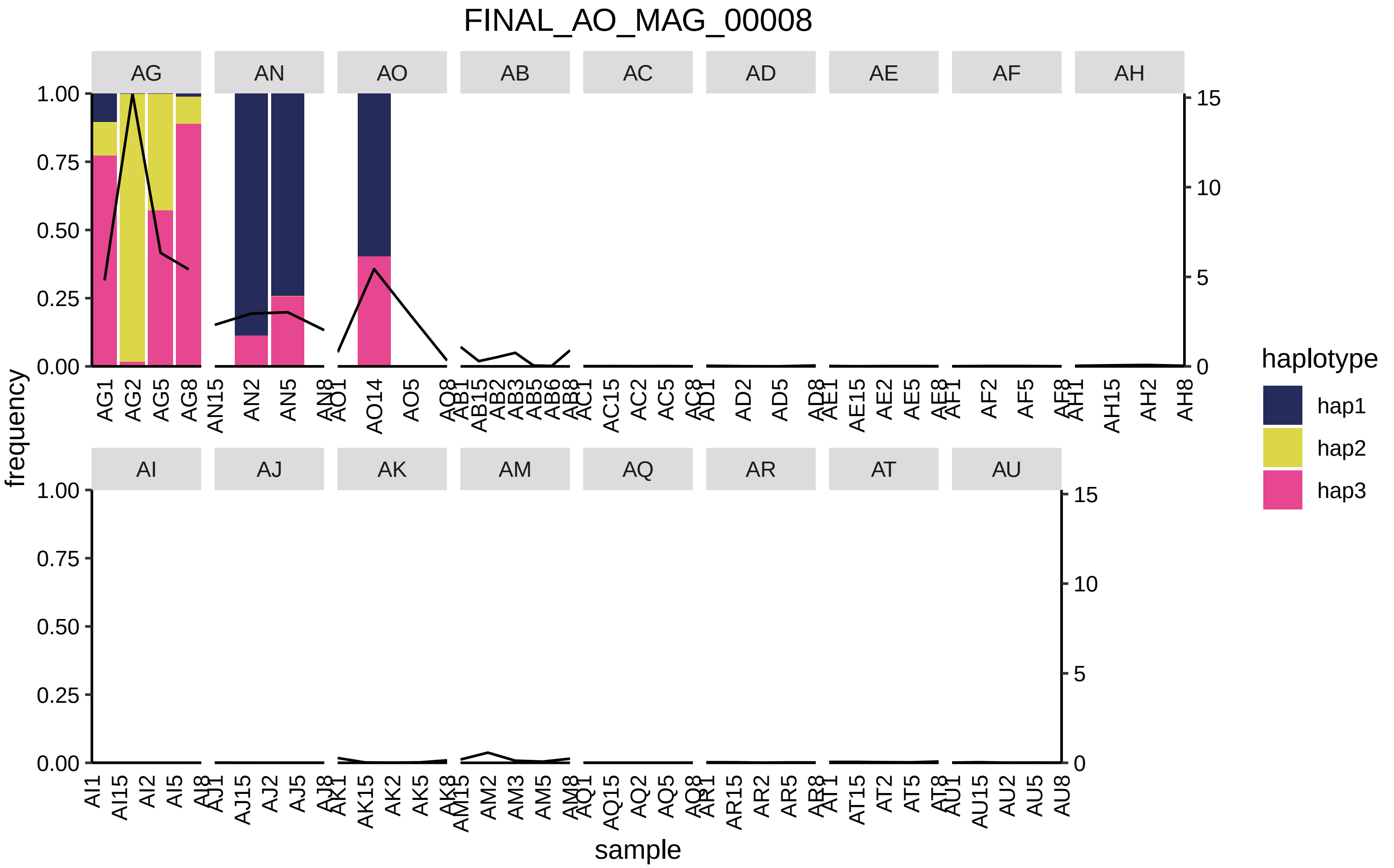
FINAL_AO_MAG_00008
AG
AN
AO
AB
AC
AD
AE
AF
AH
AI
AJ
AK
AM
AQ
AR
AT
AU
frequency
sample
haplotype
hap1
hap2
hap3

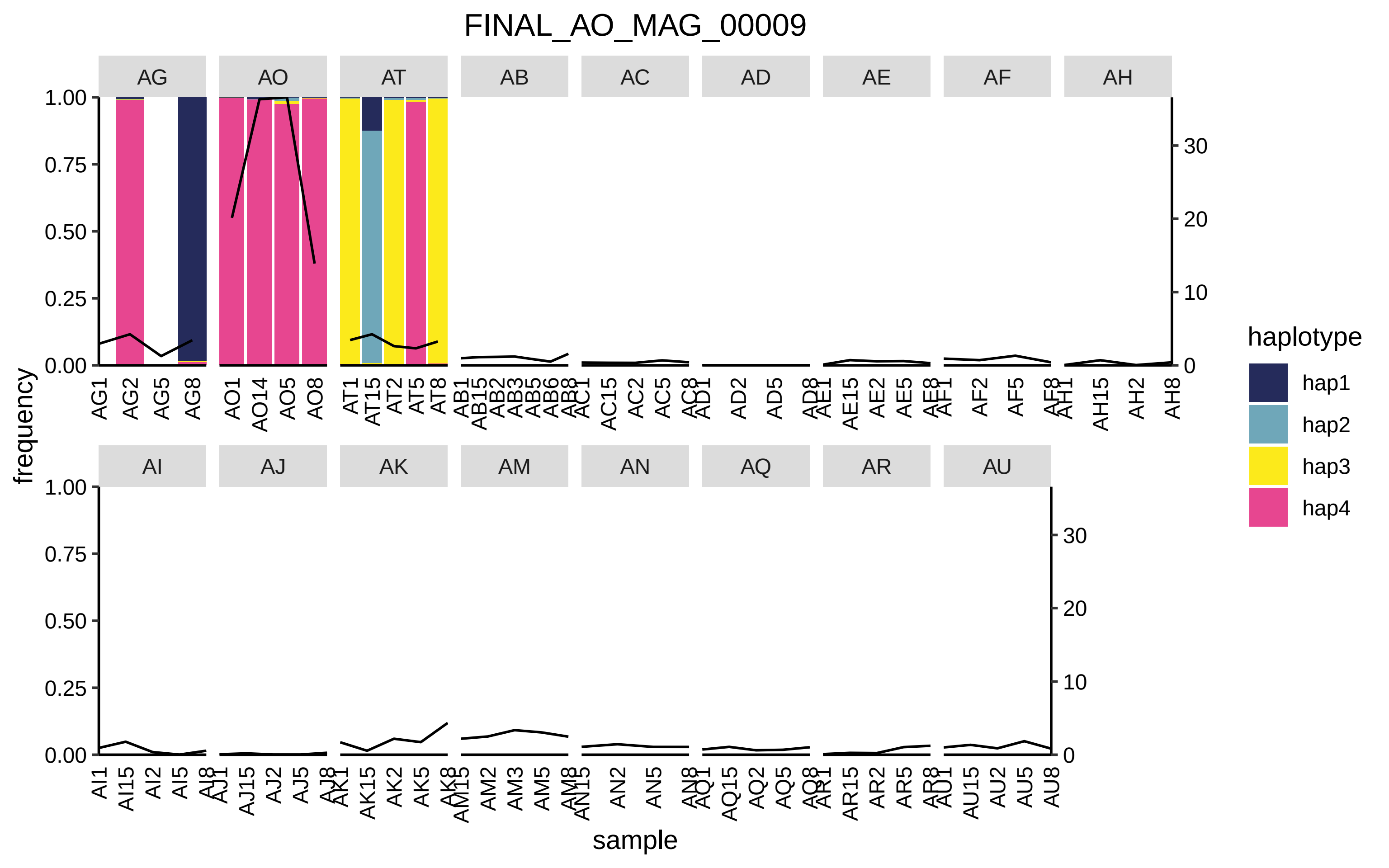

FINAL_AO_MAG_00009
AG
AO
AT
AB
AC
AD
AE
AF
AH
AI
AJ
AK
AM
AN
AQ
AR
AU
frequency
sample
haplotype
hap1
hap2
hap3
hap4

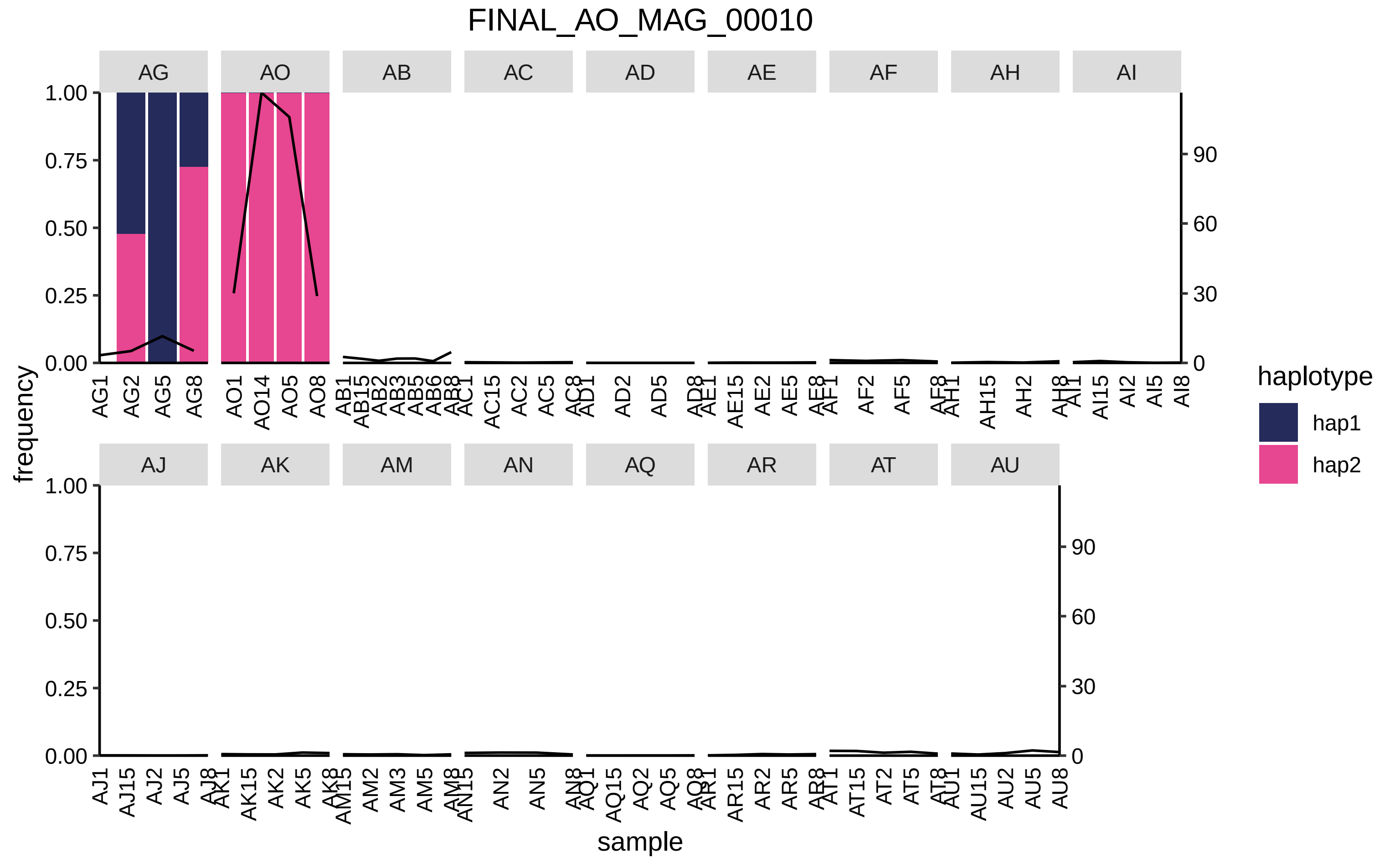

FINAL_AO_MAG_00010
AG
AO
AB
AC
AD
AE
AF
AH
AI
AJ
AK
AM
AN
AQ
AR
AT
AU
frequency
sample
haplotype
hap1
hap2

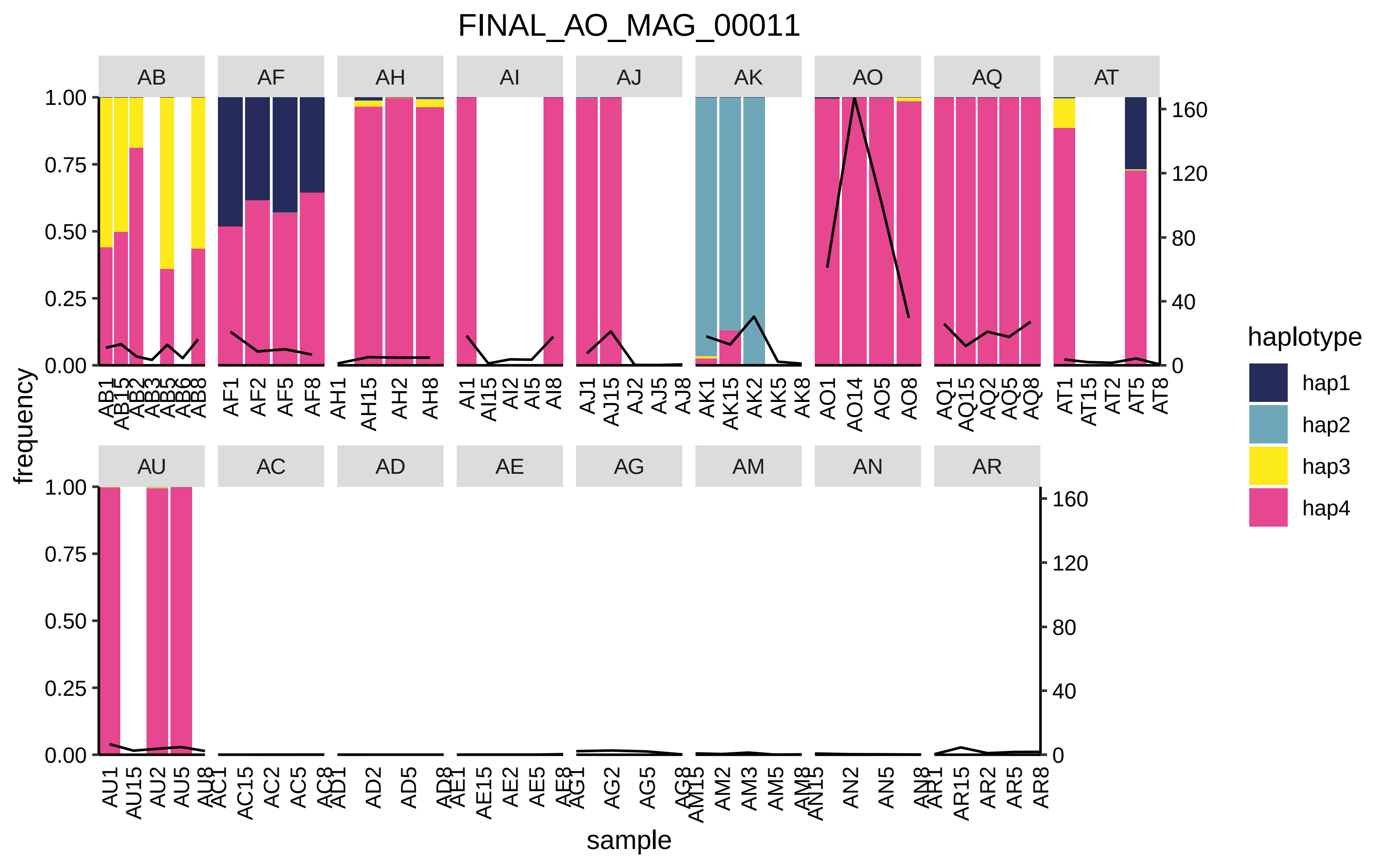

FINAL_AO_MAG_00011
AB
AF
AH
AI
AJ
AK
AO
AQ
AT
AU
AC
AD
AE
AG
AM
AN
AR
frequency
sample
haplotype
hap1
hap2
hap3
hap4

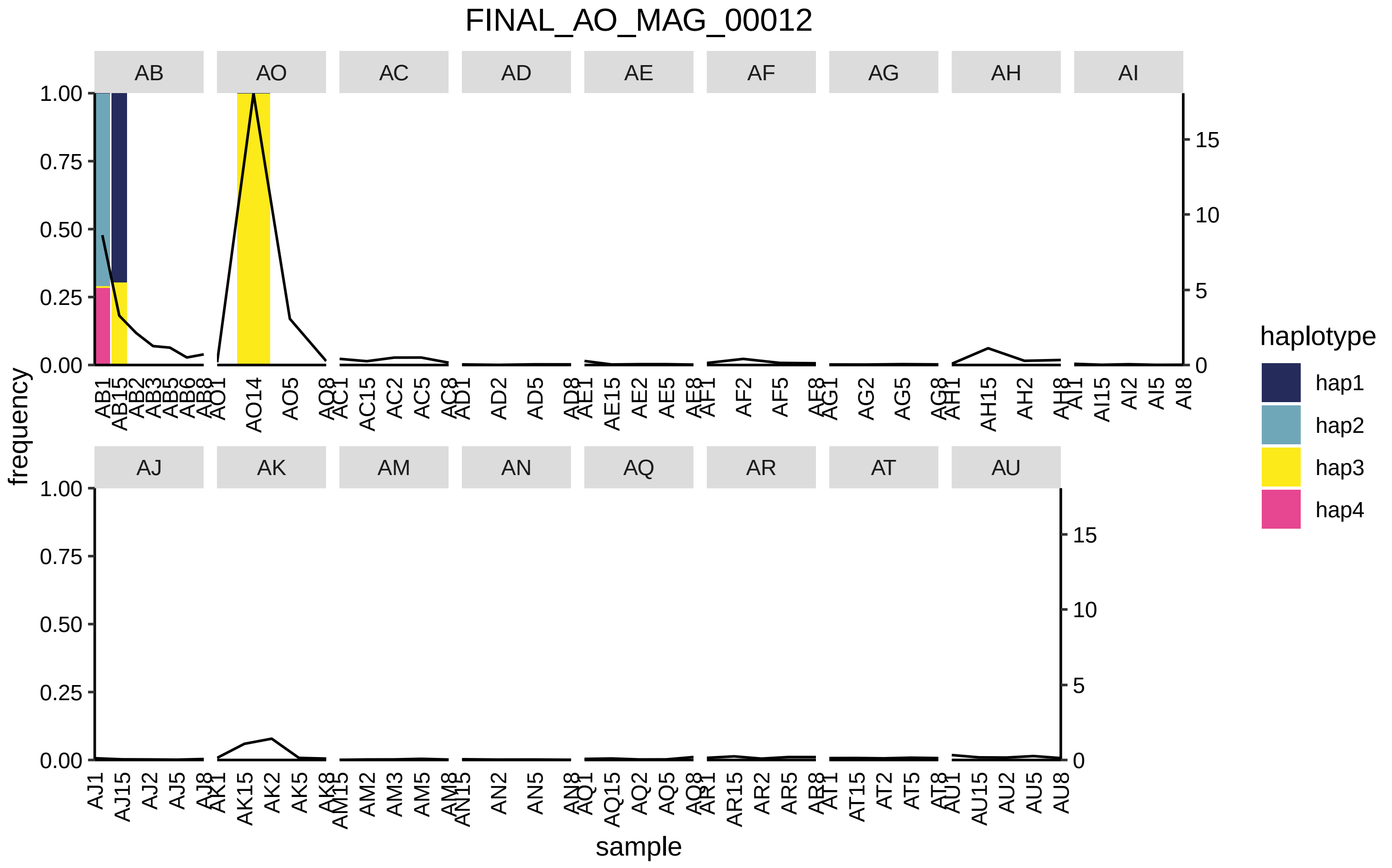

FINAL_AO_MAG_00012
AB
AO
AC
AD
AE
AF
AG
AH
AI
AJ
AK
AM
AN
AQ
AR
AT
AU
frequency
sample
haplotype
hap1
hap2
hap3
hap4

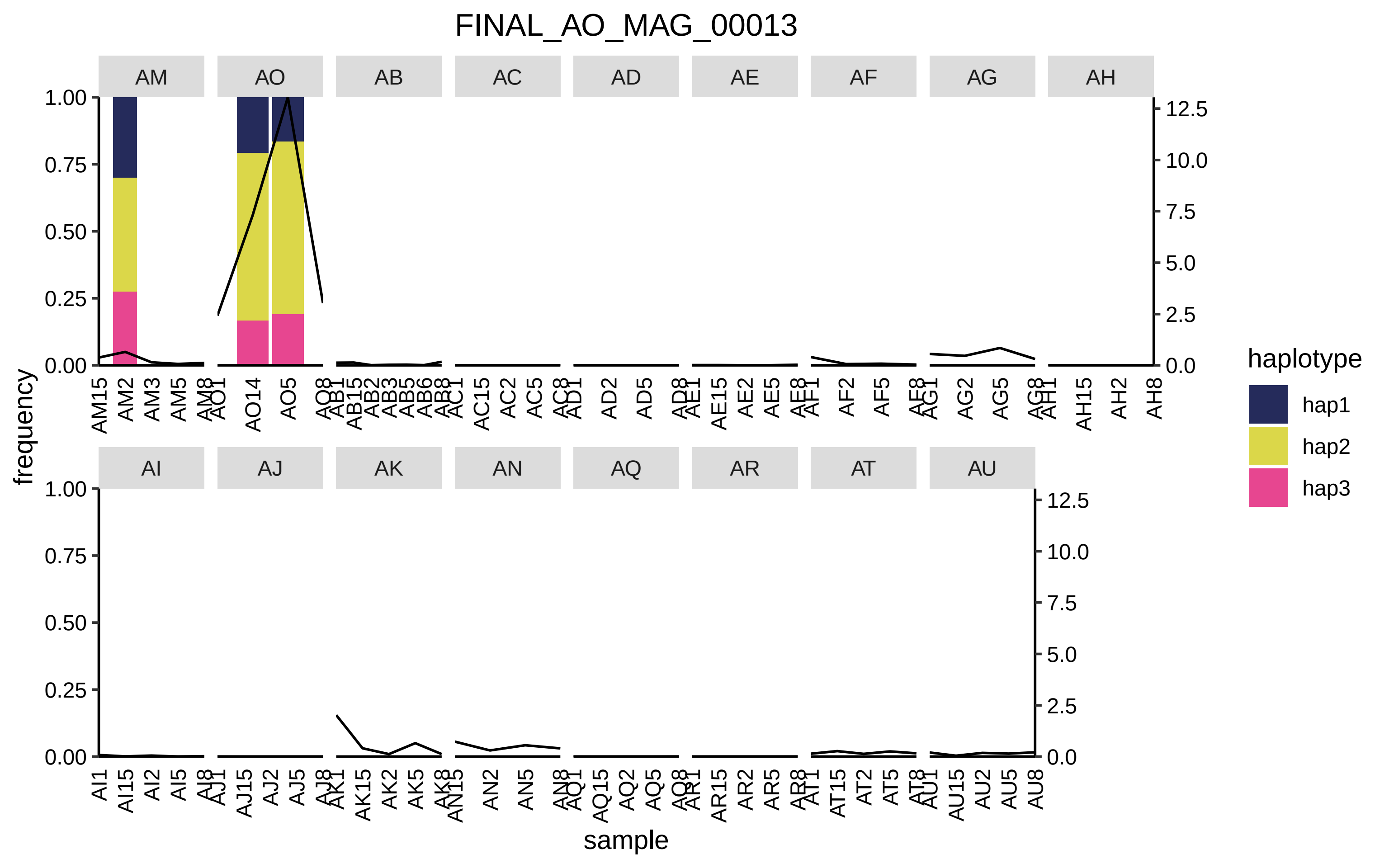

FINAL_AO_MAG_00013
AM
AO
AB
AC
AD
AE
AF
AG
AH
AI
AJ
AK
AN
AQ
AR
AT
AU
frequency
sample
haplotype
hap1
hap2
hap3

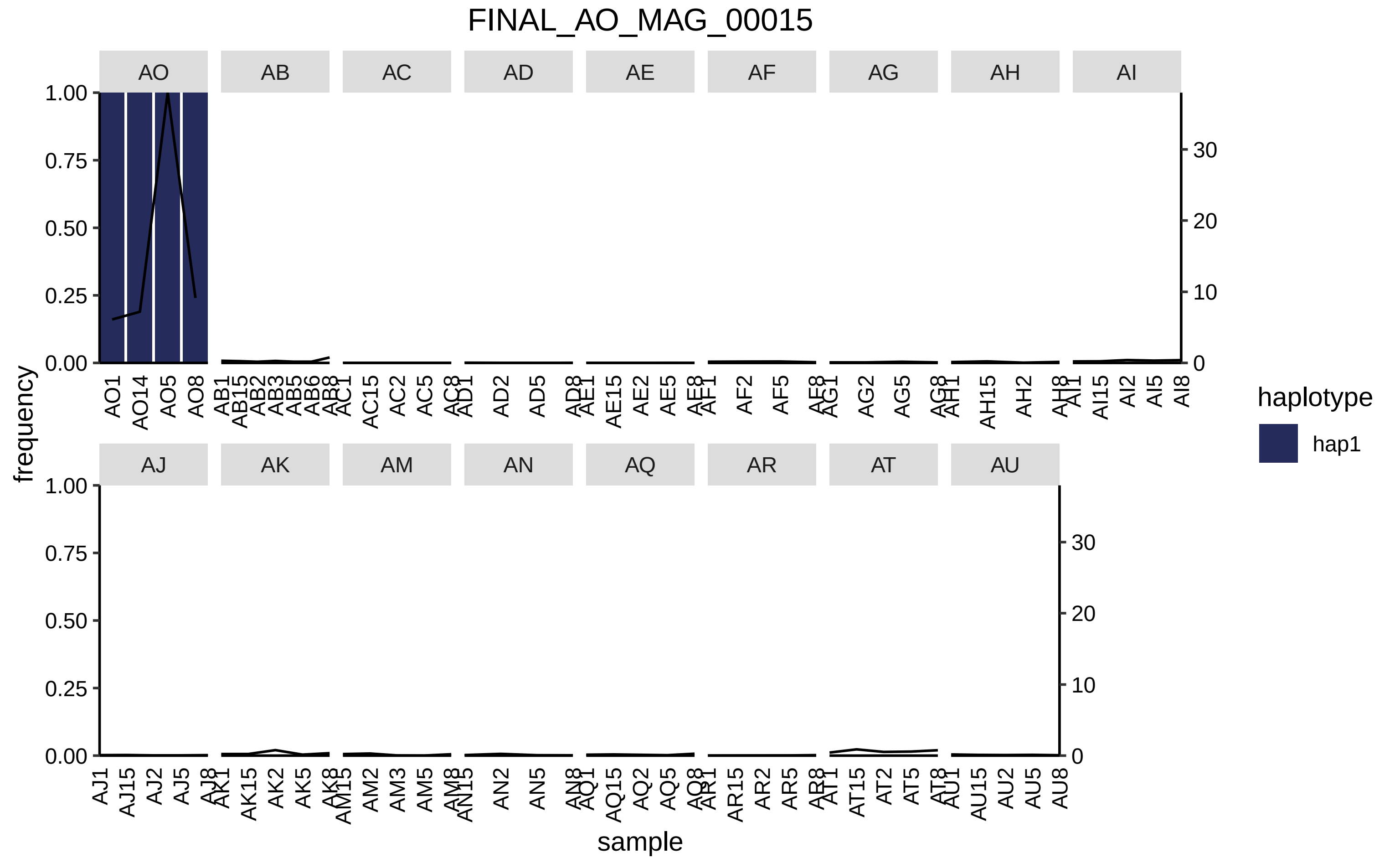

FINAL_AO_MAG_00015
AO
AB
AC
AD
AE
AF
AG
AH
AI
AJ
AK
AM
AN
AQ
AR
AT
AU
frequency
sample
haplotype
hap1
1.00
0.75
0.50
0.25
0.00
0
10
20
30
AO1 AO14 AO5 AO8
AB1 AB15 AB2 AB3 AB5 AB6 AB8
AC1 AC15 AC2 AC5 AC8
AD1 AD2 AD5 AD8
AE1 AE15 AE2 AE5 AE8
AF1 AF2 AF5 AF8
AG1 AG2 AG5 AG8
AH1 AH15 AH2 AH8
AI1 AI15 AI2 AI5 AI8
AJ1 AJ15 AJ2 AJ5 AJ8
AK1 AK15 AK2 AK5 AK8
AM15 AM2 AM3 AM5 AM8
AN15 AN2 AN5 AN8
AQ1 AQ15 AQ2 AQ5 AQ8
AR1 AR15 AR2 AR5 AR8
AT1 AT15 AT2 AT5 AT8
AU1 AU15 AU2 AU5 AU8

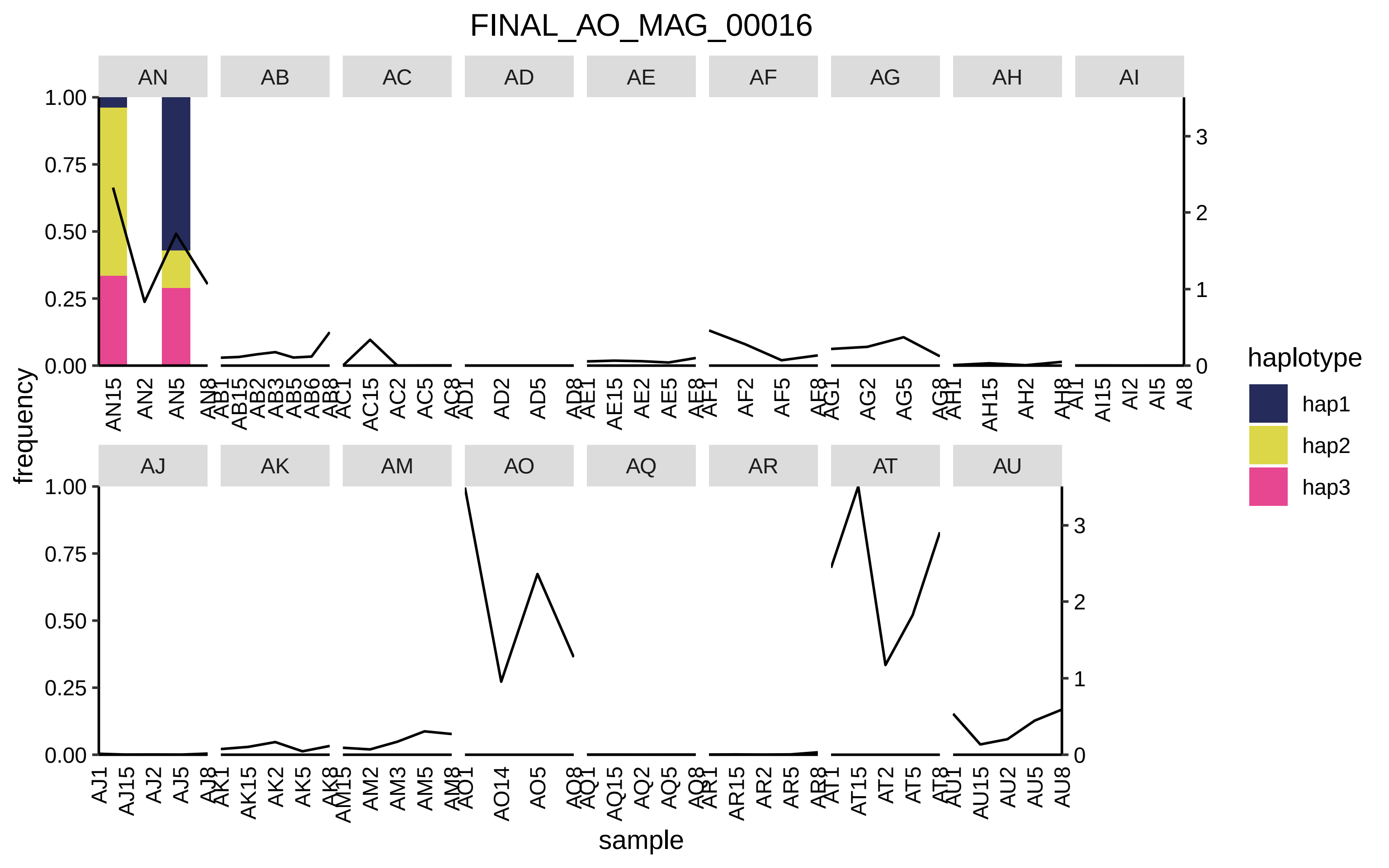
FINAL_AO_MAG_00016
AN
AB
AC
AD
AE
AF
AG
AH
AI
AJ
AK
AM
AO
AQ
AR
AT
AU
frequency
sample
haplotype
hap1
hap2
hap3

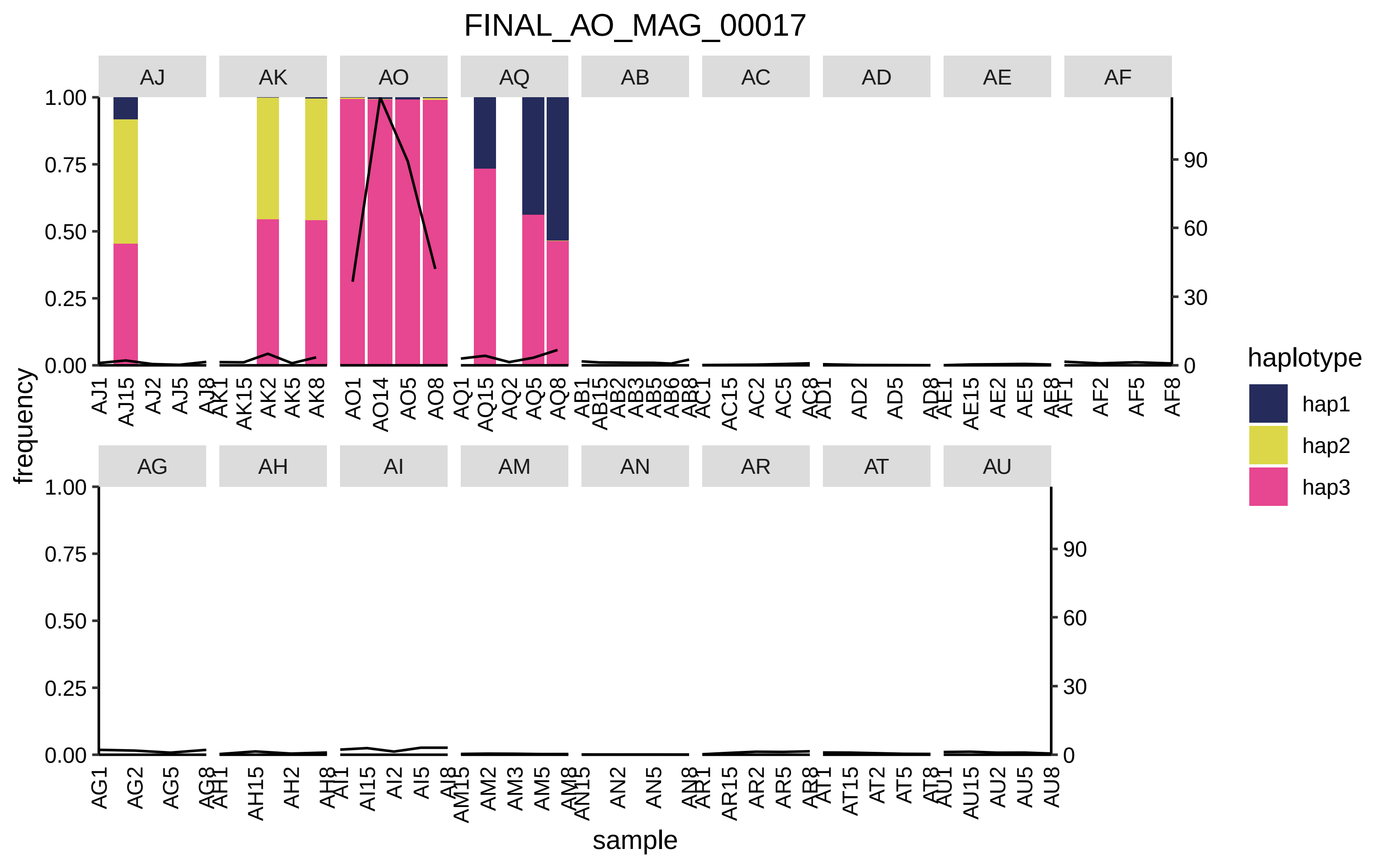

FINAL_AO_MAG_00017
AJ
AK
AO
AQ
AB
AC
AD
AE
AF
AG
AH
AI
AM
AN
AR
AT
AU
frequency
1.00
0.75
0.50
0.25
0.00
90
60
30
0
AJ1 AJ15 AJ2 AJ5 AJ8
AK1 AK15 AK2 AK5 AK8
AO1 AO14 AO5 AO8
AQ1 AQ15 AQ2 AQ5 AQ8
AB1 AB15 AB2 AB3 AB5 AB6 AB8
AC1 AC15 AC2 AC5 AC8
AD1 AD2 AD5 AD8
AE1 AE15 AE2 AE5 AE8
AF1 AF2 AF5 AF8
AG1 AG2 AG5 AG8
AH1 AH15 AH2 AH8
AI1 AI15 AI2 AI5 AI8
AM15 AM2 AM3 AM5 AM8
AN15 AN2 AN5 AN8
AR1 AR15 AR2 AR5 AR8
AT1 AT15 AT2 AT5 AT8
AU1 AU15 AU2 AU5 AU8
sample
haplotype
hap1
hap2
hap3

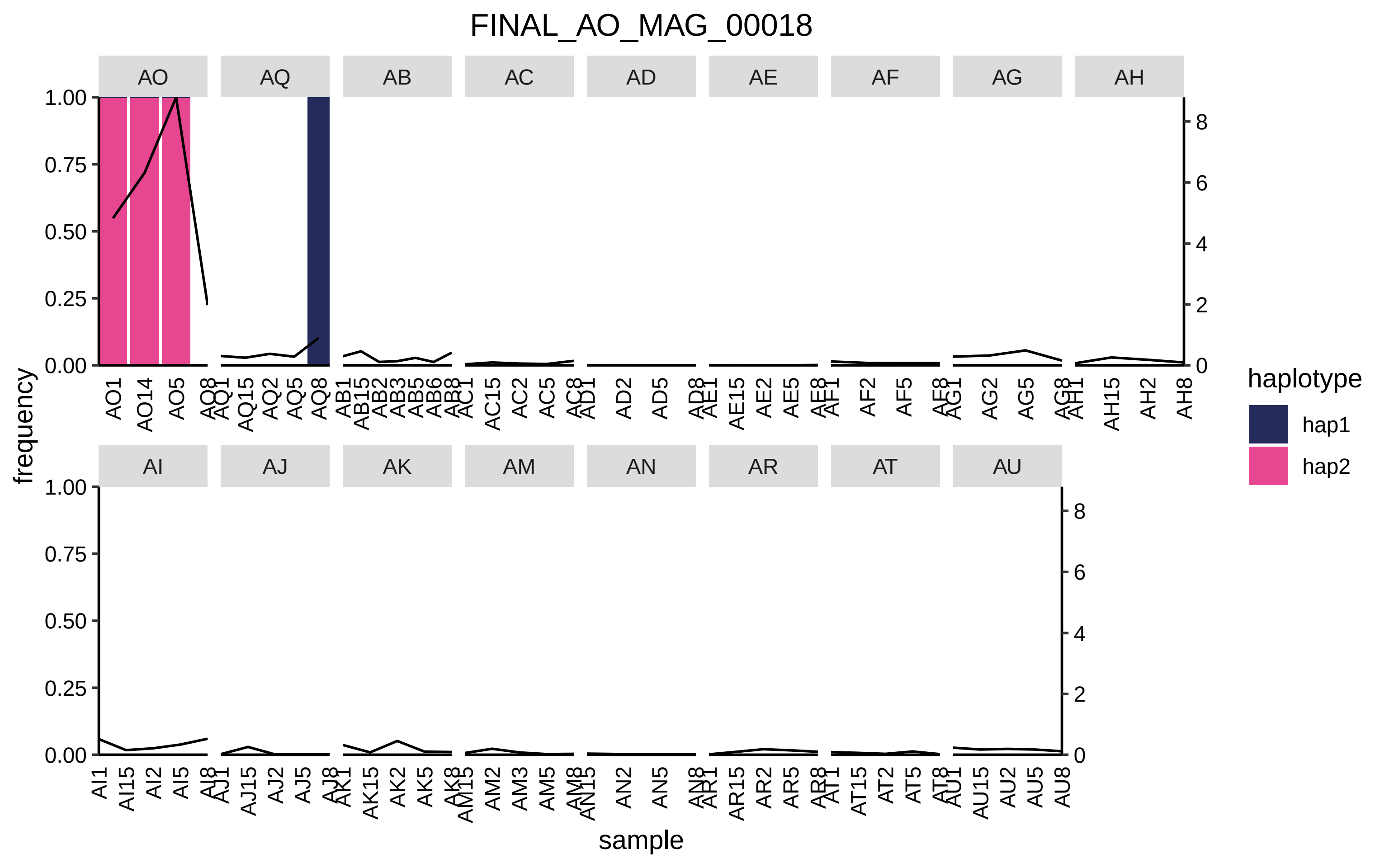

FINAL_AO_MAG_00018
AO
AQ
AB
AC
AD
AE
AF
AG
AH
AI
AJ
AK
AM
AN
AR
AT
AU
frequency
sample
haplotype
hap1
hap2

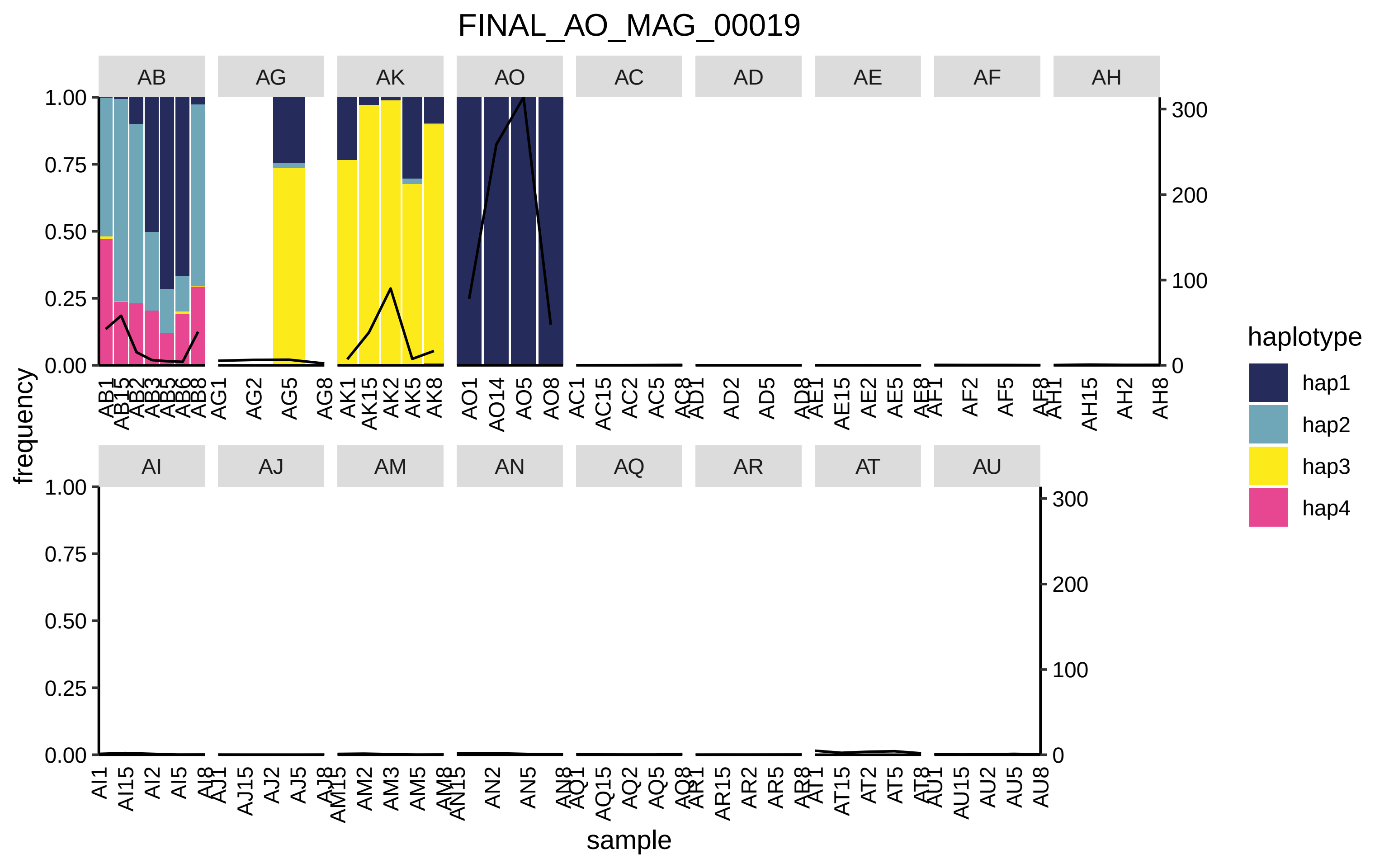
FINAL_AO_MAG_00019
AB
AG
AK
AO
AC
AD
AE
AF
AH
AI
AJ
AM
AN
AQ
AR
AT
AU
frequency
sample
haplotype
hap1
hap2
hap3
hap4

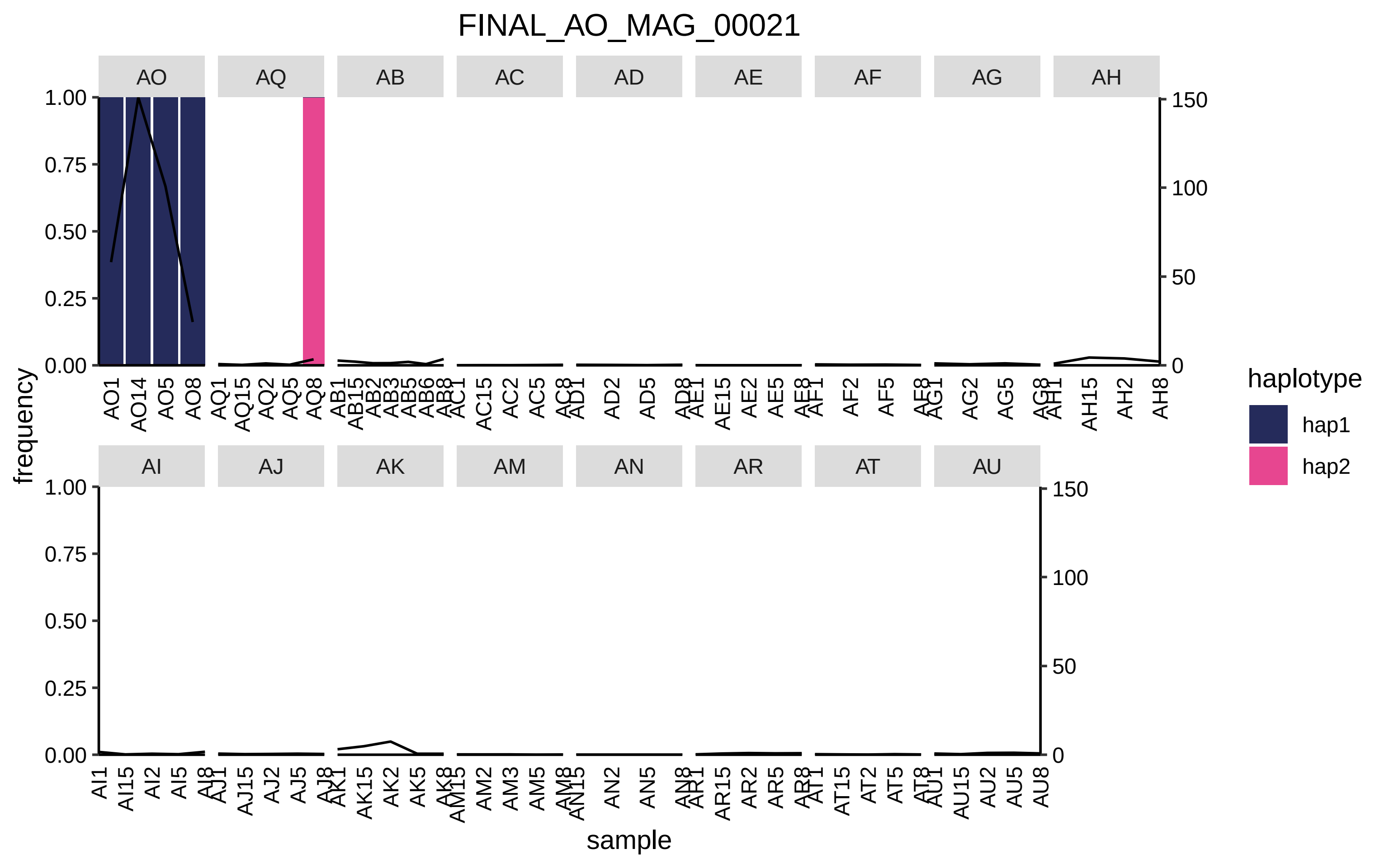
FINAL_AO_MAG_00021
AO
AQ
AB
AC
AD
AE
AF
AG
AH
AI
AJ
AK
AM
AN
AR
AT
AU
frequency
sample
haplotype
hap1
hap2

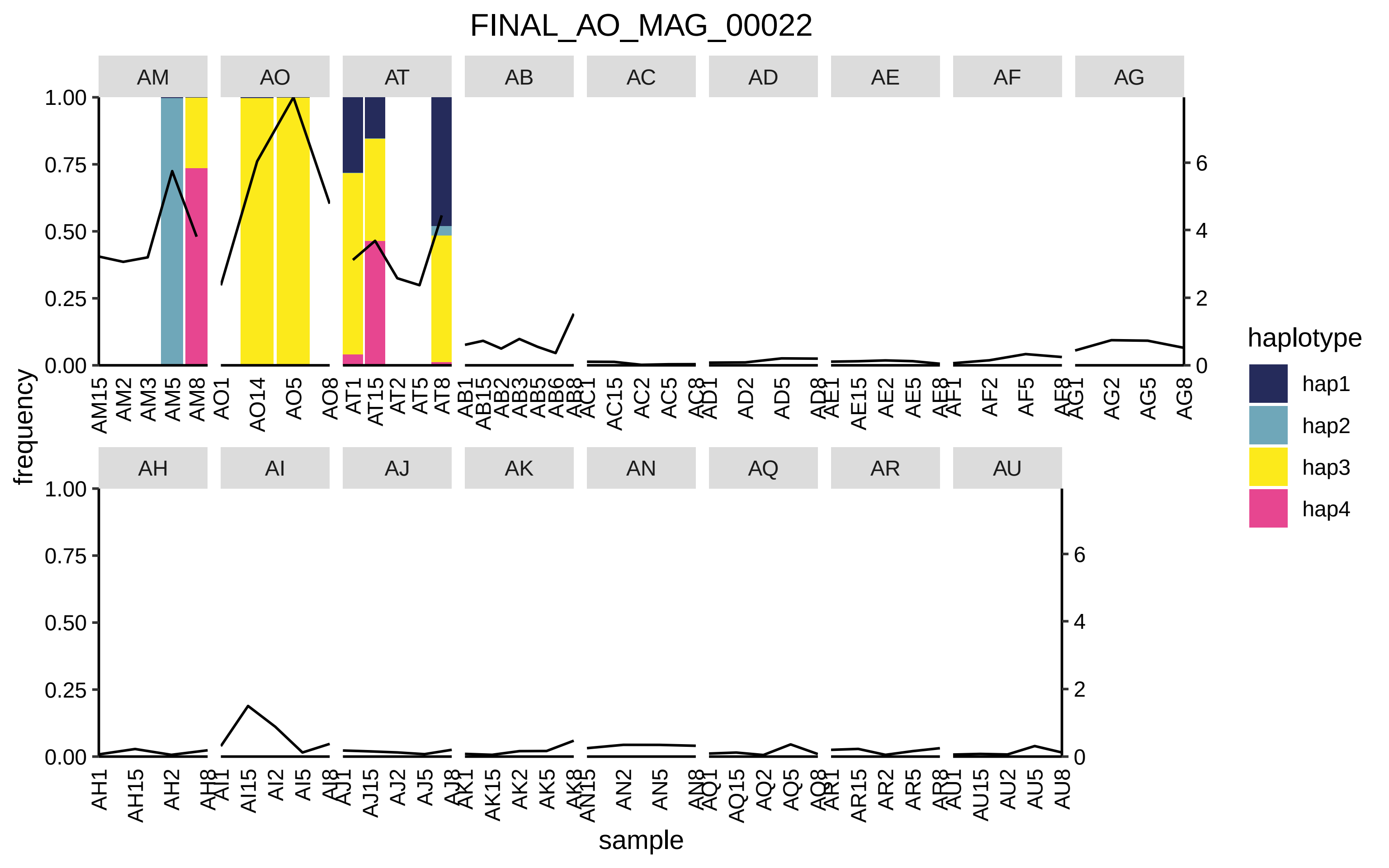

FINAL_AO_MAG_00022
AM
AO
AT
AB
AC
AD
AE
AF
AG
AH
AI
AJ
AK
AN
AQ
AR
AU
frequency
sample
haplotype
hap1
hap2
hap3
hap4

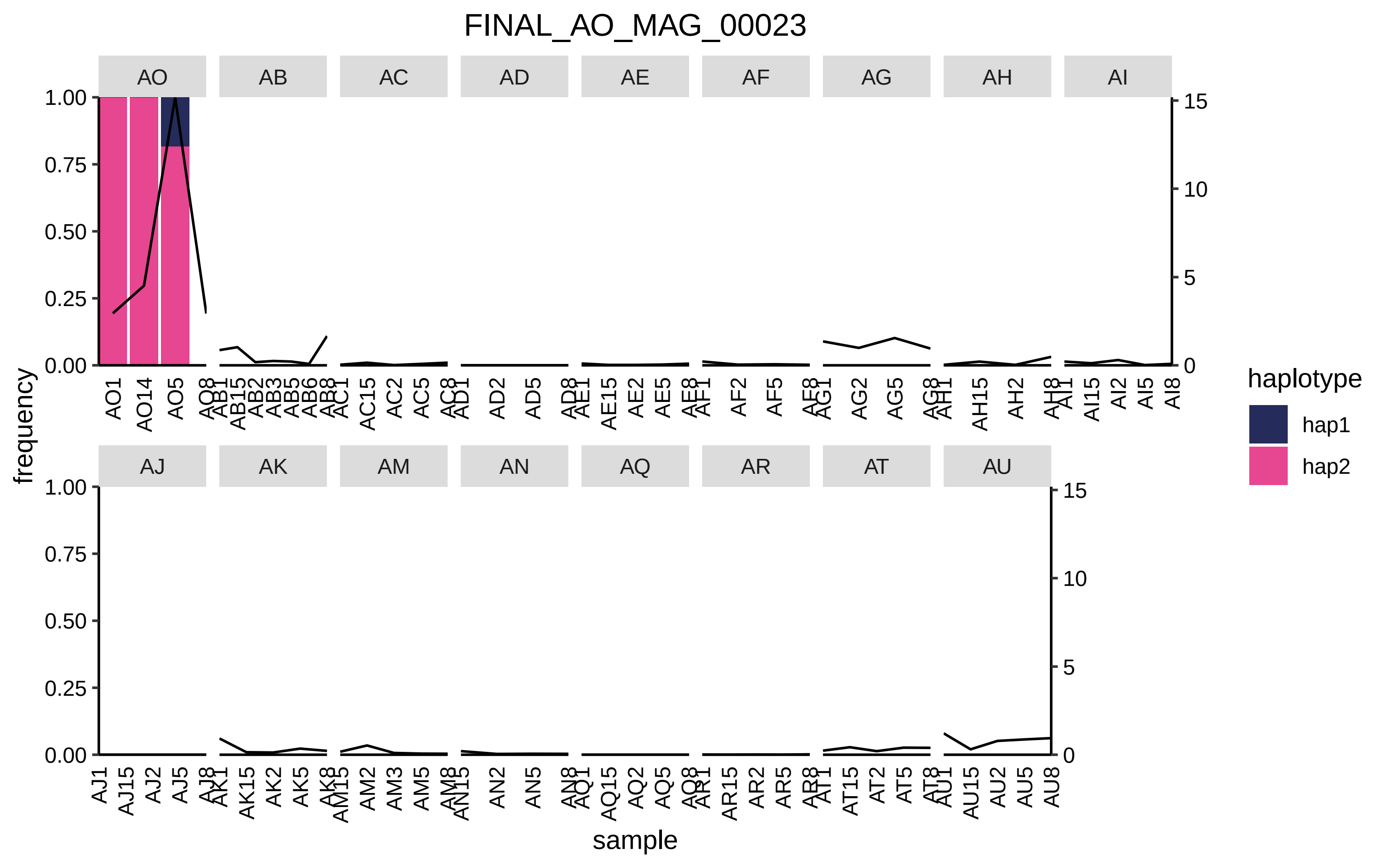

FINAL_AO_MAG_00023
AO
AB
AC
AD
AE
AF
AG
AH
AI
AJ
AK
AM
AN
AQ
AR
AT
AU
frequency
sample
haplotype
hap1
hap2

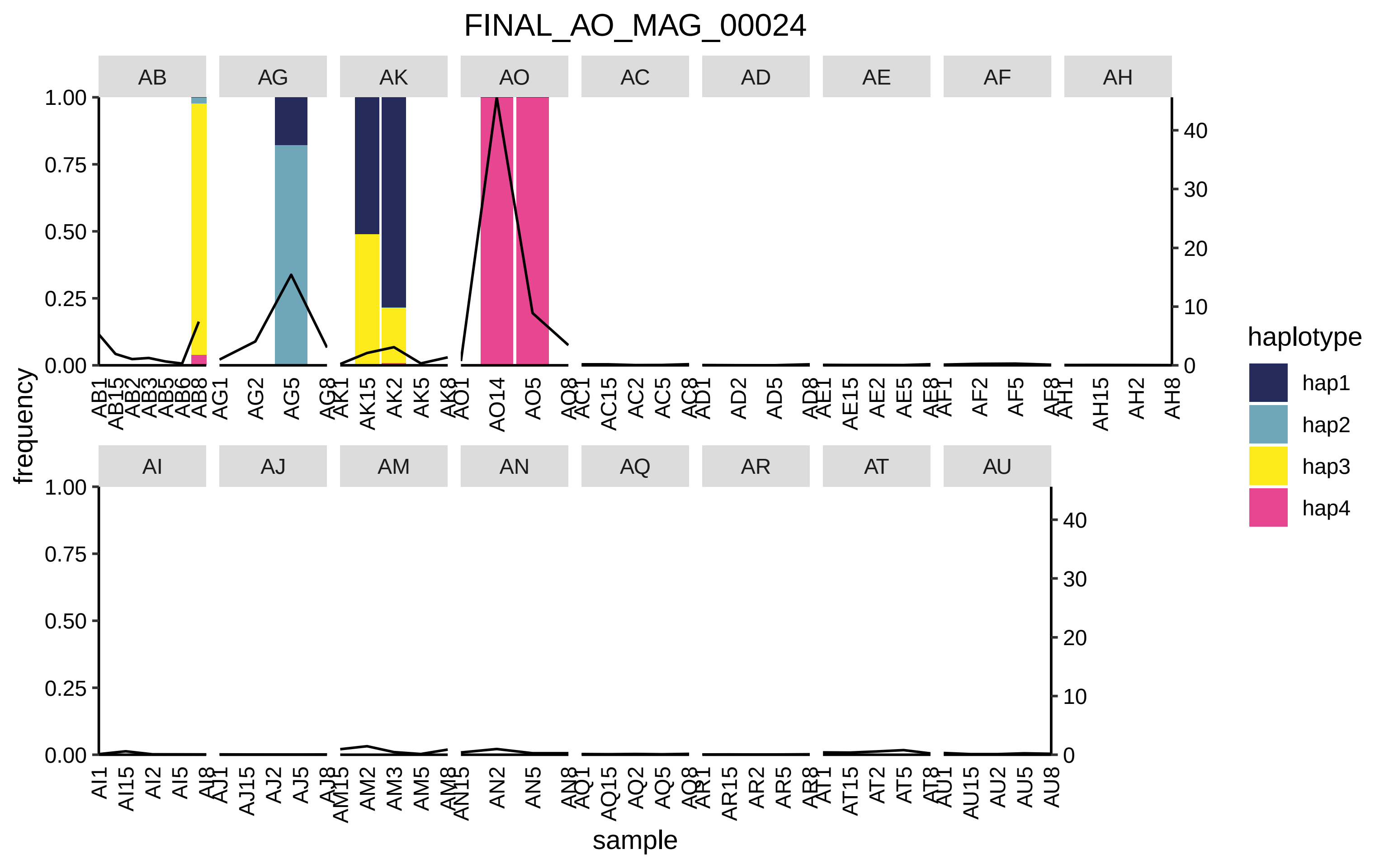

FINAL_AO_MAG_00024
AB
AG
AK
AO
AC
AD
AE
AF
AH
AI
AJ
AM
AN
AQ
AR
AT
AU
frequency
sample
haplotype
hap1
hap2
hap3
hap4

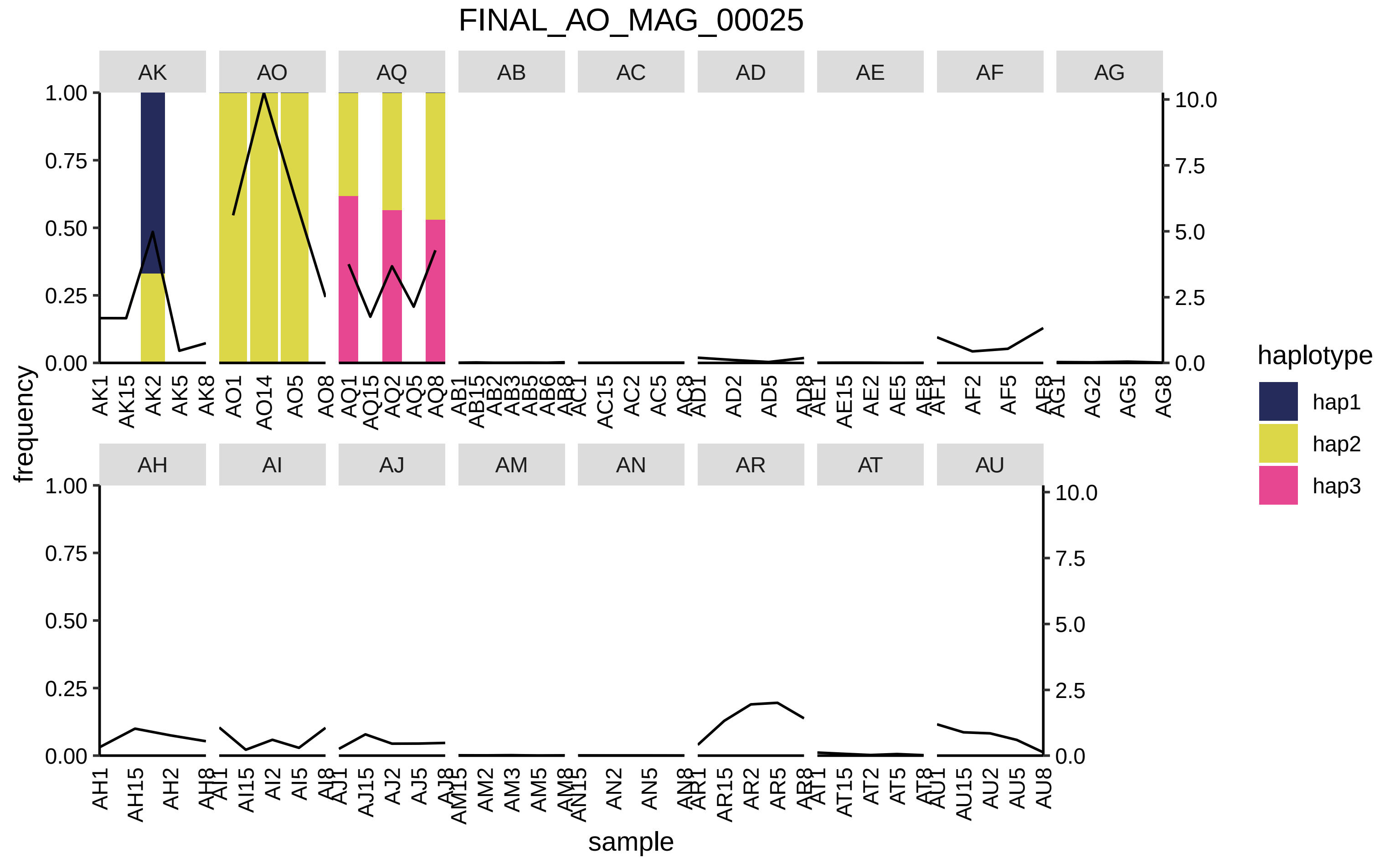

FINAL_AO_MAG_00025
AK
AO
AQ
AB
AC
AD
AE
AF
AG
AH
AI
AJ
AM
AN
AR
AT
AU
frequency
sample
haplotype
hap1
hap2
hap3
1.00
0.75
0.50
0.25
0.00
10.0
7.5
5.0
2.5
0.0
AK1 AK15 AK2 AK5 AK8
AO1 AO14 AO5 AO8
AQ1 AQ15 AQ2 AQ5 AQ8
AB1 AB15 AB2 AB3 AB5 AB6 AB8
AC1 AC15 AC2 AC5 AC8
AD1 AD2 AD5 AD8
AE1 AE15 AE2 AE5 AE8
AF1 AF2 AF5 AF8
AG1 AG2 AG5 AG8
AH1 AH15 AH2 AH8
AI1 AI15 AI2 AI5 AI8
AJ1 AJ15 AJ2 AJ5 AJ8
AM15 AM2 AM3 AM5 AM8
AN15 AN2 AN5 AN8
AR1 AR15 AR2 AR5 AR8
AT1 AT15 AT2 AT5 AT8
AU1 AU15 AU2 AU5 AU8

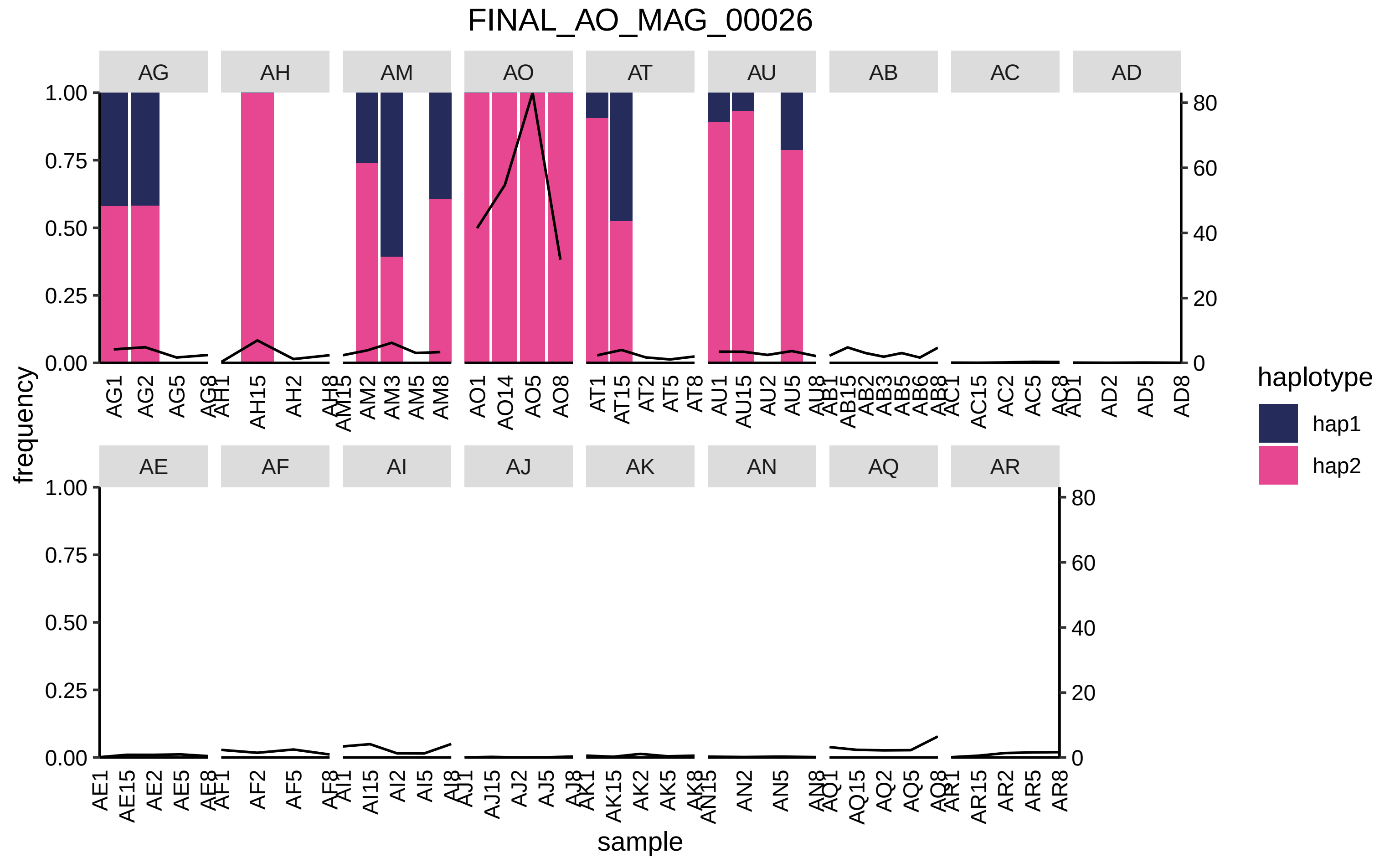
FINAL_AO_MAG_00026
AG
AH
AM
AO
AT
AU
AB
AC
AD
AE
AF
AI
AJ
AK
AN
AQ
AR
frequency
sample
haplotype
hap1
hap2
1.00
0.75
0.50
0.25
0.00
80
60
40
20
0
AG1 AG2 AG5 AG8
AH1 AH15 AH2 AH8
AM15 AM2 AM3 AM5 AM8
AO1 AO14 AO5 AO8
AT1 AT15 AT2 AT5 AT8
AU1 AU15 AU2 AU5 AU8
AB1 AB15 AB2 AB3 AB5 AB6 AB8
AC1 AC15 AC2 AC5 AC8
AD1 AD2 AD5 AD8
AE1 AE15 AE2 AE5 AE8
AF1 AF2 AF5 AF8
AI1 AI15 AI2 AI5 AI8
AJ1 AJ15 AJ2 AJ5 AJ8
AK1 AK15 AK2 AK5 AK8
AN15 AN2 AN5 AN8
AQ1 AQ15 AQ2 AQ5 AQ8
AR1 AR15 AR2 AR5 AR8

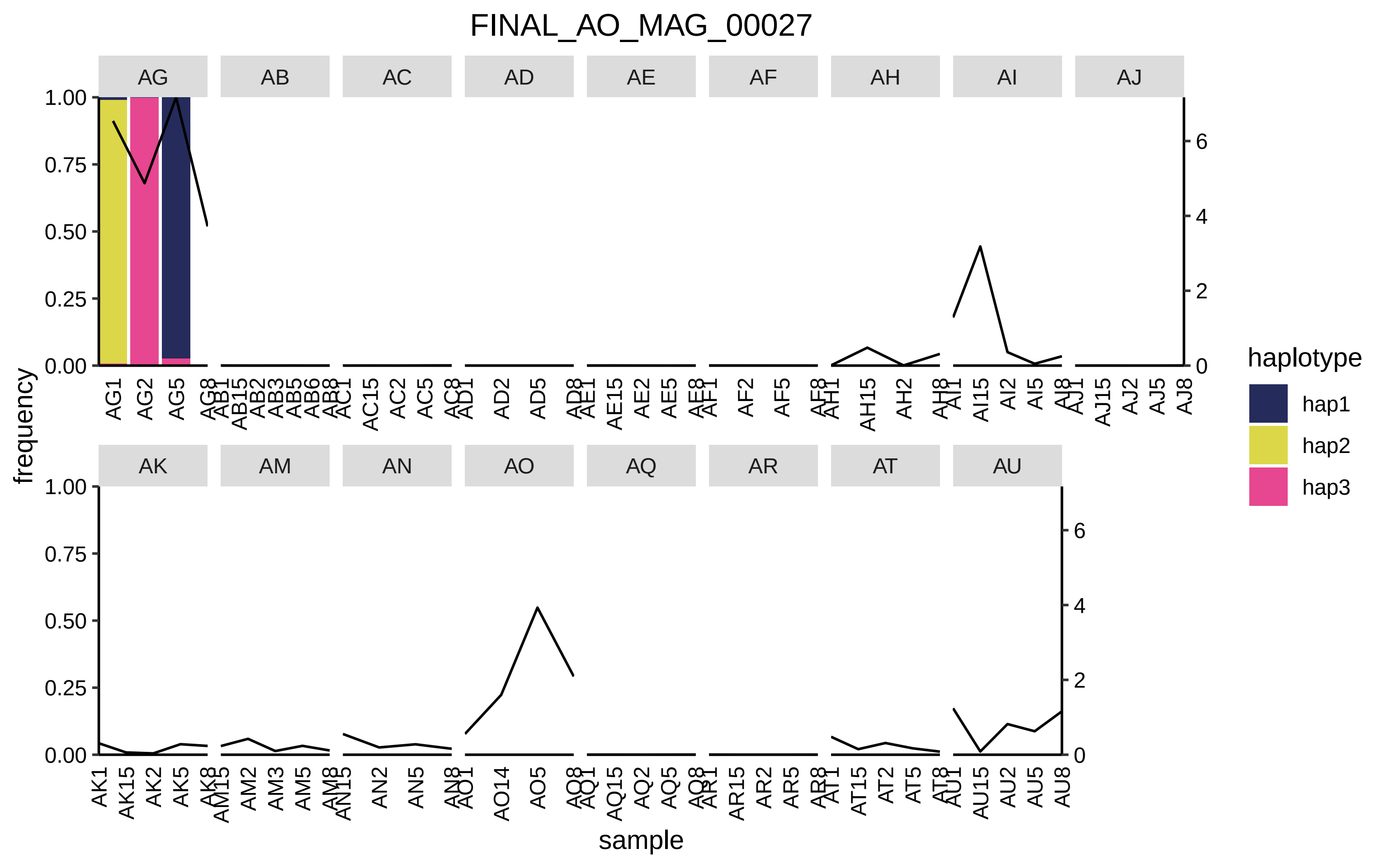
FINAL_AO_MAG_00027
AG
AB
AC
AD
AE
AF
AH
AI
AJ
AK
AM
AN
AO
AQ
AR
AT
AU
frequency
sample
haplotype
hap1
hap2
hap3
1.00
0.75
0.50
0.25
0.00
0
2
4
6

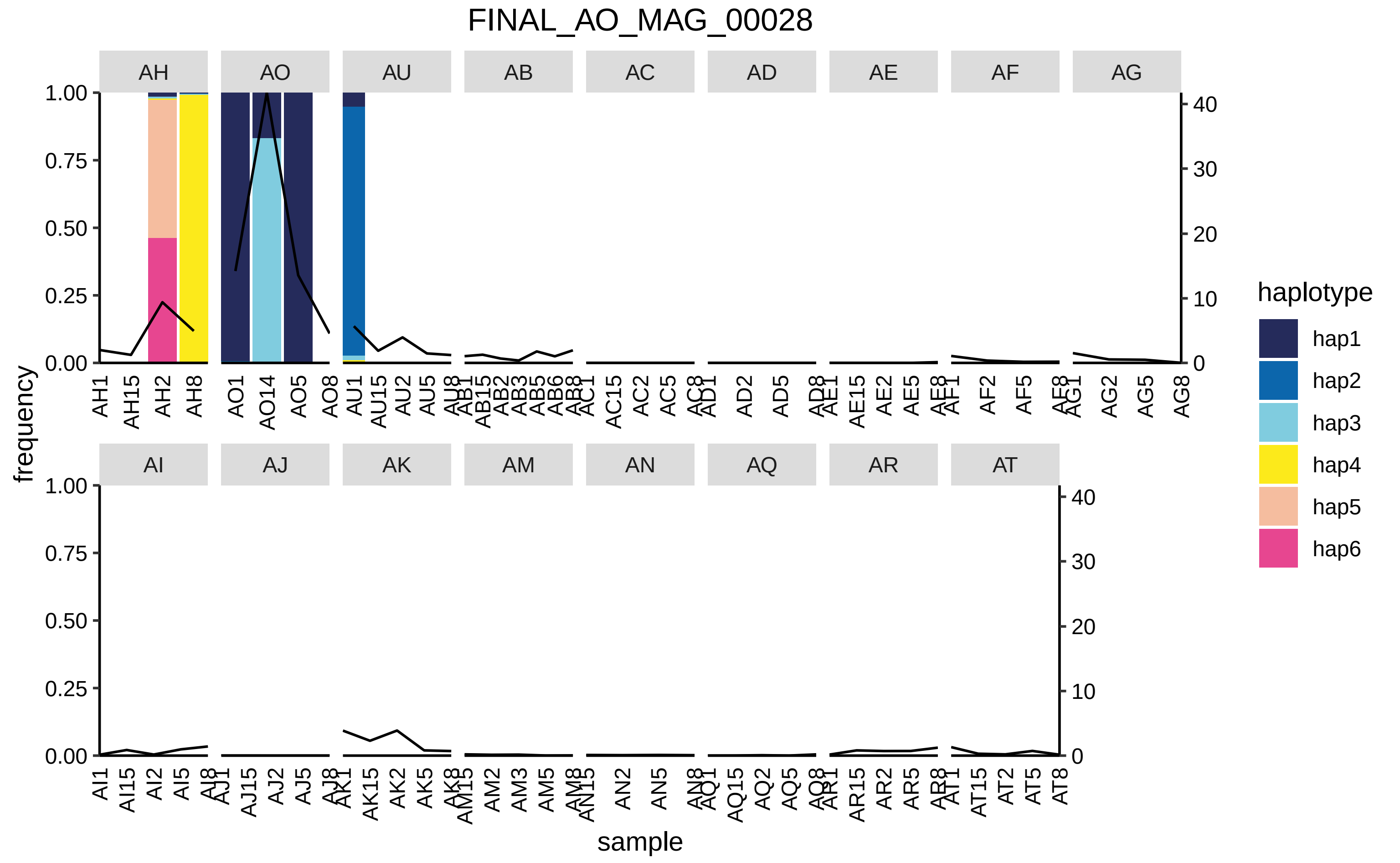
FINAL_AO_MAG_00028
AH
AO
AU
AB
AC
AD
AE
AF
AG
AI
AJ
AK
AM
AN
AQ
AR
AT
frequency
sample
haplotype
hap1
hap2
hap3
hap4
hap5
hap6

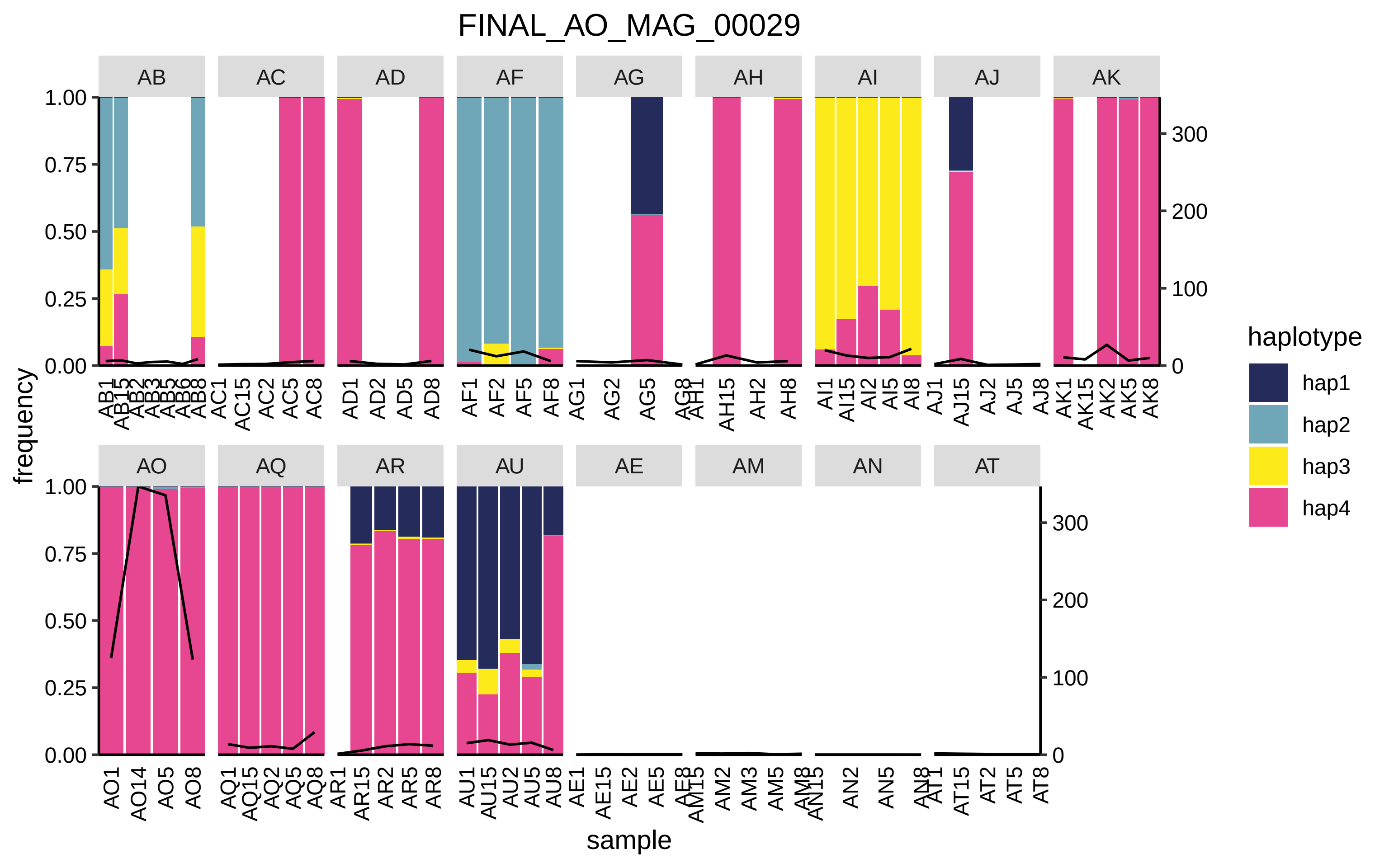
FINAL_AO_MAG_00029
AB
AC
AD
AF
AG
AH
AI
AJ
AK
AO
AQ
AR
AU
AE
AM
AN
AT
frequency
1.00
0.75
0.50
0.25
0.00
300
200
100
0
sample
haplotype
hap1
hap2
hap3
hap4
AC1
AC15
AC2
AC5
AC8
AD1
AD2
AD5
AD8
AF1
AF2
AF5
AF8
AG1
AG2
AG5
AH15
AH2
AH8
AI1
AI15
AI2
AI5
AI8
AJ1
AJ15
AJ2
AJ5
AJ8
AK1
AK15
AK2
AK5
AK8
AO1
AO14
AO5
AO8
AQ1
AQ15
AQ2
AQ5
AQ8
AR1
AR15
AR2
AR5
AR8
AU1
AU15
AU2
AU5
AU8
AE1
AE15
AE2
AE5
AM2
AM3
AM5
AN2
AN5
AT15
AT2
AT5
AT8

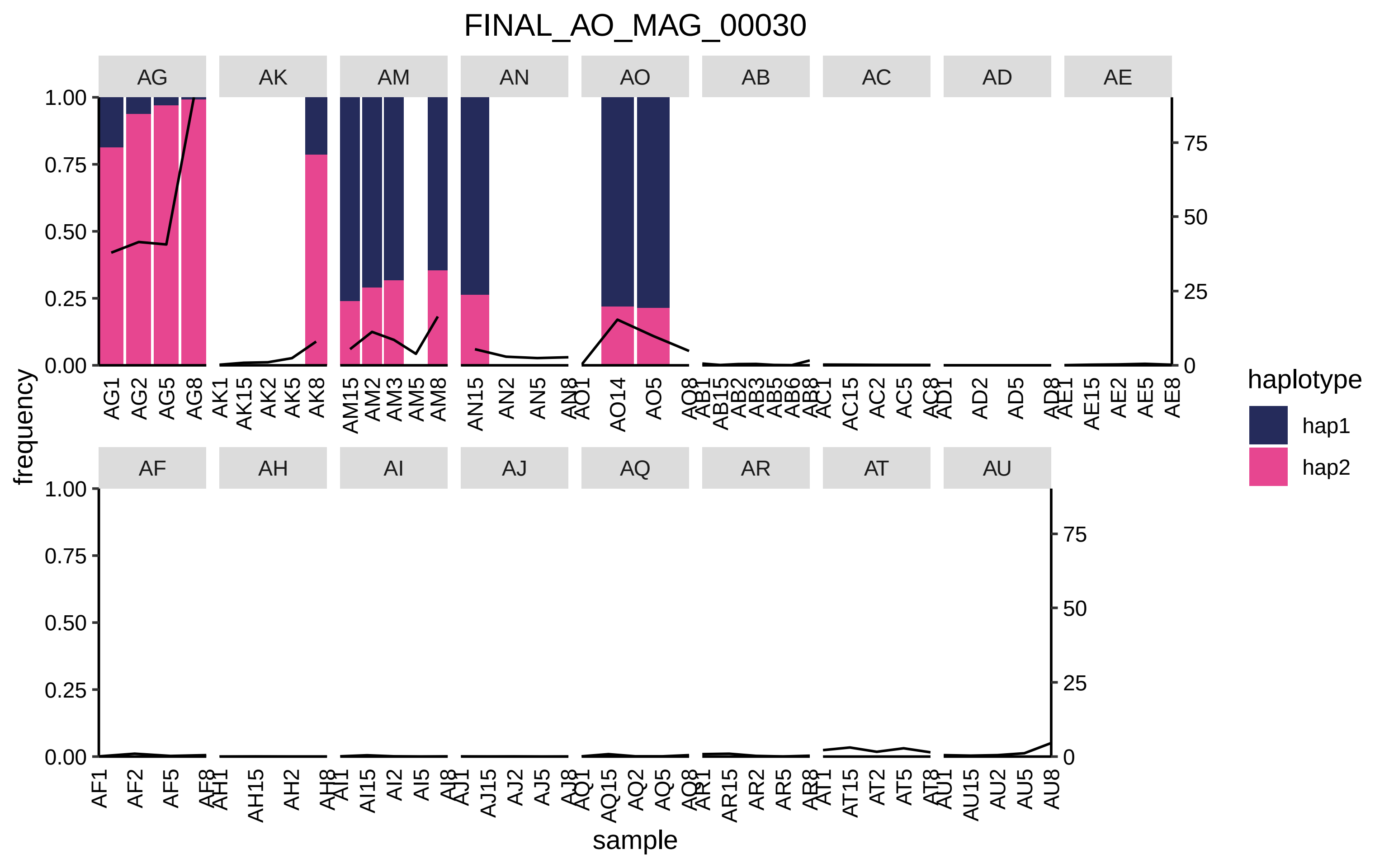

FINAL_AO_MAG_00030
AG
AK
AM
AN
AO
AB
AC
AD
AE
AF
AH
AI
AJ
AQ
AR
AT
AU
frequency
sample
haplotype
hap1
hap2

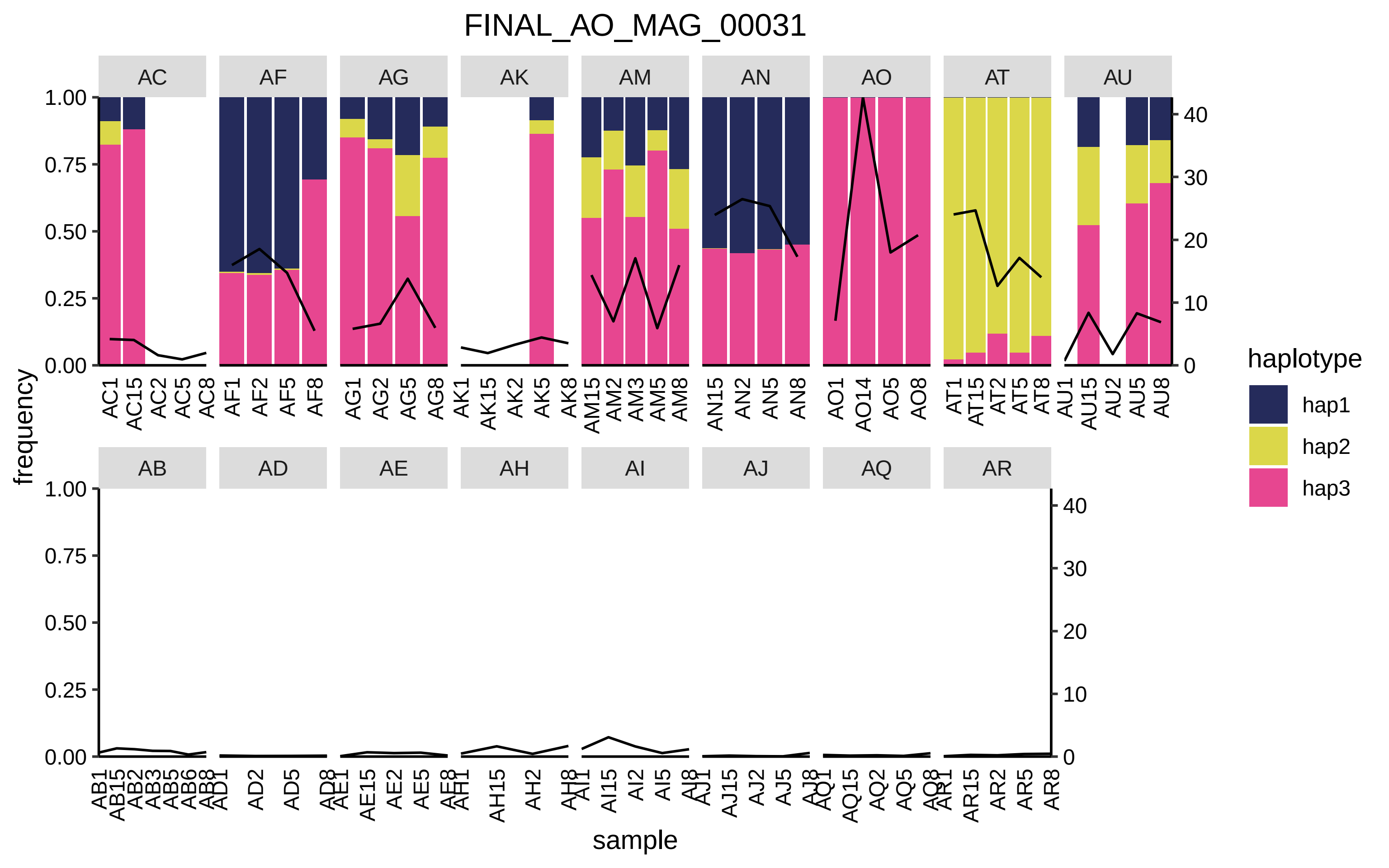
FINAL_AO_MAG_00031
AC
AF
AG
AK
AM
AN
AO
AT
AU
AB
AD
AE
AH
AI
AJ
AQ
AR
frequency
sample
haplotype
hap1
hap2
hap3
1.00
0.75
0.50
0.25
0.00
40
30
20
10
0

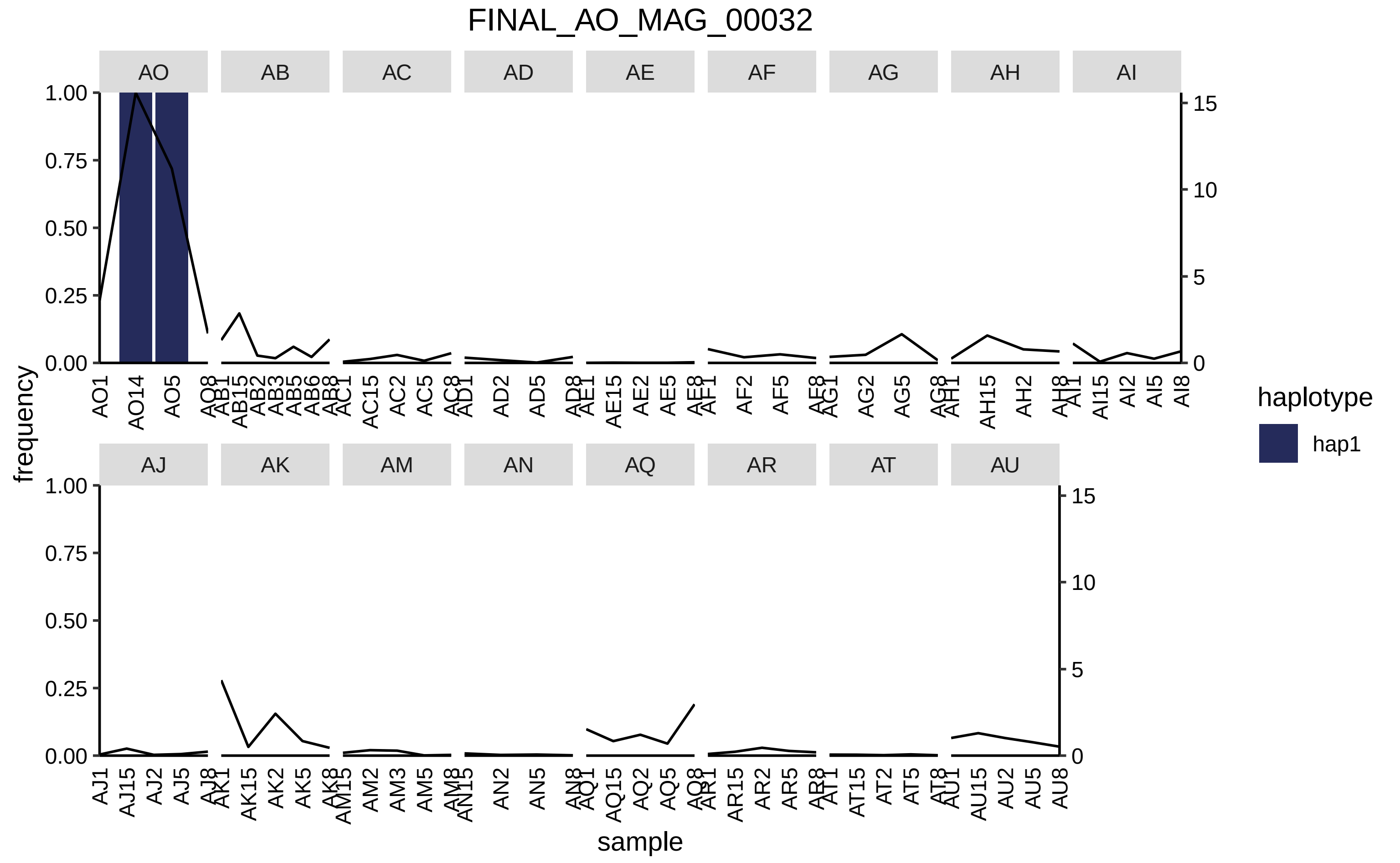

FINAL_AO_MAG_00032
AO
AB
AC
AD
AE
AF
AG
AH
AI
AJ
AK
AM
AN
AQ
AR
AT
AU
frequency
sample
haplotype
hap1

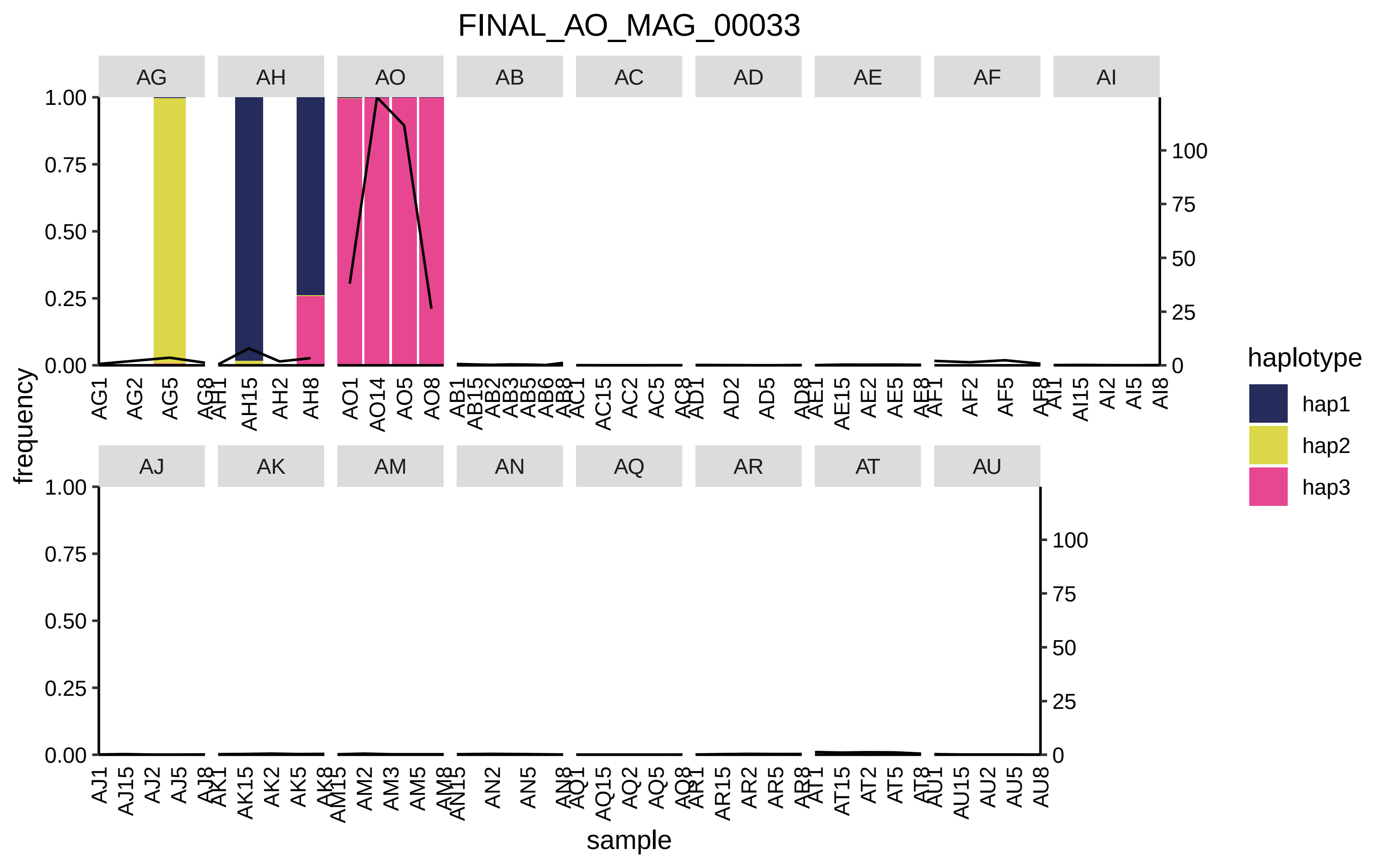
FINAL_AO_MAG_00033
AG
AH
AO
AB
AC
AD
AE
AF
AI
AJ
AK
AM
AN
AQ
AR
AT
AU
frequency
sample
haplotype
hap1
hap2
hap3
1.00
0.75
0.50
0.25
0.00
100
75
50
25
0

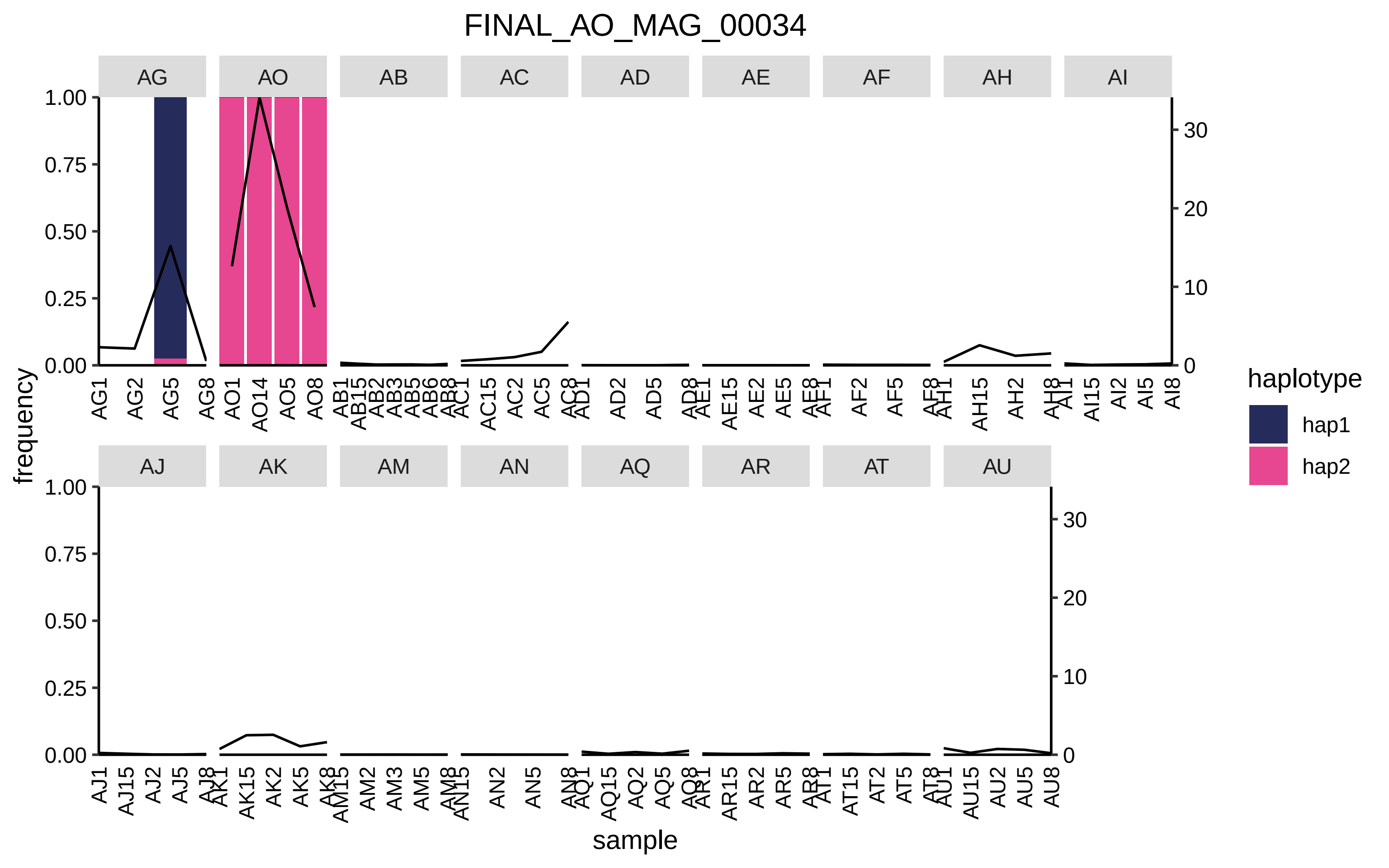
FINAL_AO_MAG_00034
AG
AO
AB
AC
AD
AE
AF
AH
AI
AJ
AK
AM
AN
AQ
AR
AT
AU
frequency
sample
haplotype
hap1
hap2

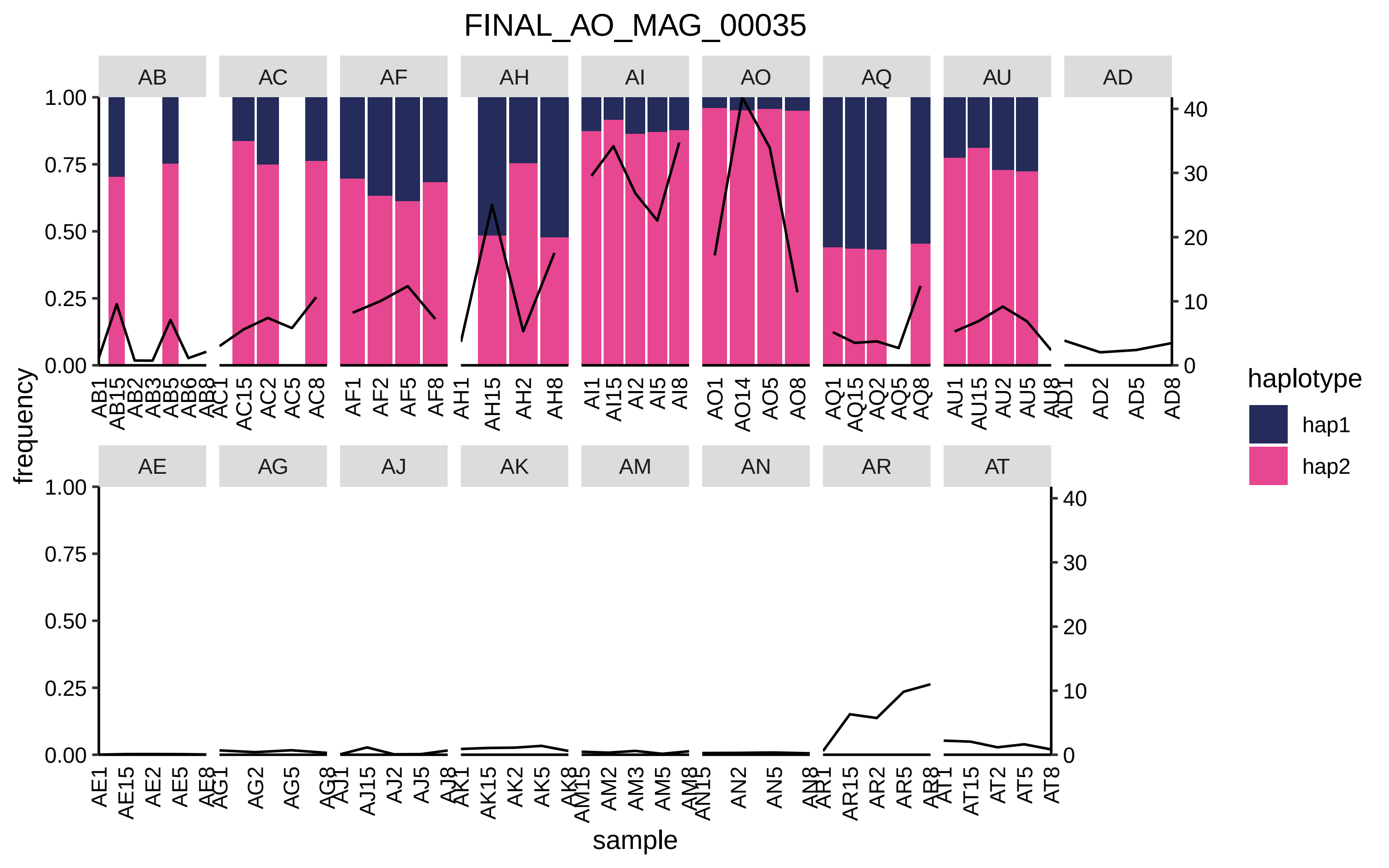

FINAL_AO_MAG_00035
AB
AC
AF
AH
AI
AO
AQ
AU
AD
AE
AG
AJ
AK
AM
AN
AR
AT
frequency
sample
haplotype
hap1
hap2

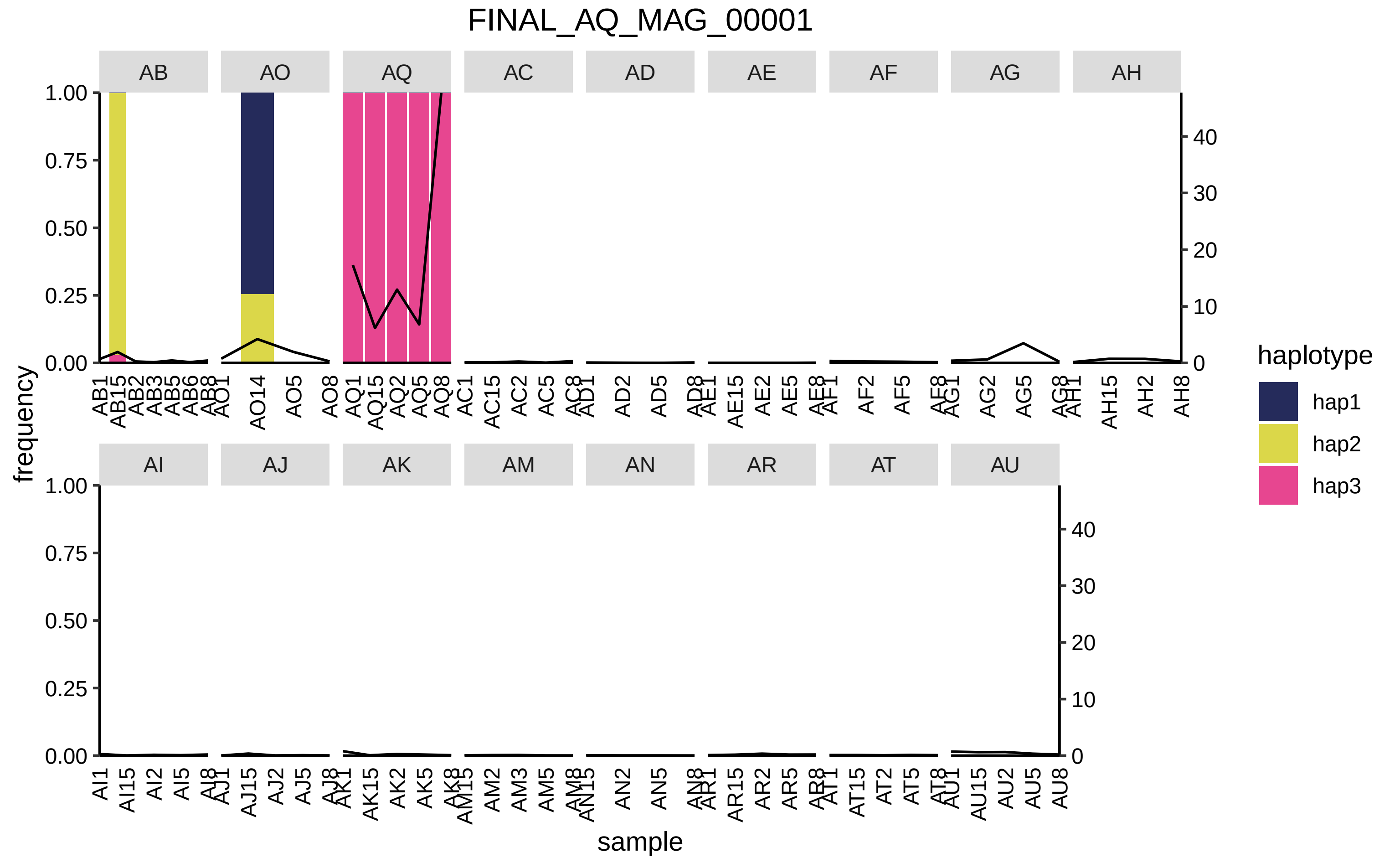

FINAL_AQ_MAG_00001
AB
AO
AQ
AC
AD
AE
AF
AG
AH
AI
AJ
AK
AM
AN
AR
AT
AU
frequency
sample
haplotype
hap1
hap2
hap3

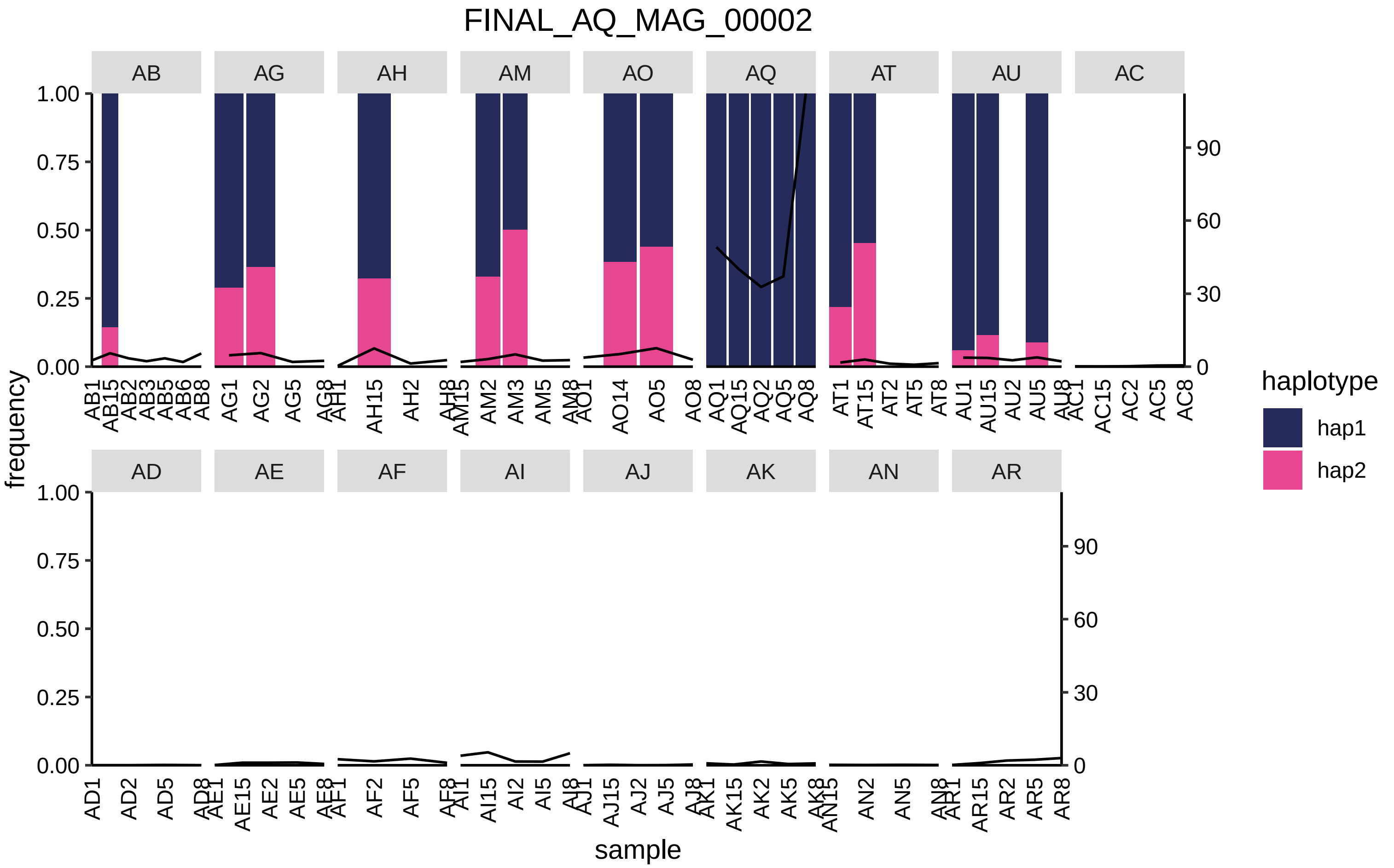

FINAL_AQ_MAG_00002
AB
AG
AH
AM
AO
AQ
AT
AU
AC
AD
AE
AF
AI
AJ
AK
AN
AR
frequency
1.00
0.75
0.50
0.25
0.00
0
30
60
90
sample
haplotype
hap1
hap2
AB1 AB15 AB2 AB3 AB5 AB6 AB8
AG1 AG2 AG5 AG8
AH1 AH15 AH2 AH8
AM15 AM2 AM3 AM5 AM8
AO1 AO14 AO5 AO8
AQ1 AQ15 AQ2 AQ5 AQ8
AT1 AT15 AT2 AT5 AT8
AU1 AU15 AU2 AU5 AU8
AC1 AC15 AC2 AC5 AC8
AD1 AD2 AD5 AD8
AE1 AE15 AE2 AE5 AE8
AF1 AF2 AF5 AF8
AI1 AI15 AI2 AI5 AI8
AJ1 AJ15 AJ2 AJ5 AJ8
AK1 AK15 AK2 AK5 AK8
AN15 AN2 AN5 AN8
AR1 AR15 AR2 AR5 AR8

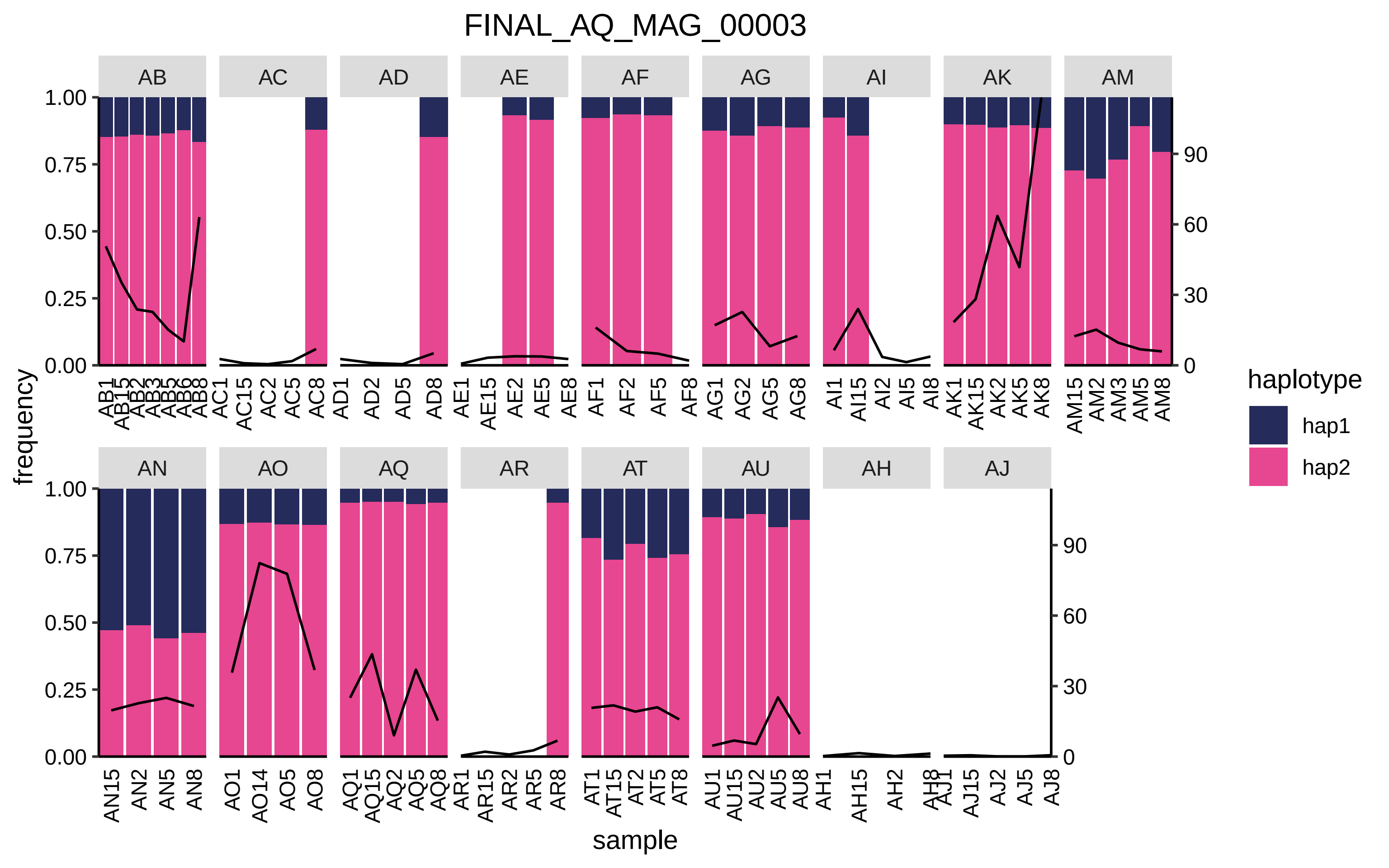

FINAL_AQ_MAG_00003
AB
AC
AD
AE
AF
AG
AI
AK
AM
AN
AO
AQ
AR
AT
AU
AH
AJ
frequency
sample
haplotype
hap1
hap2
0.00
0.25
0.50
0.75
1.00
0
30
60
90

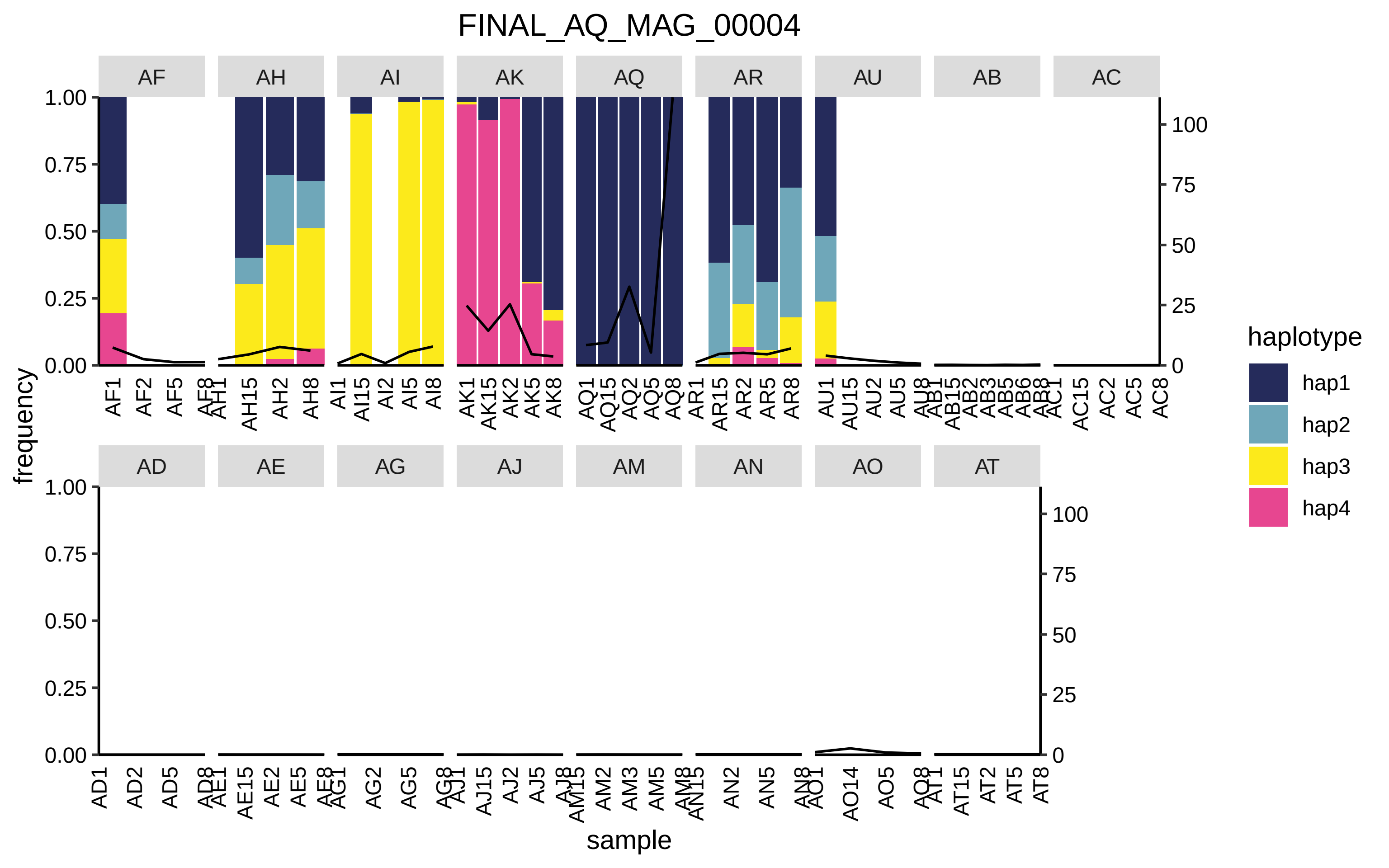
FINAL_AQ_MAG_00004
AF
AH
AI
AK
AQ
AR
AU
AB
AC
AD
AE
AG
AJ
AM
AN
AO
AT
frequency
sample
haplotype
hap1
hap2
hap3
hap4

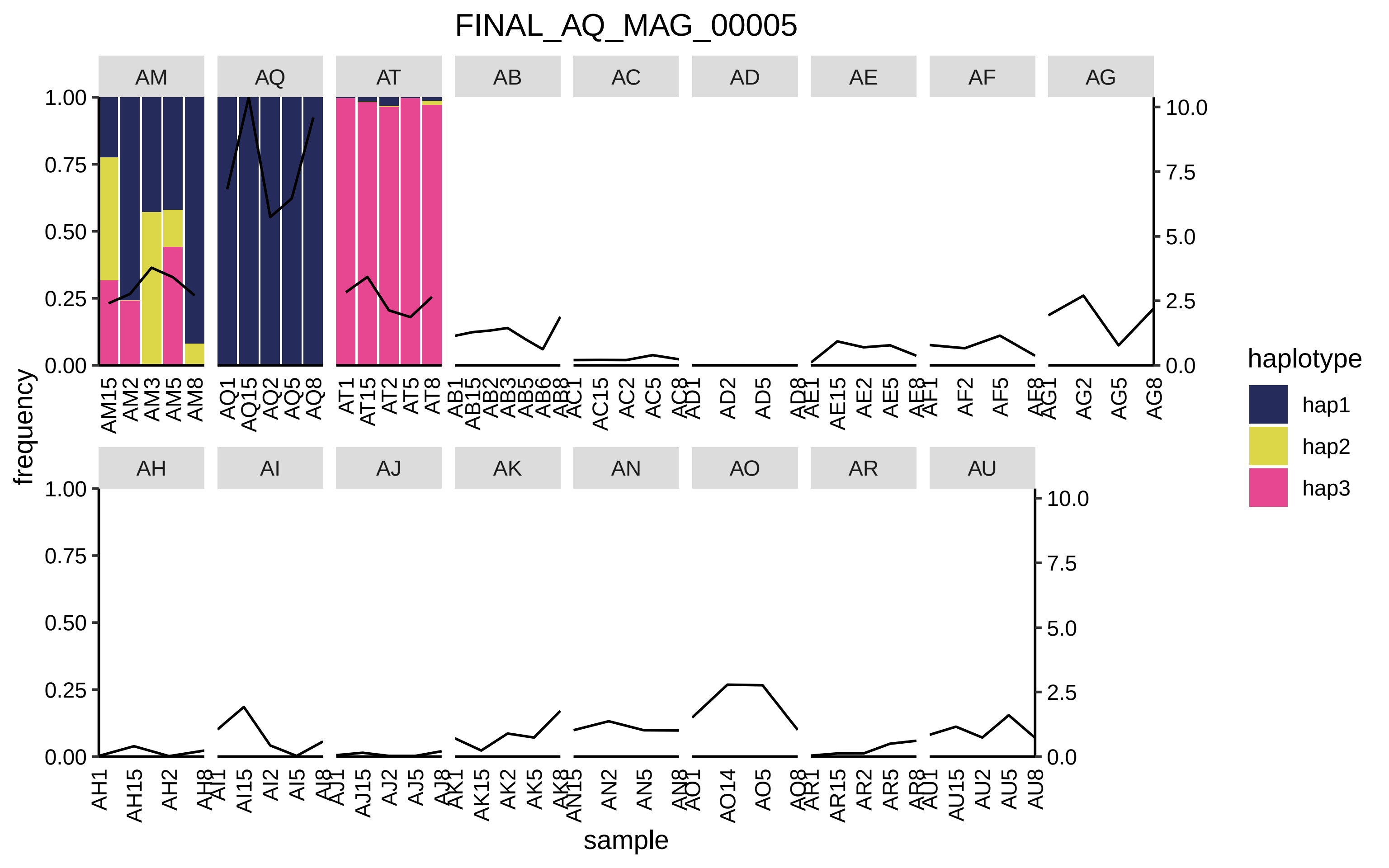

FINAL_AQ_MAG_00005
AM
AQ
AT
AB
AC
AD
AE
AF
AG
AH
AI
AJ
AK
AN
AO
AR
AU
frequency
sample
haplotype
hap1
hap2
hap3

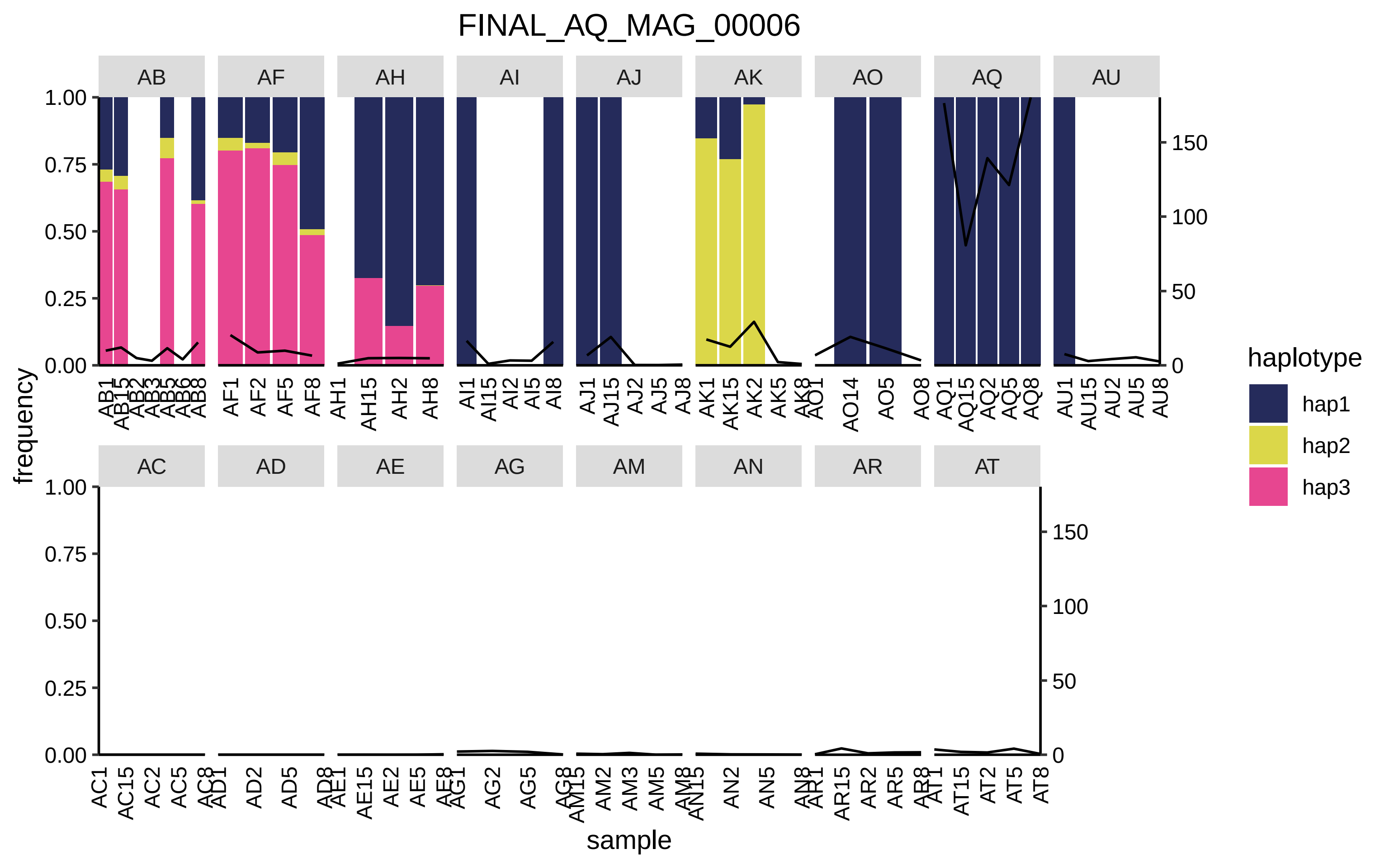

FINAL_AQ_MAG_00006
AB
AF
AH
AI
AJ
AK
AO
AQ
AU
AC
AD
AE
AG
AM
AN
AR
AT
frequency
sample
1.00
0.75
0.50
0.25
0.00
150
100
50
0
AB1 AB15 AB2 AB3 AB5 AB6 AB8
AF1 AF2 AF5 AF8
AH1 AH15 AH2 AH8
AI1 AI15 AI2 AI5 AI8
AJ1 AJ15 AJ2 AJ5 AJ8
AK1 AK15 AK2 AK5 AK8
AO1 AO14 AO5 AO8
AQ1 AQ15 AQ2 AQ5 AQ8
AU1 AU15 AU2 AU5 AU8
AC1 AC15 AC2 AC5 AC8
AD1 AD2 AD5 AD8
AE1 AE15 AE2 AE5 AE8
AG1 AG2 AG5 AG8
AM15 AM2 AM3 AM5 AM8
AN15 AN2 AN5 AN8
AR1 AR15 AR2 AR5 AR8
AT1 AT15 AT2 AT5 AT8
haplotype
hap1
hap2
hap3

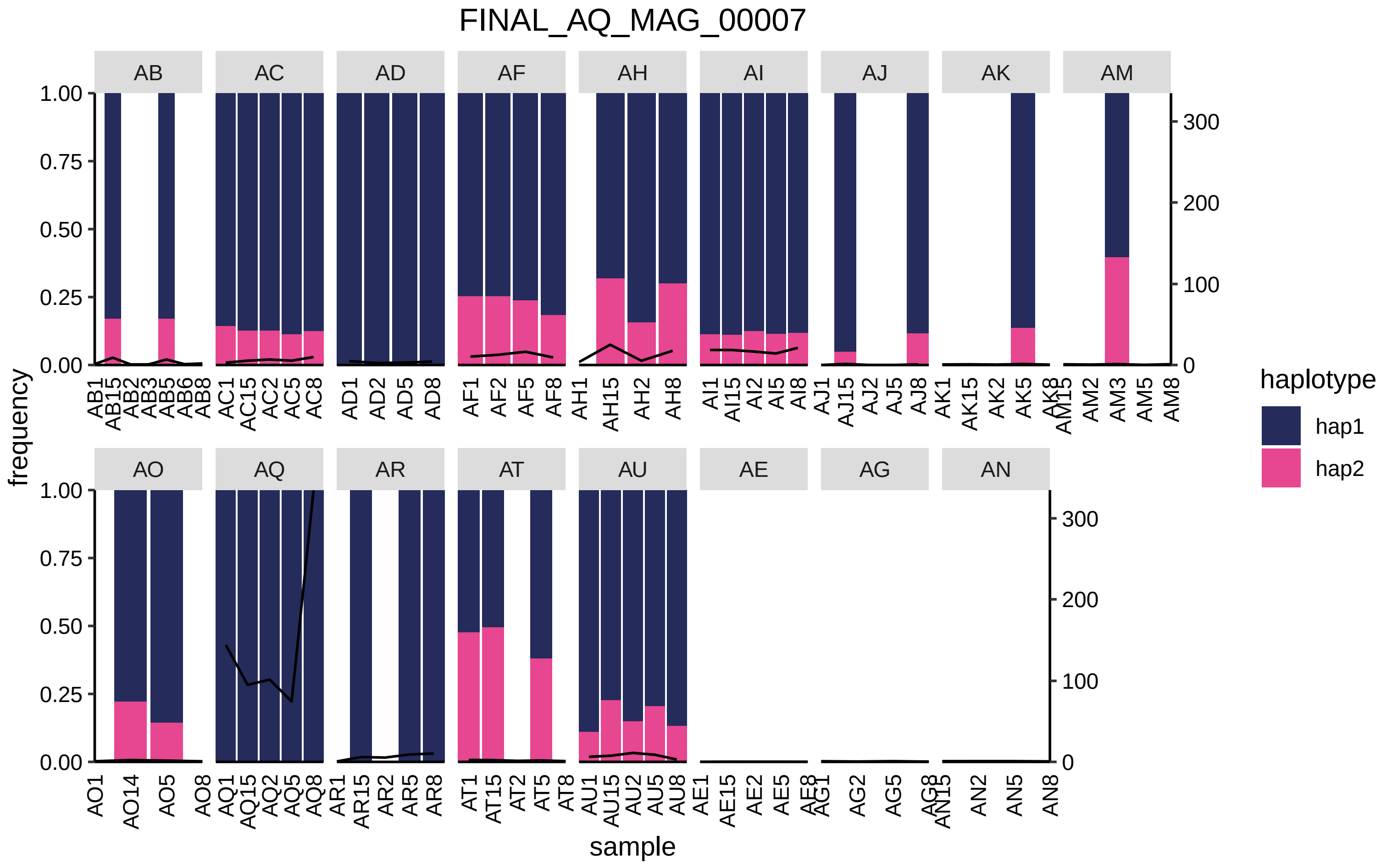
FINAL_AQ_MAG_00007
AB
AC
AD
AF
AH
AI
AJ
AK
AM
AO
AQ
AR
AT
AU
AE
AG
AN
frequency
sample
haplotype
hap1
hap2
1.00
0.75
0.50
0.25
0.00
300
200
100
0
AB1 AB15 AB2 AB3 AB5 AB6 AB8
AC1 AC15 AC2 AC5 AC8
AD1 AD2 AD5 AD8
AF1 AF2 AF5 AF8
AH1 AH15 AH2 AH8
AI1 AI15 AI2 AI5 AI8
AJ1 AJ15 AJ2 AJ5 AJ8
AK1 AK15 AK2 AK5 AK8
AM15 AM2 AM3 AM5 AM8
AO1 AO14 AO5 AO8
AQ1 AQ15 AQ2 AQ5 AQ8
AR1 AR15 AR2 AR5 AR8
AT1 AT15 AT2 AT5 AT8
AU1 AU15 AU2 AU5 AU8
AE1 AE15 AE2 AE5 AE8
AG1 AG2 AG5 AG8
AN15 AN2 AN5 AN8

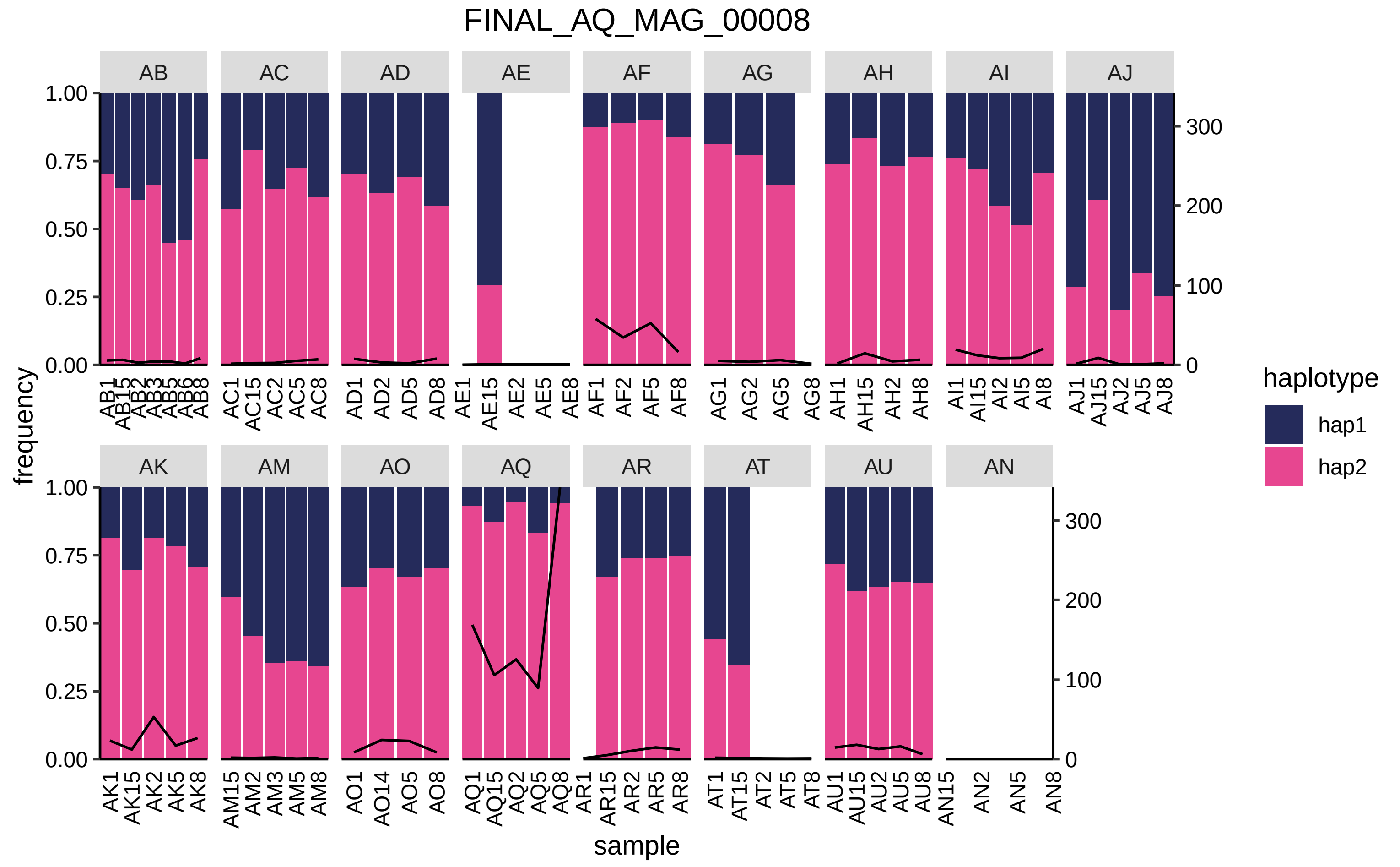
FINAL_AQ_MAG_00008
frequency
sample
haplotype
hap1
hap2
AB
AC
AD
AE
AF
AG
AH
AI
AJ
AK
AM
AO
AQ
AR
AT
AU
AN

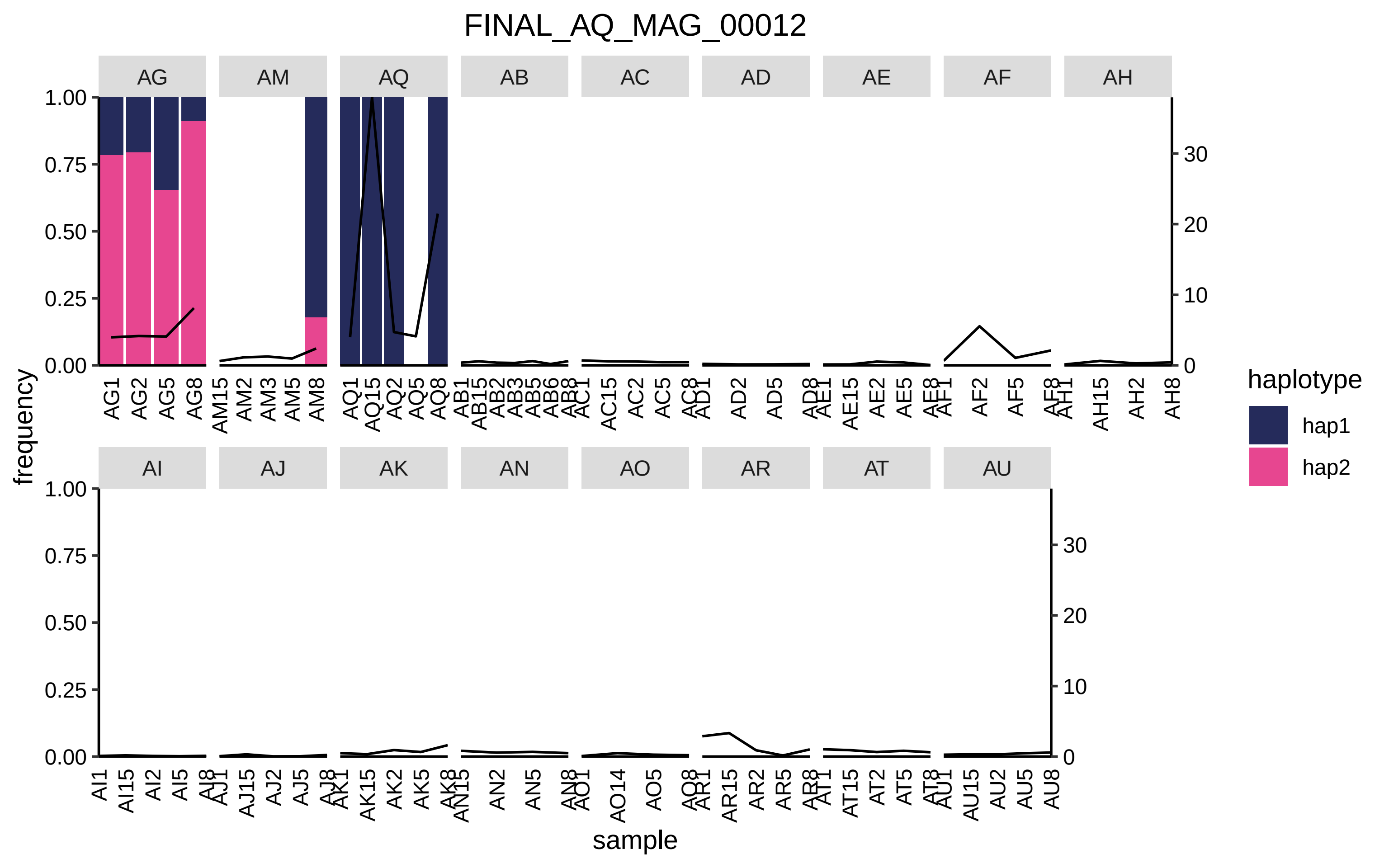

FINAL_AQ_MAG_00012
AG
AM
AQ
AB
AC
AD
AE
AF
AH
AI
AJ
AK
AN
AO
AR
AT
AU
1.00
0.75
0.50
0.25
0.00
0
10
20
30
frequency
sample
haplotype
hap1
hap2

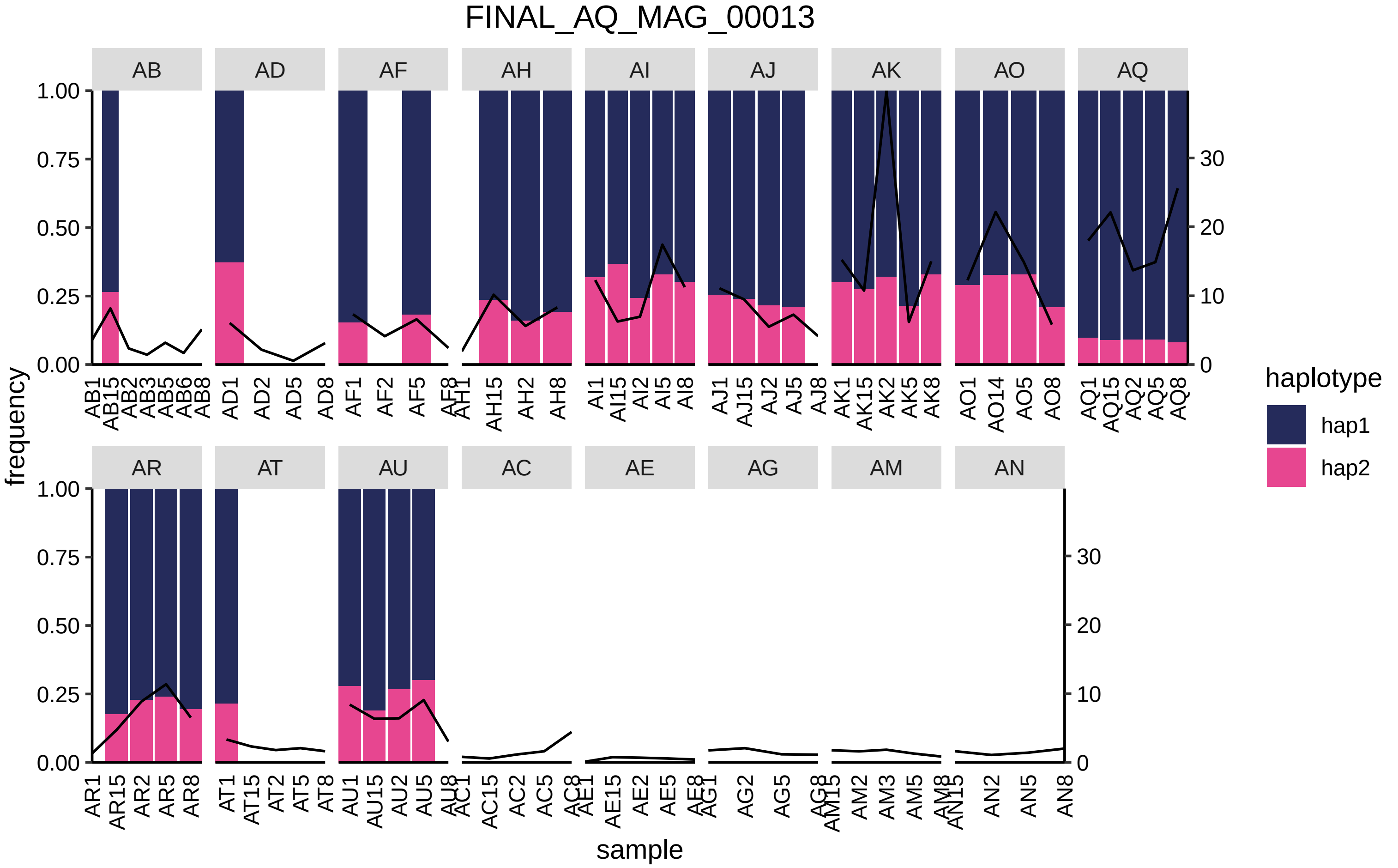

FINAL_AQ_MAG_00013
AB
AD
AF
AH
AI
AJ
AK
AO
AQ
AR
AT
AU
AC
AE
AG
AM
AN
frequency
sample
haplotype
hap1
hap2

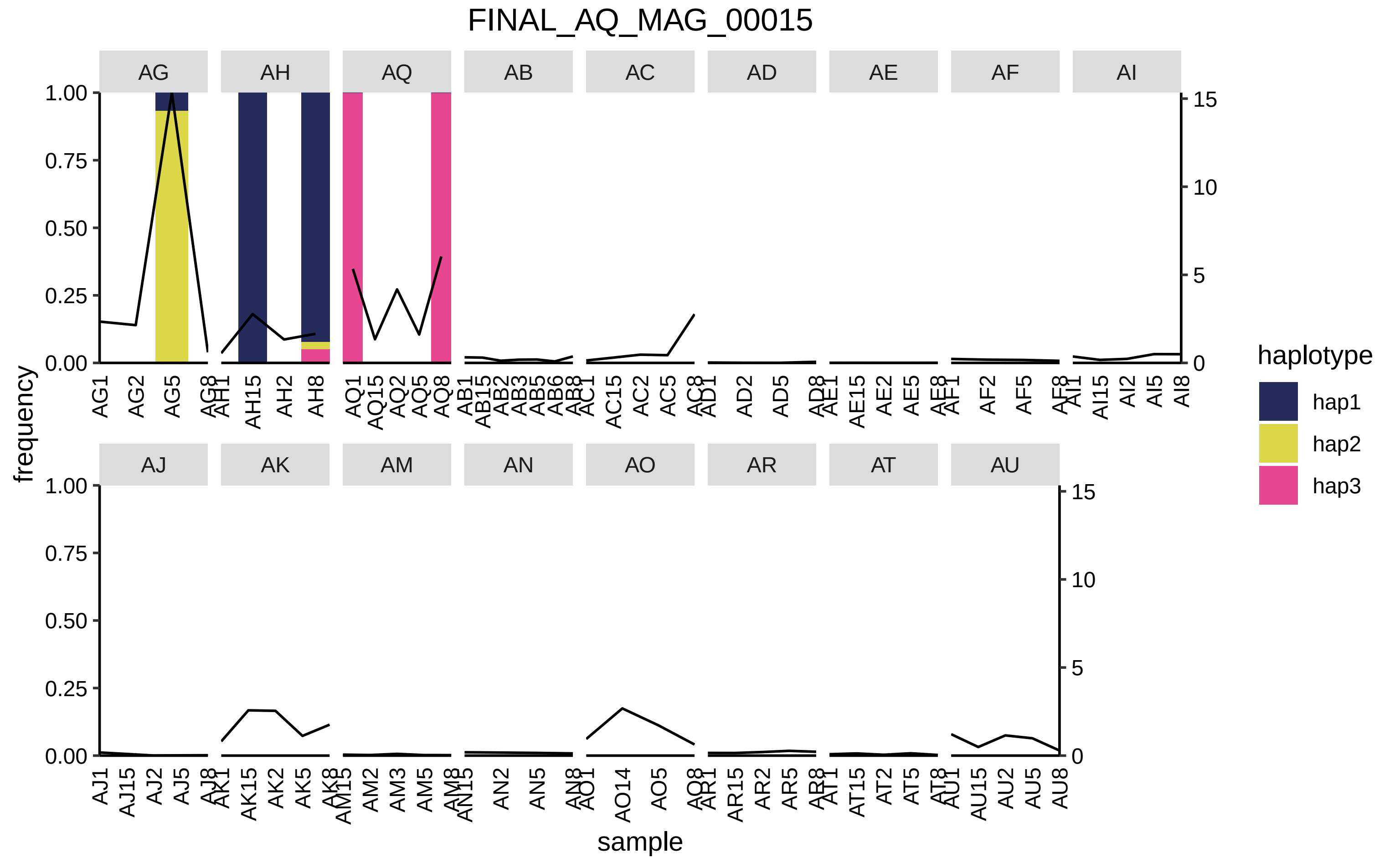

FINAL_AQ_MAG_00015
AG
AH
AQ
AB
AC
AD
AE
AF
AI
AJ
AK
AM
AN
AO
AR
AT
AU
frequency
sample
haplotype
hap1
hap2
hap3

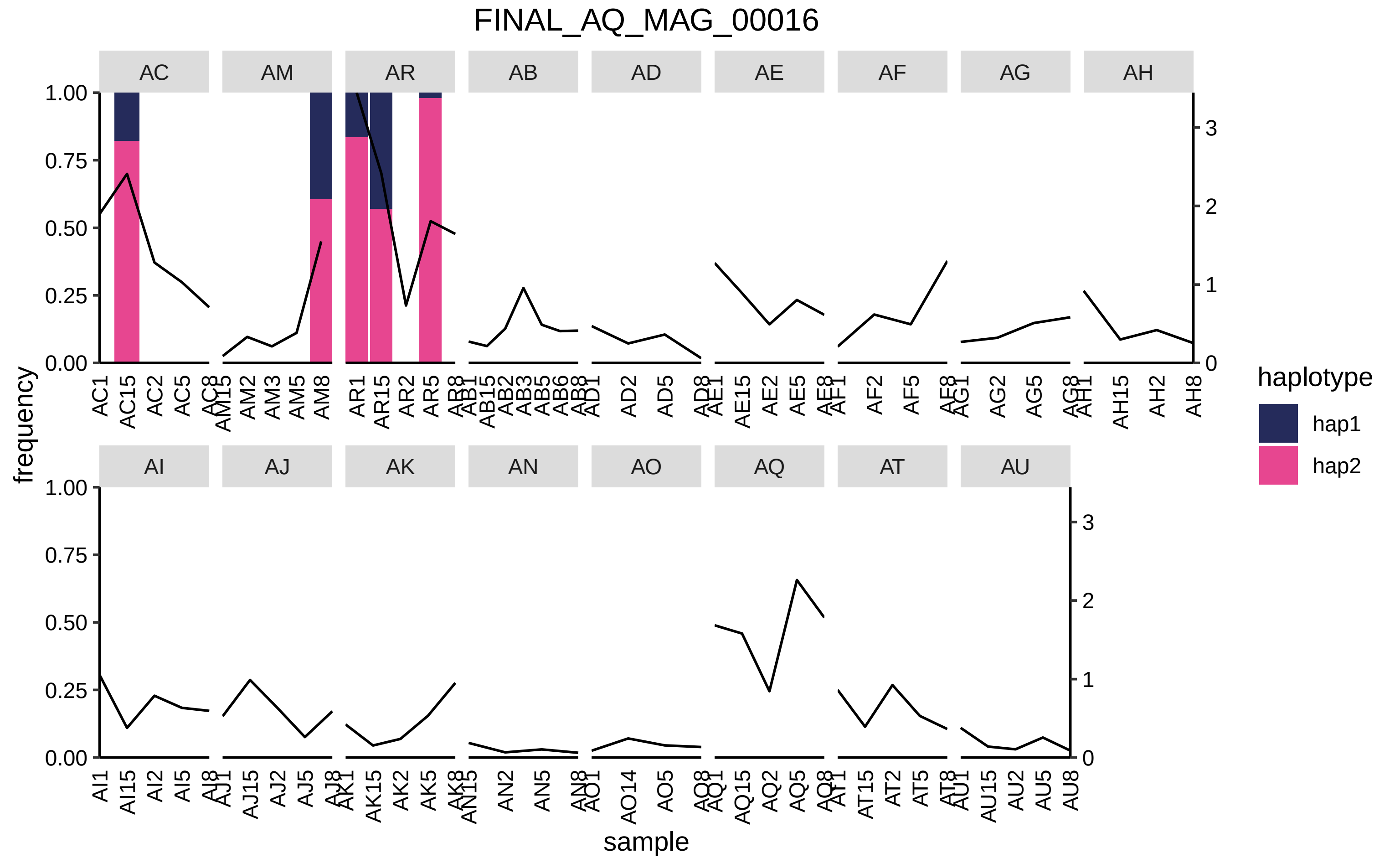
FINAL_AQ_MAG_00016
AC
AM
AR
AB
AD
AE
AF
AG
AH
AI
AJ
AK
AN
AO
AQ
AT
AU
frequency
sample
haplotype
hap1
hap2

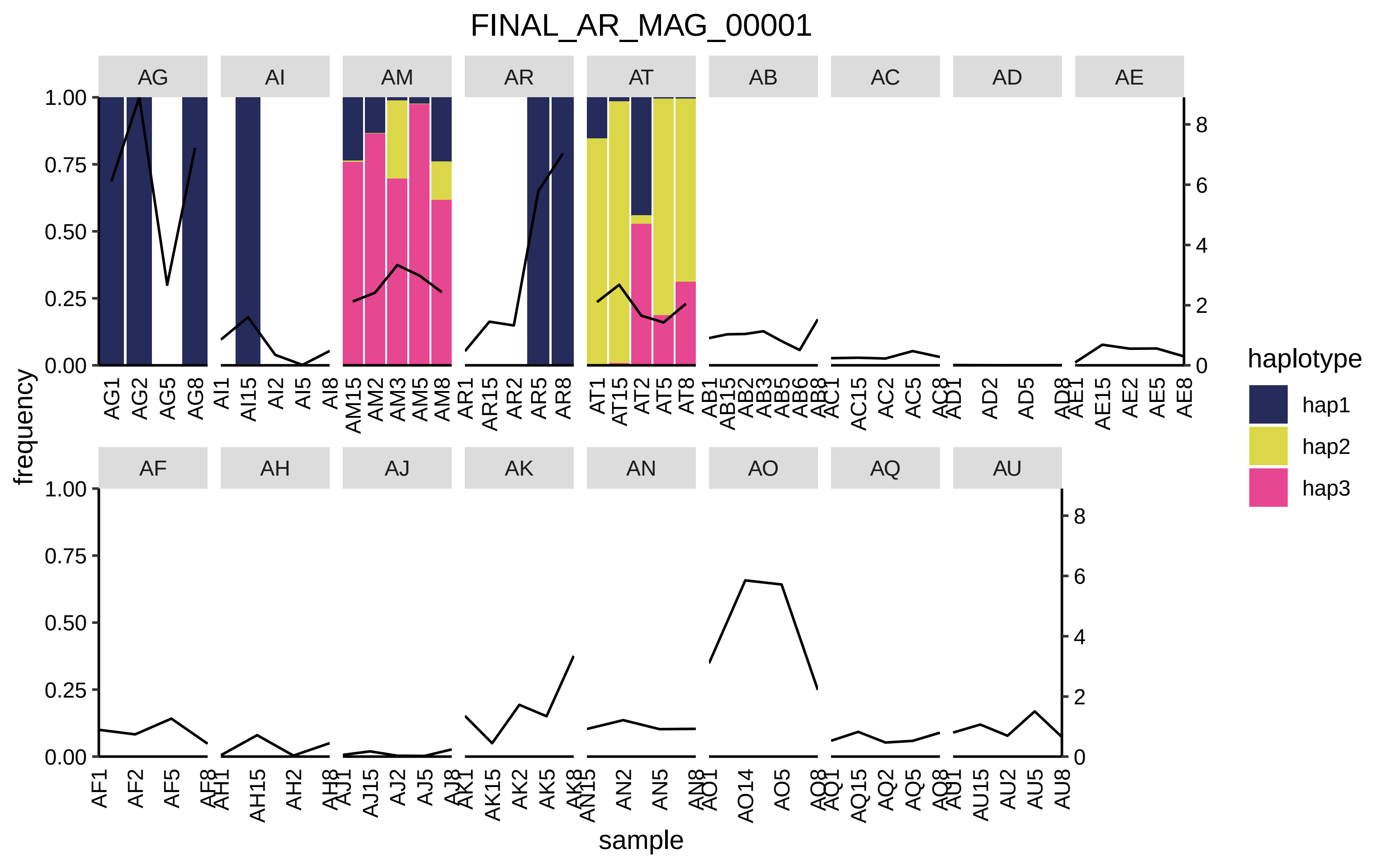

FINAL_AR_MAG_00001
AG
AI
AM
AR
AT
AB
AC
AD
AE
AF
AH
AJ
AK
AN
AO
AQ
AU
frequency
sample
1.00
0.75
0.50
0.25
0.00
8
6
4
2
0
haplotype
hap1
hap2
hap3
AG1 AG2 AG5 AG8
AI1 AI15 AI2 AI5 AI8
AM15 AM2 AM3 AM5 AM8
AR1 AR15 AR2 AR5 AR8
AT1 AT15 AT2 AT5 AT8
AB1 AB15 AB2 AB3 AB5 AB6 AB8
AC1 AC15 AC2 AC5 AC8
AD1 AD2 AD5 AD8
AE1 AE15 AE2 AE5 AE8
AF1 AF2 AF5 AF8
AH1 AH15 AH2 AH8
AJ1 AJ15 AJ2 AJ5 AJ8
AK1 AK15 AK2 AK5 AK8
AN15 AN2 AN5 AN8
AO1 AO14 AO5 AO8
AQ1 AQ15 AQ2 AQ5 AQ8
AU1 AU15 AU2 AU5 AU8

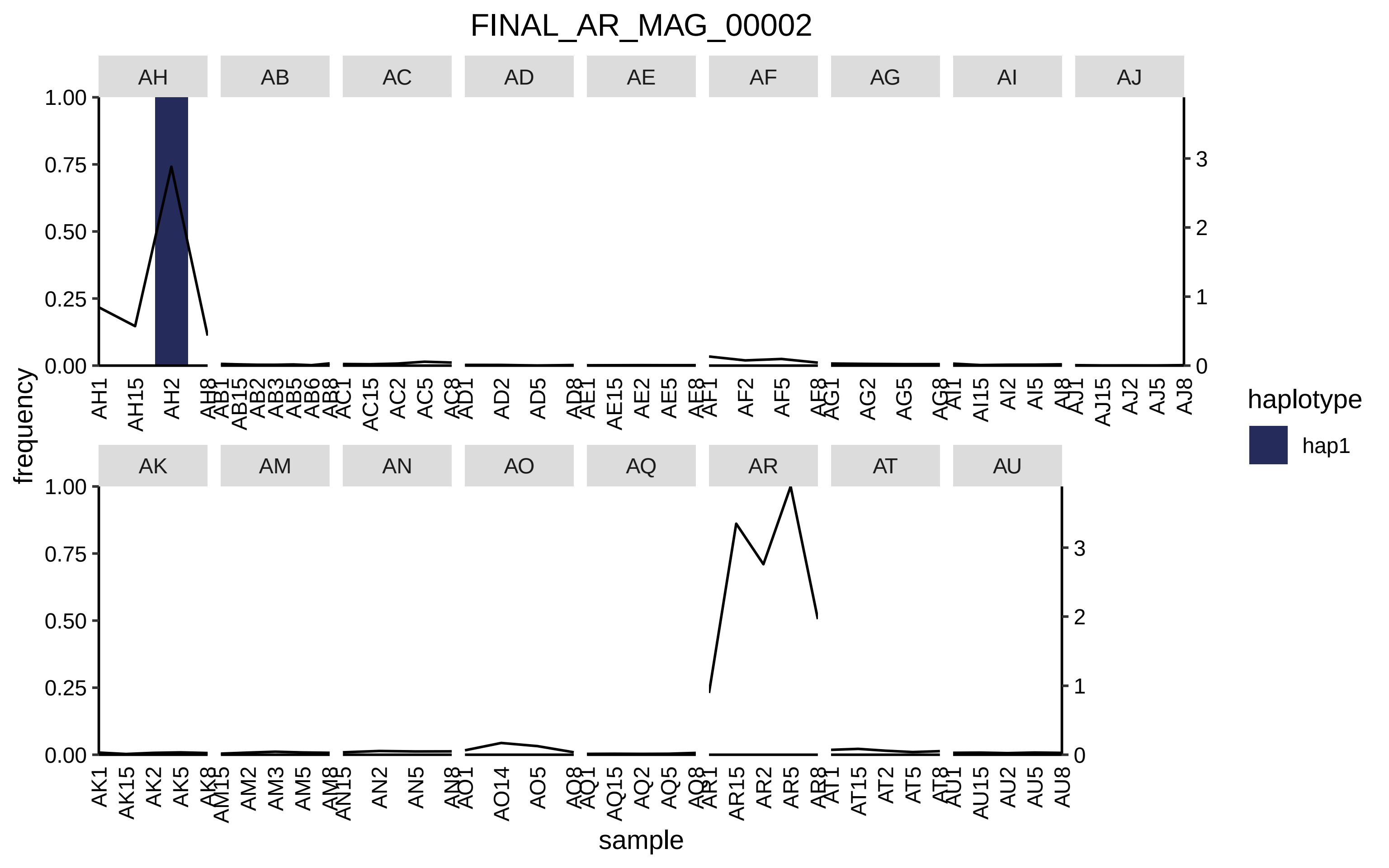

FINAL_AR_MAG_00002
AH
AB
AC
AD
AE
AF
AG
AI
AJ
AK
AM
AN
AO
AQ
AR
AT
AU
frequency
sample
haplotype
hap1
1.00
0.75
0.50
0.25
0.00
0
1
2
3

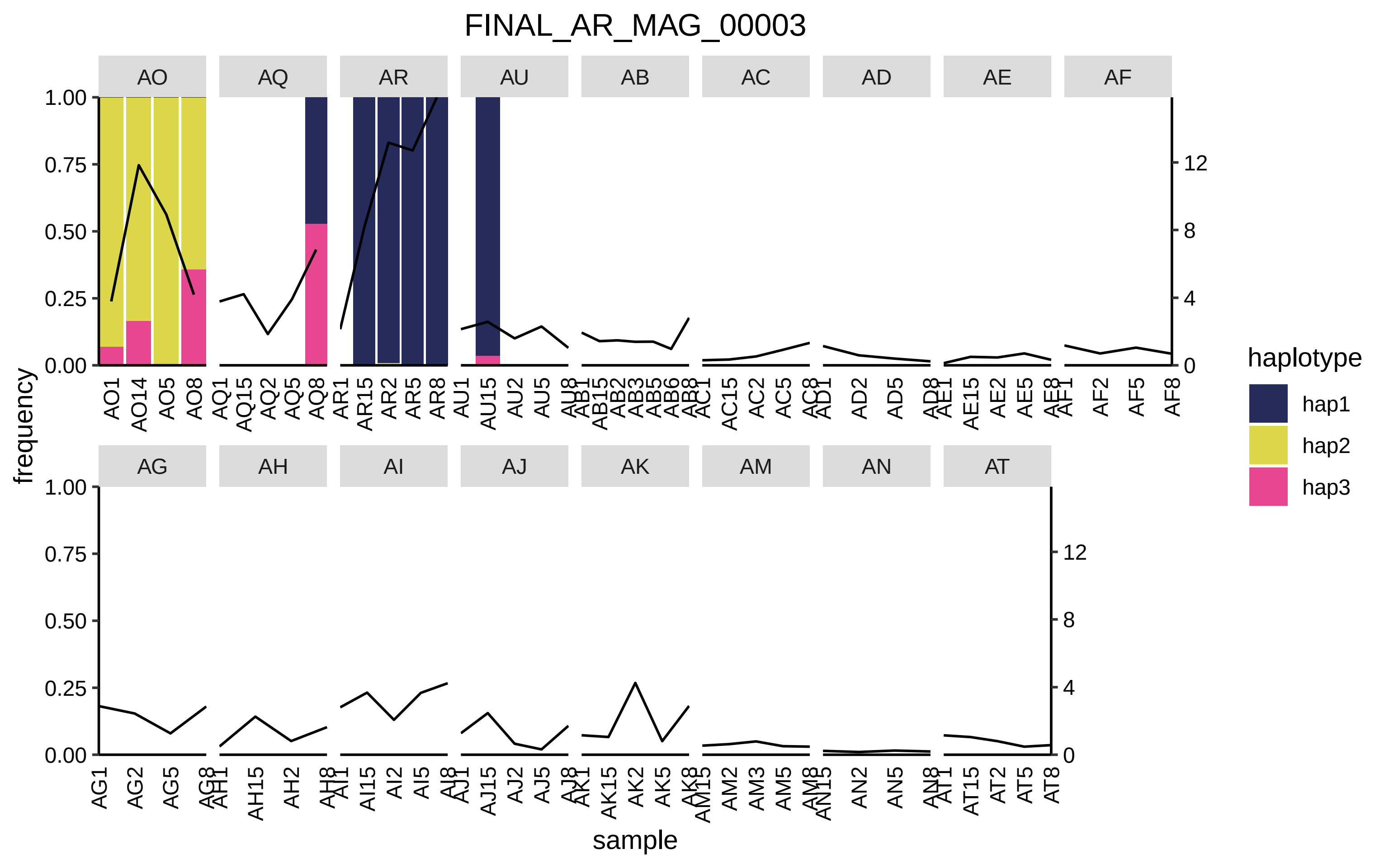

FINAL_AR_MAG_00003
AO
AQ
AR
AU
AB
AC
AD
AE
AF
AG
AH
AI
AJ
AK
AM
AN
AT
frequency
sample
haplotype
hap1
hap2
hap3

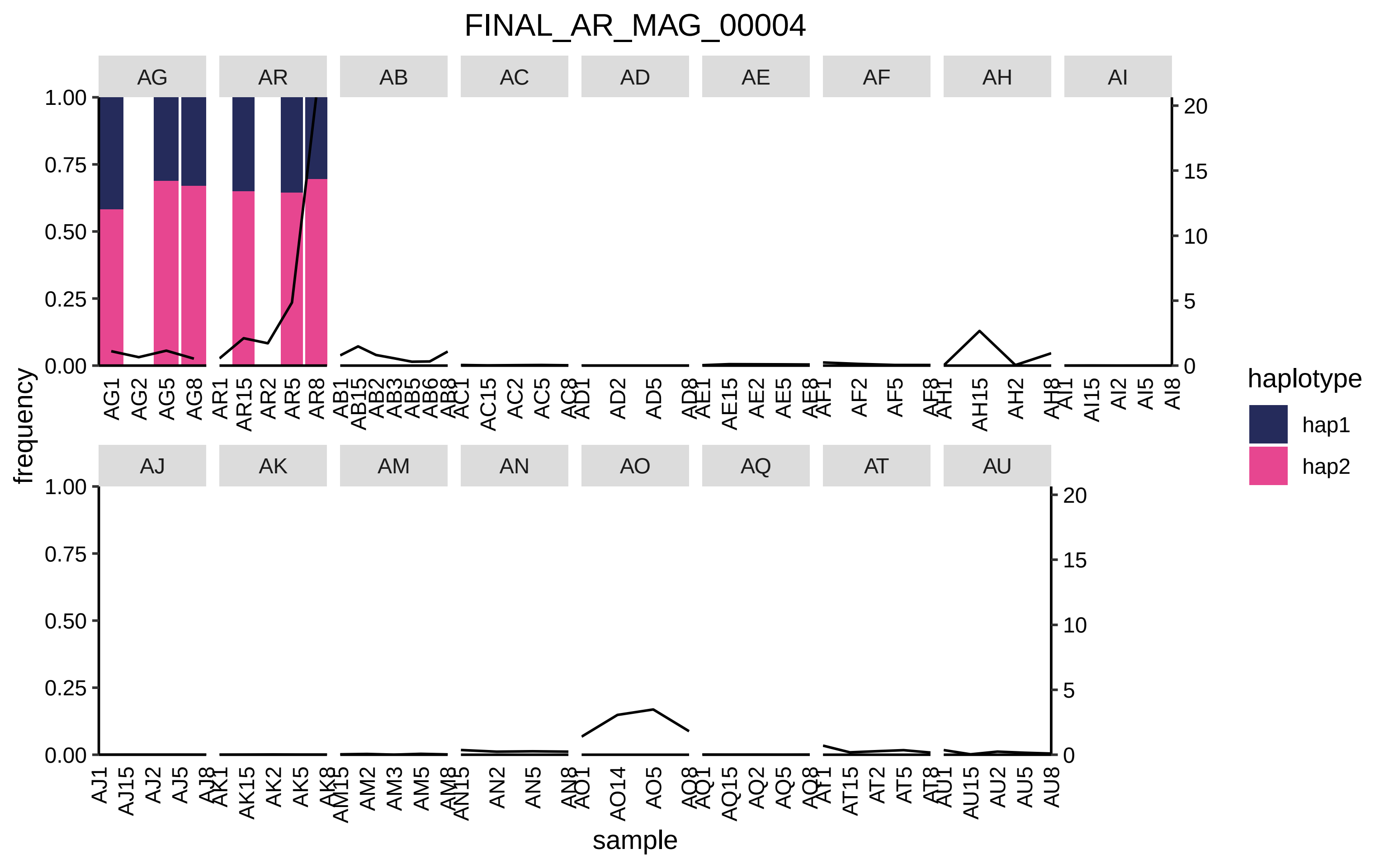
FINAL_AR_MAG_00004
AG
AR
AB
AC
AD
AE
AF
AH
AI
AJ
AK
AM
AN
AO
AQ
AT
AU
frequency
sample
haplotype
hap1
hap2

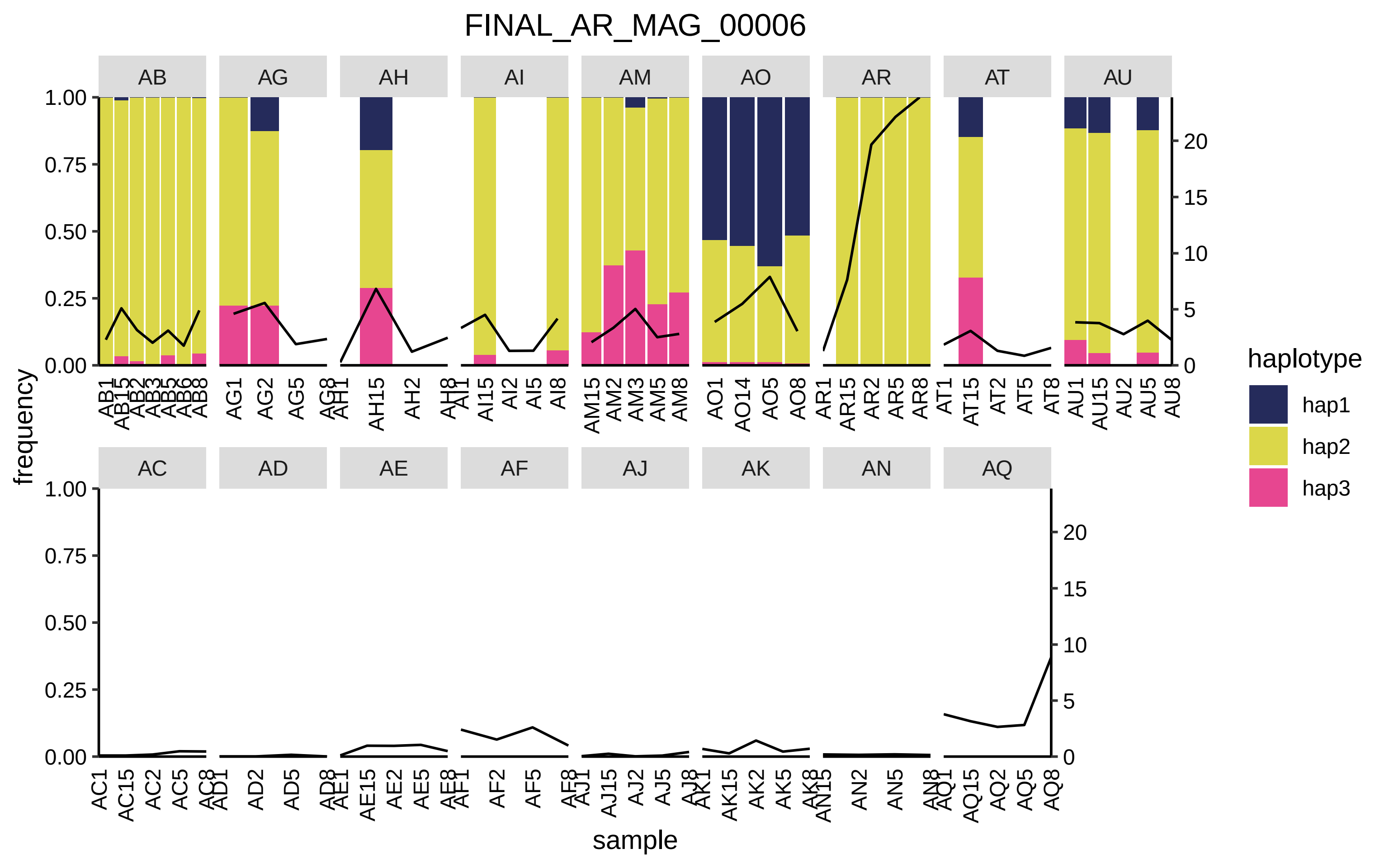

FINAL_AR_MAG_00006
AB
AG
AH
AI
AM
AO
AR
AT
AU
AC
AD
AE
AF
AJ
AK
AN
AQ
frequency
sample
haplotype
hap1
hap2
hap3
1.00
0.75
0.50
0.25
0.00
0
5
10
15
20

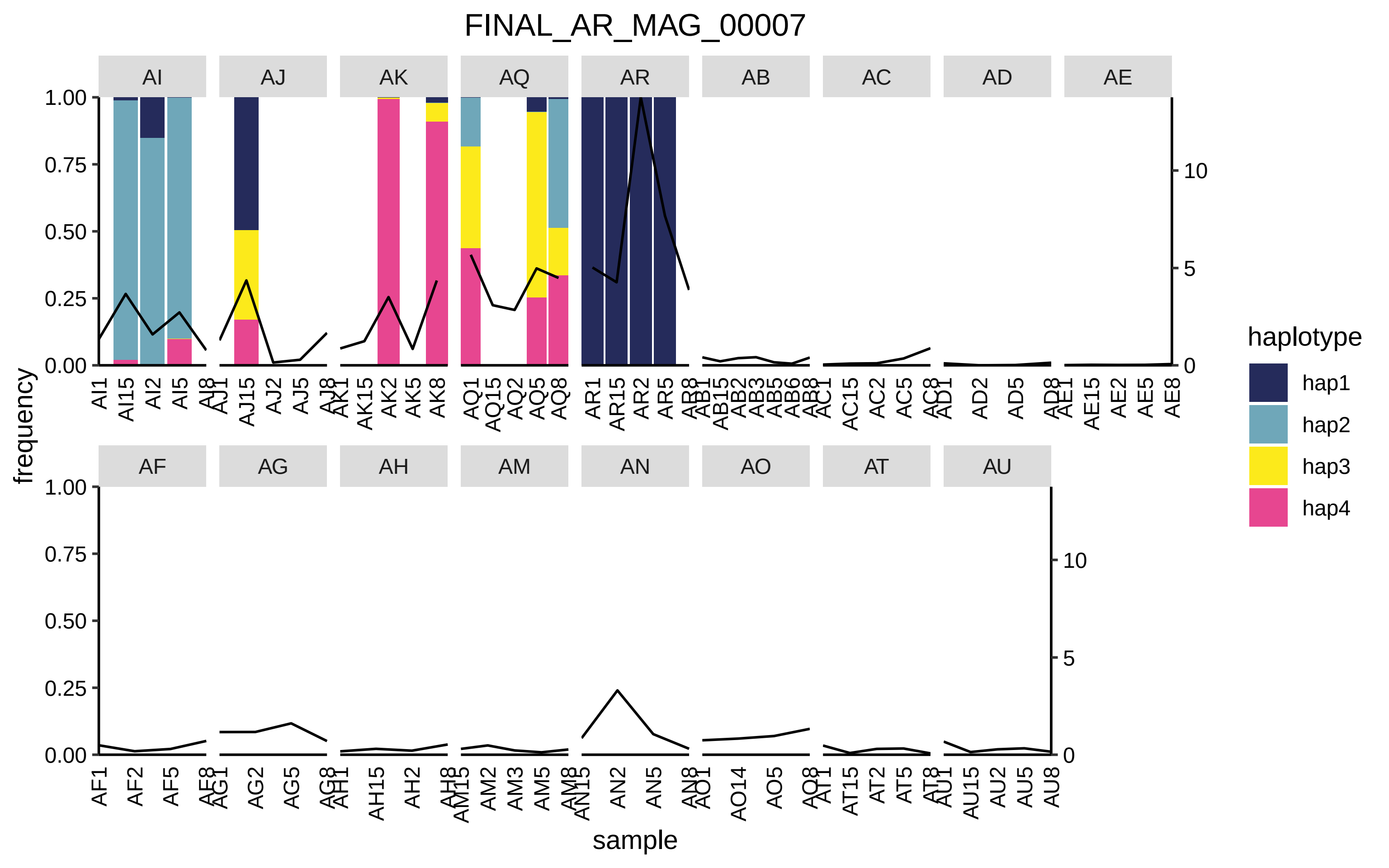
FINAL_AR_MAG_00007
AI
AJ
AK
AQ
AR
AB
AC
AD
AE
AF
AG
AH
AM
AN
AO
AT
AU
frequency
sample
haplotype
hap1
hap2
hap3
hap4
1.00
0.75
0.50
0.25
0.00
10
5
0
AI1 AI15 AI2 AI5 AI8
AJ1 AJ15 AJ2 AJ5 AJ8
AK1 AK15 AK2 AK5 AK8
AQ1 AQ15 AQ2 AQ5 AQ8
AR1 AR15 AR2 AR5 AR8
AB1 AB15 AB2 AB3 AB5 AB6 AB8
AC1 AC15 AC2 AC5 AC8
AD1 AD2 AD5 AD8
AE1 AE15 AE2 AE5 AE8
AF1 AF2 AF5 AF8
AG1 AG2 AG5 AG8
AH1 AH15 AH2 AH8
AM15 AM2 AM3 AM5 AM8
AN15 AN2 AN5 AN8
AO1 AO14 AO5 AO8
AT1 AT15 AT2 AT5 AT8
AU1 AU15 AU2 AU5 AU8

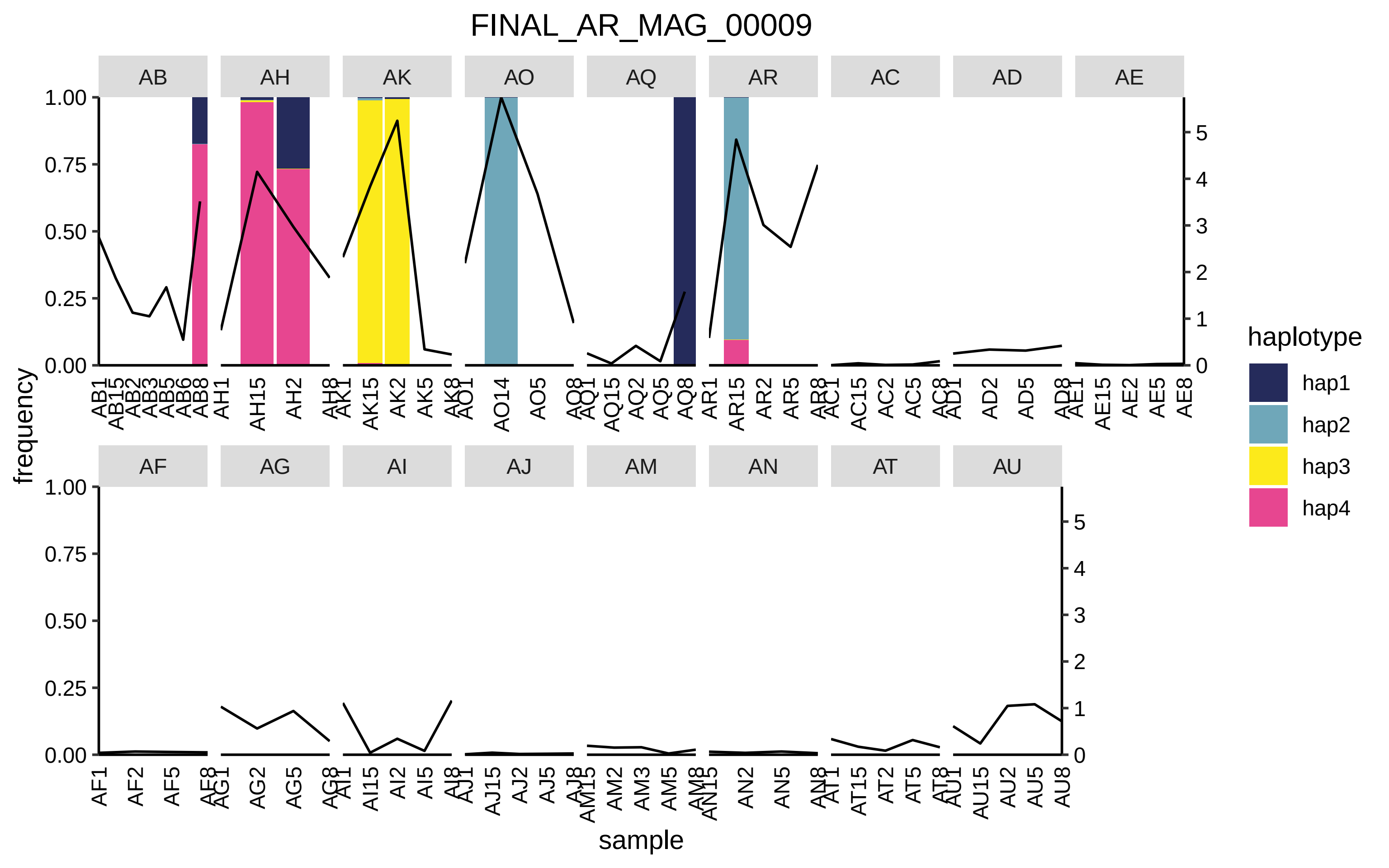
FINAL_AR_MAG_00009
AB
AH
AK
AO
AQ
AR
AC
AD
AE
AF
AG
AI
AJ
AM
AN
AT
AU
frequency
sample
haplotype
hap1
hap2
hap3
hap4

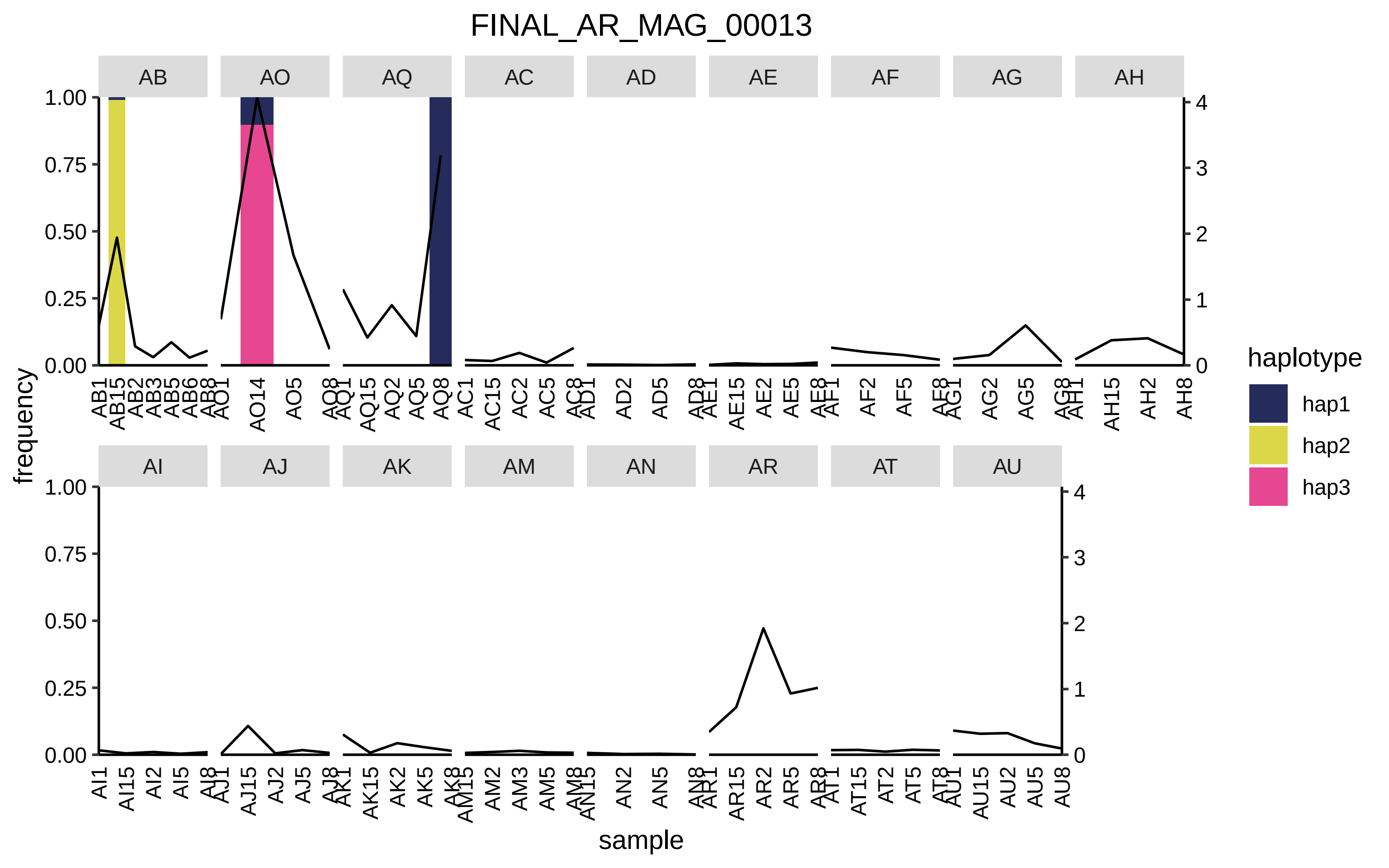
FINAL_AR_MAG_00013
AB
AO
AQ
AC
AD
AE
AF
AG
AH
AI
AJ
AK
AM
AN
AR
AT
AU
frequency
sample
haplotype
hap1
hap2
hap3
1.00
0.75
0.50
0.25
0.00
0
1
2
3
4

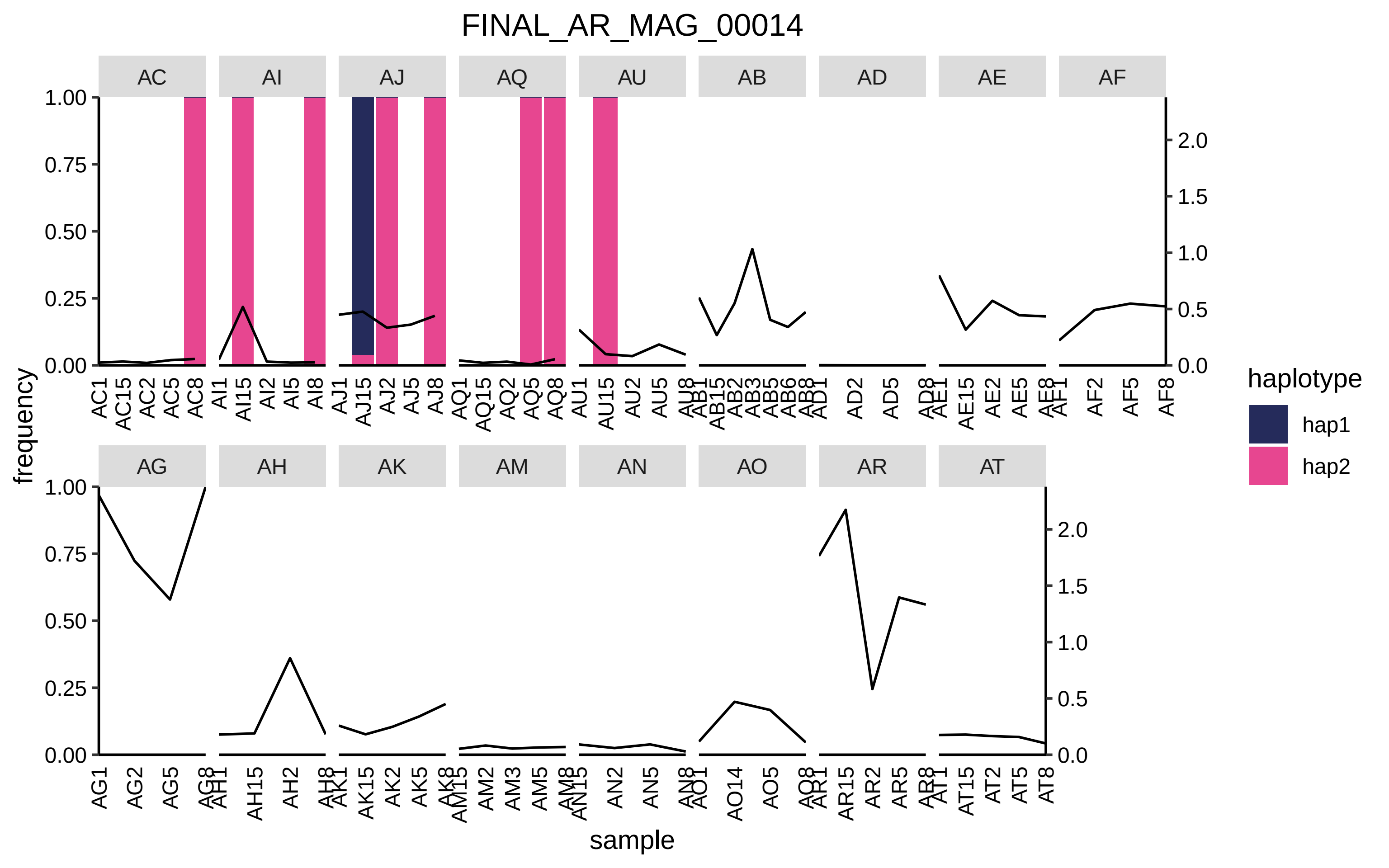

FINAL_AR_MAG_00014
AC
AI
AJ
AQ
AU
AB
AD
AE
AF
AG
AH
AK
AM
AN
AO
AR
AT
1.00
0.75
0.50
0.25
0.00
2.0
1.5
1.0
0.5
0.0
frequency
sample
AC1 AC15 AC2 AC5 AC8
AI1 AI15 AI2 AI5 AI8
AJ1 AJ15 AJ2 AJ5 AJ8
AQ1 AQ15 AQ2 AQ5 AQ8
AU1 AU15 AU2 AU5 AU8
AB1 AB15 AB2 AB3 AB5 AB6 AB8
AD1 AD2 AD5 AD8
AE1 AE15 AE2 AE5 AE8
AF1 AF2 AF5 AF8
AG1 AG2 AG5 AG8
AH1 AH15 AH2 AH8
AK1 AK15 AK2 AK5 AK8
AM15 AM2 AM3 AM5 AM8
AN15 AN2 AN5 AN8
AO1 AO14 AO5 AO8
AR1 AR15 AR2 AR5 AR8
AT1 AT15 AT2 AT5 AT8
haplotype
hap1
hap2

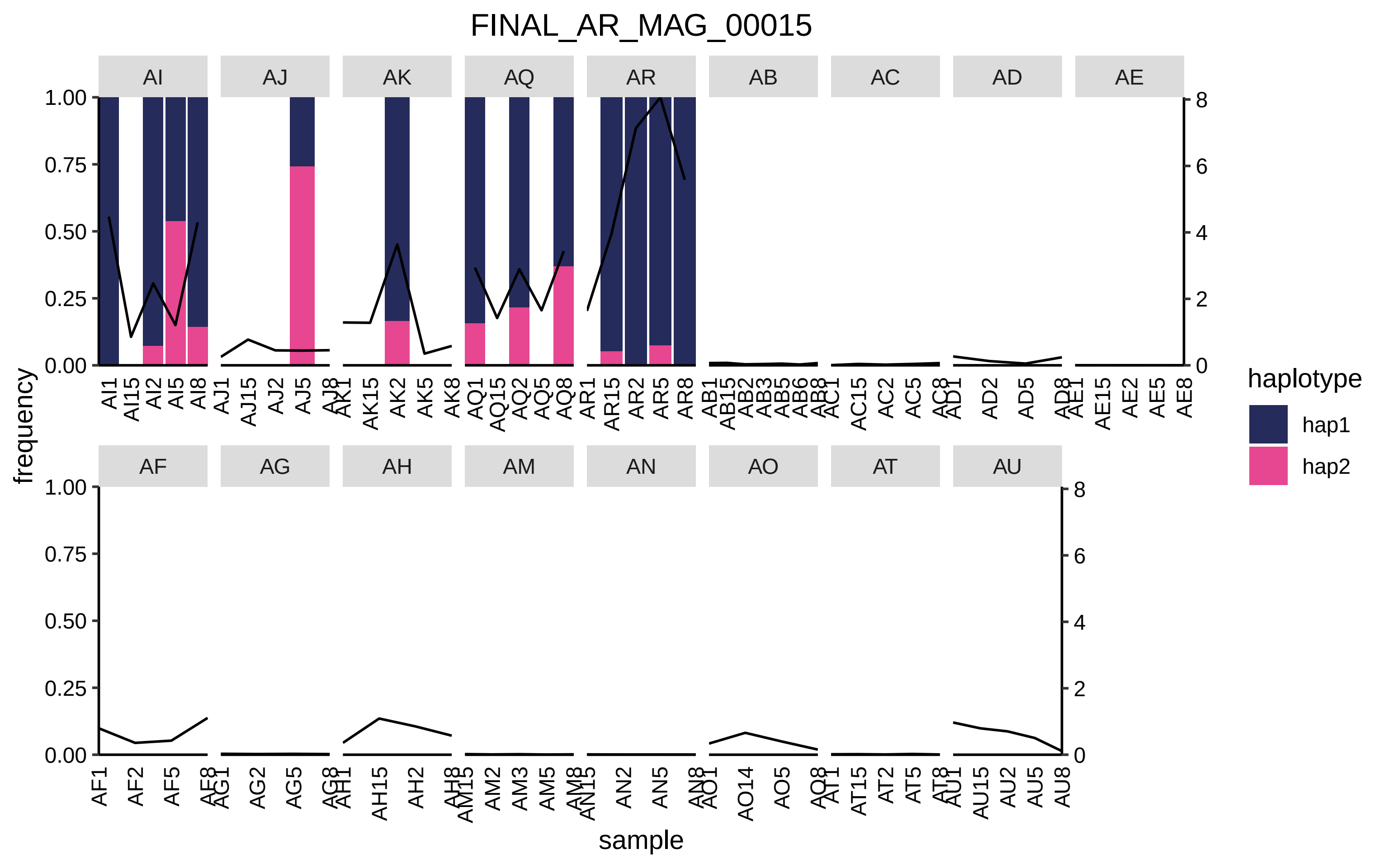
FINAL_AR_MAG_00015
AI
AJ
AK
AQ
AR
AB
AC
AD
AE
AF
AG
AH
AM
AN
AO
AT
AU
frequency
sample
haplotype
hap1
hap2
1.00
0.75
0.50
0.25
0.00
8
6
4
2
0
AI1 AI15 AI2 AI5 AI8
AJ1 AJ15 AJ2 AJ5 AJ8
AK1 AK15 AK2 AK5 AK8
AQ1 AQ15 AQ2 AQ5 AQ8
AR1 AR15 AR2 AR5 AR8
AB1 AB15 AB2 AB3 AB5 AB6 AB8
AC1 AC15 AC2 AC5 AC8
AD1 AD2 AD5 AD8
AE1 AE15 AE2 AE5 AE8
AF1 AF2 AF5 AF8
AG1 AG2 AG5 AG8
AH1 AH15 AH2 AH8
AM15 AM2 AM3 AM5 AM8
AN15 AN2 AN5 AN8
AO1 AO14 AO5 AO8
AT1 AT15 AT2 AT5 AT8
AU1 AU15 AU2 AU5 AU8

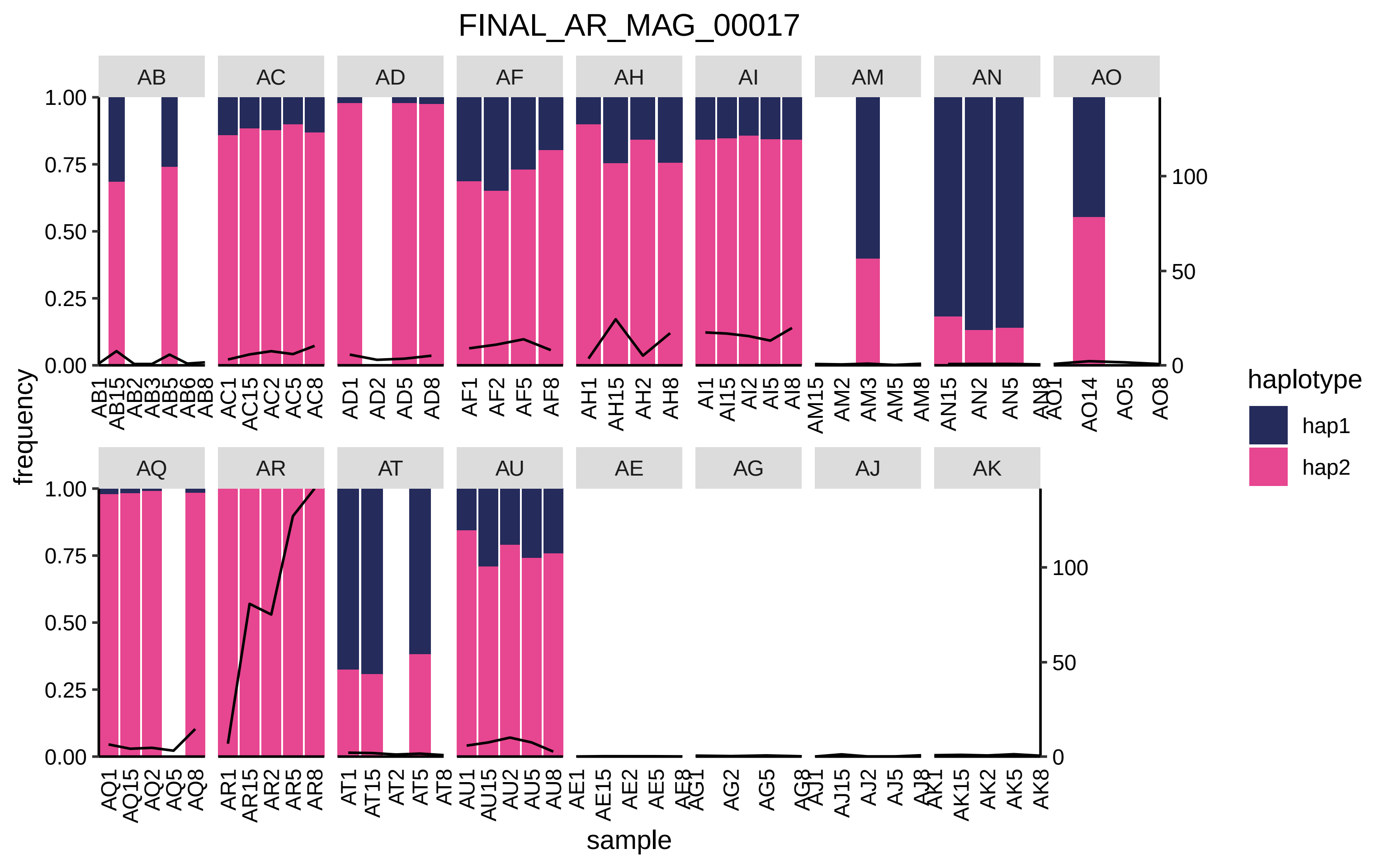
FINAL_AR_MAG_00017
AB
AC
AD
AF
AH
AI
AM
AN
AO
AQ
AR
AT
AU
AE
AG
AJ
AK
frequency
sample
haplotype
hap1
hap2
1.00
0.75
0.50
0.25
0.00
100
50
0
AB1 AB15 AB2 AB3 AB5 AB6 AB8
AC1 AC15 AC2 AC5 AC8
AD1 AD2 AD5 AD8
AF1 AF2 AF5 AF8
AH1 AH15 AH2 AH8
AI1 AI15 AI2 AI5 AI8
AM15 AM2 AM3 AM5 AM8
AN15 AN2 AN5 AN8
AO1 AO14 AO5 AO8
AQ1 AQ15 AQ2 AQ5 AQ8
AR1 AR15 AR2 AR5 AR8
AT1 AT15 AT2 AT5 AT8
AU1 AU15 AU2 AU5 AU8
AE1 AE15 AE2 AE5 AE8
AG1 AG2 AG5 AG8
AJ1 AJ15 AJ2 AJ5 AJ8
AK1 AK15 AK2 AK5 AK8

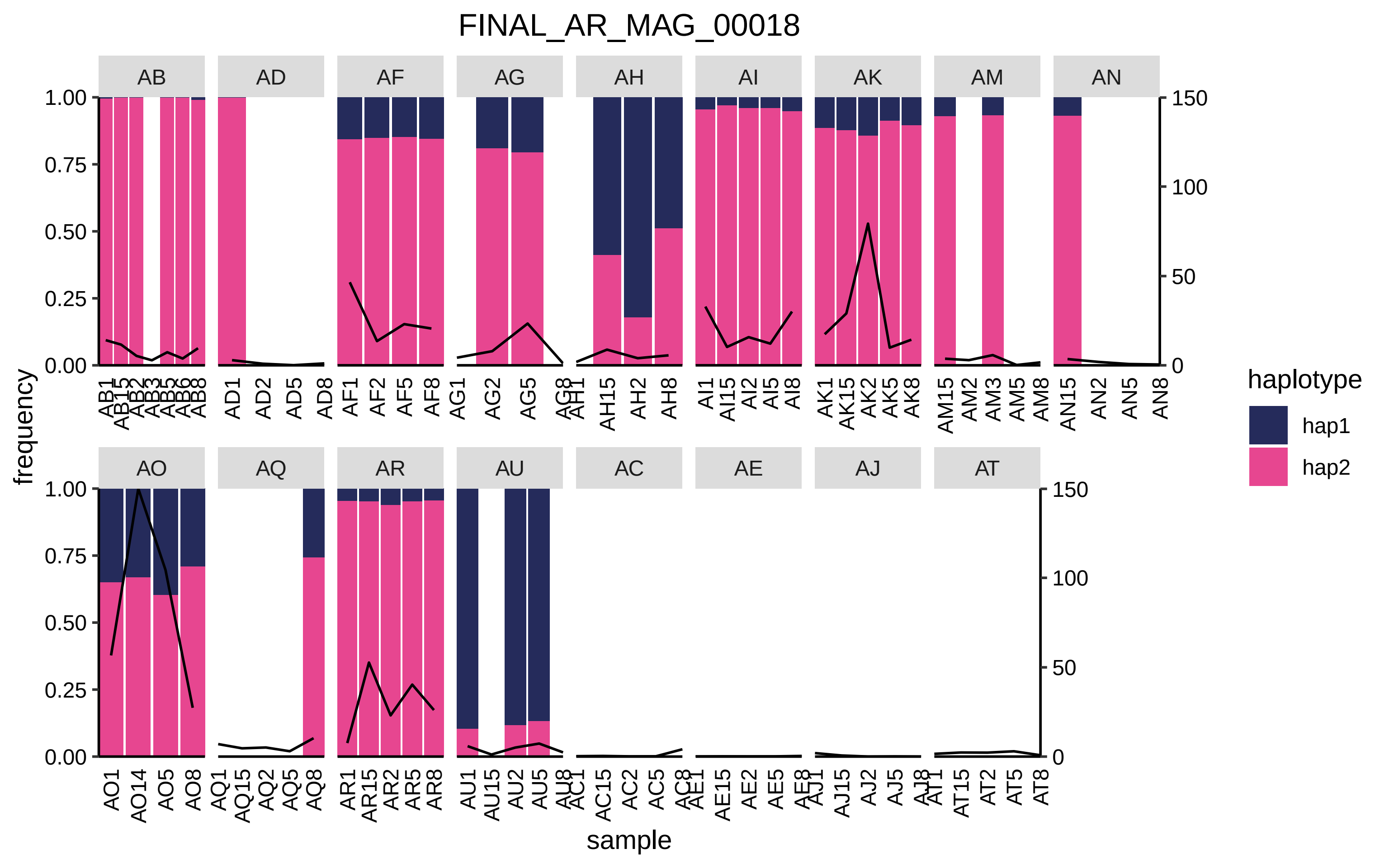
FINAL_AR_MAG_00018
AB
AD
AF
AG
AH
AI
AK
AM
AN
AO
AQ
AR
AU
AC
AE
AJ
AT
frequency
sample
1.00
0.75
0.50
0.25
0.00
150
100
50
0
AD1 AD2 AD5 AD8
AF1 AF2 AF5 AF8
AG2 AG5
AH15 AH2 AH8
AI1 AI15 AI2 AI5 AI8
AK1 AK15 AK2 AK5 AK8
AM15 AM2 AM3 AM5 AM8
AN15 AN2 AN5 AN8
AO1 AO14 AO5 AO8
AQ1 AQ15 AQ2 AQ5 AQ8
AR1 AR15 AR2 AR5 AR8
AU1 AU15 AU2 AU5
AC15 AC2 AC5
AE15 AE2 AE5
AJ15 AJ2 AJ5
AT15 AT2 AT5 AT8
haplotype
hap1
hap2

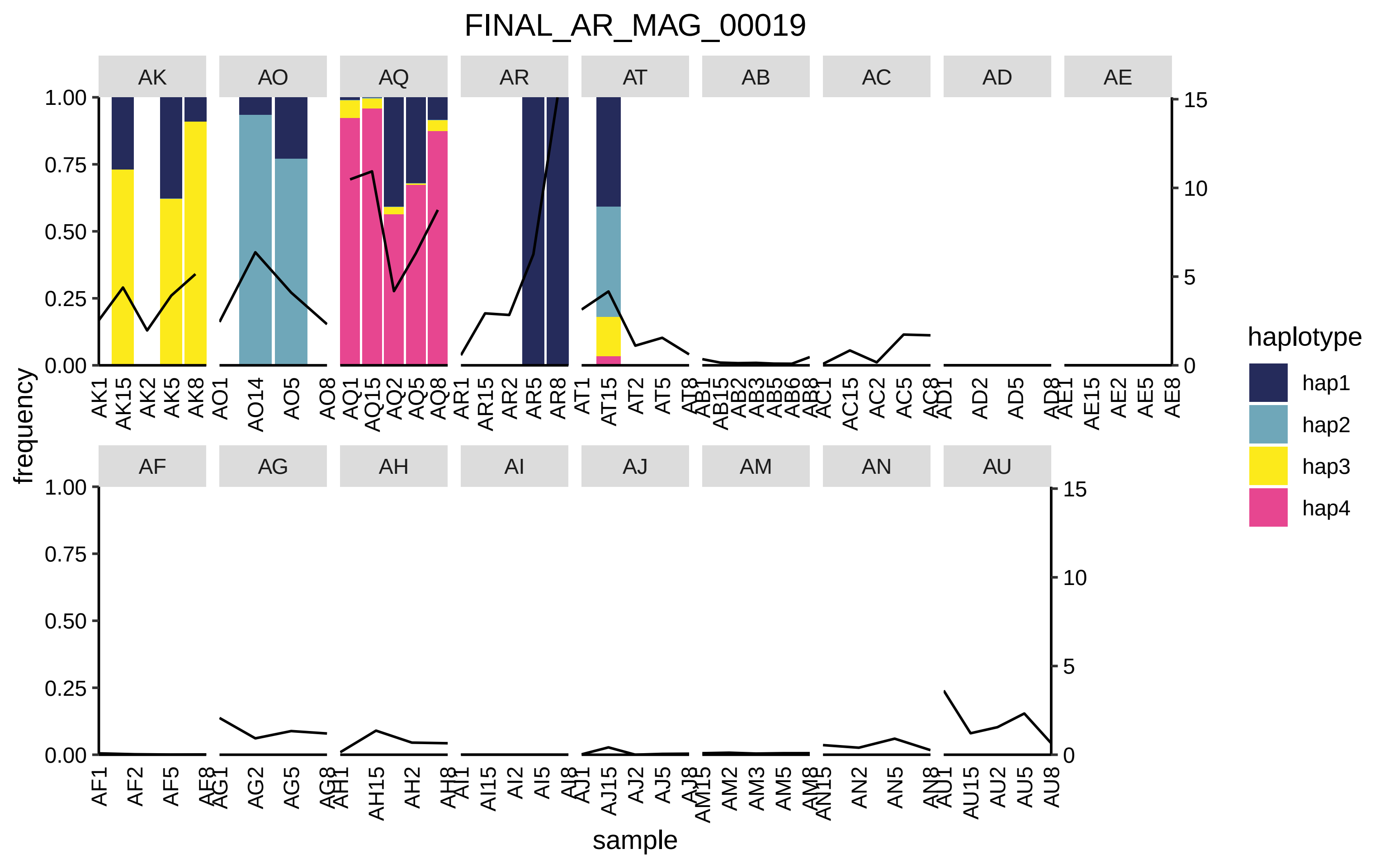
FINAL_AR_MAG_00019
AK
AO
AQ
AR
AT
AB
AC
AD
AE
AF
AG
AH
AI
AJ
AM
AN
AU
frequency
sample
haplotype
hap1
hap2
hap3
hap4
1.00
0.75
0.50
0.25
0.00
15
10
5
0

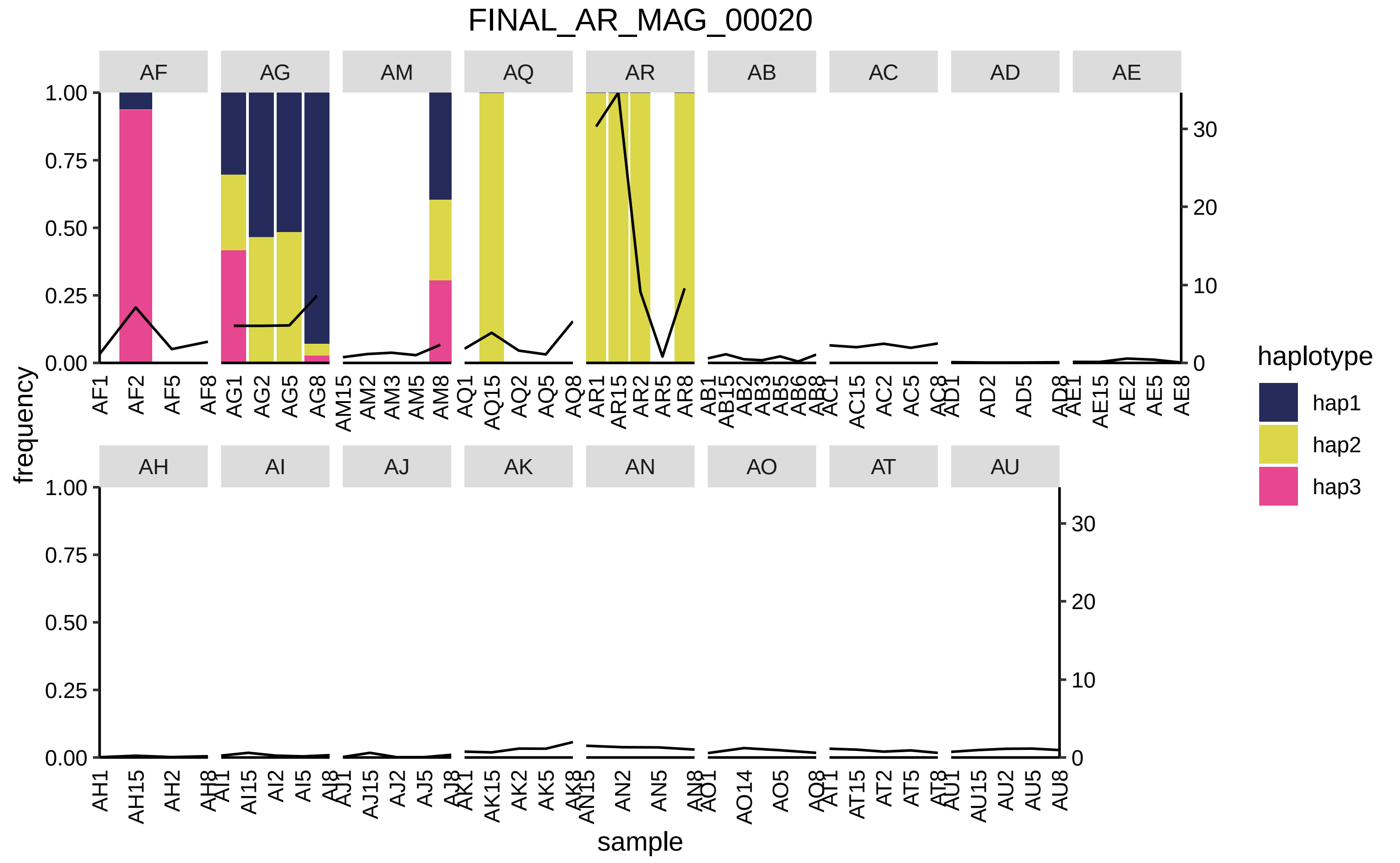

FINAL_AR_MAG_00020
AF
AG
AM
AQ
AR
AB
AC
AD
AE
AH
AI
AJ
AK
AN
AO
AT
AU
frequency
sample
haplotype
hap1
hap2
hap3

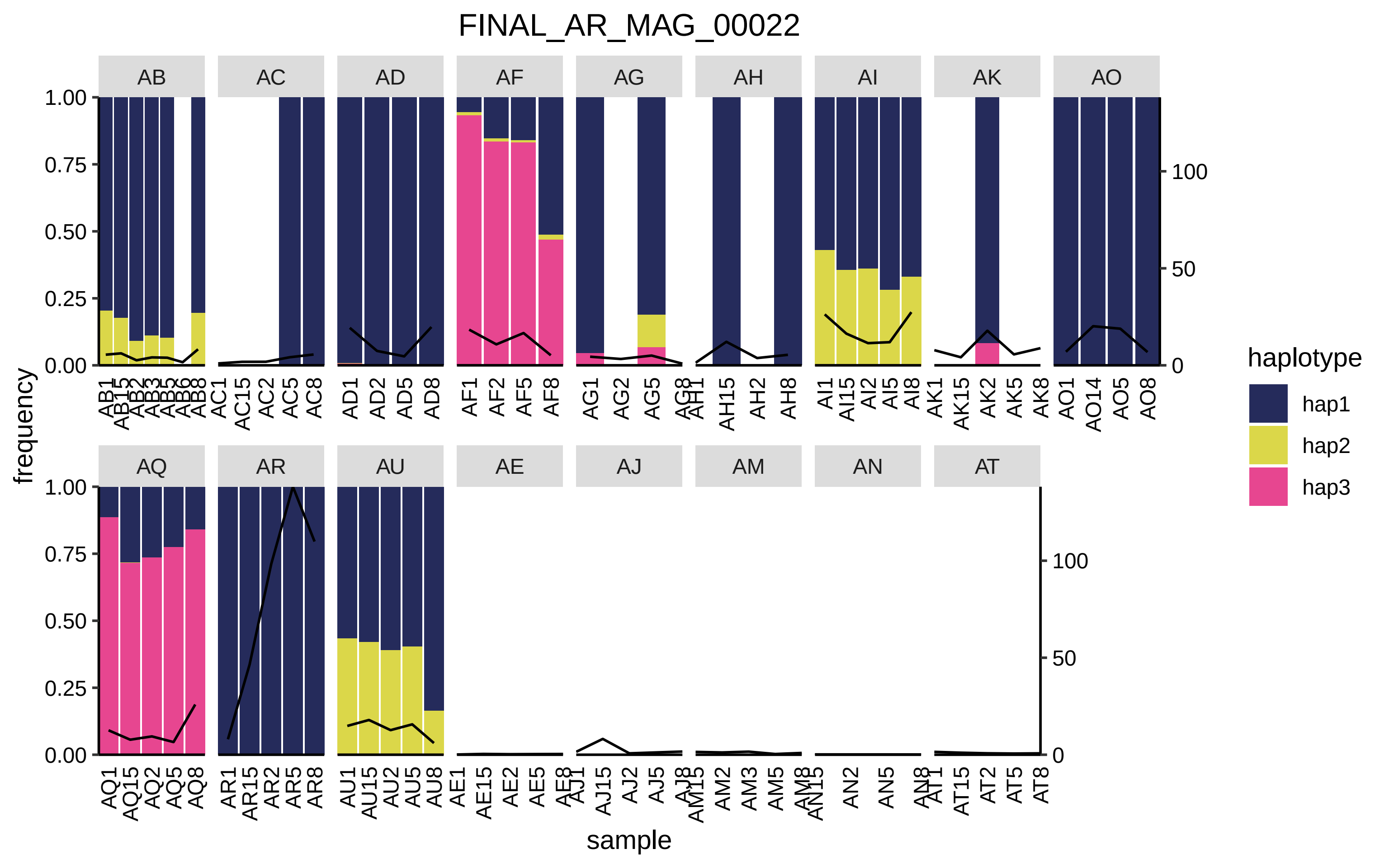

FINAL_AR_MAG_00022
AB
AC
AD
AF
AG
AH
AI
AK
AO
AQ
AR
AU
AE
AJ
AM
AN
AT
1.00
0.75
0.50
0.25
0.00
100
50
0
frequency
sample
haplotype
hap1
hap2
hap3

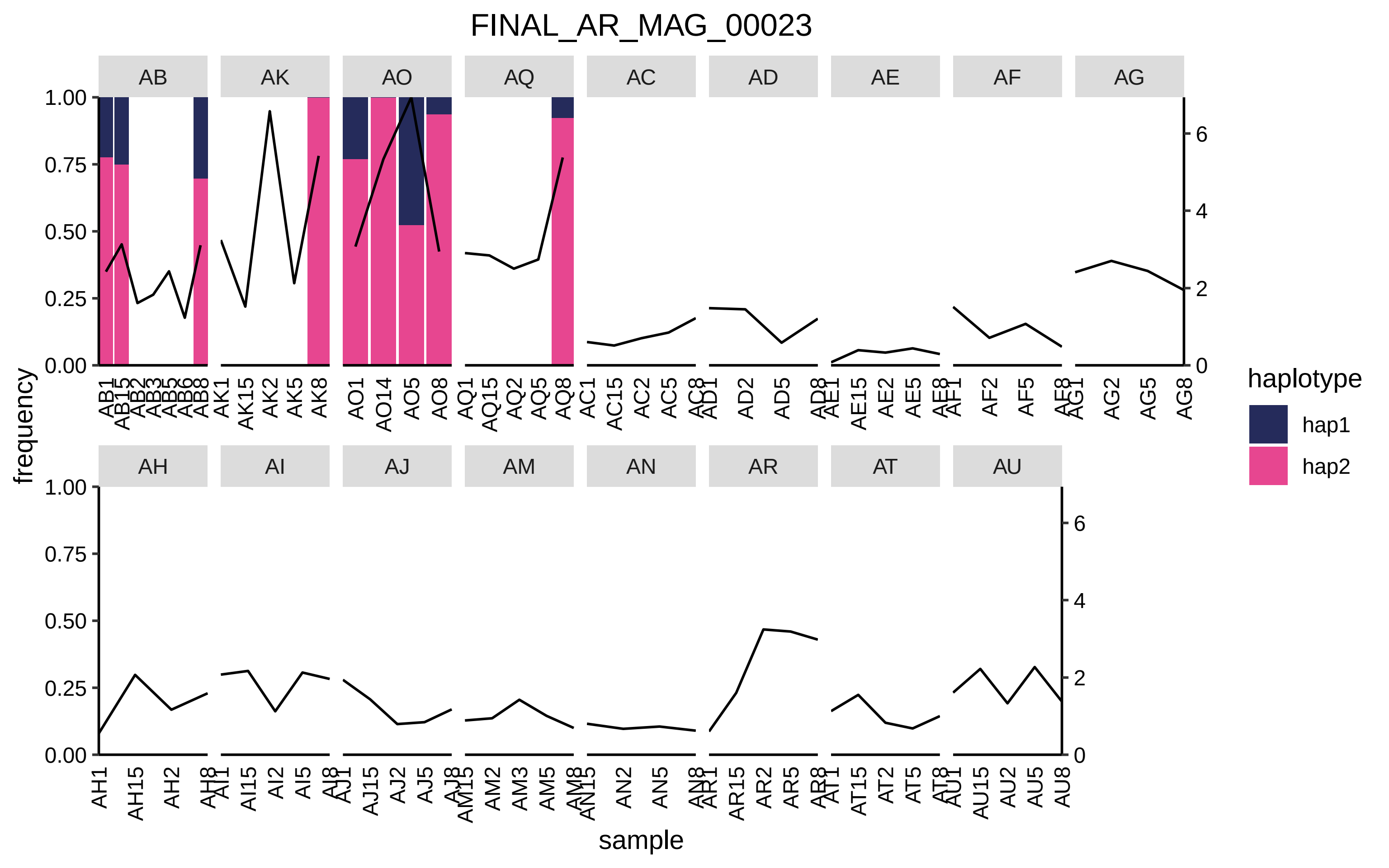
FINAL_AR_MAG_00023
AB
AK
AO
AQ
AC
AD
AE
AF
AG
AH
AI
AJ
AM
AN
AR
AT
AU
frequency
sample
haplotype
hap1
hap2

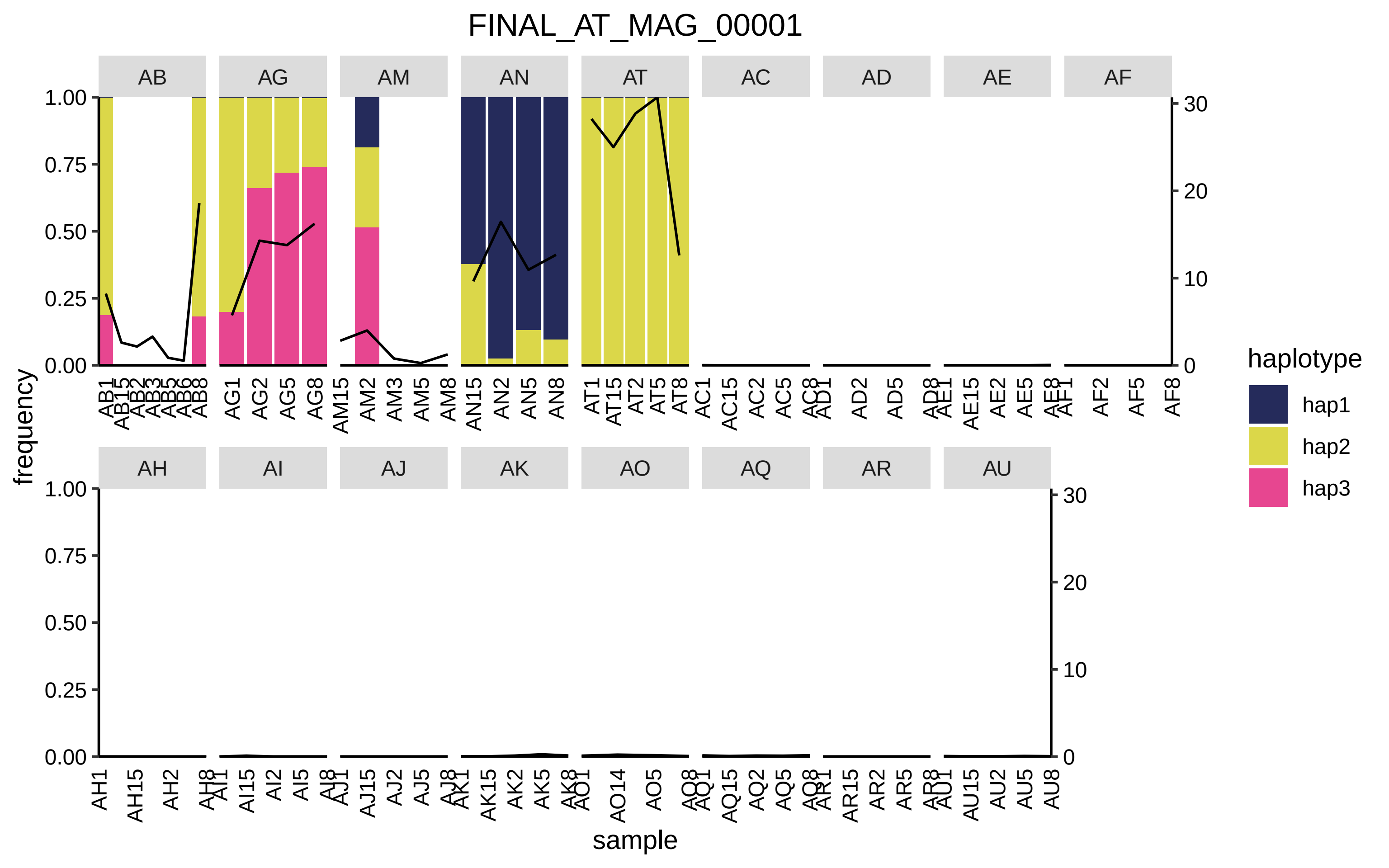

FINAL_AT_MAG_00001
AB
AG
AM
AN
AT
AC
AD
AE
AF
AH
AI
AJ
AK
AO
AQ
AR
AU
frequency
sample
haplotype
hap1
hap2
hap3

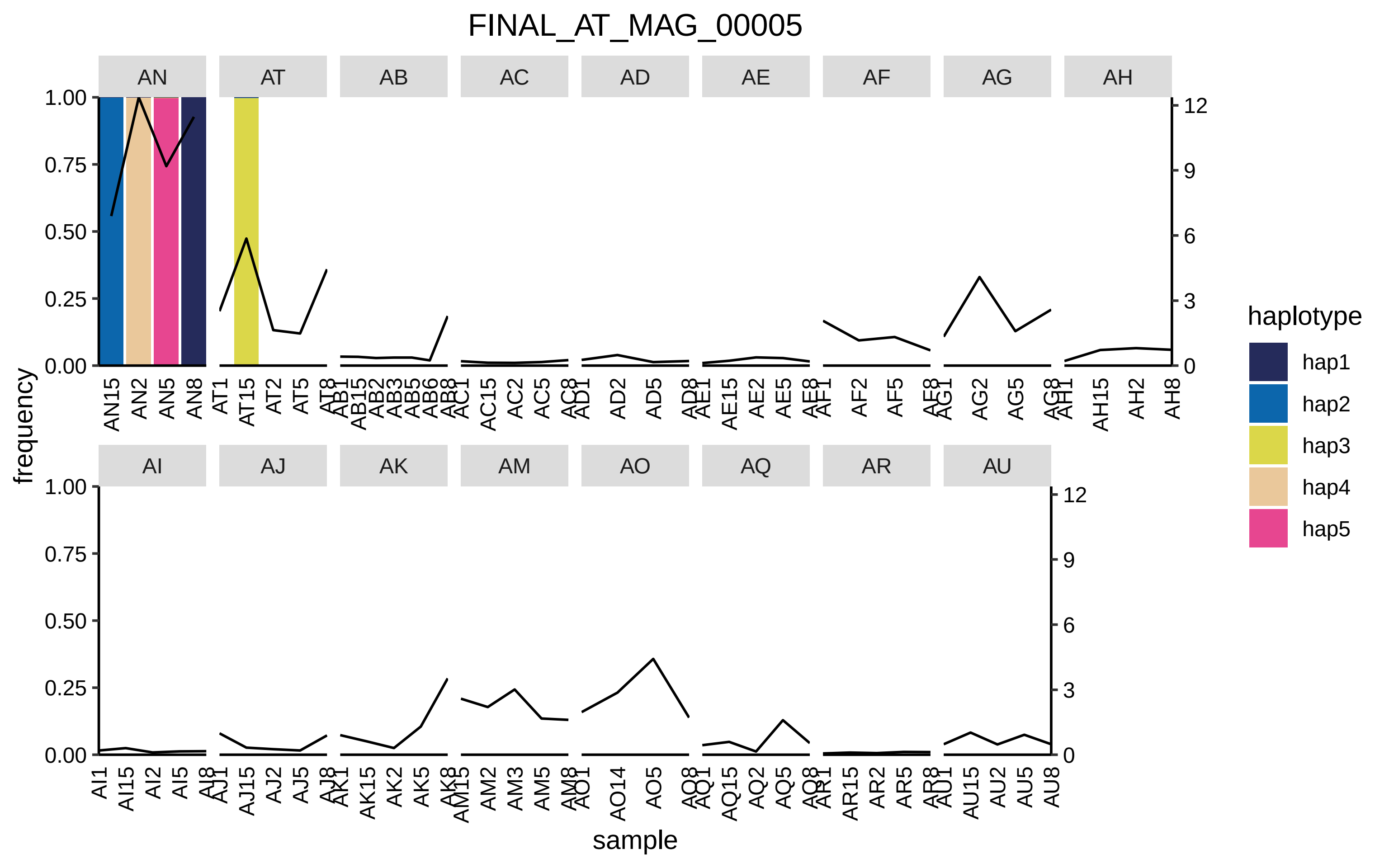

FINAL_AT_MAG_00005
AN
AT
AB
AC
AD
AE
AF
AG
AH
AI
AJ
AK
AM
AO
AQ
AR
AU
frequency
sample
haplotype
hap1
hap2
hap3
hap4
hap5
1.00
0.75
0.50
0.25
0.00
0
3
6
9
12

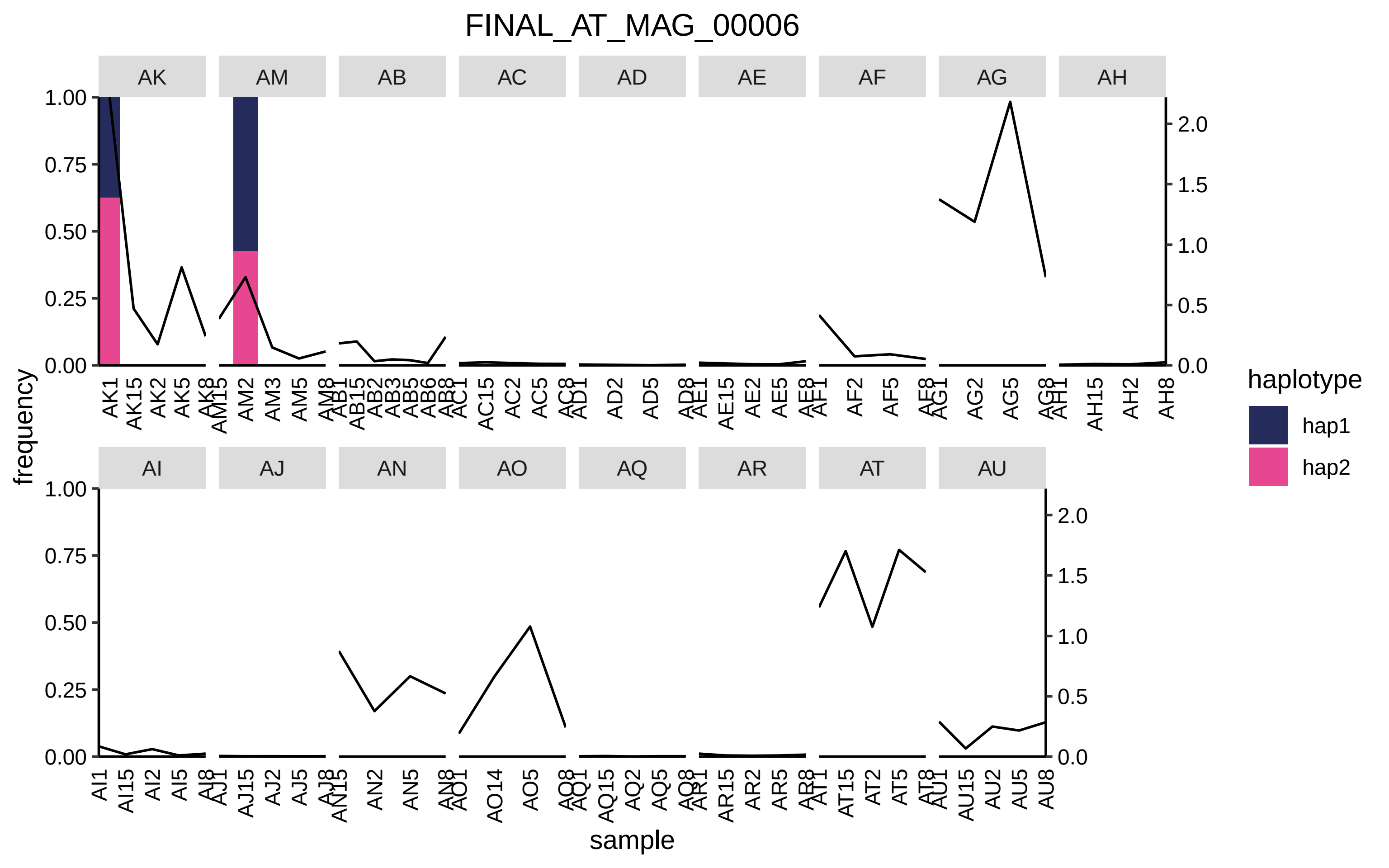
FINAL_AT_MAG_00006
AK
AM
AB
AC
AD
AE
AF
AG
AH
AI
AJ
AN
AO
AQ
AR
AT
AU
frequency
sample
haplotype
hap1
hap2

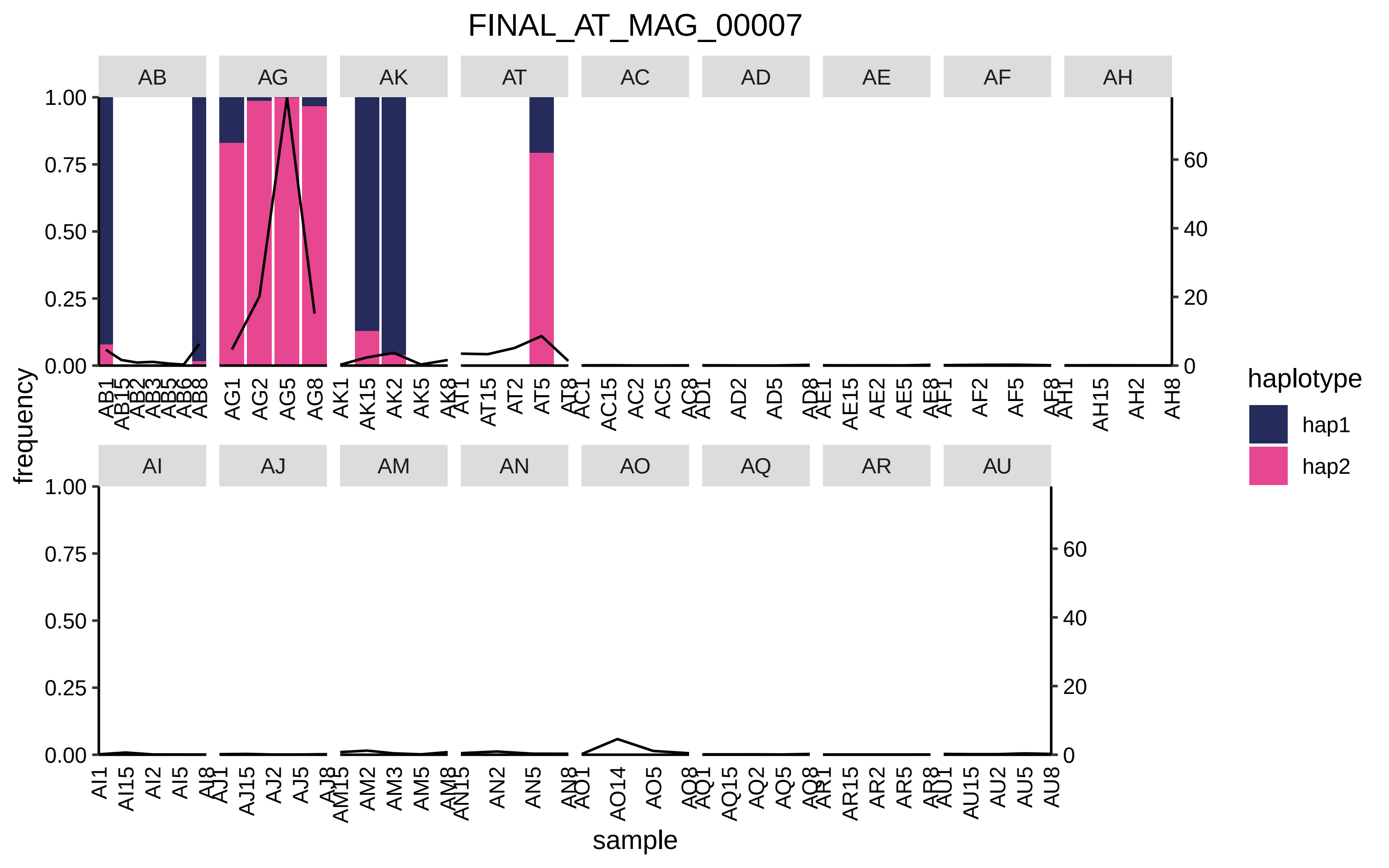
FINAL_AT_MAG_00007
AB
AG
AK
AT
AC
AD
AE
AF
AH
AI
AJ
AM
AN
AO
AQ
AR
AU
frequency
sample
haplotype
hap1
hap2

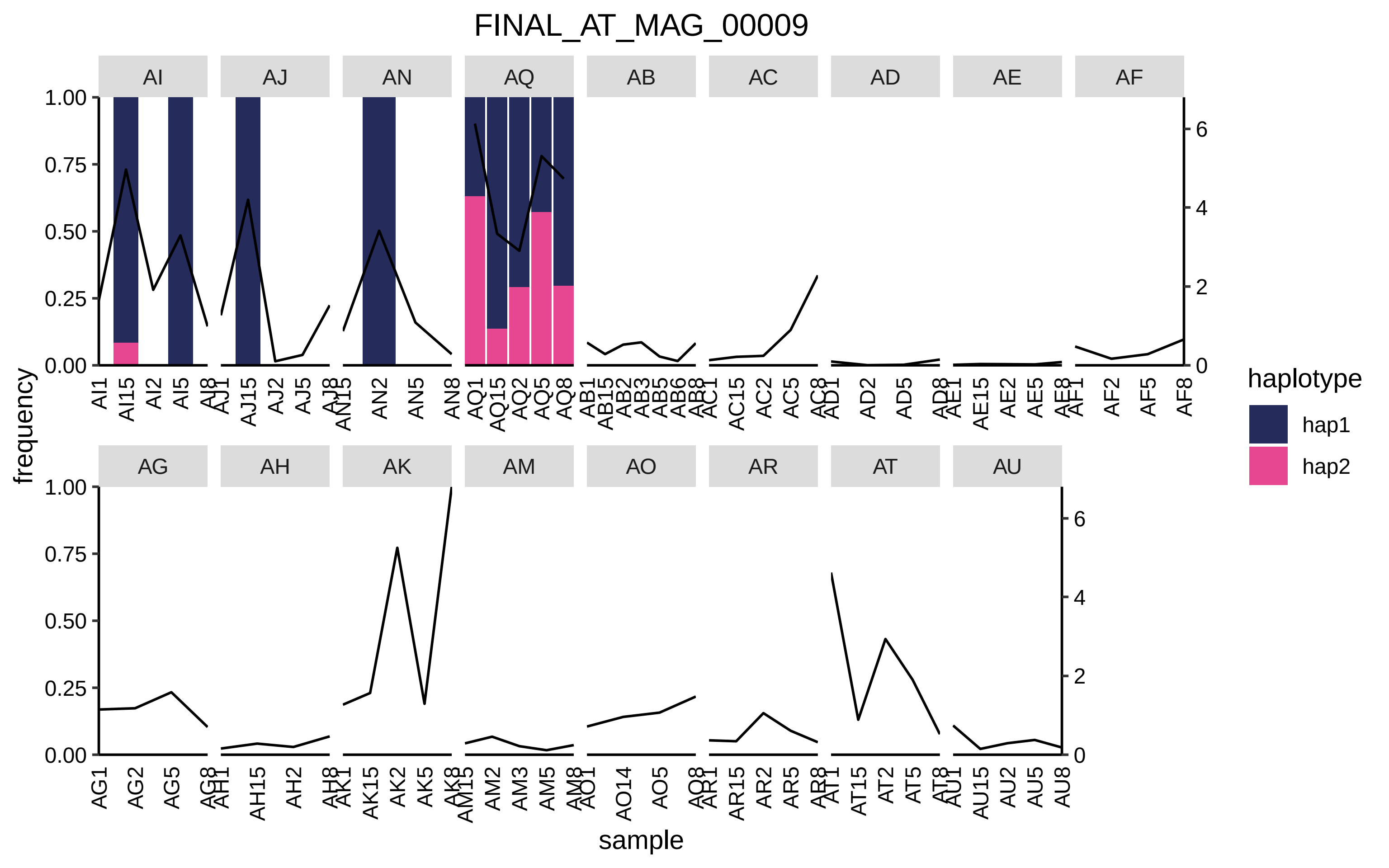
FINAL_AT_MAG_00009
AI
AJ
AN
AQ
AB
AC
AD
AE
AF
AG
AH
AK
AM
AO
AR
AT
AU
frequency
sample
haplotype
hap1
hap2

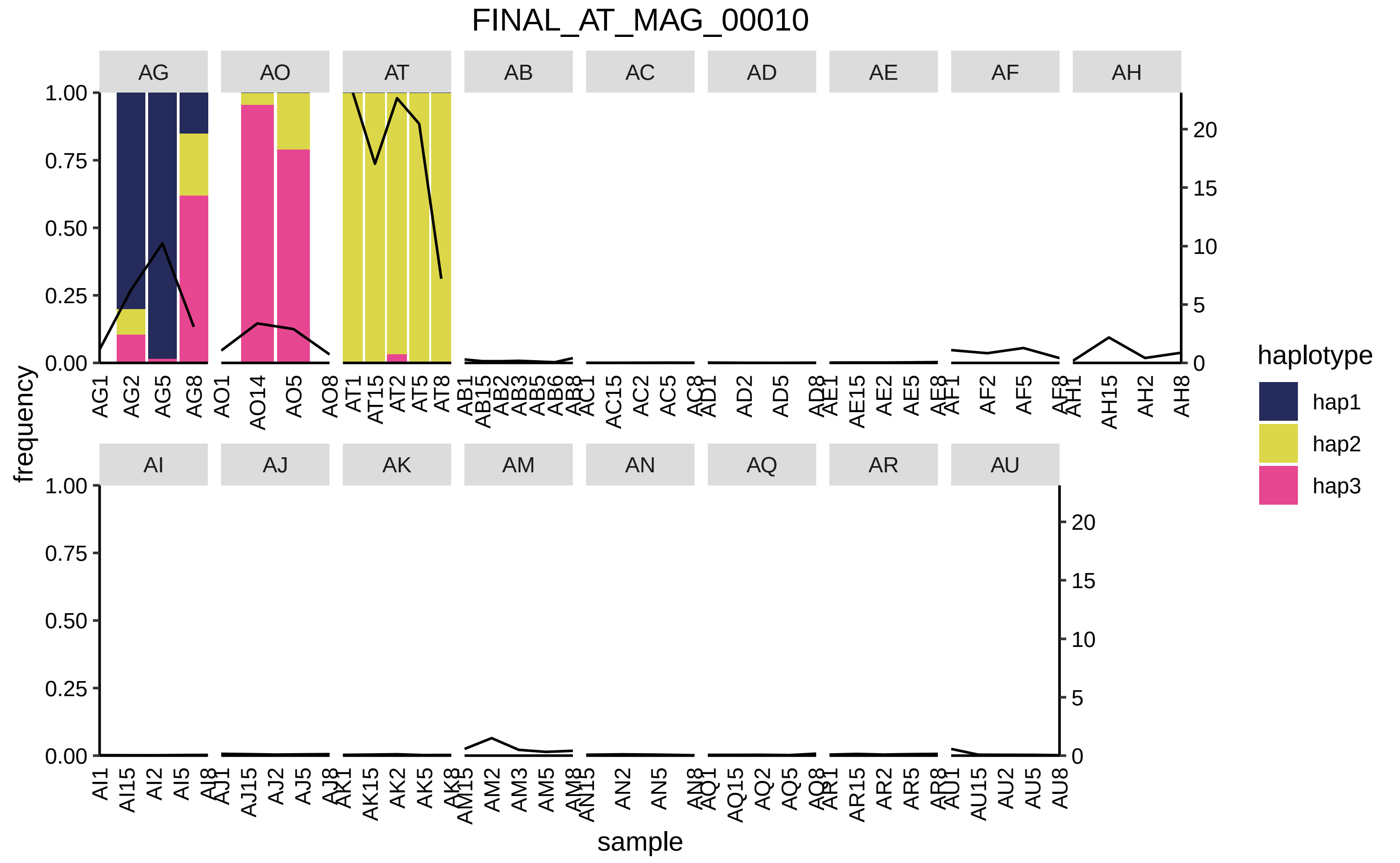

FINAL_AT_MAG_00010
AG
AO
AT
AB
AC
AD
AE
AF
AH
AI
AJ
AK
AM
AN
AQ
AR
AU
frequency
sample
haplotype
hap1
hap2
hap3
1.00
0.75
0.50
0.25
0.00
20
15
10
5
0
AG1
AG2
AG5
AG8
AO1
AO14
AO5
AO8
AT1
AT15
AT2
AT5
AT8
AB1
AB15
AB2
AB3
AB5
AB6
AB8
AC1
AC15
AC2
AC5
AC8
AD1
AD2
AD5
AD8
AE1
AE15
AE2
AE5
AE8
AF1
AF2
AF5
AF8
AH1
AH15
AH2
AH8
AI1
AI15
AI2
AI5
AI8
AJ1
AJ15
AJ2
AJ5
AJ8
AK1
AK15
AK2
AK5
AK8
AM15
AM2
AM3
AM5
AM8
AN15
AN2
AN5
AN8
AQ1
AQ15
AQ2
AQ5
AQ8
AR1
AR15
AR2
AR5
AR8
AU1
AU15
AU2
AU5
AU8

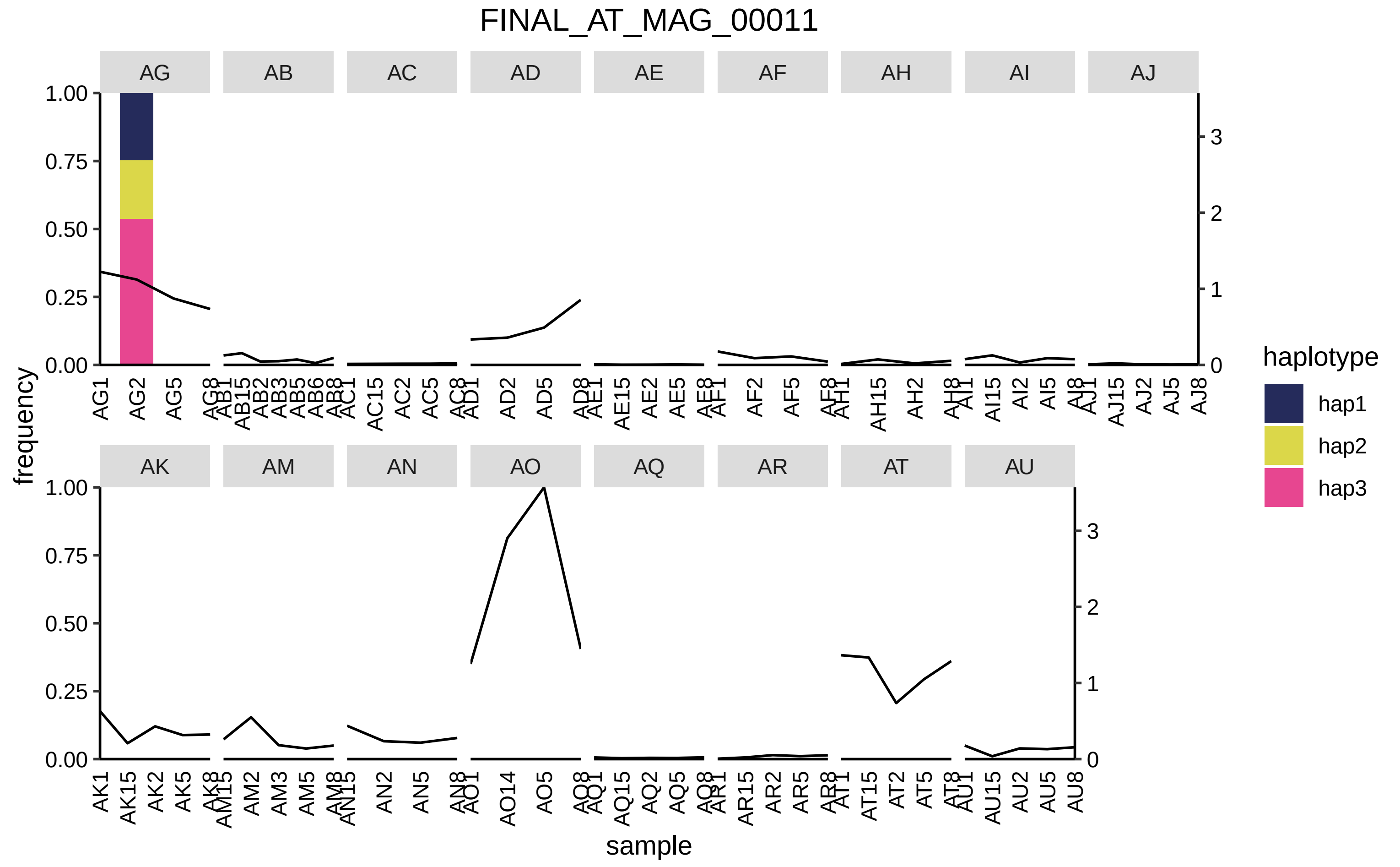

FINAL_AT_MAG_00011
AG
AB
AC
AD
AE
AF
AH
AI
AJ
AK
AM
AN
AO
AQ
AR
AT
AU
frequency
sample
haplotype
hap1
hap2
hap3
1.00
0.75
0.50
0.25
0.00
0
1
2
3
AG1
AG2
AG5
AG8
AK1
AK15
AK2
AK5
AK8
AU1
AU15
AU2
AU5
AU8

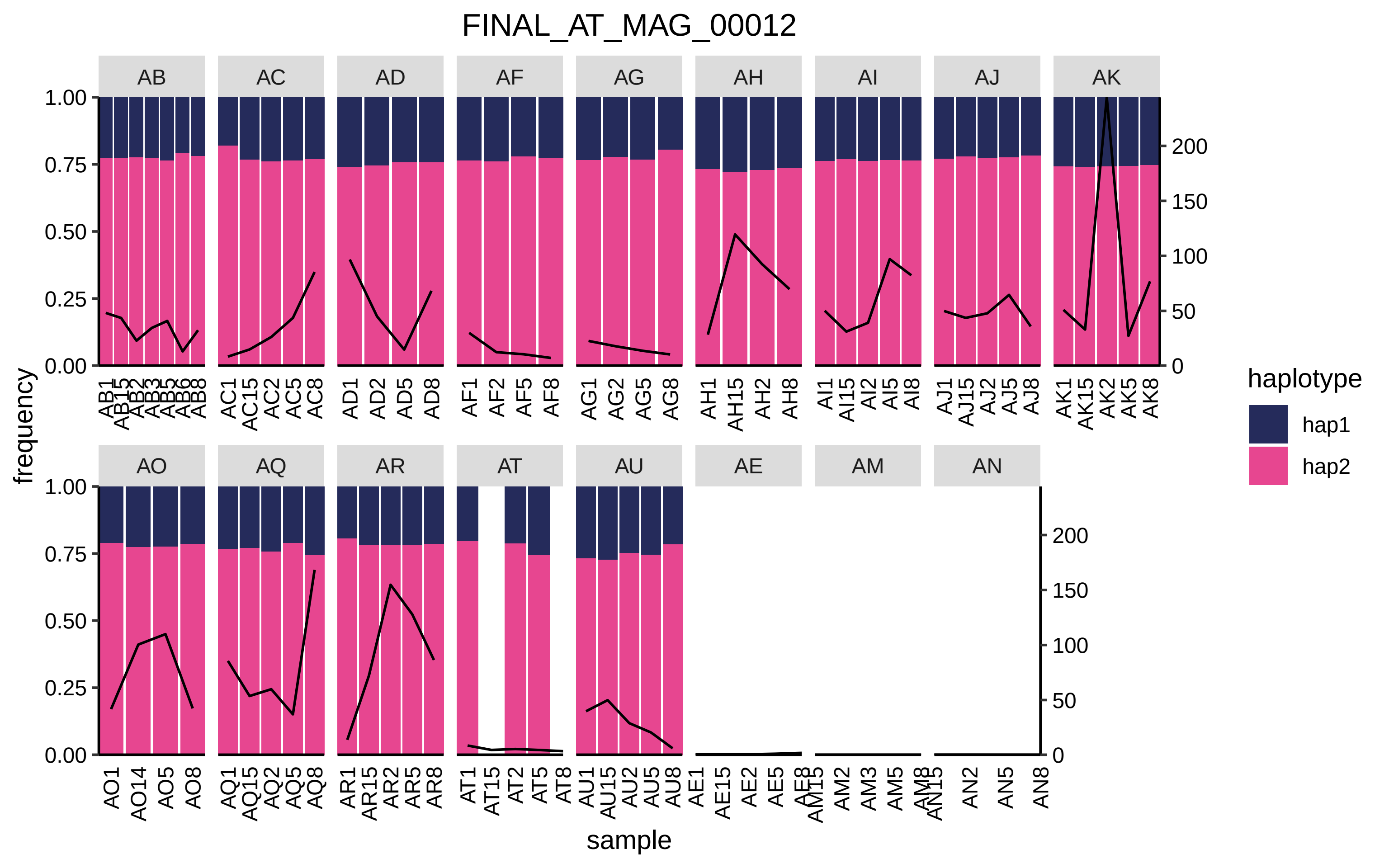

FINAL_AT_MAG_00012
AB
AC
AD
AF
AG
AH
AI
AJ
AK
AO
AQ
AR
AT
AU
AE
AM
AN
frequency
1.00
0.75
0.50
0.25
0.00
200
150
100
50
0
AB1 AB15 AB2 AB3 AB5 AB6 AB8
AC1 AC15 AC2 AC5 AC8
AD1 AD2 AD5 AD8
AF1 AF2 AF5 AF8
AG1 AG2 AG5 AG8
AH1 AH15 AH2 AH8
AI1 AI15 AI2 AI5 AI8
AJ1 AJ15 AJ2 AJ5 AJ8
AK1 AK15 AK2 AK5 AK8
AO1 AO14 AO5 AO8
AQ1 AQ15 AQ2 AQ5 AQ8
AR1 AR15 AR2 AR5 AR8
AT1 AT15 AT2 AT5 AT8
AU1 AU15 AU2 AU5 AU8
AE1 AE15 AE2 AE5 AE8
AM15 AM2 AM3 AM5 AM8
AN15 AN2 AN5 AN8
sample
haplotype
hap1
hap2

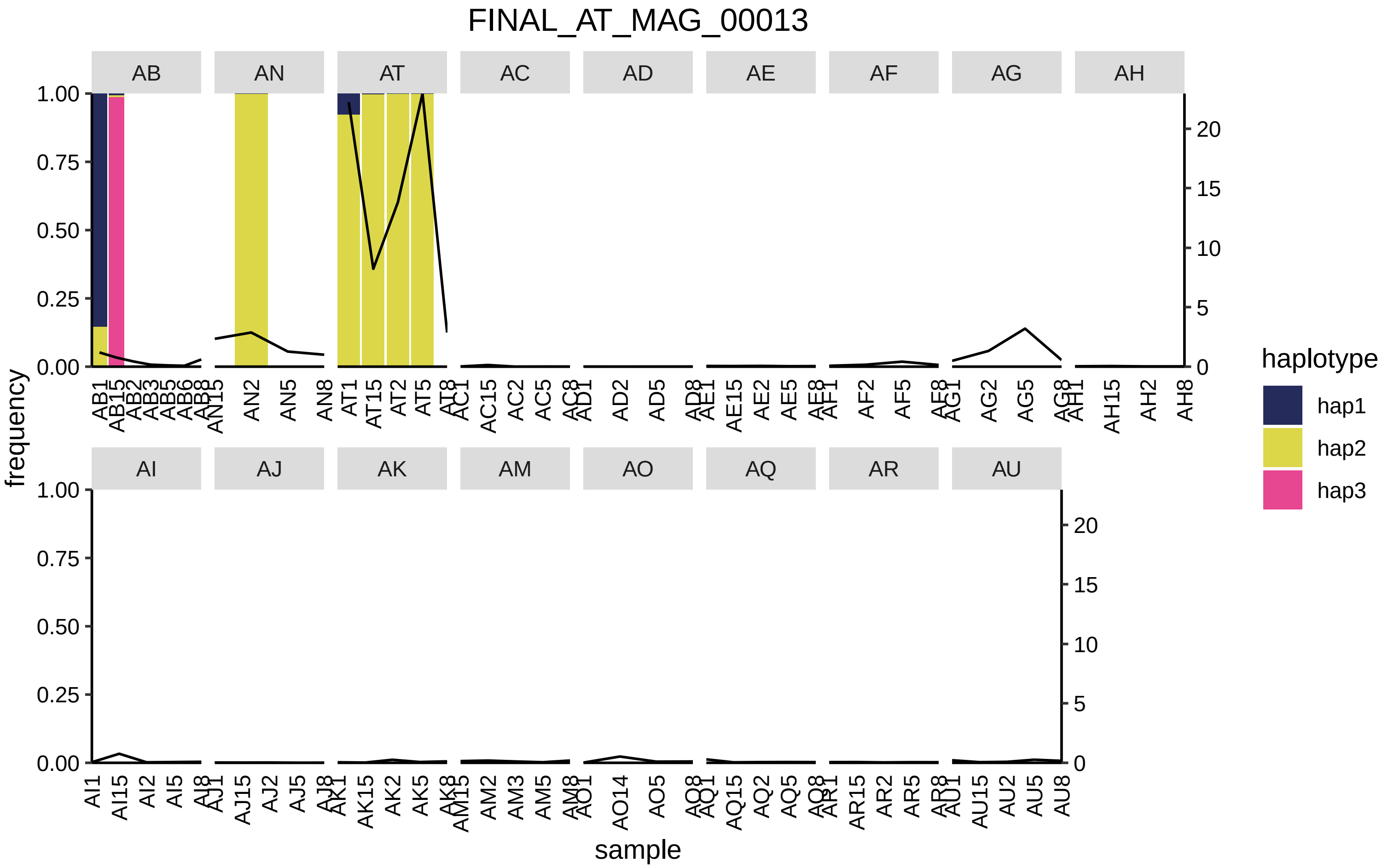
FINAL_AT_MAG_00013
AB
AN
AT
AC
AD
AE
AF
AG
AH
AI
AJ
AK
AM
AO
AQ
AR
AU
frequency
sample
haplotype
hap1
hap2
hap3

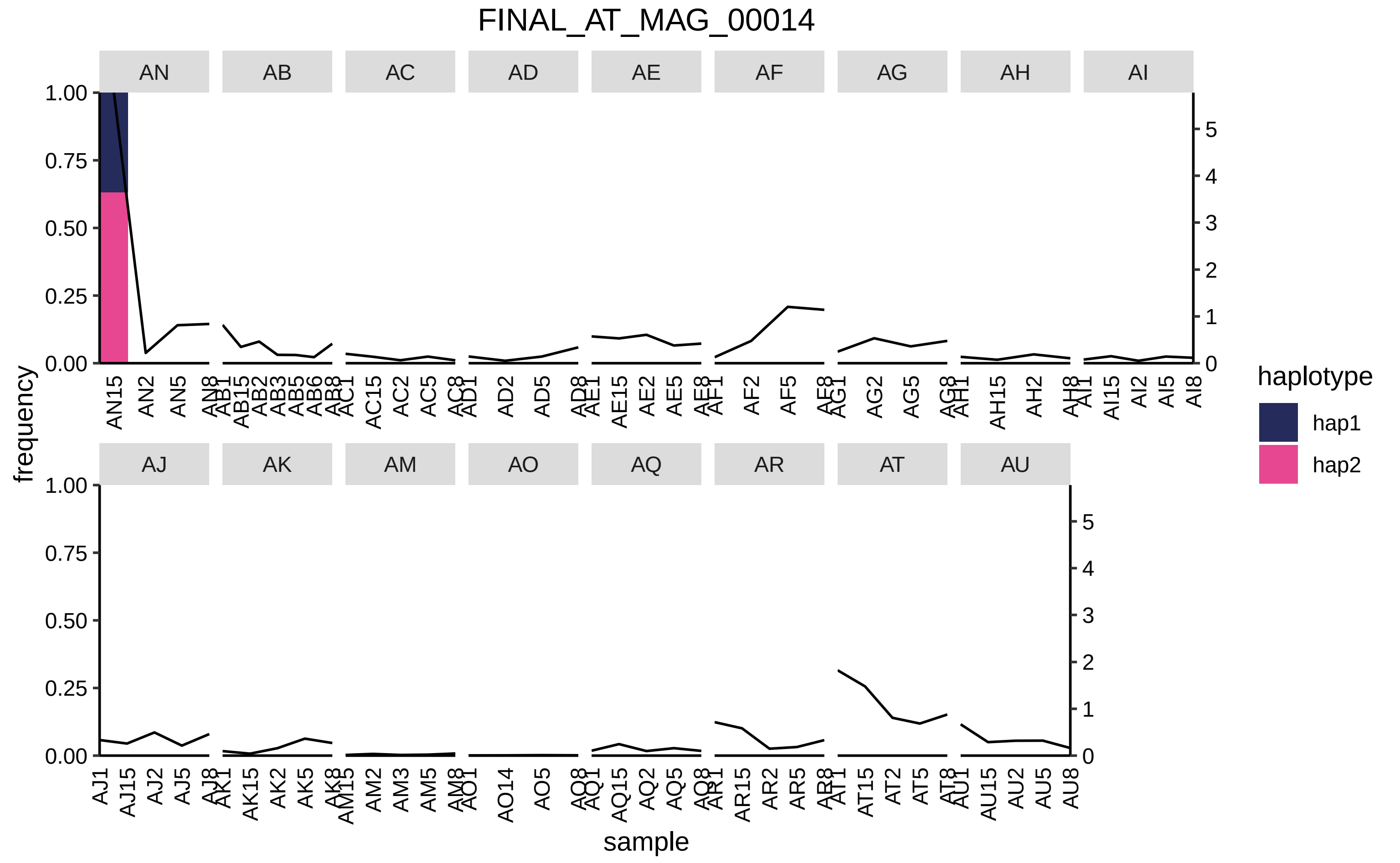
FINAL_AT_MAG_00014
AN
AB
AC
AD
AE
AF
AG
AH
AI
AJ
AK
AM
AO
AQ
AR
AT
AU
frequency
sample
haplotype
hap1
hap2
1.00
0.75
0.50
0.25
0.00
0
1
2
3
4
5

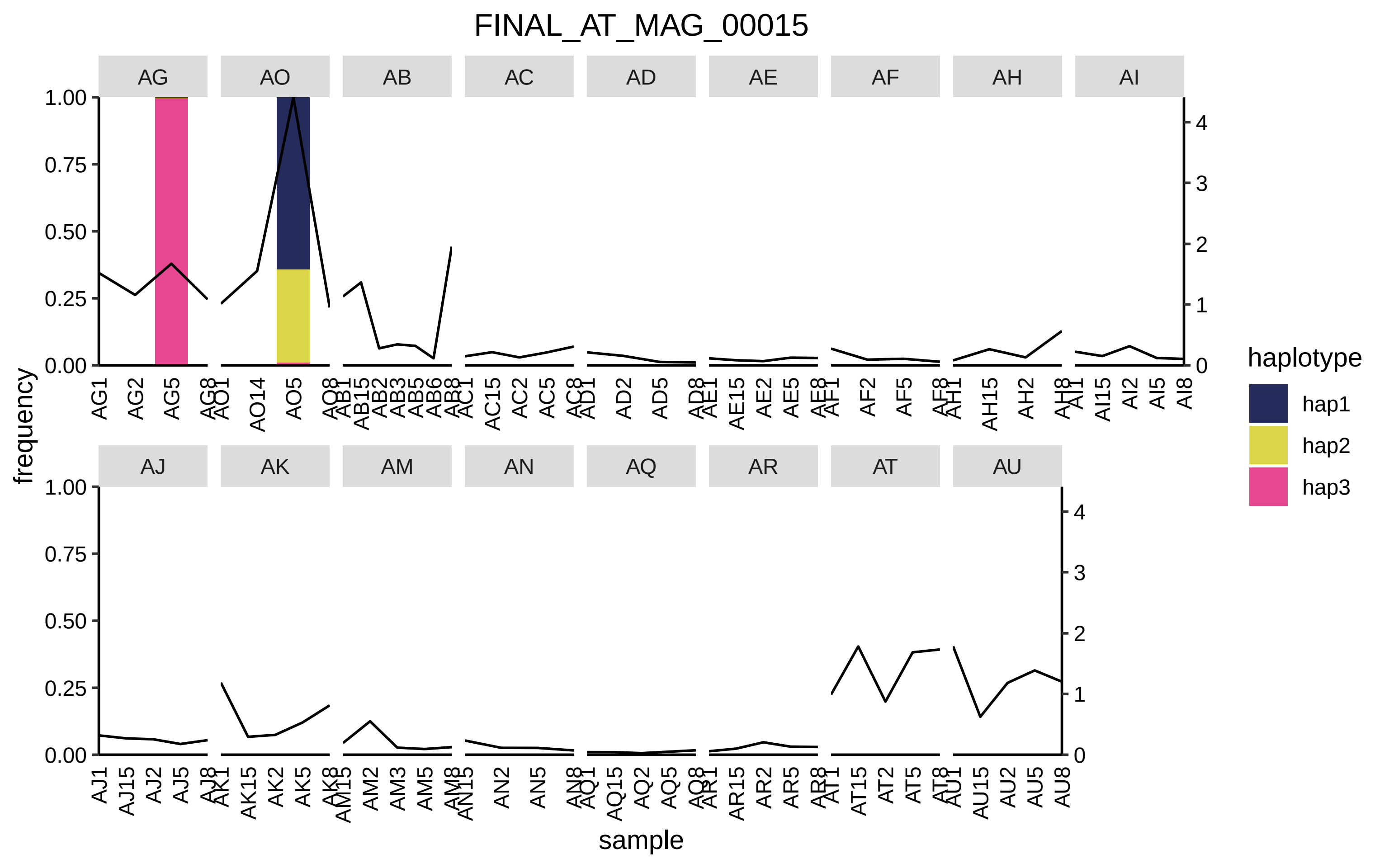

FINAL_AT_MAG_00015
AG
AO
AB
AC
AD
AE
AF
AH
AI
AJ
AK
AM
AN
AQ
AR
AT
AU
frequency
sample
haplotype
hap1
hap2
hap3

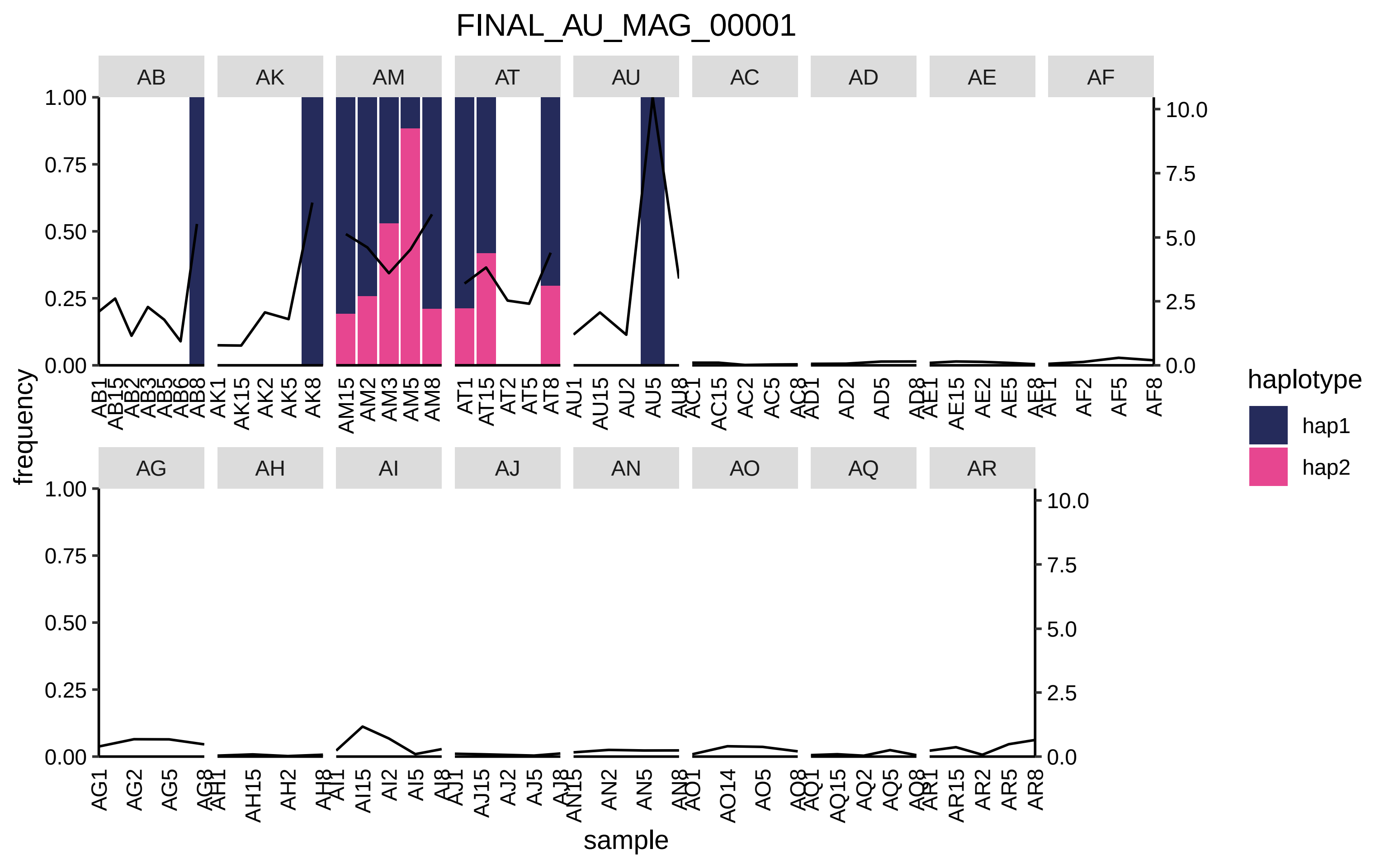
FINAL_AU_MAG_00001
AB
AK
AM
AT
AU
AC
AD
AE
AF
AG
AH
AI
AJ
AN
AO
AQ
AR
frequency
sample
haplotype
hap1
hap2
1.00
0.75
0.50
0.25
0.00
10.0
7.5
5.0
2.5
0.0

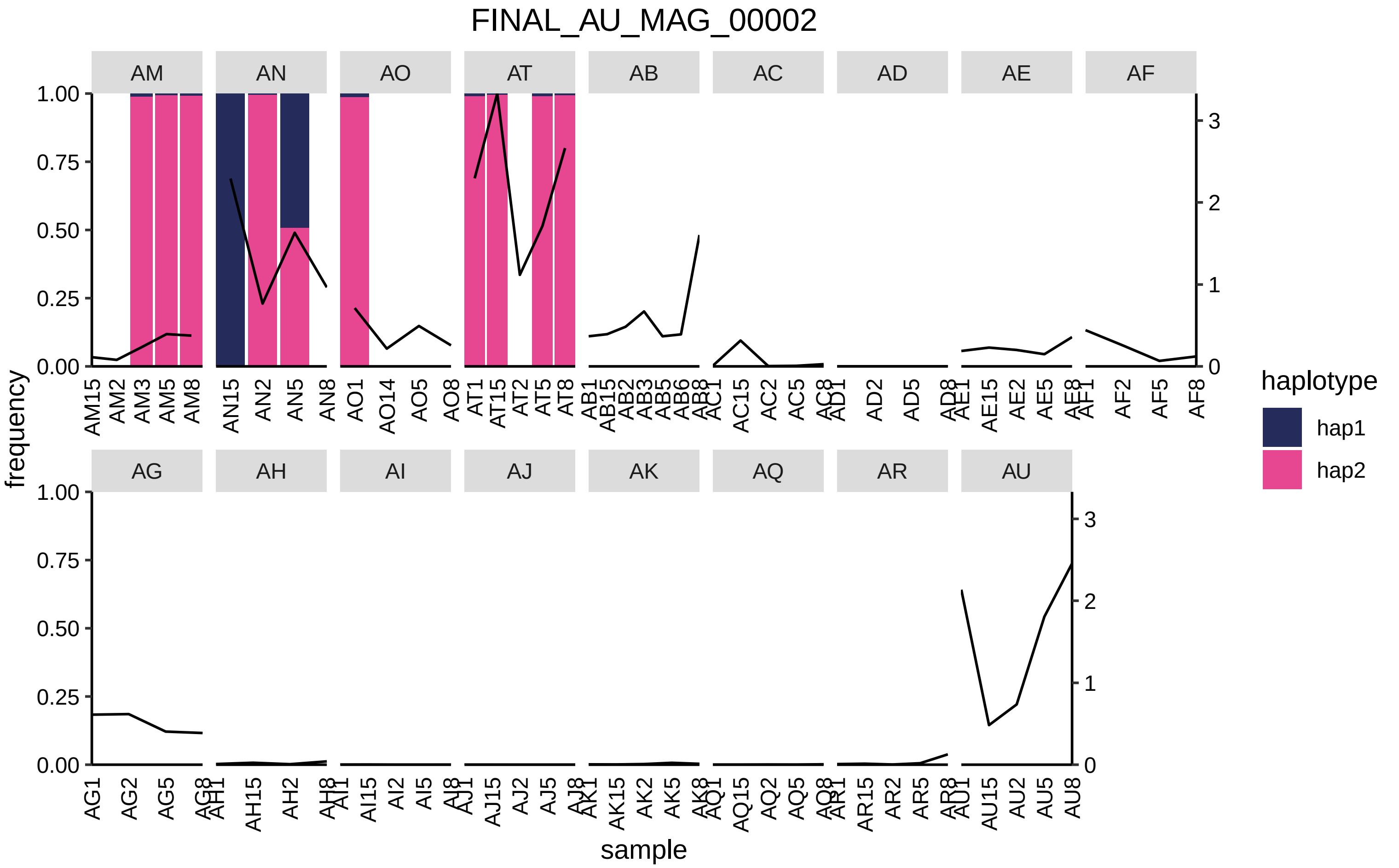
FINAL_AU_MAG_00002
AM
AN
AO
AT
AB
AC
AD
AE
AF
AG
AH
AI
AJ
AK
AQ
AR
AU
frequency
sample
haplotype
hap1
hap2
1.00
0.75
0.50
0.25
0.00
0
1
2
3
AM15 AM2 AM3 AM5 AM8
AN15 AN2 AN5 AN8
AO1 AO14 AO5 AO8
AT1 AT15 AT2 AT5 AT8
AB1 AB15 AB2 AB3 AB5 AB6 AB8
AC1 AC15 AC2 AC5 AC8
AD1 AD2 AD5 AD8
AE1 AE15 AE2 AE5 AE8
AF1 AF2 AF5 AF8
AG1 AG2 AG5 AG8
AH1 AH15 AH2 AH8
AI1 AI15 AI2 AI5 AI8
AJ1 AJ15 AJ2 AJ5 AJ8
AK1 AK15 AK2 AK5 AK8
AQ1 AQ15 AQ2 AQ5 AQ8
AR1 AR15 AR2 AR5 AR8
AU1 AU15 AU2 AU5 AU8

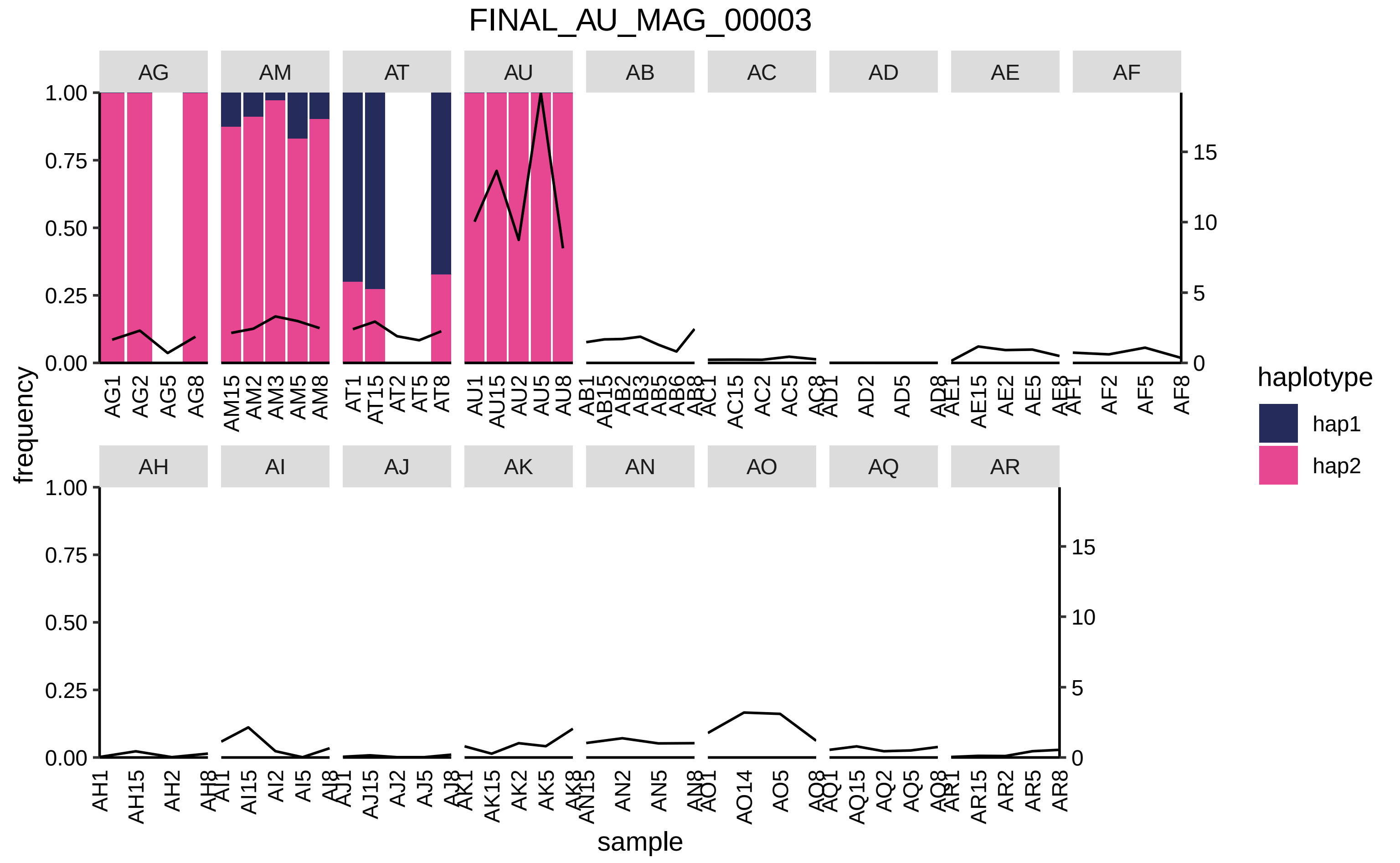
FINAL_AU_MAG_00003
AG
AM
AT
AU
AB
AC
AD
AE
AF
AH
AI
AJ
AK
AN
AO
AQ
AR
frequency
sample
haplotype
hap1
hap2
1.00
0.75
0.50
0.25
0.00
0
5
10
15
AG1 AG2 AG5 AG8
AM15 AM2 AM3 AM5 AM8
AT1 AT15 AT2 AT5 AT8
AU1 AU15 AU2 AU5 AU8
AB1 AB15 AB2 AB3 AB5 AB6 AB8
AC1 AC15 AC2 AC5 AC8
AD1 AD2 AD5 AD8
AE1 AE15 AE2 AE5 AE8
AF1 AF2 AF5 AF8
AH1 AH15 AH2 AH8
AI1 AI15 AI2 AI5 AI8
AJ1 AJ15 AJ2 AJ5 AJ8
AK1 AK15 AK2 AK5 AK8
AN15 AN2 AN5 AN8
AO1 AO14 AO5 AO8
AQ1 AQ15 AQ2 AQ5 AQ8
AR1 AR15 AR2 AR5 AR8

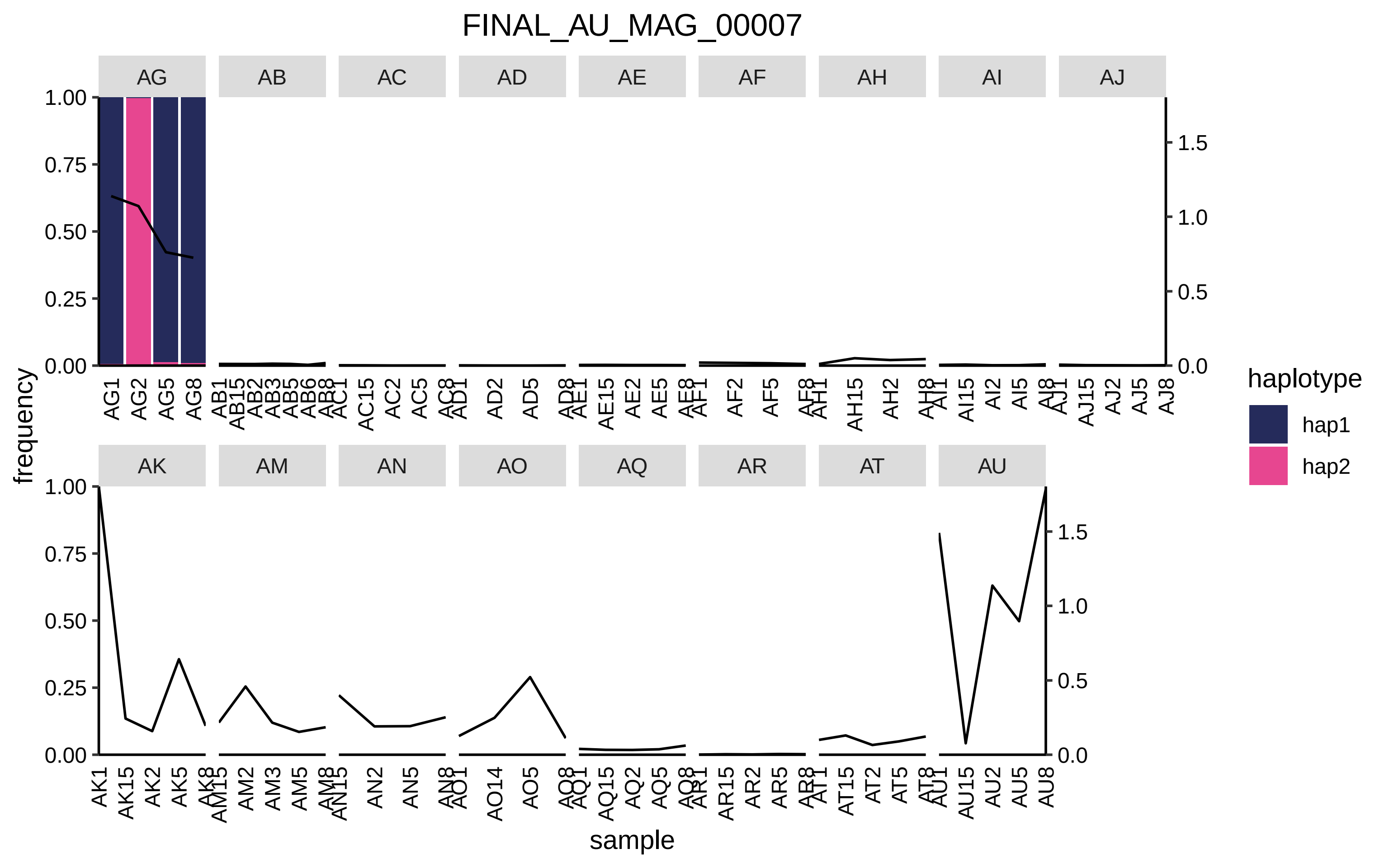

FINAL_AU_MAG_00007
AG
AB
AC
AD
AE
AF
AH
AI
AJ
AK
AM
AN
AO
AQ
AR
AT
AU
frequency
sample
haplotype
hap1
hap2

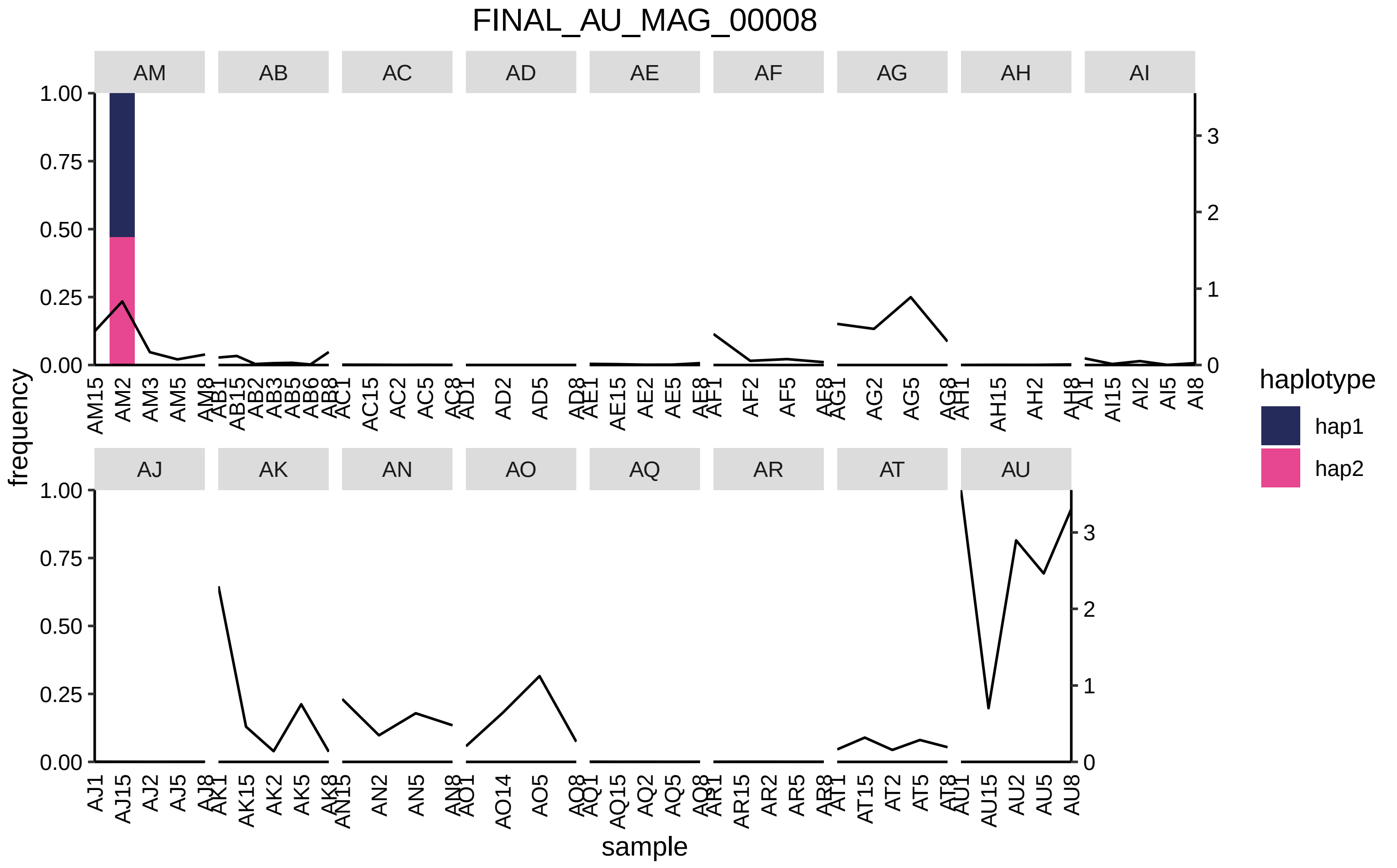

FINAL_AU_MAG_00008
AM
AB
AC
AD
AE
AF
AG
AH
AI
AJ
AK
AN
AO
AQ
AR
AT
AU
frequency
sample
1.00
0.75
0.50
0.25
0.00
0
1
2
3
AM15 AM2 AM3 AM5 AM8
AB1 AB15 AB2 AB3 AB5 AB6 AB8
AC1 AC15 AC2 AC5 AC8
AD1 AD2 AD5 AD8
AE1 AE15 AE2 AE5 AE8
AF1 AF2 AF5 AF8
AG1 AG2 AG5 AG8
AH1 AH15 AH2 AH8
AI1 AI15 AI2 AI5 AI8
AJ1 AJ15 AJ2 AJ5 AJ8
AK1 AK15 AK2 AK5 AK8
AN15 AN2 AN5 AN8
AO1 AO14 AO5 AO8
AQ1 AQ15 AQ2 AQ5 AQ8
AR1 AR15 AR2 AR5 AR8
AT1 AT15 AT2 AT5 AT8
AU1 AU15 AU2 AU5 AU8
haplotype
hap1
hap2

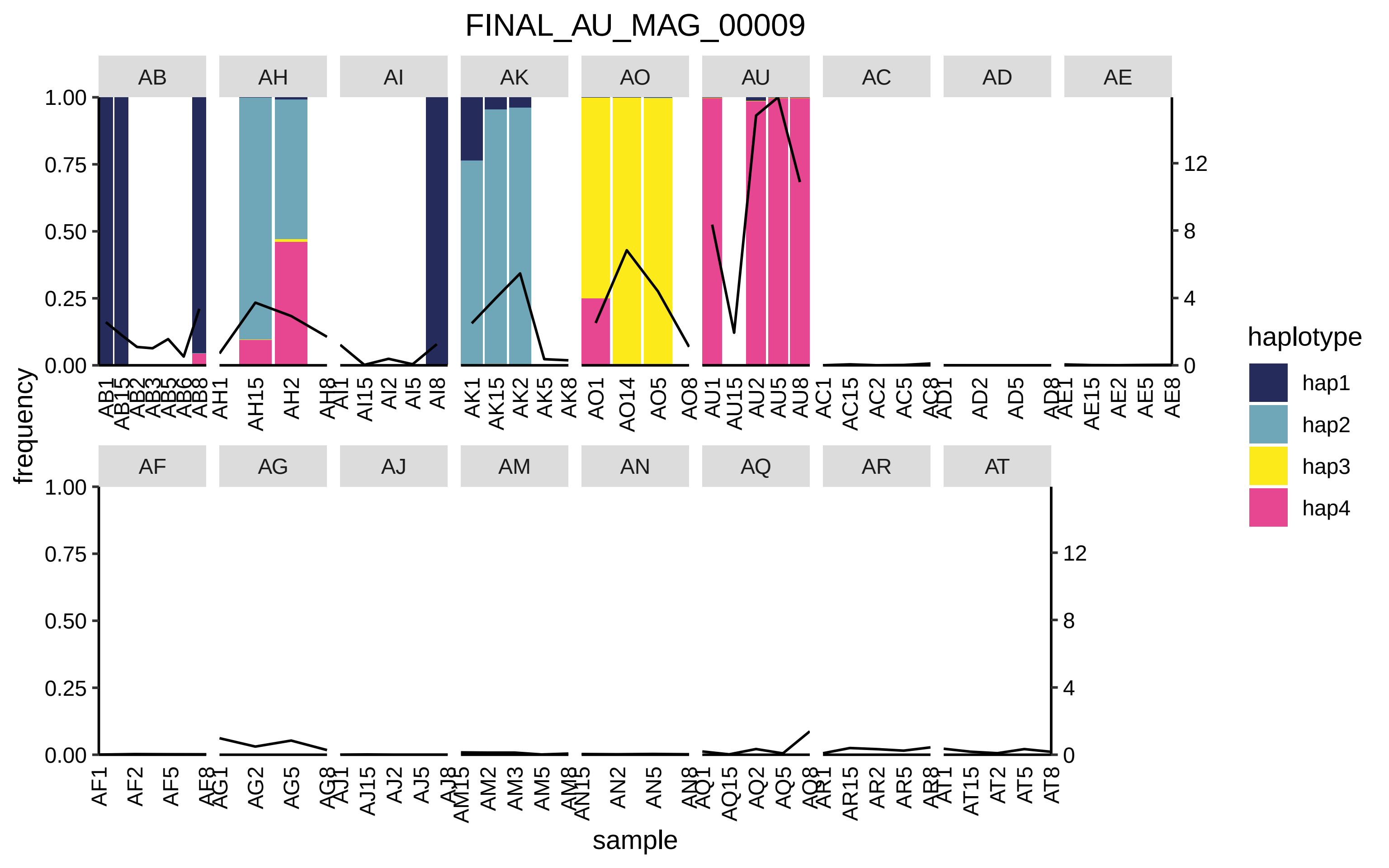
FINAL_AU_MAG_00009
AB
AH
AI
AK
AO
AU
AC
AD
AE
AF
AG
AJ
AM
AN
AQ
AR
AT
frequency
sample
haplotype
hap1
hap2
hap3
hap4
1.00
0.75
0.50
0.25
0.00
12
8
4
0

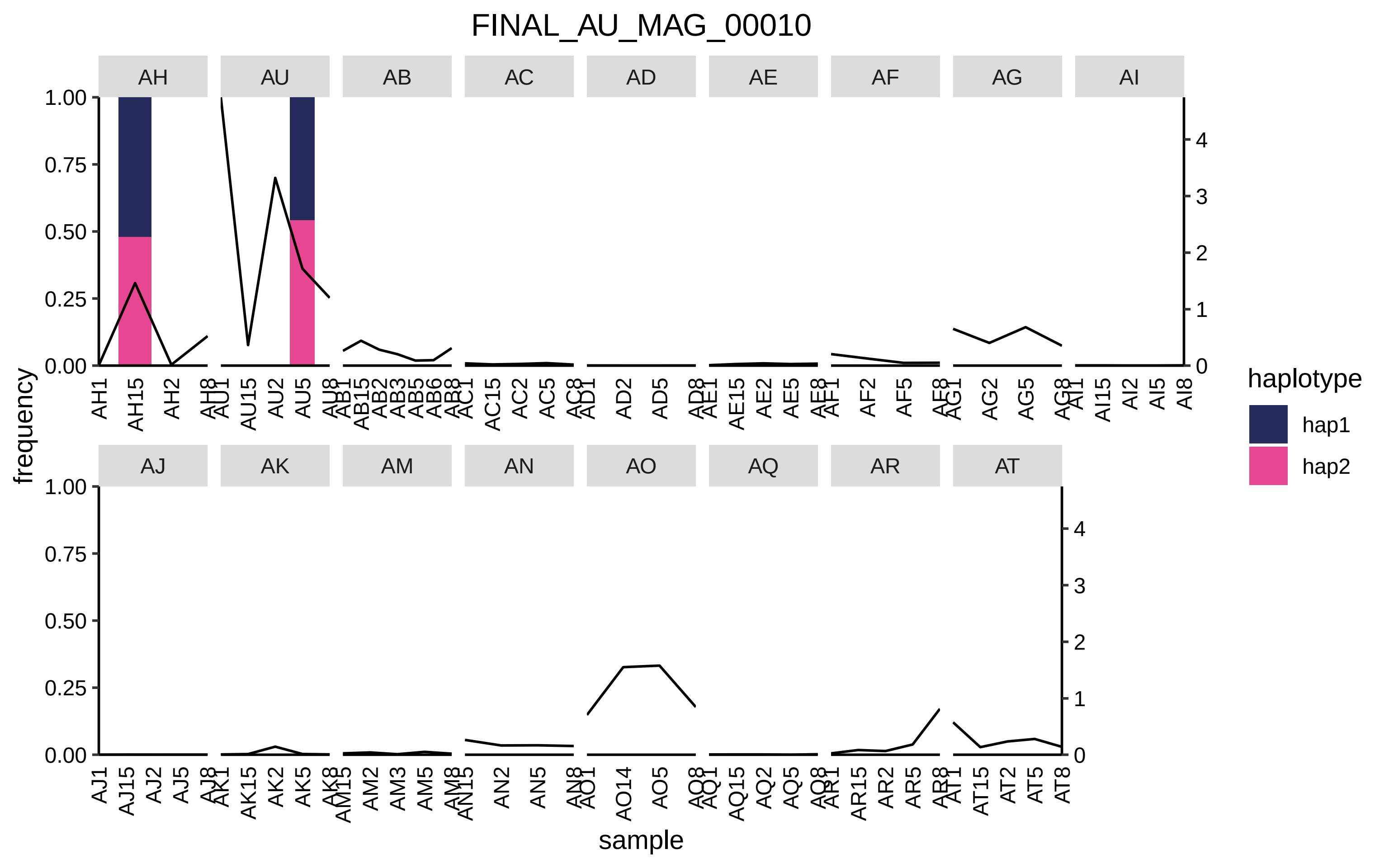
FINAL_AU_MAG_00010
AH
AU
AB
AC
AD
AE
AF
AG
AI
AJ
AK
AM
AN
AO
AQ
AR
AT
frequency
sample
haplotype
hap1
hap2
1.00
0.75
0.50
0.25
0.00
0
1
2
3
4
AH1 AH15 AH2 AH8
AU1 AU15 AU2 AU5 AU8
AB1 AB15 AB2 AB3 AB5 AB6 AB8
AC1 AC15 AC2 AC5 AC8
AD1 AD2 AD5 AD8
AE1 AE15 AE2 AE5 AE8
AF1 AF2 AF5 AF8
AG1 AG2 AG5 AG8
AI1 AI15 AI2 AI5 AI8
AJ1 AJ15 AJ2 AJ5 AJ8
AK1 AK15 AK2 AK5 AK8
AM15 AM2 AM3 AM5 AM8
AN15 AN2 AN5 AN8
AO1 AO14 AO5 AO8
AQ1 AQ15 AQ2 AQ5 AQ8
AR1 AR15 AR2 AR5 AR8
AT1 AT15 AT2 AT5 AT8

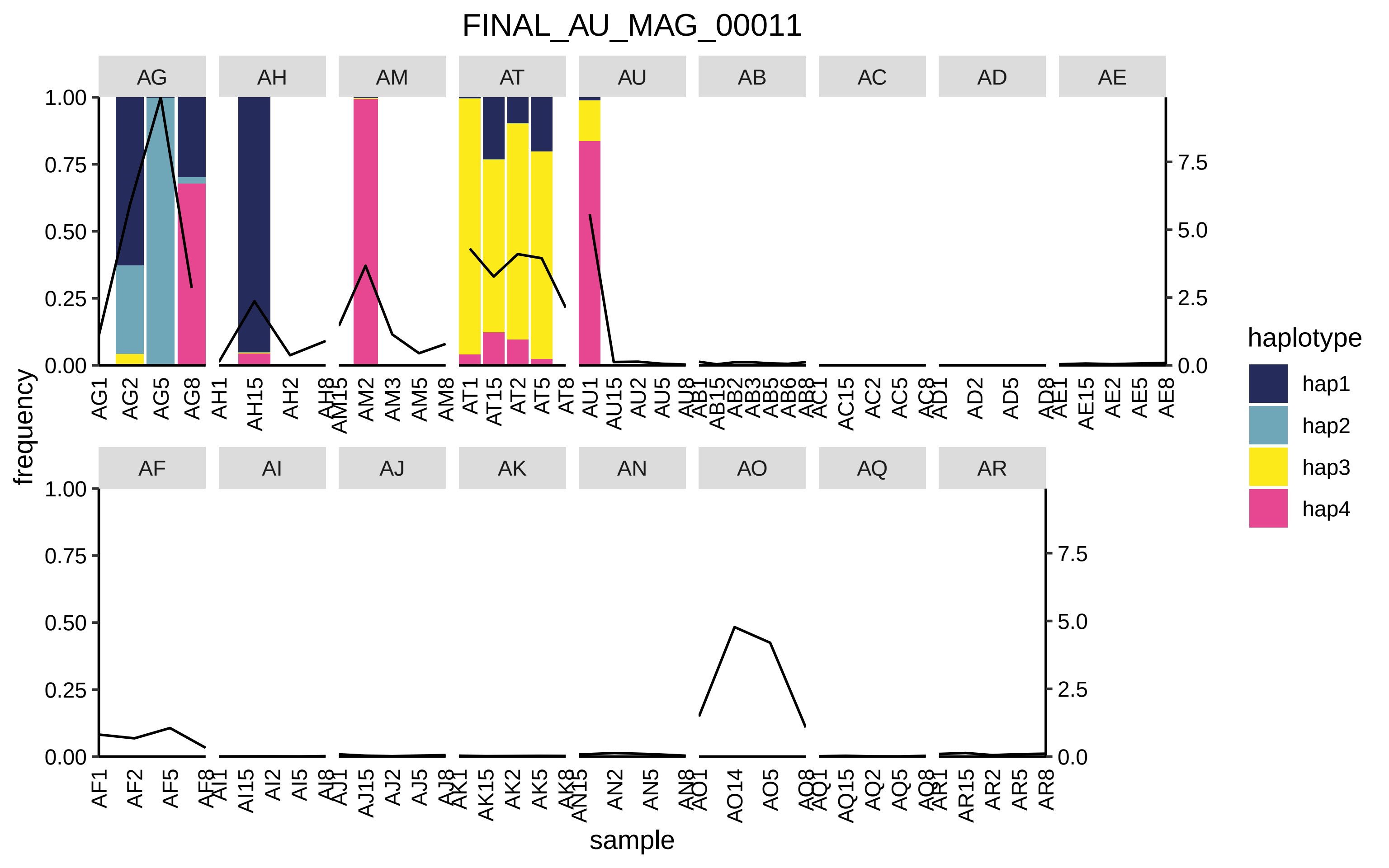

FINAL_AU_MAG_00011
AG
AH
AM
AT
AU
AB
AC
AD
AE
AF
AI
AJ
AK
AN
AO
AQ
AR
frequency
sample
haplotype
hap1
hap2
hap3
hap4

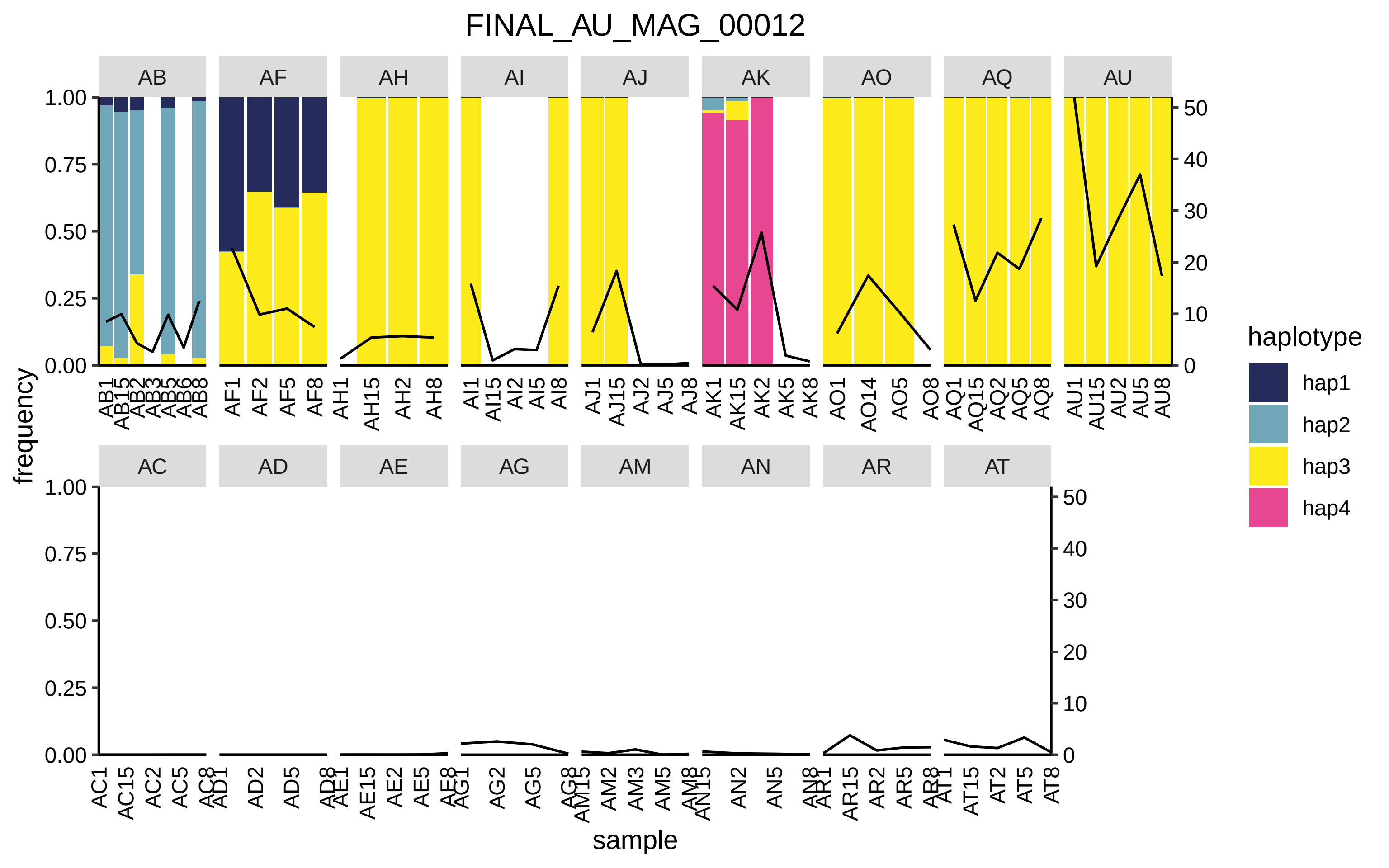
FINAL_AU_MAG_00012
frequency
sample
haplotype
hap1
hap2
hap3
hap4
AB
AF
AH
AI
AJ
AK
AO
AQ
AU
AC
AD
AE
AG
AM
AN
AR
AT

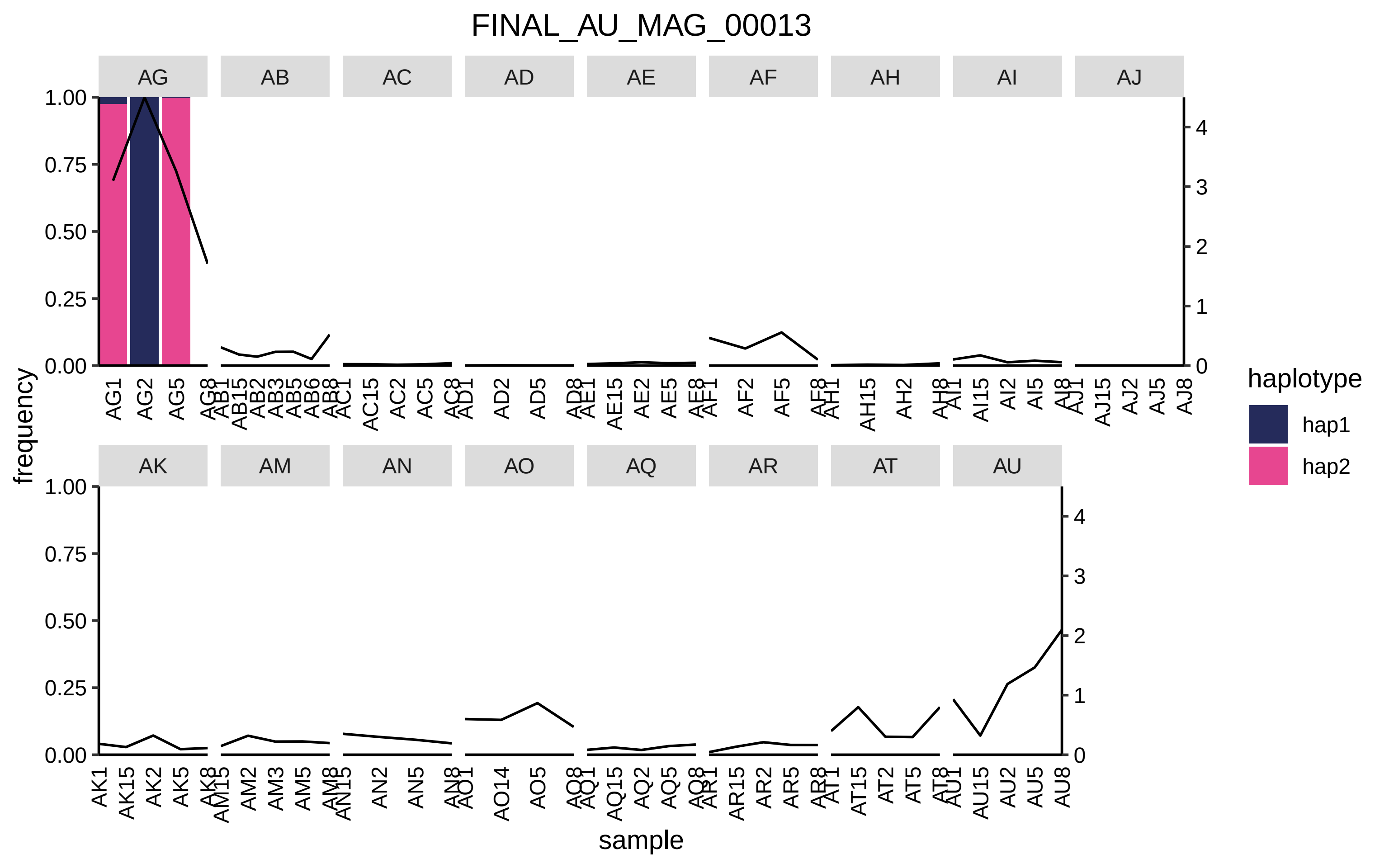
FINAL_AU_MAG_00013
AG
AB
AC
AD
AE
AF
AH
AI
AJ
AK
AM
AN
AO
AQ
AR
AT
AU
frequency
sample
haplotype
hap1
hap2

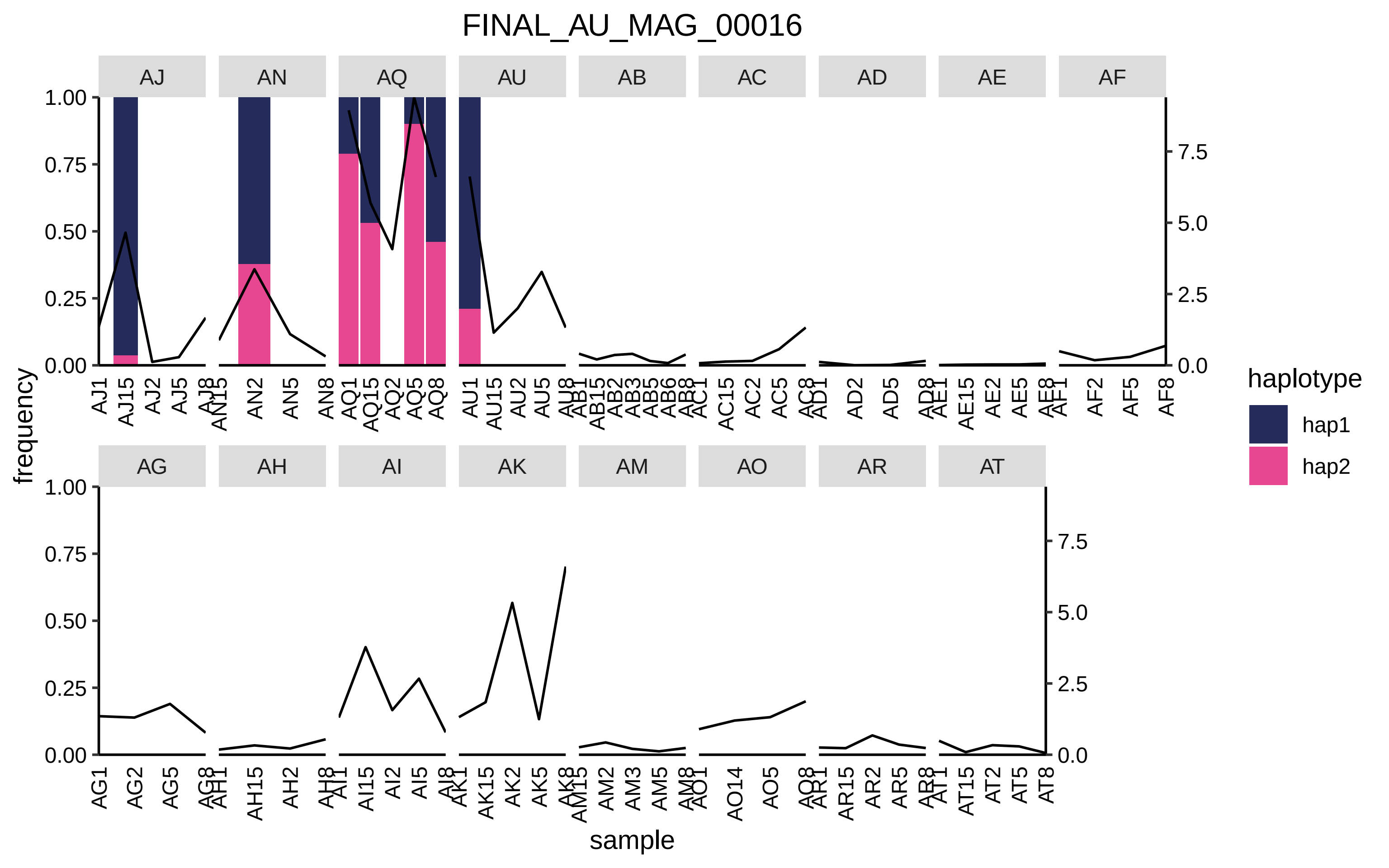

FINAL_AU_MAG_00016
AJ
AN
AQ
AU
AB
AC
AD
AE
AF
AG
AH
AI
AK
AM
AO
AR
AT
frequency
sample
haplotype
hap1
hap2
1.00
0.75
0.50
0.25
0.00
7.5
5.0
2.5
0.0

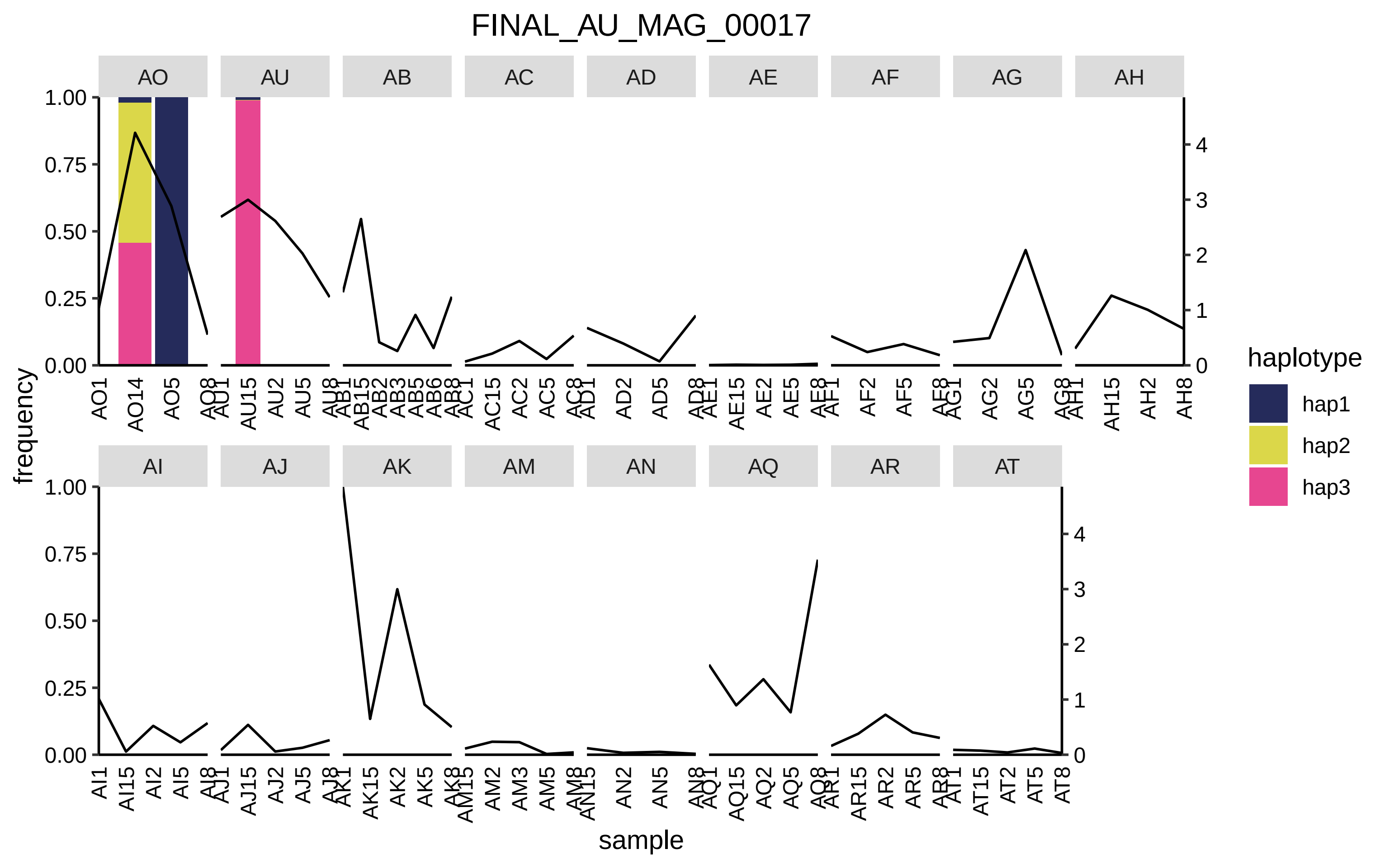

FINAL_AU_MAG_00017
AO
AU
AB
AC
AD
AE
AF
AG
AH
AI
AJ
AK
AM
AN
AQ
AR
AT
frequency
sample
haplotype
hap1
hap2
hap3
1.00
0.75
0.50
0.25
0.00
0
1
2
3
4
AO1
AO14
AO5
AO8
AU1
AU15
AU2
AU5
AU8
AB1
AB15
AB2
AB3
AB5
AB6
AB8
AC1
AC15
AC2
AC5
AC8
AD1
AD2
AD5
AD8
AE1
AE15
AE2
AE5
AE8
AF1
AF2
AF5
AF8
AG1
AG2
AG5
AG8
AH1
AH15
AH2
AH8
AI1
AI15
AI2
AI5
AI8
AJ1
AJ15
AJ2
AJ5
AJ8
AK1
AK15
AK2
AK5
AK8
AM15
AM2
AM3
AM5
AM8
AN15
AN2
AN5
AN8
AQ1
AQ15
AQ2
AQ5
AQ8
AR1
AR15
AR2
AR5
AR8
AT1
AT15
AT2
AT5
AT8

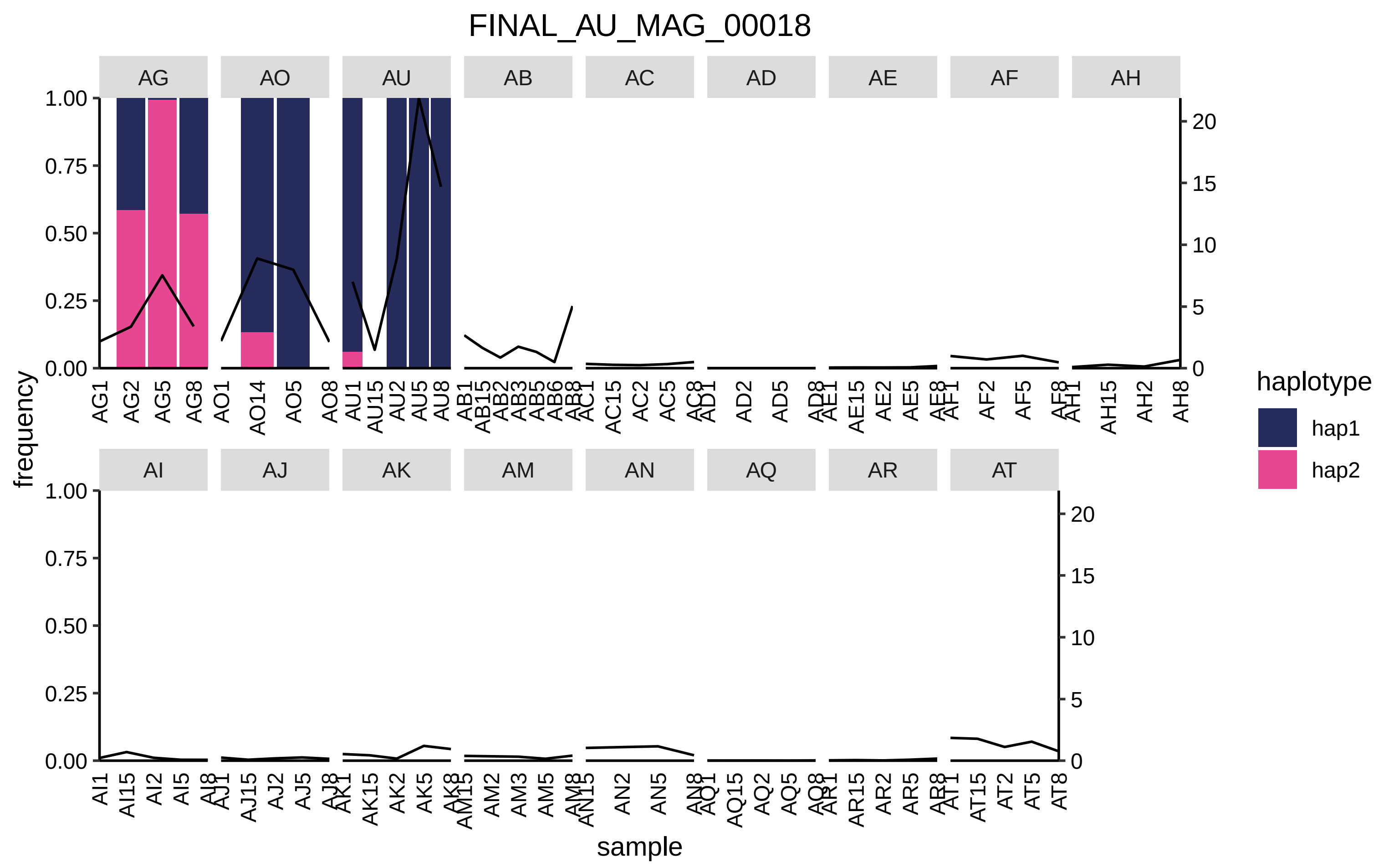

FINAL_AU_MAG_00018
AG
AO
AU
AB
AC
AD
AE
AF
AH
AI
AJ
AK
AM
AN
AQ
AR
AT
frequency
sample
haplotype
hap1
hap2
1.00
0.75
0.50
0.25
0.00
20
15
10
5
0
AG1 AG2 AG5 AG8 AO1 AO14 AO5 AO8 AU1 AU15 AU2 AU5 AU8 AB1 AB15 AB2 AB3 AB5 AB6 AB8 AC1 AC15 AC2 AC5 AC8 AD1 AD2 AD5 AD8 AE1 AE15 AE2 AE5 AE8 AF1 AF2 AF5 AF8 AH1 AH15 AH2 AH8
AI1 AI15 AI2 AI5 AI8 AJ1 AJ15 AJ2 AJ5 AJ8 AK1 AK15 AK2 AK5 AK8 AM15 AM2 AM3 AM5 AM8 AN15 AN2 AN5 AN8 AQ1 AQ15 AQ2 AQ5 AQ8 AR1 AR15 AR2 AR5 AR8 AT1 AT15 AT2 AT5 AT8

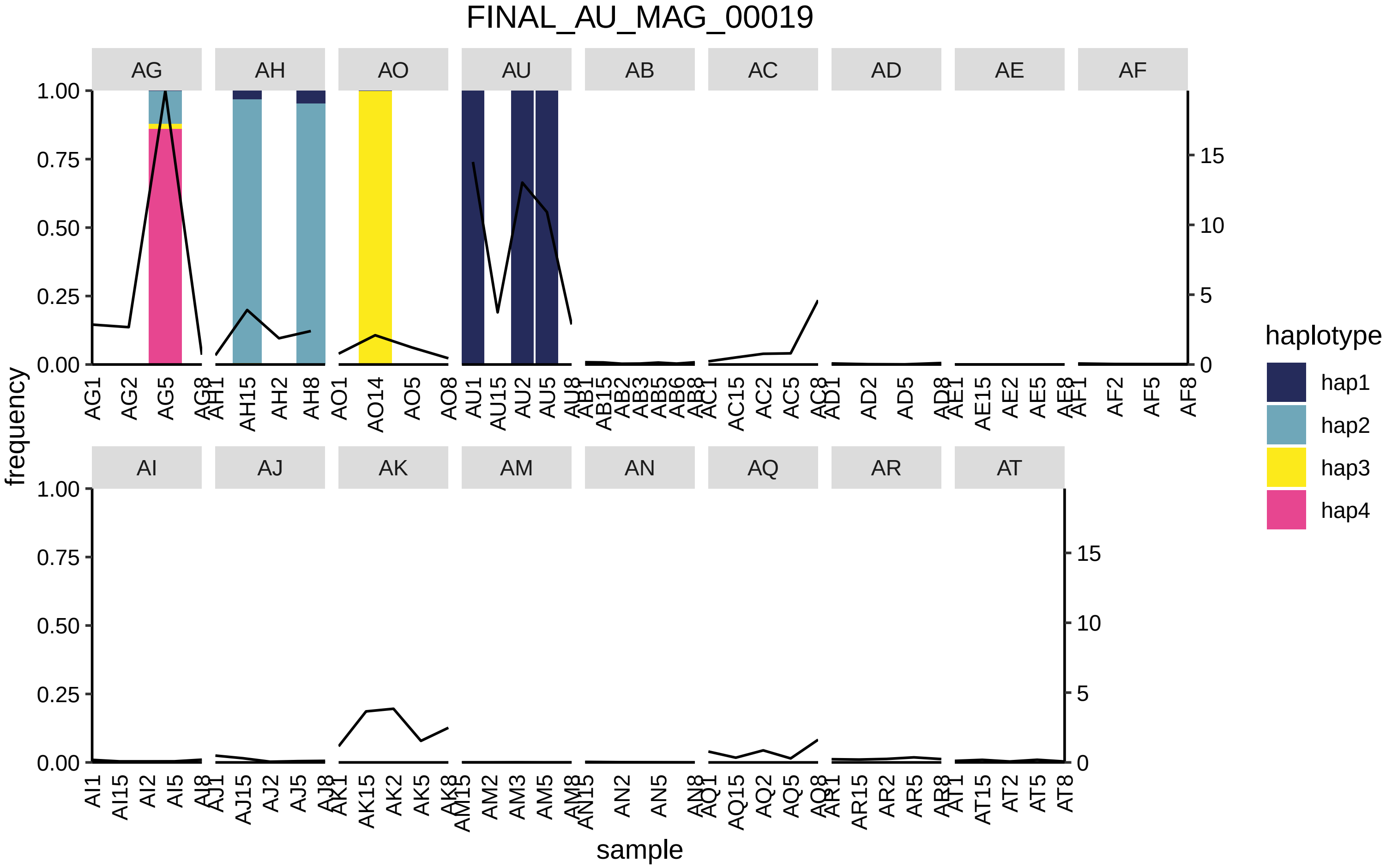
FINAL_AU_MAG_00019
AG
AH
AO
AU
AB
AC
AD
AE
AF
AI
AJ
AK
AM
AN
AQ
AR
AT
frequency
sample
haplotype
hap1
hap2
hap3
hap4
